# scOPE identifies which driver-associated expression programs transfer from bulk tumors to single cells

**DOI:** 10.64898/2026.07.24.740598

**Authors:** Andrew J. Ashford, Alex Lapadat, Emek Demir

**Affiliations:** Department of Molecular & Medical Genetics and Knight Cancer Institute, Oregon Health & Science University, Portland, OR 97239, USA; Department of Neuroscience, Amherst College, Amherst, MA 01002, USA

**Keywords:** single-cell RNA sequencing, cancer genomics, transfer learning, driver mutations, latent representations, copy-number variation

## Abstract

Single-cell RNA sequencing (scRNA-seq) resolves the phenotypic heterogeneity of tumors but rarely observes the somatic mutations that drive it: a variant is legible only where its gene is expressed, the mutant allele is transcribed, and reads span the variant site, so an absent variant read is fundamentally ambiguous. Bulk tumor cohorts have the opposite profile—matched genotype and expression for hundreds of patients, but no cellular resolution. We present scOPE (**s**ingle-**c**ell **O**ncological **P**rediction **E**xplorer), which learns cancer-specific, driver-associated expression axes from bulk tumors, freezes them, and projects single-cell transcriptomes onto the fixed axes without refitting to the target cohort. Our central finding is that this transfer is selective rather than general: of 158 audited driver–cancer models across seven malignancies, 102 met predefined claim-safety criteria and only 11 reached out-of-fold AUROC ≥ 0.90, led by acute myeloid leukemia (AML) *NPM1* (0.971), glioblastoma *IDH1* (0.963), and pancreatic adenocarcinoma *KRAS* (0.944). Determining *which* programs transfer therefore becomes the central task. We address it with a ground-truth-free confidence score—integrating bulk transferability, spatial coherence, score concentration, and copy-number (CNV) agreement—that within AML ranked the three independently supported programs above the remainder (AUROC 0.85 across 12 truth-evaluable drivers, of which three were supported), a triage signal rather than a validated genotype classifier. Against expressed-mutation labels, cell-state-residual scores were enriched in mutant-labeled cells for *NPM1*, *TP53*, and *DNMT3A*, and the *NPM1* separation survived aggregation to patients. Critically, matched genotype–score maps show that even supported programs occupy restricted transcriptional subspaces rather than uniformly marking mutation-positive tumors, and the *NPM1* program contracted during treatment across multiple patients. Transferred scores tracked inferred CNV burden yet also resolved discordant malignant populations invisible to aneuploidy alone. scOPE does not call alleles; it recovers continuous, mutation-associated transcriptional axes from existing scRNA-seq data, together with explicit diagnostics for when that reading should be withheld.

## 1 Introduction

Cancer progresses through the selection of genetically distinct clones whose phenotypes are shaped by driver mutations, lineage state, and treatment (Nowell, 1976; Greaves and Maley, 2012; Gerlinger et al., 2012; Vogelstein et al., 2013). Making sense of this process requires linking a cell’s genotype to its transcriptional state, because the transcriptome is the layer through which a mutation exerts its effects, evades therapy, and reshapes the tumor microenvironment. Single-cell RNA sequencing (scRNA-seq) has brought the phenotypic half of this link into sharp focus, resolving malignant hierarchies, developmental plasticity, and tumor–immune interactions across many cancers (Patel et al., 2014; Tirosh et al., 2016; van Galen et al., 2019; Neftel et al., 2019). The genotypic half, however, remains largely inaccessible within the same assay.

The obstacle is that scRNA-seq observes genotype only indirectly and incompletely. A somatic mutation can be detected in a given cell only when the affected gene is expressed, the mutant allele is transcribed and captured, and sequencing reads happen to span the variant position. Dropout, allelic imbalance, transcript-end bias, and shallow per-cell coverage combine so that the absence of a variant read is fundamentally ambiguous: it may indicate a wild-type cell, or merely an unobserved one (Brennecke et al., 2013; Kharchenko et al., 2014; Ziegenhain et al., 2017; Vu et al., 2019; Petti et al., 2019; Muyas et al., 2024; Marot-Lassauzaie et al., 2024). The mutations that define a tumor’s clonal architecture are thus, paradoxically, among the features that scRNA-seq is least equipped to report.

Several strategies narrow this gap, each within a bounded regime. Targeted single-cell assays amplify selected loci to place genotype and phenotype in the same cell, and computational methods integrate sparse variant reads with matched DNA-derived clonal structure (Nam et al., 2019; Rodriguez-Meira et al., 2019; Campbell et al., 2019; McCarthy et al., 2020). These are powerful where targeted panels, matched DNA, or adequate expressed-variant coverage exist—conditions that the large majority of already-archived scRNA-seq cohorts do not satisfy. Expression-derived copy-number inference offers a complementary route to identifying malignant cells, particularly in aneuploid tumors (Fan et al., 2018; R Gao et al., 2021; T Gao et al., 2023), although its accuracy depends strongly on dataset composition, copy-number architecture, and the specification of reference cells (Schmid et al., 2025). By construction, copy-number methods cannot recover the transcriptional consequence of a copy-neutral point mutation, and tumor-versus-normal classifiers or gene-set scores report malignancy or pathway activity rather than a program tied to a specific driver (Dohmen et al., 2022; Andreatta and Carmona, 2021). What remains unavailable is a way to read, from an ordinary scRNA-seq dataset, the expression program associated with a particular somatic driver.

A complementary and underused source of information exists at cohort scale. Bulk tumor profiling pairs RNA-seq with somatic genotype for hundreds of patients, and multiple studies have established that expression alone can predict selected oncogenic alterations (Way et al., 2017; Grzadkowski et al., 2021; Davis et al., 2018), as well as broader cancer-associated programs and their functional consequences (Cai et al., 2019; Jha et al., 2022). In principle, this matched resource could supply exactly the genotype–phenotype link that single-cell data lack. Moving it into single cells, however, is not a conventional classification task but a domain-transfer problem between two measurement regimes that differ in nearly every respect that matters. A bulk library averages malignant, stromal, and immune compartments into a single profile, whereas a single-cell library is a sparse, cell-type-specific snapshot; the two differ in normalization, dynamic range, dropout, and sequencing platform. A model that simply retrains on the target cohort risks discarding the very bulk-learned biology it was meant to carry over, while a model that ignores composition risks reporting lineage or tumor purity as though it were genotype.

Compounding this, drivers differ enormously in how legibly they act on the transcriptome. Some reorganize differentiation or chromatin and leave broad, reproducible signatures: *NPM1* -mutant acute myeloid leukemia (AML) adopts a characteristic HOX/MEIS-linked state (Falini et al., 2005); neomorphic *IDH1* /*IDH2* mutations remodel the epigenome through 2-hydroxyglutarate (Dang et al., 2009; Figueroa et al., 2010); and MAPK-pathway alterations define major melanoma subclasses (Davies et al., 2002; Cancer Genome Atlas Network, 2015). Others act through weak, context-dependent, or purely compositional effects that leave little cell-intrinsic transcriptional trace. This range has a direct methodological consequence. A classifier trained on bulk mutation labels can attain high accuracy for the wrong reason—by keying on molecular subtype, tumor purity, or a co-occurring alteration correlated with the label—and such a model transfers a confounder, not a driver program. The operative question is therefore not whether expression encodes genotype on average, but which individual driver programs are both reproducible in bulk and interpretable once projected into a cellular manifold.

Prior bulk expression–genotype models were not designed to answer that question. They are trained and evaluated within the bulk domain and return a performance value, not a judgment about whether—or where—the resulting program can be read in single cells. Two components are missing. The first is a transfer procedure that holds the bulk-trained axes fixed rather than silently refitting them to the target cohort, so that what is evaluated in single cells is the bulk-derived program itself. The second is an evaluation apparatus that decides, for each driver and each cohort, whether the transferred quantity can be trusted at all. This second requirement is decisive in precisely the setting that motivates the work: most archived scRNA-seq cohorts will never be genotyped, so a method whose reliability can be established only where ground truth already exists offers no guidance where it is actually needed. A usable framework must instead be able to flag its own likely failures without labels.

We developed scOPE (single-cell Oncological Prediction Explorer) to meet both requirements. For each cancer, scOPE learns a compact latent representation of bulk expression, trains driver-specific classifiers against matched genotype, and freezes the resulting gene loadings before any single-cell data are seen; no single-cell mutation label ever enters model fitting. Each transferred program is then examined through an evidence ladder that deliberately separates model performance from biological interpretation: held-out bulk discrimination; a ground-truth-free transfer-confidence score; direct mutation-label enrichment in AML; patient-level aggregation; genotype-associated per-cell localization; longitudinal treatment dynamics; cellular-state context; and comparison with expression-derived copy-number. No single rung of this ladder is treated as sufficient, and rungs that disagree are reported rather than reconciled.

This design yields three contributions. First, it establishes that only a minority of canonical drivers—not the general set—leave transcriptional structure reproducible enough to survive domain transfer, turning selectivity into an explicit, quantified result rather than an unstated caveat. Second, it supplies a label-free confidence score that ranks programs for interpretation before any genotype evidence is consulted, enabling a principled abstention policy in cohorts where ground truth will never exist. Third, it delivers a negative result of practical value: matched genotype–score maps show that even well-supported programs occupy restricted cellular subspaces, so a high transferred score is a statement about transcriptional similarity, not a per-cell genotype call. Applied across AML, breast invasive carcinoma (BRCA), colorectal cancer (CRC), glioblastoma (GBM), lung adenocarcinoma (LUAD), pancreatic adenocarcinoma (PAAD), and skin cutaneous melanoma (SKCM), scOPE identifies where driver programs transfer robustly, where they act primarily as markers of lineage or malignant state, and where the available evidence cannot support any mutation-specific interpretation.

## 2 Results

### 2.1 An evidence ladder separates driver-program transfer from genotype calling

scOPE was designed around two measurement regimes (Fig. 1). Bulk cohorts provide matched expression and mutation labels but average across cell types; scRNA-seq resolves cells but sparsely observes variant-bearing transcripts. Within each cancer, the bulk expression matrix was centered and scaled, factorized by truncated singular value decomposition (SVD), and used to train regularized driver-specific classifiers (Fig. 1A). Single cells were gene-matched, transformed with the stored bulk parameters, and projected onto the fixed bulk loadings (Fig. 1B). No single-cell mutation labels were used to fit the transfer map.

**Figure 1.**
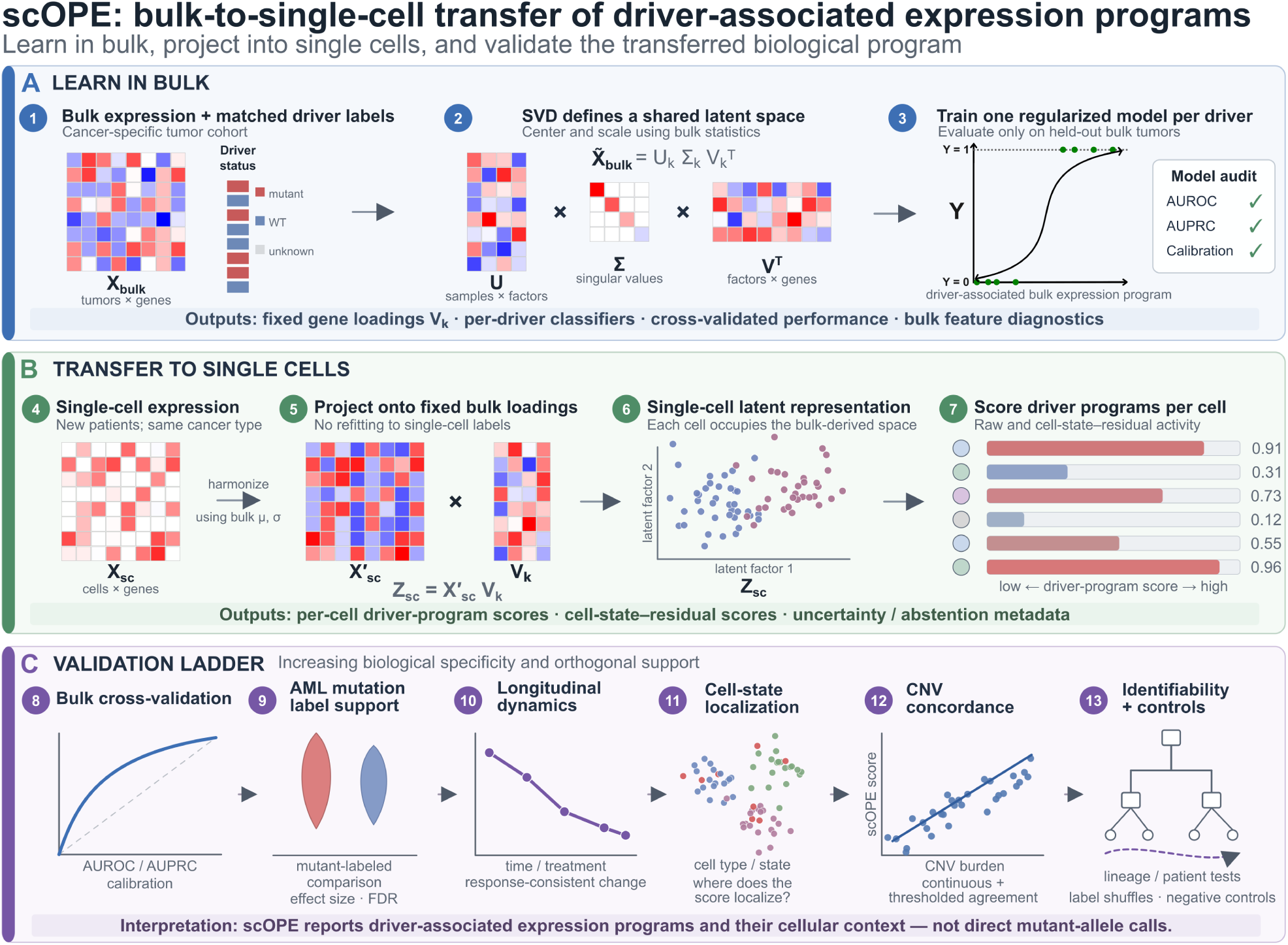
scOPE transfers bulk-derived driver-associated expression programs to single-cell transcriptomes. (A) Bulk RNA-seq expression profiles with matched driver-mutation labels are normalized and factorized by truncated singular value decomposition, producing a cancer-specific latent representation and fixed gene-loading matrix. Driver-specific classifiers are trained and evaluated in the bulk-derived latent space. (B) Single-cell expression profiles are then harmonized using the corresponding bulk-derived preprocessing parameters and projected through the fixed gene loadings to produce per-cell latent representations and driver-program scores. (C) The validation framework evaluates bulk cross-validated discrimination, enrichment in directly mutation-transcript-labeled AML cells, longitudinal treatment-associated dynamics, concordance with complementary expression-derived copy-number estimates, and nested identifiability and negative controls. scOPE estimates driver-associated transcriptional programs and their cellular context; it does not directly call mutant alleles.

The framework reports three conceptually distinct quantities. The raw classifier output measures similarity to the bulk mutation-associated program. A cell-state-residual score reduces broad lineage and healthy-reference effects and is the primary score for claim-bearing single-cell comparisons. Finally, a ground-truth-free confidence score (Fig. 1C) summarizes whether a driver is transferable in bulk, spatially coherent in the single-cell embedding, concentrated rather than diffuse, and concordant with a complementary expression-derived copy-number variation (CNV) axis. Confidence is used to prioritize interpretation; it is not a posterior probability that an individual cell carries the mutation.

### 2.2 Only a minority of drivers yield recoverable bulk expression programs

The prerequisite for transfer is that mutation status be recoverable from held-out bulk transcriptomes. scOPE evaluated 158 driver–cancer models across seven malignancies (Table 1; Fig. 2A). Of these, 102 met predefined claim-safety criteria, 75 achieved an out-of-fold area under the receiver operating characteristic curve (AUROC) ≥ 0.70, 39 achieved ≥ 0.80, and 11 achieved ≥ 0.90. Median performance was highest in CRC (0.788) and AML (0.779), followed by GBM (0.715), BRCA (0.710), LUAD (0.691), PAAD (0.652), and SKCM (0.597). This variation likely reflects both biology and study design, including mutation prevalence, cohort size, molecular-subtype structure, tumor purity, and the extent to which a driver produces a coherent downstream phenotype.

**Table 1.** Bulk driver-discrimination summary. Threshold counts refer to the full audit, which retains class-limited and failed evaluations. Claim-safe models had at least five mutation-positive tumors represented among valid pooled out-of-fold predictions, a finite AUROC whose 95% confidence-interval lower bound exceeded 0.5, and a Benjamini–Hochberg-adjusted permutation *q <* 0.05. The complete per-model audit is provided as Supplementary Table S1.

| Cancer | Evaluations | Median AUROC | Claim-safe | $\geq 0.70$ | $\geq 0.80$ | $\geq 0.90$ |
| --- | --- | --- | --- | --- | --- | --- |
| AML | 33 | 0.779 | 22 | 18 | 12 | 5 |
| BRCA | 22 | 0.710 | 15 | 9 | 4 | 1 |
| CRC | 25 | 0.788 | 23 | 20 | 11 | 2 |
| GBM | 17 | 0.715 | 8 | 6 | 3 | 1 |
| LUAD | 20 | 0.691 | 16 | 9 | 5 | 1 |
| PAAD | 15 | 0.652 | 7 | 6 | 1 | 1 |
| SKCM | 26 | 0.597 | 11 | 7 | 3 | 0 |
| Total | 158 | – | 102 | 75 | 39 | 11 |

**Figure 2.**
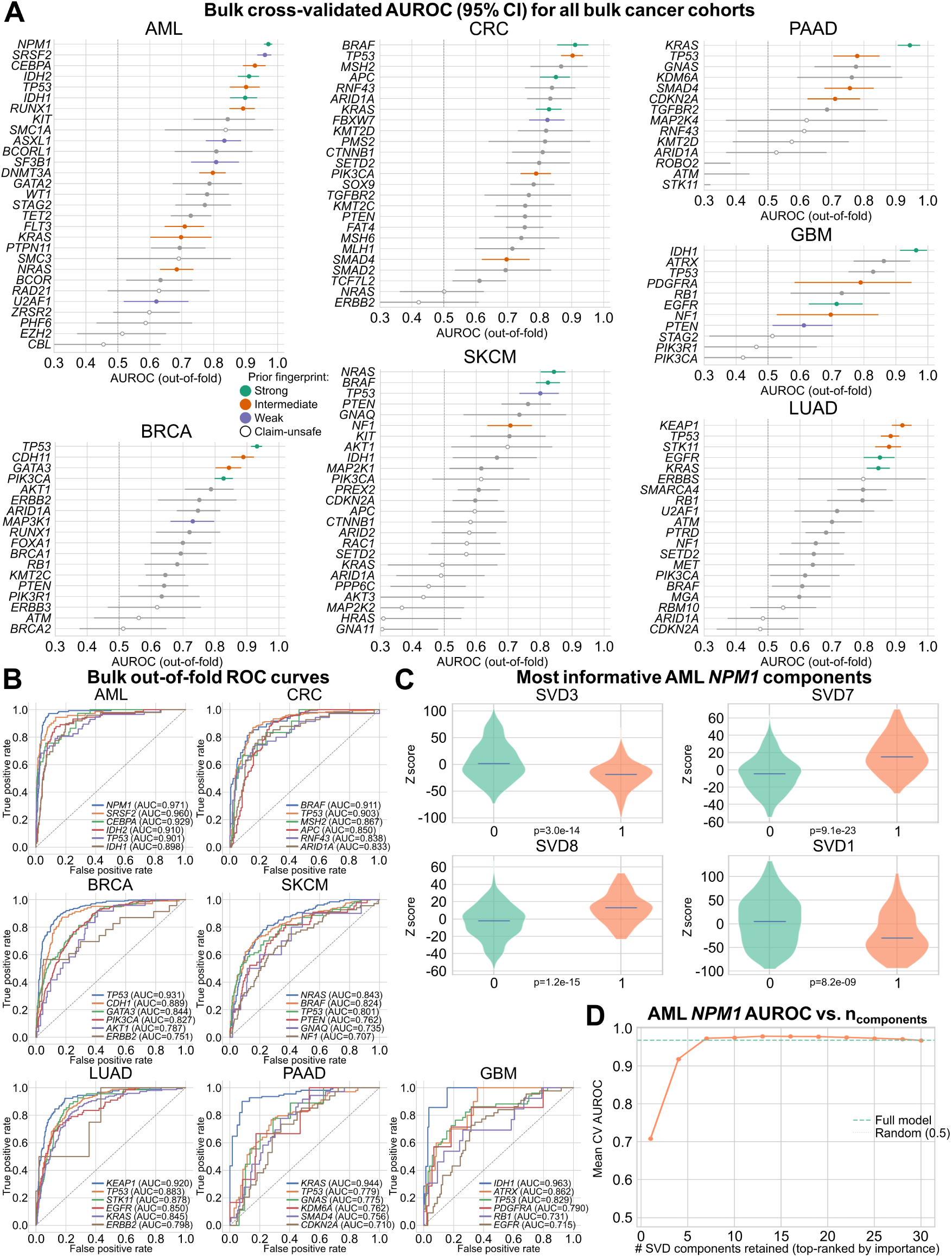
Bulk transcriptomes contain reproducible driver-associated expression programs. (A) Out-of-fold AUROC estimates and 95% confidence intervals for the complete driver–cancer audit, grouped by cancer type. Hollow symbols identify evaluations that did not meet the predefined claim-safety criteria; color denotes the prespecified biological-prior tier. (B) Out-of-fold ROC curves for representative models in all seven cancers. (C) Distribution of the four most informative latent components in *NPM1* -mutant and comparison bulk tumors. (D) AML *NPM1* performance after retaining increasing numbers of classifier-ranked latent components. Panels C–D are interpretability diagnostics from the separate driver-specific evaluator and do not replace the pooled out-of-fold benchmark in panels A–B. Complete class counts, AUPRC, calibration, permutation tests, and audit statistics are provided in Supplementary Table S1.

Canonical alterations generated several of the strongest held-out signals. In AML, *NPM1* reached AUROC 0.971 and an area under the precision–recall curve (AUPRC) of 0.872 against a prevalence baseline of 0.217, followed by *SRSF2* (0.960), *CEBPA* (0.929), and *IDH2* (0.910). Leading models in the other cohorts included GBM *IDH1* (0.963), PAAD *KRAS* (0.944), BRCA *TP53* (0.931), CRC *BRAF* (0.911), LUAD *KEAP1* (0.920), and SKCM *NRAS* (0.843). AUPRC and positive-class counts were considered alongside AUROC because several driver labels were imbalanced (Saito and Rehmsmeier, 2015); for example, the GBM *IDH1* estimate was based on 7 positive and 148 negative tumors and therefore requires wider uncertainty than its point estimate alone suggests (Fig. 2A,B). Bootstrap-interval width scaled inversely with the number of mutation-positive tumors across the audit, and AUPRC gains above prevalence, Brier scores, and claim-safety counts are summarized in Supplementary Fig. S1.

The AML *NPM1* model provided a well-powered example of how the signal was organized. Component-ablation diagnostics showed that a small set of classifier-ranked SVD components recovered approximately the full-model AUROC, consistent with a compact multicomponent program rather than dependence on a single axis (Fig. 2C,D; Supplementary Fig. S3). Performance was not an artifact of the chosen latent dimensionality: a sensitivity grid spanning component counts on either side of the canonical *k* = 30 produced comparable median AUROCs in every cancer, and this grid was used diagnostically rather than for post hoc model selection (Supplementary Fig. S2; Supplementary Table S9). Per-model label-permutation nulls for all 158 attempted models are reported in Supplementary Table S1 and summarized across controls in Supplementary Table S10.

Two features of this audit deserve emphasis, because they determine how the remainder of the study should be read. First, weak, class-limited, skipped, and failed evaluations were retained rather than removed after inspection, so the denominator of 158 reflects every model attempted rather than a surviving subset. Second, and more importantly, the distribution is heavily skewed: fewer than one model in four reached AUROC 0.80, and only 11 reached 0.90. Bulk recoverability is therefore the exception rather than the rule, and it defines the outer boundary of what can be transferred. Strong out-of-fold performance establishes that mutation-associated expression structure exists in bulk; it says nothing yet about the genotype of any single cell.

### 2.3 A ground-truth-free confidence score prioritizes, but does not prove, transfer-able programs

Bulk performance alone cannot determine whether a program remains coherent after transfer to single cells. We therefore summarized four ground-truth-free properties for each transferred driver: bulk transferability (*T*), spatial coherence in the single-cell embedding (*S*), score concentration (*C*), and agreement (*X*) with inferred expression-derived CNV. Composite confidence varied substantially among drivers and cohorts (Fig. 3A; Supplementary Fig. S4), allowing low-information programs to be down-prioritized before direct labels were examined.

**Figure 3.**
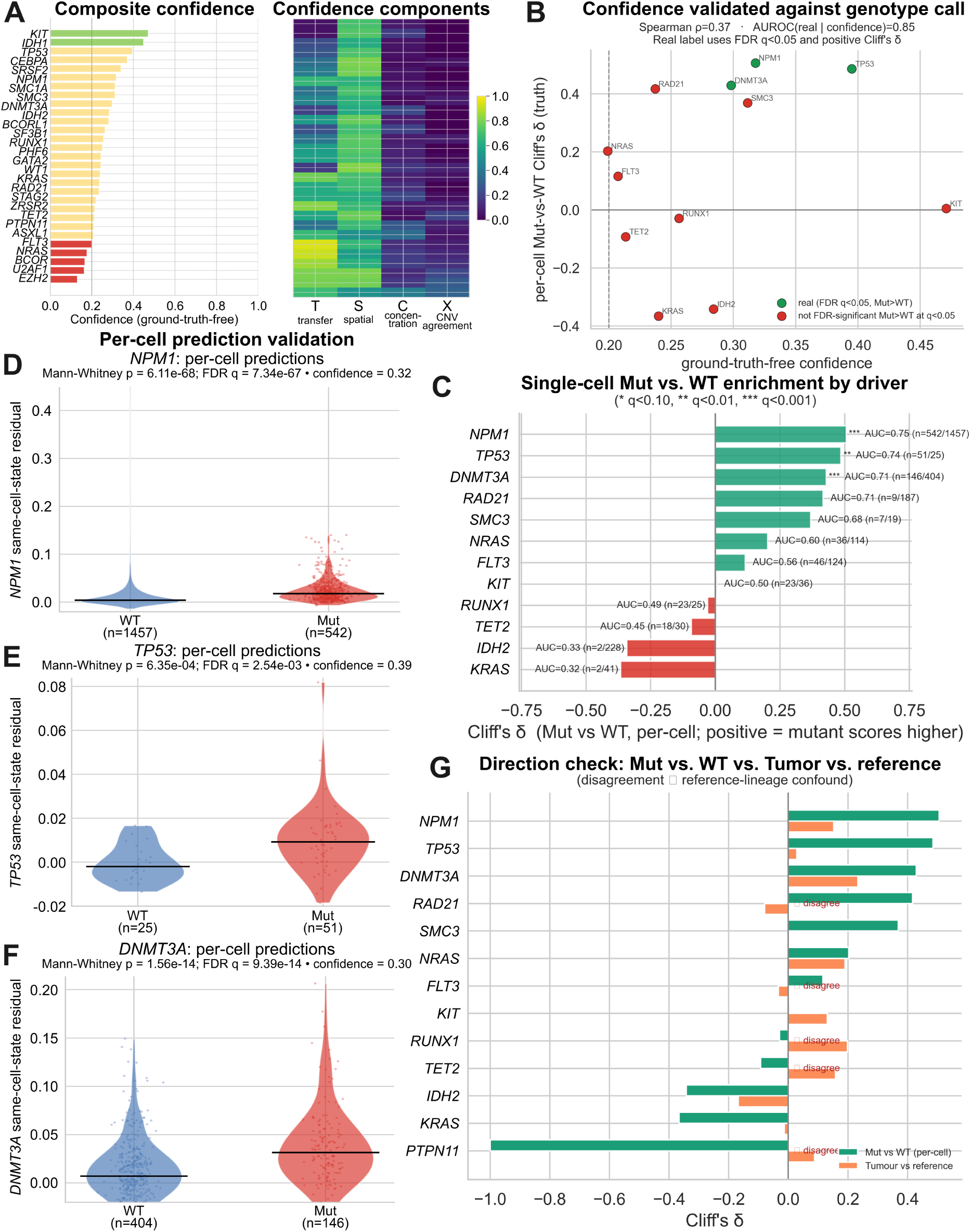
Ground-truth-free confidence prioritizes transferred programs, and AML labels support selected drivers. (A) Composite confidence (left) and its four components (right). (B) Confidence compared with direct-label effect size (*ρ* = 0.37; AUROC 0.85 for supported versus unsupported drivers). (C) Cliff’s *δ* across evaluable AML drivers. (D–F) Cell-state-residual scores for *NPM1*, *TP53*, and *DNMT3A*. (G) Directional agreement between mutation-label and malignant-reference contrasts. *PTPN11* appears only in panel G because a direction statistic was computable, whereas its direct-label coverage was insufficient for the prespecified analysis in panels B–F. Confidence profiles for the six remaining cancers and the nine remaining driver-level AML violins are in Supplementary Figs. S4–S5, respectively.

AML provided the only opportunity to test whether this unsupervised ranking anticipated separate mutation-label support. Among truth-evaluable drivers, confidence correlated moderately with the direct-label effect size, although the correlation was not statistically significant at this sample size (Spearman *ρ* = 0.37, *p* = 0.24, *n* = 12; Fig. 3B). When a supported program was defined as a positive Cliff’s *δ* with a false-discovery rate (FDR) *q <* 0.05, confidence discriminated supported from unsupported drivers with AUROC 0.85 (Fig. 3B). Both figures rest on 12 truth-evaluable drivers containing only three supported programs (Supplementary Table S3), so they should be read as evidence that the ranking is useful within this cohort and not as an externally validated performance estimate; a comparably labeled second cohort would be required for the latter.

The relationship was also not one-to-one, and the exceptions are instructive. *KIT* and *IDH1* received high confidence because their transferred scores were coherent by the ground-truth-free criteria, yet *KIT* showed no direct-label enrichment whatsoever (*δ* = 0.005). Conversely, *NPM1* carried only moderate composite confidence (0.32) despite having by far the strongest direct mutation-label evidence, because its score distribution was diffuse rather than concentrated (*C* = 0.14) and its CNV agreement was low (*X* = 0.08; Fig. 3A). Confidence therefore measures whether a transferred program is internally coherent, which is related to, but distinct from, whether it tracks the intended genotype. It is a triage device indicating where to inspect a program, not a surrogate genotype label or a substitute for direct validation.

### 2.4 AML mutation labels support transfer for *NPM1*, *TP53*, and *DNMT3A*

We next evaluated cell-state-residual scores against reported mutation annotations in the van Galen AML cohort. Because expressed mutation calls are incomplete, this analysis was framed as positive-unlabeled: mutant-labeled cells were treated as high-confidence positives, whereas comparison cells were required to carry an explicit wild-type transcript annotation and were not assumed to represent exhaustive genomic wild type. Across 12 evaluable drivers, three programs showed positive FDR-corrected enrichment (Table 2; Fig. 3C). *NPM1* separated 542 mutant-labeled from 1,457 comparison cells with apparent AUROC 0.753 (95% confidence interval [CI] 0.730–0.776), Cliff’s *δ* = 0.506, and FDR *q* = 7.3 × 10*^−^*^67^. *TP53* showed similar separation despite fewer cells (51 versus 25; AUROC 0.743; *δ* = 0.485; *q* = 2.5 × 10*^−^*^3^), and *DNMT3A* showed AUROC 0.714 and *δ* = 0.429 across 146 versus 404 cells (*q* = 9.4 × 10*^−^*^14^). The corresponding per-cell score distributions are shown in Fig. 3D–F.

**Table 2.** AML per-cell mutation-label validation using cell-state-residual scores. “Comparison” required an explicit wild-type transcript annotation without a contradictory mutant annotation; it does not establish complete genomic wild type.

| Driver | Mutant | Comparison | Apparent AUROC | Cliff’s $\delta$ | FDR $q$ |
| --- | --- | --- | --- | --- | --- |
| *NPM1* | 542 | 1,457 | 0.753 | 0.506 | $7.3 \times 10^{-67}$ |
| *TP53* | 51 | 25 | 0.743 | 0.485 | $2.5 \times 10^{-3}$ |
| *DNMT3A* | 146 | 404 | 0.714 | 0.429 | $9.4 \times 10^{-14}$ |
| *RAD21* | 9 | 187 | 0.708 | 0.417 | 0.106 |
| *SMC3* | 7 | 19 | 0.684 | 0.368 | 0.338 |
| *NRAS* | 36 | 114 | 0.601 | 0.203 | 0.162 |
| *FLT3* | 46 | 124 | 0.558 | 0.116 | 0.425 |
| *KIT* | 23 | 36 | 0.502 | 0.005 | 0.981 |
| *RUNX1* | 23 | 25 | 0.485 | -0.030 | 0.948 |
| *TET2* | 18 | 30 | 0.454 | -0.093 | 0.722 |
| *IDH2* | 2 | 228 | 0.329 | -0.342 | 0.585 |
| *KRAS* | 2 | 41 | 0.317 | -0.366 | 0.585 |

The remaining drivers defined useful boundaries. *RAD21* and *SMC3* were directionally positive but underpowered; *NRAS* and *FLT3* showed weak positive effects; *KIT* and *RUNX1* were near null; and *TET2*, *IDH2*, and *KRAS* were directionally negative, with the latter two based on only two mutant-labeled cells each. A direction diagnostic compared mutant-versus-comparison effects with malignant-versus-reference effects (Fig. 3G). The three FDR-supported drivers agreed in sign, whereas several unsupported drivers showed disagreement consistent with lineage or reference-set confounding. Together, these results support transfer of selected mutation-associated programs while arguing against universal per-cell genotyping.

These cell-level tests are descriptive because cells are nested within patients. We therefore complemented them with patient-level aggregation using matched sample genotype, reported below. Cell-level effect sizes remain useful for localization and positive-unlabeled discrimination, but patient-level effects carry the more conservative inferential interpretation.

### 2.5 Patient-level aggregation retains *NPM1* in AML and supports *TP53* in CRC

To reduce within-patient pseudoreplication, we aggregated cell-state-residual scores by patient and compared genotype-positive with genotype-negative patients (Table 3). In AML, *NPM1* showed complete rank separation between five mutation-positive and five comparison patients (Cliff’s *δ* = 1.00, FDR *q* = 0.040). *DNMT3A* and *NRAS* were directionally strong but narrowly missed FDR significance (*q* = 0.060 for each), whereas *TP53* did not separate the small patient groups. Thus, the cell-level *NPM1* result survives a substantially more conservative unit of analysis; the *TP53* and *DNMT3A* cell-level results remain supported as cellular enrichment but not as definitive AML patient-level associations in this cohort.

**Table 3.** Patient-level genotype bridge using median cell-state-residual scores. FDR was controlled within each cohort’s evaluable driver family.

| Cohort | Driver | Mutant patients | Comparison patients | Cliff’s $\delta$ | FDR $q$ |
| --- | --- | --- | --- | --- | --- |
| AML | *NPM1* | 5 | 5 | 1.000 | 0.040 |
| AML | *DNMT3A* | 6 | 6 | 0.778 | 0.060 |
| AML | *NRAS* | 3 | 5 | 1.000 | 0.060 |
| AML | *FLT3* | 2 | 5 | 0.400 | 0.714 |
| AML | *TP53* | 3 | 5 | −0.067 | 1.000 |
| CRC | *TP53* | 14 | 19 | 0.571 | 0.030 |
| CRC | *APC* | 17 | 16 | 0.404 | 0.124 |
| CRC | *SMAD4* | 4 | 29 | 0.397 | 0.373 |
| CRC | *BRAF* | 4 | 29 | −0.293 | 0.472 |
| CRC | *KRAS* | 10 | 23 | 0.087 | 0.710 |
| BRCA | *ERBB2* | 5 | 21 | 0.200 | 0.527 |

The same bridge was evaluable for selected solid-tumor drivers with matched patient metadata. In CRC, *TP53* scores were higher in 14 mutation-positive than 19 comparison patients (*δ* = 0.571, FDR *q* = 0.030); *APC* was nominally positive but did not survive correction. BRCA *ERBB2* showed no patient-level separation. These results provide a patient-level aggregation layer while also emphasizing that patient-level validation remains driver- and metadata-dependent. The complete bridge, including all evaluable drivers in AML, CRC, and BRCA, is given in Supplementary Table S4.

### 2.6 Matched genotype and score maps reveal state-restricted cellular localization

Aggregate effect sizes do not show whether a driver-associated signal is distributed broadly across cells from a genotype-positive sample or concentrated within a restricted transcriptional state. We therefore displayed patient identity, available mutation annotation, and the corresponding cell-state-residual score on identical expression-derived uniform manifold approximation and projection (UMAP) coordinates (Fig. 4). In AML, mutation status was assigned from direct per-cell expressed-mutation annotations. In CRC, mutation status was available only at the patient level and was assigned to the constituent cells of each tumor for visualization; these colors provide sample-level genotype context and were not treated as per-cell truth.

**Figure 4.**
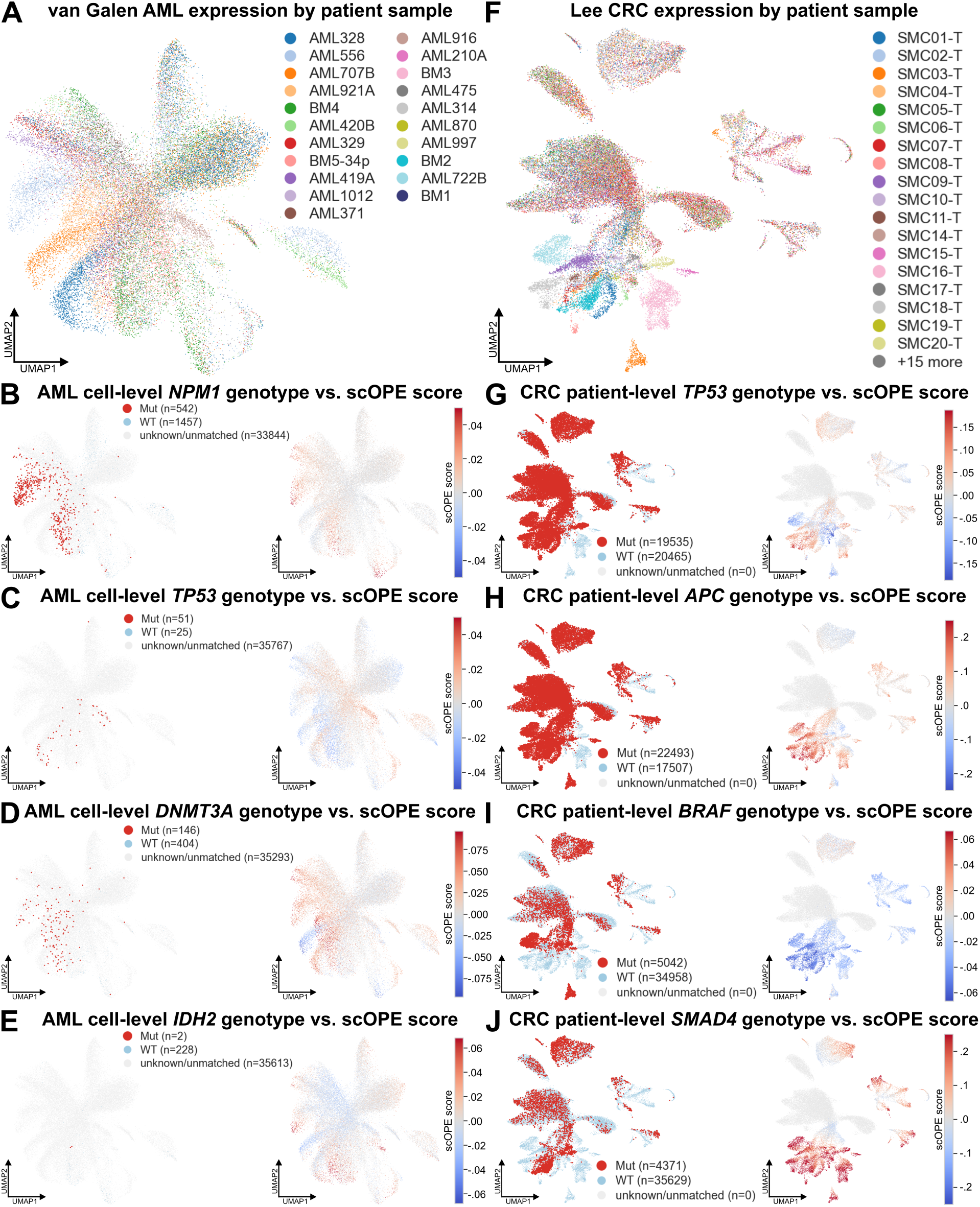
Driver-associated scOPE scores localize to heterogeneous, genotype-associated cellular subspaces. (A,F) AML and CRC UMAPs colored by patient. (B–E) AML per-cell mutation annotations and matched residual-score maps for *NPM1*, *TP53*, *DNMT3A*, and *IDH2*. (G–J) CRC patient-level mutation-status overlays and matched score maps for *TP53*, *APC*, *BRAF*, and *SMAD4*. AML annotations reflect expressed mutant or wild-type transcripts; CRC annotations reflect tumor-level genotype and are not per-cell truth. Scores are continuous, driver-specific phenotypes, with separately centered color scales. Complete driver atlases are shown in Supplementary Fig. S7.

The AML maps made the direct-label results spatially interpretable (Fig. 4A–E). *NPM1* -annotated mutant cells occupied coherent regions that overlapped elevated *NPM1* residual scores, although the score gradient extended beyond cells with detected mutant transcripts (Fig. 4B). The *TP53* and *DNMT3A* programs showed partial rather than complete spatial correspondence between directly annotated mutant cells and elevated scores (Fig. 4C,D). These broader score gradients are consistent with scOPE measuring a continuous transcriptional phenotype, whereas expressed-mutation annotations represent sparse observations of driver-specific transcripts. By contrast, only two cells carried an unambiguous mutant *IDH2* annotation, preventing a stable spatial comparison and illustrating the limited interpretability of localization when positive labels are rare (Fig. 4E).

CRC provided a complementary but weaker level of genotype context (Fig. 4F–J). The *TP53* program, which showed FDR-supported patient-level separation, formed regional score gradients within areas populated by mutation-positive tumors rather than uniformly coloring every cell from those patients (Fig. 4G). *APC* scores were likewise localized to selected manifold regions, consistent with the stronger patient-level association observed after restricting the analysis to near-diploid tumors (Fig. 4H). The *BRAF* and *SMAD4* maps provided boundary examples in which the spatial score patterns did not simply recapitulate patient mutation status (Fig. 4I,J).

Together, these maps show that scOPE transfers spatially structured, context-dependent driver-associated phenotypes rather than binary per-cell genotype labels. Concordant localization identifies the cellular states in which a bulk-derived driver program is most strongly expressed, whereas partial or discordant localization exposes within-patient heterogeneity and potential lineage or cell-state contributions that are obscured by cohort-level summary statistics. Complete AML and CRC driver triptychs are provided in Supplementary Fig. S7.

### 2.7 The *NPM1* program contracts during treatment in multiple AML patients

Longitudinal samples provide a second test that does not depend on complete per-cell variant capture. For each patient and time point, we quantified the separation between the driver score and a healthy bone-marrow reference using Cliff’s *δ*. The *NPM1* program was the clearest example because it had strong bulk performance, direct mutation-label support, and repeated treatment-associated changes (Fig. 5A). The largest first-to-last contractions occurred in AML997 (Δ*δ* = −0.840; 0.706 to −0.134), AML556 (−0.529; 0.634 to 0.104), and AML329 (−0.371; 0.551 to 0.180). In each case, the high-score distribution present at the initial sample was markedly reduced at the later treatment time point (Fig. 5B–D).

**Figure 5.**
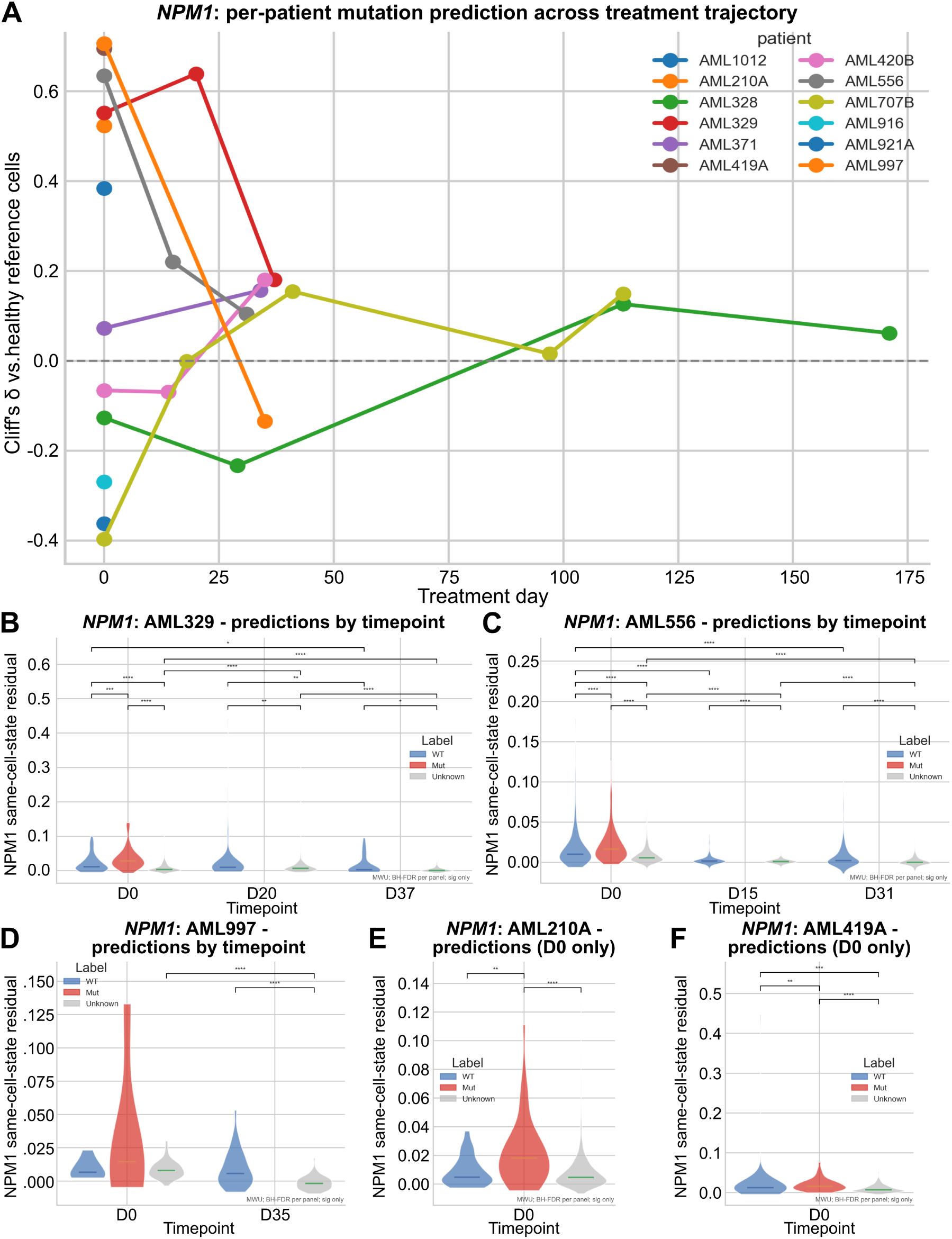
The transferred *NPM1* program contracts during treatment in multiple AML patients. (A) Per-patient effect sizes relative to healthy bone-marrow references. (B–D) Cell-level score distributions at successive time points for AML329, AML556, and AML997. (E–F) Cell-level score distributions at diagnosis for AML210A and AML419A. The complete trajectory summaries for the remaining drivers, together with additional *NPM1* patient distributions, are provided in Supplementary Fig. S6; the exhaustive per-patient, per-driver violin and density set is retained in `Supplemental_Code.zip`.

Not every patient followed the same trajectory, and this heterogeneity is expected in a cohort spanning different genotypes, treatments, response states, and sampling intervals. For example, first-to-last *NPM1* effects increased in AML707B (Δ*δ* = 0.545), AML420B (0.246), and AML328 (0.188), rather than contracting. Several other drivers, including *FLT3*, *RUNX1*, and *TET2*, also showed large longitudinal shifts in individual patients, but lacked comparably strong direct-label support. We therefore treat those changes as exploratory driver-associated state dynamics; the complete patient-by-driver trajectory summary, including trend tests and evaluation status, is provided in Supplementary Table S7. Diagnosis-only samples AML210A and AML419A were retained as cross-sectional guardrails (Fig. 5E,F): they show how mutant-, wild-type-, and unlabeled cells contribute to the baseline score distribution even when a within-patient trajectory is unavailable. A decreasing score can reflect contraction of a mutant clone, depletion of a correlated differentiation state, or both; longitudinal behavior strengthens the *NPM1* result but does not by itself establish clonal identity.

### 2.8 Transferred programs occupy structured, non-uniform cellular contexts

Beyond the genotype-aligned examples above, a transferable program should have interpretable context across the full cellular composition of each tumor. We therefore visualized cell-state-residual scores on expression-derived UMAPs and summarized them by annotated cell type (Fig. 6). In AML, the highest-confidence programs occupied distinct regions rather than uniformly coloring the entire embedding (Fig. 6A–C). *NPM1*, *TP53*, and *DNMT3A* showed different patterns across immature, myeloid, erythroid, and immune compartments, consistent with the idea that the bulk-trained models capture partially overlapping but nonidentical malignant-state axes (Fig. 6D). Residualization reduced broad healthy-reference effects, but did not erase biologically meaningful lineage structure.

**Figure 6.**
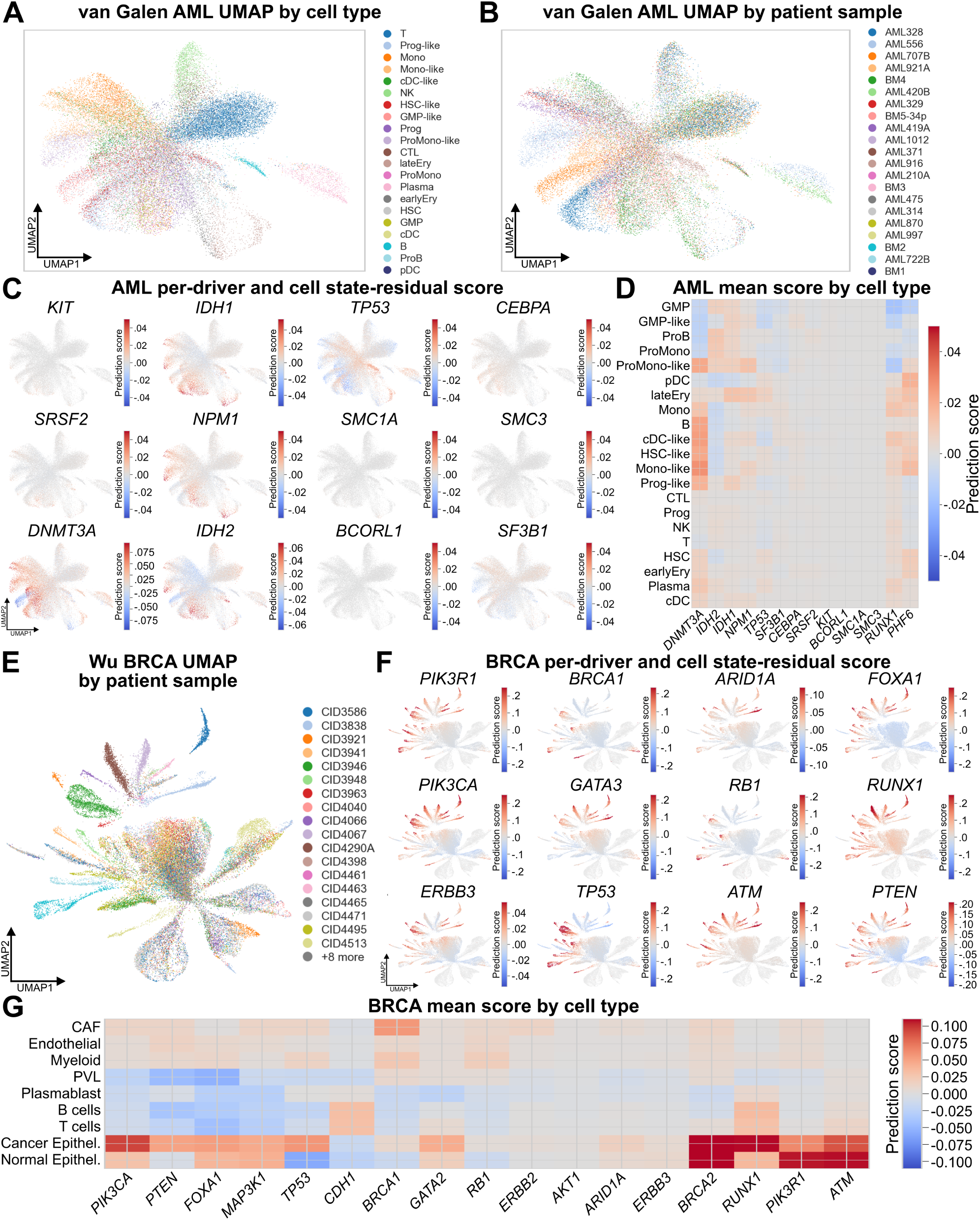
Transferred driver programs resolve structured cellular phenotypes. (A–B) AML expression-space UMAPs colored by cell type and patient sample. (C) Cell-state-residual localization for selected AML programs. (D) Mean residual score by AML cell type and driver. (E) BRCA expression-space UMAP colored by patient sample. (F) Residual-score localization for selected BRCA programs. (G) Mean residual score by BRCA cell type and driver, illustrating both tumor-epithelial enrichment and residual lineage structure. Equivalent context and score atlases for CRC, GBM, LUAD, PAAD, and SKCM are in Supplementary Fig. S8.

The same analysis across solid tumors revealed both coherent and cautionary patterns. Several programs were concentrated in epithelial or malignant-annotated compartments, whereas others remained elevated in normal epithelial or stromal populations (Fig. 6E–G). These patterns are not treated as independent validation: spatial localization can reflect a mutation-linked state, a cancer lineage, tumor purity in the source bulk cohort, or a shared stress program. Instead, the UMAP and cell-type atlases make the transferred phenotype inspectable and expose precisely the confounding that a single scalar performance metric would hide. Cellular-state context, score, and cell-type heatmap outputs for the five cohorts not shown here (CRC, GBM, LUAD, PAAD, and SKCM) are reported in Supplementary Fig. S8; the AML and CRC driver triptychs are in Supplementary Fig. S7.

### 2.9 Copy-number burden provides complementary evidence and exposes discordant populations

We compared scOPE with an infercnvpy-derived malignant-state axis using cancer-specific non-malignant references. The continuous maximum locked-driver score correlated positively with CNV burden in every cohort (Table 4; Fig. 7A). Concordance was strongest in GBM (*ρ* = 0.787; 24,131 cells) and BRCA (0.761; 39,998 cells), followed by PAAD (0.598), AML (0.559), LUAD (0.321), SKCM (0.268), and CRC (0.065). Given the very large cell counts, all correlations were nominally significant; effect size, not the *p*-value, is the relevant comparison. Complete four-way discordance counts and group-wise mean scores for every cohort are reported in Supplementary Table S5.

**Table 4.** Continuous and thresholded agreement between scOPE and inferred CNV burden. ARI is calculated from the binary malignant calls shown in the confusion matrices of Fig. 7B,C and Supplementary Fig. S9A.

| Dataset | Cells | Spearman $\rho$ | $p$ | Binary ARI |
| --- | --- | --- | --- | --- |
| GBM Neftel | 24,131 | 0.787 | $< 10^{-300}$ | 0.079 |
| BRCA Wu | 39,998 | 0.761 | $< 10^{-300}$ | 0.477 |
| PAAD Moncada | 3,659 | 0.598 | $< 10^{-300}$ | -0.075 |
| AML van Galen | 35,555 | 0.559 | $< 10^{-300}$ | 0.287 |
| LUAD Lambrechts | 40,000 | 0.321 | $< 10^{-300}$ | -0.006 |
| SKCM Jerby-Arnon | 7,186 | 0.268 | $2.6 \times 10^{-118}$ | 0.261 |
| CRC Lee | 40,000 | 0.065 | $8.0 \times 10^{-39}$ | 0.017 |

**Figure 7.**
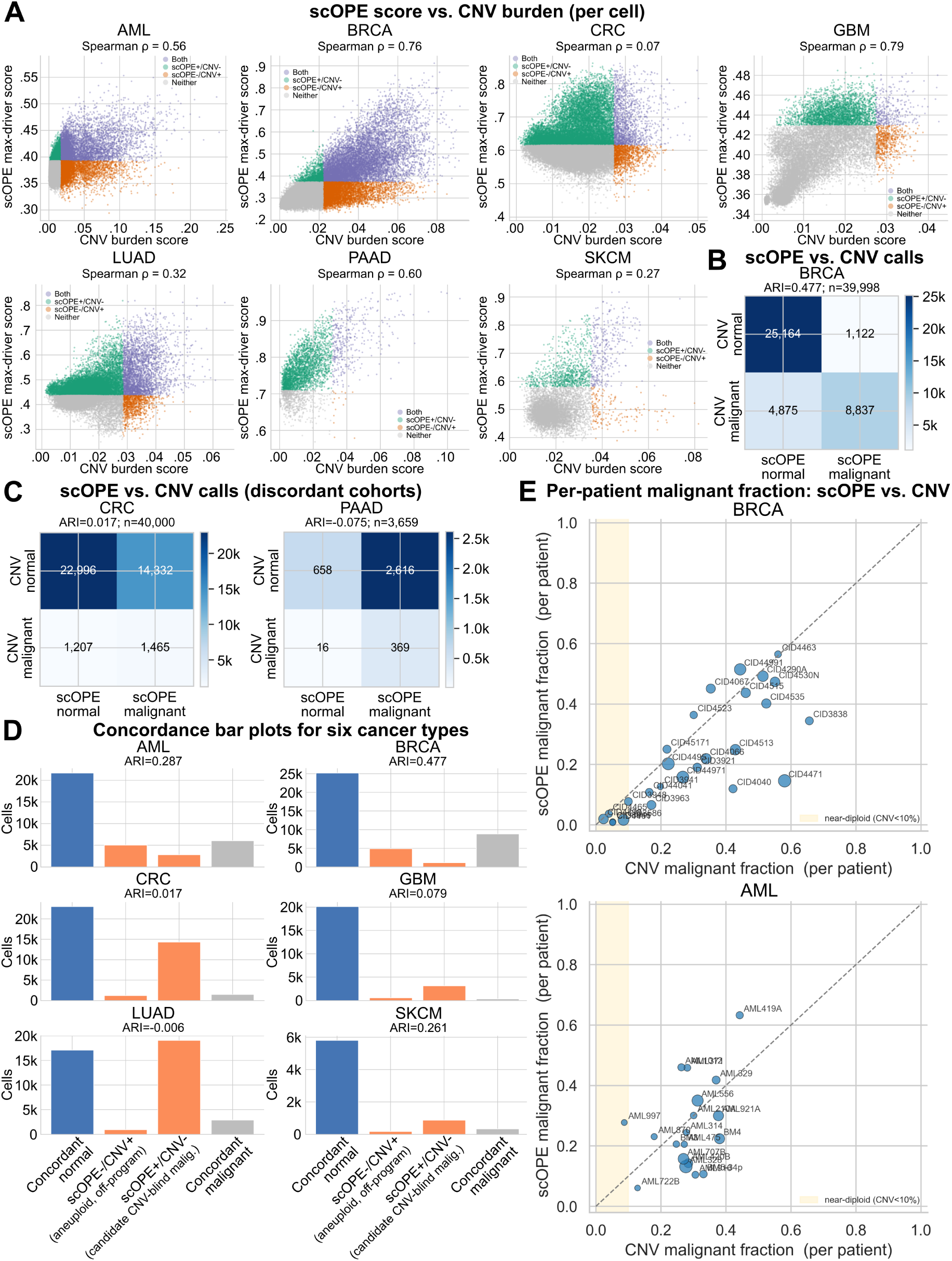
scOPE and inferred CNV capture overlapping but nonidentical malignant-state axes. (A) Continuous per-cell concordance across seven cancers, colored by thresholded agreement category. (B) BRCA confusion matrix, the strongest thresholded-agreement example. (C) CRC and PAAD confusion matrices illustrate discordant cohorts. (D) Counts in the concordant-normal, CNV-only, scOPE-only, and concordant-malignant quadrants. (E) Patient-level malignant fractions in BRCA and AML; shaded regions mark patients with inferred-CNV malignant fraction below 10%. CNV is treated as complementary expression-derived evidence rather than absolute ground truth.

Thresholded agreement was more heterogeneous than the continuous correlations. The adjusted Rand index (ARI) ranged from −0.075 in PAAD to 0.477 in BRCA, demonstrating that the two methods should not be treated as interchangeable binary classifiers (Fig. 7B–D); confusion matrices and patient-level malignant fractions for the cohorts not displayed in Fig. 7 are provided in Supplementary Fig. S9. In BRCA, 25,164 cells were concordant normal and 8,837 concordant malignant, whereas 4,875 were CNV-positive/scOPE-negative and 1,122 were scOPE-positive/CNV-negative. Mean scOPE score increased from concordant-normal cells (0.297) to the scOPE-positive/CNV-negative group (0.403) and concordant-malignant cells (0.478; Fig. 7B). The discordant quadrant is compatible with copy-neutral malignant programs, CNV-blind malignant cells, thresholding differences, or false-positive scOPE scores; marker, pathology, and ideally matched DNA data are needed to distinguish these explanations.

The low CRC correlation is equally informative (Fig. 7A,C). scOPE classified many cells along an epithelial driver-associated expression axis that was not accompanied by broad inferred aneuploidy. More generally, CNV methods detect dosage imbalance, whereas scOPE detects similarity to a bulk genotype-associated transcriptomic program. Concordance therefore provides complementary evidence, while discordance identifies the biological and technical boundary between the two measurement axes.

We additionally asked whether patient-level driver separation persisted in tumors with a low inferred-CNV malignant fraction (*<* 10%). In this near-diploid CRC subset, *APC* (14 mutation-positive versus 11 comparison patients; *δ* = 0.623, FDR *q* = 0.019) and *TP53* (12 versus 13; *δ* = 0.654, *q* = 0.019) remained separated, whereas *KRAS* and *SMAD4* did not (Supplementary Table S6). Near-diploid AML and BRCA comparisons involved at most two mutation-positive patients per driver and are reported for completeness only. The CRC result argues that at least some transferred programs are not wholly explained by large-scale aneuploidy, but it remains a small, secondary analysis rather than proof of copy-neutral clone identity.

## 3 Discussion

scOPE addresses a common asymmetry in cancer single-cell studies: genotype-rich bulk cohorts are abundant, whereas the single-cell datasets needed to resolve malignant phenotypes often lack matched genotyping. By fixing a cancer-specific latent transfer map after bulk training, scOPE converts matched bulk genotype and expression into a continuous coordinate that can be evaluated in existing scRNA-seq cohorts. The principal finding is not that mutations are broadly inferable from expression. It is the opposite: most audited driver–cancer combinations did not yield a transferable program, a minority of canonical drivers did, and the difference between the two is measurable in advance of any genotype label.

This reframing is what separates scOPE from earlier bulk expression–genotype models (Way et al., 2017; Grzadkowski et al., 2021; Davis et al., 2018). Those studies established that expression predicts selected alterations within the bulk domain. The harder question for single-cell biology is not average predictability but per-program admissibility: given an archived scRNA-seq cohort that will never be genotyped, which driver axes may be read, and which must be withheld? By freezing the transfer map, refusing to refit it on the target cohort, and reporting every failed and class-limited evaluation alongside the successful ones, scOPE makes that question answerable rather than assumed.

The AML results illustrate the value of an evidence hierarchy. *NPM1* was strong in held-out bulk tumors, enriched in directly mutant-transcript-labeled cells, localized to specific hematopoietic states, and contracted in multiple longitudinal samples. No single analysis is definitive, but their agreement makes *NPM1* the most compelling example of a transferable mutation-associated program. Matched genotype–score maps added a cellular-resolution spatial interpretation: supported programs occupied restricted regions of the manifold and remained heterogeneous within mutation-positive samples. *TP53* and *DNMT3A* also showed direct-label enrichment, although their cellular context and sample sizes differ. Conversely, weak, diffuse, and inverted maps for other drivers establish an important negative boundary: a high score can represent lineage, malignancy, co-mutation, or cohort composition rather than the target mutation itself.

The ground-truth-free confidence score formalizes this distinction. It asks whether a transferred program is strong in bulk, coherent in single-cell space, concentrated in a subset of cells, and compatible with a complementary expression-derived CNV axis. Its ability to prioritize the directly supported AML programs is encouraging, but the moderate correlation with truth and the high ranking of some unsupported drivers show why it must remain a triage score. In future applications, confidence can define an abstention policy: low-confidence programs should not be interpreted, moderate-confidence programs should be reported as exploratory, and high-confidence programs should still require independent or complementary validation.

The CNV comparison further clarifies method scope. Strong continuous concordance in GBM and BRCA indicates that driver-associated expression states often track aneuploid malignant cells. The persistence of *APC* and *TP53* patient-level effects in a small near-diploid CRC subset suggests that some transferred programs are not wholly reducible to large-scale CNV. Variable binary agreement and large discordant populations are not necessarily failures, because the methods measure different biology. Expression-based CNV inference is sensitive to reference choice, dataset structure, and the magnitude of dosage alterations (Schmid et al., 2025). scOPE can, in principle, respond to copy-neutral point-mutation programs, but can also be confounded by lineage and purity. Accordingly, the scOPE-positive/CNV-negative quadrant should be framed as a testable candidate population rather than evidence of copy-neutral malignancy by itself.

Several limitations define the work that is required for a mutation-specific claim. First, bulk labels are associated with tumor composition, molecular subtype, co-mutation, and purity; a classifier can exploit any of these correlates. Cell-state residualization reduces broad reference effects but cannot guarantee causal or cell-intrinsic specificity. Second, direct AML mutation labels are sparse and nested within patients. Patient-level aggregation reduces pseudoreplication and retained the *NPM1* association, but the number of genotype-evaluable patients remains small. Patient-blocked permutation or a mixed-effects analysis would further strengthen population-level inference. Third, rare bulk classes can yield unstable estimates even under cross-validation, making class counts, AUPRC, confidence intervals, and permutation results essential. Fourth, domain shift between bulk and single-cell assays remains a fundamental limitation: gene overlap, normalization, sequencing platform, and cell-type composition all influence projection. Finally, the confidence score and CNV validation must be kept analytically noncircular; CNV-derived confidence components cannot be used to construct the score that is subsequently tested against CNV.

The operating interpretation of a scOPE score is therefore deliberately narrower than “mutation probability.” Within a cancer type, the score ranks cells by similarity to a bulk-derived driver-associated transcriptional program. A high score is most persuasive when bulk discrimination is robust, the program is not dominated by healthy-reference or lineage effects, direct or patient-level genotype evidence agrees in direction, and an independent assay or complementary analysis supports the implicated population. When those layers disagree, the disagreement is itself useful: it identifies a program that is composition-mediated, context-dependent, technically unstable, or biologically uncoupled from the available genotype annotation.

These constraints define a practical role for scOPE. It is a discovery and prioritization framework for mutation-associated phenotypes in scRNA-seq cohorts that cannot otherwise be genotyped, of which a large fraction of published cancer atlases are examples. Its immediate uses are concrete: nominating cells for targeted variant recovery, selecting samples worth matched DNA profiling, defining populations for spatial or protein validation, and supplying a principled covariate for clone-aware differential expression. Because the confidence score is computed without labels, it can be applied to a new cohort before any validation is commissioned, which is precisely when prioritization has value. A natural extension is a jointly calibrated model that incorporates transcriptome, chromatin, protein, CNV, spatial context, and sparse variant reads while explicitly separating driver identity from lineage and patient effects. Until such data are available, scOPE provides a transparent intermediate layer: more specific than a generic malignant-cell score, broader than an allele call, and accompanied by diagnostics that reveal when the interpretation is likely to fail.

## 4 Methods

### 4.1 Study design and evidence hierarchy

The study was organized to prevent single-cell validation labels from influencing model training. Bulk preprocessing, latent-space fitting, and driver-classifier estimation were performed separately for each cancer. The fitted gene set, bulk means and standard deviations, latent loadings, and classifier parameters were then frozen. Single-cell mutation labels were used only for downstream evaluation in AML. The claim hierarchy was: (i) recoverability in held-out bulk tumors; (ii) ground-truth-free transfer confidence; (iii) direct single-cell mutation-label enrichment where available; (iv) longitudinal behavior; (v) localization to annotated cellular states; and (vi) concordance with a complementary expression-derived CNV axis. No single layer was treated as sufficient evidence of per-cell genotype.

### 4.2 Bulk cohorts, mutation labels, and candidate drivers

AML models used BeatAML expression and mutation resources (Tyner et al., 2018; Bottomly et al., 2022). Solid-tumor models used cancer-specific RNA-seq and somatic mutation calls derived from The Cancer Genome Atlas (TCGA) for BRCA, CRC, GBM, LUAD, PAAD, and SKCM (Cancer Genome Atlas Network, 2012b; Cancer Genome Atlas Network, 2012a; Verhaak et al., 2010; Cancer Genome Atlas Research Network, 2014; Cancer Genome Atlas Research Network, 2017; Cancer Genome Atlas Network, 2015). The source projects are identified at project level as BeatAML (GDC project BEATAML1.0-COHORT; dbGaP phs001657), TCGA-BRCA, TCGA-COAD/TCGA-READ, TCGA-GBM, TCGA-LUAD, TCGA-PAAD, and TCGA-SKCM. The frozen archive preserves the exact derived input-matrix filenames and numerical configuration, but the original portal-specific download manifests and file UUIDs used to construct those previously derived matrices were not retained. Supplementary Table S2 therefore reports project-level source identifiers rather than unsupported file-level accessions. Models were trained independently within cancer type. The source matrices contained up to 651 AML, 903 BRCA, 408 CRC, 155 GBM, 567 LUAD, 172 PAAD, and 466 SKCM tumors; driver-specific sample counts were smaller when mutation labels were unavailable or a cohort-specific subset was required. The audit considered 33 AML, 22 BRCA, 25 CRC, 17 GBM, 20 LUAD, 15 PAAD, and 26 SKCM driver labels. The v10.3.2.2 run used an exploratory all-label mode and attempted training when the mutation-positive fraction was at least 0.01; labels below this threshold or lacking both classes were retained as explicit skipped entries.

For each driver, the audit reports the mutation-positive and comparison counts among valid pooled out-of-fold predictions, fold-level prediction omissions, performance metrics, permutation results, and any skip reason. A result was designated claim-safe only when at least five mutation-positive tumors were represented among valid pooled out-of-fold predictions, with a finite AUROC, a 95% AUROC confidence-interval lower bound greater than 0.5, and a Benjamini–Hochberg-adjusted permutation *q <* 0.05. Claim safety was assigned before biological interpretation.

### 4.3 Expression preprocessing and gene harmonization

Bulk and single-cell expression matrices were harmonized to a common gene-symbol namespace within each cancer type. For the AML bulk cohort, Ensembl version suffixes were removed before mapping Ensembl identifiers to GENCODE v44 gene names. Genes that could not be matched to a GENCODE gene record, as well as records for which the mapped `gene_name remained` an Ensembl-like placeholder rather than a readable gene symbol, were excluded. A complete audit of excluded identifiers was retained by the preprocessing script.

Duplicate mapped gene symbols were not summed, averaged, or otherwise numerically collapsed in the canonical analysis. Instead, all mapped columns were retained as separate features. The first occurrence retained the unsuffixed gene symbol, and subsequent occurrences were assigned deterministic suffixes of the form `--2, --3`, and so forth, in their original column order. Thus, for example, two columns mapping to the same symbol were represented as `GENE` and `GENE_ _ 2`. This rule prevented expression values from distinct source identifiers from being combined after symbol mapping while ensuring that every matrix column had a unique identifier.

The supplied bulk expression matrices had already undergone cohort-specific normalization and log transformation upstream. The canonical analysis therefore applied no additional library-size normalization or logarithmic transformation (`norm_method=“none“` and `log1p=false`). Within each cancer type, the fitted bulk preprocessor established the ordered feature set and stored the bulk-derived transformation parameters used during model fitting. This same fitted pre-processing object and feature order were subsequently used during single-cell alignment and projection. The canonical `AML model used the symbol-mapped, duplicate-preserving matrix AML_bulk_expression_samples_by_genes_SYMBOLS_MAPPED_ONLY.tsv.gz`.

### 4.4 Latent-space construction and driver classifiers

For cancer *c*, the processed bulk matrix *X_c_*∈ R*^nc×pc^* was standardized and factorized by truncated singular value decomposition,

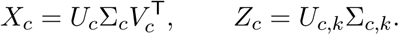

The columns of *V_c,k_* define the fixed bulk-derived gene loadings and *Z_c_* the sample latent representation. The canonical figure-generation archive was `scope_analysis_v10.3.2.2`. Bulk models fitted within that run used *k* = 30 components and L2-regularized logistic regression with *C* = 1.0, class weighting set to “balanced”, the liblinear solver, 2,000 maximum iterations, and random seed 67. These settings generated the bulk audit, interpretability outputs, and the bulk-derived transferability component used by the confidence analysis.

The canonical single-cell analysis was executed as a fresh forced rescore using `drivers/config_ canonical_v3.json`. Bulk pipelines were refit, curated single-cell expression objects were reprojected through the corresponding 30-component bulk models using `moment_matching`, and all raw `mutation_prob_*` columns were regenerated. The run used `–force-rescore-singlecell, –overwrite-scored-h5ad`, and `–clean-output-root`, so the newly scored objects replaced the configured target H5ADs and stale figure/table outputs were removed. Historical z_score_bulk strings in those target filenames are legacy path names and do not describe the alignment used by this run. At the start of downstream analysis, stale calibrated, residual, percentile, and confidence-adjusted columns were cleared and regenerated from the fresh raw scores using the canonical references and settings. Claim-bearing single-cell, patient-bridge, localization, and longitudinal comparisons used `mutation_resid_*`; the CNV comparison used an unadjusted maximum over raw `mutation_prob_*` columns. The archived score-audit label `loaded_existing_scored_h5ad` reflects a subsequent reload of the overwritten score objects and should not be interpreted as evidence that an older alignment or model produced the claim-bearing scores. The exact command is archived with the analysis package.

For interpretability, component importance was summarized using the absolute logistic coefficient multiplied by the corresponding singular value. Component ablation retained the top-ranked components in increasing number and used a separate driver-specific evaluator on the full-cohort fitted latent representation. Gene-loading tables, component-label correlations, SHapley Additive exPlanations (SHAP) feature-attribution summaries (Lundberg and SI Lee, 2017), calibration plots, and biplots were exported as diagnostics; they are not used as independent validation or as replacements for the leakage-free pooled out-of-fold benchmark.

### 4.5 Bulk cross-validation, performance metrics, and model audit

Bulk discrimination used one shared set of five shuffled folds (random seed 67), stratified by the most balanced available driver label so that all drivers were evaluated on a common partition. Within every fold, a new bulk pipeline was fit exclusively to the training tumors: fitted preprocessing, standardization, truncated SVD, and eligible driver classifiers therefore never used the held-out samples. The fitted fold-specific pipeline was then applied unchanged to the held-out tumors. If a driver classifier was not trained in a fold because one class was absent or its training prevalence was below 0.01, predictions for that driver–fold combination remained missing. AUROC, AUPRC, and reported class counts were calculated from the valid pooled out-of-fold predictions, with AUPRC interpreted relative to positive-class prevalence (Saito and Rehmsmeier, 2015). A separate driver-specific evaluator, used only for interpretability plots such as calibration, permutation, and component-ablation panels, adapted the number of folds to rare class counts and operated on the full-cohort fitted latent representation; its diagnostic AUROCs can therefore differ slightly from the claim-bearing pooled out-of-fold estimates. Brier score quantified probabilistic error (Brier, 1950). The archived configuration specifies 2,000 bootstrap replicates for confidence intervals and 2,000 label permutations for null assessment. Bootstrap AUROC intervals used case–control-stratified resampling with replacement, preserving the observed number of positive and negative tumors in each replicate; 95% intervals were the 2.5th and 97.5th percentiles of the valid bootstrap distribution. Label-permutation *p*-values were one-sided empirical probabilities with the (*b* + 1)*/*(*n*_perm_ + 1) correction, where *b* is the number of permuted AUROCs at least as large as the observed AUROC, and used random seed 67. Permutation *p*-values were adjusted with the Benjamini–Hochberg procedure (Benjamini and Hochberg, 1995).

### 4.6 Single-cell cohorts

The transferred models were evaluated in seven publicly documented scRNA-seq cohorts: AML (van Galen; GEO GSE116256), BRCA (Wu; GEO GSE176078; controlled raw data EGAS00001005173), CRC (Lee; GEO GSE132465), GBM (Neftel; GEO GSE131928), LUAD (Lambrechts; BioStudies/ArrayExpress E-MTAB-6149), PAAD (Moncada; GEO GSE111672), and SKCM (Jerby-Arnon; GEO GSE115978) (van Galen et al., 2019; Wu et al., 2021; HO Lee et al., 2020; Neftel et al., 2019; Lambrechts et al., 2018; Moncada et al., 2020; Jerby-Arnon et al., 2018). The analysis used processed, quality-controlled, normalized, and log-transformed expression objects; internal filtering, renormalization, log transformation, and doublet detection were disabled during rescoring. Final retained cell counts were 35,843 AML, 39,998 BRCA, 40,000 CRC, 24,131 GBM, 40,000 LUAD, 3,659 PAAD, and 7,186 SKCM. Large cohorts were subsampled with a fixed seed and, where patient labels were available, proportional representation by patient. Supplementary Table S2 reports source publications and DOIs, repository and project accessions, processed/raw-data access status, analysis-input provenance, retained and CNV-matched cell counts, reference-cell counts, and the source annotation categories used as references.

### 4.7 Fixed bulk-to-single-cell projection

Let *X*_sc_ ∈ R*^m×p^*^sc^ denote the expression matrix for a single-cell cohort from the same cancer type as the corresponding fitted bulk model. The canonical analysis used the `SingleCellPipeline` implementation in `scope-bio v0.2.0`, with `alignment_method=“moment_matching“`. The input single-cell AnnData objects were treated as previously quality-controlled, normalized, and log-transformed. Consequently, no additional cell filtering, normalization, logarithmic transformation, or doublet detection was performed during the forced-rescore analysis.

Gene matching, ordering, and construction of the aligned single-cell feature matrix were performed by the pinned scope-bio single-cell pipeline using the fitted bulk model and the supplied single-cell object. The manuscript-specific scope_analysis v10.3.2.2 wrapper imposed no additional numerical minimum-gene-overlap threshold. Genes not represented as matched features in the fitted projection did not contribute observed expression information to the projected representation, and the analysis wrapper did not perform expression-level imputation of absent genes.

For the matched features, moment matching aligned the gene-wise location and scale of the single-cell expression distribution to those of the corresponding preprocessed bulk reference. This alignment was label-free but target-cohort-dependent: it used gene-wise moments estimated from the supplied single-cell cohort, while leaving the bulk SVD loadings and classifier parameters unchanged. Denoting the resulting aligned single-cell matrix by 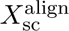, cells were projected through the frozen bulk-derived truncated-SVD loadings *V_k_*:

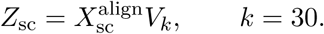

The bulk SVD loadings and driver classifiers remained fixed throughout this step. No single-cell mutation labels, patient labels, cell-type annotations, reference labels, or embedding coordinates were used to refit *V_k_*, select components, or update the classifier parameters. Each fitted driver-specific logistic classifier was applied directly to *Z*_sc_, producing the raw per-cell score column `mutation_prob_<DRIVER>`.

Before each classifier prediction, the projected latent matrix was checked for nonfinite values. Under the configured `singlecell_projection_nan_policy=“zero_fill“`, any `NaN`, positive-infinite, or negative-infinite latent coordinate was replaced by zero, which represents the centered or neutral value of that bulk latent component. The same rule was applied to nonfinite entries in stored projected latent matrices after transformation so that downstream neighborhood, embedding, and plotting procedures received finite coordinates. Importantly, this `zero_fill` policy applied only to nonfinite values in the projected latent representation; it was not a rule for imputing genes absent from the original single-cell expression matrix.

The canonical `scope_analysis v10.3.2.2` run regenerated this entire projection from the con-figured curated AnnData objects under moment matching. Previously stored `mutation_prob_*`, calibrated, adjusted, residual, and reference-percentile score columns were removed before rescoring. Fresh raw probabilities were then generated from the newly projected latent coordinates and used to reconstruct all derived score columns reported in the manuscript.

### 4.8 Healthy-reference guardrails and cell-state-residual scores

Raw transferred probabilities can be high in healthy cells when the bulk signature tracks a lineage represented differently across assays. Reference cells were therefore identified from cohort-specific source annotations while excluding malignant, epithelial, and ambiguous compartments. In AML, 5,946 cells from BM1, BM2, BM3, BM4, and BM5-34p formed the primary reference set. The BM5 CD34^+^CD38*^−^* (“34p38n”) subset and AML cell lines were excluded; the exact case-insensitive sample regular expression is archived in Supplementary Methods.

For driver *d* in cohort *c*, let *p_id_* be the raw score of cell *i*, *s*(*i*) its canonicalized cell state, and *R_c_* the cohort reference cells. The cell-state-residual score was computed separately within each cohort as

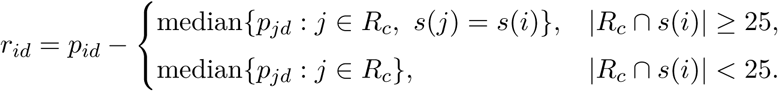

Thus, states represented by at least 25 reference cells used a same-state reference median; otherwise the cohort-wide reference median was the explicit fallback. No shrinkage or cross-cohort pooling was applied. Raw probabilities were retained for score audits and visualization but were not interpreted as allele probabilities. The numerical single-cell, patient-bridge, and longitudinal results reported here use the supplied “mutation_resid_*” tables.

### 4.9 Ground-truth-free driver confidence

A composite driver-level confidence score was computed without per-cell mutation labels from four components evaluated on raw transferred probabilities. Bulk transferability was *T* = max[0.05, min{1, 2(AUROC_bulk_ − 0.5)}]. Spatial coherence *S* was Moran’s *I* on the expression-neighbor graph, clipped to [0, 1]. Concentration *C* was the Gini coefficient of the raw score across all retained cells, clipped to [0, 1]. CNV-axis agreement was *X* = min{1, max[0, 2(AUROC_CNV_ − 0.5)]}, where the score was used to classify the separately inferred expression-derived CNV label. Available components were combined as a weighted geometric mean,

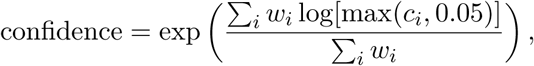

with weights *w_T_* = 1, *w_S_* = 1.5, *w_C_* = 1, and *w_X_* = 1. These canonical weights were fixed in the archived implementation and were not estimated or optimized against the AML mutation-label results. Components that could not be computed were omitted and the remaining weights renormalized. Display tiers were low (*<* 0.2), moderate (0.2–*<* 0.4), high (0.4–*<* 0.6), and very high (≥ 0.6). Confidence-adjusted visualization columns were calculated as raw probability multiplied by driver confidence; inferential comparisons used the explicitly named raw or residual score specified for each analysis. Confidence was validated in AML by merging the driver-level table with direct-label effect sizes. Spearman correlation quantified ranking agreement, and AUROC assessed whether confidence separated drivers with positive Cliff’s *δ* and FDR *q <* 0.05 from the remaining truth-evaluable drivers. This internal validation involved 12 drivers, of which three met the support definition, and was interpreted as a ranking check rather than an external performance benchmark.

### 4.10 AML mutation-label and patient-level validation

The processed van Galen AML object contained per-cell lists of detected mutant and wild-type transcripts for a subset of evaluated drivers (van Galen et al., 2019). For driver *d*, a cell was assigned a reported-mutant label when *d* appeared in its mutant-transcript list but not its wild-type transcript list. A cell was assigned a reported-wild-type comparison label when *d* appeared in its wild-type transcript list but not its mutant-transcript list. Cells in which the driver appeared in both lists, in neither list, or for which the corresponding score was nonfinite were excluded from the direct per-cell comparison for that driver.

Because failure to detect a variant transcript in a single cell does not establish genomic wild type, these comparisons were interpreted as positive-unlabeled validation. Discrimination was therefore reported as apparent AUROC rather than as a fully supervised genomic-classification AUROC. For each evaluable driver, the analysis calculated apparent AUROC with a 95% bootstrap confidence interval, the two-sided Mann–Whitney *U* statistic, and Cliff’s *δ* with a bootstrap confidence interval (Mann and Whitney, 1947; Cliff, 1993). Driver-level Mann–Whitney *p*-values were adjusted by the Benjamini–Hochberg procedure within the family of AML drivers having evaluable per-cell labels.

A complementary patient-level bridge was used to reduce within-patient pseudoreplication. Patient mutation or alteration labels were supplied in tidy tables containing the fields `patient, mutation`, and `label`, where `label=1` represented a reported positive call, `label=0` represented a reported negative call, and missing or unknown calls were excluded.

For AML, patient-level labels were derived from the mutation-transcript truth annotations in the same van Galen single-cell object and were supplemented with the configured healthy bone-marrow reference records. Consequently, this analysis changes the statistical unit and reduces within-patient pseudoreplication, but it is not an independent genotype assay. Within each patient and driver, the patient was called positive when at least one truth-evaluable cell had a reported-mutant label. A patient was called negative only when no reported-mutant cell was present and at least one truth-evaluable cell had a reported-wild-type label. Patients with no evaluable truth annotation for a driver were assigned unknown status and excluded for that driver.

For CRC and BRCA, the canonical run read cohort-specific curated patient-status tables configured for the Lee colorectal-cancer and Wu breast-cancer single-cell cohorts, respectively (HO Lee et al., 2020; Wu et al., 2021). The CRC table contained patient-level calls for the drivers evaluated in the bridge analysis, including *APC*, *KRAS*, *BRAF*, *SMAD4*, and *TP53*. When separate tumor and normal sample records were available for the same CRC patient, the patient was called positive if any matched tumor sample was positive; otherwise, an evaluable negative call was retained, and unresolved cases were treated as unknown. The BRCA status table provided cohort-reported *ERBB2* status for the 26 evaluable primary tumors; no patient-level calls were inferred for other BRCA drivers from population-level mutation frequencies.

Patient identifiers were matched conservatively in three successive tiers: first by exact string equality; second after removing punctuation and standardizing case; and third after removing one recognized, separator-delimited sample-type or time-point suffix. The final tier allowed, for example, `SMC01-T` and `SMC01-N` to match patient `SMC01`, and `AML329-D0` to match patient `AML329`. Characters fused directly to the patient identifier were not removed. A normalized identifier was accepted only when it mapped unambiguously to one patient in the single-cell object; ambiguous or nonoverlapping records were left unmatched and excluded.

For the AML patient bridge, patient identifiers were created by removing the terminal `-D <NUMBER>` time-point suffix from the Sample field. No single AML visit was selected for this analysis. Instead, cells from all retained time points for a patient were pooled, and the median `mutation_resid_<DRIVER>` score was calculated across the patient’s retained cells. The diagnosis-only D0 inclusion rules used for selected longitudinal panels were applied after the patient-bridge stage and therefore did not restrict this patient-level validation.

A patient contributed a median score only when at least five cells with finite scores were available for the corresponding driver. The bridge test was performed only when at least two positive and two negative patients remained. Mutation-positive and mutation-negative patient medians were compared using a two-sided Mann–Whitney test and Cliff’s *δ* with a 95% bootstrap confidence interval. Benjamini–Hochberg correction was applied across the evaluable driver family within each single-cell cohort.

### 4.11 Longitudinal AML analysis

Sample identifiers were parsed into patient and treatment day. The primary trajectory set excluded AML870 and retained the prespecified patient/sample filters recorded in the configuration. For each driver, patient, and time point, the residual-score distribution was compared with the fixed 5,946-cell healthy bone-marrow reference using Cliff’s *δ*. The primary change statistic was Δ*δ* = *δ*_last_ − *δ*_first_. Spearman trend tests were reported when at least three time points were available. Pairwise time-point comparisons in the violin panels used Mann–Whitney tests with within-panel Benjamini–Hochberg adjustment. Drivers lacking direct-label support were interpreted as exploratory state dynamics rather than clone tracking. Cell counts and driver-level annotations for every patient–time-point combination are provided in the exported trajectory tables.

### 4.12 Expression-space embeddings and cell-type summaries

UMAPs (McInnes et al., 2018) used an expression-derived highly variable gene representation rather than the bulk SVD coordinates being evaluated. The canonical command requested 3,000 highly variable genes, 50 principal components, 15 neighbors, minimum distance 0.5, and random seed 42. The automatic preprocessing branch detected the curated objects as already normalized and log-transformed, avoiding a second normalization/log transformation. Reference cells were greyed in score maps. For driver-level triptychs, the same UMAP coordinates were reused for patient identity, mutation context, and score. AML mutation context used the guarded direct per-cell labels described above. CRC mutation context used patient-level genotype assigned to that patient’s cells solely for visualization; no CRC cell was treated as directly genotyped. Residual scores were displayed on a centered symmetric ±0.05 color range for visual comparison, whereas all statistical analyses used the original untruncated values. Drivers were ordered by confidence, and low-confidence programs were suppressed or relegated to the supplement. Cell-type heatmaps report mean cell-state-residual score within source annotation categories using a global centered scale for cross-driver comparison.

### 4.13 CNV inference and complementary comparison

CNV burden was inferred from scRNA-seq expression using infercnvpy with genomic gene positions and dataset-specific non-malignant reference categories (Fan et al., 2018; Sturm, 2024; Schmid et al., 2025). References were drawn from each cohort’s own annotations and excluded malignant, epithelial, and ambiguous compartments. Reference counts were 5,946 AML, 9,529 BRCA, 20,435 CRC, 10,680 GBM, 1,783 LUAD, 208 PAAD, and 618 SKCM cells; category-level counts are reported in Supplementary Table S2. The archived CNV-generation script specifies a 100-gene window, step size 10, log-fold-change clipping at 3, dynamic threshold 1.5, exclusion of chromosomes X and Y, and a 5,000-cell processing chunk. Per-cell CNV burden was the genome-wide mean absolute inferred deviation, and CNV-malignant cells were called above the 95th percentile of the cohort-specific reference burden distribution.

For the complementary comparison, v10.3.2.2 explicitly constructed `scope_max_driver_score` as the maximum raw `mutation_prob_*` value over the cancer’s locked primary-driver set. This was an unadjusted score: neither cell-state residualization nor multiplication by the confidence score entered the CNV comparison. The corresponding binary scOPE call used the 80th percentile of the reference-cell maximum-score distribution (`reference_quantile=0.8`). CNV calls and scOPE scores were joined by dataset and cell barcode using exact and progressively normalized barcode matching, with match diagnostics exported for each cohort. Continuous concordance was quantified by Spearman correlation. Binary agreement was summarized by four-way confusion tables and adjusted Rand index (Hubert and Arabie, 1985); patient-level malignant fractions were also compared. Near-diploid patients were defined as those with an inferred-CNV malignant fraction below 0.10, and matched patient-genotype groups were compared using the same patient-level Mann–Whitney/Cliff’s-*δ* framework. This separation prevents circular validation through the CNV-derived component of the driver-confidence score.

### 4.14 Statistical analysis and reproducibility

All hypothesis tests were two-sided unless an explicitly directional permutation null was specified. Bulk mutation-label permutation tests used the one-sided alternative that the observed AUROC exceeded the label-permuted null. Tumor-versus-reference and mutation-positive-versus-mutation-negative permutation analyses similarly tested whether the observed directed statistic exceeded its permuted null. Empirical permutation *p*-values used the finite-sampling correction (*b* + 1)*/*(*B* + 1), where *b* is the number of permuted statistics at least as large as the observed statistic and *B* is the number of valid permutations. Other group comparisons used two-sided Mann–Whitney *U* or Kolmogorov–Smirnov tests as described above. Effect sizes were reported as Cliff’s *δ*, median differences, or Spearman correlations, with bootstrap confidence intervals where applicable.

Multiple-testing correction used the Benjamini–Hochberg procedure at a false-discovery rate of 0.05. Correction was applied once within each prespecified analysis family, rather than separately within individual figure panels. Bulk permutation *p*-values were adjusted across the evaluable cancer–driver tests in the bulk audit. Single-cell validation, patient-bridge, longitudinal, and pan-cancer analyses were adjusted across the drivers evaluated within the corresponding cohort and analysis stage. Missing or non-evaluable *p*-values were retained as missing and excluded from the denominator used for adjustment.

The canonical analysis was performed in Python 3.10 using scOPE release `scope-bio==0.2.0` and the frozen manuscript-analysis snapshot `scope_analysis_v10.3.2.2`. Release `v0.2.0` corresponds to Git commit `23d50b9`. Core data structures and analyses used AnnData, Scanpy, NumPy, SciPy, pandas, scikit-learn, statsmodels, matplotlib, and infercnvpy (Virshup et al., 2024; Wolf et al., 2018; Pedregosa et al., 2011; Sturm, 2024). The manuscript archive includes a minimal machine-readable environment specification that pins Python and scope-bio; exact build-level versions of additional transitive packages were not recorded by the original run and are therefore not asserted here.

The canonical configuration was `drivers/config_canonical_v3.json`. Exact SHA-256 checksums for the configuration, manifest, and Supplemental Code archive are provided in the distributed `SHA256SUMS.txt` file. Statistical resampling and model evaluation used random seed 67; expression-derived UMAP construction used random seed 42. The configuration requested five shared folds for the claim-bearing pooled out-of-fold benchmark, 2,000 bootstrap replicates, 2,000 permutation replicates, a minimum of five mutation-positive bulk samples represented among valid out-of-fold predictions for claim-bearing inference, and a minimum of five cells for patient-level summaries. The separate interpretability evaluator recorded any rare-driver reduction in its diagnostic cross-validation folds; fold-specific missing predictions in the claim-bearing benchmark were retained in the bulk audit.

The canonical analysis was executed as a clean, forced rescore with the following command:

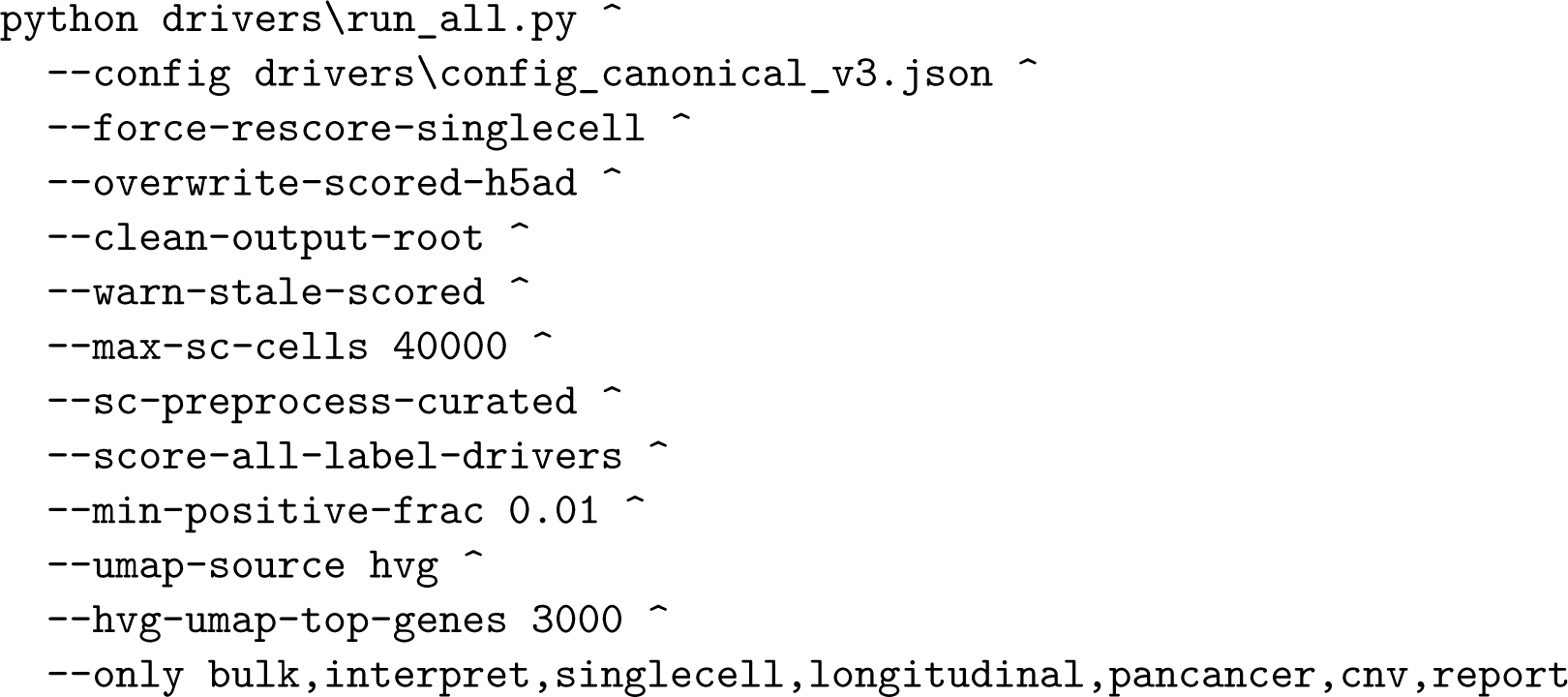

The forced-rescore flags removed stale score columns, refitted the cancer-specific bulk pipelines, reprojected the configured single-cell objects using moment matching, overwrote the target scored AnnData files, cleared the previous output directory, and regenerated the downstream tables and figures. The analysis archive contains the exact configuration, canonical command, output manifest, driver-training audits, single-cell score audits, reference-composition tables, panel-provenance table, and generated result tables.

Negative controls and reference guardrails are summarized in Supplementary Fig. S10 and Supplementary Table S10, and every main-text panel is mapped to its source artifact, score column, statistical unit, correction family, and random seed in Supplementary Table S8. Negative controls present in the canonical output include bulk mutation-label permutations; tumor/reference-label permutation nulls for the pan-cancer score shifts; same-prevalence random patient-label controls where patient genotype metadata were available; reference-composition and reference-direction sensitivity analyses; and latent-dimension sensitivity analyses for every cancer. The supplied code also implements patient-block label permutation for group-level mutation labels and broader decomposition, classifier, and alignment ablations. These implemented but non-claim-bearing utilities are included in `Supplemental_Code.zip` so that additional sensitivity analyses can be reproduced without altering the primary results.

## Supporting information

scOPE_supplement

## Data and code availability

The source studies are publicly documented, but access to the underlying human data is not uniform. Source processed expression matrices and annotations for the single-cell cohorts are available from the repositories listed in Supplementary Table S2. Some raw human sequence data require controlled access (for example, BRCA raw data in EGA), whereas other source records provide processed data without releasing raw reads directly. BeatAML and TCGA source projects likewise contain a mixture of open processed resources and controlled sequence-level data. Supplementary Table S2 reports the source publication and DOI, repository and project accessions, access status, analysis-input description, modeled or retained analysis size, and reference annotations for every bulk and single-cell cohort.

The bulk expression and mutation-label matrices supplied to the canonical analysis were previously derived from the listed BeatAML and TCGA source projects. The frozen archive preserves their exact local filenames, cancer assignment, preprocessing settings, and numerical configuration. The original portal-specific download manifests and file UUIDs used to construct these derived matrices were not retained; accordingly, Supplementary Table S2 reports verified project-level source identifiers rather than guessed file-level accessions. Original cohort expression matrices, raw human sequence data, and fitted AnnData objects are not redistributed in the Supplemental Code archive. The canonical analysis can be rerun once equivalent derived input matrices are constructed from the listed source deposits; the archive preserves the preprocessing assumptions and analysis scripts but not a complete portal-download manifest, and all source-data access terms continue to apply.

The maintained scOPE implementation, installation instructions, documentation, and versioned releases are openly available from:

- **GitHub repository and releases:** https://github.com/Ashford-A/scOPE
- **Python Package Index:** https://pypi.org/project/scope-bio/
- **Conda-forge package:** https://anaconda.org/conda-forge/scope-bio
- **Frozen manuscript-analysis archive:** Supplemental_Code.zip

The numerical analyses reported here used scOPE release `scope-bio==0.2.0`, GitHub tag `v0.2.0`, and commit `23d50b9`. The frozen `Supplemental_Code.zip` archive contains the complete `scope_analysis_v10.3.2.2` directory used for the manuscript, including its source modules, cohort drivers, exact numerical configuration, execution command, statistical and plotting utilities, pathway resources, generated analysis tables, audit tables, negative-control outputs, sensitivity outputs, source-data metadata, and machine-readable manifest. The canonical source identifiers use PAAD Moncada (GSE111672) and SKCM Jerby-Arnon (GSE115978) consistently across the manuscript, configuration, manifests, output filenames, and plotting source. The snapshot identifier `scope_analysis_v10.3.2.2` denotes the manuscript analysis bundle and is distinct from the public `scope-bio` package version.

The Supplemental Code archive is the frozen record of the manuscript-generating analysis, whereas the GitHub repository is the maintained source for installation, documentation, issue tracking, and post-publication updates.

## Acknowledgements

This work was supported in part by the Oregon Health & Science University (OHSU) Advanced Computing Center (ACC) via computational infrastructure support, including Exacloud and the ACC research cluster (ARC). This infrastructure is supported by the Office of Research Infrastructure Programs, Office of the Director, National Institutes of Health, under Award Number S10OD034224. Analyses were conducted using a combination of institutional computing resources and local workstation hardware for profiling experiments. We would like to thank the investigators who generated the publicly available datasets analyzed in this study, as well as the maintainers of public repositories that enable open data sharing and open-source software.

## Author contributions

Conceptualization: AJA, ED. Methodology: AJA, ED, AL. Investigation: AJA, AL. Software: AJA, AL. Formal analysis: AJA. Visualization: AJA. Writing–original draft: AJA, ED. Writing–review & editing: AJA, ED. Supervision: ED.

## References

Andreatta, M and SJ Carmona (2021). “UCell: Robust and scalable single-cell gene signature scoring”. In: Computational and Structural Biotechnology Journal 19, pp. 3796–3798. doi: 10.1016/j.csbj.2021.06.043.

Benjamini, Y and Y Hochberg (1995). “Controlling the false discovery rate: a practical and powerful approach to multiple testing”. In: Journal of the Royal Statistical Society: Series B (Methodological) 57.1, pp. 289–300. doi: 10.1111/j.2517-6161.1995.tb02031.x.

Bottomly, D et al. (2022). “Integrative analysis of drug response and clinical outcome in acute myeloid leukemia”. In: Cancer Cell 40.8, 850–864.e9. doi: 10.1016/j.ccell.2022.07.002.

Brennecke, P, S Anders, J Kim, A Kolodziejczyk, X Zhang, V Proserpio, F Zanini, D McCarthy, J Marioni, and S Teichmann (2013). “Accounting for technical noise in single-cell RNA-seq experiments”. In: Nature Methods 10.11, pp. 1093–1095. doi: 10.1038/nmeth.2645.

Brier, GW (1950). “Verification of forecasts expressed in terms of probability”. In: Monthly Weather Review 78.1, pp. 1–3. doi: 10.1175/1520-0493(1950)078<0001:VOFEIT>2.0.CO;2.

Cai, C, GF Cooper, KN Lu, X Ma, S Xu, Z Zhao, X Chen, Y Xue, AV Lee, N Clark, V Chen, S Lu, L Chen, and X Lu (2019). “Systematic discovery of the functional impact of somatic genome alterations in individual tumors through tumor-specific causal inference”. In: PLoS Computational Biology 15.7, e1007088. doi: 10.1371/journal.pcbi.1007088.

Campbell, KR, A Steif, E Laks, H Zahn, D Lai, A McPherson, H Farahani, F Kabeer, C O’Flanagan, J Biele, J Brimhall, B Wang, P Walters, et al. (2019). “clonealign: statistical integration of independent single-cell RNA and DNA sequencing data from human cancers”. In: Genome Biology 20, p. 54. doi: 10.1186/s13059-019-1645-z.

Cancer Genome Atlas Network (2012a). “Comprehensive molecular characterization of human colon and rectal cancer”. In: Nature 487.7407, pp. 330–337. doi: 10.1038/nature11252.

Cancer Genome Atlas Network (2012b). “Comprehensive molecular portraits of human breast tumours”. In: Nature 490.7418, pp. 61–70. doi: 10.1038/nature11412.

Cancer Genome Atlas Network (2015). “Genomic classification of cutaneous melanoma”. In: Cell 161.7, pp. 1681–1696. doi: 10.1016/j.cell.2015.05.044.

Cancer Genome Atlas Research Network (2014). “Comprehensive molecular profiling of lung adenocarcinoma”. In: Nature 511.7511, pp. 543–550. doi: 10.1038/nature13385.

Cancer Genome Atlas Research Network (2017). “Integrated genomic characterization of pancreatic ductal adenocarcinoma”. In: Cancer Cell 32.2, 185–203.e13. doi: 10.1016/j.ccell.2017.07.007.

Cliff, N (1993). “Dominance statistics: ordinal analyses to answer ordinal questions”. In: Psychological Bulletin 114.3, pp. 494–509. doi: 10.1037/0033-2909.114.3.494.

Dang, L, DW White, S Gross, BD Bennett, MA Bittinger, EM Driggers, VR Fantin, HG Jang, S Jin, MC Keenan, KM Marks, RM Prins, PS Ward, KE Yen, LM Liau, JD Rabinowitz, LC Cantley, CB Thompson, MG Vander Heiden, and SM Su (2009). “Cancer-associated IDH1 mutations produce 2-hydroxyglutarate”. In: Nature 462.7274, pp. 739–744. doi: 10.1038/nature08617.

Davies, H et al. (2002). “Mutations of the BRAF gene in human cancer”. In: Nature 417.6892, pp. 949–954. doi: 10.1038/nature00766.

Davis, RJ, M Gönen, DH Margineantu, and BE Clurman (2018). “Pan-cancer transcriptional signatures predictive of oncogenic mutations reveal that Fbw7 regulates cancer cell oxidative metabolism”. In: Proceedings of the National Academy of Sciences 115.21, pp. 5462–5467. doi: 10.1073/pnas.1718338115.

Dohmen, J, A Baranovskii, J Ronen, B Uyar, V Franke, and A Akalin (2022). “Identifying tumor cells at the single-cell level using machine learning”. In: Genome Biology 23, p. 123. doi: 10.1186/s13059-022-02683-1.

Falini, B et al. (2005). “Cytoplasmic nucleophosmin in acute myelogenous leukemia with a normal karyotype”. In: New England Journal of Medicine 352.3, pp. 254–266. doi: 10.1056/NEJMoa041974.

Fan, J, HO Lee, S Lee, DE Ryu, S Lee, C Xue, SJ Kim, K Kim, N Barkas, PJ Park, WY Park, and PV Kharchenko (2018). “Linking transcriptional and genetic tumor heterogeneity through allele analysis of single-cell RNA-seq data”. In: Genome Research 28, pp. 1217–1227. doi: 10.1101/gr.228080.117.

Figueroa, ME et al. (2010). “Leukemic IDH1 and IDH2 mutations result in a hypermethylation phenotype, disrupt TET2 function, and impair hematopoietic differentiation”. In: Cancer Cell 18.6, pp. 553–567. doi: 10.1016/j.ccr.2010.11.015.

Gao, R, S Bai, YC Henderson, Y Lin, A Schalck, Y Yan, T Kumar, M Hu, E Sei, A Davis, F Wang, SF Shaitelman, JR Wang, K Chen, S Moulder, SY Lai, and NE Navin (2021). “Delineating copy number and clonal substructure in human tumors from single-cell transcriptomes”. In: Nature Biotechnology 39, pp. 599–608. doi: 10.1038/s41587-020-00795-2.

Gao, T, R Soldatov, H Sarkar, A Kurkiewicz, E Biederstedt, PR Loh, and PV Kharchenko (2023). “Haplotype-aware analysis of somatic copy number variations from single-cell transcriptomes”. In: Nature Biotechnology 41, pp. 417–426. doi: 10.1038/s41587-022-01468-y.

Gerlinger, M et al. (2012). “Intratumor heterogeneity and branched evolution revealed by multiregion sequencing”. In: New England Journal of Medicine 366.10, pp. 883–892. doi: 10.1056/NEJMoa1113205.

Greaves, M and CC Maley (2012). “Clonal evolution in cancer”. In: Nature 481.7381, pp. 306–313. doi: 10.1038/nature10762.

Grzadkowski, MR, HD Holly, J Somers, and E Demir (2021). “Systematic interrogation of mutation groupings reveals divergent downstream expression programs within key cancer genes”. In: BMC Bioinformatics 22, p. 233. doi: 10.1186/s12859-021-04147-y.

Hubert, L and P Arabie (1985). “Comparing partitions”. In: Journal of Classification 2.1, pp. 193–218. doi: 10.1007/bf01908075.

Jerby-Arnon, L et al. (2018). “A cancer cell program promotes T cell exclusion and resistance to checkpoint blockade”. In: Cell 175.4, 984–997.e24. doi: 10.1016/j.cell.2018.09.006.

Jha, A, M Quesnel-Vallières, D Wang, A Thomas-Tikhonenko, KW Lynch, and Y Barash (2022). “Identifying common transcriptome signatures of cancer by interpreting deep learning models”. In: Genome Biology 23, p. 117. doi: 10.1186/s13059-022-02681-3.

Kharchenko, PV, L Silberstein, and DT Scadden (2014). “Bayesian approach to single-cell differential expression analysis”. In: Nature Methods 11.7, pp. 740–742. doi: 10.1038/nmeth.2967.

Lambrechts, D, E Wauters, B Boeckx, S Aibar, D Nittner, O Burton, A Bassez, H Decaluwé, A Pircher, K Van den Eynde, B Weynand, E Verbeken, P De Leyn, A Liston, J Vansteenkiste, P Carmeliet, S Aerts, and B Thienpont (2018). “Phenotype molding of stromal cells in the lung tumor microenvironment”. In: Nature Medicine 24.8, pp. 1277–1289. doi: 10.1038/s41591-018-0096-5.

Lee, HO et al. (2020). “Lineage-dependent gene expression programs influence the immune landscape of colorectal cancer”. In: Nature Genetics 52.6, pp. 594–603. doi: 10.1038/s41588-020-0636-z.

Lundberg, SM and SI Lee (2017). “A Unified Approach to Interpreting Model Predictions”. In: Advances in Neural Information Processing Systems 30, pp. 4765–4774.

Mann, HB and DR Whitney (1947). “On a test of whether one of two random variables is stochastically larger than the other”. In: The Annals of Mathematical Statistics 18.1, pp. 50–60. doi: 10.1214/aoms/1177730491.

Marot-Lassauzaie, V, S Beneyto-Calabuig, B Obermayer, L Velten, D Beule, and L Haghverdi (2024). “Identifying cancer cells from calling single-nucleotide variants in scRNA-seq data”. In: Bioinfor-matics 40.9, btae512. doi: 10.1093/bioinformatics/btae512.

McCarthy, DJ, R Rostom, Y Huang, DJ Kunz, P Danecek, MJ Bonder, T Hagai, R Lyu, et al. (2020). “Cardelino: computational integration of somatic clonal substructure and single-cell transcriptomes”. In: Nature Methods 17, pp. 414–421. doi: 10.1038/s41592-020-0766-3.

McInnes, L, J Healy, N Saul, and L Großberger (2018). “UMAP: Uniform Manifold Approximation and Projection for Dimension Reduction”. In: Journal of Open Source Software 3.29, p. 861. doi: 10.21105/joss.00861.

Moncada, R, D Barkley, F Wagner, M Chiodin, JC Devlin, M Baron, CH Hajdu, DM Simeone, and I Yanai (2020). “Integrating microarray-based spatial transcriptomics and single-cell RNA-seq reveals tissue architecture in pancreatic ductal adenocarcinomas”. In: Nature Biotechnology 38.3, pp. 333–342. doi: 10.1038/s41587-019-0392-8.

Muyas, F, CM Sauer, J Espejo Valle-Inclán, R Li, R Rahbari, TJ Mitchell, S Hormoz, and I Cortés-Ciriano (2024). “De novo detection of somatic mutations in high-throughput single-cell profiling data sets”. In: Nature Biotechnology 42, pp. 758–767. doi: 10.1038/s41587-023-01863-z.

Nam, AS, KT Kim, R Chaligne, F Izzo, C Ahn, J Taylor, RM Myers, V Volpe, D Marenstein, R Yelin, et al. (2019). “Somatic mutations and cell identity linked by Genotyping of Transcriptomes”. In: Nature 571, pp. 355–360. doi: 10.1038/s41586-019-1367-0.

Neftel, C et al. (2019). “An Integrative Model of Cellular States, Plasticity, and Genetics for Glioblastoma”. In: Cell 178.4, 835–849.e21. doi: 10.1016/j.cell.2019.06.024.

Nowell, PC (1976). “The clonal evolution of tumor cell populations”. In: Science 194.4260, pp. 23–28. doi: 10.1126/science.959840.

Patel, AP, I Tirosh, JJ Trombetta, AK Shalek, SM Gillespie, H Wakimoto, DP Cahill, BV Nahed, WT Curry, RL Martuza, et al. (2014). “Single-cell RNA-seq highlights intratumoral heterogeneity in primary glioblastoma”. In: Science 344.6190, pp. 1396–1401. doi: 10.1126/science.1254257.

Pedregosa, F, G Varoquaux, A Gramfort, V Michel, B Thirion, O Grisel, M Blondel, P Prettenhofer, R Weiss, V Dubourg, J Vanderplas, A Passos, D Cournapeau, M Brucher, M Perrot, and E Duchesnay (2011). “Scikit-learn: Machine Learning in Python”. In: Journal of Machine Learning Research 12, pp. 2825–2830.

Petti, AA, SR Williams, CA Miller, IT Fiddes, SN Srivatsan, DY Chen, CC Fronick, RS Fulton, DM Church, and TJ Ley (2019). “A general approach for detecting expressed mutations in AML cells using single cell RNA-sequencing”. In: Nature Communications 10, p. 3660. doi: 10.1038/s41467-019-11591-1.

Rodriguez-Meira, A, G Buck, SA Clark, BJ Povinelli, V Alcolea, E Louka, S McGowan, A Hamblin, N Sousos, N Barkas, A Giustacchini, B Psaila, SEW Jacobsen, S Thongjuea, and AJ Mead (2019). “Unravelling Intratumoral Heterogeneity through High-Sensitivity Single-Cell Mutational Analysis and Parallel RNA Sequencing”. In: Molecular Cell 73.6, 1292–1305.e8. doi: 10.1016/j.molcel.2019.01.009.

Saito, T and M Rehmsmeier (2015). “The Precision–Recall Plot Is More Informative than the ROC Plot When Evaluating Binary Classifiers on Imbalanced Datasets”. In: PLOS ONE 10.3, e0118432. doi: 10.1371/journal.pone.0118432.

Schmid, KT et al. (2025). “Benchmarking scRNA-seq copy number variation callers”. In: Nature Communications 16, p. 8777. doi: 10.1038/s41467-025-62359-9.

Sturm, G (2024). infercnvpy: Scanpy plugin to infer copy number variation (CNV) from single-cell transcriptomics data. Software documentation.

Tirosh, I, AS Venteicher, C Hebert, LE Escalante, AP Patel, K Yizhak, JM Fisher, C Rodman, C Mount, MG Filbin, et al. (2016). “Single-cell RNA-seq supports a developmental hierarchy in human oligodendroglioma”. In: Nature 539.7628, pp. 309–313. doi: 10.1038/nature20123.

Tyner, JW et al. (2018). “Functional genomic landscape of acute myeloid leukaemia”. In: Nature 562, pp. 526–531. doi: 10.1038/s41586-018-0623-z.

van Galen, P, V Hovestadt, I Wadsworth M. H., TK Hughes, GK Griffin, S Battaglia, JA Verga, J Stephansky, TJ Pastika, J Lombardi Story, GS Pinkus, O Pozdnyakova, I Galinsky, RM Stone, TA Graubert, AK Shalek, JC Aster, AA Lane, and BE Bernstein (2019). “Single-Cell RNA-Seq Reveals AML Hierarchies Relevant to Disease Progression and Immunity”. In: Cell 176.6, 1265–1281.e24. doi: 10.1016/j.cell.2019.01.031.

Verhaak, RGW et al. (2010). “Integrated genomic analysis identifies clinically relevant subtypes of glioblastoma characterized by abnormalities in PDGFRA, IDH1, EGFR, and NF1”. In: Cancer Cell 17.1, pp. 98–110. doi: 10.1016/j.ccr.2009.12.020.

Virshup, I, S Rybakov, FJ Theis, P Angerer, and FA Wolf (2024). “anndata: Access and store annotated data matrices”. In: Journal of Open Source Software 9.101, p. 4371. doi: 10.21105/joss.04371.

Vogelstein, B, N Papadopoulos, VE Velculescu, S Zhou, LA Diaz, and KW Kinzler (2013). “Cancer genome landscapes”. In: Science 339.6127, pp. 1546–1558. doi: 10.1126/science.1235122.

Vu, TN, HN Nguyen, S Calza, KR Kalari, L Wang, and Y Pawitan (2019). “Cell-level somatic mutation detection from single-cell RNA sequencing”. In: Bioinformatics 35.22, pp. 4679–4687. doi: 10.1093/bioinformatics/btz288.

Way, GP, RJ Allaway, SJ Bouley, CE Fadul, Y Sanchez, and CS Greene (2017). “A machine learning classifier trained on cancer transcriptomes detects NF1 inactivation signal in glioblastoma”. In: BMC Genomics 18, p. 127. doi: 10.1186/s12864-017-3519-7.

Wolf, FA, P Angerer, and FJ Theis (2018). “SCANPY: Large-scale single-cell gene expression data analysis”. In: Genome Biology 19, p. 15. doi: 10.1186/s13059-017-1382-0.

Wu, SZ et al. (2021). “A single-cell and spatially resolved atlas of human breast cancers”. In: Nature Genetics 53.9, pp. 1334–1347. doi: 10.1038/s41588-021-00911-1.

Ziegenhain, C et al. (2017). “Comparative analysis of single-cell RNA sequencing methods”. In: Molecular Cell 65.4, 631–643.e4. doi: 10.1016/j.molcel.2017.01.023.

