## Supplemental_Code for "scOPE identifies which driver-associated expression programs transfer from bulk tumors to single cells": check_all_celltype_cols.py

```
# check_all_celltype_cols.py
#
# Runs the SAME check that caught the GBM bug, across all 7 CNV datasets at once:
# for each dataset's celltype_col, does that column actually exist on its scored
# h5ad? If yes, does the configured reference_regex actually match anything, and
# is the matched fraction plausible (not 0%, not implausibly close to 100%)?
#
# This does NOT modify anything. It only reads and reports. Run it once, paste the
# output back, and any dataset flagged [SUSPECT] or [BROKEN] gets the same
# treatment GBM just got before touching make_infercnvpy_cnvs.py further.

import re
import anndata as ad
import make_infercnvpy_cnvs as M   # reuse the real DEFAULT_DATASETS + regexes, no copy/paste drift

print(f"Checking {len(M.DEFAULT_DATASETS)} dataset specs from make_infercnvpy_cnvs.DEFAULT_DATASETS\n")

for name, spec in M.DEFAULT_DATASETS.items():
    print("=" * 88)
    print(f"{name}  ->  {spec['path']}")
    path = spec["path"]
    if not path.exists():
        print(f"  [SKIP] file not found on this machine")
        continue

    try:
        a = ad.read_h5ad(path, backed="r")  # backed=r: fast, obs-only read
    except Exception as e:
        print(f"  [ERROR] could not open: {e}")
        continue

    n = a.n_obs
    ct_col = spec.get("celltype_col")
    obs_cols = list(a.obs.columns)
    print(f"  n_obs={n:,} | configured celltype_col={ct_col!r}")

    if ct_col not in obs_cols:
        print(f"  [BROKEN] '{ct_col}' NOT IN obs. This is the exact GBM failure mode:")
        print(f"           _standardize_obs will silently fall back to an all-cell reference.")
        # surface likely alternative columns so the fix is fast
        candidates = [c for c in obs_cols if re.search(r"cell.?type|celltype|annotation|cluster", c, re.I)]
        print(f"           obs columns that LOOK like cell-type annotations: {candidates}")
        print(f"           full obs columns: {obs_cols}")
        continue

    vc = a.obs[ct_col].astype(str).value_counts()
    print(f"  '{ct_col}' has {len(vc)} categories. Top values:")
    print("    " + vc.head(15).to_string().replace("\n", "\n    "))

    ref_re = spec.get("reference_regex") or ""
    is_ref = a.obs[ct_col].astype(str).str.contains(ref_re, case=False, regex=True, na=False)
    n_ref = int(is_ref.sum())
    frac = n_ref / max(n, 1)
    print(f"  reference_regex matches: {n_ref:,}/{n:,}  ({frac:.1%})")

    # Flag anything that looks like GBM's failure mode or its near-miss cousins.
    flags = []
    if n_ref == 0:
        flags.append("0 reference cells -> WILL error out or silently use all-cell reference")
    elif frac < 0.03:
        flags.append(f"reference fraction very low ({frac:.1%}) -- check if regex/column is actually matching this vocabulary")
    elif frac > 0.85:
        flags.append(f"reference fraction very high ({frac:.1%}) -- check whether malignant cells are leaking into 'reference'")

    # Look for a suspiciously large non-reference "stromal/fibroblast"-labeled bucket,
    # which is what actually happened in GBM (malignant cells mislabeled as stroma).
    stromal_like = vc[vc.index.str.contains(r"strom|fibroblast|caf", case=False, regex=True)]
    if not stromal_like.empty:
        st_frac = stromal_like.sum() / n
        tag = " <-- large fraction; verify these are real stroma, not a malignant-cell-state confound" if st_frac > 0.15 else ""
        print(f"  stromal/fibroblast-labeled cells: {int(stromal_like.sum()):,} ({st_frac:.1%}){tag}")
        if st_frac > 0.15 and name in {"GBM", "LUAD", "PAAD", "SKCM"}:
            flags.append(f"large stromal/fibroblast bucket ({st_frac:.1%}) in a cancer where a mesenchymal-like "
                          f"or EMT malignant state can masquerade as fibroblast -- same failure mode as GBM")

    if flags:
        print(f"  [SUSPECT] " + " | ".join(flags))
    else:
        print(f"  [OK] celltype_col exists, reference fraction plausible")

print("\n" + "=" * 88)
print("Done. Any [BROKEN] or [SUSPECT] dataset above should be treated like GBM:")
print("  1. inspect its FULL celltype_col value_counts (not just top 15)")
print("  2. check for a companion 'is_reference'-style boolean already on the object")
print("  3. only then decide whether to repoint celltype_col or rewrite reference_regex")
```
