## Supplemental_Code for "scOPE identifies which driver-associated expression programs transfer from bulk tumors to single cells": check_gbm.py

```
# check_gbm2.py
import anndata as ad
p = r"C:\Users\ashfo\OneDrive\Desktop\Pathways+Omics\scOPE\scOPE\data\models\GBM_scored_z_score_bulk_with_eval_labels.h5ad"
a = ad.read_h5ad(p)

print("== full celltype_coarse value_counts ==")
print(a.obs["celltype_coarse"].value_counts(dropna=False).to_string())

print("\n== full celltype_marker value_counts ==")
print(a.obs["celltype_marker"].value_counts(dropna=False).to_string())

print("\n== is_reference distribution ==")
print(a.obs["is_reference"].value_counts(dropna=False).to_string())

print("\n== is_reference crosstab vs celltype_coarse ==")
print(a.obs.groupby("celltype_coarse")["is_reference"].mean().sort_values().to_string())

for col in ["is_reference_lymphoid_BM1_BM4", "is_reference_all_healthy_BM1_BM4",
            "is_reference_myeloid_prog_BM1_BM4", "is_reference_hspc_prog_BM1_BM4",
            "is_normal_display", "scope_malignant"]:
    print(f"\n== {col} distribution ==")
    print(a.obs[col].value_counts(dropna=False).to_string())
```
