## Supplemental_Code for "scOPE identifies which driver-associated expression programs transfer from bulk tumors to single cells": diagnose_bulk_id_overlap.py

```
"""
Diagnose why bulk expression sample IDs do not overlap mutation-label sample IDs.

Run from the suite root:
  cd C:\\Users\\ashfo\\OneDrive\\Desktop\\Pathways+Omics\\scOPE\\scope_analysis
  python diagnose_bulk_id_overlap.py --config drivers\\config_canonical_v3.json
"""
from pathlib import Path
import argparse
import json
import re
import pandas as pd


def sep_for(path):
    name = str(path).lower()
    return "\t" if name.endswith((".tsv", ".tsv.gz", ".txt", ".txt.gz")) else ","


def norm_id(x):
    s = str(x).strip()
    s = re.sub(r"^X", "", s)
    s = s.replace(".", "-").replace("_", "-")
    s = re.sub(r"\s+", "", s)
    s = s.upper()
    if s.startswith("TCGA-"):
        parts = s.split("-")
        if len(parts) >= 4:
            s = "-".join(parts[:3] + [parts[3][:2]])
    return s


def read_expr_ids(path):
    path = Path(path)
    if path.suffix == ".h5ad":
        import anndata as ad
        a = ad.read_h5ad(path, backed="r")
        return list(map(str, a.obs_names[:])), "h5ad"
    df_auto = pd.read_csv(path, sep=sep_for(path), index_col=0)
    # Also mimic the old/broken loader for useful diagnosis.
    try:
        df_old = pd.read_csv(path, index_col=0)
        old_shape = df_old.shape
    except Exception:
        old_shape = None
    return list(map(str, df_auto.index)), f"table_auto_sep_shape={df_auto.shape}; old_csv_default_shape={old_shape}"


def choose_label_col(df, requested):
    if requested in df.columns:
        return requested
    candidates = [
        "sample_id", "sample", "Sample", "SAMPLE", "id", "ID",
        "barcode", "Barcode", "BARCODE", "patient_id", "patient", "case_id",
        "tumor_sample_barcode", "Tumor_Sample_Barcode", "TumorSampleBarcode",
    ]
    for c in candidates:
        if c in df.columns:
            return c
    return df.columns[0]


def main():
    ap = argparse.ArgumentParser()
    ap.add_argument("--config", required=True)
    ap.add_argument("--out", default="bulk_id_overlap_diagnostic.csv")
    args = ap.parse_args()

    cfg = json.loads(Path(args.config).read_text())
    rows = []
    for cancer, bc in cfg.get("bulk_cohorts", {}).items():
        expr = Path(bc["expression"])
        mut = Path(bc["mutations"])
        sample_col = bc.get("sample_col", "sample_id")
        try:
            expr_ids, expr_read_info = read_expr_ids(expr)
            lab_df = pd.read_csv(mut, sep=sep_for(mut))
            used_col = choose_label_col(lab_df, sample_col)
            lab_ids = lab_df[used_col].astype(str).str.strip().tolist()

            exact = len(set(expr_ids) & set(lab_ids))
            norm_expr = [norm_id(x) for x in expr_ids]
            norm_lab = [norm_id(x) for x in lab_ids]
            norm = len(set(norm_expr) & set(norm_lab))
            rows.append({
                "cancer": cancer,
                "expr_read_info": expr_read_info,
                "requested_sample_col": sample_col,
                "used_label_col": used_col,
                "n_expr_ids": len(expr_ids),
                "n_label_ids": len(lab_ids),
                "exact_overlap": exact,
                "normalized_overlap": norm,
                "expr_examples": "; ".join(expr_ids[:5]),
                "label_examples": "; ".join(lab_ids[:5]),
                "expr_norm_examples": "; ".join(norm_expr[:5]),
                "label_norm_examples": "; ".join(norm_lab[:5]),
                "expression": str(expr),
                "mutations": str(mut),
            })
        except Exception as e:
            rows.append({"cancer": cancer, "expression": str(expr), "mutations": str(mut), "error": repr(e)})

    out = Path(args.out)
    df = pd.DataFrame(rows)
    df.to_csv(out, index=False)
    print(df.to_string(index=False))
    print(f"\nWrote: {out.resolve()}")


if __name__ == "__main__":
    main()
```
