## Supplemental_Code for "scOPE identifies which driver-associated expression programs transfer from bulk tumors to single cells": make_pathway_analysis.py

```
#!/usr/bin/env python
"""
make_pathway_analysis.py
========================
Pathway / gene-set over-representation analysis (ORA) on the gene programmes scOPE
learned for each driver, using the `genes_<cancer>_<driver>.csv` tables the pipeline
already wrote. Offline and reproducible: needs only a local `.gmt`.

Get a gene-set file once (symbols), e.g. MSigDB Hallmark:
    https://www.gsea-msigdb.org/gsea/msigdb  ->  h.all.<ver>.symbols.gmt

Usage
-----
python drivers/make_pathway_analysis.py \
    --tables-dir scope_analysis_outputs/tables \
    --gmt h.all.v2024.1.Hs.symbols.gmt \
    --background-genes scope_analysis_outputs/tables/bulk_gene_universe.txt \
    --outdir scope_analysis_outputs/pathways \
    --top-n 100

--background-genes is optional but recommended (one symbol per line: your assayed
gene universe). Without it the union of gene-set members is used as background.
"""

from __future__ import annotations
import argparse, glob, re
from pathlib import Path

import pandas as pd

from scope_analysis import pathways as pw


def _load_background(path):
    if path and Path(path).exists():
        with open(path) as fh:
            return [ln.strip() for ln in fh if ln.strip()]
    return None


def main():
    ap = argparse.ArgumentParser()
    ap.add_argument("--tables-dir", required=True, help="scope_analysis_outputs/tables")
    ap.add_argument("--gmt", required=True, help="local .gmt gene-set file (symbols)")
    ap.add_argument("--background-genes", default=None, help="optional txt, one symbol/line")
    ap.add_argument("--outdir", default="pathways")
    ap.add_argument("--top-n", type=int, default=100, help="leading genes per programme")
    ap.add_argument("--by-direction", action="store_true",
                    help="also test up/down gene sets separately")
    ap.add_argument("--use-gseapy", action="store_true",
                    help="use gseapy.enrichr (online) instead of offline hypergeometric")
    args = ap.parse_args()

    outdir = Path(args.outdir); outdir.mkdir(parents=True, exist_ok=True)
    background = _load_background(args.background_genes)
    files = sorted(glob.glob(str(Path(args.tables_dir) / "genes_*.csv")))
    if not files:
        # fall back to the deeper component scan tables
        files = sorted(glob.glob(str(Path(args.tables_dir).parent /
                                     "fig2_bulk" / "*" / "component_signature_scan.csv")))
    if not files:
        raise SystemExit(f"No genes_*.csv or component_signature_scan.csv under {args.tables_dir}")

    combined = []
    for f in files:
        m = re.search(r"genes_([A-Za-z0-9]+)_([A-Za-z0-9]+)\.csv$", Path(f).name)
        if m:
            cancer, driver = m.group(1), m.group(2)
        else:  # component_signature_scan.csv -> split per driver below
            cancer, driver = Path(f).parent.name, None
        df = pd.read_csv(f)
        if "gene" not in df.columns:
            continue
        groups = [(driver, df)] if driver else [(d, sub) for d, sub in df.groupby("driver")]
        for drv, sub in groups:
            enr = pw.enrich_driver_programs(
                sub, args.gmt, background=background, top_n=args.top_n,
                by_direction=args.by_direction, use_gseapy=args.use_gseapy)
            if enr.empty:
                print(f"[pathways] {cancer}/{drv}: no enrichment")
                continue
            enr.insert(0, "driver", drv); enr.insert(0, "cancer", cancer)
            enr.to_csv(outdir / f"pathways_{cancer}_{drv}.csv", index=False)
            pw.plot_pathway_bar(enr, outdir / f"pathways_{cancer}_{drv}.png",
                                title=f"{cancer} · {drv} — enriched gene sets", direction="combined")
            sig = enr[enr["fdr"] < 0.05]
            print(f"[pathways] {cancer}/{drv}: {len(sig)} sets FDR<0.05; "
                  f"top: {enr.iloc[0]['term'][:40]} (FDR={enr.iloc[0]['fdr']:.1e})")
            combined.append(enr)

    if combined:
        allt = pd.concat(combined, ignore_index=True)
        allt.to_csv(outdir / "pathways_ALL.csv", index=False)
        print(f"\n[pathways] wrote {len(combined)} driver tables + pathways_ALL.csv to {outdir}")


if __name__ == "__main__":
    main()
```
