## Supplemental_Code for "scOPE identifies which driver-associated expression programs transfer from bulk tumors to single cells": make_pathway_analysis_updated.py

```
#!/usr/bin/env python
"""
make_pathway_analysis.py
========================
Pathway / gene-set over-representation analysis (ORA) on the gene programmes scOPE
learned for each driver, using the `genes_<cancer>_<driver>.csv` tables the pipeline
already wrote.

This version supports multiple pathway/gene-set resources in one run:
  - local GMT files: Hallmark, GO BP/MF/CC, KEGG, Reactome, WikiPathways, etc.
  - a directory of GMT files
  - optional online Enrichr libraries through gseapy
  - ranked summary figures across all pathway databases, sorted by FDR/significance

Recommended reproducible/offline usage
--------------------------------------
Put your MSigDB GMTs in one folder, e.g.:
  h.all.v2024.1.Hs.symbols.gmt
  c5.go.bp.v2024.1.Hs.symbols.gmt
  c5.go.mf.v2024.1.Hs.symbols.gmt
  c5.go.cc.v2024.1.Hs.symbols.gmt
  c2.cp.kegg*.v2024.1.Hs.symbols.gmt
  c2.cp.reactome.v2024.1.Hs.symbols.gmt

Then run:
  python drivers/make_pathway_analysis.py \
      --tables-dir scope_analysis_outputs/tables \
      --gmt-dir path_to_msigdbr_gmts \
      --background-genes scope_analysis_outputs/tables/bulk_gene_universe.txt \
      --outdir scope_analysis_outputs/pathways \
      --top-n 100 \
      --by-direction

Single-GMT legacy usage still works:
  python drivers/make_pathway_analysis.py \
      --tables-dir scope_analysis_outputs/tables \
      --gmt h.all.v2024.1.Hs.symbols.gmt

Online Enrichr usage, requires gseapy + internet:
  python drivers/make_pathway_analysis.py \
      --tables-dir scope_analysis_outputs/tables \
      --enrichr-preset core \
      --outdir scope_analysis_outputs/pathways_enrichr

Used this most recently to run:
python drivers\make_pathway_analysis_updated.py --tables-dir scope_analysis_outputs\tables --background-genes scope_analysis_outputs/tables/bulk_gene_universe.txt --gmt-dir msigdb_gmts --background-genes scope_analysis_outputs\tables\bulk_gene_universe.txt --outdir scope_analysis_outputs\pathways --top-n 500 --by-direction
"""

from __future__ import annotations

import argparse
import glob
import math
import re
import sys
import textwrap
from pathlib import Path
from typing import Sequence, Tuple

import numpy as np
import pandas as pd


# Make imports work when launched either from repo root or from drivers/.
_REPO_ROOT = Path(__file__).resolve().parents[1] if len(Path(__file__).resolve().parents) > 1 else Path.cwd()
if str(_REPO_ROOT) not in sys.path:
    sys.path.insert(0, str(_REPO_ROOT))


Resource = Tuple[str, str, bool]  # collection_label, gmt_or_library, use_gseapy


# -----------------------------
# Input/resource helpers
# -----------------------------

def _load_background(path: str | None) -> list[str] | None:
    if path and Path(path).exists():
        with open(path, encoding="utf-8") as fh:
            return [ln.strip() for ln in fh if ln.strip()]
    return None


def _flatten_cli_values(values: Sequence[Sequence[str]] | Sequence[str] | None) -> list[str]:
    """Flatten argparse values and split comma-separated entries."""
    if not values:
        return []
    out: list[str] = []
    for item in values:
        if isinstance(item, (list, tuple)):
            raw_items = item
        else:
            raw_items = [item]
        for raw in raw_items:
            for piece in str(raw).split(","):
                piece = piece.strip().strip('"').strip("'")
                if piece:
                    out.append(piece)
    return out


def _safe_name(x: str) -> str:
    x = str(x).replace(".symbols.gmt", "").replace(".gmt", "")
    x = re.sub(r"[^A-Za-z0-9_.+-]+", "_", x)
    x = re.sub(r"_+", "_", x).strip("_")
    return x[:120] or "gene_sets"


def _collection_label(source: str, online: bool = False) -> str:
    if online:
        return _safe_name(source)
    name = Path(source).name
    name = name.replace(".symbols.gmt", "").replace(".gmt", "")
    return _safe_name(name)


def _glob_gmts(gmt_dirs: Sequence[str], gmt_glob: str) -> list[str]:
    files: list[str] = []
    for d in gmt_dirs:
        p = Path(d)
        if p.is_file() and p.suffix.lower() == ".gmt":
            files.append(str(p))
        elif p.is_dir():
            files.extend(str(x) for x in sorted(p.glob(gmt_glob)))
        else:
            # Glob patterns passed directly to --gmt-dir are allowed too.
            files.extend(sorted(glob.glob(str(d))))
    return files


def _find_gene_tables(tables_dir: str) -> list[str]:
    tables_path = Path(tables_dir)
    files = sorted(glob.glob(str(tables_path / "genes_*.csv")))
    if not files:
        # Fall back to the deeper component scan tables.
        files = sorted(glob.glob(str(tables_path.parent / "fig2_bulk" / "*" / "component_signature_scan.csv")))
    if not files:
        raise SystemExit(f"No genes_*.csv or component_signature_scan.csv under {tables_dir}")
    return files


def _driver_groups(path: str) -> tuple[str, list[tuple[str, pd.DataFrame]]]:
    f = Path(path)
    m = re.search(r"genes_([A-Za-z0-9]+)_([A-Za-z0-9]+)\.csv$", f.name)
    df = pd.read_csv(f)
    if "gene" not in df.columns:
        return f.parent.name, []

    if m:
        cancer, driver = m.group(1), m.group(2)
        return cancer, [(driver, df)]

    # component_signature_scan.csv -> split per driver if possible.
    cancer = f.parent.name
    if "driver" in df.columns:
        return cancer, [(str(d), sub.copy()) for d, sub in df.groupby("driver", dropna=False)]
    return cancer, [("program", df)]


def _resolve_enrichr_preset(preset: str, explicit_libs: Sequence[str]) -> list[str]:
    """
    Resolve online Enrichr libraries.

    The exact Enrichr library names drift over time, so for presets we try to query
    the local gseapy installation / Enrichr and pick the newest matching library.
    If that fails, we fall back to common library names.
    """
    libs = list(explicit_libs)
    if preset in (None, "none"):
        return libs

    fallback = {
        "go": [
            "GO_Biological_Process_2025",
            "GO_Molecular_Function_2025",
            "GO_Cellular_Component_2025",
        ],
        "core": [
            "GO_Biological_Process_2025",
            "GO_Molecular_Function_2025",
            "GO_Cellular_Component_2025",
            "KEGG_2021_Human",
            "Reactome_2022",
            "WikiPathways_2024_Human",
            "MSigDB_Hallmark_2020",
        ],
        "go_kegg": [
            "GO_Biological_Process_2025",
            "GO_Molecular_Function_2025",
            "GO_Cellular_Component_2025",
            "KEGG_2021_Human",
        ],
    }

    requested_patterns = {
        "go": [
            r"^GO_Biological_Process_\d{4}$",
            r"^GO_Molecular_Function_\d{4}$",
            r"^GO_Cellular_Component_\d{4}$",
        ],
        "go_kegg": [
            r"^GO_Biological_Process_\d{4}$",
            r"^GO_Molecular_Function_\d{4}$",
            r"^GO_Cellular_Component_\d{4}$",
            r"^KEGG_\d{4}_Human$",
        ],
        "core": [
            r"^GO_Biological_Process_\d{4}$",
            r"^GO_Molecular_Function_\d{4}$",
            r"^GO_Cellular_Component_\d{4}$",
            r"^KEGG_\d{4}_Human$",
            r"^Reactome_\d{4}$",
            r"^WikiPathways_\d{4}_Human$",
            r"^MSigDB_Hallmark_\d{4}$",
        ],
    }

    selected: list[str] = []
    try:
        import gseapy as gp  # type: ignore

        try:
            available = gp.get_library_name(organism="Human")
        except TypeError:
            available = gp.get_library_name()

        def year_key(name: str) -> tuple[int, str]:
            years = [int(y) for y in re.findall(r"(?:19|20)\d{2}", name)]
            return (max(years) if years else -1, name)

        for pat in requested_patterns[preset]:
            hits = sorted([x for x in available if re.search(pat, x)], key=year_key)
            if hits:
                selected.append(hits[-1])
            else:
                print(f"[pathways] WARNING: no Enrichr library matched {pat!r}; using fallback if available")
    except Exception as e:
        print(f"[pathways] WARNING: could not query Enrichr library names ({e}); using fallback names")
        selected = []

    if not selected:
        selected = fallback[preset]

    # Preserve explicit libraries and remove duplicates while keeping order.
    out: list[str] = []
    for lib in list(libs) + selected:
        if lib not in out:
            out.append(lib)
    return out


def _build_resources(args: argparse.Namespace) -> list[Resource]:
    gmt_files = _flatten_cli_values(args.gmt)
    gmt_files.extend(_glob_gmts(_flatten_cli_values(args.gmt_dir), args.gmt_glob))

    # De-duplicate while preserving order.
    seen = set()
    unique_gmts: list[str] = []
    for g in gmt_files:
        key = str(Path(g))
        if key not in seen:
            seen.add(key)
            unique_gmts.append(g)

    missing = [g for g in unique_gmts if not Path(g).exists()]
    if missing:
        raise SystemExit("Missing GMT file(s):\n  " + "\n  ".join(missing))

    enrichr_libs = _flatten_cli_values(args.enrichr_libraries)
    enrichr_libs = _resolve_enrichr_preset(args.enrichr_preset, enrichr_libs)

    resources: list[Resource] = []
    for g in unique_gmts:
        resources.append((_collection_label(g, online=False), g, bool(args.use_gseapy)))
    for lib in enrichr_libs:
        resources.append((_collection_label(lib, online=True), lib, True))

    if not resources:
        raise SystemExit(
            "No pathway/gene-set resources provided. Use --gmt, --gmt-dir, "
            "--enrichr-libraries, or --enrichr-preset."
        )
    return resources


# -----------------------------
# Output/plot helpers
# -----------------------------

def _fdr_column(enr: pd.DataFrame) -> str | None:
    for c in ["fdr", "FDR", "qvalue", "q_value", "adj_p", "adj_pvalue", "Adjusted P-value", "Adjusted P"]:
        if c in enr.columns:
            return c
    return None


def _p_column(enr: pd.DataFrame) -> str | None:
    for c in ["pval", "p_value", "pvalue", "P-value", "P-value", "p"]:
        if c in enr.columns:
            return c
    return None


def _term_column(enr: pd.DataFrame) -> str | None:
    for c in ["term", "Term", "name", "gene_set", "Gene_set", "pathway", "Pathway"]:
        if c in enr.columns:
            return c
    return None


def _direction_column(enr: pd.DataFrame) -> str | None:
    for c in ["direction", "Direction", "set", "subset", "contrast"]:
        if c in enr.columns:
            return c
    return None


def _first_term(enr: pd.DataFrame) -> str:
    c = _term_column(enr)
    if c and len(enr):
        return str(enr.iloc[0][c])
    return "NA"


def _wrap_label(text: str, width: int = 72, max_chars: int = 150) -> str:
    text = re.sub(r"\s+", " ", str(text)).strip()
    if len(text) > max_chars:
        text = text[: max_chars - 1].rstrip() + "…"
    return "\n".join(textwrap.wrap(text, width=width, break_long_words=False, break_on_hyphens=False))


def _standardize_for_summary(enr: pd.DataFrame) -> pd.DataFrame:
    """Add robust term/FDR/significance/display columns for plotting summaries."""
    df = enr.copy()
    term_col = _term_column(df)
    fdr_col = _fdr_column(df)
    p_col = _p_column(df)
    dir_col = _direction_column(df)

    if term_col is None:
        df["summary_term"] = df.index.astype(str)
    else:
        df["summary_term"] = df[term_col].astype(str)

    if fdr_col is not None:
        df["summary_fdr"] = pd.to_numeric(df[fdr_col], errors="coerce")
    elif p_col is not None:
        # Fallback only. This is not adjusted significance, so label it clearly.
        df["summary_fdr"] = pd.to_numeric(df[p_col], errors="coerce")
    else:
        df["summary_fdr"] = np.nan

    # Avoid inf from exact zeros while keeping very strong hits very strong.
    df["summary_fdr_clipped"] = df["summary_fdr"].clip(lower=1e-300, upper=1.0)
    df["summary_neglog10_fdr"] = -np.log10(df["summary_fdr_clipped"])

    pieces = []
    for _, row in df.iterrows():
        context = f"{row.get('cancer', 'NA')}/{row.get('driver', 'NA')} | {row.get('collection', 'NA')}"
        if dir_col is not None and pd.notna(row.get(dir_col)):
            direction = str(row.get(dir_col))
            if direction and direction.lower() not in {"nan", "none", "combined"}:
                context += f" | {direction}"
        pieces.append(f"{row['summary_term']} [{context}]")
    df["summary_display"] = pieces

    df = df.replace([np.inf, -np.inf], np.nan)
    df = df.dropna(subset=["summary_neglog10_fdr"])
    return df


def _savefig_all_formats(fig, base_path: Path, formats: Sequence[str], dpi: int = 220) -> list[Path]:
    written: list[Path] = []
    for fmt in formats:
        fmt = fmt.lower().lstrip(".")
        path = base_path.with_suffix(f".{fmt}")
        fig.savefig(path, bbox_inches="tight", dpi=dpi)
        written.append(path)
    return written


def _plot_ranked_pathways(
    df: pd.DataFrame,
    base_path: Path,
    title: str,
    formats: Sequence[str],
    top_n: int = 50,
    fdr_cutoff: float | None = None,
    max_label_chars: int = 160,
) -> list[Path]:
    """Horizontal barplot ranked by -log10(FDR), regardless of database."""
    import matplotlib.pyplot as plt

    if df.empty:
        return []

    plot_df = df.copy()
    plot_df = plot_df.sort_values(["summary_fdr", "summary_neglog10_fdr"], ascending=[True, False]).head(top_n)
    if plot_df.empty:
        return []

    # Reverse so the strongest hit is at the top in barh.
    plot_df = plot_df.iloc[::-1].copy()
    labels = [_wrap_label(x, width=68, max_chars=max_label_chars) for x in plot_df["summary_display"]]
    values = plot_df["summary_neglog10_fdr"].astype(float).values

    height = max(4.5, min(0.34 * len(plot_df) + 1.7, 24))
    fig, ax = plt.subplots(figsize=(12, height))
    ax.barh(range(len(plot_df)), values)
    ax.set_yticks(range(len(plot_df)))
    ax.set_yticklabels(labels, fontsize=8)
    ax.set_xlabel("-log10(FDR)")
    ax.set_title(title)
    ax.grid(axis="x", alpha=0.25)

    if fdr_cutoff is not None and 0 < fdr_cutoff < 1:
        ax.axvline(-math.log10(fdr_cutoff), linestyle="--", linewidth=1)
        ax.text(
            -math.log10(fdr_cutoff),
            len(plot_df) - 0.35,
            f" FDR={fdr_cutoff:g}",
            va="top",
            fontsize=8,
        )

    # Annotate a few values without crowding.
    for y, (_, row) in enumerate(plot_df.iterrows()):
        val = float(row["summary_neglog10_fdr"])
        fdr = float(row["summary_fdr"])
        label = f"{fdr:.1e}" if fdr < 0.001 else f"{fdr:.3g}"
        ax.text(val + max(values) * 0.006, y, label, va="center", fontsize=7)

    fig.tight_layout()
    written = _savefig_all_formats(fig, base_path, formats)
    plt.close(fig)
    return written


def _plot_collection_counts(
    df: pd.DataFrame,
    base_path: Path,
    title: str,
    formats: Sequence[str],
    fdr_cutoff: float = 0.05,
) -> list[Path]:
    """Barplot counting significant sets per collection/database."""
    import matplotlib.pyplot as plt

    if df.empty or "collection" not in df.columns:
        return []
    sig = df[df["summary_fdr"] < fdr_cutoff].copy()
    if sig.empty:
        return []
    counts = sig["collection"].astype(str).value_counts().sort_values(ascending=True)

    height = max(4, min(0.35 * len(counts) + 1.8, 16))
    fig, ax = plt.subplots(figsize=(9, height))
    ax.barh(range(len(counts)), counts.values)
    ax.set_yticks(range(len(counts)))
    ax.set_yticklabels([_wrap_label(x, width=45, max_chars=90) for x in counts.index], fontsize=8)
    ax.set_xlabel(f"Number of enriched sets, FDR < {fdr_cutoff:g}")
    ax.set_title(title)
    ax.grid(axis="x", alpha=0.25)
    for y, v in enumerate(counts.values):
        ax.text(v + max(counts.values) * 0.01, y, str(int(v)), va="center", fontsize=8)
    fig.tight_layout()
    written = _savefig_all_formats(fig, base_path, formats)
    plt.close(fig)
    return written


def _plot_top_heatmap(
    df: pd.DataFrame,
    base_path: Path,
    title: str,
    formats: Sequence[str],
    top_n_terms: int = 40,
    cap_neglog10: float = 50.0,
) -> list[Path]:
    """Heatmap of top pathway terms across cancer/driver contexts."""
    import matplotlib.pyplot as plt

    if df.empty:
        return []

    work = df.copy()
    work["context"] = work["cancer"].astype(str) + "/" + work["driver"].astype(str)
    work["term_collection"] = work["summary_term"].astype(str) + " [" + work["collection"].astype(str) + "]"

    # Keep the globally strongest unique term/database pairs.
    keep_terms = (
        work.sort_values("summary_fdr", ascending=True)
        .drop_duplicates("term_collection")
        .head(top_n_terms)["term_collection"]
        .tolist()
    )
    work = work[work["term_collection"].isin(keep_terms)].copy()
    if work.empty:
        return []

    piv = work.pivot_table(
        index="term_collection",
        columns="context",
        values="summary_neglog10_fdr",
        aggfunc="max",
        fill_value=0.0,
    )
    # Reorder rows by max significance.
    piv = piv.loc[piv.max(axis=1).sort_values(ascending=False).index]
    piv = piv.clip(upper=cap_neglog10)
    piv.to_csv(base_path.with_suffix(".csv"))

    height = max(5, min(0.32 * piv.shape[0] + 2.2, 26))
    width = max(7, min(0.48 * piv.shape[1] + 7.0, 26))
    fig, ax = plt.subplots(figsize=(width, height))
    im = ax.imshow(piv.values, aspect="auto")
    cbar = fig.colorbar(im, ax=ax)
    cbar.set_label(f"-log10(FDR), capped at {cap_neglog10:g}")
    ax.set_xticks(range(piv.shape[1]))
    ax.set_xticklabels(piv.columns, rotation=55, ha="right", fontsize=7)
    ax.set_yticks(range(piv.shape[0]))
    ax.set_yticklabels([_wrap_label(x, width=60, max_chars=130) for x in piv.index], fontsize=7)
    ax.set_title(title)
    fig.tight_layout()
    written = _savefig_all_formats(fig, base_path, formats)
    plt.close(fig)
    return written


def _manifest_add(
    manifest: list[dict],
    path: Path,
    kind: str,
    description: str,
    cancer: str | None = None,
    driver: str | None = None,
    collection: str | None = None,
) -> None:
    """Track important outputs in a machine-readable manifest."""
    manifest.append(
        {
            "kind": kind,
            "path": str(path),
            "cancer": cancer or "",
            "driver": driver or "",
            "collection": collection or "",
            "description": description,
        }
    )


def _layout_roots(outdir: Path, flat_output: bool) -> dict[str, Path]:
    """Return root directories for tables/figures/index files."""
    if flat_output:
        return {
            "tables": outdir,
            "figures": outdir,
            "summary_tables": outdir / "summary_figures",
            "summary_figures": outdir / "summary_figures",
            "index": outdir,
        }

    return {
        "tables": outdir / "tables",
        "figures": outdir / "figures",
        "summary_tables": outdir / "tables" / "summary",
        "summary_figures": outdir / "figures" / "summary",
        "index": outdir / "index",
    }


def _individual_output_paths(
    outdir: Path,
    flat_output: bool,
    cancer: str,
    driver: str,
    collection: str,
) -> tuple[Path, Path]:
    """CSV and PNG path for one cancer/driver/database result."""
    safe_cancer = _safe_name(cancer)
    safe_driver = _safe_name(driver)
    safe_collection = _safe_name(collection)

    if flat_output:
        stem = f"pathways_{safe_cancer}_{safe_driver}_{safe_collection}"
        return outdir / f"{stem}.csv", outdir / f"{stem}.png"

    table_dir = outdir / "tables" / "by_driver" / safe_cancer / safe_driver
    fig_dir = outdir / "figures" / "by_driver" / safe_cancer / safe_driver
    table_dir.mkdir(parents=True, exist_ok=True)
    fig_dir.mkdir(parents=True, exist_ok=True)
    return table_dir / f"{safe_collection}.csv", fig_dir / f"{safe_collection}.png"


def _all_collections_path(
    outdir: Path,
    flat_output: bool,
    cancer: str,
    driver: str,
) -> Path:
    safe_cancer = _safe_name(cancer)
    safe_driver = _safe_name(driver)
    if flat_output:
        return outdir / f"pathways_{safe_cancer}_{safe_driver}_ALL_COLLECTIONS.csv"
    d = outdir / "tables" / "by_driver" / safe_cancer / safe_driver
    d.mkdir(parents=True, exist_ok=True)
    return d / "ALL_COLLECTIONS.csv"


def _summary_table_dir(outdir: Path, flat_output: bool) -> Path:
    d = (outdir / "summary_figures") if flat_output else (outdir / "tables" / "summary")
    d.mkdir(parents=True, exist_ok=True)
    return d


def _summary_figure_dir(outdir: Path, flat_output: bool) -> Path:
    d = (outdir / "summary_figures") if flat_output else (outdir / "figures" / "summary")
    d.mkdir(parents=True, exist_ok=True)
    return d


def _write_grouped_tables(
    all_results: pd.DataFrame,
    outdir: Path,
    flat_output: bool,
    manifest: list[dict],
) -> None:
    """Write browseable aggregate tables by cancer and by pathway database."""
    if all_results.empty or flat_output:
        return

    by_cancer_root = outdir / "tables" / "by_cancer"
    by_database_root = outdir / "tables" / "by_database"
    by_cancer_root.mkdir(parents=True, exist_ok=True)
    by_database_root.mkdir(parents=True, exist_ok=True)

    for cancer, sub in all_results.groupby("cancer", dropna=False):
        safe_cancer = _safe_name(str(cancer))
        d = by_cancer_root / safe_cancer
        d.mkdir(parents=True, exist_ok=True)
        path = d / "ALL_DRIVERS_ALL_COLLECTIONS.csv"
        sub.to_csv(path, index=False)
        _manifest_add(
            manifest, path, "table",
            "All pathway enrichment rows for one cancer across drivers and collections.",
            cancer=str(cancer),
        )

    if "collection" in all_results.columns:
        for collection, sub in all_results.groupby("collection", dropna=False):
            safe_collection = _safe_name(str(collection))
            d = by_database_root / safe_collection
            d.mkdir(parents=True, exist_ok=True)
            path = d / "ALL_CANCERS_ALL_DRIVERS.csv"
            sub.to_csv(path, index=False)
            _manifest_add(
                manifest, path, "table",
                "All pathway enrichment rows for one pathway database/collection.",
                collection=str(collection),
            )


def _write_top_hit_indexes(
    all_results: pd.DataFrame,
    outdir: Path,
    flat_output: bool,
    manifest: list[dict],
    fdr_cutoff: float,
) -> None:
    """Write compact index tables that are nice to open first."""
    roots = _layout_roots(outdir, flat_output)
    index_dir = roots["index"]
    index_dir.mkdir(parents=True, exist_ok=True)

    std = _standardize_for_summary(all_results)
    if std.empty:
        return

    std = std.sort_values(["summary_fdr", "summary_neglog10_fdr"], ascending=[True, False]).copy()

    top_global = std.head(250)
    path = index_dir / "top_250_pathways_ranked_all_databases.csv"
    top_global.to_csv(path, index=False)
    _manifest_add(
        manifest, path, "index",
        "Quick-open ranked index of the top 250 pathway hits across every cancer, driver, and database.",
    )

    sig = std[std["summary_fdr"] < fdr_cutoff].copy()
    path = index_dir / f"significant_pathways_FDR_lt_{str(fdr_cutoff).replace('.', 'p')}.csv"
    sig.to_csv(path, index=False)
    _manifest_add(
        manifest, path, "index",
        f"All enrichment rows passing FDR < {fdr_cutoff:g}, sorted by significance.",
    )

    # Best row for each cancer/driver/database.
    group_cols = [c for c in ["cancer", "driver", "collection"] if c in std.columns]
    if group_cols:
        best = std.sort_values("summary_fdr", ascending=True).groupby(group_cols, dropna=False).head(1)
        path = index_dir / "best_hit_per_cancer_driver_database.csv"
        best.to_csv(path, index=False)
        _manifest_add(
            manifest, path, "index",
            "One strongest pathway hit per cancer/driver/database combination.",
        )

    # Best row for each cancer/driver regardless of database.
    group_cols = [c for c in ["cancer", "driver"] if c in std.columns]
    if group_cols:
        best = std.sort_values("summary_fdr", ascending=True).groupby(group_cols, dropna=False).head(1)
        path = index_dir / "best_hit_per_cancer_driver_any_database.csv"
        best.to_csv(path, index=False)
        _manifest_add(
            manifest, path, "index",
            "One strongest pathway hit per cancer/driver across all databases.",
        )


def _write_summary_figures(
    all_results: pd.DataFrame,
    outdir: Path,
    summary_top_n: int,
    heatmap_top_n: int,
    fig_formats: Sequence[str],
    fdr_cutoff: float,
    write_per_driver_summary: bool,
    max_label_chars: int,
    flat_output: bool,
    manifest: list[dict],
) -> None:
    summary_table_dir = _summary_table_dir(outdir, flat_output)
    summary_fig_dir = _summary_figure_dir(outdir, flat_output)

    std = _standardize_for_summary(all_results)
    if std.empty:
        print("[pathways] summary figures skipped: no usable FDR/p-value column found")
        return

    std = std.sort_values(["summary_fdr", "summary_neglog10_fdr"], ascending=[True, False]).copy()

    ranked_path = summary_table_dir / "summary_all_ranked_pathways.csv"
    std.to_csv(ranked_path, index=False)
    _manifest_add(
        manifest, ranked_path, "summary_table",
        "All pathway enrichment rows ranked by FDR across all databases.",
    )

    sig = std[std["summary_fdr"] < fdr_cutoff].copy()
    sig_path = summary_table_dir / f"summary_all_FDR_lt_{str(fdr_cutoff).replace('.', 'p')}.csv"
    sig.to_csv(sig_path, index=False)
    _manifest_add(
        manifest, sig_path, "summary_table",
        f"All significant pathway hits at FDR < {fdr_cutoff:g}.",
    )

    written: list[Path] = []
    written.extend(
        _plot_ranked_pathways(
            std,
            summary_fig_dir / "summary_top_pathways_ALL_DATABASES",
            title=f"Top enriched gene sets across all databases and drivers (top {summary_top_n})",
            formats=fig_formats,
            top_n=summary_top_n,
            fdr_cutoff=fdr_cutoff,
            max_label_chars=max_label_chars,
        )
    )
    written.extend(
        _plot_collection_counts(
            std,
            summary_fig_dir / "summary_significant_counts_by_database",
            title="Significant enriched gene sets by database/collection",
            formats=fig_formats,
            fdr_cutoff=fdr_cutoff,
        )
    )
    written.extend(
        _plot_top_heatmap(
            std,
            summary_fig_dir / "summary_top_pathways_heatmap_by_driver",
            title=f"Top pathway signatures across cancer/driver contexts (top {heatmap_top_n})",
            formats=fig_formats,
            top_n_terms=heatmap_top_n,
        )
    )

    for p in written:
        _manifest_add(
            manifest, p, "summary_figure",
            "Global summary figure across databases/drivers.",
        )

    # Per-cancer ranked summaries.
    for cancer, sub in std.groupby("cancer", dropna=False):
        safe_cancer = _safe_name(str(cancer))
        if flat_output:
            fig_base = summary_fig_dir / f"summary_top_pathways_{safe_cancer}_ALL_DATABASES"
            table_path = summary_table_dir / f"summary_ranked_{safe_cancer}_ALL_DATABASES.csv"
        else:
            cancer_table_dir = outdir / "tables" / "by_cancer" / safe_cancer
            cancer_fig_dir = outdir / "figures" / "by_cancer" / safe_cancer
            cancer_table_dir.mkdir(parents=True, exist_ok=True)
            cancer_fig_dir.mkdir(parents=True, exist_ok=True)
            fig_base = cancer_fig_dir / "top_pathways_ALL_DATABASES"
            table_path = cancer_table_dir / "ranked_ALL_DATABASES.csv"

        sub.to_csv(table_path, index=False)
        _manifest_add(
            manifest, table_path, "summary_table",
            "Cancer-specific pathway hits ranked across all drivers and databases.",
            cancer=str(cancer),
        )
        cancer_written = _plot_ranked_pathways(
            sub,
            fig_base,
            title=f"{cancer}: top enriched gene sets across all databases",
            formats=fig_formats,
            top_n=summary_top_n,
            fdr_cutoff=fdr_cutoff,
            max_label_chars=max_label_chars,
        )
        for p in cancer_written:
            _manifest_add(
                manifest, p, "summary_figure",
                "Cancer-specific ranked pathway summary figure.",
                cancer=str(cancer),
            )

    # Per-driver ranked summaries across databases, so NRAS/TP53/etc can be browsed directly.
    if write_per_driver_summary:
        if flat_output:
            per_driver_table_root = summary_table_dir / "per_driver"
            per_driver_fig_root = summary_fig_dir / "per_driver"
            per_driver_table_root.mkdir(parents=True, exist_ok=True)
            per_driver_fig_root.mkdir(parents=True, exist_ok=True)

        for (cancer, driver), sub in std.groupby(["cancer", "driver"], dropna=False):
            safe_cancer = _safe_name(str(cancer))
            safe_driver = _safe_name(str(driver))
            if flat_output:
                table_path = per_driver_table_root / f"summary_ranked_{safe_cancer}_{safe_driver}_ALL_DATABASES.csv"
                fig_base = per_driver_fig_root / f"summary_top_pathways_{safe_cancer}_{safe_driver}_ALL_DATABASES"
            else:
                d_table = outdir / "tables" / "by_driver" / safe_cancer / safe_driver
                d_fig = outdir / "figures" / "by_driver" / safe_cancer / safe_driver
                d_table.mkdir(parents=True, exist_ok=True)
                d_fig.mkdir(parents=True, exist_ok=True)
                table_path = d_table / "ranked_ALL_DATABASES.csv"
                fig_base = d_fig / "top_pathways_ALL_DATABASES"

            sub.to_csv(table_path, index=False)
            _manifest_add(
                manifest, table_path, "summary_table",
                "Driver-specific pathway hits ranked across all databases.",
                cancer=str(cancer),
                driver=str(driver),
            )
            driver_written = _plot_ranked_pathways(
                sub,
                fig_base,
                title=f"{cancer}/{driver}: top enriched gene sets across all databases",
                formats=fig_formats,
                top_n=summary_top_n,
                fdr_cutoff=fdr_cutoff,
                max_label_chars=max_label_chars,
            )
            for p in driver_written:
                _manifest_add(
                    manifest, p, "summary_figure",
                    "Driver-specific ranked pathway summary figure.",
                    cancer=str(cancer),
                    driver=str(driver),
                )

    print(f"[pathways] summary tables -> {summary_table_dir}")
    print(f"[pathways] summary figures -> {summary_fig_dir}")
    if written:
        preview = ", ".join(p.name for p in written[:6])
        more = "..." if len(written) > 6 else ""
        print(f"[pathways] global summary figures: wrote {len(written)} files ({preview}{more})")
    else:
        print("[pathways] summary figures: no global figures written")


def _write_and_plot(
    pw,
    enr: pd.DataFrame,
    out_csv: Path,
    out_png: Path,
    title: str,
    manifest: list[dict],
    cancer: str,
    driver: str,
    collection: str,
) -> None:
    out_csv.parent.mkdir(parents=True, exist_ok=True)
    out_png.parent.mkdir(parents=True, exist_ok=True)

    enr.to_csv(out_csv, index=False)
    _manifest_add(
        manifest, out_csv, "table",
        "Pathway enrichment table for one cancer/driver/database.",
        cancer=cancer,
        driver=driver,
        collection=collection,
    )

    try:
        pw.plot_pathway_bar(enr, out_png, title=title, direction="combined")
        _manifest_add(
            manifest, out_png, "figure",
            "Individual pathway enrichment barplot for one cancer/driver/database.",
            cancer=cancer,
            driver=driver,
            collection=collection,
        )
    except Exception as e:
        print(f"[pathways] WARNING: plot failed for {out_csv.name}: {e}")


def _write_output_readme(
    outdir: Path,
    flat_output: bool,
    resources: list[Resource],
    gene_table_files: list[str],
    background: list[str] | None,
    args: argparse.Namespace,
    manifest: list[dict],
) -> None:
    """Write a small human-readable guide to the output directory."""
    layout = "flat legacy" if flat_output else "organized"
    lines = [
        "scOPE pathway analysis outputs",
        "==============================",
        "",
        f"Layout: {layout}",
        f"Output directory: {outdir}",
        f"Input gene tables: {len(gene_table_files)}",
        f"Background genes: {len(background):,}" if background else "Background genes: not provided",
        f"Top genes per programme: {args.top_n}",
        f"By-direction analysis: {bool(args.by_direction)}",
        f"Summary FDR cutoff: {args.summary_fdr_cutoff:g}",
        "",
        "Pathway resources:",
    ]
    for label, src, online in resources:
        mode = "gseapy/enrichr" if online else "offline-hypergeom"
        lines.append(f"  - {label} [{mode}] <- {src}")

    lines.extend(["", "Where to look first:"])
    if flat_output:
        lines.extend(
            [
                "  - pathways_ALL.csv",
                "  - summary_figures/summary_top_pathways_ALL_DATABASES.png",
                "  - summary_figures/summary_top_pathways_heatmap_by_driver.png",
                "  - outputs_manifest.csv",
            ]
        )
    else:
        lines.extend(
            [
                "  - index/top_250_pathways_ranked_all_databases.csv",
                "  - figures/summary/summary_top_pathways_ALL_DATABASES.png",
                "  - figures/summary/summary_top_pathways_heatmap_by_driver.png",
                "  - tables/by_driver/<CANCER>/<DRIVER>/ranked_ALL_DATABASES.csv",
                "  - figures/by_driver/<CANCER>/<DRIVER>/top_pathways_ALL_DATABASES.png",
                "  - outputs_manifest.csv",
            ]
        )

    if not flat_output:
        lines.extend(
            [
                "",
                "Organized directory map:",
                "  index/                         compact CSVs to open first",
                "  tables/pathways_ALL.csv          complete combined table",
                "  tables/summary/                  global ranked/significant summary tables",
                "  tables/by_cancer/<CANCER>/       cancer-level tables",
                "  tables/by_driver/<CANCER>/<DRIVER>/",
                "                                  per-driver tables and ranked all-database hits",
                "  tables/by_database/<DATABASE>/   database-specific combined tables",
                "  figures/summary/                 global summary figures",
                "  figures/by_cancer/<CANCER>/      cancer-level summary figures",
                "  figures/by_driver/<CANCER>/<DRIVER>/",
                "                                  individual database plots and all-database summaries",
            ]
        )

    lines.extend(
        [
            "",
            "Manifest:",
            "  outputs_manifest.csv records the important files with kind/cancer/driver/database labels.",
            "",
        ]
    )

    readme_path = outdir / "README_outputs.txt"
    readme_path.write_text("\n".join(lines), encoding="utf-8")
    _manifest_add(manifest, readme_path, "readme", "Human-readable guide to the output directory.")


def _write_manifest(outdir: Path, manifest: list[dict]) -> Path:
    path = outdir / "outputs_manifest.csv"
    if manifest:
        pd.DataFrame(manifest).drop_duplicates().to_csv(path, index=False)
    else:
        pd.DataFrame(columns=["kind", "path", "cancer", "driver", "collection", "description"]).to_csv(path, index=False)
    return path


# -----------------------------
# Main
# -----------------------------

def main() -> None:
    ap = argparse.ArgumentParser(formatter_class=argparse.ArgumentDefaultsHelpFormatter)
    ap.add_argument("--tables-dir", required=True, help="scope_analysis_outputs/tables")
    ap.add_argument(
        "--gmt",
        action="append",
        nargs="+",
        default=[],
        help="one or more local .gmt gene-set files; can be repeated",
    )
    ap.add_argument(
        "--gmt-dir",
        action="append",
        nargs="+",
        default=[],
        help="directory/directories containing GMTs, or a glob pattern such as msigdb/*.gmt",
    )
    ap.add_argument("--gmt-glob", default="*.gmt", help="glob used inside each --gmt-dir directory")
    ap.add_argument(
        "--enrichr-libraries",
        action="append",
        nargs="+",
        default=[],
        help="online Enrichr libraries, e.g. GO_Biological_Process_2025 KEGG_2021_Human Reactome_2022",
    )
    ap.add_argument(
        "--enrichr-preset",
        choices=["none", "go", "go_kegg", "core"],
        default="none",
        help="online Enrichr preset; requires gseapy + internet",
    )
    ap.add_argument(
        "--list-enrichr-libraries",
        action="store_true",
        help="print available Enrichr libraries and exit; requires gseapy + internet",
    )
    ap.add_argument("--background-genes", default=None, help="optional txt, one symbol/line")
    ap.add_argument("--outdir", default="pathways")
    ap.add_argument("--top-n", type=int, default=100, help="leading genes per programme")
    ap.add_argument("--by-direction", action="store_true", help="also test up/down gene sets separately")
    ap.add_argument(
        "--use-gseapy",
        action="store_true",
        help="use gseapy.enrichr for local GMTs too; online Enrichr libraries always use gseapy",
    )
    ap.add_argument(
        "--summary-top-n",
        type=int,
        default=50,
        help="number of top enrichment rows to show in ranked summary barplots",
    )
    ap.add_argument(
        "--heatmap-top-n",
        type=int,
        default=40,
        help="number of unique term/database pairs to show in the summary heatmap",
    )
    ap.add_argument(
        "--summary-fdr-cutoff",
        type=float,
        default=0.05,
        help="FDR cutoff shown as a dashed line and used for database count plots",
    )
    ap.add_argument(
        "--fig-formats",
        nargs="+",
        default=["png", "pdf"],
        help="figure formats for summary figures",
    )
    ap.add_argument(
        "--no-per-driver-summary-figures",
        action="store_true",
        help="skip per-driver all-database ranked summary figures if there are too many outputs",
    )
    ap.add_argument(
        "--max-label-chars",
        type=int,
        default=160,
        help="truncate long pathway labels in summary figures after this many characters",
    )
    ap.add_argument(
        "--flat-output",
        action="store_true",
        help="use the older single-folder output layout instead of the organized tables/figures/index layout",
    )
    args = ap.parse_args()

    if args.list_enrichr_libraries:
        import gseapy as gp  # type: ignore

        try:
            libs = gp.get_library_name(organism="Human")
        except TypeError:
            libs = gp.get_library_name()
        for lib in libs:
            print(lib)
        return

    from scope_analysis import pathways as pw

    outdir = Path(args.outdir)
    outdir.mkdir(parents=True, exist_ok=True)

    roots = _layout_roots(outdir, flat_output=args.flat_output)
    for d in roots.values():
        d.mkdir(parents=True, exist_ok=True)

    background = _load_background(args.background_genes)
    resources = _build_resources(args)
    files = _find_gene_tables(args.tables_dir)
    manifest: list[dict] = []

    print(f"[pathways] output layout: {'flat legacy' if args.flat_output else 'organized'}")
    print(f"[pathways] output root: {outdir}")
    print(f"[pathways] gene tables: {len(files)}")
    print("[pathways] resources:")
    for label, src, online in resources:
        mode = "gseapy/enrichr" if online else "offline-hypergeom"
        print(f"  - {label} [{mode}] <- {src}")
    if background:
        print(f"[pathways] background genes: {len(background):,}")
    else:
        print("[pathways] background genes: not provided; enrichment function will use its default")

    # Save resource and input manifests early.
    resource_manifest = roots["index"] / "resources_manifest.csv"
    pd.DataFrame(
        [
            {"collection": label, "source": src, "mode": "gseapy/enrichr" if online else "offline-hypergeom"}
            for label, src, online in resources
        ]
    ).to_csv(resource_manifest, index=False)
    _manifest_add(manifest, resource_manifest, "index", "Pathway resources used in this run.")

    input_manifest = roots["index"] / "input_gene_tables.csv"
    pd.DataFrame({"input_gene_table": files}).to_csv(input_manifest, index=False)
    _manifest_add(manifest, input_manifest, "index", "Input gene-program tables used in this run.")

    combined_all: list[pd.DataFrame] = []
    n_driver_resource_tables = 0

    for f in files:
        cancer, groups = _driver_groups(f)
        if not groups:
            print(f"[pathways] {f}: skipped because no 'gene' column")
            continue

        for drv, sub in groups:
            cancer_s = str(cancer)
            driver_s = str(drv)
            per_driver: list[pd.DataFrame] = []

            for collection, gene_sets, use_gseapy in resources:
                collection_s = str(collection)
                try:
                    enr = pw.enrich_driver_programs(
                        sub,
                        gene_sets,
                        background=background,
                        top_n=args.top_n,
                        by_direction=args.by_direction,
                        use_gseapy=use_gseapy,
                    )
                except Exception as e:
                    print(f"[pathways] ERROR {cancer_s}/{driver_s}/{collection_s}: {e}")
                    continue

                if enr is None or enr.empty:
                    print(f"[pathways] {cancer_s}/{driver_s}/{collection_s}: no enrichment")
                    continue

                enr = enr.copy()
                enr.insert(0, "gene_set_source", str(gene_sets))
                enr.insert(0, "collection", collection_s)
                enr.insert(0, "driver", driver_s)
                enr.insert(0, "cancer", cancer_s)

                out_csv, out_png = _individual_output_paths(
                    outdir,
                    flat_output=args.flat_output,
                    cancer=cancer_s,
                    driver=driver_s,
                    collection=collection_s,
                )
                _write_and_plot(
                    pw,
                    enr,
                    out_csv,
                    out_png,
                    title=f"{cancer_s} · {driver_s} — {collection_s}",
                    manifest=manifest,
                    cancer=cancer_s,
                    driver=driver_s,
                    collection=collection_s,
                )

                fdr_col = _fdr_column(enr)
                if fdr_col:
                    fdrs = pd.to_numeric(enr[fdr_col], errors="coerce")
                    sig = int((fdrs < args.summary_fdr_cutoff).sum())
                    top_fdr = fdrs.iloc[0]
                    print(
                        f"[pathways] {cancer_s}/{driver_s}/{collection_s}: "
                        f"{sig} sets FDR<{args.summary_fdr_cutoff:g}; "
                        f"top: {_first_term(enr)[:60]} (FDR={top_fdr:.1e})"
                    )
                else:
                    print(f"[pathways] {cancer_s}/{driver_s}/{collection_s}: wrote {len(enr)} rows")

                per_driver.append(enr)
                combined_all.append(enr)
                n_driver_resource_tables += 1

            if per_driver:
                driver_all = pd.concat(per_driver, ignore_index=True)
                driver_path = _all_collections_path(
                    outdir,
                    flat_output=args.flat_output,
                    cancer=cancer_s,
                    driver=driver_s,
                )
                driver_all.to_csv(driver_path, index=False)
                _manifest_add(
                    manifest,
                    driver_path,
                    "table",
                    "Combined all-database pathway enrichment table for one cancer/driver.",
                    cancer=cancer_s,
                    driver=driver_s,
                )

    if combined_all:
        allt = pd.concat(combined_all, ignore_index=True)

        if args.flat_output:
            all_path = outdir / "pathways_ALL.csv"
        else:
            all_path = outdir / "tables" / "pathways_ALL.csv"
            all_path.parent.mkdir(parents=True, exist_ok=True)

        allt.to_csv(all_path, index=False)
        _manifest_add(
            manifest, all_path, "table",
            "Complete combined pathway enrichment table across all cancers, drivers, and databases.",
        )

        _write_grouped_tables(
            all_results=allt,
            outdir=outdir,
            flat_output=args.flat_output,
            manifest=manifest,
        )
        _write_top_hit_indexes(
            all_results=allt,
            outdir=outdir,
            flat_output=args.flat_output,
            manifest=manifest,
            fdr_cutoff=args.summary_fdr_cutoff,
        )
        _write_summary_figures(
            all_results=allt,
            outdir=outdir,
            summary_top_n=args.summary_top_n,
            heatmap_top_n=args.heatmap_top_n,
            fig_formats=args.fig_formats,
            fdr_cutoff=args.summary_fdr_cutoff,
            write_per_driver_summary=not args.no_per_driver_summary_figures,
            max_label_chars=args.max_label_chars,
            flat_output=args.flat_output,
            manifest=manifest,
        )

        _write_output_readme(
            outdir=outdir,
            flat_output=args.flat_output,
            resources=resources,
            gene_table_files=files,
            background=background,
            args=args,
            manifest=manifest,
        )
        manifest_path = _write_manifest(outdir, manifest)

        print(
            f"\n[pathways] wrote {n_driver_resource_tables} driver/resource tables + "
            f"combined summaries to {outdir}"
        )
        print(f"[pathways] manifest -> {manifest_path}")
        if not args.flat_output:
            print("[pathways] start here:")
            print(f"  - {outdir / 'README_outputs.txt'}")
            print(f"  - {outdir / 'index' / 'top_250_pathways_ranked_all_databases.csv'}")
            print(f"  - {outdir / 'figures' / 'summary' / 'summary_top_pathways_ALL_DATABASES.png'}")
    else:
        _write_output_readme(
            outdir=outdir,
            flat_output=args.flat_output,
            resources=resources,
            gene_table_files=files,
            background=background,
            args=args,
            manifest=manifest,
        )
        manifest_path = _write_manifest(outdir, manifest)
        print("\n[pathways] no enrichment results were produced")
        print(f"[pathways] manifest -> {manifest_path}")


if __name__ == "__main__":
    main()
```
