## Supplemental_Code for "scOPE identifies which driver-associated expression programs transfer from bulk tumors to single cells": make_umap_panels.py

```
#!/usr/bin/env python
"""
make_umap_panels.py
===================
Render the high-quality multi-panel UMAPs (patient | ground-truth genotype |
predicted score) from a scored single-cell h5ad, WITHOUT re-running the pipeline.

Usage
-----
# single object (must contain mutation_calib_*/mutation_prob_* in obs):
python drivers/make_umap_panels.py \
    --scored-h5ad data/.../AML_scored_with_muttranscript_eval_labels.h5ad \
    --confidence-table scope_analysis_outputs/tables/Fig3_confidence_AML_vanGalen.csv \
    --patient-col Sample --celltype-col CellType \
    --umap-source hvg --hvg-top-genes 3000 \
    --outdir scope_analysis_outputs/figures/fig4_umap_panels --key AML_vanGalen

# or iterate every cohort in your config that has a scored_h5ad:
python drivers/make_umap_panels.py --config drivers/config_canonical_v3.json \
    --umap-source hvg --hvg-top-genes 3000 --outdir scope_analysis_outputs/figures/fig4_umap_panels
"""
from __future__ import annotations
import argparse, json
from pathlib import Path

import numpy as np
import pandas as pd
import anndata as ad

import scope_analysis as sa
from scope_analysis import umap_panels as up
from scope_analysis.config import AnalysisConfig


def _matrix_quick_stats(X, *, max_cells=5000, max_genes=3000):
    import numpy as np
    try:
        import scipy.sparse as sp
    except Exception:
        sp = None
    out = {
        "min": float("nan"), "max": float("nan"), "p99": float("nan"), "p999": float("nan"),
        "median_library": float("nan"), "integer_fraction": float("nan"),
        "has_negative": False, "is_sparse": bool(sp is not None and sp.issparse(X)),
    }
    try:
        nr = min(max_cells, int(X.shape[0])); nc = min(max_genes, int(X.shape[1]))
        xs = X[:nr, :nc]
        lib = np.asarray(X[:nr, :].sum(axis=1)).ravel()
        out["median_library"] = float(np.nanmedian(lib)) if lib.size else float("nan")
        if sp is not None and sp.issparse(xs):
            data = np.asarray(xs.data, dtype=float)
            if data.size:
                out["min"] = float(np.nanmin(data)); out["max"] = float(np.nanmax(data))
                out["p99"] = float(np.nanpercentile(data, 99.0)); out["p999"] = float(np.nanpercentile(data, 99.9))
                out["has_negative"] = bool(np.nanmin(data) < -1e-8)
                out["integer_fraction"] = float(np.nanmean(np.isclose(data, np.round(data), atol=1e-8)))
            else:
                out.update({"min": 0.0, "max": 0.0, "p99": 0.0, "p999": 0.0, "integer_fraction": 1.0})
        else:
            flat = np.asarray(xs, dtype=float).ravel()
            flat = flat[np.isfinite(flat)]
            if flat.size:
                out["min"] = float(np.nanmin(flat)); out["max"] = float(np.nanmax(flat))
                out["p99"] = float(np.nanpercentile(flat, 99.0)); out["p999"] = float(np.nanpercentile(flat, 99.9))
                out["has_negative"] = bool(np.nanmin(flat) < -1e-8)
                nz = flat[np.abs(flat) > 1e-12]
                out["integer_fraction"] = float(np.nanmean(np.isclose(nz, np.round(nz), atol=1e-8))) if nz.size else 1.0
    except Exception:
        pass
    return out


def _looks_count_like(stats):
    max_val = float(stats.get("max", np.nan)); p999 = float(stats.get("p999", np.nan))
    p99 = float(stats.get("p99", np.nan)); med_lib = float(stats.get("median_library", np.nan))
    int_frac = float(stats.get("integer_fraction", np.nan))
    if bool(stats.get("has_negative", False)):
        return False
    if np.isfinite(max_val) and max_val > 50:
        return True
    if np.isfinite(p999) and p999 > 30:
        return True
    if np.isfinite(int_frac) and int_frac > 0.98:
        if np.isfinite(p99) and p99 > 5:
            return True
        if np.isfinite(med_lib) and med_lib > 5000:
            return True
    return False


def _choose_hvg_source(adata):
    """Use raw when adata.X appears scaled; otherwise use adata.X."""
    try:
        if getattr(adata, "n_vars", 0) >= 500:
            stats = _matrix_quick_stats(adata.X)
            if stats.get("has_negative", False) and adata.raw is not None and adata.raw.shape[1] >= 500:
                print("[umap] adata.X has negative/scaled values; using adata.raw for HVG UMAP source.")
                return adata.raw.to_adata()
            return adata
    except Exception:
        pass
    try:
        if adata.raw is not None and adata.raw.shape[1] >= 500:
            print("[umap] using adata.raw for HVG UMAP source.")
            return adata.raw.to_adata()
    except Exception:
        pass
    return adata


def _hvg_umap(adata, top_genes=3000, n_pcs=50, n_neighbors=15, min_dist=0.5, seed=42,
              preprocess="auto", scale="auto", scale_max_value=10.0):
    import scanpy as sc
    source = _choose_hvg_source(adata)
    a = source.copy()

    preprocess = str(preprocess or "auto").lower().replace("-", "_")
    stats = _matrix_quick_stats(a.X)
    print(
        "[umap] HVG source quick stats: "
        f"min={stats.get('min', float('nan')):.3g}, max={stats.get('max', float('nan')):.3g}, "
        f"p99={stats.get('p99', float('nan')):.3g}, p999={stats.get('p999', float('nan')):.3g}, "
        f"median_lib={stats.get('median_library', float('nan')):.3g}, "
        f"integer_frac={stats.get('integer_fraction', float('nan')):.3g}"
    )
    if preprocess in {"normalize_log1p", "normalize", "log1p", "counts"}:
        print("[umap] forced normalize_total + log1p for HVG UMAP only.")
        sc.pp.normalize_total(a, target_sum=1e4)
        sc.pp.log1p(a)
        preprocess_mode = "normalize_log1p_forced"
    elif preprocess in {"none", "false", "off", "no"}:
        print("[umap] HVG preprocessing disabled; using expression as-is.")
        preprocess_mode = "none"
    elif stats.get("has_negative", False):
        print("[umap] expression has negative values; assuming already scaled/transformed and skipping normalize/log1p.")
        preprocess_mode = "as_is_negative_scaled"
    elif _looks_count_like(stats):
        print("[umap] expression looks count-like; running normalize_total + log1p for UMAP only.")
        sc.pp.normalize_total(a, target_sum=1e4)
        sc.pp.log1p(a)
        preprocess_mode = "normalize_log1p_auto"
    else:
        print("[umap] expression looks log-normalized-ish; using as-is before HVG selection.")
        preprocess_mode = "as_is_logish"

    try:
        sc.pp.highly_variable_genes(a, n_top_genes=min(int(top_genes), a.n_vars), flavor="seurat")
        mask = np.asarray(a.var.get("highly_variable", False), dtype=bool)
        if mask.sum() >= 10:
            a = a[:, mask].copy()
        else:
            raise ValueError(f"only {int(mask.sum())} HVGs")
    except Exception as exc:
        print(f"[umap] scanpy HVG failed ({type(exc).__name__}: {exc}); using variance-ranked genes.")
        X = a.X
        try:
            import scipy.sparse as sp
            if sp.issparse(X):
                mean = np.asarray(X.mean(axis=0)).ravel()
                mean_sq = np.asarray(X.multiply(X).mean(axis=0)).ravel()
                var = mean_sq - mean ** 2
            else:
                var = np.nanvar(np.asarray(X), axis=0)
        except Exception:
            var = np.nanvar(np.asarray(X), axis=0)
        idx = np.argsort(np.asarray(var, dtype=float))[::-1][:min(int(top_genes), a.n_vars)]
        a = a[:, idx].copy()

    scale = str(scale or "auto").lower()
    dense_gb = a.n_obs * a.n_vars * 4 / (1024 ** 3)
    did_scale = False
    if scale in {"never", "none", "false", "off", "no"}:
        print("[umap] HVG scaling disabled; sparse PCA zero_center=False.")
    elif scale in {"auto", "safe"} and dense_gb > 8.0:
        print(f"[umap] skipping HVG scaling to avoid dense memory blowup (estimated dense={dense_gb:.2f} GB).")
    else:
        try:
            print(f"[umap] scaling HVGs for UMAP/PCA only (max_value={float(scale_max_value):g}, estimated dense={dense_gb:.2f} GB).")
            sc.pp.scale(a, max_value=float(scale_max_value))
            did_scale = True
        except Exception as exc:
            print(f"[umap] scaling failed ({type(exc).__name__}: {exc}); using sparse PCA zero_center=False.")

    pcs_eff = max(2, min(int(n_pcs), a.n_vars - 1, a.n_obs - 1))
    sc.tl.pca(a, n_comps=pcs_eff, svd_solver="arpack", zero_center=did_scale, random_state=int(seed))
    sc.pp.neighbors(a, n_pcs=pcs_eff, n_neighbors=int(n_neighbors), random_state=int(seed))
    sc.tl.umap(a, min_dist=float(min_dist), random_state=int(seed))
    if a.obsm["X_umap"].shape[0] != adata.n_obs:
        raise ValueError(f"UMAP row mismatch: got {a.obsm['X_umap'].shape[0]}, expected {adata.n_obs}")
    if "X_umap" in adata.obsm and "X_umap_original_once" not in adata.obsm:
        adata.obsm["X_umap_original_once"] = adata.obsm["X_umap"].copy()
    adata.obsm[f"X_umap_hvg{int(top_genes)}"] = np.asarray(a.obsm["X_umap"], dtype=np.float32)
    adata.obsm["X_umap"] = adata.obsm[f"X_umap_hvg{int(top_genes)}"].copy()
    adata.uns["scope_umap_source"] = {
        "mode": "hvg", "preprocess_mode": preprocess_mode, "scaled": bool(did_scale),
        "n_top_genes": int(top_genes), "n_pcs": int(pcs_eff),
        "n_neighbors": int(n_neighbors), "min_dist": float(min_dist),
    }
    return adata

def _prep(adata, cfg, spec, args):
    # Standalone plotting must be current-config deterministic too: keep raw
    # mutation_prob_* but wipe stale derived score columns from older runs.
    adata, _ = sa.singlecell.clear_scope_score_columns(
        adata, include_raw=False, include_derived=True, label="umap-panels derived reset")
    adata = sa.data.ensure_patient_metadata(
        adata, sample_col=spec.get("sample_col") or None,
        patient_strip_regex=spec.get("patient_strip_regex", r"-D\d+$"),
        timepoint_regex=spec.get("timepoint_regex", r"-(D\d+)$"),
    )
    celltype_col = spec.get("celltype_col", args.celltype_col)
    try:
        adata, celltype_col = sa.annotate.ensure_celltype(adata, celltype_col)
    except Exception as e:
        print(f"[umap-panels] marker-based celltype fallback skipped ({e})")
    try:
        adata, _coarse_col = sa.reference.ensure_coarse_cellstate(
            adata, celltype_col=celltype_col, out_col="celltype_coarse")
        if spec.get("use_coarse_celltype", False):
            celltype_col = _coarse_col
    except Exception as e:
        print(f"[umap-panels] coarse cell-state labels skipped ({e})")
    adata = sa.data.mark_reference_cells(
        adata, celltype_col=celltype_col,
        reference_celltypes=tuple(spec.get("reference_celltypes", ("T", "B", "NK"))),
        sample_col=spec.get("sample_col", "Sample"),
        reference_samples=spec.get("reference_samples"))
    drivers = sa.data.scored_drivers(adata)
    if getattr(cfg, "enable_reference_guardrails", True) and drivers:
        try:
            # Standalone UMAP rendering should match run_all: add same-cell-state
            # guarded scores before choosing score columns for score panels.
            adata, _guard_tbl = sa.reference.add_cellstate_guarded_scores(
                adata, cfg, drivers=drivers,
                reference_mask_col="is_reference",
                cellstate_col=getattr(cfg, "cellstate_col", "celltype_coarse"),
                min_reference_per_state=getattr(cfg, "cellstate_min_reference_cells", 25),
            )
        except Exception as e:
            print(f"[umap-panels] guarded scores skipped ({e}); falling back to raw/configured.")
    if cfg.use_calibrated_scores:
        adata = sa.singlecell.calibrate_scores(adata, cfg, drivers=drivers)
    else:
        print("[umap-panels] use_calibrated_scores=false; plotting raw/adjusted scores.")
    try:
        adata, _ = sa.singlecell.derive_malignant_compartment(adata, cfg, drivers=drivers)
    except Exception:
        pass
    if args.umap_source == "hvg" and adata.X is not None:
        try:
            adata = _hvg_umap(
                adata,
                top_genes=args.hvg_top_genes,
                n_pcs=args.hvg_n_pcs,
                n_neighbors=args.hvg_n_neighbors,
                min_dist=args.hvg_min_dist,
                seed=args.umap_random_state,
                preprocess=args.hvg_preprocess,
                scale=args.hvg_scale,
                scale_max_value=args.hvg_scale_max_value,
            )
        except Exception as e:
            print(f"[umap] HVG UMAP failed ({e}); falling back to existing/X_svd.")
    return adata, drivers


def _patient_status(path, adata=None):
    if path and Path(path).exists():
        status = sa.data.load_patient_status(path)
        if adata is not None and "patient" in adata.obs.columns:
            status = sa.data.harmonize_patient_status(status, adata.obs["patient"].astype(str).values)
            status = sa.data.validate_patient_status_for_obs(status, adata.obs["patient"].astype(str).values, cohort_name="UMAP")
        return status
    return None


def _confidence_map(path):
    if path and Path(path).exists():
        df = pd.read_csv(path)
        if {"driver", "confidence"}.issubset(df.columns):
            return dict(zip(df["driver"], df["confidence"]))
    return {}


def _cfg_from_raw(raw):
    r = raw.get("rigor", {})
    plotting = raw.get("plotting", {})
    return AnalysisConfig(
        use_calibrated_scores=r.get("use_calibrated_scores", False),
        calibration_method=r.get("calibration_method", "reference_ecdf"),
        longitudinal_use_calibrated_scores=r.get("longitudinal_use_calibrated_scores", False),
        plot_score_scale=plotting.get("score_scale", r.get("plot_score_scale", "auto_robust")),
        plot_robust_lower_q=plotting.get("robust_lower_q", r.get("plot_robust_lower_q", 0.01)),
        plot_robust_upper_q=plotting.get("robust_upper_q", r.get("plot_robust_upper_q", 0.99)),
        plot_robust_min_range=plotting.get("robust_min_range", r.get("plot_robust_min_range", 0.05)),
        heatmap_score_scale=plotting.get("heatmap_score_scale", r.get("heatmap_score_scale", "driver_robust")),
        score_umap_grey_mask=plotting.get("score_umap_grey_mask", r.get("score_umap_grey_mask", "reference")),
        enable_reference_guardrails=r.get("enable_reference_guardrails", True),
        reference_guardrail_samples=tuple(r.get("reference_guardrail_samples", ("BM1", "BM2", "BM3", "BM4"))),
        cellstate_min_reference_cells=r.get("cellstate_min_reference_cells", 25),
        longitudinal_guarded_mode=r.get("longitudinal_guarded_mode", "cellstate_residual"),
    )


def _render(adata, drivers, key, conf_map, outdir, patient_col, cfg=None, patient_status=None):
    outdir = Path(outdir); outdir.mkdir(parents=True, exist_ok=True)
    if conf_map:
        try:
            conf_table = pd.DataFrame({"driver": list(conf_map.keys()), "confidence": list(conf_map.values())})
            adata = sa.confidence.add_confidence_adjusted_scores(adata, conf_table)
            print(f"[umap-panels] recomputed mutation_adj_* for {len(conf_map)} drivers from confidence table.")
        except Exception as e:
            print(f"[umap-panels] confidence-adjusted scores skipped ({e}); using raw/guarded scores.")
    guarded_available = any((f"mutation_resid_{d}" in adata.obs.columns) or (f"mutation_refpct_{d}" in adata.obs.columns) for d in drivers)
    score_prefer = ("resid", "refpct", "adj", "raw", "calib") if guarded_available else ("adj", "raw", "calib")
    pcol = patient_col if patient_col in adata.obs.columns else (
        "patient" if "patient" in adata.obs.columns else None)
    audit = up.triptych_genotype_audit(adata, drivers, patient_status=patient_status, patient_col=pcol or "patient")
    if audit is not None and not audit.empty:
        ap = outdir / f"umap_triptych_genotype_audit_{key}.csv"
        audit.to_csv(ap, index=False)
        print(f"[umap-panels] wrote {ap}")
    p1 = up.driver_triptych_grid(
        adata, drivers, outdir / f"umap_triptych_{key}.png", cfg=cfg,
        confidence=conf_map, patient_col=pcol or "patient", only_genotyped=True,
        score_prefer=score_prefer, patient_status=patient_status)
    p2 = up.context_strip(adata, outdir / f"umap_context_{key}.png",
                          patient_col=pcol or "patient")
    # score-only grid for ALL drivers (no genotype needed)
    p3 = up.driver_triptych_grid(
        adata, drivers, outdir / f"umap_triptych_alldrivers_{key}.png", cfg=cfg,
        confidence=conf_map, patient_col=pcol or "patient", only_genotyped=False,
        max_rows=24, score_prefer=score_prefer, patient_status=patient_status)
    for p in (p1, p2, p3):
        if p:
            print(f"[umap-panels] wrote {p}")


def main():
    ap = argparse.ArgumentParser()
    ap.add_argument("--config", default=None)
    ap.add_argument("--scored-h5ad", default=None)
    ap.add_argument("--key", default="cohort")
    ap.add_argument("--confidence-table", default=None)
    ap.add_argument("--patient-status-csv", default=None)
    ap.add_argument("--patient-col", default="patient")
    ap.add_argument("--celltype-col", default="CellType")
    ap.add_argument("--umap-source", default="auto", choices=("auto", "hvg"))
    ap.add_argument("--hvg-top-genes", type=int, default=3000)
    ap.add_argument("--hvg-n-pcs", type=int, default=50)
    ap.add_argument("--hvg-n-neighbors", type=int, default=15)
    ap.add_argument("--hvg-min-dist", type=float, default=0.5)
    ap.add_argument("--hvg-preprocess", default="auto", choices=("auto", "none", "normalize_log1p"))
    ap.add_argument("--hvg-scale", default="auto", choices=("auto", "never", "always"))
    ap.add_argument("--hvg-scale-max-value", type=float, default=10.0)
    ap.add_argument("--umap-random-state", type=int, default=42)
    ap.add_argument("--outdir", default="umap_panels")
    args = ap.parse_args()
    cfg = _cfg_from_raw(json.load(open(args.config))) if args.config else AnalysisConfig(use_calibrated_scores=False)

    if args.scored_h5ad:
        adata = sa.data.load_scored_singlecell(args.scored_h5ad)
        spec = {"sample_col": args.patient_col if args.patient_col != "patient" else None,
                "celltype_col": args.celltype_col}
        adata, drivers = _prep(adata, cfg, spec, args)
        status = _patient_status(args.patient_status_csv, adata)
        _render(adata, drivers, args.key, _confidence_map(args.confidence_table),
                args.outdir, args.patient_col, cfg=cfg, patient_status=status)
        return

    if not args.config:
        raise SystemExit("Provide --scored-h5ad or --config.")
    raw = json.load(open(args.config))
    cfg = _cfg_from_raw(raw)
    for key, spec in raw.get("singlecell_cohorts", {}).items():
        sp = spec.get("scored_h5ad")
        if not (sp and Path(sp).exists()):
            print(f"[umap-panels] {key}: no scored_h5ad on disk; skipping "
                  "(point scored_h5ad at a scored object, or save one from the run).")
            continue
        print(f"[umap-panels] {key}: {sp}")
        adata = sa.data.load_scored_singlecell(sp)
        adata, drivers = _prep(adata, cfg, spec, args)
        conf = _confidence_map(
            Path(args.outdir).parent.parent / "tables" / f"Fig3_confidence_{key}.csv")
        status = _patient_status(spec.get("patient_status_csv"), adata)
        _render(adata, drivers, key, conf, args.outdir,
                spec.get("sample_col", args.patient_col), cfg=cfg, patient_status=status)


if __name__ == "__main__":
    main()
```
