## Supplemental_Code for "scOPE identifies which driver-associated expression programs transfer from bulk tumors to single cells": run_all.py

```
"""
drivers/run_all.py
==================
Single entry point. Reads a JSON config (see config_template.json), runs every
stage in the correct order against your real data, and writes all tables, figures,
and a MANIFEST.json into the output root.

    python drivers/run_all.py --config drivers/config_template.json
    python drivers/run_all.py --config my.json --only bulk,longitudinal
    python drivers/run_all.py --config my.json --skip robustness

RUN ORDER (and the chapter figure each stage serves)
----------------------------------------------------
    1. bulk          Table 1 / Fig 2   (leakage-free OOF discrimination)
    2. interpret     Fig 2             (SVD gene programs per driver)
    3. singlecell    Fig 3             (per-cell PU + patient bridge)
    4. longitudinal  Fig 3b            (treatment-response trajectory)  <- the gold
    5. pancancer     Fig 5 / Table 2   (tiered validation + null controls)
    6. cnv           Fig 6             (scOPE vs CNV complementarity)
    7. robustness    Robustness Table  (ablations + calibration)
    8. report        manifest + driver-prior table

Every stage is independently guarded: a cohort missing the files a stage needs is
skipped with a printed reason, never a crash, so a partial data set still yields a
partial-but-correct manuscript package.
"""
from __future__ import annotations
import argparse, json, warnings, sys, shutil, tempfile
from pathlib import Path

# Force imports to come from THIS checkout, not an older installed/cached copy.
# This matters when running e.g. `python scope_analysis_v3/drivers/run_all.py`
# from the parent scOPE folder: by default Python puts only drivers/ on sys.path.
PROJECT_ROOT = Path(__file__).resolve().parents[1]
if str(PROJECT_ROOT) not in sys.path:
    sys.path.insert(0, str(PROJECT_ROOT))

warnings.filterwarnings("ignore")
import anndata as ad
import pandas as pd

import scope_analysis as sa
from scope_analysis.config import AnalysisConfig, cancer_drivers, prior_of
print(f"[run_all] imported scope_analysis from: {Path(sa.__file__).resolve()}")


def _parse_driver_list(value):
    """Parse JSON/CLI driver selectors. Returns 'all' or a tuple of gene names."""
    if value is None or value == "":
        return "all"
    if isinstance(value, str):
        v = value.strip()
        if not v or v.lower() == "all":
            return "all"
        return tuple(x.strip().upper() for x in v.split(",") if x.strip())
    if isinstance(value, (list, tuple, set)):
        if len(value) == 1 and str(next(iter(value))).lower() == "all":
            return "all"
        return tuple(str(x).strip().upper() for x in value if str(x).strip())
    return "all"


def load_config(path) -> tuple[AnalysisConfig, dict]:
    raw = json.loads(Path(path).read_text())
    m, r = raw.get("modelling", {}), raw.get("rigor", {})
    interp = raw.get("interpretation", {})
    plotting = raw.get("plotting", {})
    scprep = raw.get("singlecell_preprocessing", raw.get("singlecell_preprocess", {}))
    target_drivers = _parse_driver_list(raw.get("target_drivers", m.get("target_drivers", "all")))
    score_all_label_drivers = bool(raw.get("score_all_label_drivers", m.get("score_all_label_drivers", False)))
    # SVD scaling: honour either nested decomposition_kwargs or a top-level flag,
    # defaulting to the manuscript convention Z = U·Σ (scale_by_singular_values=False).
    dkw = dict(m.get("decomposition_kwargs", {}))
    if "scale_by_singular_values" in m:
        dkw.setdefault("scale_by_singular_values", m["scale_by_singular_values"])
    dkw.setdefault("scale_by_singular_values", False)
    cfg = AnalysisConfig(
        output_root=Path(raw.get("output_root", "scope_analysis_outputs")),
        norm_method=m.get("norm_method", "none"), log1p=m.get("log1p", False),
        decomposition=m.get("decomposition", "svd"),
        n_components=m.get("n_components", 50), classifier=m.get("classifier", "logistic"),
        alignment_method=m.get("alignment_method", "z_score_bulk"),
        min_positive_frac=m.get("min_positive_frac", r.get("min_positive_frac", 0.05)),
        decomposition_kwargs=dkw,
        n_boot=r.get("n_boot", 2000), n_perm=r.get("n_perm", 2000),
        fdr_alpha=r.get("fdr_alpha", 0.10), min_pos=r.get("min_pos", 10),
        min_neg=r.get("min_neg", 10), cv_folds=r.get("cv_folds", 5),
        min_cells_per_patient=r.get("min_cells_per_patient", 10),
        reference_quantile=r.get("reference_quantile", 0.95),
        random_state=r.get("random_state", 42),
        singlecell_preprocessing=scprep or None,
        singlecell_projection_nan_policy=raw.get("singlecell_projection_nan_policy", r.get("singlecell_projection_nan_policy", "zero_fill")),
        max_singlecell_cells=raw.get("max_singlecell_cells", r.get("max_singlecell_cells", 0)),
        score_all_label_drivers=score_all_label_drivers,
        target_drivers=target_drivers,
        interpret_top_components=interp.get("top_components", interp.get("interpret_top_components", r.get("interpret_top_components", 25))),
        interpret_genes_per_component=interp.get("genes_per_component", interp.get("interpret_genes_per_component", r.get("interpret_genes_per_component", 50))),
        interpret_plot_top_components=interp.get("plot_top_components", interp.get("interpret_plot_top_components", r.get("interpret_plot_top_components", 6))),
        interpret_plot_genes_per_component=interp.get("plot_genes_per_component", interp.get("interpret_plot_genes_per_component", r.get("interpret_plot_genes_per_component", 30))),
        interpret_heatmap_top_components=interp.get("heatmap_top_components", interp.get("interpret_heatmap_top_components", r.get("interpret_heatmap_top_components", 10))),
        interpret_heatmap_genes_per_component=interp.get("heatmap_genes_per_component", interp.get("interpret_heatmap_genes_per_component", r.get("interpret_heatmap_genes_per_component", 15))),
        # calibration + malignant-compartment knobs (rigor block, with defaults)
        use_calibrated_scores=r.get("use_calibrated_scores", False),
        calibration_method=r.get("calibration_method", "reference_ecdf"),
        longitudinal_use_calibrated_scores=r.get("longitudinal_use_calibrated_scores", False),
        malignant_call_quantile=r.get("malignant_call_quantile", 0.95),
        restrict_longitudinal_to_malignant=r.get("restrict_longitudinal_to_malignant", True),
        plot_score_scale=plotting.get("score_scale", r.get("plot_score_scale", "auto_robust")),
        plot_robust_lower_q=plotting.get("robust_lower_q", r.get("plot_robust_lower_q", 0.01)),
        plot_robust_upper_q=plotting.get("robust_upper_q", r.get("plot_robust_upper_q", 0.99)),
        plot_robust_min_range=plotting.get("robust_min_range", r.get("plot_robust_min_range", 0.05)),
        plot_residual_min_abs_vmax=plotting.get("residual_min_abs_vmax", r.get("plot_residual_min_abs_vmax", 0.05)),
        plot_residual_abs_vmax=plotting.get("residual_abs_vmax", r.get("plot_residual_abs_vmax", 0.25)),
        heatmap_residual_abs_vmax=plotting.get("heatmap_residual_abs_vmax", r.get("heatmap_residual_abs_vmax", 0.20)),
        heatmap_score_scale=plotting.get("heatmap_score_scale", r.get("heatmap_score_scale", "global_robust")),
        confidence_display_min=plotting.get("confidence_display_min", r.get("confidence_display_min", 0.35)),
        plot_driver_set=plotting.get("plot_driver_set", r.get("plot_driver_set", "primary_if_all_labels")),
        score_umap_grey_mask=plotting.get("score_umap_grey_mask", r.get("score_umap_grey_mask", "reference")),
        enable_reference_guardrails=r.get("enable_reference_guardrails", True),
        reference_guardrail_samples=tuple(r.get("reference_guardrail_samples", ["BM1", "BM2", "BM3", "BM4"])),
        cellstate_col=r.get("cellstate_col", "celltype_coarse"),
        cellstate_min_reference_cells=r.get("cellstate_min_reference_cells", 25),
        longitudinal_guarded_mode=r.get("longitudinal_guarded_mode", "cellstate_residual"),
        target_cancers=tuple(raw.get("bulk_cohorts", {}).keys()) or "all",
    )
    return cfg, raw


def _exists(p) -> bool:
    return bool(p) and Path(p).exists()


def _slug(x) -> str:
    """Filesystem-safe token for per-patient/per-driver output names."""
    return "".join(c if c.isalnum() or c in "._-" else "_" for c in str(x))


def _selected_drivers_for_labels(cancer: str, labels: pd.DataFrame, cfg: AnalysisConfig) -> list[str]:
    """Resolve driver columns to train/score for this cancer.

    Primary mode is the locked DRIVER_PRIORS table. Exploratory mode
    (--score-all-label-drivers / cfg.score_all_label_drivers) uses every binary
    mutation-label column present in the cohort labels file.
    """
    label_cols = [str(c).strip().upper() for c in labels.columns]
    # Preserve original column spelling/order but compare uppercase.
    original_by_upper = {str(c).strip().upper(): str(c).strip() for c in labels.columns}
    if getattr(cfg, "score_all_label_drivers", False):
        wanted_upper = label_cols
    elif getattr(cfg, "target_drivers", "all") == "all":
        wanted_upper = [str(d).strip().upper() for d in cancer_drivers(cancer)]
    else:
        wanted_upper = [str(d).strip().upper() for d in getattr(cfg, "target_drivers", ())]
    seen = set()
    present = []
    for d in wanted_upper:
        if d in seen:
            continue
        seen.add(d)
        if d in original_by_upper:
            present.append(original_by_upper[d])
    return present


def _plot_drivers_for_stage(cancer_type: str | None, drivers: list[str], cfg: AnalysisConfig, conf_map: dict | None = None) -> list[str]:
    """Choose which drivers should appear in crowded Fig4 panels.

    Broad exploratory scoring can train every mutation-label column, but the main
    UMAPs/heatmaps should not automatically include rare/noisy labels.  Scores and
    audit tables still contain all trained drivers; this only controls display.
    """
    drivers = list(drivers or [])
    mode = str(getattr(cfg, "plot_driver_set", "primary_if_all_labels") or "primary_if_all_labels").lower()
    conf_map = conf_map or {}
    if mode in {"all", "all_scored", "scored"}:
        out = drivers
    elif mode in {"confidence", "confident"}:
        thr = float(getattr(cfg, "confidence_display_min", 0.35))
        out = [d for d in drivers if float(conf_map.get(d, -1)) >= thr]
    else:
        use_primary = mode == "primary" or (mode in {"primary_if_all_labels", "primary_if_exploratory"} and getattr(cfg, "score_all_label_drivers", False))
        if use_primary and cancer_type:
            primary = {str(d).upper() for d in cancer_drivers(cancer_type)}
            out = [d for d in drivers if str(d).upper() in primary]
            if not out:
                out = drivers
        else:
            out = drivers
    if conf_map:
        out = sorted(out, key=lambda d: float(conf_map.get(d, -1)), reverse=True)
    if len(out) != len(drivers):
        print(f"[plot-drivers] display set={mode!r}: plotting {len(out)}/{len(drivers)} scored drivers: {out}")
    return out


def _driver_selection_audit(cancer: str, labels: pd.DataFrame, selected: list[str],
                            trained: list[str] | None = None, cfg: AnalysisConfig | None = None,
                            stage: str = "fit") -> pd.DataFrame:
    """Auditable table of label columns, selected drivers, and trained classifiers."""
    import numpy as np
    selected_u = {str(d).strip().upper() for d in (selected or [])}
    trained_u = None if trained is None else {str(d).strip().upper() for d in trained}
    rows = []
    min_frac = float(getattr(cfg, "min_positive_frac", 0.05)) if cfg is not None else 0.05
    for col in labels.columns:
        d = str(col).strip()
        du = d.upper()
        y = pd.to_numeric(labels[col], errors="coerce").fillna(0).astype(int)
        n = int(y.shape[0])
        n_pos = int((y == 1).sum())
        n_neg = int((y == 0).sum())
        pos_frac = float(n_pos / n) if n else float("nan")
        selected_flag = du in selected_u
        trained_flag = (du in trained_u) if trained_u is not None else None
        if not selected_flag:
            reason = "not_selected_primary_prior_or_target_list"
        elif trained_u is None:
            reason = "selected_training_not_yet_run"
        elif trained_flag:
            reason = "trained_and_scored"
        elif n_pos <= 0 or n_neg <= 0:
            reason = "skipped_one_class"
        elif pos_frac < min_frac:
            reason = f"skipped_min_positive_frac<{min_frac:g}"
        else:
            reason = "selected_but_not_trained_by_scope_classifier"
        rows.append(dict(
            stage=stage,
            cancer=cancer,
            driver=d,
            prior=prior_of(cancer, d),
            label_column_present=True,
            selected_for_training=bool(selected_flag),
            trained_classifier=("" if trained_flag is None else bool(trained_flag)),
            n_samples=n,
            n_pos=n_pos,
            n_neg=n_neg,
            positive_fraction=pos_frac,
            min_positive_frac=min_frac,
            driver_selection_mode=("all_label_drivers_exploratory" if getattr(cfg, "score_all_label_drivers", False) else "primary_prespecified"),
            reason=reason,
        ))
    # Also report locked-primary drivers missing from the label table in primary mode.
    if cfg is not None and not getattr(cfg, "score_all_label_drivers", False):
        have_u = {str(c).strip().upper() for c in labels.columns}
        for d in cancer_drivers(cancer):
            du = str(d).strip().upper()
            if du not in have_u:
                rows.append(dict(
                    stage=stage, cancer=cancer, driver=d, prior=prior_of(cancer, d),
                    label_column_present=False, selected_for_training=False,
                    trained_classifier=("" if trained_u is None else False),
                    n_samples=int(labels.shape[0]), n_pos=np.nan, n_neg=np.nan,
                    positive_fraction=np.nan, min_positive_frac=min_frac,
                    driver_selection_mode="primary_prespecified",
                    reason="primary_driver_missing_from_label_matrix",
                ))
    return pd.DataFrame(rows)


def _resolve_h5ad_path(p):
    """Prefer a repaired *_WITH_LUAD_OBS.h5ad sibling when present.

    The Lambrechts LUAD objects in older configs can have patient/celltype metadata
    stripped or barcode-like; the repair script writes a metadata-restored sibling.
    This keeps old configs working while automatically taking the repaired file once
    it exists.
    """
    if not p:
        return p
    path = Path(p)
    if path.exists():
        repaired = path.with_name(path.stem + "_WITH_LUAD_OBS" + path.suffix)
        if repaired.exists() and "LUAD" in path.name.upper() and "WITH_LUAD_OBS" not in path.name.upper():
            print(f"[singlecell-input] using repaired LUAD metadata object: {repaired}")
            return str(repaired)
        return str(path)
    # If the config already points to a repaired object that has not been made yet,
    # fall back to the original path so the run can still proceed (patient plots will
    # be skipped with a warning rather than bogus).
    marker = "_WITH_LUAD_OBS"
    if marker in path.stem:
        original = path.with_name(path.stem.replace(marker, "") + path.suffix)
        if original.exists():
            print(f"[singlecell-input] repaired metadata object not found; falling back to: {original}")
            return str(original)
    return p


def _as_nonempty_str_list(value) -> list[str]:
    """Normalize JSON scalars/lists into clean string lists for filters."""
    if value is None or value == "":
        return []
    if isinstance(value, str):
        return [value.strip()] if value.strip() else []
    return [str(x).strip() for x in value if x is not None and str(x).strip()]


def _preserve_obs_mask(adata, preserve_mask_cols=None) -> pd.Series:
    """Cells that should survive sensitivity filters, usually healthy references."""
    preserve = pd.Series(False, index=adata.obs.index)
    for col in _as_nonempty_str_list(preserve_mask_cols):
        if col in adata.obs.columns:
            preserve |= adata.obs[col].astype(bool)
    return preserve


def _filter_patients(adata, include=None, exclude=None, patient_col: str = "patient",
                     label: str = "analysis", preserve_mask_cols=None):
    """Return an AnnData copy filtered to requested patients, with diagnostics.

    For longitudinal sensitivity plots, ``preserve_mask_cols`` keeps healthy/reference
    cells even when they are not one of the selected tumour patients.  Otherwise a
    patient-only include such as ["AML328"] would also remove the BM reference ruler.
    """
    include = _as_nonempty_str_list(include)
    exclude = _as_nonempty_str_list(exclude)
    if not include and not exclude:
        return adata
    if patient_col not in adata.obs.columns:
        print(f"[filter] {label}: patient filter requested but {patient_col!r} is absent; not filtering.")
        return adata
    pat = adata.obs[patient_col].astype(str)
    analysis_mask = pd.Series(True, index=adata.obs.index)
    if include:
        analysis_mask &= pat.isin(include)
    if exclude:
        analysis_mask &= ~pat.isin(exclude)
    preserve = _preserve_obs_mask(adata, preserve_mask_cols)
    mask = analysis_mask | preserve
    kept = int(mask.sum())
    if kept == 0:
        print(f"[filter] {label}: requested patient filter would remove all cells; not filtering. "
              f"include={include or None}, exclude={exclude or None}")
        return adata
    kept_analysis = int(analysis_mask.sum())
    kept_preserve = int((preserve & ~analysis_mask).sum())
    dropped_pats = sorted(set(pat[~mask]))[:20]
    preserve_note = f"; preserved_reference_cells={kept_preserve:,}" if kept_preserve else ""
    print(f"[filter] {label}: kept {kept:,}/{adata.n_obs:,} cells "
          f"({kept_analysis:,} matched analysis cells{preserve_note}); "
          f"include={include or None}, exclude={exclude or None}; dropped examples={dropped_pats}")
    return adata[mask.values].copy()


def _filter_longitudinal_samples(adata, *, include_samples=None, exclude_samples=None,
                                 include_regex=None, exclude_regex=None,
                                 sample_col: str = "Sample", timepoint_col: str = "timepoint",
                                 preserve_mask_cols=None, label: str = "longitudinal"):
    """Filter longitudinal tumour timepoint cells by full sample/timepoint names.

    Exact filters use values from ``sample_col`` (for van Galen, e.g.
    ``AML328-D0`` or ``AML556-D31``).  Healthy/reference cells named by
    ``preserve_mask_cols`` are kept so the fixed reference distribution remains
    available for Cliff's delta, KS, and density overlays.
    """
    include_samples = _as_nonempty_str_list(include_samples)
    exclude_samples = _as_nonempty_str_list(exclude_samples)
    include_regex = str(include_regex or "").strip()
    exclude_regex = str(exclude_regex or "").strip()
    if not (include_samples or exclude_samples or include_regex or exclude_regex):
        return adata
    if sample_col not in adata.obs.columns:
        print(f"[filter] {label}: longitudinal sample filter requested but sample_col {sample_col!r} is absent; not filtering.")
        return adata

    samp = adata.obs[sample_col].astype(str)
    if timepoint_col in adata.obs.columns:
        has_tp = adata.obs[timepoint_col].notna()
    else:
        has_tp = pd.Series(True, index=adata.obs.index)
    preserve = _preserve_obs_mask(adata, preserve_mask_cols)

    # Include filters are additive/OR:
    #   longitudinal_include_samples keeps exact named samples
    #   longitudinal_include_samples_regex keeps regex-matched samples
    # This lets you keep the main longitudinal whitelist AND add one-off samples
    # like AML1012-D0 or AML210A-D0.
    if include_samples or include_regex:
        analysis_mask = pd.Series(False, index=adata.obs.index)
        if include_samples:
            analysis_mask |= samp.isin(include_samples)
        if include_regex:
            try:
                analysis_mask |= samp.str.contains(include_regex, regex=True, na=False)
            except Exception as e:
                print(f"[filter] {label}: invalid longitudinal_include_samples_regex={include_regex!r}: {e}; ignoring regex.")
    else:
        analysis_mask = pd.Series(True, index=adata.obs.index)
    if exclude_samples:
        analysis_mask &= ~samp.isin(exclude_samples)
    if exclude_regex:
        try:
            analysis_mask &= ~samp.str.contains(exclude_regex, regex=True, na=False)
        except Exception as e:
            print(f"[filter] {label}: invalid longitudinal_exclude_samples_regex={exclude_regex!r}: {e}; ignoring regex.")

    # Only sample-filter cells that belong to detected longitudinal timepoints.
    # Non-timepoint cells are irrelevant to trajectories, except preserved reference
    # cells which are intentionally retained as the comparison ruler.
    mask = (has_tp & analysis_mask) | preserve
    kept = int(mask.sum())
    kept_tp = int((has_tp & analysis_mask).sum())
    kept_ref = int((preserve & ~has_tp).sum())
    if kept_tp == 0:
        present_examples = sorted(pd.unique(samp[has_tp]).tolist())[:30]
        print(f"[filter] {label}: sample filter matched 0 longitudinal timepoint cells; not filtering. "
              f"sample_col={sample_col!r}, include_samples={include_samples or None}, "
              f"exclude_samples={exclude_samples or None}, include_regex={include_regex or None}, "
              f"exclude_regex={exclude_regex or None}; present examples={present_examples}")
        return adata
    kept_samples = sorted(pd.unique(samp[has_tp & analysis_mask]).tolist())
    dropped_samples = sorted(pd.unique(samp[has_tp & ~analysis_mask]).tolist())[:30]
    print(f"[filter] {label}: kept {kept:,}/{adata.n_obs:,} cells after sample/timepoint filter "
          f"({kept_tp:,} longitudinal cells across {len(kept_samples)} sample(s); "
          f"preserved_reference_cells={kept_ref:,}). kept_samples={kept_samples}; "
          f"dropped_sample_examples={dropped_samples}")
    return adata[mask.values].copy()


def _driver_stats_from_tables(driver: str, conf_map: dict | None = None, percell_table: pd.DataFrame | None = None) -> dict:
    """Collect per-driver annotations for figure titles/tables when available."""
    out = {}
    if conf_map and driver in conf_map:
        out["confidence"] = conf_map.get(driver)
    if (percell_table is not None and isinstance(percell_table, pd.DataFrame)
            and not percell_table.empty and "driver" in percell_table.columns):
        sub = percell_table[percell_table["driver"].astype(str) == str(driver)]
        if not sub.empty:
            r = sub.iloc[0]
            for c in ["mannwhitney_p", "fdr_q", "cliffs_delta", "apparent_auroc", "n_mut", "n_wt", "score_col", "score_kind"]:
                if c in r.index:
                    out[c] = r[c]
    return out


def _add_driver_stats_columns(df: pd.DataFrame, conf_map: dict | None = None,
                              percell_table: pd.DataFrame | None = None) -> pd.DataFrame:
    """Annotate longitudinal tables with confidence and van-Galen truth stats."""
    if df is None or df.empty or "driver" not in df.columns:
        return df
    out = df.copy()
    if conf_map:
        out["driver_confidence"] = out["driver"].map(conf_map)
    if percell_table is not None and isinstance(percell_table, pd.DataFrame) and not percell_table.empty:
        cols = [c for c in ["driver", "mannwhitney_p", "fdr_q", "cliffs_delta", "apparent_auroc", "n_mut", "n_wt"] if c in percell_table.columns]
        if "driver" in cols and len(cols) > 1:
            ann = percell_table[cols].drop_duplicates("driver").rename(columns={
                "mannwhitney_p": "ground_truth_mannwhitney_p",
                "fdr_q": "ground_truth_fdr_q",
                "cliffs_delta": "ground_truth_cliffs_delta",
                "apparent_auroc": "ground_truth_apparent_auroc",
                "n_mut": "ground_truth_n_mut_cells",
                "n_wt": "ground_truth_n_wt_cells",
            })
            out = out.merge(ann, on="driver", how="left")
    return out


def _subsample_for_memory(adata, cap: int, seed: int = 0, patient_col: str | None = None):
    """Memory safeguard: if adata has more than `cap` cells, return a seeded
    subsample (stratified by patient when available so every patient/timepoint is
    represented). cap<=0 disables. scope-bio densifies cell x gene to float64, so
    huge cohorts (e.g. 200k x 26684 = 40 GiB) OOM during scoring; this caps peak RAM
    to roughly cap x n_genes x 8 bytes."""
    import numpy as np
    if not cap or adata.n_obs <= cap:
        return adata
    rng = np.random.default_rng(seed)
    if patient_col and patient_col in adata.obs.columns:
        # proportional allocation per patient, >=1 cell each, total ~cap
        grp = adata.obs[patient_col].astype(str).values
        uniq, counts = np.unique(grp, return_counts=True)
        frac = cap / adata.n_obs
        take = []
        for g, c in zip(uniq, counts):
            idx = np.where(grp == g)[0]
            k = max(1, int(round(c * frac)))
            take.append(rng.choice(idx, size=min(k, len(idx)), replace=False))
        sel = np.sort(np.concatenate(take))
        if len(sel) > cap:
            sel = np.sort(rng.choice(sel, size=cap, replace=False))
    else:
        sel = np.sort(rng.choice(adata.n_obs, size=cap, replace=False))
    print(f"[memory] subsampled {adata.n_obs:,} -> {len(sel):,} cells "
          f"(cap={cap:,}{', stratified by '+patient_col if patient_col else ''})")
    return adata[sel].copy()


CURATED_SC_PREPROCESS_KWARGS = {
    "filter_strategy": "none",
    "already_qc_filtered": True,
    "already_normalized": True,
    "already_log_transformed": True,
    "run_doublet_detection": False,
    "auto_flag_mito": True,
}


def _singlecell_preprocess_kwargs(cfg, spec, args=None) -> dict:
    """Resolve upstream scope.SingleCellPreprocessor kwargs for this run/cohort."""
    out = dict(getattr(cfg, "singlecell_preprocessing", None) or {})
    # Per-cohort override lets one dataset use raw-count preprocessing while another
    # is treated as already curated/log-normalized.
    for key in ("singlecell_preprocessing", "singlecell_preprocess", "preprocess_kwargs"):
        if isinstance(spec.get(key), dict):
            out.update(spec[key])
    if args is not None and getattr(args, "sc_preprocess_curated", False):
        out.update(CURATED_SC_PREPROCESS_KWARGS)
    if args is not None and getattr(args, "sc_preprocess_raw", False):
        out = {}
    # Drop JSON comments and empty values.
    return {str(k): v for k, v in out.items() if not str(k).startswith("_") and v is not None}


def _score_input_path(scored_path, raw_path):
    """Choose the expression-bearing h5ad to use for rescoring.

    Prefer raw_h5ad when provided, but allow the existing scored_h5ad to serve as
    the input matrix too. This is the important escape hatch for pre-scored files:
    we strip all stale score columns before projection, then write the new scores
    back to scored_h5ad if requested.
    """
    if _exists(raw_path):
        return raw_path, "raw_h5ad"
    if _exists(scored_path):
        return scored_path, "scored_h5ad_as_raw_input"
    return None, "missing"


def _safe_project(bp, bulk_pp, raw_path, cfg, spec, args):
    """Read an expression h5ad, remove stale score columns, cap cells if requested,
    then project through the current bulk pipeline.

    This intentionally replaces the earlier implementation: force-rescore must not
    let mutation_prob_*/mutation_adj_*/mutation_resid_* columns from the input h5ad
    leak into the transformed object.
    """
    import gc
    cap = args.max_sc_cells if getattr(args, "max_sc_cells", -1) >= 0 else getattr(cfg, "max_singlecell_cells", 0)
    try:
        raw = ad.read_h5ad(raw_path)
        raw, _ = sa.singlecell.clear_scope_score_columns(
            raw, include_raw=True, include_derived=True,
            label=f"pre-rescore {Path(raw_path).name}",
        )
        raw = _subsample_for_memory(raw, cap, seed=getattr(args, "umap_random_state", 42),
                                    patient_col=spec.get("sample_col"))
        prep_kwargs = _singlecell_preprocess_kwargs(cfg, spec, args)
        if prep_kwargs:
            print(f"[singlecell-input] {Path(raw_path).name}: using configured scope single-cell preprocessing overrides: {prep_kwargs}")
        out = sa.singlecell.project_singlecell(
            bp, bulk_pp, raw, cfg,
            preprocess_kwargs=prep_kwargs,
            nan_policy=getattr(cfg, "singlecell_projection_nan_policy", "zero_fill"),
        )
        # transform() should create fresh mutation_prob_* columns.  Any derived columns
        # at this point can only be leftovers carried through obs, so drop them and let
        # the current run rebuild calibration/guardrails/confidence-adjusted scores.
        out, _ = sa.singlecell.clear_scope_score_columns(
            out, include_raw=False, include_derived=True,
            label=f"post-rescore {Path(raw_path).name}",
        )
        del raw; gc.collect()
        return out
    except MemoryError as e:
        gc.collect()
        approx = ""
        try:
            import h5py
            with h5py.File(raw_path, "r") as f:
                if "X" in f and hasattr(f["X"], "shape"):
                    approx = f" (raw X ~{f['X'].shape})"
        except Exception:
            pass
        print(f"[memory] OUT OF MEMORY scoring this cohort{approx}: {e}")
        print(f"[memory] -> skipping. Re-run with a cap, e.g.  --max-sc-cells 40000  "
              f"(or set max_singlecell_cells in the config).")
        return None


def _score_audit_rows(key, cancer, adata, *, score_source, input_h5ad=None,
                      scored_h5ad=None, write_h5ad=None, cfg=None):
    """Summarize the exact score columns used by this run.

    The goal is practical debugging: if plots do not change, this table makes it
    obvious whether the run loaded existing scores, rescored from raw, wrote back to
    disk, and which score families are present.
    """
    import numpy as np
    rows = []
    cols = sa.singlecell.scope_score_columns(adata, include_raw=True, include_derived=True)
    for c in cols:
        vals = pd.to_numeric(adata.obs[c], errors="coerce")
        finite = vals[np.isfinite(vals)]
        kind = str(c).split("_")[1] if "_" in str(c) else "score"
        driver = str(c).split("_", 2)[2] if len(str(c).split("_", 2)) >= 3 else ""
        row = dict(
            dataset=key,
            cancer=cancer,
            score_source=score_source,
            input_h5ad=str(input_h5ad or ""),
            scored_h5ad=str(scored_h5ad or ""),
            write_h5ad=str(write_h5ad or ""),
            n_cells=int(adata.n_obs),
            score_col=str(c),
            score_kind=kind,
            driver=driver,
            n_finite=int(finite.shape[0]),
            mean=float(finite.mean()) if finite.shape[0] else float("nan"),
            sd=float(finite.std()) if finite.shape[0] else float("nan"),
            min=float(finite.min()) if finite.shape[0] else float("nan"),
            q01=float(finite.quantile(0.01)) if finite.shape[0] else float("nan"),
            q50=float(finite.quantile(0.50)) if finite.shape[0] else float("nan"),
            q99=float(finite.quantile(0.99)) if finite.shape[0] else float("nan"),
            max=float(finite.max()) if finite.shape[0] else float("nan"),
        )
        if cfg is not None:
            row.update(
                use_calibrated_scores=bool(getattr(cfg, "use_calibrated_scores", False)),
                calibration_method=str(getattr(cfg, "calibration_method", "")),
                enable_reference_guardrails=bool(getattr(cfg, "enable_reference_guardrails", False)),
                longitudinal_guarded_mode=str(getattr(cfg, "longitudinal_guarded_mode", "")),
                score_umap_grey_mask=str(getattr(cfg, "score_umap_grey_mask", "")),
                malignant_call_quantile=float(getattr(cfg, "malignant_call_quantile", float("nan"))),
            )
        rows.append(row)
    if not rows:
        rows.append(dict(
            dataset=key, cancer=cancer, score_source=score_source,
            input_h5ad=str(input_h5ad or ""), scored_h5ad=str(scored_h5ad or ""),
            write_h5ad=str(write_h5ad or ""), n_cells=int(adata.n_obs),
            score_col="", note="NO mutation_* score columns present after scoring/loading",
        ))
    return rows


def _write_h5ad_atomic(adata, path, *, label="cohort"):
    """Write an AnnData safely by writing a sibling temp file then replacing."""
    if not path:
        return None
    path = Path(path)
    path.parent.mkdir(parents=True, exist_ok=True)
    tmp = path.with_name(path.name + ".tmp_rescore_write.h5ad")
    if tmp.exists():
        tmp.unlink()
    print(f"[singlecell-output] {label}: writing current run scores to {path}")
    adata.write_h5ad(tmp)
    tmp.replace(path)
    print(f"[singlecell-output] {label}: overwrote {path}")
    return path


# === HVG UMAP PATCH START ===
def _scope_hvg_umap_memory_gb() -> str:
    try:
        import os, psutil
        rss = psutil.Process(os.getpid()).memory_info().rss / (1024 ** 3)
        return f" RSS={rss:.2f} GB"
    except Exception:
        return ""


def _scope_matrix_quick_stats(X, *, max_cells: int = 5000, max_genes: int = 3000) -> dict:
    """Sparse-safe quick matrix stats for deciding UMAP-only preprocessing.

    This intentionally samples only a slice so it is safe on large h5ad files.
    """
    import numpy as np
    try:
        import scipy.sparse as sp
    except Exception:
        sp = None

    out = {
        "min": float("nan"),
        "max": float("nan"),
        "p99": float("nan"),
        "p999": float("nan"),
        "median_library": float("nan"),
        "integer_fraction": float("nan"),
        "nonzero_n": 0,
        "has_negative": False,
        "is_sparse": bool(sp is not None and sp.issparse(X)),
    }
    try:
        n_obs = int(getattr(X, "shape", (0, 0))[0])
        n_vars = int(getattr(X, "shape", (0, 0))[1])
        nr = min(max_cells, n_obs)
        nc = min(max_genes, n_vars)
        if nr <= 0 or nc <= 0:
            return out
        xs = X[:nr, :nc]
        lib = X[:nr, :].sum(axis=1)
        lib = np.asarray(lib).ravel()
        out["median_library"] = float(np.nanmedian(lib)) if lib.size else float("nan")
        if sp is not None and sp.issparse(xs):
            data = np.asarray(xs.data, dtype=float)
            out["nonzero_n"] = int(data.size)
            if data.size:
                out["min"] = float(np.nanmin(data))
                out["max"] = float(np.nanmax(data))
                out["p99"] = float(np.nanpercentile(data, 99.0))
                out["p999"] = float(np.nanpercentile(data, 99.9))
                out["has_negative"] = bool(np.nanmin(data) < -1e-8)
                out["integer_fraction"] = float(np.nanmean(np.isclose(data, np.round(data), atol=1e-8)))
            else:
                out["min"] = out["max"] = out["p99"] = out["p999"] = 0.0
                out["has_negative"] = False
                out["integer_fraction"] = 1.0
        else:
            arr = np.asarray(xs, dtype=float)
            flat = arr[np.isfinite(arr)]
            out["nonzero_n"] = int(np.count_nonzero(flat)) if flat.size else 0
            if flat.size:
                out["min"] = float(np.nanmin(flat))
                out["max"] = float(np.nanmax(flat))
                out["p99"] = float(np.nanpercentile(flat, 99.0))
                out["p999"] = float(np.nanpercentile(flat, 99.9))
                out["has_negative"] = bool(np.nanmin(flat) < -1e-8)
                nz = flat[np.abs(flat) > 1e-12]
                out["integer_fraction"] = float(np.nanmean(np.isclose(nz, np.round(nz), atol=1e-8))) if nz.size else 1.0
    except Exception:
        pass
    return out


def _scope_stats_look_count_like(stats: dict) -> bool:
    """Heuristic for raw UMI/count-like expression matrices.

    The goal is not perfect classification; it is to avoid re-log-transforming
    already log-normalized/scaled matrices while catching obvious raw counts.
    """
    import numpy as np
    if bool(stats.get("has_negative", False)):
        return False
    max_val = float(stats.get("max", np.nan))
    p999 = float(stats.get("p999", np.nan))
    p99 = float(stats.get("p99", np.nan))
    med_lib = float(stats.get("median_library", np.nan))
    int_frac = float(stats.get("integer_fraction", np.nan))

    # Obvious raw counts: large integer-ish nonzero entries.
    if np.isfinite(max_val) and max_val > 50:
        return True
    if np.isfinite(p999) and p999 > 30:
        return True
    if np.isfinite(int_frac) and int_frac > 0.98:
        if np.isfinite(p99) and p99 > 5:
            return True
        if np.isfinite(med_lib) and med_lib > 5000:
            return True
    return False


def _scope_choose_hvg_source_adata(adata, *, cohort_key: str = ""):
    """Return an AnnData-like object with gene expression for HVG UMAP.

    Prefer adata.X when it looks like expression. If scored adata.X looks already
    scaled/z-scored and adata.raw exists, use raw for HVG detection instead. This
    is visualization-only and never changes mutation scores.
    """
    try:
        if getattr(adata, "n_vars", 0) >= 500:
            stats = _scope_matrix_quick_stats(adata.X)
            if stats.get("has_negative", False):
                try:
                    if adata.raw is not None and adata.raw.shape[1] >= 500:
                        print(
                            f"[umap] {cohort_key}: adata.X contains negative/scaled values; using adata.raw for HVG UMAP source.",
                            flush=True,
                        )
                        return adata.raw.to_adata()
                except Exception:
                    pass
            return adata
    except Exception:
        pass
    try:
        if adata.raw is not None and adata.raw.shape[1] >= 500:
            print(f"[umap] {cohort_key}: using adata.raw for HVG UMAP source.", flush=True)
            return adata.raw.to_adata()
    except Exception:
        pass
    return adata


def _scope_var_rank_hvg_mask(adata_expr, n_top_genes: int):
    """Fallback HVG selector by variance, sparse-safe."""
    import numpy as np
    try:
        import scipy.sparse as sp
    except Exception:
        sp = None
    X = adata_expr.X
    if sp is not None and sp.issparse(X):
        mean = np.asarray(X.mean(axis=0)).ravel()
        mean_sq = np.asarray(X.multiply(X).mean(axis=0)).ravel()
        var = mean_sq - mean ** 2
    else:
        arr = np.asarray(X)
        var = np.nanvar(arr, axis=0)
    var = np.asarray(var, dtype=float)
    var[~np.isfinite(var)] = -np.inf
    n = min(int(n_top_genes), int(var.size))
    idx = np.argsort(var)[::-1][:n]
    mask = np.zeros(var.size, dtype=bool)
    mask[idx] = True
    return mask


def _scope_preprocess_hvg_umap_expression(work, *, cohort_key: str, preprocess: str = "auto"):
    """Normalize/log only for the fresh HVG UMAP branch.

    preprocess:
      auto            raw/count-like -> normalize_total+log1p; log-ish/scaled -> as-is
      normalize_log1p force normalize_total+log1p
      none            never normalize/log
    """
    import scanpy as sc
    preprocess = str(preprocess or "auto").lower().replace("-", "_")
    stats = _scope_matrix_quick_stats(work.X)
    looks_counts = _scope_stats_look_count_like(stats)
    summary = (
        f"min={stats.get('min', float('nan')):.3g}, max={stats.get('max', float('nan')):.3g}, "
        f"p99={stats.get('p99', float('nan')):.3g}, p999={stats.get('p999', float('nan')):.3g}, "
        f"median_lib={stats.get('median_library', float('nan')):.3g}, "
        f"integer_frac={stats.get('integer_fraction', float('nan')):.3g}"
    )
    print(f"[umap] {cohort_key}: HVG source quick stats: {summary}", flush=True)

    if preprocess in {"none", "false", "off", "no"}:
        print(f"[umap] {cohort_key}: --hvg-umap-preprocess=none; using expression as-is before HVG selection.", flush=True)
        return work, "none"
    if preprocess in {"normalize_log1p", "normalize", "log1p", "counts"}:
        print(f"[umap] {cohort_key}: forced normalize_total + log1p for HVG UMAP only.", flush=True)
        sc.pp.normalize_total(work, target_sum=1e4)
        sc.pp.log1p(work)
        return work, "normalize_log1p_forced"
    if preprocess != "auto":
        print(f"[umap] {cohort_key}: unknown --hvg-umap-preprocess={preprocess!r}; using auto.", flush=True)

    if stats.get("has_negative", False):
        print(f"[umap] {cohort_key}: expression has negative values; assuming already scaled/transformed and skipping normalize/log1p.", flush=True)
        return work, "as_is_negative_scaled"
    if looks_counts:
        print(f"[umap] {cohort_key}: expression looks count-like; running normalize_total + log1p for UMAP only.", flush=True)
        sc.pp.normalize_total(work, target_sum=1e4)
        sc.pp.log1p(work)
        return work, "normalize_log1p_auto"
    print(f"[umap] {cohort_key}: expression looks log-normalized-ish; using as-is before HVG selection.", flush=True)
    return work, "as_is_logish"


def _scope_scale_hvg_for_umap(work, *, cohort_key: str, scale: str = "auto", max_value: float = 10.0, dense_limit_gb: float = 8.0):
    """Scale HVGs when safe. Scanpy scaling may densify sparse matrices."""
    import scanpy as sc
    scale = str(scale or "auto").lower()
    try:
        max_value = float(max_value)
    except Exception:
        max_value = 10.0
    dense_gb = float(work.n_obs) * float(work.n_vars) * 4.0 / (1024.0 ** 3)

    if scale in {"none", "false", "off", "no", "never"}:
        print(f"[umap] {cohort_key}: HVG scaling disabled; sparse PCA zero_center=False.", flush=True)
        return work, False, "not_scaled_disabled"

    if scale in {"auto", "safe"} and dense_gb > dense_limit_gb:
        print(
            f"[umap] {cohort_key}: skipping HVG scaling to avoid dense memory blowup "
            f"(estimated dense HVG matrix={dense_gb:.2f} GB > {dense_limit_gb:.2f} GB); sparse PCA zero_center=False.",
            flush=True,
        )
        return work, False, "not_scaled_memory_guard"

    if scale not in {"auto", "safe", "always", "true", "yes"}:
        print(f"[umap] {cohort_key}: unknown --hvg-umap-scale={scale!r}; using auto.", flush=True)

    try:
        print(
            f"[umap] {cohort_key}: scaling HVGs for UMAP/PCA only "
            f"(max_value={max_value:g}, estimated dense={dense_gb:.2f} GB).",
            flush=True,
        )
        sc.pp.scale(work, max_value=max_value)
        return work, True, "scaled"
    except Exception as exc:
        print(
            f"[umap] {cohort_key}: HVG scaling failed ({type(exc).__name__}: {exc}); "
            "falling back to sparse PCA zero_center=False.",
            flush=True,
        )
        return work, False, "not_scaled_error"


def _scope_compute_hvg_umap_for_plotting(
    adata,
    *,
    cohort_key: str,
    n_top_genes: int = 2000,
    n_pcs: int = 50,
    n_neighbors: int = 15,
    min_dist: float = 0.5,
    random_state: int = 42,
    flavor: str = "seurat",
    batch_key: str = "",
    preprocess: str = "auto",
    scale: str = "auto",
    scale_max_value: float = 10.0,
    save_basis: bool = True,
):
    """Compute an HVG-expression UMAP and make plotting functions use it.

    Existing X_umap is preserved as X_umap_original_once. The new coordinates are
    stored as X_umap_hvg{n_top_genes} and copied into X_umap because the existing
    plotting helpers use X_umap internally.

    This branch now performs standard visualization-only preprocessing when needed:
    raw/count-like expression gets normalize_total + log1p, HVGs are selected from
    that log expression, and the HVG subset is scaled before PCA when memory-safe.
    """
    import numpy as np
    import scanpy as sc

    n_top_genes = int(n_top_genes)
    n_pcs = int(n_pcs)
    n_neighbors = int(n_neighbors)
    min_dist = float(min_dist)
    random_state = int(random_state)
    basis_key = f"X_umap_hvg{n_top_genes}"

    if not save_basis and basis_key in adata.obsm:
        print(f"[umap] {cohort_key}: using existing {basis_key}", flush=True)
        if "X_umap" in adata.obsm and "X_umap_original_once" not in adata.obsm:
            adata.obsm["X_umap_original_once"] = adata.obsm["X_umap"].copy()
        adata.obsm["X_umap"] = adata.obsm[basis_key].copy()
        return adata

    source = _scope_choose_hvg_source_adata(adata, cohort_key=cohort_key)
    if getattr(source, "n_vars", 0) < 50:
        raise ValueError(f"Too few genes/variables for HVG UMAP: n_vars={getattr(source, 'n_vars', 'NA')}")

    print(
        f"[umap] {cohort_key}: computing HVG UMAP from expression "
        f"n_cells={source.n_obs:,}, n_genes={source.n_vars:,}, top_hvg={n_top_genes}, "
        f"pcs={n_pcs}, neighbors={n_neighbors}, min_dist={min_dist}, "
        f"preprocess={preprocess}, scale={scale}" + _scope_hvg_umap_memory_gb(),
        flush=True,
    )

    # Copy only metadata + matrix needed for visualization. This avoids mutating scores.
    work = source.copy()
    work, preprocess_mode = _scope_preprocess_hvg_umap_expression(work, cohort_key=cohort_key, preprocess=preprocess)

    hv_kwargs = dict(n_top_genes=min(n_top_genes, int(work.n_vars)), flavor=flavor)
    if batch_key and batch_key in work.obs.columns:
        hv_kwargs["batch_key"] = batch_key
        print(f"[umap] {cohort_key}: using batch-aware HVG selection with batch_key={batch_key!r}.", flush=True)
    elif batch_key:
        print(f"[umap] {cohort_key}: requested batch_key={batch_key!r} not found in obs; HVG selection is not batch-aware.", flush=True)
    try:
        sc.pp.highly_variable_genes(work, **hv_kwargs)
        hvg_mask = np.asarray(work.var.get("highly_variable", False), dtype=bool)
        if hvg_mask.sum() < 10:
            raise ValueError(f"HVG selection returned only {int(hvg_mask.sum())} genes")
    except Exception as exc:
        print(f"[umap] {cohort_key}: scanpy HVG failed ({type(exc).__name__}: {exc}); using variance-ranked genes", flush=True)
        hvg_mask = _scope_var_rank_hvg_mask(work, n_top_genes)
        work.var["highly_variable"] = hvg_mask

    work = work[:, np.asarray(work.var["highly_variable"], dtype=bool)].copy()
    print(f"[umap] {cohort_key}: HVG subset = {work.n_vars:,} genes" + _scope_hvg_umap_memory_gb(), flush=True)

    work, did_scale, scale_mode = _scope_scale_hvg_for_umap(
        work,
        cohort_key=cohort_key,
        scale=scale,
        max_value=scale_max_value,
    )

    pcs_eff = max(2, min(n_pcs, int(work.n_obs) - 1, int(work.n_vars) - 1))
    # zero_center=True only after scaling; otherwise keep sparse matrices sparse.
    sc.tl.pca(work, n_comps=pcs_eff, svd_solver="arpack", zero_center=bool(did_scale), random_state=random_state)
    print(f"[umap] {cohort_key}: PCA done" + _scope_hvg_umap_memory_gb(), flush=True)

    sc.pp.neighbors(work, n_neighbors=n_neighbors, n_pcs=pcs_eff, random_state=random_state)
    print(f"[umap] {cohort_key}: neighbors done" + _scope_hvg_umap_memory_gb(), flush=True)

    sc.tl.umap(work, min_dist=min_dist, random_state=random_state)
    coords = np.asarray(work.obsm["X_umap"], dtype=np.float32)
    if coords.shape[0] != adata.n_obs:
        raise ValueError(f"UMAP row mismatch: got {coords.shape[0]}, expected {adata.n_obs}")

    if "X_umap" in adata.obsm and "X_umap_original_once" not in adata.obsm:
        adata.obsm["X_umap_original_once"] = adata.obsm["X_umap"].copy()
    adata.obsm[basis_key] = coords
    adata.obsm["X_umap"] = coords.copy()
    adata.uns["scope_umap_source"] = {
        "mode": "hvg",
        "basis_key": basis_key,
        "preprocess_mode": str(preprocess_mode),
        "scale_mode": str(scale_mode),
        "n_top_genes": int(n_top_genes),
        "n_pcs": int(pcs_eff),
        "n_neighbors": int(n_neighbors),
        "min_dist": float(min_dist),
        "random_state": int(random_state),
        "flavor": str(flavor),
        "batch_key": str(batch_key),
    }
    print(f"[umap] {cohort_key}: wrote {basis_key} and replaced X_umap for plotting" + _scope_hvg_umap_memory_gb(), flush=True)
    return adata


def _scope_compute_svd_umap_for_plotting(
    adata,
    *,
    cohort_key: str,
    n_neighbors: int = 15,
    min_dist: float = 0.5,
    random_state: int = 42,
):
    """Explicitly recompute UMAP from X_svd and use it for plotting."""
    import numpy as np
    import scanpy as sc
    import anndata as ad

    if "X_svd" not in adata.obsm:
        raise KeyError("adata.obsm['X_svd'] not found")
    Z = np.asarray(adata.obsm["X_svd"], dtype=np.float32)
    work = ad.AnnData(Z)
    sc.pp.neighbors(work, use_rep="X", n_neighbors=int(n_neighbors), random_state=int(random_state))
    sc.tl.umap(work, min_dist=float(min_dist), random_state=int(random_state))
    coords = np.asarray(work.obsm["X_umap"], dtype=np.float32)
    if "X_umap" in adata.obsm and "X_umap_original_once" not in adata.obsm:
        adata.obsm["X_umap_original_once"] = adata.obsm["X_umap"].copy()
    adata.obsm["X_umap_svd_recomputed"] = coords
    adata.obsm["X_umap"] = coords.copy()
    adata.uns["scope_umap_source"] = {
        "mode": "svd",
        "basis_key": "X_umap_svd_recomputed",
        "n_neighbors": int(n_neighbors),
        "min_dist": float(min_dist),
        "random_state": int(random_state),
    }
    print(f"[umap] {cohort_key}: recomputed SVD UMAP and replaced X_umap for plotting", flush=True)
    return adata


def _scope_prepare_umap_for_plotting(adata, *, cohort_key: str, spec: dict, args):
    """Select the UMAP coordinate source before calling existing plotting helpers."""
    mode = getattr(args, "umap_source", "cached")
    if mode in {None, "", "cached"}:
        return adata

    # Allow per-cohort JSON overrides without requiring separate commands.
    n_top_genes = int(spec.get("hvg_umap_top_genes", getattr(args, "hvg_umap_top_genes", 2000)))
    n_pcs = int(spec.get("hvg_umap_n_pcs", getattr(args, "hvg_umap_n_pcs", 50)))
    n_neighbors = int(spec.get("hvg_umap_n_neighbors", getattr(args, "hvg_umap_n_neighbors", 15)))
    min_dist = float(spec.get("hvg_umap_min_dist", getattr(args, "hvg_umap_min_dist", 0.5)))
    random_state = int(getattr(args, "umap_random_state", 42))
    flavor = str(getattr(args, "hvg_umap_flavor", "seurat"))
    batch_key = str(spec.get("hvg_umap_batch_key", getattr(args, "hvg_umap_batch_key", "")) or "")
    preprocess = str(spec.get("hvg_umap_preprocess", getattr(args, "hvg_umap_preprocess", "auto")) or "auto")
    scale = str(spec.get("hvg_umap_scale", getattr(args, "hvg_umap_scale", "auto")) or "auto")
    scale_max_value = float(spec.get("hvg_umap_scale_max_value", getattr(args, "hvg_umap_scale_max_value", 10.0)))

    try:
        if mode == "hvg":
            return _scope_compute_hvg_umap_for_plotting(
                adata,
                cohort_key=cohort_key,
                n_top_genes=n_top_genes,
                n_pcs=n_pcs,
                n_neighbors=n_neighbors,
                min_dist=min_dist,
                random_state=random_state,
                flavor=flavor,
                batch_key=batch_key,
                preprocess=preprocess,
                scale=scale,
                scale_max_value=scale_max_value,
                save_basis=True,
            )
        if mode == "svd":
            return _scope_compute_svd_umap_for_plotting(
                adata,
                cohort_key=cohort_key,
                n_neighbors=n_neighbors,
                min_dist=min_dist,
                random_state=random_state,
            )
        if mode == "auto":
            if "X_umap" in adata.obsm:
                print(f"[umap] {cohort_key}: using cached X_umap", flush=True)
                return adata
            if "X_svd" in adata.obsm:
                return _scope_compute_svd_umap_for_plotting(
                    adata,
                    cohort_key=cohort_key,
                    n_neighbors=n_neighbors,
                    min_dist=min_dist,
                    random_state=random_state,
                )
            return _scope_compute_hvg_umap_for_plotting(
                adata,
                cohort_key=cohort_key,
                n_top_genes=n_top_genes,
                n_pcs=n_pcs,
                n_neighbors=n_neighbors,
                min_dist=min_dist,
                random_state=random_state,
                flavor=flavor,
                batch_key=batch_key,
                preprocess=preprocess,
                scale=scale,
                scale_max_value=scale_max_value,
                save_basis=True,
            )
    except Exception as exc:
        print(f"[umap] {cohort_key}: requested UMAP source {mode!r} failed: {type(exc).__name__}: {exc}", flush=True)
        print(f"[umap] {cohort_key}: falling back to existing plotting coordinates", flush=True)
        return adata

    print(f"[umap] {cohort_key}: unknown --umap-source={mode!r}; using cached", flush=True)
    return adata
# === HVG UMAP PATCH END ===

def main():
    import shutil

    ap = argparse.ArgumentParser()
    ap.add_argument("--config", required=True)
    ap.add_argument("--only", default="", help="comma list of stages to run")
    ap.add_argument("--skip", default="", help="comma list of stages to skip")
    ap.add_argument("--clean-interpret", action="store_true", help="delete existing Fig2 per-driver SVD eval folders before re-writing them")
    ap.add_argument("--umap-source", default="cached", choices=("cached", "hvg", "svd", "auto"), help="Coordinate source for Fig4 UMAP panels: cached X_umap, fresh HVG expression UMAP, fresh X_svd UMAP, or auto fallback.")
    ap.add_argument("--hvg-umap-top-genes", type=int, default=2000, help="Number of highly variable genes for --umap-source hvg.")
    ap.add_argument("--hvg-umap-n-pcs", type=int, default=50, help="Number of PCs for --umap-source hvg.")
    ap.add_argument("--hvg-umap-n-neighbors", type=int, default=15, help="Neighbors for fresh UMAP computation.")
    ap.add_argument("--hvg-umap-min-dist", type=float, default=0.5, help="UMAP min_dist for fresh UMAP computation.")
    ap.add_argument("--hvg-umap-flavor", default="seurat", help="scanpy.pp.highly_variable_genes flavor for HVG UMAP; seurat is robust without extra deps.")
    ap.add_argument("--hvg-umap-batch-key", default="", help="Optional obs column for batch-aware HVG selection.")
    ap.add_argument("--hvg-umap-preprocess", default="auto", choices=("auto", "none", "normalize_log1p"), help="For --umap-source hvg: auto-detect count-like expression and run normalize_total+log1p, never preprocess, or force normalize_total+log1p.")
    ap.add_argument("--hvg-umap-scale", default="auto", choices=("auto", "never", "always"), help="For --umap-source hvg: scale HVGs before PCA when memory-safe, never scale, or force scaling.")
    ap.add_argument("--hvg-umap-scale-max-value", type=float, default=10.0, help="Max value clipping used by scanpy.pp.scale for --umap-source hvg.")
    ap.add_argument("--umap-random-state", type=int, default=42, help="Random seed for fresh UMAP computation.")
    ap.add_argument(
        "--force-rescore-singlecell",
        action="store_true",
        help=(
            "Ignore existing scored_h5ad files and re-project raw_h5ad through the current "
            "bulk pipeline. Use this when code/model settings changed and old data/models/* "
            "scored objects should not be reused."
        ),
    )
    ap.add_argument(
        "--overwrite-scored-h5ad",
        action="store_true",
        help=(
            "With --force-rescore-singlecell, write the current run's freshly recomputed "
            "scores/derived score columns back to the cohort's scored_h5ad path, replacing "
            "that same .h5ad atomically. If raw_h5ad is missing, the existing scored_h5ad "
            "is used as the expression input after stripping stale score columns."
        ),
    )
    ap.add_argument(
        "--clean-output-root",
        action="store_true",
        help=(
            "Delete output_root before running. Useful when changing score settings so old "
            "PNGs/CSVs from prior runs cannot be mistaken for current-run outputs."
        ),
    )
    ap.add_argument(
        "--warn-stale-scored",
        action="store_true",
        help="Print an extra warning when a single-cell cohort is loaded from data/models/* scored_h5ad.",
    )
    ap.add_argument(
        "--max-sc-cells", type=int, default=-1,
        help=("Memory safeguard: cap the number of single cells scored per cohort "
              "(subsample, seeded, stratified by patient if available). scope-bio "
              "densifies the cell x gene matrix to float64, so very large cohorts "
              "(e.g. BRCA 200k x 26684 = 40 GiB) OOM. -1 uses cfg.max_singlecell_cells; "
              "0 means no cap."),
    )
    ap.add_argument(
        "--sc-preprocess-curated", action="store_true",
        help=("Treat scRNA h5ads as already QC-filtered/normalized/log-transformed during "
              "force-rescoring: disables scope-bio's internal QC/doublet filters. "
              "Equivalent to singlecell_preprocessing curated settings in the JSON."),
    )
    ap.add_argument(
        "--sc-preprocess-raw", action="store_true",
        help=("Ignore JSON singlecell_preprocessing overrides and let upstream scope-bio use "
              "its raw-count preprocessing defaults."),
    )
    ap.add_argument(
        "--score-all-label-drivers", action="store_true",
        help=("Exploratory mode: train/score every binary mutation column present in each "
              "bulk labels file, instead of only the pre-specified DRIVER_PRIORS list. "
              "Unlisted genes are marked prior='unspecified' in audit tables."),
    )
    ap.add_argument(
        "--target-drivers", default="",
        help=("Comma-separated explicit driver list to train/score for every cancer where "
              "the labels exist, e.g. NPM1,DNMT3A,TP53. Mutually exclusive with "
              "--score-all-label-drivers."),
    )
    ap.add_argument(
        "--min-positive-frac", type=float, default=None,
        help=("Override scope-bio classifier min_positive_frac for this run. Useful with "
              "--score-all-label-drivers; e.g. 0.01 scores rarer exploratory mutations. "
              "Genes with too few positives/negatives can still be skipped by the classifier."),
    )
    ap.add_argument(
        "--plot-driver-set", default=None,
        choices=("primary_if_all_labels", "primary", "all", "confidence"),
        help=("Which scored drivers to show in crowded Fig4 UMAP/heatmap/triptych panels. "
              "Default keeps primary drivers when --score-all-label-drivers is on; use 'all' "
              "for full exploratory atlases."),
    )
    ap.add_argument("--confidence-display-min", type=float, default=None,
                    help="Minimum ground-truth-free confidence for confidence-filtered heatmaps/plots.")
    ap.add_argument("--plot-residual-min-abs-vmax", type=float, default=None,
                    help="Plot-only minimum +/- color limit for residual UMAPs; prevents tiny noise ranges from looking bright.")
    ap.add_argument("--plot-residual-abs-vmax", type=float, default=None,
                    help="Plot-only maximum +/- color limit for residual UMAPs; high values are clipped for readability.")
    ap.add_argument("--heatmap-score-scale", default=None,
                    choices=("global_robust", "driver_robust", "absolute"),
                    help="Celltype heatmap color scaling. global_robust is comparable across drivers; driver_robust is exploratory.")
    ap.add_argument("--interpret-top-components", type=int, default=None,
                    help="Number of top LR-important SVD components to export in gene_program_table/component_signature_scan.")
    ap.add_argument("--interpret-genes-per-component", type=int, default=None,
                    help="Number of top absolute-loading genes per component to export in gene_program_table/component_signature_scan.")
    ap.add_argument("--interpret-plot-top-components", type=int, default=None,
                    help="Number of components shown in top_genes_per_component.png.")
    ap.add_argument("--interpret-plot-genes-per-component", type=int, default=None,
                    help="Number of genes shown per component in top_genes_per_component.png.")
    ap.add_argument("--interpret-heatmap-top-components", type=int, default=None,
                    help="Number of components included in gene_loading_heatmap.png.")
    ap.add_argument("--interpret-heatmap-genes-per-component", type=int, default=None,
                    help="Number of top genes per component contributing to gene_loading_heatmap.png.")
    args = ap.parse_args()
    if args.sc_preprocess_curated and args.sc_preprocess_raw:
        ap.error("--sc-preprocess-curated and --sc-preprocess-raw are mutually exclusive")
    if args.score_all_label_drivers and str(args.target_drivers).strip():
        ap.error("--score-all-label-drivers and --target-drivers are mutually exclusive")
    if args.overwrite_scored_h5ad and not args.force_rescore_singlecell:
        print("[run_all] --overwrite-scored-h5ad implies --force-rescore-singlecell; enabling force rescore.")
        args.force_rescore_singlecell = True
    cfg, raw = load_config(args.config)
    if args.score_all_label_drivers:
        cfg.score_all_label_drivers = True
    if str(args.target_drivers).strip():
        cfg.target_drivers = _parse_driver_list(args.target_drivers)
    if args.min_positive_frac is not None:
        if args.min_positive_frac < 0 or args.min_positive_frac >= 1:
            ap.error("--min-positive-frac must be >=0 and <1")
        cfg.min_positive_frac = float(args.min_positive_frac)
    if args.plot_driver_set is not None:
        cfg.plot_driver_set = args.plot_driver_set
    if args.confidence_display_min is not None:
        cfg.confidence_display_min = float(args.confidence_display_min)
    if args.plot_residual_min_abs_vmax is not None:
        cfg.plot_residual_min_abs_vmax = float(args.plot_residual_min_abs_vmax)
    if args.plot_residual_abs_vmax is not None:
        cfg.plot_residual_abs_vmax = float(args.plot_residual_abs_vmax)
    if args.heatmap_score_scale is not None:
        cfg.heatmap_score_scale = args.heatmap_score_scale
    for _name in [
        "interpret_top_components",
        "interpret_genes_per_component",
        "interpret_plot_top_components",
        "interpret_plot_genes_per_component",
        "interpret_heatmap_top_components",
        "interpret_heatmap_genes_per_component",
    ]:
        _val = getattr(args, _name, None)
        if _val is not None:
            if int(_val) < 1:
                ap.error(f"--{_name.replace('_', '-')} must be >= 1")
            setattr(cfg, _name, int(_val))
    if args.clean_output_root and cfg.output_root.exists():
        print(f"[run_all] --clean-output-root: removing {cfg.output_root}")
        shutil.rmtree(cfg.output_root, ignore_errors=True)
    driver_mode = "all label columns (exploratory)" if getattr(cfg, "score_all_label_drivers", False) else ("configured target_drivers" if getattr(cfg, "target_drivers", "all") != "all" else "pre-specified DRIVER_PRIORS")
    print(f"[run_all] config: n_components={cfg.n_components}, interpret_top_components={getattr(cfg, 'interpret_top_components', None)}, interpret_genes_per_component={getattr(cfg, 'interpret_genes_per_component', None)}, output_root={cfg.output_root}")
    print(f"[run_all] driver_selection={driver_mode}; min_positive_frac={cfg.min_positive_frac:g}")
    only = {s for s in args.only.split(",") if s}
    skip = {s for s in args.skip.split(",") if s}

    def run(stage): return (not only or stage in only) and stage not in skip

    M = sa.report.Manifest(cfg)
    fitted: dict[str, object] = {}          # cancer -> fitted BulkPipeline
    bulk_pp: dict[str, object] = {}         # cancer -> preprocessed bulk
    bulk_table = pd.DataFrame()

    # ---------- bulk cohort loader bound to the config ----------
    def bulk_loader(cancer):
        spec = raw["bulk_cohorts"][cancer]
        if not (_exists(spec["expression"]) and _exists(spec["mutations"])):
            raise FileNotFoundError(f"{cancer}: bulk files missing")
        return sa.data.load_bulk_cohort(spec["expression"], spec["mutations"],
                                        sample_col=spec.get("sample_col", "sample_id"))

    # ================= 1. BULK =================
    if run("bulk"):
        print("\n########## STAGE 1: BULK ##########")
        bulk_table = sa.bulk.evaluate_all_cancers_bulk(bulk_loader, cfg)
        if not bulk_table.empty:
            M.save_table("Table1_bulk_discrimination", bulk_table)
            M.register_figure("Fig2_Table1_bulk",
                              sa.plots.forest_bulk_auroc(bulk_table,
                                  cfg.figdir("fig2_bulk") / "Fig2_forest_bulk_auroc.png"))
            for cancer in bulk_table.cancer.unique():
                drivers = bulk_table[bulk_table.cancer == cancer].driver.tolist()[:6]
                M.register_figure("Fig2_Table1_bulk",
                                  sa.plots.roc_overlay(bulk_table.attrs["oof_store"],
                                      cancer, drivers, cfg.figdir("fig2_bulk") / f"Fig2_roc_{cancer}.png"))

    # ============ FIT pipelines for downstream stages ============
    if run("interpret") or run("singlecell") or run("longitudinal") or \
       run("pancancer") or run("cnv") or run("robustness"):
        for cancer in cfg.cancers():
            try:
                a_bulk, labels = bulk_loader(cancer)
            except FileNotFoundError as e:
                print(f"[run_all] {e}; downstream stages will skip {cancer}.")
                continue
            present = _selected_drivers_for_labels(cancer, labels, cfg)
            labels_selected = labels[present]
            if not present:
                print(f"[run_all] {cancer}: no selected mutation labels available; downstream stages will skip this cancer.")
                continue
            print(f"[run_all] {cancer}: selected {len(present)}/{labels.shape[1]} label column(s) for training/scoring "
                  f"({ 'all-label exploratory' if getattr(cfg, 'score_all_label_drivers', False) else 'primary/targeted' }).")
            bp = sa.bulk.fit_bulk_pipeline(a_bulk, labels_selected, cfg)
            trained = list(getattr(bp.classifier_set_, "classifiers_", {}).keys())
            M.save_table(f"Driver_training_audit_{cancer}",
                         _driver_selection_audit(cancer, labels, present, trained=trained, cfg=cfg, stage="downstream_fit"))
            if len(trained) != len(present):
                skipped = [d for d in present if str(d).upper() not in {str(t).upper() for t in trained}]
                print(f"[run_all] {cancer}: trained/scored {len(trained)}/{len(present)} selected drivers; "
                      f"skipped examples={skipped[:12]}")
            fitted[cancer] = (bp, a_bulk, labels_selected)
            bulk_pp[cancer] = bp.preprocessor_.transform(a_bulk)

    # ================= 2. INTERPRET =================
    if run("interpret"):
        print("\n########## STAGE 2: INTERPRET ##########")
        if args.clean_interpret:
            import shutil
            fig2 = cfg.figdir("fig2_bulk")
            for old in fig2.glob("*/svd_eval_*"):
                print(f"[run_all] clean-interpret: removing {old}")
                shutil.rmtree(old, ignore_errors=True)
        for cancer, (bp, a_bulk, labels) in fitted.items():
            drivers = list(bp.classifier_set_.classifiers_.keys())
            sa.interpret.interpret_drivers(bp, a_bulk, drivers, cfg, cancer, full_suite=True, labels=labels)
            # deeper signature mining across many top components (for gene-signature scanning)
            sa.interpret.component_signature_scan(
                bp, a_bulk, drivers, cfg, cancer,
                top_components=getattr(cfg, "interpret_top_components", 25),
                genes_per_component=getattr(cfg, "interpret_genes_per_component", 50))
            for d in drivers:
                lead = sa.interpret.leading_gene_programs(
                    bp, a_bulk, d, top_components=3,
                    genes_per_component=getattr(cfg, "interpret_genes_per_component", 50))
                M.save_table(f"genes_{cancer}_{d}", lead)

    # ================= 3-6. SINGLE-CELL STAGES (per cohort) =================
    for key, spec in raw.get("singlecell_cohorts", {}).items():
        cancer = spec["cancer"]
        conf_table = pd.DataFrame()
        conf_map = {}
        percell_stats_for_longitudinal = pd.DataFrame()
        malig_col = None
        guard_ref_col = "is_reference"
        # obtain a single-cell object. Default keeps prior behavior (use scored_h5ad
        # if present), but --force-rescore-singlecell makes this deterministic after
        # code/model updates by ignoring stale scored objects and re-projecting raw_h5ad.
        adata = None
        scored_path = _resolve_h5ad_path(spec.get("scored_h5ad"))
        raw_path = _resolve_h5ad_path(spec.get("raw_h5ad"))
        rescore_input_path, rescore_input_kind = _score_input_path(scored_path, raw_path)
        can_rescore = _exists(rescore_input_path) and cancer in fitted
        score_source = "not_loaded"
        score_input_used = ""
        h5ad_write_path = None

        if args.force_rescore_singlecell and can_rescore:
            print(f"[singlecell-input] {key}: FORCE-RESCORE from {rescore_input_kind}: {rescore_input_path}")
            bp, _, _ = fitted[cancer]
            adata = _safe_project(bp, bulk_pp[cancer], rescore_input_path, cfg, spec, args)
            score_source = f"rescored_from_{rescore_input_kind}"
            score_input_used = str(rescore_input_path)
        elif args.force_rescore_singlecell and not can_rescore:
            print(f"[singlecell-input] WARNING {key}: --force-rescore-singlecell requested, "
                  f"but neither raw_h5ad nor scored_h5ad can be used for rescoring, or {cancer} "
                  "has no fitted bulk pipeline. Falling back to scored_h5ad only if present.")
            if _exists(scored_path):
                print(f"[singlecell-input] {key}: using EXISTING scored_h5ad fallback: {scored_path}")
                adata = sa.data.load_scored_singlecell(scored_path)
                score_source = "loaded_existing_scored_h5ad_after_failed_rescore"
                score_input_used = str(scored_path)
        elif _exists(scored_path):
            print(f"[singlecell-input] {key}: using EXISTING scored_h5ad: {scored_path}")
            if args.warn_stale_scored and "data/models" in str(scored_path).replace("\\", "/"):
                print(f"[singlecell-input] WARNING {key}: scored_h5ad is under data/models; "
                      "this may be an older precomputed object. Use --force-rescore-singlecell "
                      "or point scored_h5ad to the new derived file if you changed scoring/model settings.")
            adata = sa.data.load_scored_singlecell(scored_path)
            score_source = "loaded_existing_scored_h5ad"
            score_input_used = str(scored_path)
        elif can_rescore:
            print(f"[singlecell-input] {key}: no scored_h5ad found; projecting {rescore_input_kind}: {rescore_input_path}")
            bp, _, _ = fitted[cancer]
            adata = _safe_project(bp, bulk_pp[cancer], rescore_input_path, cfg, spec, args)
            score_source = f"rescored_from_{rescore_input_kind}"
            score_input_used = str(rescore_input_path)

        if adata is None:
            print(f"[run_all] cohort {key}: no usable scored/raw h5ad (+fitted bulk); skipping.")
            continue

        # At the start of every run, wipe run-derived score columns so plots/tables
        # cannot use stale calibration, confidence-adjusted, residual, or percentile
        # scores from a previous parameter setting. Raw mutation_prob_* is preserved
        # unless this was a forced rescore, where _safe_project already rebuilt it.
        adata, _dropped_derived_scores = sa.singlecell.clear_scope_score_columns(
            adata, include_raw=False, include_derived=True, label=f"{key} run-derived reset")

        # standard metadata derivation
        # NEW: drop contaminating samples (e.g. sorted-progenitor BM5* controls and
        # cell lines) before anything else, so they can't seed false positives or
        # distort concordance / calibration downstream.
        if spec.get("sample_col") and spec["sample_col"] in adata.obs.columns:
            adata = sa.data.filter_samples(
                adata, sample_col=spec["sample_col"],
                exclude_samples=spec.get("exclude_samples"),
                exclude_regex=spec.get("exclude_samples_regex"),
                keep_samples=spec.get("keep_samples"))
        # Robust patient/sample metadata. Works for van Galen timepoints, solid
        # tumour patient-level cohorts, and new datasets with common obs names.
        adata = sa.data.ensure_patient_metadata(
            adata, sample_col=spec.get("sample_col") or None,
            patient_strip_regex=spec.get("patient_strip_regex", r"-D\d+$"),
            timepoint_regex=spec.get("timepoint_regex", r"-(D\d+)$"),
        )
        has_patient = ("patient" in adata.obs.columns and adata.obs["patient"].notna().any()
                       and sa.data.is_usable_patient_series(adata.obs["patient"], colname="patient"))
        celltype_col = spec.get("celltype_col", "CellType")
        # Weak-celltyping guardrail: if the requested annotation is missing or
        # uninformative, create a coarse marker-based label before references are
        # built. This keeps weakly celltyped data from silently becoming all
        # non-reference or lineage-mismatched.
        try:
            adata, celltype_col = sa.annotate.ensure_celltype(adata, celltype_col)
        except Exception as e:
            print(f"[annotate] marker-based fallback skipped for {key}: {e}")
        # Always create a coarse display/state label. For PAAD and other public
        # cohorts with very granular/janky labels this gives readable heatmaps,
        # while preserving the original annotation in obs.
        try:
            adata, _coarse_col = sa.reference.ensure_coarse_cellstate(adata, celltype_col=celltype_col, out_col="celltype_coarse")
            if spec.get("use_coarse_celltype", key.upper().startswith("PAAD") or cancer == "PAAD"):
                celltype_col = _coarse_col
                print(f"[annotate] {key}: using coarse celltype labels for heatmaps/context ({celltype_col}).")
        except Exception as e:
            print(f"[annotate] coarse celltype labels skipped for {key}: {e}")
        adata = sa.data.mark_reference_cells(
            adata, celltype_col=celltype_col,
            reference_celltypes=tuple(spec.get("reference_celltypes", ("T", "B", "NK"))),
            sample_col=spec.get("sample_col", "Sample"),
            reference_samples=spec.get("reference_samples"))
        if "is_reference" in adata.obs.columns:
            ref_include = [str(x) for x in spec.get("reference_include_patients", []) if str(x).strip()]
            ref_exclude = [str(x) for x in spec.get("reference_exclude_patients", []) if str(x).strip()]
            if (ref_include or ref_exclude) and "patient" in adata.obs.columns:
                pat = adata.obs["patient"].astype(str)
                ref_mask = adata.obs["is_reference"].astype(bool).copy()
                before_ref = int(ref_mask.sum())
                if ref_include:
                    ref_mask &= pat.isin(ref_include)
                if ref_exclude:
                    ref_mask &= ~pat.isin(ref_exclude)
                if int(ref_mask.sum()) == 0:
                    print(f"[data] WARNING {key}: reference patient filter removed all reference cells; "
                          f"keeping original reference. include={ref_include or None}, exclude={ref_exclude or None}")
                else:
                    adata.obs["is_reference"] = ref_mask.values
                    print(f"[data] {key}: reference patient filter {before_ref:,} -> {int(ref_mask.sum()):,} cells; "
                          f"include={ref_include or None}, exclude={ref_exclude or None}")
            elif (ref_include or ref_exclude):
                print(f"[data] WARNING {key}: reference patient filter requested but 'patient' is absent; not filtering reference.")

        if celltype_col in adata.obs.columns and "is_reference" in adata.obs.columns:
            ref_comp = (adata.obs.groupby([celltype_col, "is_reference"], observed=False)
                        .size().reset_index(name="n_cells")
                        .sort_values(["is_reference", "n_cells"], ascending=[False, False]))
            M.save_table(f"Reference_composition_{key}", ref_comp)
            if has_patient:
                ref_patient_comp = (adata.obs.loc[adata.obs["is_reference"].astype(bool)]
                                    .groupby("patient", observed=False).size()
                                    .reset_index(name="n_reference_cells")
                                    .sort_values("n_reference_cells", ascending=False))
                M.save_table(f"Reference_patient_composition_{key}", ref_patient_comp)
            ref_total = int(adata.obs["is_reference"].sum())
            top_ref = (ref_comp[ref_comp["is_reference"]]
                       .head(8)[[celltype_col, "n_cells"]].to_dict("records"))
            print(f"[data] {key}: reference composition saved; {ref_total:,} reference cells; top={top_ref}")
        status = (sa.data.load_patient_status(spec["patient_status_csv"])
                  if _exists(spec.get("patient_status_csv")) else None)
        if status is not None and has_patient:
            status = sa.data.harmonize_patient_status(status, adata.obs["patient"].dropna().astype(str).values)
            status = sa.data.validate_patient_status_for_obs(status, adata.obs["patient"].dropna().astype(str).values, cohort_name=key)
        elif status is not None:
            print(f"[data] WARNING {key}: patient_status present but no usable scRNA patient column; patient-level overlays/bridges skipped.")
            status = None
        drivers = sa.data.scored_drivers(adata)
        # van Galen stores per-cell expressed-mutation truth in generic
        # MutTranscript/WtTranscript string columns. Parse those into per-driver
        # <GENE>_genotype columns before Fig3 validation and Fig4 triptychs,
        # otherwise AML falls back to patient-level mutation status.
        try:
            adata, mt_parse = sa.singlecell.add_muttranscript_genotype_columns(
                adata, drivers=drivers, overwrite=True)
            if mt_parse is not None and not mt_parse.empty:
                M.save_table(f"MutTranscript_genotype_parse_{key}", mt_parse)
        except Exception as e:
            print(f"[singlecell] {key}: MutTranscript/WtTranscript genotype parse skipped ({e})")

        # ---- Reference-set sensitivity guardrails (NPM1 / progenitor-program issue).
        # Adds multiple healthy reference masks plus a coarse cell-state column.
        guard_ref_col = "is_reference"
        if getattr(cfg, "enable_reference_guardrails", True):
            try:
                guard_samples = spec.get("reference_guardrail_samples", list(getattr(cfg, "reference_guardrail_samples", ())))
                if not guard_samples:
                    guard_samples = spec.get("reference_samples")
                adata, refset_tbl, refset_cols = sa.reference.add_reference_sets(
                    adata, celltype_col=celltype_col, sample_col=spec.get("sample_col", "Sample"),
                    reference_samples=guard_samples)
                if not refset_tbl.empty:
                    M.save_table(f"Reference_sensitivity_sets_{key}", refset_tbl)
                preferred = "is_reference_all_healthy_BM1_BM4"
                guard_ref_col = preferred if preferred in adata.obs.columns and adata.obs[preferred].astype(bool).sum() > 0 else "is_reference"
            except Exception as e:
                print(f"[reference] guardrail reference sets skipped for {key}: {e}")

        # Score-UMAP greying mask. v9.1 auto-switched AML to is_normal_display, which
        # greyed every non-"-like" annotated cell and hid too many cell types in the
        # prediction UMAPs. Keep building is_normal_display for diagnostics, but use
        # the explicit config knob below for plotting. Default = true reference cells only.
        grey_col = "is_reference"
        try:
            mal_regex = spec.get("malignant_label_regex")
            if not mal_regex:
                # auto-detect van Galen "-like" malignant annotation
                ctv = adata.obs[celltype_col].astype(str) if celltype_col in adata.obs.columns else None
                if ctv is not None and ctv.str.contains(r"-like$", case=False, regex=True).any():
                    mal_regex = r"-like$"
            adata, _ndinfo = sa.reference.add_normal_display_mask(
                adata, celltype_col=celltype_col, malignant_label_regex=mal_regex)
        except Exception as e:
            print(f"[reference] normal-display mask skipped for {key}: {e}")

        grey_mode = str(getattr(cfg, "score_umap_grey_mask", "reference") or "reference").strip().lower()
        if grey_mode in {"normal_display", "normal", "all_normal", "context", "all_context"}:
            if "is_normal_display" in adata.obs.columns and adata.obs["is_normal_display"].astype(bool).any():
                grey_col = "is_normal_display"
            else:
                print(f"[reference] {key}: score_umap_grey_mask={grey_mode!r} requested, but "
                      "is_normal_display is unavailable/empty; falling back to is_reference.")
                grey_col = "is_reference"
        elif grey_mode in {"none", "off", "false", "0"}:
            grey_col = None
        else:
            grey_col = "is_reference"
        if grey_col is None:
            print(f"[reference] {key}: score UMAP greying disabled (score_umap_grey_mask='none').")
        elif grey_col in adata.obs.columns:
            print(f"[reference] {key}: score UMAP greying uses {grey_col!r} "
                  f"({int(adata.obs[grey_col].astype(bool).sum()):,}/{adata.n_obs:,} cells greyed).")

        # ---- Optional score calibration. When use_calibrated_scores=false, stale
        #      mutation_calib_* columns may still exist on reused h5ads, but every
        #      cfg-aware downstream step resolves to raw mutation_prob_* instead. ----
        if cfg.use_calibrated_scores:
            adata = sa.singlecell.calibrate_scores(adata, cfg, drivers=drivers)
        else:
            print(f"[singlecell] {key}: use_calibrated_scores=false; using raw mutation_prob_* "
                  "for cfg-aware statistics and plots.")
        # malignant-compartment call (for within-tumour longitudinal / shift)
        malig_col = None
        if drivers:
            adata, _mthr = sa.singlecell.derive_malignant_compartment(adata, cfg, drivers=drivers)
            if cfg.restrict_longitudinal_to_malignant:
                # Prefer the dataset's own malignant annotation (van Galen "-like") over
                # the threshold-based scope_malignant call: it cleanly excludes normal
                # T/NK from the trajectory rather than relying on a score cutoff.
                if "is_malignant_label" in adata.obs.columns and adata.obs["is_malignant_label"].astype(bool).any():
                    malig_col = "is_malignant_label"
                    print("[run_all] malignant compartment for longitudinal = 'is_malignant_label' "
                          "(label-based, excludes normal lineages).")
                else:
                    malig_col = "scope_malignant"

        # Same-cell-state guarded raw scores. These are the main guardrail for
        # NPM1-like cases where healthy HSPC/myeloid cells share the learned program
        # and make global tumour-vs-reference Cliff's deltas negative.
        if getattr(cfg, "enable_reference_guardrails", True) and drivers:
            try:
                adata, guard_tbl = sa.reference.add_cellstate_guarded_scores(
                    adata, cfg, drivers=drivers, reference_mask_col=guard_ref_col,
                    cellstate_col=getattr(cfg, "cellstate_col", "celltype_coarse"),
                    min_reference_per_state=getattr(cfg, "cellstate_min_reference_cells", 25))
                if not guard_tbl.empty:
                    M.save_table(f"Reference_cellstate_guardrail_{key}", guard_tbl)
                diag_tables = []
                for rcol in ["is_reference", guard_ref_col, "is_reference_lymphoid_BM1_BM4",
                             "is_reference_myeloid_prog_BM1_BM4", "is_reference_hspc_prog_BM1_BM4"]:
                    if rcol in adata.obs.columns and adata.obs[rcol].astype(bool).sum() > 0:
                        dd = sa.reference.reference_direction_diagnostics(
                            adata, cfg, drivers=drivers, reference_mask_col=rcol,
                            malignant_mask_col=("scope_malignant" if "scope_malignant" in adata.obs.columns else None))
                        if not dd.empty:
                            dd["reference_set"] = rcol
                            diag_tables.append(dd)
                if diag_tables:
                    diag_guard = pd.concat(diag_tables, ignore_index=True)
                    M.save_table(f"Reference_direction_sensitivity_{key}", diag_guard)
                    flagged = diag_guard[diag_guard.get("reference_high_flag", False).astype(bool)]
                    flagged_drivers = sorted(flagged['driver'].dropna().astype(str).unique().tolist()) if not flagged.empty else []
                    adata.uns["reference_high_drivers"] = flagged_drivers
                    if flagged_drivers:
                        print(f"[reference] {key}: {len(flagged_drivers)} driver(s) are below at least one healthy reference set; "
                              "use cell-state residual/percentile outputs for these drivers.")
            except Exception as e:
                print(f"[reference] cell-state guardrails skipped for {key}: {e}")

        # Primary claim-bearing score mode. Once guarded columns exist, use them as
        # the default everywhere Fig3/Fig4/Fig5 compare cells or patients against
        # normal/reference baselines. Raw/configured scores remain only in low-level
        # diagnostics that construct the guardrail flags.
        primary_sc_score_mode = "configured"
        if getattr(cfg, "enable_reference_guardrails", True) and drivers:
            guarded_mode = getattr(cfg, "longitudinal_guarded_mode", "cellstate_residual")
            if guarded_mode and guarded_mode != "none" and any(
                (f"mutation_resid_{d}" in adata.obs.columns) or (f"mutation_refpct_{d}" in adata.obs.columns)
                for d in drivers
            ):
                primary_sc_score_mode = guarded_mode
                print(f"[reference] {key}: primary single-cell/reference comparisons use guarded score_mode='{primary_sc_score_mode}'.")

        # ---- CONFIDENCE: ground-truth-free per-driver confidence (the signal/noise
        #      filter). Combines bulk transferability, spatial coherence, score
        #      concentration, and cross-modal CNV agreement. ----
        conf_map = {}
        if run("singlecell") or run("umap"):
            bulk_auroc = (bulk_table[bulk_table.cancer == cancer].set_index("driver")["auroc"].to_dict()
                          if not bulk_table.empty else None)
            # If the bulk REPORT stage didn't run, the discrimination table is empty
            # and T would collapse to its floor. Recover it directly from the fitted
            # bulk pipeline (cached per cancer) so transferability always counts.
            if not bulk_auroc and cancer in fitted:
                try:
                    _bp, _abulk, _labels = fitted[cancer]
                    bulk_auroc = sa.bulk.bulk_auroc_map(_abulk, _labels, cfg)
                    print(f"[confidence] {key}: computed bulk AUROC for {len(bulk_auroc)} drivers "
                          f"from the fitted pipeline (bulk report stage not run).")
                except Exception as e:
                    print(f"[confidence] {key}: could not compute bulk AUROC ({e}); T will be floored.")
            cnv_for_conf = (sa.cnv.load_cnv_calls(spec["cnv_calls_csv"], dataset=cancer)
                            if _exists(spec.get("cnv_calls_csv")) else None)
            conf_table = sa.confidence.driver_confidence(
                adata, cfg, drivers=drivers, bulk_auroc=bulk_auroc, cnv_calls=cnv_for_conf)
            if not conf_table.empty:
                # surface how much real signal each component carries (helps see when
                # X is dead because CNV didn't match, or AML is simply CNV-quiet)
                import numpy as _np
                cov = {c: int(_np.isfinite(conf_table[c]).sum()) for c in
                       ["T_transferability", "S_spatial", "C_concentration", "X_crossmodal"]
                       if c in conf_table.columns}
                print(f"[confidence] {key}: component coverage (non-NaN drivers) = {cov}")
                M.save_table(f"Fig3_confidence_{key}", conf_table)
                M.register_figure("Fig3_singlecell",
                    sa.plots.confidence_bar(conf_table,
                        cfg.figdir("fig3_singlecell") / f"Fig3_confidence_{key}.png",
                        title_suffix=f"  ({key})"))
                conf_map = conf_table.set_index("driver")["confidence"].to_dict()
                hi = conf_table[conf_table["confidence"] >= 0.4]["driver"].tolist()
                print(f"[run_all] {key}: confidence computed for {len(conf_table)} drivers; "
                      f"{len(hi)} at confidence>=0.4 (likely real): {hi[:12]}")
                # merge per-cell scores with per-driver confidence -> mutation_adj_*
                # (raw x confidence); suppresses low-confidence overcalling on the
                # UMAPs/violins. Uses RAW probabilities, so the reference-ECDF
                # 'uniform reference' artefact that makes healthy cells look lit up
                # for myeloid drivers (NPM1) is avoided.
                try:
                    adata = sa.confidence.add_confidence_adjusted_scores(adata, conf_table)
                except Exception as e:
                    print(f"[run_all] {key}: adjusted scores skipped ({e})")

        # Score provenance audit: this is the sanity check table for the exact scores
        # all downstream plots/tables are about to consume in this run.
        if sa.singlecell.scope_score_columns(adata, include_raw=True, include_derived=True):
            if args.overwrite_scored_h5ad and str(score_source).startswith("rescored"):
                if scored_path:
                    h5ad_write_path = scored_path
                else:
                    print(f"[singlecell-output] WARNING {key}: --overwrite-scored-h5ad requested, "
                          "but this cohort has no scored_h5ad path. Not overwriting raw_h5ad.")
            audit_rows = _score_audit_rows(
                key, cancer, adata, score_source=score_source, input_h5ad=score_input_used,
                scored_h5ad=scored_path, write_h5ad=h5ad_write_path, cfg=cfg)
            M.save_table(f"SingleCell_score_audit_{key}", pd.DataFrame(audit_rows))
            if h5ad_write_path:
                try:
                    _write_h5ad_atomic(adata, h5ad_write_path, label=key)
                except Exception as e:
                    print(f"[singlecell-output] ERROR {key}: could not overwrite {h5ad_write_path}: {e}")

        # ---- 3. SINGLECELL (Fig 3) ----
        if run("singlecell"):
            print(f"\n########## STAGE 3: SINGLE-CELL [{key}] ##########")
            pc, cell_long = sa.singlecell.per_cell_validation(adata, cfg, drivers=drivers, score_mode=primary_sc_score_mode)
            if not pc.empty:
                M.save_table(f"Fig3_percell_{key}", pc)
                percell_stats_for_longitudinal = pc.copy()
                for d in pc.driver:
                    _pc_idx = pc.set_index("driver")
                    pv = _pc_idx["mannwhitney_p"].get(d)
                    qv = _pc_idx["fdr_q"].get(d) if "fdr_q" in _pc_idx.columns else None
                    refsc = None
                    rc = pc.set_index("driver")["score_col"].get(d, cfg.score_col(adata, d))
                    ref_mask_col = guard_ref_col if str(rc).startswith(("mutation_resid_", "mutation_refpct_")) and guard_ref_col in adata.obs.columns else "is_reference"
                    if ref_mask_col in adata.obs.columns and adata.obs[ref_mask_col].any():
                        refsc = adata.obs.loc[adata.obs[ref_mask_col], rc].astype(float).dropna().values
                    M.register_figure("Fig3_singlecell",
                        sa.plots.percell_violin(cell_long, d,
                            cfg.figdir("fig3_singlecell") / f"Fig3_violin_{key}_{d}.png",
                            p_annot=pv, q_annot=qv, reference_scores=refsc,
                            describe=cfg.score_kind(rc), score_label=cfg.score_label_for_col(rc, d),
                            confidence=conf_map.get(d)))
                # one compact summary across all genotyped drivers
                M.register_figure("Fig3_singlecell",
                    sa.plots.percell_summary_heatmap(pc,
                        cfg.figdir("fig3_singlecell") / f"Fig3_summary_{key}.png"))
                # GUARDED Fig 3: same violins on the within-cell-state guarded score,
                # so normal cells (which a myeloid-leaning driver inflates) are
                # de-emphasised and Mut-vs-WT is read against same-lineage healthy cells.
                if primary_sc_score_mode == "configured" and getattr(cfg, "enable_reference_guardrails", True) and \
                   any(f"mutation_refpct_{d}" in adata.obs.columns or f"mutation_resid_{d}" in adata.obs.columns for d in drivers):
                    gmode = getattr(cfg, "longitudinal_guarded_mode", "cellstate_residual")
                    pcg, cell_long_g = sa.singlecell.per_cell_validation(adata, cfg, drivers=drivers, score_mode=gmode)
                    if not pcg.empty:
                        M.save_table(f"Fig3_percell_guarded_{key}", pcg)
                        for d in pcg.driver:
                            _pcg_idx = pcg.set_index("driver")
                            pvg = _pcg_idx["mannwhitney_p"].get(d)
                            qvg = _pcg_idx["fdr_q"].get(d) if "fdr_q" in _pcg_idx.columns else None
                            M.register_figure("Fig3_singlecell",
                                sa.plots.percell_violin(cell_long_g, d,
                                    cfg.figdir("fig3_singlecell") / f"Fig3_violin_guarded_{key}_{d}.png",
                                    p_annot=pvg, q_annot=qvg, describe="guarded",
                                    score_label=f"{d} guarded score (within-lineage)",
                                    confidence=conf_map.get(d)))
                # VALIDATE the ground-truth-free confidence against this cohort's
                # per-cell genotype, where it exists (e.g. van Galen).
                if not conf_table.empty:
                    val = sa.confidence.validate_confidence(conf_table, pc, fdr_alpha=getattr(cfg, "fdr_alpha", 0.10))
                    if val.get("spearman_rho") is not None and val.get("merged") is not None \
                       and "is_real" in val["merged"].columns:
                        M.save_table(f"Fig3_confidence_validation_{key}", val["merged"])
                        M.register_figure("Fig3_singlecell",
                            sa.plots.confidence_vs_truth(val,
                                cfg.figdir("fig3_singlecell") / f"Fig3_confidence_validation_{key}.png"))
                        rho, auc = val.get("spearman_rho"), val.get("auroc_real_vs_conf")
                        print(f"[run_all] {key}: confidence VALIDATED vs ground truth — "
                              f"Spearman ρ={rho:.2f}" + (f", AUROC(real)={auc:.2f}" if auc==auc else ""))
            # direction / lineage-confound diagnostic (the "anything else" finding)
            diag = sa.singlecell.reference_direction_diagnostic(adata, cfg, drivers=drivers, score_mode=primary_sc_score_mode)
            if not diag.empty:
                M.save_table(f"Fig3_direction_diagnostic_{key}", diag)
                M.register_figure("Fig3_singlecell",
                    sa.plots.reference_direction_bar(diag,
                        cfg.figdir("fig3_singlecell") / f"Fig3_direction_{key}.png"))
                n_flag = int(diag["lineage_confound_flag"].sum())
                if n_flag:
                    print(f"[run_all] ⚠ {n_flag} driver(s) show tumour-vs-reference direction "
                          f"DISAGREEING with per-cell Mut-vs-WT — likely lineage-confounded "
                          f"reference. See Fig3_direction_diagnostic_{key}.csv; the per-cell "
                          f"Mut-vs-WT result is the trustworthy one.")
            if status is not None and has_patient:
                br = sa.singlecell.patient_bridge(adata, status, cfg, drivers=drivers, score_mode=primary_sc_score_mode)
                if not br.empty:
                    M.save_table(f"Fig3_bridge_{key}", br)

        # ---- 3b. UMAP MAPS (Fig 4) ----
        if run("singlecell") or run("umap"):
            print(f"\n########## STAGE 4: UMAP MAPS [{key}] ##########")
            adata = _scope_prepare_umap_for_plotting(adata, cohort_key=key, spec=spec, args=args)
            cancer_type_for_plots = sa.priors.cancer_type_for_cohort(key, explicit=spec.get("cancer_type"))
            plot_drivers = _plot_drivers_for_stage(cancer_type_for_plots, drivers, cfg, conf_map)
            try:
                M.register_figure("Fig4_umap",
                    sa.plots.umap_probability_panels(adata, plot_drivers,
                        cfg.figdir("fig4_umap") / f"Fig4_umap_prob_{key}.png", cfg=cfg,
                        confidence=conf_map, reference_mask_col=grey_col, score_mode=primary_sc_score_mode))
                M.register_figure("Fig4_umap",
                    sa.plots.umap_context_panel(adata,
                        cfg.figdir("fig4_umap") / f"Fig4_umap_context_{key}.png",
                        celltype_col=celltype_col, reference_mask_col=grey_col,
                        extra_cols=["patient"] if has_patient else None))
                ck = celltype_col
                if ck in adata.obs.columns:
                    M.register_figure("Fig4_umap",
                        sa.plots.cluster_probability_heatmap(adata, plot_drivers, ck,
                            cfg.figdir("fig4_umap") / f"Fig4_celltype_heatmap_{key}.png", cfg=cfg,
                            confidence=conf_map, score_mode=primary_sc_score_mode,
                            min_confidence=getattr(cfg, "confidence_display_min", 0.2),
                            cluster=True))
                    if primary_sc_score_mode == "configured" and getattr(cfg, "enable_reference_guardrails", True) and any(f"mutation_resid_{d}" in adata.obs.columns for d in drivers):
                        M.register_figure("Fig4_umap",
                            sa.plots.cluster_probability_heatmap(adata, plot_drivers, ck,
                                cfg.figdir("fig4_umap") / f"Fig4_celltype_heatmap_guarded_{key}.png",
                                cfg=cfg, score_mode=getattr(cfg, "longitudinal_guarded_mode", "cellstate_residual"),
                                confidence=conf_map, min_confidence=getattr(cfg, "confidence_display_min", 0.2),
                                cluster=True))
                if primary_sc_score_mode == "configured" and getattr(cfg, "enable_reference_guardrails", True) and any(f"mutation_resid_{d}" in adata.obs.columns for d in drivers):
                    M.register_figure("Fig4_umap",
                        sa.plots.umap_probability_panels(adata, plot_drivers,
                            cfg.figdir("fig4_umap") / f"Fig4_umap_cellstate_guarded_{key}.png", cfg=cfg,
                            confidence=conf_map, score_mode=getattr(cfg, "longitudinal_guarded_mode", "cellstate_residual"),
                            reference_mask_col=grey_col))
                # NEW: per-driver triptych [patient | ground-truth genotype | score]
                # + a patient/celltype/malignant context strip, with legends.
                try:
                    pcol = "patient" if has_patient else spec.get("sample_col", "patient")
                    # Cancer-type prior: which mutations to expect when a cohort lacks
                    # per-patient genotype for a scored driver (BRCA/GBM/PAAD). Saved so
                    # the expectation is explicit; never used to fake per-patient calls.
                    cancer_type = cancer_type_for_plots
                    if cancer_type and plot_drivers:
                        M.save_table(f"Expected_drivers_{key}",
                                     sa.priors.expected_drivers_table(cancer_type, list(plot_drivers)))
                    # Paginate the genotype/score triptych so drivers like NPM1 are
                    # never dropped from the bottom of a giant grid.
                    gdrivers = []
                    _src = {"per-cell": [], "patient": [], "prior/none": []}
                    audit = sa.umap_panels.triptych_genotype_audit(
                        adata, plot_drivers, patient_status=status, patient_col=pcol)
                    if audit is not None and not audit.empty:
                        M.save_table(f"Fig4_triptych_genotype_audit_{key}", audit)
                    for d in plot_drivers:
                        ar = audit[audit["driver"].astype(str).str.upper().eq(str(d).upper())] if audit is not None and not audit.empty else pd.DataFrame()
                        source = str(ar.iloc[0]["source"]) if not ar.empty else "prior/none"
                        gcol = (ar.iloc[0].get("genotype_col") if not ar.empty else None)
                        has_cell_truth = source == "per-cell"
                        has_patient_truth = source == "patient"
                        if has_cell_truth or has_patient_truth:
                            gdrivers.append(d)
                        (_src["per-cell"] if has_cell_truth else
                         _src["patient"] if has_patient_truth else _src["prior/none"]).append(
                            f"{d}({gcol})" if has_cell_truth and pd.notna(gcol) else d)
                    print(f"[triptych] {key} genotype source — per-cell: {_src['per-cell']} | "
                          f"patient-status: {_src['patient']} | prior/none: {_src['prior/none']}")
                    if audit is not None and not audit.empty:
                        used = audit[audit["source"].isin(["per-cell", "patient"])]
                        n_patient = int((used["source"] == "patient").sum())
                        n_cell = int((used["source"] == "per-cell").sum())
                        print(f"[triptych] {key}: audit saved with {n_cell} per-cell and "
                              f"{n_patient} patient-level mutation panels. "
                              f"See Fig4_triptych_genotype_audit_{key}.csv.")
                    if not gdrivers:
                        gdrivers = list(plot_drivers)
                    page_size = 10
                    for page_i in range(0, len(gdrivers), page_size):
                        page = gdrivers[page_i: page_i + page_size]
                        suffix_page = "" if len(gdrivers) <= page_size and page_i == 0 else f"_p{page_i // page_size + 1:02d}"
                        M.register_figure("Fig4_umap",
                            sa.umap_panels.driver_triptych_grid(adata, page,
                                cfg.figdir("fig4_umap") / f"Fig4_umap_triptych_{key}{suffix_page}.png",
                                cfg=cfg, confidence=conf_map, patient_col=pcol, only_genotyped=True,
                                max_rows=page_size, reference_col=grey_col,
                                score_prefer=(("resid", "refpct", "adj", "raw", "calib") if primary_sc_score_mode != "configured" else ("adj", "raw", "calib")),
                                patient_status=status, cancer_type=cancer_type))
                    M.register_figure("Fig4_umap",
                        sa.umap_panels.context_strip(adata,
                            cfg.figdir("fig4_umap") / f"Fig4_umap_context_strip_{key}.png",
                            patient_col=pcol, celltype_col=ck))
                except Exception as e:
                    print(f"[run_all] triptych/context UMAP panels skipped for {key}: {e}")
            except Exception as e:
                print(f"[run_all] UMAP stage skipped for {key}: {e}")

        # ---- 4. LONGITUDINAL (Fig 3b) — the gold ----
        if run("longitudinal") and spec.get("is_longitudinal") and \
           adata.obs.get("timepoint") is not None and adata.obs["timepoint"].notna().any():
            print(f"\n########## STAGE 5: LONGITUDINAL [{key}] ##########")
            if malig_col:
                print(f"[run_all] longitudinal restricted to malignant compartment "
                      f"('{malig_col}') to avoid lineage-admixture confounding.")
            long_ref_col = spec.get("longitudinal_reference_mask_col", "is_reference")
            guarded_ref_for_long = spec.get(
                "longitudinal_guarded_reference_mask_col",
                guard_ref_col if guard_ref_col in adata.obs.columns else long_ref_col,
            )
            long_primary_ref_col = guarded_ref_for_long if primary_sc_score_mode != "configured" else long_ref_col
            long_preserve_cols = [long_ref_col, guarded_ref_for_long, long_primary_ref_col]

            adata_long = _filter_patients(
                adata,
                include=spec.get("longitudinal_include_patients"),
                exclude=spec.get("longitudinal_exclude_patients"),
                patient_col="patient",
                label=f"longitudinal {key}",
                preserve_mask_cols=long_preserve_cols,
            )
            long_sample_col = spec.get("longitudinal_sample_col", spec.get("sample_col", "Sample"))
            adata_long = _filter_longitudinal_samples(
                adata_long,
                include_samples=spec.get("longitudinal_include_samples"),
                exclude_samples=spec.get("longitudinal_exclude_samples"),
                include_regex=spec.get("longitudinal_include_samples_regex"),
                exclude_regex=spec.get("longitudinal_exclude_samples_regex"),
                sample_col=long_sample_col,
                timepoint_col="timepoint",
                preserve_mask_cols=long_preserve_cols,
                label=f"longitudinal {key}",
            )
            include_singletons = bool(spec.get("longitudinal_include_single_timepoint_patients",
                                                spec.get("longitudinal_include_singletons", False)))
            if include_singletons:
                print(f"[longitudinal] {key}: including one-timepoint samples as singleton points; "
                      "trend rho/p will be NaN for singleton patient-driver pairs.")
            traj = sa.longitudinal.longitudinal_trajectory(adata_long, cfg, drivers=drivers,
                                                           malignant_only_col=malig_col,
                                                           reference_mask_col=long_primary_ref_col,
                                                           score_mode=primary_sc_score_mode,
                                                           include_single_timepoint_patients=include_singletons)
            if not traj.empty:
                traj = _add_driver_stats_columns(traj, conf_map, percell_stats_for_longitudinal)
                M.save_table(f"Fig3b_trajectory_{key}", traj)
                trend = sa.longitudinal.trajectory_trend_test(traj, cfg)
                trend = _add_driver_stats_columns(trend, conf_map, percell_stats_for_longitudinal)
                M.save_table(f"Fig3b_trend_{key}", trend)
                suffix_parts = []
                if spec.get("longitudinal_include_patients") or spec.get("longitudinal_exclude_patients"):
                    suffix_parts.append("patient_filtered")
                if (spec.get("longitudinal_include_samples") or spec.get("longitudinal_exclude_samples")
                        or spec.get("longitudinal_include_samples_regex") or spec.get("longitudinal_exclude_samples_regex")):
                    suffix_parts.append("sample_filtered")
                suffix = ("_" + "_".join(suffix_parts)) if suffix_parts else ""
                for d in traj.driver.unique():
                    M.register_figure("Fig3b_longitudinal",
                        sa.plots.treatment_response_trajectory(traj, d,
                            cfg.figdir("fig3b_longitudinal") / f"Fig3b_traj_{key}_{d}{suffix}.png"))

                # Guarded longitudinal: same-cell-state residual/percentile scores.
                # This is the primary plot when global healthy reference is higher
                # than AML cells (e.g. NPM1/HSPC programme overlap).
                guarded_mode = getattr(cfg, "longitudinal_guarded_mode", "cellstate_residual")
                if primary_sc_score_mode == "configured" and guarded_mode and guarded_mode != "none" and any(f"mutation_resid_{d}" in adata_long.obs.columns for d in drivers):
                    gref = spec.get("longitudinal_guarded_reference_mask_col", guard_ref_col if guard_ref_col in adata_long.obs.columns else long_ref_col)
                    gtraj = sa.longitudinal.longitudinal_trajectory(
                        adata_long, cfg, drivers=drivers, malignant_only_col=malig_col,
                        reference_mask_col=gref, score_mode=guarded_mode,
                        include_single_timepoint_patients=include_singletons)
                    if not gtraj.empty:
                        M.save_table(f"Fig3b_trajectory_guarded_{key}", gtraj)
                        gmetric = "median_diff"
                        gtrend = sa.longitudinal.trajectory_trend_test(gtraj, cfg, shift_metric=gmetric)
                        M.save_table(f"Fig3b_trend_guarded_{key}", gtrend)
                        for d in gtraj.driver.unique():
                            M.register_figure("Fig3b_longitudinal",
                                sa.plots.treatment_response_trajectory(gtraj, d,
                                    cfg.figdir("fig3b_longitudinal") / f"Fig3b_traj_guarded_{key}_{d}{suffix}.png",
                                    metric=gmetric))

                # per-patient density overlays + timepoint violins, in per-patient subdirs
                for pat in traj.patient.unique():
                    pdir = cfg.figdir("fig3b_longitudinal", str(pat))
                    for d in traj[traj.patient == pat].driver.unique():
                        dstats = _driver_stats_from_tables(d, conf_map, percell_stats_for_longitudinal)
                        ov = sa.longitudinal.density_overlay_table(adata_long, pat, d, cfg,
                                                                    reference_mask_col=long_primary_ref_col, score_mode=primary_sc_score_mode)
                        ov["driver_stats"] = dstats
                        M.register_figure("Fig3b_longitudinal",
                            sa.plots.density_overlay(ov, pdir / f"density_{d}{suffix}.png", driver_stats=dstats))
                        sp = sa.longitudinal.percell_timepoint_split(adata_long, pat, d, cfg=cfg, score_mode=primary_sc_score_mode)
                        sp.attrs["driver_stats"] = dstats
                        score_label = sp.attrs.get("score_label", cfg.score_label_for_col(sp.attrs.get("score_col", cfg.longitudinal_score_col(adata_long, d)), d))
                        sp_sig = sa.plots.timepoint_genotype_violin_stats(sp)
                        if not sp_sig.empty:
                            sp_sig.insert(0, "driver", d)
                            sp_sig.insert(0, "patient", pat)
                            sp_sig.insert(0, "dataset", key)
                            sp_sig["score_col"] = sp.attrs.get("score_col", "")
                            sp_sig["score_label"] = score_label
                            M.save_table(f"Fig3b_tpviolin_significance_{_slug(key)}_{_slug(pat)}_{_slug(d)}{suffix}", sp_sig)
                        M.register_figure("Fig3b_longitudinal",
                            sa.plots.timepoint_genotype_violin(sp, pat, d,
                                pdir / f"tpviolin_{d}{suffix}.png", score_label=score_label,
                                driver_stats=dstats, significance_stats=sp_sig))
                        emit_secondary_guarded_density = bool(
                            spec.get("emit_secondary_guarded_density", False)
                            or getattr(cfg, "emit_secondary_guarded_density", False)
                        )
                        if emit_secondary_guarded_density and getattr(cfg, "enable_reference_guardrails", True) and f"mutation_resid_{d}" in adata_long.obs.columns:
                            gov = sa.longitudinal.density_overlay_table(adata_long, pat, d, cfg,
                                                                        reference_mask_col=guarded_ref_for_long,
                                                                        score_mode=getattr(cfg, "longitudinal_guarded_mode", "cellstate_residual"))
                            if gov.get("timepoints"):
                                M.register_figure("Fig3b_longitudinal",
                                    sa.plots.density_overlay(gov, pdir / f"density_guarded_{d}{suffix}.png",
                                                             score_label=gov.get("score_label"), driver_stats=dstats))
                            spg = sa.longitudinal.percell_timepoint_split(adata_long, pat, d, cfg=cfg,
                                                                          score_mode=getattr(cfg, "longitudinal_guarded_mode", "cellstate_residual"))
                            if not spg.empty:
                                spg_score_label = spg.attrs.get("score_label", f"{d} guarded score")
                                spg_sig = sa.plots.timepoint_genotype_violin_stats(spg)
                                if not spg_sig.empty:
                                    spg_sig.insert(0, "driver", d)
                                    spg_sig.insert(0, "patient", pat)
                                    spg_sig.insert(0, "dataset", key)
                                    spg_sig["score_col"] = spg.attrs.get("score_col", "")
                                    spg_sig["score_label"] = spg_score_label
                                    M.save_table(f"Fig3b_tpviolin_significance_guarded_{_slug(key)}_{_slug(pat)}_{_slug(d)}{suffix}", spg_sig)
                                M.register_figure("Fig3b_longitudinal",
                                    sa.plots.timepoint_genotype_violin(spg, pat, d,
                                        pdir / f"tpviolin_guarded_{d}{suffix}.png",
                                        score_label=spg_score_label,
                                        driver_stats=dstats, significance_stats=spg_sig))

        # ---- 5. PANCANCER (Fig 5 / Table 2) ----
        if run("pancancer"):
            print(f"\n########## STAGE 6: PANCANCER [{key}] ##########")
            res = sa.pancancer.validate_cohort_by_tier(adata, cancer, cfg,
                                                       patient_status=status, drivers=drivers,
                                                       malignant_mask_col=malig_col,
                                                       score_mode=primary_sc_score_mode)
            for name, tbl in res.items():
                if isinstance(tbl, pd.DataFrame) and not tbl.empty:
                    M.save_table(f"Fig5_{key}_{name}", tbl)

        # ---- 6. CNV (Fig 6) ----
        if run("cnv") and _exists(spec.get("cnv_calls_csv")):
            print(f"\n########## STAGE 7: CNV [{key}] ##########")
            cnv_calls = sa.cnv.load_cnv_calls(spec["cnv_calls_csv"], dataset=spec.get("cancer"))
            # v10.3.3 FIX (CNV concordance artifact): in all-label exploratory mode,
            # scored_drivers() includes rare/noisy heads trained on very few positives.
            # Taking max() over them makes scope_max_driver_score track raw signal
            # magnitude and spuriously co-vary with CNV burden (GBM Spearman flipped
            # -0.36 -> +0.84 with an UNCHANGED CNV side; BRCA collapsed +0.81 -> +0.19).
            # Restrict the scOPE malignant-call / CNV-concordance driver set to the locked
            # primary drivers. No-op in primary/target-driver runs (manuscript mode).
            cnv_drivers = drivers
            if getattr(cfg, "score_all_label_drivers", False):
                _primary = {str(d).upper() for d in cancer_drivers(cancer)}
                _restricted = [d for d in drivers if str(d).upper() in _primary]
                if _restricted:
                    cnv_drivers = _restricted
                    print(f"[cnv] {key}: CNV concordance restricted to "
                          f"{len(cnv_drivers)}/{len(drivers)} locked primary drivers "
                          f"(all-label exploratory scoring detected): {cnv_drivers}")
                else:
                    print(f"[cnv] {key}: no locked primary drivers among scored set; "
                          f"using all {len(drivers)} scored drivers for CNV concordance.")
            adata, thr = sa.cnv.derive_scope_malignant_call(adata, cfg, drivers=cnv_drivers)
            conc = sa.cnv.cnv_concordance(adata, cnv_calls)
            if conc.get("n", 0) > 0:
                disc = sa.cnv.discordance_analysis(conc["merged"])
                M.save_table(f"Fig6_discordance_{key}", disc)
                M.register_figure("Fig6_cnv",
                    sa.plots.cnv_discordance_bar(disc,
                        cfg.figdir("fig6_cnv") / f"Fig6_discordance_{key}.png", ari=conc["ari"]))
                M.register_figure("Fig6_cnv",
                    sa.plots.cnv_confusion_heatmap(conc,
                        cfg.figdir("fig6_cnv") / f"Fig6_confusion_{key}.png"))
                # continuous score-vs-burden
                corr = sa.cnv.cnv_score_correlation(adata, cnv_calls)
                M.save_table(f"Fig6_score_correlation_{key}", pd.DataFrame([corr]))
                M.register_figure("Fig6_cnv",
                    sa.plots.cnv_score_scatter(conc["merged"],
                        cfg.figdir("fig6_cnv") / f"Fig6_scorescatter_{key}.png",
                        rho=corr.get("spearman_rho")))
                # per-patient malignant-fraction agreement
                frac = sa.cnv.per_patient_malignant_fraction(adata, cnv_calls, patient_col=("patient" if has_patient else spec.get("sample_col", "patient")))
                if not frac.empty:
                    M.save_table(f"Fig6_patient_fraction_{key}", frac)
                    M.register_figure("Fig6_cnv",
                        sa.plots.cnv_malignant_fraction_scatter(frac,
                            cfg.figdir("fig6_cnv") / f"Fig6_patientfrac_{key}.png"))
                else:
                    print(f"[cnv] {key}: per-patient malignant fraction skipped because no usable patient column survived the CNV merge.")
                if status is not None:
                    sa_tbl = sa.cnv.diploid_patient_standalone(adata, cnv_calls, status, cfg)
                    if not sa_tbl.empty:
                        M.save_table(f"Fig6_diploid_standalone_{key}", sa_tbl)
            else:
                # not silent: record why CNV produced nothing
                M.save_table(f"Fig6_cnv_match_diagnostics_{key}",
                             pd.DataFrame([conc.get("match_diagnostics", {"note": conc.get("note")})]))

    # ================= 7. ROBUSTNESS =================
    if run("robustness"):
        print("\n########## STAGE 7: ROBUSTNESS ##########")
        for cancer, (bp, a_bulk, labels) in fitted.items():
            ld = sa.robustness.latent_dim_ablation(a_bulk, labels, cfg)
            if not ld.empty:
                M.save_table(f"Robustness_latentdim_{cancer}", ld)

    # ================= 8. REPORT =================
    if run("report") or run("bulk"):
        print("\n########## STAGE 8: REPORT ##########")
        prior_table = sa.report.build_driver_prior_table(cfg)
        if not bulk_table.empty:
            prior_table = sa.report.merge_priors_with_results(prior_table, bulk_table)
        M.save_table("Table0_driver_priors", prior_table)
    M.write_manifest()
    print("\n✔ run_all complete. See", cfg.output_root)


if __name__ == "__main__":
    main()
```
