## Supplemental_Code for "scOPE identifies which driver-associated expression programs transfer from bulk tumors to single cells": validate_confidence_vangalen.py

```
#!/usr/bin/env python
"""
validate_confidence_vangalen.py
===============================
Run the ground-truth-free confidence score on your van Galen scored h5ad and
VALIDATE it against the per-cell genotype you already have, so you can trust it on
the cohorts that LACK ground truth (BRCA / CRC / GBM / LUAD).

The spatial (Moran's I) and cross-modal (CNV) components need the actual cells, so
this runs in YOUR environment where the scored h5ad lives — it is not something the
suite can do from the exported tables alone.

What it does
------------
1. loads the scored van Galen h5ad (per-cell mutation_prob_* already in obs),
2. marks lymphoid reference cells + calibrates (monotone; ranks unchanged),
3. builds the bulk-AUROC prior from Table1_bulk_discrimination.csv (AML rows),
4. loads cnv_calls.csv (dataset=AML) for the cross-modal term,
5. computes driver_confidence(...) WITHOUT touching the genotype,
6. validates against Fig3_percell_AML_vanGalen.csv (Mut-vs-WT) and writes a scatter.

Expected outcome (from your tables): high confidence for NPM1, TP53, DNMT3A, NRAS,
RAD21, SMC3; low for RUNX1, KRAS, IDH2, SETD2, TET2 and the untrainable BCOR /
BCORL1 / EZH2 / KMT2A.

Usage
-----
python drivers/validate_confidence_vangalen.py \
    --scored-h5ad /path/AML_scored_with_muttranscript_eval_labels.h5ad \
    --bulk-table  scope_analysis_outputs/tables/Table1_bulk_discrimination.csv \
    --percell     scope_analysis_outputs/tables/Fig3_percell_AML_vanGalen.csv \
    --cnv-calls   /path/cnv_calls.csv \
    --celltype-col CellType \
    --outdir      confidence_validation
"""
from __future__ import annotations
import argparse
from pathlib import Path

import numpy as np
import pandas as pd

import scope_analysis as sa
from scope_analysis.config import AnalysisConfig


def main() -> None:
    ap = argparse.ArgumentParser()
    ap.add_argument("--scored-h5ad", required=True)
    ap.add_argument("--bulk-table", required=True, help="Table1_bulk_discrimination.csv")
    ap.add_argument("--percell", required=True, help="Fig3_percell_<cohort>.csv (ground truth)")
    ap.add_argument("--cnv-calls", default=None, help="cnv_calls.csv (optional, enables cross-modal term)")
    ap.add_argument("--cancer", default="AML")
    ap.add_argument("--celltype-col", default="CellType")
    ap.add_argument("--reference-celltypes", default="T,B,NK")
    ap.add_argument("--outdir", default="confidence_validation")
    args = ap.parse_args()

    import anndata as ad
    out = Path(args.outdir); out.mkdir(parents=True, exist_ok=True)
    cfg = AnalysisConfig(use_calibrated_scores=True)

    print(f"[load] {args.scored_h5ad}")
    adata = ad.read_h5ad(args.scored_h5ad)
    adata = sa.data.mark_reference_cells(
        adata, celltype_col=args.celltype_col,
        reference_celltypes=tuple(args.reference_celltypes.split(",")))
    drivers = sa.data.scored_drivers(adata)
    adata = sa.singlecell.calibrate_scores(adata, cfg, drivers=drivers)
    print(f"[info] {len(drivers)} scored drivers; {int(adata.obs['is_reference'].sum())} reference cells")

    bulk = pd.read_csv(args.bulk_table)
    bulk_auroc = (bulk[bulk.cancer == args.cancer].set_index("driver")["auroc"].to_dict()
                  if "cancer" in bulk.columns else bulk.set_index("driver")["auroc"].to_dict())

    cnv_calls = None
    if args.cnv_calls:
        try:
            cnv_calls = sa.cnv.load_cnv_calls(args.cnv_calls, dataset=args.cancer)
            print(f"[info] loaded {len(cnv_calls):,} CNV calls for {args.cancer} (cross-modal term ON)")
        except Exception as e:
            print(f"[warn] could not load CNV calls ({e}); cross-modal term OFF")

    conf_table = sa.confidence.driver_confidence(
        adata, cfg, drivers=drivers, bulk_auroc=bulk_auroc, cnv_calls=cnv_calls)
    conf_table.to_csv(out / "confidence_table.csv", index=False)
    print("\n=== confidence (ground-truth-free) ===")
    cols = ["driver", "T_transferability", "S_spatial", "C_concentration",
            "X_crossmodal", "confidence", "tier"]
    print(conf_table[[c for c in cols if c in conf_table.columns]].round(3).to_string(index=False))

    sa.plots.confidence_bar(conf_table, out / "confidence_bar.png", title_suffix=f"  ({args.cancer} van Galen)")

    truth = pd.read_csv(args.percell)
    val = sa.confidence.validate_confidence(conf_table, truth)
    if val.get("merged") is not None and "is_real" in val["merged"].columns:
        val["merged"].to_csv(out / "confidence_validation.csv", index=False)
        sa.plots.confidence_vs_truth(val, out / "confidence_vs_truth.png")
        print(f"\n=== VALIDATION ===\n  overlapping drivers : {val['n']}"
              f"\n  Spearman(conf, truth Cliffs δ) : {val['spearman_rho']:.3f} (p={val['spearman_p']:.3g})"
              f"\n  AUROC(real | confidence)       : {val['auroc_real_vs_conf']}"
              f"\n  real drivers (sig. Mut>WT)     : {val['n_real']}")
        print(f"\n[done] wrote {out}/ (confidence_table.csv, confidence_bar.png, "
              f"confidence_vs_truth.png, confidence_validation.csv)")
    else:
        print(f"\n[warn] only {val.get('n')} overlapping drivers — too few to validate; "
              "confidence_table.csv + confidence_bar.png still written.")


if __name__ == "__main__":
    main()
```
