## Supplemental_Code for "scOPE identifies which driver-associated expression programs transfer from bulk tumors to single cells": environment_minimal.yml

```
name: scope-manuscript
channels:
  - conda-forge
dependencies:
  - python=3.10
  - scope-bio=0.2.0
  - anndata
  - scanpy
  - numpy
  - scipy
  - pandas
  - scikit-learn
  - statsmodels
  - matplotlib
  - infercnvpy
```
