## Supplemental_Code for "scOPE identifies which driver-associated expression programs transfer from bulk tumors to single cells": export_cnv_arm_chromosome_clustermaps_all_cancers.py

```
#!/usr/bin/env python
"""
Export inferCNVpy chromosome- and chromosome-arm-level CNV tables and plots.

This is a drop-in replacement for the chromosome-only exporter. Main changes:
  1) Adds chromosome-arm summaries: chr1p, chr1q, chr2p, chr2q, ... chr22p, chr22q.
  2) Keeps heatmap columns in genomic order by default; columns do NOT cluster.
  3) Still supports batch mode across cancer-specific inferCNVpy .h5ad files.
  4) Writes both chromosome-level and arm-level tables/figures by default.

Expected AnnData content from inferCNVpy:
  adata.obsm["X_cnv"]
  adata.uns["cnv"]["chr_pos"]
  adata.var[["chromosome", "start", "end"]]

Example batch run on Windows:
  python export_cnv_arm_chromosome_clustermaps_all_cancers.py ^
    --h5ad-dir "C:\\Users\\ashfo\\OneDrive\\Desktop\\Pathways+Omics\\scOPE\\scOPE\\data\\scope_robust_real_inputs\\infercnvpy_new" ^
    --datasets AML,BRCA,CRC,GBM,LUAD,PAAD,SKCM ^
    --outdir "C:\\Users\\ashfo\\OneDrive\\Desktop\\Pathways+Omics\\scOPE\\scope_analysis_v2\\cnv_arm_outputs" ^
    --region-level both ^
    --threshold 0.05 ^
    --max-cells-plot 3000

Important notes:
  - Arm assignment uses the midpoint of each inferred CNV window.
  - Default centromere coordinates are approximate GRCh38/hg38 cytoband-style boundaries.
  - If your GTF/gene coordinates are hg19/GRCh37, pass --genome-build hg19.
  - For maximum rigor, pass --centromere-csv with columns:
        chromosome, centromere_start, centromere_end
    or:
        chromosome, start, end
"""

from __future__ import annotations

import argparse
import contextlib
import gc
import json
import math
import os
import re
import threading
import time
import warnings
from pathlib import Path
from typing import Iterable, Sequence

import numpy as np
import pandas as pd

try:
    import matplotlib
    matplotlib.use("Agg", force=True)
    import matplotlib.pyplot as plt
except Exception as e:  # pragma: no cover
    raise RuntimeError("matplotlib is required for plotting") from e

try:
    import seaborn as sns
except Exception as e:  # pragma: no cover
    raise RuntimeError("seaborn is required for plotting") from e

try:
    import scipy.sparse as sp
except Exception:  # pragma: no cover
    sp = None

try:
    from tqdm.auto import tqdm
except Exception:  # pragma: no cover
    class tqdm:
        def __init__(self, iterable=None, total=None, desc=None, unit=None, disable=False, leave=True, **kwargs):
            self.iterable = iterable
            self.total = total
            self.desc = desc or "progress"
            self.disable = disable
            self.n = 0
            if not disable:
                print(f"[progress] {self.desc}", flush=True)
        def __iter__(self):
            if self.iterable is None:
                return iter(())
            for x in self.iterable:
                yield x
                self.update(1)
        def update(self, n=1):
            self.n += n
            if not self.disable and self.total:
                print(f"[progress] {self.desc}: {self.n}/{self.total}", flush=True)
        def set_description(self, desc):
            self.desc = desc
            if not self.disable:
                print(f"[progress] {desc}", flush=True)
        def set_postfix(self, *args, **kwargs):
            pass
        def close(self):
            pass
        def __enter__(self):
            return self
        def __exit__(self, exc_type, exc, tb):
            self.close()


# -----------------------------------------------------------------------------
# Progress / diagnostics
# -----------------------------------------------------------------------------

HEARTBEAT_SECONDS = 30


def _print(msg: str) -> None:
    print(msg, flush=True)


def _format_seconds(seconds: float) -> str:
    seconds = float(seconds)
    if seconds < 60:
        return f"{seconds:.1f}s"
    minutes = seconds / 60
    if minutes < 60:
        return f"{minutes:.1f}m"
    return f"{minutes / 60:.1f}h"


def _rss_gb() -> float | None:
    try:
        import psutil
        return psutil.Process(os.getpid()).memory_info().rss / (1024 ** 3)
    except Exception:
        return None


def _mem_suffix() -> str:
    rss = _rss_gb()
    return "" if rss is None else f", RSS={rss:.2f} GB"


@contextlib.contextmanager
def _timed_step(label: str):
    t0 = time.time()
    _print(f"[step] START {label}{_mem_suffix()}")
    try:
        yield
    finally:
        _print(f"[step] DONE  {label} in {_format_seconds(time.time() - t0)}{_mem_suffix()}")


@contextlib.contextmanager
def _heartbeat(label: str, interval: int | None = None):
    if interval is None:
        interval = HEARTBEAT_SECONDS
    stop_event = threading.Event()
    t0 = time.time()

    def _worker():
        while not stop_event.wait(interval):
            _print(f"[heartbeat] {label}: still running after {_format_seconds(time.time() - t0)}{_mem_suffix()}")

    thread = threading.Thread(target=_worker, daemon=True)
    thread.start()
    try:
        yield
    finally:
        stop_event.set()
        thread.join(timeout=2)


# -----------------------------------------------------------------------------
# Genomic ordering and centromeres
# -----------------------------------------------------------------------------

CANONICAL_CHROMS = [f"chr{i}" for i in range(1, 23)]
CANONICAL_CHROMS_WITH_SEX = CANONICAL_CHROMS + ["chrX", "chrY"]

# Approximate cytoband-style centromere boundaries. These are used only to split
# inferred CNV windows into p/q arms. Passing --centromere-csv overrides them.
CENTROMERES: dict[str, dict[str, tuple[int, int]]] = {
    "hg38": {
        "chr1": (121700000, 125100000), "chr2": (91800000, 96000000),
        "chr3": (87800000, 94000000), "chr4": (48200000, 51800000),
        "chr5": (46100000, 51400000), "chr6": (58500000, 62600000),
        "chr7": (58100000, 62100000), "chr8": (43200000, 47200000),
        "chr9": (42200000, 45500000), "chr10": (38000000, 41600000),
        "chr11": (51000000, 55800000), "chr12": (33200000, 37800000),
        "chr13": (16500000, 18900000), "chr14": (16100000, 18200000),
        "chr15": (17500000, 20500000), "chr16": (35300000, 38400000),
        "chr17": (22700000, 27400000), "chr18": (15400000, 21500000),
        "chr19": (24200000, 28100000), "chr20": (25700000, 30400000),
        "chr21": (10900000, 13000000), "chr22": (13700000, 17400000),
        "chrX": (58100000, 63800000), "chrY": (10300000, 10600000),
    },
    "grch38": {},
    "hg19": {
        "chr1": (121535434, 124535434), "chr2": (92326171, 95326171),
        "chr3": (90504854, 93504854), "chr4": (49660117, 52660117),
        "chr5": (46405641, 49405641), "chr6": (58830166, 61830166),
        "chr7": (58054331, 61054331), "chr8": (43838887, 46838887),
        "chr9": (47367679, 50367679), "chr10": (39254935, 42254935),
        "chr11": (51644205, 54644205), "chr12": (34856694, 37856694),
        "chr13": (16000000, 19000000), "chr14": (16000000, 19000000),
        "chr15": (17000000, 20000000), "chr16": (35335801, 38335801),
        "chr17": (22263006, 25263006), "chr18": (15460898, 18460898),
        "chr19": (24681782, 27681782), "chr20": (26369569, 29369569),
        "chr21": (11288129, 14288129), "chr22": (13000000, 16000000),
        "chrX": (58632012, 61632012), "chrY": (10104553, 13104553),
    },
    "grch37": {},
}
CENTROMERES["grch38"] = CENTROMERES["hg38"]
CENTROMERES["grch37"] = CENTROMERES["hg19"]


def _normalize_chrom(x: object) -> str:
    s = str(x)
    if s.lower().startswith("chr"):
        core = s[3:]
    else:
        core = s
    core = core.upper()
    if core in {"M", "MT", "MITO", "MITOCHONDRIA"}:
        return "chrM"
    return f"chr{core}"


def _chrom_sort_key(x: object) -> tuple[int, str]:
    c = _normalize_chrom(x)
    m = re.fullmatch(r"chr(\d+)", c)
    if m:
        return (int(m.group(1)), "")
    if c == "chrX":
        return (23, "")
    if c == "chrY":
        return (24, "")
    if c == "chrM":
        return (25, "")
    return (99, c)


def _arm_sort_key(x: object) -> tuple[int, int, str]:
    s = str(x)
    m = re.fullmatch(r"(chr(?:\d+|X|Y))([pq])", s)
    if m:
        chrom, arm = m.group(1), m.group(2)
        return (_chrom_sort_key(chrom)[0], 0 if arm == "p" else 1, "")
    if s.endswith("cen"):
        return (_chrom_sort_key(s[:-3])[0], 2, "")
    return (99, 99, s)


def _ordered_chrom_columns(cols: Iterable[object]) -> list[str]:
    return sorted([str(c) for c in cols], key=_chrom_sort_key)


def _ordered_arm_columns(cols: Iterable[object]) -> list[str]:
    return sorted([str(c) for c in cols], key=_arm_sort_key)


def _ordered_region_columns(cols: Iterable[object], region_level: str) -> list[str]:
    if region_level == "arm":
        return _ordered_arm_columns(cols)
    return _ordered_chrom_columns(cols)


def _safe_name(x: object) -> str:
    s = str(x)
    s = re.sub(r"[^\w.\-]+", "_", s).strip("_")
    return s[:120] if s else "NA"


def _read_centromere_csv(path: str | Path | None) -> dict[str, tuple[int, int]] | None:
    if path is None or str(path).strip() == "":
        return None
    p = Path(path)
    if not p.exists():
        raise FileNotFoundError(p)
    df = pd.read_csv(p)
    lower = {c.lower(): c for c in df.columns}
    chrom_col = lower.get("chromosome") or lower.get("chrom") or lower.get("chr")
    start_col = lower.get("centromere_start") or lower.get("cen_start") or lower.get("start")
    end_col = lower.get("centromere_end") or lower.get("cen_end") or lower.get("end")
    if chrom_col is None or start_col is None or end_col is None:
        raise ValueError(
            "--centromere-csv must contain chromosome/chrom/chr and "
            "centromere_start+centromere_end or start+end columns"
        )
    cent = {}
    for _, row in df.iterrows():
        chrom = _normalize_chrom(row[chrom_col])
        cent[chrom] = (int(row[start_col]), int(row[end_col]))
    return cent


def _get_centromeres(genome_build: str = "hg38", centromere_csv: str | Path | None = None) -> dict[str, tuple[int, int]]:
    custom = _read_centromere_csv(centromere_csv)
    if custom is not None:
        return custom
    key = str(genome_build).lower()
    if key == "auto":
        key = "hg38"
    if key not in CENTROMERES:
        raise ValueError(f"Unknown genome build {genome_build!r}; use hg38, hg19, or pass --centromere-csv")
    return dict(CENTROMERES[key])


# -----------------------------------------------------------------------------
# Matrix helpers
# -----------------------------------------------------------------------------


def _maybe_dense(x):
    if sp is not None and sp.issparse(x):
        return x.toarray()
    if hasattr(x, "A"):
        return np.asarray(x.A)
    return np.asarray(x)


def _series_from_matrix_mean(x, axis: int = 1) -> np.ndarray:
    if sp is not None and sp.issparse(x):
        arr = np.asarray(x.mean(axis=axis)).ravel()
    else:
        arr = np.asarray(np.nanmean(x, axis=axis)).ravel()
    return arr.astype(np.float32, copy=False)


def _write_df(df: pd.DataFrame, path: Path) -> None:
    path.parent.mkdir(parents=True, exist_ok=True)
    if path.name.endswith(".csv.gz"):
        df.to_csv(path, compression="gzip")
    else:
        df.to_csv(path)


def _save_current_fig(path_base: Path, dpi: int = 220) -> None:
    path_base.parent.mkdir(parents=True, exist_ok=True)
    plt.savefig(path_base.with_suffix(".png"), dpi=dpi, bbox_inches="tight")
    plt.savefig(path_base.with_suffix(".pdf"), bbox_inches="tight")
    plt.close()


def _save_clustermap(g, path_base: Path, dpi: int = 220) -> None:
    path_base.parent.mkdir(parents=True, exist_ok=True)
    g.savefig(path_base.with_suffix(".png"), dpi=dpi, bbox_inches="tight")
    g.savefig(path_base.with_suffix(".pdf"), bbox_inches="tight")
    plt.close(g.fig)


def _resolve_obsm_key(adata, use_rep: str) -> str:
    candidates = [f"X_{use_rep}", use_rep]
    if use_rep == "cnv":
        candidates += ["X_cnv", "cnv"]
    for k in candidates:
        if k in adata.obsm:
            return k
    raise KeyError(f"Could not find CNV matrix for use_rep={use_rep!r}. Available obsm keys: {list(adata.obsm.keys())}")


def get_cnv_matrix(adata, use_rep: str = "cnv"):
    return adata.obsm[_resolve_obsm_key(adata, use_rep)]


def get_chr_pos(adata, use_rep: str = "cnv") -> dict[str, int]:
    if use_rep not in adata.uns:
        raise KeyError(f"Expected adata.uns[{use_rep!r}], but did not find it.")
    uns = adata.uns[use_rep]
    if isinstance(uns, dict) and "chr_pos" in uns:
        chr_pos = dict(uns["chr_pos"])
    elif hasattr(uns, "keys") and "chr_pos" in uns.keys():
        chr_pos = dict(uns["chr_pos"])
    else:
        raise KeyError(
            f"Expected adata.uns[{use_rep!r}]['chr_pos'], but did not find it. "
            f"Available keys: {list(uns.keys()) if hasattr(uns, 'keys') else type(uns)}"
        )
    return {_normalize_chrom(k): int(v) for k, v in chr_pos.items()}


# -----------------------------------------------------------------------------
# Reconstruct inferCNV windows and assign arms
# -----------------------------------------------------------------------------


def reconstruct_cnv_windows(
    adata,
    *,
    use_rep: str = "cnv",
    window_size: int = 100,
    step: int = 10,
    exclude_chromosomes: Sequence[str] | None = ("chrX", "chrY"),
) -> pd.DataFrame:
    X_cnv = get_cnv_matrix(adata, use_rep=use_rep)
    n_windows = int(X_cnv.shape[1])
    chr_pos = get_chr_pos(adata, use_rep=use_rep)

    missing = {"chromosome", "start", "end"} - set(adata.var.columns)
    if missing:
        raise ValueError(
            f"adata.var is missing required genomic position columns: {missing}. "
            "Need these to reconstruct inferCNV windows and arm-level summaries."
        )

    var = adata.var.loc[:, ["chromosome", "start", "end"]].copy()
    var["chromosome"] = var["chromosome"].map(_normalize_chrom)
    var["gene"] = adata.var_names.astype(str)
    var = var.dropna(subset=["chromosome", "start", "end"]).copy()

    if exclude_chromosomes is not None:
        exclude = {_normalize_chrom(c) for c in exclude_chromosomes}
        var = var.loc[~var["chromosome"].isin(exclude)].copy()
        chr_pos = {c: v for c, v in chr_pos.items() if c not in exclude}

    chromosomes_from_uns = sorted(chr_pos.keys(), key=lambda c: chr_pos[c])
    chromosomes_from_var = set(var["chromosome"].unique())
    chromosomes = [c for c in chromosomes_from_uns if c in chromosomes_from_var]
    if not chromosomes:
        chromosomes = chromosomes_from_uns

    starts = [chr_pos[c] for c in chromosomes]
    ends = starts[1:] + [n_windows]
    chr_counts = {c: int(e - s) for c, s, e in zip(chromosomes, starts, ends)}

    records = []
    running_window_id = 0

    for chrom in chromosomes:
        n_for_chrom = chr_counts.get(chrom, 0)
        if n_for_chrom <= 0:
            continue

        g = var.loc[var["chromosome"].eq(chrom)].copy()
        if g.empty:
            for j in range(n_for_chrom):
                records.append({
                    "window_id": running_window_id,
                    "chromosome": chrom,
                    "window_in_chromosome": j,
                    "start": np.nan,
                    "end": np.nan,
                    "midpoint": np.nan,
                    "first_gene": "",
                    "last_gene": "",
                    "n_genes_in_window": np.nan,
                    "approximate_interval": True,
                })
                running_window_id += 1
            continue

        g["start"] = pd.to_numeric(g["start"], errors="coerce")
        g["end"] = pd.to_numeric(g["end"], errors="coerce")
        g = g.dropna(subset=["start", "end"]).sort_values(["start", "end"])
        genes = g["gene"].to_numpy()
        starts_bp = g["start"].to_numpy()
        ends_bp = g["end"].to_numpy()
        n_genes = len(g)

        if n_genes > window_size:
            win_starts = np.arange(0, n_genes - window_size + 1, step, dtype=int)
        else:
            win_starts = np.array([0], dtype=int)

        used_approx = False
        if len(win_starts) != n_for_chrom:
            max_start = max(0, n_genes - min(window_size, n_genes))
            if n_for_chrom == 1:
                win_starts = np.array([0], dtype=int)
            else:
                win_starts = np.round(np.linspace(0, max_start, n_for_chrom)).astype(int)
            used_approx = True
            warnings.warn(
                f"{chrom}: reconstructed windows to match X_cnv using approximate spacing. "
                f"Check --window-size {window_size} and --step {step} match original infercnvpy run."
            )

        for j, sidx in enumerate(win_starts):
            eidx = min(sidx + window_size - 1, n_genes - 1)
            start_bp = float(starts_bp[sidx])
            end_bp = float(ends_bp[eidx])
            records.append({
                "window_id": running_window_id,
                "chromosome": chrom,
                "window_in_chromosome": j,
                "start": start_bp,
                "end": end_bp,
                "midpoint": (start_bp + end_bp) / 2.0,
                "first_gene": str(genes[sidx]),
                "last_gene": str(genes[eidx]),
                "n_genes_in_window": int(eidx - sidx + 1),
                "approximate_interval": used_approx,
            })
            running_window_id += 1

    windows = pd.DataFrame.from_records(records)
    if len(windows) != n_windows:
        raise RuntimeError(
            f"Window metadata length ({len(windows)}) does not match X_cnv columns ({n_windows}). "
            "Most likely --window-size/--step/excluded chromosomes do not match the original infercnvpy run."
        )
    return windows


def add_chromosome_arm_annotations(
    windows: pd.DataFrame,
    *,
    genome_build: str = "hg38",
    centromere_csv: str | Path | None = None,
    centromere_windows: str = "nearest",
) -> pd.DataFrame:
    """
    Add columns: arm, chromosome_arm, centromere_start, centromere_end.

    centromere_windows:
      - nearest: assign centromeric-spanning windows to nearest p/q by midpoint.
      - cen: label midpoint-inside-centromere windows as chrNcen.
      - drop: set midpoint-inside-centromere windows to NaN; ignored in arm means.
    """
    if centromere_windows not in {"nearest", "cen", "drop"}:
        raise ValueError("centromere_windows must be one of: nearest, cen, drop")

    cent = _get_centromeres(genome_build, centromere_csv)
    out = windows.copy()
    arms = []
    arm_labels = []
    cen_starts = []
    cen_ends = []

    for _, row in out.iterrows():
        chrom = _normalize_chrom(row["chromosome"])
        mid = row.get("midpoint", np.nan)
        cstart, cend = cent.get(chrom, (np.nan, np.nan))
        cen_starts.append(cstart)
        cen_ends.append(cend)

        if not np.isfinite(mid) or not np.isfinite(cstart) or not np.isfinite(cend):
            arm = np.nan
            label = np.nan
        elif mid < cstart:
            arm = "p"
            label = f"{chrom}p"
        elif mid > cend:
            arm = "q"
            label = f"{chrom}q"
        else:
            if centromere_windows == "cen":
                arm = "cen"
                label = f"{chrom}cen"
            elif centromere_windows == "drop":
                arm = np.nan
                label = np.nan
            else:
                cen_mid = (float(cstart) + float(cend)) / 2.0
                arm = "p" if mid < cen_mid else "q"
                label = f"{chrom}{arm}"
        arms.append(arm)
        arm_labels.append(label)

    out["centromere_start"] = cen_starts
    out["centromere_end"] = cen_ends
    out["arm"] = arms
    out["chromosome_arm"] = arm_labels
    return out


# -----------------------------------------------------------------------------
# Tables
# -----------------------------------------------------------------------------


def compute_cell_region_means(
    adata,
    windows: pd.DataFrame,
    *,
    use_rep: str = "cnv",
    region_col: str = "chromosome_arm",
) -> pd.DataFrame:
    X = get_cnv_matrix(adata, use_rep=use_rep)
    if region_col not in windows.columns:
        raise KeyError(f"windows does not contain {region_col!r}")

    win_region = windows[region_col].astype("object")
    keep = win_region.notna()
    regions = list(pd.Series(win_region[keep]).drop_duplicates())
    if region_col == "chromosome_arm":
        regions = _ordered_arm_columns(regions)
    elif region_col == "chromosome":
        regions = _ordered_chrom_columns(regions)
    else:
        regions = sorted(map(str, regions))

    data = {}
    for region in tqdm(regions, desc=f"cell means by {region_col}", unit="region", leave=False):
        cols = windows.index[win_region.astype(str).eq(str(region))].to_numpy()
        if len(cols) == 0:
            continue
        data[str(region)] = _series_from_matrix_mean(X[:, cols], axis=1)

    df = pd.DataFrame(data, index=adata.obs_names.astype(str))
    df.index.name = "cell_id"
    return df.loc[:, _ordered_region_columns(df.columns, "arm" if region_col == "chromosome_arm" else "chromosome")]


def classify_cnv_types(region_df: pd.DataFrame, threshold: float = 0.05) -> pd.DataFrame:
    types = pd.DataFrame("neutral", index=region_df.index, columns=region_df.columns)
    types[region_df > threshold] = "gain"
    types[region_df < -threshold] = "loss"
    types.index.name = region_df.index.name
    return types


def _choose_cancer_col(obs: pd.DataFrame, requested: str | None = None) -> str | None:
    if requested and str(requested).lower() not in {"auto", "none", "null", ""} and requested in obs.columns:
        return requested
    if requested and str(requested).lower() not in {"auto", "none", "null", ""} and requested not in obs.columns:
        warnings.warn(f"Requested cancer_col={requested!r} not in adata.obs. Falling back to auto-detect.")
    candidates = [
        "cancer", "cancer_type", "tumor_type", "dataset", "study", "cohort",
        "Sample", "sample", "patient", "donor", "Patient", "PatientID",
    ]
    for c in candidates:
        if c in obs.columns and obs[c].nunique(dropna=True) > 1:
            return c
    return None


def _choose_groupby(obs: pd.DataFrame, requested: str | None = None) -> str | None:
    if requested and str(requested).lower() not in {"auto", "none", "null", ""} and requested in obs.columns:
        return requested
    if requested and str(requested).lower() not in {"auto", "none", "null", ""} and requested not in obs.columns:
        warnings.warn(f"Requested groupby={requested!r} not in adata.obs. Falling back to auto-detect.")
    candidates = [
        "cnv_leiden", "cnv_status", "malignant_call", "malignant",
        "tumor_call", "clone", "subclone", "celltype", "cell_type", "CellType",
        "major_celltype", "patient", "sample", "Sample",
    ]
    for c in candidates:
        if c in obs.columns and 1 < obs[c].nunique(dropna=True) <= 200:
            return c
    return None


def make_obs_table(adata, cancer_col: str | None, groupby: str | None) -> pd.DataFrame:
    cols = []
    candidates = [cancer_col, groupby, "cnv_score", "malignant_call", "cnv_status", "celltype", "cell_type", "CellType", "Sample", "sample", "patient"]
    for c in candidates:
        if c is not None and c in adata.obs.columns and c not in cols:
            cols.append(c)
    obs = adata.obs.loc[:, cols].copy() if cols else pd.DataFrame(index=adata.obs_names)
    obs.index = obs.index.astype(str)
    obs.index.name = "cell_id"
    return obs


def compute_group_tables(
    region_df: pd.DataFrame,
    type_df: pd.DataFrame,
    obs: pd.DataFrame,
    *,
    cancer_col: str | None,
    groupby: str | None,
    region_name: str,
) -> tuple[pd.DataFrame, pd.DataFrame, pd.DataFrame]:
    tmp = obs.copy()
    tmp["_cell_id"] = tmp.index

    if cancer_col is None or cancer_col not in tmp.columns:
        tmp["_cancer"] = "all"
        cancer_col_eff = "_cancer"
    else:
        cancer_col_eff = cancer_col

    if groupby is None or groupby not in tmp.columns:
        tmp["_group"] = "all"
        groupby_eff = "_group"
    else:
        groupby_eff = groupby

    joined = tmp.join(region_df, how="inner")
    group_mean = (
        joined.groupby([cancer_col_eff, groupby_eff], observed=False)[region_df.columns]
        .mean()
        .reset_index()
        .rename(columns={cancer_col_eff: "cancer", groupby_eff: "group"})
    )

    type_long = (
        type_df.reset_index()
        .melt(id_vars="cell_id", var_name=region_name, value_name="cnv_type")
        .merge(
            tmp.reset_index().rename(columns={cancer_col_eff: "cancer", groupby_eff: "group"}),
            on="cell_id",
            how="left",
        )
    )

    type_fraction = (
        type_long
        .groupby(["cancer", "group", region_name, "cnv_type"], observed=False)
        .size()
        .rename("n_cells")
        .reset_index()
    )
    denom = (
        type_long
        .groupby(["cancer", "group", region_name], observed=False)
        .size()
        .rename("n_total")
        .reset_index()
    )
    type_fraction = type_fraction.merge(denom, on=["cancer", "group", region_name], how="left")
    type_fraction["fraction"] = type_fraction["n_cells"] / type_fraction["n_total"].replace(0, np.nan)

    cancer_fraction = (
        type_long
        .groupby(["cancer", region_name, "cnv_type"], observed=False)
        .size()
        .rename("n_cells")
        .reset_index()
    )
    denom2 = (
        type_long
        .groupby(["cancer", region_name], observed=False)
        .size()
        .rename("n_total")
        .reset_index()
    )
    cancer_fraction = cancer_fraction.merge(denom2, on=["cancer", region_name], how="left")
    cancer_fraction["fraction"] = cancer_fraction["n_cells"] / cancer_fraction["n_total"].replace(0, np.nan)

    return group_mean, type_fraction, cancer_fraction


# -----------------------------------------------------------------------------
# Plotting
# -----------------------------------------------------------------------------


def _infer_region_level_from_columns(cols: Sequence[str]) -> str:
    if any(re.fullmatch(r"chr(?:\d+|X|Y)[pq]", str(c)) for c in cols):
        return "arm"
    return "chromosome"


def _reorder_matrix_columns(matrix: pd.DataFrame, region_level: str | None = None) -> pd.DataFrame:
    if region_level is None:
        region_level = _infer_region_level_from_columns(list(matrix.columns))
    cols = _ordered_region_columns(matrix.columns, "arm" if region_level == "arm" else "chromosome")
    cols = [c for c in cols if c in matrix.columns]
    return matrix.loc[:, cols]


def _plot_matrix_clustermap(
    matrix: pd.DataFrame,
    path_base: Path,
    *,
    title: str,
    center: float | None = 0.0,
    cmap: str = "vlag",
    vlim: float | None = None,
    figsize_scale: tuple[float, float] = (0.45, 0.22),
    min_figsize: tuple[float, float] = (8, 6),
    max_figsize: tuple[float, float] = (26, 30),
    row_cluster: bool | None = None,
    col_cluster: bool = False,
    row_colors=None,
    cbar_label: str = "Mean inferCNV signal",
    region_level: str | None = None,
    dpi: int = 220,
) -> bool:
    if matrix.empty:
        return False

    matrix = matrix.copy()
    matrix = matrix.dropna(axis=0, how="all").dropna(axis=1, how="all")
    if matrix.empty:
        return False

    matrix = _reorder_matrix_columns(matrix, region_level=region_level)
    matrix = matrix.astype(float).replace([np.inf, -np.inf], np.nan).fillna(0.0)

    if vlim is None and center == 0:
        vals = matrix.to_numpy().ravel()
        vals = vals[np.isfinite(vals)]
        vlim = float(np.nanquantile(np.abs(vals), 0.99)) if len(vals) else 1.0
        vlim = max(vlim, 1e-6)

    if row_cluster is None:
        row_cluster = matrix.shape[0] >= 2
    row_cluster = bool(row_cluster and matrix.shape[0] >= 2)
    col_cluster = bool(col_cluster and matrix.shape[1] >= 2)  # intentionally false by default

    width = min(max(min_figsize[0], matrix.shape[1] * figsize_scale[0]), max_figsize[0])
    height = min(max(min_figsize[1], matrix.shape[0] * figsize_scale[1]), max_figsize[1])

    g = sns.clustermap(
        matrix,
        cmap=cmap,
        center=center,
        vmin=-vlim if center == 0 and vlim is not None else (0 if center is None and cmap == "viridis" else None),
        vmax=vlim if center == 0 and vlim is not None else None,
        row_cluster=row_cluster,
        col_cluster=col_cluster,
        row_colors=row_colors,
        xticklabels=True,
        yticklabels=matrix.shape[0] <= 80,
        linewidths=0.0,
        figsize=(width, height),
        cbar_kws={"label": cbar_label},
    )
    g.fig.suptitle(title, y=1.02, fontsize=14)
    _save_clustermap(g, path_base, dpi=dpi)
    return True


def _plot_type_fraction_heatmap(
    type_fraction: pd.DataFrame,
    path_base: Path,
    *,
    title: str,
    region_name: str,
    region_level: str,
    dpi: int = 220,
) -> bool:
    if type_fraction.empty:
        return False

    p = type_fraction.pivot_table(
        index=region_name,
        columns="cnv_type",
        values="fraction",
        aggfunc="mean",
        fill_value=0.0,
    )
    for c in ["loss", "neutral", "gain"]:
        if c not in p.columns:
            p[c] = 0.0
    p = p.loc[_ordered_region_columns(p.index, region_level), ["loss", "neutral", "gain"]]

    h = max(6, min(26, 0.28 * p.shape[0]))
    plt.figure(figsize=(8, h))
    sns.heatmap(
        p,
        vmin=0,
        vmax=1,
        cmap="viridis",
        annot=p.shape[0] <= 60,
        fmt=".2f",
        cbar_kws={"label": "Fraction of cells"},
    )
    plt.title(title)
    plt.xlabel("CNV type")
    plt.ylabel("Chromosome arm" if region_level == "arm" else "Chromosome")
    plt.tight_layout()
    _save_current_fig(path_base, dpi=dpi)
    return True


def _sample_cells_for_plot(df: pd.DataFrame, obs: pd.DataFrame, max_cells: int, stratify_col: str | None = None, seed: int = 13) -> pd.DataFrame:
    if df.shape[0] <= max_cells:
        return df
    rng = np.random.default_rng(seed)
    if stratify_col and stratify_col in obs.columns:
        keep = []
        obs_aligned = obs.reindex(df.index)
        groups = obs_aligned[stratify_col].astype(str).fillna("NA")
        per_group = max(10, int(math.ceil(max_cells / max(1, groups.nunique()))))
        for _, idx in groups.groupby(groups).groups.items():
            idx = np.array(list(idx))
            n = min(len(idx), per_group)
            keep.extend(rng.choice(idx, size=n, replace=False).tolist())
        if len(keep) > max_cells:
            keep = rng.choice(np.array(keep), size=max_cells, replace=False).tolist()
        return df.loc[keep]
    keep = rng.choice(df.index.to_numpy(), size=max_cells, replace=False)
    return df.loc[keep]


def _make_region_output_prefix(region_level: str) -> str:
    return "arm" if region_level == "arm" else "chromosome"


def make_region_plots(
    adata,
    region_df: pd.DataFrame,
    type_df: pd.DataFrame,
    group_mean: pd.DataFrame,
    type_fraction: pd.DataFrame,
    cancer_fraction: pd.DataFrame,
    obs: pd.DataFrame,
    *,
    outfig: Path,
    cancer_col: str | None,
    groupby: str | None,
    region_level: str,
    region_name: str,
    max_cells_plot: int = 3000,
    dpi: int = 220,
) -> None:
    outfig.mkdir(parents=True, exist_ok=True)
    prefix = _make_region_output_prefix(region_level)
    pretty_region = "chromosome arms" if region_level == "arm" else "chromosomes"

    if cancer_col and cancer_col in obs.columns:
        cancers = list(pd.Series(obs[cancer_col]).dropna().astype(str).unique())
    else:
        cancers = ["all"]

    # Across-cancer summary when a single AnnData contains multiple cancer labels.
    if cancer_col and cancer_col in obs.columns and obs[cancer_col].nunique(dropna=True) >= 2:
        joined = obs[[cancer_col]].join(region_df, how="inner")
        cancer_mean = joined.groupby(cancer_col, observed=False)[region_df.columns].mean()
        cancer_mean = _reorder_matrix_columns(cancer_mean, region_level=region_level)
        _plot_matrix_clustermap(
            cancer_mean,
            outfig / f"all_cancers_{prefix}_mean_clustermap",
            title=f"All cancers: mean {prefix}-level inferCNV signal",
            cmap="vlag",
            center=0,
            dpi=dpi,
            figsize_scale=(0.35 if region_level == "arm" else 0.55, 0.55),
            min_figsize=(12, 7),
            col_cluster=False,
            region_level=region_level,
        )

    for cancer in cancers:
        if cancer == "all":
            idx = region_df.index
        else:
            idx = obs.index[obs[cancer_col].astype(str).eq(str(cancer))]
            idx = region_df.index.intersection(idx)
        if len(idx) == 0:
            continue

        safe_cancer = _safe_name(cancer)
        cfig = outfig / safe_cancer / prefix
        cfig.mkdir(parents=True, exist_ok=True)

        c_region = region_df.loc[idx]
        c_type = type_df.loc[idx]
        c_obs = obs.reindex(idx)

        c_plot = _sample_cells_for_plot(c_region, c_obs, max_cells=max_cells_plot, stratify_col=groupby)
        _plot_matrix_clustermap(
            c_plot,
            cfig / f"cells_x_{prefix}s_mean_cnv_clustermap",
            title=f"{cancer}: cells × {pretty_region} mean inferCNV signal",
            cmap="vlag",
            center=0,
            dpi=dpi,
            figsize_scale=(0.30 if region_level == "arm" else 0.50, 0.008 if c_plot.shape[0] > 500 else 0.08),
            min_figsize=(12 if region_level == "arm" else 10, 8),
            max_figsize=(28, 28),
            col_cluster=False,
            region_level=region_level,
        )

        gm = group_mean.loc[group_mean["cancer"].astype(str).eq(str(cancer))].copy()
        if not gm.empty:
            mat = gm.set_index("group")[region_df.columns]
            mat = _reorder_matrix_columns(mat, region_level=region_level)
            _plot_matrix_clustermap(
                mat,
                cfig / f"groups_x_{prefix}s_mean_cnv_clustermap",
                title=f"{cancer}: CNV groups × {pretty_region} mean inferCNV signal",
                cmap="vlag",
                center=0,
                dpi=dpi,
                figsize_scale=(0.30 if region_level == "arm" else 0.55, 0.45),
                min_figsize=(12 if region_level == "arm" else 10, 6),
                max_figsize=(28, 20),
                col_cluster=False,
                region_level=region_level,
            )

        tf = type_fraction.loc[type_fraction["cancer"].astype(str).eq(str(cancer))].copy()
        if not tf.empty:
            for cnv_type in ["gain", "loss"]:
                sub = tf.loc[tf["cnv_type"].eq(cnv_type)]
                if sub.empty:
                    continue
                mat = sub.pivot_table(
                    index="group",
                    columns=region_name,
                    values="fraction",
                    aggfunc="mean",
                    fill_value=0.0,
                )
                mat = _reorder_matrix_columns(mat, region_level=region_level)
                _plot_matrix_clustermap(
                    mat,
                    cfig / f"groups_x_{prefix}s_{cnv_type}_fraction_clustermap",
                    title=f"{cancer}: fraction of cells with {prefix} {cnv_type}",
                    cmap="viridis",
                    center=None,
                    dpi=dpi,
                    figsize_scale=(0.30 if region_level == "arm" else 0.55, 0.45),
                    min_figsize=(12 if region_level == "arm" else 10, 6),
                    max_figsize=(28, 20),
                    row_cluster=mat.shape[0] >= 2,
                    col_cluster=False,
                    cbar_label="Fraction of cells",
                    region_level=region_level,
                )

        cf = cancer_fraction.loc[cancer_fraction["cancer"].astype(str).eq(str(cancer))].copy()
        _plot_type_fraction_heatmap(
            cf,
            cfig / f"{prefix}s_x_cnv_type_fraction_heatmap",
            title=f"{cancer}: {prefix}-level CNV type fractions",
            region_name=region_name,
            region_level=region_level,
            dpi=dpi,
        )


def _make_batch_summary_from_exported_tables(
    *,
    batch_outdir: Path,
    successful: Sequence[str],
    region_level: str,
    dpi: int = 220,
) -> None:
    prefix = _make_region_output_prefix(region_level)
    region_name = "chromosome_arm" if region_level == "arm" else "chromosome"
    summary_dir = batch_outdir / "BATCH_SUMMARY" / prefix
    summary_tables = summary_dir / "tables"
    summary_figures = summary_dir / "figures"
    summary_tables.mkdir(parents=True, exist_ok=True)
    summary_figures.mkdir(parents=True, exist_ok=True)

    mean_rows = []
    type_records = []

    for cancer in successful:
        tables = batch_outdir / _safe_name(cancer) / "tables" / prefix
        mean_path = tables / f"per_cell_{prefix}_mean_cnv.csv.gz"
        type_path = tables / f"per_cell_{prefix}_cnv_type.csv.gz"

        if mean_path.exists():
            m = pd.read_csv(mean_path, index_col=0)
            row = m.mean(axis=0, numeric_only=True)
            row.name = cancer
            mean_rows.append(row)

        if type_path.exists():
            t = pd.read_csv(type_path, index_col=0)
            for region in t.columns:
                vc = t[region].astype(str).value_counts(normalize=True)
                for cnv_type in ["loss", "neutral", "gain"]:
                    type_records.append({
                        "cancer": cancer,
                        region_name: region,
                        "cnv_type": cnv_type,
                        "fraction": float(vc.get(cnv_type, 0.0)),
                    })

    if not mean_rows:
        warnings.warn(f"No exported per-cell {prefix} mean tables found for batch summary.")
        return

    cancer_mean = pd.DataFrame(mean_rows).fillna(0.0)
    cancer_mean = _reorder_matrix_columns(cancer_mean, region_level=region_level)
    cancer_mean.index.name = "cancer"
    cancer_mean.to_csv(summary_tables / f"batch_cancer_{prefix}_mean_cnv.csv")

    _plot_matrix_clustermap(
        cancer_mean,
        summary_figures / f"batch_all_cancers_{prefix}_mean_clustermap",
        title=f"All cancers: mean {prefix}-level inferCNV signal",
        cmap="vlag",
        center=0,
        dpi=dpi,
        figsize_scale=(0.30 if region_level == "arm" else 0.55, 0.55),
        min_figsize=(12 if region_level == "arm" else 10, 7),
        col_cluster=False,
        region_level=region_level,
    )

    type_fraction = pd.DataFrame(type_records)
    if not type_fraction.empty:
        type_fraction.to_csv(summary_tables / f"batch_cancer_{prefix}_cnv_type_fraction.csv", index=False)

        p = type_fraction.pivot_table(
            index=["cancer", region_name],
            columns="cnv_type",
            values="fraction",
            fill_value=0.0,
        ).reset_index()
        for c in ["gain", "loss"]:
            if c not in p.columns:
                p[c] = 0.0
        p["signed_gain_minus_loss_fraction"] = p["gain"] - p["loss"]

        signed = p.pivot_table(
            index="cancer",
            columns=region_name,
            values="signed_gain_minus_loss_fraction",
            fill_value=0.0,
        )
        signed = _reorder_matrix_columns(signed, region_level=region_level)
        _plot_matrix_clustermap(
            signed,
            summary_figures / f"batch_all_cancers_{prefix}_signed_gain_minus_loss_fraction_clustermap",
            title=f"All cancers: signed gain-minus-loss fraction by {prefix}",
            cmap="vlag",
            center=0,
            vlim=1.0,
            dpi=dpi,
            figsize_scale=(0.30 if region_level == "arm" else 0.55, 0.55),
            min_figsize=(12 if region_level == "arm" else 10, 7),
            col_cluster=False,
            region_level=region_level,
        )

        for cnv_type in ["gain", "loss"]:
            mat = type_fraction.loc[type_fraction["cnv_type"].eq(cnv_type)].pivot_table(
                index="cancer",
                columns=region_name,
                values="fraction",
                fill_value=0.0,
            )
            if not mat.empty:
                mat = _reorder_matrix_columns(mat, region_level=region_level)
                _plot_matrix_clustermap(
                    mat,
                    summary_figures / f"batch_all_cancers_{prefix}_{cnv_type}_fraction_clustermap",
                    title=f"All cancers: {prefix} {cnv_type} fraction",
                    cmap="viridis",
                    center=None,
                    dpi=dpi,
                    figsize_scale=(0.30 if region_level == "arm" else 0.55, 0.55),
                    min_figsize=(12 if region_level == "arm" else 10, 7),
                    col_cluster=False,
                    cbar_label="Fraction of cells",
                    region_level=region_level,
                )

    _print(f"[batch-summary:{prefix}] tables:  {summary_tables}")
    _print(f"[batch-summary:{prefix}] figures: {summary_figures}")


# -----------------------------------------------------------------------------
# Export function
# -----------------------------------------------------------------------------


def export_cnv_arm_chromosome_clustermaps(
    adata,
    *,
    outdir: str | Path,
    cancer_col: str | None = None,
    groupby: str | None = None,
    use_rep: str = "cnv",
    window_size: int = 100,
    step: int = 10,
    cnv_type_threshold: float = 0.05,
    region_level: str = "both",
    genome_build: str = "hg38",
    centromere_csv: str | Path | None = None,
    centromere_windows: str = "nearest",
    exclude_chromosomes: Sequence[str] | None = ("chrX", "chrY"),
    max_cells_plot: int = 3000,
    dpi: int = 220,
    progress: bool = True,
) -> dict[str, str]:
    if region_level not in {"chromosome", "arm", "both"}:
        raise ValueError("region_level must be chromosome, arm, or both")

    outdir = Path(outdir)
    outtab = outdir / "tables"
    outfig = outdir / "figures"
    outtab.mkdir(parents=True, exist_ok=True)
    outfig.mkdir(parents=True, exist_ok=True)

    levels = ["chromosome", "arm"] if region_level == "both" else [region_level]
    n_steps = 4 + 2 * len(levels)
    step_bar = tqdm(total=n_steps, desc="export steps", unit="step", disable=not progress)

    outputs: dict[str, str] = {"outdir": str(outdir), "tables": str(outtab), "figures": str(outfig)}

    try:
        with _timed_step("detect metadata columns"):
            cancer_col = _choose_cancer_col(adata.obs, cancer_col)
            groupby = _choose_groupby(adata.obs, groupby)
            _print(f"[cnv-export] outdir: {outdir}")
            _print(f"[cnv-export] use_rep: {use_rep}")
            _print(f"[cnv-export] cancer_col: {cancer_col}")
            _print(f"[cnv-export] groupby: {groupby}")
            _print(f"[cnv-export] region_level={region_level}; genome_build={genome_build}; threshold={cnv_type_threshold}")
            _print(f"[cnv-export] adata: cells={adata.n_obs:,}, genes={adata.n_vars:,}")
        step_bar.update(1)

        with _timed_step("reconstruct inferCNV windows and assign chromosome arms"):
            windows = reconstruct_cnv_windows(
                adata,
                use_rep=use_rep,
                window_size=window_size,
                step=step,
                exclude_chromosomes=exclude_chromosomes,
            )
            windows = add_chromosome_arm_annotations(
                windows,
                genome_build=genome_build,
                centromere_csv=centromere_csv,
                centromere_windows=centromere_windows,
            )
            windows_path = outtab / "cnv_windows_with_arms.csv.gz"
            _write_df(windows, windows_path)
            outputs["windows"] = str(windows_path)
            _print(
                f"[cnv-export] windows={len(windows):,}; "
                f"chromosomes={windows['chromosome'].nunique():,}; "
                f"arms={windows['chromosome_arm'].dropna().nunique():,}"
            )
        step_bar.update(1)

        with _timed_step("make obs metadata table"):
            obs = make_obs_table(adata, cancer_col, groupby)
            obs_path = outtab / "obs_metadata_used.csv.gz"
            _write_df(obs, obs_path)
            outputs["obs_metadata_used"] = str(obs_path)
        step_bar.update(1)

        metadata = {
            "use_rep": use_rep,
            "obsm_key": _resolve_obsm_key(adata, use_rep),
            "uns_key": use_rep,
            "n_cells": int(adata.n_obs),
            "n_genes": int(adata.n_vars),
            "n_cnv_windows": int(get_cnv_matrix(adata, use_rep=use_rep).shape[1]),
            "cancer_col": cancer_col,
            "groupby": groupby,
            "window_size": int(window_size),
            "step": int(step),
            "cnv_type_threshold": float(cnv_type_threshold),
            "region_level": region_level,
            "genome_build": genome_build,
            "centromere_csv": str(centromere_csv) if centromere_csv else None,
            "centromere_windows": centromere_windows,
            "exclude_chromosomes": list(exclude_chromosomes) if exclude_chromosomes is not None else None,
            "max_cells_plot": int(max_cells_plot),
            "column_clustering": False,
        }

        for level in levels:
            prefix = _make_region_output_prefix(level)
            region_col = "chromosome_arm" if level == "arm" else "chromosome"
            region_name = "chromosome_arm" if level == "arm" else "chromosome"
            level_tab = outtab / prefix
            level_tab.mkdir(parents=True, exist_ok=True)

            with _timed_step(f"compute {prefix}-level CNV tables"):
                region_df = compute_cell_region_means(adata, windows, use_rep=use_rep, region_col=region_col)
                type_df = classify_cnv_types(region_df, threshold=cnv_type_threshold)
                group_mean, type_fraction, cancer_fraction = compute_group_tables(
                    region_df,
                    type_df,
                    obs,
                    cancer_col=cancer_col,
                    groupby=groupby,
                    region_name=region_name,
                )
                _print(f"[cnv-export:{prefix}] per-cell matrix: {region_df.shape[0]:,} cells × {region_df.shape[1]:,} {prefix}s")
                _print(f"[cnv-export:{prefix}] group mean rows={len(group_mean):,}; type fraction rows={len(type_fraction):,}")
            step_bar.update(1)

            with _timed_step(f"write {prefix}-level tables"):
                paths = {
                    f"per_cell_{prefix}_mean_cnv": level_tab / f"per_cell_{prefix}_mean_cnv.csv.gz",
                    f"per_cell_{prefix}_cnv_type": level_tab / f"per_cell_{prefix}_cnv_type.csv.gz",
                    f"per_group_{prefix}_mean_cnv": level_tab / f"per_group_{prefix}_mean_cnv.csv.gz",
                    f"per_group_{prefix}_cnv_type_fraction": level_tab / f"per_group_{prefix}_cnv_type_fraction.csv.gz",
                    f"per_cancer_{prefix}_cnv_type_fraction": level_tab / f"per_cancer_{prefix}_cnv_type_fraction.csv.gz",
                }
                table_items = [
                    ("per-cell means", region_df, paths[f"per_cell_{prefix}_mean_cnv"]),
                    ("per-cell types", type_df, paths[f"per_cell_{prefix}_cnv_type"]),
                    ("per-group means", group_mean, paths[f"per_group_{prefix}_mean_cnv"]),
                    ("per-group type fractions", type_fraction, paths[f"per_group_{prefix}_cnv_type_fraction"]),
                    ("per-cancer type fractions", cancer_fraction, paths[f"per_cancer_{prefix}_cnv_type_fraction"]),
                ]
                for label, df, path in tqdm(table_items, desc=f"write {prefix} tables", unit="table", disable=not progress, leave=False):
                    _write_df(df, path)
                    _print(f"[write:{prefix}] {label}: {path}")
                outputs.update({k: str(v) for k, v in paths.items()})
            step_bar.update(1)

            with _timed_step(f"make {prefix}-level figures"):
                with _heartbeat(f"{prefix}-level figure generation", interval=45):
                    make_region_plots(
                        adata,
                        region_df,
                        type_df,
                        group_mean,
                        type_fraction,
                        cancer_fraction,
                        obs,
                        outfig=outfig,
                        cancer_col=cancer_col,
                        groupby=groupby,
                        region_level=level,
                        region_name=region_name,
                        max_cells_plot=max_cells_plot,
                        dpi=dpi,
                    )
            # Not counted separately because n_steps expects compute/write per level plus initial/final.
            # But we want visible movement; update if room remains.
            if step_bar.n < n_steps - 1:
                step_bar.update(1)

            del region_df, type_df, group_mean, type_fraction, cancer_fraction
            gc.collect()

        with _timed_step("write export metadata"):
            metadata_path = outtab / "export_metadata.json"
            metadata_path.write_text(json.dumps(metadata, indent=2), encoding="utf-8")
            outputs["metadata"] = str(metadata_path)
            _print(f"[write] metadata: {metadata_path}")
            _print("[cnv-export] done.")
            _print(f"[cnv-export] tables:  {outtab}")
            _print(f"[cnv-export] figures: {outfig}")
        if step_bar.n < n_steps:
            step_bar.update(n_steps - step_bar.n)

    finally:
        step_bar.close()

    return outputs


# -----------------------------------------------------------------------------
# Optional automatic CNV Leiden clustering
# -----------------------------------------------------------------------------


def _compute_cnv_reduced_embedding(
    adata,
    *,
    obsm_key: str,
    out_key: str = "X_cnv_svd",
    n_components: int = 30,
    random_state: int = 0,
) -> str:
    if out_key in adata.obsm:
        _print(f"[cluster] using existing adata.obsm[{out_key!r}] shape={adata.obsm[out_key].shape}")
        return out_key
    try:
        from sklearn.decomposition import TruncatedSVD
    except Exception as e:
        raise RuntimeError("Automatic CNV Leiden clustering requires scikit-learn") from e

    X = adata.obsm[obsm_key]
    n_obs, n_features = int(X.shape[0]), int(X.shape[1])
    n_components = int(max(2, min(n_components, n_obs - 1, n_features - 1)))
    _print(f"[cluster] reducing {obsm_key}: {n_obs:,} cells × {n_features:,} windows -> {n_components} components")
    with _heartbeat(f"SVD reduction of {obsm_key}", interval=30):
        if sp is not None and sp.issparse(X):
            emb = TruncatedSVD(n_components=n_components, random_state=random_state).fit_transform(X)
        else:
            X_arr = np.asarray(X, dtype=np.float32)
            emb = TruncatedSVD(n_components=n_components, random_state=random_state).fit_transform(X_arr)
    adata.obsm[out_key] = emb.astype(np.float32, copy=False)
    return out_key


def _ensure_groupby_column(
    adata,
    *,
    requested_groupby: str | None,
    make_cnv_leiden: bool,
    use_rep: str,
    leiden_resolution: float,
    neighbors_n: int,
    cnv_leiden_rep: str = "svd",
    cnv_svd_components: int = 30,
    progress: bool = True,
) -> str | None:
    if requested_groupby and requested_groupby.lower() not in {"auto", "none", "null"}:
        if requested_groupby in adata.obs.columns:
            _print(f"[groupby] using requested obs column: {requested_groupby}")
            return requested_groupby
        warnings.warn(f"Requested --groupby {requested_groupby!r} is not in adata.obs; falling back to auto.")

    if "cnv_leiden" in adata.obs.columns:
        _print("[groupby] using existing obs column: cnv_leiden")
        return "cnv_leiden"

    if make_cnv_leiden:
        try:
            import scanpy as sc
            obsm_key = _resolve_obsm_key(adata, use_rep)
            with tqdm(total=3, desc="CNV Leiden", unit="step", disable=not progress) as pbar:
                if cnv_leiden_rep.lower() in {"svd", "pca", "reduced"}:
                    with _timed_step("CNV dimensionality reduction"):
                        cluster_rep = _compute_cnv_reduced_embedding(
                            adata,
                            obsm_key=obsm_key,
                            out_key=f"X_{use_rep}_svd",
                            n_components=cnv_svd_components,
                        )
                else:
                    cluster_rep = obsm_key
                    _print(f"[cluster] using raw CNV windows for neighbors: {cluster_rep}")
                pbar.update(1)

                with _timed_step(f"build kNN graph on {cluster_rep}"):
                    with _heartbeat("scanpy.pp.neighbors", interval=30):
                        sc.pp.neighbors(adata, use_rep=cluster_rep, n_neighbors=neighbors_n, random_state=0)
                pbar.update(1)

                with _timed_step("run Leiden clustering"):
                    with _heartbeat("scanpy.tl.leiden", interval=30):
                        try:
                            sc.tl.leiden(
                                adata,
                                resolution=leiden_resolution,
                                key_added="cnv_leiden",
                                random_state=0,
                                flavor="igraph",
                                n_iterations=2,
                                directed=False,
                            )
                        except TypeError:
                            sc.tl.leiden(adata, resolution=leiden_resolution, key_added="cnv_leiden", random_state=0)
                pbar.update(1)
            counts = adata.obs["cnv_leiden"].astype(str).value_counts().to_dict()
            _print("[cluster] cnv_leiden counts: " + json.dumps(counts, indent=None))
            return "cnv_leiden"
        except Exception as e:
            warnings.warn(f"Could not create cnv_leiden automatically: {type(e).__name__}: {e}. Falling back to auto grouping.")

    return None


# -----------------------------------------------------------------------------
# CLI batch/single mode
# -----------------------------------------------------------------------------

DEFAULT_INFERCNV_DIR = Path(
    r"C:\Users\ashfo\OneDrive\Desktop\Pathways+Omics\scOPE\scOPE\data\scope_robust_real_inputs\infercnvpy_new"
)
DEFAULT_DATASETS = ["AML", "BRCA", "CRC", "GBM", "LUAD", "PAAD", "SKCM"]


def _split_csv_arg(x: str | None, default: Sequence[str] | None = None) -> list[str]:
    if x is None or str(x).strip() == "":
        return list(default or [])
    if str(x).strip().lower() in {"all", "*"}:
        return list(default or [])
    return [v.strip() for v in str(x).split(",") if v.strip()]


def _find_dataset_h5ad(h5ad_dir: Path, dataset: str, *, pattern: str = "{dataset}_infercnvpy.h5ad", recursive: bool = False) -> Path | None:
    exact = h5ad_dir / pattern.format(dataset=dataset)
    if exact.exists():
        return exact
    globber = h5ad_dir.rglob if recursive else h5ad_dir.glob
    candidates = list(globber(pattern.format(dataset="*")))
    candidates += list(globber(f"{dataset}*infercnvpy*.h5ad"))
    candidates += list(globber(f"*{dataset}*infercnvpy*.h5ad"))
    unique = []
    seen = set()
    for p in candidates:
        if p in seen:
            continue
        seen.add(p)
        if p.is_file() and p.suffix == ".h5ad":
            unique.append(p)
    if not unique:
        return None
    return sorted(unique, key=lambda p: (len(str(p)), str(p).lower()))[0]


def _discover_auto_datasets(h5ad_dir: Path, recursive: bool = False) -> list[tuple[str, Path]]:
    globber = h5ad_dir.rglob if recursive else h5ad_dir.glob
    files = sorted(globber("*.h5ad"))
    discovered = []
    for p in files:
        stem = p.stem
        dataset = re.sub(r"_?infercnvpy.*$", "", stem, flags=re.IGNORECASE) or stem
        discovered.append((dataset, p))
    return discovered


def _add_dataset_obs_column(adata, dataset: str, cancer_col: str | None) -> None:
    if "dataset" not in adata.obs.columns:
        adata.obs["dataset"] = str(dataset)
    if cancer_col and cancer_col not in {"auto", "None", "none", ""}:
        if cancer_col not in adata.obs.columns:
            adata.obs[cancer_col] = str(dataset)


def _parse_args() -> argparse.Namespace:
    p = argparse.ArgumentParser(
        description=(
            "Export chromosome- and arm-level inferCNVpy tables/plots from one .h5ad "
            "or a directory of per-cancer inferCNVpy .h5ad files."
        )
    )

    mode = p.add_argument_group("Input mode")
    mode.add_argument("--h5ad", default=None, help="Single inferCNVpy AnnData .h5ad file.")
    mode.add_argument("--h5ad-dir", default=str(DEFAULT_INFERCNV_DIR), help="Directory containing per-cancer inferCNVpy h5ads.")
    mode.add_argument("--h5ad-pattern", default="{dataset}_infercnvpy.h5ad", help="Pattern for per-cancer h5ads inside --h5ad-dir.")
    mode.add_argument("--recursive", action="store_true", help="Search --h5ad-dir recursively.")
    mode.add_argument("--datasets", default="AML,BRCA,CRC,GBM,LUAD,PAAD,SKCM", help="Comma-separated datasets, or auto/all.")

    out = p.add_argument_group("Outputs")
    out.add_argument("--outdir", default=None, help="Output directory. Batch mode creates one subfolder per dataset.")
    out.add_argument("--skip-existing", action="store_true", help="Skip dataset when tables/export_metadata.json exists.")
    out.add_argument("--no-batch-summary", action="store_true", help="Do not make BATCH_SUMMARY plots/tables.")

    meta = p.add_argument_group("Metadata/grouping")
    meta.add_argument("--cancer-col", default="dataset", help="adata.obs cancer/dataset column. Use auto for auto-detect.")
    meta.add_argument("--groupby", default="auto", help="obs column for group-level rows; auto prefers cnv_leiden.")
    meta.add_argument("--make-cnv-leiden", action="store_true", help="If cnv_leiden is missing, build Leiden clusters from X_cnv.")
    meta.add_argument("--leiden-resolution", type=float, default=1.0)
    meta.add_argument("--neighbors-n", type=int, default=15)
    meta.add_argument("--cnv-leiden-rep", default="svd", choices=["svd", "raw"])
    meta.add_argument("--cnv-svd-components", type=int, default=30)
    meta.add_argument("--heartbeat-seconds", type=int, default=30)
    meta.add_argument("--quiet-progress", action="store_true", help="Disable tqdm progress bars.")

    cnv_args = p.add_argument_group("CNV/window settings")
    cnv_args.add_argument("--use-rep", default="cnv", help="inferCNVpy key; default expects X_cnv/cnv.")
    cnv_args.add_argument("--window-size", type=int, default=100, help="window_size used in original inferCNVpy run.")
    cnv_args.add_argument("--step", type=int, default=10, help="step used in original inferCNVpy run.")
    cnv_args.add_argument("--threshold", type=float, default=0.05, help="Mean > threshold gain; < -threshold loss.")
    cnv_args.add_argument("--region-level", default="both", choices=["chromosome", "arm", "both"], help="Export chromosome, arm, or both. Default: both.")
    cnv_args.add_argument("--genome-build", default="hg38", choices=["hg38", "GRCh38", "grch38", "hg19", "GRCh37", "grch37", "auto"], help="Genome build for centromere split. Default: hg38.")
    cnv_args.add_argument("--centromere-csv", default=None, help="Optional centromere file overriding built-in coordinates.")
    cnv_args.add_argument("--centromere-windows", default="nearest", choices=["nearest", "cen", "drop"], help="How to handle windows whose midpoint lies inside centromere interval.")
    cnv_args.add_argument("--include-sex-chromosomes", action="store_true", help="Do not exclude chrX/chrY.")

    plot = p.add_argument_group("Plot settings")
    plot.add_argument("--max-cells-plot", type=int, default=3000)
    plot.add_argument("--dpi", type=int, default=220)

    return p.parse_args()


def _run_single_h5ad(args: argparse.Namespace) -> None:
    import scanpy as sc
    h5ad = Path(args.h5ad)
    if not h5ad.exists():
        raise FileNotFoundError(h5ad)
    outdir = Path(args.outdir) if args.outdir else h5ad.with_suffix("").parent / f"{h5ad.stem}_cnv_arm_chromosome_clustermaps"
    exclude = None if args.include_sex_chromosomes else ("chrX", "chrY")
    cancer_col = None if str(args.cancer_col).lower() == "auto" else args.cancer_col
    dataset = h5ad.stem.replace("_infercnvpy", "")

    _print(f"[single] reading: {h5ad}")
    with _timed_step("read single h5ad"):
        adata = sc.read_h5ad(h5ad)
        _print(f"[read] loaded cells={adata.n_obs:,}, genes={adata.n_vars:,}, obsm={list(adata.obsm.keys())}")
    _add_dataset_obs_column(adata, dataset=dataset, cancer_col=cancer_col)

    groupby = _ensure_groupby_column(
        adata,
        requested_groupby=args.groupby,
        make_cnv_leiden=args.make_cnv_leiden,
        use_rep=args.use_rep,
        leiden_resolution=args.leiden_resolution,
        neighbors_n=args.neighbors_n,
        cnv_leiden_rep=args.cnv_leiden_rep,
        cnv_svd_components=args.cnv_svd_components,
        progress=not args.quiet_progress,
    )

    export_cnv_arm_chromosome_clustermaps(
        adata,
        outdir=outdir,
        cancer_col=cancer_col,
        groupby=groupby,
        use_rep=args.use_rep,
        window_size=args.window_size,
        step=args.step,
        cnv_type_threshold=args.threshold,
        region_level=args.region_level,
        genome_build=args.genome_build,
        centromere_csv=args.centromere_csv,
        centromere_windows=args.centromere_windows,
        exclude_chromosomes=exclude,
        max_cells_plot=args.max_cells_plot,
        dpi=args.dpi,
        progress=not args.quiet_progress,
    )


def _run_batch(args: argparse.Namespace) -> None:
    import scanpy as sc
    h5ad_dir = Path(args.h5ad_dir)
    if not h5ad_dir.exists():
        raise FileNotFoundError(h5ad_dir)
    batch_outdir = Path(args.outdir) if args.outdir else h5ad_dir / "cnv_arm_chromosome_clustermaps"
    batch_outdir.mkdir(parents=True, exist_ok=True)
    exclude = None if args.include_sex_chromosomes else ("chrX", "chrY")
    cancer_col = None if str(args.cancer_col).lower() == "auto" else args.cancer_col

    if str(args.datasets).strip().lower() in {"auto", "*", "allh5ad", "all-h5ad"}:
        dataset_paths = _discover_auto_datasets(h5ad_dir, recursive=args.recursive)
    else:
        dataset_paths = []
        for dataset in _split_csv_arg(args.datasets, DEFAULT_DATASETS):
            p = _find_dataset_h5ad(h5ad_dir, dataset, pattern=args.h5ad_pattern, recursive=args.recursive)
            if p is None:
                _print(f"[skip] {dataset}: no matching h5ad found in {h5ad_dir}")
                continue
            dataset_paths.append((dataset, p))

    if not dataset_paths:
        raise SystemExit("No input h5ad files found. Check --h5ad-dir, --datasets, and --h5ad-pattern.")

    _print(f"[batch] h5ad_dir: {h5ad_dir}")
    _print(f"[batch] outdir:   {batch_outdir}")
    _print(f"[batch] datasets: {[d for d, _ in dataset_paths]}")

    successful = []
    failures = []

    dataset_iter = tqdm(dataset_paths, desc="datasets", unit="dataset", disable=args.quiet_progress)
    for dataset, h5ad in dataset_iter:
        try:
            dataset_iter.set_description(f"dataset {dataset}")
        except Exception:
            pass
        safe_dataset = _safe_name(dataset)
        dataset_outdir = batch_outdir / safe_dataset
        metadata_path = dataset_outdir / "tables" / "export_metadata.json"

        if args.skip_existing and metadata_path.exists():
            _print(f"\n[skip] {dataset}: existing export found at {metadata_path}")
            successful.append(dataset)
            continue

        _print("\n" + "=" * 96)
        _print(f"[dataset] {dataset}")
        _print(f"[read] {h5ad}")

        try:
            with _timed_step(f"read h5ad for {dataset}"):
                adata = sc.read_h5ad(h5ad)
                _print(f"[read] loaded cells={adata.n_obs:,}, genes={adata.n_vars:,}, obsm={list(adata.obsm.keys())}")
            _add_dataset_obs_column(adata, dataset=dataset, cancer_col=cancer_col)

            groupby = _ensure_groupby_column(
                adata,
                requested_groupby=args.groupby,
                make_cnv_leiden=args.make_cnv_leiden,
                use_rep=args.use_rep,
                leiden_resolution=args.leiden_resolution,
                neighbors_n=args.neighbors_n,
                cnv_leiden_rep=args.cnv_leiden_rep,
                cnv_svd_components=args.cnv_svd_components,
                progress=not args.quiet_progress,
            )

            export_cnv_arm_chromosome_clustermaps(
                adata,
                outdir=dataset_outdir,
                cancer_col=cancer_col,
                groupby=groupby,
                use_rep=args.use_rep,
                window_size=args.window_size,
                step=args.step,
                cnv_type_threshold=args.threshold,
                region_level=args.region_level,
                genome_build=args.genome_build,
                centromere_csv=args.centromere_csv,
                centromere_windows=args.centromere_windows,
                exclude_chromosomes=exclude,
                max_cells_plot=args.max_cells_plot,
                dpi=args.dpi,
                progress=not args.quiet_progress,
            )
            successful.append(dataset)
            _print(f"[done] {dataset}: {dataset_outdir}")

        except Exception as e:
            failures.append((dataset, type(e).__name__, str(e)))
            _print(f"[ERROR] {dataset}: {type(e).__name__}: {e}")

        finally:
            try:
                del adata
            except Exception:
                pass
            gc.collect()

    if successful and not args.no_batch_summary:
        _print("\n" + "=" * 96)
        _print("[batch-summary] making cross-cancer summary plots")
        levels = ["chromosome", "arm"] if args.region_level == "both" else [args.region_level]
        for level in levels:
            _make_batch_summary_from_exported_tables(batch_outdir=batch_outdir, successful=successful, region_level=level, dpi=args.dpi)

    _print("\n" + "=" * 96)
    _print(f"[batch] successful: {successful}")
    if failures:
        _print("[batch] failures:")
        for dataset, err_type, msg in failures:
            _print(f"  - {dataset}: {err_type}: {msg}")
    _print("[done]")


def main() -> None:
    global HEARTBEAT_SECONDS
    args = _parse_args()
    HEARTBEAT_SECONDS = int(args.heartbeat_seconds)
    if args.h5ad:
        _run_single_h5ad(args)
    else:
        _run_batch(args)


if __name__ == "__main__":
    main()
```
