## Supplemental_Code for "scOPE identifies which driver-associated expression programs transfer from bulk tumors to single cells": export_cnv_chromosome_clustermaps_all_cancers_PROGRESS.py

```
#!/usr/bin/env python
"""
Export inferCNVpy chromosome-level tables and clustermap plots.

Use case
--------
Run this after infercnvpy has already populated an AnnData object with:
    adata.obsm["X_cnv"]
    adata.uns["cnv"]["chr_pos"]

Typical notebook usage:
    from export_cnv_chromosome_clustermaps import export_cnv_chromosome_clustermaps

    export_cnv_chromosome_clustermaps(
        annotated,
        outdir=ctx.figures / "cnv_chromosome_clustermaps",
        cancer_col="dataset",          # or "cancer"; auto-detected if None
        groupby=None,                  # auto: cnv_leiden > malignant_call > cnv_status > celltype
        use_rep="cnv",
        window_size=100,
        step=10,
        cnv_type_threshold=0.05,
        max_cells_plot=3000,
        make_builtin_infercnvpy_heatmap=False,
    )

Command-line usage:
    python export_cnv_chromosome_clustermaps.py \
        --h5ad path/to/infercnv_annotated.h5ad \
        --outdir path/to/cnv_chromosome_clustermaps \
        --cancer-col dataset \
        --window-size 100 \
        --step 10

Outputs
-------
outdir/
  tables/
    cnv_windows.csv.gz
    per_cell_chromosome_mean_cnv.csv.gz
    per_cell_chromosome_cnv_type.csv.gz
    per_group_chromosome_mean_cnv.csv.gz
    per_group_chromosome_cnv_type_fraction.csv.gz
    per_cancer_chromosome_cnv_type_fraction.csv.gz
  figures/
    all_cancers_chromosome_mean_clustermap.png/.pdf
    all_cancers_signed_gain_loss_fraction_clustermap.png/.pdf
    <cancer>/
      cells_x_chromosomes_mean_cnv_clustermap.png/.pdf
      groups_x_chromosomes_mean_cnv_clustermap.png/.pdf
      groups_x_chromosomes_gain_fraction_clustermap.png/.pdf
      groups_x_chromosomes_loss_fraction_clustermap.png/.pdf
      chromosomes_x_cnv_type_fraction_heatmap.png/.pdf
"""

from __future__ import annotations

import argparse
import contextlib
import gc
import json
import math
import os
import re
import sys
import threading
import time
import warnings
from pathlib import Path
from typing import Iterable, Sequence

import numpy as np
import pandas as pd

try:
    import matplotlib
    matplotlib.use("Agg", force=True)
    import matplotlib.pyplot as plt
except Exception as e:  # pragma: no cover
    raise RuntimeError("matplotlib is required for plotting") from e

try:
    import seaborn as sns
except Exception as e:  # pragma: no cover
    raise RuntimeError("seaborn is required for clustermap plotting") from e

try:
    import scipy.sparse as sp
except Exception:  # pragma: no cover
    sp = None


# -----------------------------
# progress / diagnostics helpers
# -----------------------------

try:
    from tqdm.auto import tqdm
except Exception:  # pragma: no cover
    class tqdm:  # minimal fallback if tqdm is unavailable
        def __init__(self, iterable=None, total=None, desc=None, unit=None, **kwargs):
            self.iterable = iterable
            self.total = total
            self.desc = desc or ""
            self.n = 0
            if self.desc:
                print(f"[progress] {self.desc}", flush=True)

        def __iter__(self):
            if self.iterable is None:
                return iter(())
            for x in self.iterable:
                yield x
                self.update(1)

        def update(self, n=1):
            self.n += n
            if self.total:
                print(f"[progress] {self.desc}: {self.n}/{self.total}", flush=True)

        def set_description(self, desc):
            self.desc = desc
            print(f"[progress] {desc}", flush=True)

        def set_postfix(self, *args, **kwargs):
            pass

        def close(self):
            pass

        def __enter__(self):
            return self

        def __exit__(self, exc_type, exc, tb):
            self.close()


def _print(msg: str) -> None:
    print(msg, flush=True)


def _format_seconds(seconds: float) -> str:
    seconds = float(seconds)
    if seconds < 60:
        return f"{seconds:.1f}s"
    minutes = seconds / 60
    if minutes < 60:
        return f"{minutes:.1f}m"
    return f"{minutes / 60:.1f}h"


def _rss_gb() -> float | None:
    try:
        import psutil
        return psutil.Process(os.getpid()).memory_info().rss / (1024 ** 3)
    except Exception:
        return None


def _mem_suffix() -> str:
    rss = _rss_gb()
    return "" if rss is None else f", RSS={rss:.2f} GB"


@contextlib.contextmanager
def _timed_step(label: str):
    t0 = time.time()
    _print(f"[step] START {label}{_mem_suffix()}")
    try:
        yield
    finally:
        _print(f"[step] DONE  {label} in {_format_seconds(time.time() - t0)}{_mem_suffix()}")


HEARTBEAT_SECONDS = 30


@contextlib.contextmanager
def _heartbeat(label: str, interval: int | None = None):
    """
    Print periodic progress during library calls that do not expose native progress bars
    such as scanpy.pp.neighbors or seaborn.clustermap.
    """
    if interval is None:
        interval = HEARTBEAT_SECONDS
    stop_event = threading.Event()
    t0 = time.time()

    def _worker():
        while not stop_event.wait(interval):
            _print(f"[heartbeat] {label}: still running after {_format_seconds(time.time() - t0)}{_mem_suffix()}")

    thread = threading.Thread(target=_worker, daemon=True)
    thread.start()
    try:
        yield
    finally:
        stop_event.set()
        thread.join(timeout=2)


def _matrix_shape_string(x) -> str:
    try:
        return f"{int(x.shape[0]):,} × {int(x.shape[1]):,}"
    except Exception:
        return "unknown shape"


def _resolve_obsm_key(adata, use_rep: str) -> str:
    candidates = [f"X_{use_rep}", use_rep]
    if use_rep == "cnv":
        candidates += ["X_cnv", "cnv"]
    for k in candidates:
        if k in adata.obsm:
            return k
    raise KeyError(f"Could not find CNV matrix for use_rep={use_rep!r}. Available obsm keys: {list(adata.obsm.keys())}")


# -----------------------------
# small utilities
# -----------------------------

def _safe_name(x: object) -> str:
    s = str(x)
    s = re.sub(r"[^\w.\-]+", "_", s).strip("_")
    return s[:120] if s else "NA"


def _natural_key(s: object):
    return [int(t) if t.isdigit() else t.lower() for t in re.split(r"([0-9]+)", str(s))]


def _natural_chromosomes(chroms: Iterable[object]) -> list[str]:
    chroms = [str(c) for c in pd.Series(list(chroms)).dropna().unique()]
    # Match inferCNVpy's default behavior: chromosomes that start with "chr", excluding chrM.
    chroms = [c for c in chroms if c.startswith("chr") and c != "chrM"]
    return sorted(chroms, key=_natural_key)


def _maybe_dense(x):
    if sp is not None and sp.issparse(x):
        return x.toarray()
    if hasattr(x, "A"):
        return np.asarray(x.A)
    return np.asarray(x)


def _series_from_matrix_mean(x, axis: int = 1) -> np.ndarray:
    """Memory-safe mean for dense or sparse matrices."""
    if sp is not None and sp.issparse(x):
        arr = np.asarray(x.mean(axis=axis)).ravel()
    else:
        arr = np.asarray(np.nanmean(x, axis=axis)).ravel()
    return arr


def _write_df(df: pd.DataFrame, path: Path) -> None:
    path.parent.mkdir(parents=True, exist_ok=True)
    if path.suffix == ".gz" or path.name.endswith(".csv.gz"):
        df.to_csv(path, compression="gzip")
    else:
        df.to_csv(path)


def _save_current_fig(path_base: Path, dpi: int = 220) -> None:
    path_base.parent.mkdir(parents=True, exist_ok=True)
    plt.savefig(path_base.with_suffix(".png"), dpi=dpi, bbox_inches="tight")
    plt.savefig(path_base.with_suffix(".pdf"), bbox_inches="tight")
    plt.close()


def _save_clustermap(g, path_base: Path, dpi: int = 220) -> None:
    path_base.parent.mkdir(parents=True, exist_ok=True)
    g.savefig(path_base.with_suffix(".png"), dpi=dpi, bbox_inches="tight")
    g.savefig(path_base.with_suffix(".pdf"), bbox_inches="tight")
    plt.close(g.fig)


def _has_min_shape(df: pd.DataFrame, min_rows: int = 2, min_cols: int = 2) -> bool:
    return df.shape[0] >= min_rows and df.shape[1] >= min_cols


def _cluster_flags(df: pd.DataFrame) -> tuple[bool, bool]:
    """seaborn.clustermap dies on single-row/col matrices; disable clustering as needed."""
    row_cluster = df.shape[0] >= 2
    col_cluster = df.shape[1] >= 2
    return row_cluster, col_cluster


def _choose_cancer_col(obs: pd.DataFrame, requested: str | None = None) -> str | None:
    if requested and requested in obs.columns:
        return requested
    if requested and requested not in obs.columns:
        warnings.warn(f"Requested cancer_col={requested!r} not in adata.obs. Falling back to auto-detect.")
    candidates = [
        "cancer", "cancer_type", "tumor_type", "dataset", "study", "cohort",
        "Sample", "sample", "patient", "donor", "Patient", "PatientID",
    ]
    for c in candidates:
        if c in obs.columns and obs[c].nunique(dropna=True) > 1:
            return c
    return None


def _choose_groupby(obs: pd.DataFrame, requested: str | None = None) -> str | None:
    if requested and requested in obs.columns:
        return requested
    if requested and requested not in obs.columns:
        warnings.warn(f"Requested groupby={requested!r} not in adata.obs. Falling back to auto-detect.")
    candidates = [
        "cnv_leiden", "cnv_status", "malignant_call", "malignant",
        "tumor_call", "clone", "subclone", "celltype", "cell_type", "CellType",
        "major_celltype", "patient", "sample", "Sample",
    ]
    for c in candidates:
        if c in obs.columns and 1 < obs[c].nunique(dropna=True) <= 100:
            return c
    return None


# -----------------------------
# extracting CNV matrix / windows
# -----------------------------

def get_cnv_matrix(adata, use_rep: str = "cnv"):
    obsm_key = f"X_{use_rep}"
    if obsm_key not in adata.obsm:
        available = ", ".join(map(str, adata.obsm.keys()))
        raise KeyError(
            f"Expected adata.obsm[{obsm_key!r}], but did not find it. "
            f"Available adata.obsm keys: {available}"
        )
    return adata.obsm[obsm_key]


def get_chr_pos(adata, use_rep: str = "cnv") -> dict[str, int]:
    if use_rep not in adata.uns:
        raise KeyError(f"Expected adata.uns[{use_rep!r}], but did not find it.")
    uns = adata.uns[use_rep]
    if isinstance(uns, dict) and "chr_pos" in uns:
        chr_pos = dict(uns["chr_pos"])
    elif hasattr(uns, "keys") and "chr_pos" in uns.keys():
        chr_pos = dict(uns["chr_pos"])
    else:
        raise KeyError(
            f"Expected adata.uns[{use_rep!r}]['chr_pos'], but did not find it. "
            f"Available keys: {list(uns.keys()) if hasattr(uns, 'keys') else type(uns)}"
        )
    return {str(k): int(v) for k, v in chr_pos.items()}


def reconstruct_cnv_windows(
    adata,
    *,
    use_rep: str = "cnv",
    window_size: int = 100,
    step: int = 10,
    exclude_chromosomes: Sequence[str] | None = ("chrX", "chrY"),
) -> pd.DataFrame:
    """
    Reconstruct per-column CNV window metadata.

    inferCNVpy stores chromosome start positions, not every window's genomic interval.
    This reconstructs intervals from adata.var["chromosome", "start", "end"] using the
    same broad window logic as inferCNVpy:
      - filter genes without positions
      - drop excluded chromosomes
      - sort genes by chromosome and genomic start
      - compute running windows independently per chromosome
    """
    X_cnv = get_cnv_matrix(adata, use_rep=use_rep)
    n_windows = int(X_cnv.shape[1])
    chr_pos = get_chr_pos(adata, use_rep=use_rep)

    missing = {"chromosome", "start", "end"} - set(adata.var.columns)
    if missing:
        raise ValueError(
            f"adata.var is missing required genomic position columns: {missing}. "
            "You need these to reconstruct chromosome-level windows."
        )

    var = adata.var.loc[:, ["chromosome", "start", "end"]].copy()
    var["gene"] = adata.var_names.astype(str)
    var = var.dropna(subset=["chromosome", "start", "end"]).copy()

    if exclude_chromosomes is not None:
        var = var.loc[~var["chromosome"].astype(str).isin(set(exclude_chromosomes))].copy()

    chromosomes_from_var = _natural_chromosomes(var["chromosome"])
    chromosomes_from_uns = sorted(chr_pos.keys(), key=lambda c: chr_pos[c])
    chromosomes = [c for c in chromosomes_from_uns if c in set(chromosomes_from_var)]

    # If var/uns naming is imperfect, trust uns order and still try to use matching var rows.
    if not chromosomes:
        chromosomes = chromosomes_from_uns

    # Convert chromosome start positions to per-chromosome window counts.
    starts = [chr_pos[c] for c in chromosomes]
    ends = starts[1:] + [n_windows]
    chr_counts = {c: int(e - s) for c, s, e in zip(chromosomes, starts, ends)}

    records = []
    running_window_id = 0

    for chrom in chromosomes:
        n_for_chrom = chr_counts.get(chrom, 0)
        if n_for_chrom <= 0:
            continue

        g = var.loc[var["chromosome"].astype(str).eq(chrom)].copy()
        if g.empty:
            # Last-ditch placeholder: still allow chromosome aggregation, but no genomic interval labels.
            for _ in range(n_for_chrom):
                records.append({
                    "window_id": running_window_id,
                    "chromosome": chrom,
                    "window_in_chromosome": running_window_id - chr_pos[chrom],
                    "start": np.nan,
                    "end": np.nan,
                    "midpoint": np.nan,
                    "first_gene": "",
                    "last_gene": "",
                    "n_genes_in_window": np.nan,
                    "approximate_interval": True,
                })
                running_window_id += 1
            continue

        g["start"] = pd.to_numeric(g["start"], errors="coerce")
        g["end"] = pd.to_numeric(g["end"], errors="coerce")
        g = g.dropna(subset=["start", "end"]).sort_values(["start", "end"])
        genes = g["gene"].to_numpy()
        starts_bp = g["start"].to_numpy()
        ends_bp = g["end"].to_numpy()
        n_genes = len(g)

        # Expected starts from inferCNVpy's running mean logic.
        if n_genes > window_size:
            win_starts = np.arange(0, n_genes - window_size + 1, step, dtype=int)
        else:
            win_starts = np.array([0], dtype=int)

        # Robust correction if the user passes a different window_size/step than was actually used.
        if len(win_starts) != n_for_chrom:
            max_start = max(0, n_genes - min(window_size, n_genes))
            if n_for_chrom == 1:
                win_starts = np.array([0], dtype=int)
            else:
                win_starts = np.round(np.linspace(0, max_start, n_for_chrom)).astype(int)
            warnings.warn(
                f"{chrom}: reconstructed {len(win_starts)} windows to match X_cnv, "
                f"but this did not match the nominal window_size={window_size}, step={step}. "
                "Check that these match the infercnvpy parameters originally used."
            )

        for j, sidx in enumerate(win_starts):
            eidx = min(sidx + window_size - 1, n_genes - 1)
            start_bp = float(starts_bp[sidx])
            end_bp = float(ends_bp[eidx])
            records.append({
                "window_id": running_window_id,
                "chromosome": chrom,
                "window_in_chromosome": j,
                "start": start_bp,
                "end": end_bp,
                "midpoint": (start_bp + end_bp) / 2.0,
                "first_gene": str(genes[sidx]),
                "last_gene": str(genes[eidx]),
                "n_genes_in_window": int(eidx - sidx + 1),
                "approximate_interval": len(win_starts) != n_for_chrom,
            })
            running_window_id += 1

    windows = pd.DataFrame.from_records(records)
    if len(windows) != n_windows:
        raise RuntimeError(
            f"Window metadata length ({len(windows)}) does not match X_cnv columns ({n_windows}). "
            "Most likely window_size/step/excluded chromosomes do not match the original infercnvpy run."
        )

    return windows


# -----------------------------
# chromosome-level tables
# -----------------------------

def compute_cell_chromosome_means(
    adata,
    windows: pd.DataFrame,
    *,
    use_rep: str = "cnv",
) -> pd.DataFrame:
    X = get_cnv_matrix(adata, use_rep=use_rep)
    chromosomes = list(windows["chromosome"].drop_duplicates())
    data = {}

    for chrom in chromosomes:
        cols = windows.index[windows["chromosome"].astype(str).eq(str(chrom))].to_numpy()
        if len(cols) == 0:
            continue
        data[str(chrom)] = _series_from_matrix_mean(X[:, cols], axis=1)

    chrom_df = pd.DataFrame(data, index=adata.obs_names.astype(str))
    chrom_df.index.name = "cell_id"
    return chrom_df


def classify_cnv_types(chrom_df: pd.DataFrame, threshold: float = 0.05) -> pd.DataFrame:
    types = pd.DataFrame("neutral", index=chrom_df.index, columns=chrom_df.columns)
    types[chrom_df > threshold] = "gain"
    types[chrom_df < -threshold] = "loss"
    types.index.name = chrom_df.index.name
    return types


def make_obs_table(adata, cancer_col: str | None, groupby: str | None) -> pd.DataFrame:
    cols = []
    for c in [cancer_col, groupby, "cnv_score", "malignant_call", "cnv_status", "celltype", "cell_type", "Sample", "sample", "patient"]:
        if c is not None and c in adata.obs.columns and c not in cols:
            cols.append(c)
    obs = adata.obs.loc[:, cols].copy() if cols else pd.DataFrame(index=adata.obs_names)
    obs.index = obs.index.astype(str)
    obs.index.name = "cell_id"
    return obs


def compute_group_tables(
    chrom_df: pd.DataFrame,
    type_df: pd.DataFrame,
    obs: pd.DataFrame,
    *,
    cancer_col: str | None,
    groupby: str | None,
) -> tuple[pd.DataFrame, pd.DataFrame, pd.DataFrame]:
    tmp = obs.copy()
    tmp["_cell_id"] = tmp.index

    if cancer_col is None:
        tmp["_cancer"] = "all"
        cancer_col_eff = "_cancer"
    else:
        cancer_col_eff = cancer_col

    if groupby is None:
        tmp["_group"] = "all"
        groupby_eff = "_group"
    else:
        groupby_eff = groupby

    # Mean CNV by cancer/group/chromosome.
    joined = tmp.join(chrom_df, how="inner")
    group_mean = (
        joined.groupby([cancer_col_eff, groupby_eff], observed=False)[chrom_df.columns]
        .mean()
        .reset_index()
        .rename(columns={cancer_col_eff: "cancer", groupby_eff: "group"})
    )

    # Type fractions by cancer/group/chromosome/type.
    type_long = (
        type_df.reset_index()
        .melt(id_vars="cell_id", var_name="chromosome", value_name="cnv_type")
    )
    type_long = type_long.merge(
        tmp.reset_index().rename(columns={cancer_col_eff: "cancer", groupby_eff: "group"}),
        on="cell_id",
        how="left",
    )
    type_fraction = (
        type_long
        .groupby(["cancer", "group", "chromosome", "cnv_type"], observed=False)
        .size()
        .rename("n_cells")
        .reset_index()
    )
    denom = (
        type_long
        .groupby(["cancer", "group", "chromosome"], observed=False)
        .size()
        .rename("n_total")
        .reset_index()
    )
    type_fraction = type_fraction.merge(denom, on=["cancer", "group", "chromosome"], how="left")
    type_fraction["fraction"] = type_fraction["n_cells"] / type_fraction["n_total"].replace(0, np.nan)

    # Type fractions by cancer/chromosome/type.
    cancer_fraction = (
        type_long
        .groupby(["cancer", "chromosome", "cnv_type"], observed=False)
        .size()
        .rename("n_cells")
        .reset_index()
    )
    denom2 = (
        type_long
        .groupby(["cancer", "chromosome"], observed=False)
        .size()
        .rename("n_total")
        .reset_index()
    )
    cancer_fraction = cancer_fraction.merge(denom2, on=["cancer", "chromosome"], how="left")
    cancer_fraction["fraction"] = cancer_fraction["n_cells"] / cancer_fraction["n_total"].replace(0, np.nan)

    return group_mean, type_fraction, cancer_fraction


# -----------------------------
# plotting
# -----------------------------

def _plot_matrix_clustermap(
    matrix: pd.DataFrame,
    path_base: Path,
    *,
    title: str,
    center: float = 0.0,
    cmap: str = "vlag",
    vlim: float | None = None,
    figsize_scale: tuple[float, float] = (0.45, 0.22),
    min_figsize: tuple[float, float] = (8, 6),
    max_figsize: tuple[float, float] = (26, 30),
    row_cluster: bool | None = None,
    col_cluster: bool | None = None,
    row_colors=None,
    dpi: int = 220,
) -> bool:
    if matrix.empty:
        return False

    # Drop empty/all-NaN rows and cols.
    matrix = matrix.copy()
    matrix = matrix.dropna(axis=0, how="all").dropna(axis=1, how="all")
    if matrix.empty:
        return False

    # Fill rare NaNs with 0 for clustering/plotting.
    matrix = matrix.astype(float).replace([np.inf, -np.inf], np.nan).fillna(0.0)

    if vlim is None:
        vals = matrix.to_numpy().ravel()
        vals = vals[np.isfinite(vals)]
        vlim = float(np.nanquantile(np.abs(vals), 0.99)) if len(vals) else 1.0
        vlim = max(vlim, 1e-6)

    rc_default, cc_default = _cluster_flags(matrix)
    row_cluster = rc_default if row_cluster is None else bool(row_cluster and rc_default)
    col_cluster = cc_default if col_cluster is None else bool(col_cluster and cc_default)

    width = min(max(min_figsize[0], matrix.shape[1] * figsize_scale[0]), max_figsize[0])
    height = min(max(min_figsize[1], matrix.shape[0] * figsize_scale[1]), max_figsize[1])

    g = sns.clustermap(
        matrix,
        cmap=cmap,
        center=center,
        vmin=-vlim if center == 0 else None,
        vmax=vlim if center == 0 else None,
        row_cluster=row_cluster,
        col_cluster=col_cluster,
        row_colors=row_colors,
        xticklabels=True,
        yticklabels=matrix.shape[0] <= 80,
        linewidths=0.0,
        figsize=(width, height),
        cbar_kws={"label": "Mean inferCNV signal"},
    )
    g.fig.suptitle(title, y=1.02, fontsize=14)
    _save_clustermap(g, path_base, dpi=dpi)
    return True


def _plot_type_fraction_heatmap(
    type_fraction: pd.DataFrame,
    path_base: Path,
    *,
    title: str,
    dpi: int = 220,
) -> bool:
    if type_fraction.empty:
        return False

    # Rows: chromosome. Columns: loss/neutral/gain. Values: fraction.
    p = type_fraction.pivot_table(
        index="chromosome",
        columns="cnv_type",
        values="fraction",
        aggfunc="mean",
        fill_value=0.0,
    )
    for c in ["loss", "neutral", "gain"]:
        if c not in p.columns:
            p[c] = 0.0
    p = p.loc[sorted(p.index, key=_natural_key), ["loss", "neutral", "gain"]]

    h = max(6, min(18, 0.35 * p.shape[0]))
    plt.figure(figsize=(8, h))
    sns.heatmap(
        p,
        vmin=0,
        vmax=1,
        cmap="viridis",
        annot=p.shape[0] <= 30,
        fmt=".2f",
        cbar_kws={"label": "Fraction of cells"},
    )
    plt.title(title)
    plt.xlabel("CNV type")
    plt.ylabel("Chromosome")
    plt.tight_layout()
    _save_current_fig(path_base, dpi=dpi)
    return True


def _sample_cells_for_plot(df: pd.DataFrame, obs: pd.DataFrame, max_cells: int, stratify_col: str | None = None, seed: int = 13) -> pd.DataFrame:
    if df.shape[0] <= max_cells:
        return df

    rng = np.random.default_rng(seed)

    if stratify_col and stratify_col in obs.columns:
        # Stratified sample so rare malignant/clone groups do not disappear.
        keep = []
        obs_aligned = obs.reindex(df.index)
        groups = obs_aligned[stratify_col].astype(str).fillna("NA")
        per_group = max(10, int(math.ceil(max_cells / max(1, groups.nunique()))))
        for _, idx in groups.groupby(groups).groups.items():
            idx = np.array(list(idx))
            n = min(len(idx), per_group)
            keep.extend(rng.choice(idx, size=n, replace=False).tolist())
        if len(keep) > max_cells:
            keep = rng.choice(np.array(keep), size=max_cells, replace=False).tolist()
        return df.loc[keep]

    keep = rng.choice(df.index.to_numpy(), size=max_cells, replace=False)
    return df.loc[keep]


def _plot_all_cancer_summaries(
    chrom_df: pd.DataFrame,
    type_df: pd.DataFrame,
    obs: pd.DataFrame,
    *,
    outfig: Path,
    cancer_col: str | None,
    dpi: int = 220,
) -> None:
    if cancer_col is None or cancer_col not in obs.columns:
        return

    joined = obs[[cancer_col]].join(chrom_df, how="inner")
    cancer_mean = joined.groupby(cancer_col, observed=False)[chrom_df.columns].mean()
    if cancer_mean.shape[0] >= 2:
        _plot_matrix_clustermap(
            cancer_mean,
            outfig / "all_cancers_chromosome_mean_clustermap",
            title="All cancers: mean chromosome-level inferCNV signal",
            cmap="vlag",
            center=0,
            dpi=dpi,
            figsize_scale=(0.55, 0.55),
            min_figsize=(10, 7),
        )

    # Signed alteration fraction: gain fraction minus loss fraction.
    type_long = (
        type_df.reset_index()
        .melt(id_vars="cell_id", var_name="chromosome", value_name="cnv_type")
        .merge(obs[[cancer_col]].reset_index(), on="cell_id", how="left")
    )
    frac = (
        type_long
        .groupby([cancer_col, "chromosome", "cnv_type"], observed=False)
        .size()
        .rename("n")
        .reset_index()
    )
    denom = (
        type_long
        .groupby([cancer_col, "chromosome"], observed=False)
        .size()
        .rename("N")
        .reset_index()
    )
    frac = frac.merge(denom, on=[cancer_col, "chromosome"], how="left")
    frac["fraction"] = frac["n"] / frac["N"].replace(0, np.nan)
    p = frac.pivot_table(
        index=[cancer_col, "chromosome"],
        columns="cnv_type",
        values="fraction",
        fill_value=0.0,
    ).reset_index()
    for c in ["gain", "loss"]:
        if c not in p.columns:
            p[c] = 0.0
    p["signed_gain_minus_loss_fraction"] = p["gain"] - p["loss"]
    signed = p.pivot_table(
        index=cancer_col,
        columns="chromosome",
        values="signed_gain_minus_loss_fraction",
        fill_value=0.0,
    )
    signed = signed.loc[:, sorted(signed.columns, key=_natural_key)]
    if signed.shape[0] >= 2:
        _plot_matrix_clustermap(
            signed,
            outfig / "all_cancers_signed_gain_loss_fraction_clustermap",
            title="All cancers: signed alteration fraction by chromosome",
            cmap="vlag",
            center=0,
            vlim=1.0,
            dpi=dpi,
            figsize_scale=(0.55, 0.55),
            min_figsize=(10, 7),
        )


def _maybe_plot_builtin_infercnvpy_heatmap(
    adata_sub,
    *,
    outpath_base: Path,
    groupby: str | None,
    use_rep: str = "cnv",
    dpi: int = 220,
) -> None:
    if groupby is None or groupby not in adata_sub.obs.columns:
        return
    try:
        import infercnvpy as cnv
        figdir_old = None
        try:
            import scanpy as sc
            figdir_old = sc.settings.figdir
            sc.settings.figdir = outpath_base.parent
        except Exception:
            pass

        # infercnvpy's save argument appends to its default file name; this is less predictable.
        # show=False gives axes; then save current figure ourselves.
        cnv.pl.chromosome_heatmap(
            adata_sub,
            groupby=groupby,
            use_rep=use_rep,
            show=False,
            figsize=(18, 10),
            dendrogram=False,
        )
        plt.gcf().suptitle(f"inferCNVpy chromosome heatmap grouped by {groupby}", y=1.02)
        _save_current_fig(outpath_base, dpi=dpi)

        if figdir_old is not None:
            import scanpy as sc
            sc.settings.figdir = figdir_old
    except Exception as e:
        warnings.warn(f"Could not make built-in inferCNVpy chromosome_heatmap: {e}")


def make_all_plots(
    adata,
    chrom_df: pd.DataFrame,
    type_df: pd.DataFrame,
    group_mean: pd.DataFrame,
    type_fraction: pd.DataFrame,
    cancer_fraction: pd.DataFrame,
    obs: pd.DataFrame,
    *,
    outfig: Path,
    cancer_col: str | None,
    groupby: str | None,
    use_rep: str = "cnv",
    max_cells_plot: int = 3000,
    make_builtin_infercnvpy_heatmap: bool = False,
    dpi: int = 220,
) -> None:
    outfig.mkdir(parents=True, exist_ok=True)

    _plot_all_cancer_summaries(chrom_df, type_df, obs, outfig=outfig, cancer_col=cancer_col, dpi=dpi)

    if cancer_col and cancer_col in obs.columns:
        cancers = list(pd.Series(obs[cancer_col]).dropna().astype(str).unique())
    else:
        cancers = ["all"]

    for cancer in cancers:
        if cancer == "all":
            idx = chrom_df.index
        else:
            idx = obs.index[obs[cancer_col].astype(str).eq(str(cancer))]
            idx = chrom_df.index.intersection(idx)

        if len(idx) == 0:
            continue

        safe_cancer = _safe_name(cancer)
        cfig = outfig / safe_cancer
        cfig.mkdir(parents=True, exist_ok=True)

        c_chrom = chrom_df.loc[idx]
        c_type = type_df.loc[idx]
        c_obs = obs.reindex(idx)

        # 1) single-cell chromosome-level clustermap
        c_plot = _sample_cells_for_plot(c_chrom, c_obs, max_cells=max_cells_plot, stratify_col=groupby)
        _plot_matrix_clustermap(
            c_plot,
            cfig / "cells_x_chromosomes_mean_cnv_clustermap",
            title=f"{cancer}: cells × chromosomes mean inferCNV signal",
            cmap="vlag",
            center=0,
            dpi=dpi,
            figsize_scale=(0.50, 0.008 if c_plot.shape[0] > 500 else 0.08),
            min_figsize=(10, 8),
            max_figsize=(24, 28),
        )

        # 2) group-level mean CNV clustermap
        gm = group_mean.loc[group_mean["cancer"].astype(str).eq(str(cancer))].copy()
        if not gm.empty:
            mat = gm.set_index("group")[chrom_df.columns]
            _plot_matrix_clustermap(
                mat,
                cfig / "groups_x_chromosomes_mean_cnv_clustermap",
                title=f"{cancer}: CNV groups × chromosomes mean inferCNV signal",
                cmap="vlag",
                center=0,
                dpi=dpi,
                figsize_scale=(0.55, 0.45),
                min_figsize=(10, 6),
                max_figsize=(24, 20),
            )

        # 3) group-level gain/loss fractions
        tf = type_fraction.loc[type_fraction["cancer"].astype(str).eq(str(cancer))].copy()
        if not tf.empty:
            for cnv_type, pretty in [("gain", "gain"), ("loss", "loss")]:
                sub = tf.loc[tf["cnv_type"].eq(cnv_type)]
                if sub.empty:
                    continue
                mat = sub.pivot_table(
                    index="group",
                    columns="chromosome",
                    values="fraction",
                    aggfunc="mean",
                    fill_value=0.0,
                )
                mat = mat.loc[:, sorted(mat.columns, key=_natural_key)]
                if not mat.empty:
                    _plot_matrix_clustermap(
                        mat,
                        cfig / f"groups_x_chromosomes_{cnv_type}_fraction_clustermap",
                        title=f"{cancer}: fraction of cells with chromosome {pretty}",
                        cmap="viridis",
                        center=None,
                        vlim=None,
                        dpi=dpi,
                        figsize_scale=(0.55, 0.45),
                        min_figsize=(10, 6),
                        max_figsize=(24, 20),
                        row_cluster=mat.shape[0] >= 2,
                        col_cluster=mat.shape[1] >= 2,
                    )

        # 4) cancer-level chromosome × type fraction heatmap
        cf = cancer_fraction.loc[cancer_fraction["cancer"].astype(str).eq(str(cancer))].copy()
        _plot_type_fraction_heatmap(
            cf,
            cfig / "chromosomes_x_cnv_type_fraction_heatmap",
            title=f"{cancer}: chromosome-level CNV type fractions",
            dpi=dpi,
        )

        # 5) optional raw inferCNVpy window-level heatmap
        if make_builtin_infercnvpy_heatmap:
            try:
                adata_sub = adata[idx, :].copy()
                _maybe_plot_builtin_infercnvpy_heatmap(
                    adata_sub,
                    outpath_base=cfig / f"infercnvpy_window_heatmap_groupby_{_safe_name(groupby)}",
                    groupby=groupby,
                    use_rep=use_rep,
                    dpi=dpi,
                )
            except Exception as e:
                warnings.warn(f"{cancer}: skipping built-in inferCNVpy heatmap because subsetting failed: {e}")


# -----------------------------
# main export function
# -----------------------------

def export_cnv_chromosome_clustermaps(
    adata,
    *,
    outdir: str | Path,
    cancer_col: str | None = None,
    groupby: str | None = None,
    use_rep: str = "cnv",
    window_size: int = 100,
    step: int = 10,
    cnv_type_threshold: float = 0.05,
    exclude_chromosomes: Sequence[str] | None = ("chrX", "chrY"),
    max_cells_plot: int = 3000,
    make_builtin_infercnvpy_heatmap: bool = False,
    dpi: int = 220,
    progress: bool = True,
) -> dict[str, str]:
    """
    Export all tables and figures needed for chromosome-level CNV clustermaps.

    This version is intentionally chatty: it prints step timings and uses tqdm for
    long per-chromosome/per-dataset work so command-line runs do not look frozen.
    """
    outdir = Path(outdir)
    outtab = outdir / "tables"
    outfig = outdir / "figures"
    outtab.mkdir(parents=True, exist_ok=True)
    outfig.mkdir(parents=True, exist_ok=True)

    step_bar = tqdm(total=6, desc="export steps", unit="step", disable=not progress)

    try:
        with _timed_step("detect metadata columns"):
            cancer_col = _choose_cancer_col(adata.obs, cancer_col)
            groupby = _choose_groupby(adata.obs, groupby)

            _print(f"[cnv-export] outdir: {outdir}")
            _print(f"[cnv-export] use_rep: {use_rep}")
            _print(f"[cnv-export] cancer_col: {cancer_col}")
            _print(f"[cnv-export] groupby: {groupby}")
            _print(f"[cnv-export] window_size={window_size}, step={step}, threshold={cnv_type_threshold}")
            _print(f"[cnv-export] adata: cells={adata.n_obs:,}, genes={adata.n_vars:,}")
        step_bar.update(1)

        with _timed_step("reconstruct inferCNV windows"):
            windows = reconstruct_cnv_windows(
                adata,
                use_rep=use_rep,
                window_size=window_size,
                step=step,
                exclude_chromosomes=exclude_chromosomes,
            )
            _print(f"[cnv-export] windows: {len(windows):,}; chromosomes={windows['chromosome'].nunique():,}")
        step_bar.update(1)

        with _timed_step("compute chromosome-level CNV tables"):
            chrom_df = compute_cell_chromosome_means(adata, windows, use_rep=use_rep)
            type_df = classify_cnv_types(chrom_df, threshold=cnv_type_threshold)
            obs = make_obs_table(adata, cancer_col, groupby)
            group_mean, type_fraction, cancer_fraction = compute_group_tables(
                chrom_df, type_df, obs, cancer_col=cancer_col, groupby=groupby
            )
            _print(f"[cnv-export] per-cell chromosome matrix: {chrom_df.shape[0]:,} cells × {chrom_df.shape[1]:,} chromosomes")
            _print(f"[cnv-export] group mean rows: {len(group_mean):,}; type fraction rows: {len(type_fraction):,}")
        step_bar.update(1)

        with _timed_step("write tables"):
            windows_path = outtab / "cnv_windows.csv.gz"
            chrom_path = outtab / "per_cell_chromosome_mean_cnv.csv.gz"
            type_path = outtab / "per_cell_chromosome_cnv_type.csv.gz"
            group_mean_path = outtab / "per_group_chromosome_mean_cnv.csv.gz"
            type_frac_path = outtab / "per_group_chromosome_cnv_type_fraction.csv.gz"
            cancer_frac_path = outtab / "per_cancer_chromosome_cnv_type_fraction.csv.gz"
            metadata_path = outtab / "export_metadata.json"

            table_items = [
                ("windows", windows, windows_path),
                ("per-cell chromosome means", chrom_df, chrom_path),
                ("per-cell chromosome types", type_df, type_path),
                ("per-group chromosome means", group_mean, group_mean_path),
                ("per-group type fractions", type_fraction, type_frac_path),
                ("per-cancer type fractions", cancer_fraction, cancer_frac_path),
            ]
            for label, df, path in tqdm(table_items, desc="write tables", unit="table", disable=not progress, leave=False):
                _write_df(df, path)
                _print(f"[write] {label}: {path}")

            metadata = {
                "use_rep": use_rep,
                "obsm_key": f"X_{use_rep}",
                "uns_key": use_rep,
                "n_cells": int(adata.n_obs),
                "n_genes": int(adata.n_vars),
                "n_cnv_windows": int(get_cnv_matrix(adata, use_rep=use_rep).shape[1]),
                "n_chromosomes": int(chrom_df.shape[1]),
                "chromosomes": list(map(str, chrom_df.columns)),
                "cancer_col": cancer_col,
                "groupby": groupby,
                "window_size": int(window_size),
                "step": int(step),
                "cnv_type_threshold": float(cnv_type_threshold),
                "exclude_chromosomes": list(exclude_chromosomes) if exclude_chromosomes is not None else None,
                "max_cells_plot": int(max_cells_plot),
                "make_builtin_infercnvpy_heatmap": bool(make_builtin_infercnvpy_heatmap),
            }
            metadata_path.write_text(json.dumps(metadata, indent=2), encoding="utf-8")
            _print(f"[write] metadata: {metadata_path}")
        step_bar.update(1)

        with _timed_step("make figures"):
            with _heartbeat("figure generation", interval=45):
                make_all_plots(
                    adata,
                    chrom_df,
                    type_df,
                    group_mean,
                    type_fraction,
                    cancer_fraction,
                    obs,
                    outfig=outfig,
                    cancer_col=cancer_col,
                    groupby=groupby,
                    use_rep=use_rep,
                    max_cells_plot=max_cells_plot,
                    make_builtin_infercnvpy_heatmap=make_builtin_infercnvpy_heatmap,
                    dpi=dpi,
                )
        step_bar.update(1)

        with _timed_step("finalize export"):
            _print("[cnv-export] done.")
            _print(f"[cnv-export] tables:  {outtab}")
            _print(f"[cnv-export] figures: {outfig}")
        step_bar.update(1)

    finally:
        step_bar.close()

    return {
        "outdir": str(outdir),
        "tables": str(outtab),
        "figures": str(outfig),
        "windows": str(windows_path),
        "per_cell_chromosome_mean_cnv": str(chrom_path),
        "per_cell_chromosome_cnv_type": str(type_path),
        "per_group_chromosome_mean_cnv": str(group_mean_path),
        "per_group_chromosome_cnv_type_fraction": str(type_frac_path),
        "per_cancer_chromosome_cnv_type_fraction": str(cancer_frac_path),
        "metadata": str(metadata_path),
    }


# -----------------------------
# CLI: single-file or all-cancers batch mode
# -----------------------------

DEFAULT_INFERCNV_DIR = Path(
    r"C:\Users\ashfo\OneDrive\Desktop\Pathways+Omics\scOPE\scOPE\data\scope_robust_real_inputs\infercnvpy_new"
)
DEFAULT_DATASETS = ["AML", "BRCA", "CRC", "GBM", "LUAD", "PAAD", "SKCM"]


def _split_csv_arg(x: str | None, default: Sequence[str] | None = None) -> list[str]:
    if x is None or str(x).strip() == "":
        return list(default or [])
    if str(x).strip().lower() in {"all", "*"}:
        return list(default or [])
    return [v.strip() for v in str(x).split(",") if v.strip()]


def _find_dataset_h5ad(
    h5ad_dir: Path,
    dataset: str,
    *,
    pattern: str = "{dataset}_infercnvpy.h5ad",
    recursive: bool = False,
) -> Path | None:
    """Find one inferCNVpy h5ad for a dataset using exact pattern first, then safe fallbacks."""
    exact = h5ad_dir / pattern.format(dataset=dataset)
    if exact.exists():
        return exact

    globber = h5ad_dir.rglob if recursive else h5ad_dir.glob
    candidates = list(globber(pattern.format(dataset="*")))
    candidates += list(globber(f"{dataset}*infercnvpy*.h5ad"))
    candidates += list(globber(f"*{dataset}*infercnvpy*.h5ad"))

    # De-duplicate and prefer shorter / closer names.
    unique = []
    seen = set()
    for p in candidates:
        if p in seen:
            continue
        seen.add(p)
        if p.is_file() and p.suffix == ".h5ad":
            unique.append(p)

    if not unique:
        return None

    unique = sorted(unique, key=lambda p: (len(str(p)), str(p).lower()))
    return unique[0]


def _add_dataset_obs_column(adata, dataset: str, cancer_col: str | None) -> None:
    """
    Ensure there is a dataset/cancer column for the exporter.

    This does not alter the input file on disk unless the user explicitly writes it later.
    """
    if "dataset" not in adata.obs.columns:
        adata.obs["dataset"] = str(dataset)

    if cancer_col and cancer_col not in {"auto", "None", "none", ""}:
        if cancer_col not in adata.obs.columns:
            adata.obs[cancer_col] = str(dataset)


def _compute_cnv_reduced_embedding(
    adata,
    *,
    obsm_key: str,
    out_key: str = "X_cnv_svd",
    n_components: int = 30,
    random_state: int = 0,
    progress: bool = True,
) -> str:
    """
    Create a low-dimensional embedding from X_cnv for faster neighbors/Leiden.

    Why this exists:
    scanpy.pp.neighbors(..., use_rep="X_cnv") can look frozen on large cancer
    objects because it builds kNN directly on thousands of CNV windows. Reducing
    to 20-50 components first is faster and usually cleaner for broad CNV clones.
    """
    if out_key in adata.obsm:
        _print(f"[cluster] using existing adata.obsm[{out_key!r}] with shape {_matrix_shape_string(adata.obsm[out_key])}")
        return out_key

    X = adata.obsm[obsm_key]
    n_obs, n_features = int(X.shape[0]), int(X.shape[1])
    n_components = int(max(2, min(n_components, n_obs - 1, n_features - 1)))

    _print(f"[cluster] reducing {obsm_key}: {n_obs:,} cells × {n_features:,} CNV windows -> {n_components} components")

    # Convert to float32 only when needed. Dense X_cnv is usually already dense.
    try:
        from sklearn.decomposition import TruncatedSVD
    except Exception as e:
        raise RuntimeError("Automatic CNV Leiden clustering requires scikit-learn. Install with: pip install scikit-learn") from e

    with _heartbeat(f"SVD/PCA reduction of {obsm_key}", interval=30):
        if sp is not None and sp.issparse(X):
            model = TruncatedSVD(n_components=n_components, random_state=random_state)
            emb = model.fit_transform(X).astype(np.float32, copy=False)
        else:
            # Keep this memory-safe: avoid StandardScaler/PCA copies on large dense matrices.
            # TruncatedSVD on raw CNV windows is usually enough for broad CNV clone grouping.
            X_arr = np.asarray(X, dtype=np.float32)
            model = TruncatedSVD(n_components=n_components, random_state=random_state)
            emb = model.fit_transform(X_arr).astype(np.float32, copy=False)

    adata.obsm[out_key] = emb
    _print(f"[cluster] wrote adata.obsm[{out_key!r}] shape={emb.shape}")
    return out_key


def _ensure_groupby_column(
    adata,
    *,
    requested_groupby: str | None,
    make_cnv_leiden: bool,
    use_rep: str,
    leiden_resolution: float,
    neighbors_n: int,
    cnv_leiden_rep: str = "svd",
    cnv_svd_components: int = 30,
    progress: bool = True,
) -> str | None:
    """
    Resolve the groupby column.

    - If requested_groupby is a real obs column, use it.
    - If requested_groupby is auto/None, prefer cnv_leiden if present.
    - Optionally create cnv_leiden from reduced CNV space with visible progress.
    - Otherwise return None and let export_cnv_chromosome_clustermaps auto-detect.
    """
    if requested_groupby and requested_groupby.lower() not in {"auto", "none", "null"}:
        if requested_groupby in adata.obs.columns:
            _print(f"[groupby] using requested obs column: {requested_groupby}")
            return requested_groupby
        warnings.warn(f"Requested --groupby {requested_groupby!r} is not in adata.obs; falling back to auto.")

    if "cnv_leiden" in adata.obs.columns:
        _print("[groupby] using existing obs column: cnv_leiden")
        return "cnv_leiden"

    if make_cnv_leiden:
        try:
            import scanpy as sc

            obsm_key = _resolve_obsm_key(adata, use_rep)
            _print(f"[cluster] requested automatic CNV Leiden clustering from adata.obsm[{obsm_key!r}]")
            _print(f"[cluster] input CNV matrix shape: {_matrix_shape_string(adata.obsm[obsm_key])}")

            with tqdm(total=3, desc="CNV Leiden", unit="step", disable=not progress) as pbar:
                if cnv_leiden_rep.lower() in {"svd", "pca", "reduced"}:
                    with _timed_step("CNV dimensionality reduction"):
                        cluster_rep = _compute_cnv_reduced_embedding(
                            adata,
                            obsm_key=obsm_key,
                            out_key=f"X_{use_rep}_svd",
                            n_components=cnv_svd_components,
                            progress=progress,
                        )
                elif cnv_leiden_rep.lower() in {"raw", "x_cnv", "cnv"}:
                    cluster_rep = obsm_key
                    _print(f"[cluster] using raw CNV windows for neighbors: {cluster_rep}")
                else:
                    raise ValueError("--cnv-leiden-rep must be 'svd' or 'raw'")
                pbar.update(1)

                with _timed_step(f"build kNN graph on {cluster_rep}"):
                    with _heartbeat("scanpy.pp.neighbors", interval=30):
                        sc.pp.neighbors(
                            adata,
                            use_rep=cluster_rep,
                            n_neighbors=neighbors_n,
                            random_state=0,
                        )
                pbar.update(1)

                with _timed_step("run Leiden clustering"):
                    with _heartbeat("scanpy.tl.leiden", interval=30):
                        try:
                            sc.tl.leiden(
                                adata,
                                resolution=leiden_resolution,
                                key_added="cnv_leiden",
                                random_state=0,
                                flavor="igraph",
                                n_iterations=2,
                                directed=False,
                            )
                        except TypeError:
                            # Older scanpy versions do not support all kwargs.
                            sc.tl.leiden(
                                adata,
                                resolution=leiden_resolution,
                                key_added="cnv_leiden",
                                random_state=0,
                            )
                pbar.update(1)

            counts = adata.obs["cnv_leiden"].astype(str).value_counts().to_dict()
            _print("[cluster] cnv_leiden counts: " + json.dumps(counts, indent=None))
            return "cnv_leiden"

        except Exception as e:
            warnings.warn(
                f"Could not create cnv_leiden automatically: {type(e).__name__}: {e}. "
                "Falling back to existing obs columns."
            )

    # Let export_cnv_chromosome_clustermaps auto-detect from cnv_status/malignant_call/celltype/etc.
    _print("[groupby] no CNV Leiden; exporter will auto-detect fallback grouping")
    return None


def _make_batch_summary_from_exported_tables(
    *,
    batch_outdir: Path,
    successful: Sequence[str],
    dpi: int = 220,
) -> None:
    """
    Build across-cancer summary plots from each dataset's exported chromosome tables.

    This works even when each cancer was processed as a separate AnnData file.
    """
    summary_dir = batch_outdir / "BATCH_SUMMARY"
    summary_tables = summary_dir / "tables"
    summary_figures = summary_dir / "figures"
    summary_tables.mkdir(parents=True, exist_ok=True)
    summary_figures.mkdir(parents=True, exist_ok=True)

    mean_rows = []
    type_records = []

    for cancer in successful:
        tables = batch_outdir / _safe_name(cancer) / "tables"
        mean_path = tables / "per_cell_chromosome_mean_cnv.csv.gz"
        type_path = tables / "per_cell_chromosome_cnv_type.csv.gz"

        if mean_path.exists():
            m = pd.read_csv(mean_path, index_col=0)
            row = m.mean(axis=0, numeric_only=True)
            row.name = cancer
            mean_rows.append(row)

        if type_path.exists():
            t = pd.read_csv(type_path, index_col=0)
            for chrom in t.columns:
                vc = t[chrom].astype(str).value_counts(normalize=True)
                for cnv_type in ["loss", "neutral", "gain"]:
                    type_records.append({
                        "cancer": cancer,
                        "chromosome": chrom,
                        "cnv_type": cnv_type,
                        "fraction": float(vc.get(cnv_type, 0.0)),
                    })

    if not mean_rows:
        warnings.warn("No exported per-cell chromosome mean tables found for batch summary.")
        return

    cancer_mean = pd.DataFrame(mean_rows).fillna(0.0)
    cancer_mean = cancer_mean.loc[:, sorted(cancer_mean.columns, key=_natural_key)]
    cancer_mean.index.name = "cancer"
    cancer_mean.to_csv(summary_tables / "batch_cancer_chromosome_mean_cnv.csv")

    _plot_matrix_clustermap(
        cancer_mean,
        summary_figures / "batch_all_cancers_chromosome_mean_clustermap",
        title="All cancers: mean chromosome-level inferCNV signal",
        cmap="vlag",
        center=0,
        dpi=dpi,
        figsize_scale=(0.55, 0.55),
        min_figsize=(10, 7),
    )

    type_fraction = pd.DataFrame(type_records)
    if not type_fraction.empty:
        type_fraction.to_csv(summary_tables / "batch_cancer_chromosome_cnv_type_fraction.csv", index=False)

        p = type_fraction.pivot_table(
            index=["cancer", "chromosome"],
            columns="cnv_type",
            values="fraction",
            fill_value=0.0,
        ).reset_index()

        for c in ["gain", "loss"]:
            if c not in p.columns:
                p[c] = 0.0

        p["signed_gain_minus_loss_fraction"] = p["gain"] - p["loss"]

        signed = p.pivot_table(
            index="cancer",
            columns="chromosome",
            values="signed_gain_minus_loss_fraction",
            fill_value=0.0,
        )
        signed = signed.loc[:, sorted(signed.columns, key=_natural_key)]

        _plot_matrix_clustermap(
            signed,
            summary_figures / "batch_all_cancers_signed_gain_loss_fraction_clustermap",
            title="All cancers: signed gain-minus-loss fraction by chromosome",
            cmap="vlag",
            center=0,
            vlim=1.0,
            dpi=dpi,
            figsize_scale=(0.55, 0.55),
            min_figsize=(10, 7),
        )

        for cnv_type in ["gain", "loss"]:
            mat = type_fraction.loc[type_fraction["cnv_type"].eq(cnv_type)].pivot_table(
                index="cancer",
                columns="chromosome",
                values="fraction",
                fill_value=0.0,
            )
            if not mat.empty:
                mat = mat.loc[:, sorted(mat.columns, key=_natural_key)]
                _plot_matrix_clustermap(
                    mat,
                    summary_figures / f"batch_all_cancers_{cnv_type}_fraction_clustermap",
                    title=f"All cancers: chromosome {cnv_type} fraction",
                    cmap="viridis",
                    center=None,
                    dpi=dpi,
                    figsize_scale=(0.55, 0.55),
                    min_figsize=(10, 7),
                )

    print(f"[batch-summary] tables:  {summary_tables}")
    print(f"[batch-summary] figures: {summary_figures}")


def _parse_args() -> argparse.Namespace:
    p = argparse.ArgumentParser(
        description=(
            "Export chromosome-level inferCNVpy tables and clustermap plots from either "
            "one AnnData .h5ad or all per-cancer inferCNVpy .h5ad files in a directory."
        )
    )

    mode = p.add_argument_group("Input mode")
    mode.add_argument("--h5ad", default=None, help="Single inferCNVpy AnnData .h5ad file.")
    mode.add_argument(
        "--h5ad-dir",
        default=str(DEFAULT_INFERCNV_DIR),
        help="Directory containing per-cancer inferCNVpy h5ads. Used when --h5ad is omitted.",
    )
    mode.add_argument(
        "--h5ad-pattern",
        default="{dataset}_infercnvpy.h5ad",
        help="Pattern for per-cancer h5ads inside --h5ad-dir. Must include {dataset}.",
    )
    mode.add_argument(
        "--recursive",
        action="store_true",
        help="Search --h5ad-dir recursively for per-cancer h5ads.",
    )
    mode.add_argument(
        "--datasets",
        default="AML,BRCA,CRC,GBM,LUAD,PAAD,SKCM",
        help="Comma-separated datasets to process, or 'auto' to use every *.h5ad in --h5ad-dir.",
    )

    out = p.add_argument_group("Outputs")
    out.add_argument(
        "--outdir",
        default=None,
        help=(
            "Output directory. For batch mode, each cancer gets a subfolder. "
            "Default: <h5ad-dir>/cnv_chromosome_clustermaps"
        ),
    )
    out.add_argument(
        "--skip-existing",
        action="store_true",
        help="Skip a dataset when <outdir>/<dataset>/tables/export_metadata.json already exists.",
    )
    out.add_argument(
        "--no-batch-summary",
        action="store_true",
        help="Do not make BATCH_SUMMARY plots/tables after processing all cancers.",
    )

    meta = p.add_argument_group("Metadata/grouping")
    meta.add_argument(
        "--cancer-col",
        default="dataset",
        help="adata.obs column for cancer/dataset. Use 'auto' for auto-detect. Default: dataset.",
    )
    meta.add_argument(
        "--groupby",
        default="auto",
        help=(
            "adata.obs column for rows/groups in group-level plots. "
            "Use 'auto' to prefer cnv_leiden, then malignant_call/celltype/etc."
        ),
    )
    meta.add_argument(
        "--make-cnv-leiden",
        action="store_true",
        help="If cnv_leiden is missing, build Leiden clusters from X_cnv using scanpy.",
    )
    meta.add_argument("--leiden-resolution", type=float, default=1.0)
    meta.add_argument("--neighbors-n", type=int, default=15)
    meta.add_argument(
        "--cnv-leiden-rep",
        default="svd",
        choices=["svd", "raw"],
        help="Representation for automatic CNV Leiden. Default 'svd' is much faster than raw X_cnv.",
    )
    meta.add_argument(
        "--cnv-svd-components",
        type=int,
        default=30,
        help="Number of SVD/PCA components for --make-cnv-leiden when --cnv-leiden-rep svd.",
    )
    meta.add_argument(
        "--heartbeat-seconds",
        type=int,
        default=30,
        help="Print still-running messages this often during slow library calls.",
    )
    meta.add_argument(
        "--quiet-progress",
        action="store_true",
        help="Disable tqdm progress bars; step start/done logs still print.",
    )

    cnv_args = p.add_argument_group("CNV/window settings")
    cnv_args.add_argument("--use-rep", default="cnv", help="inferCNVpy key; default expects adata.obsm['X_cnv'].")
    cnv_args.add_argument("--window-size", type=int, default=100, help="window_size used in the original inferCNVpy run.")
    cnv_args.add_argument("--step", type=int, default=10, help="step used in the original inferCNVpy run.")
    cnv_args.add_argument("--threshold", type=float, default=0.05, help="Chromosome mean > threshold is gain; < -threshold is loss.")
    cnv_args.add_argument("--include-sex-chromosomes", action="store_true", help="Do not exclude chrX/chrY during window reconstruction.")

    plot = p.add_argument_group("Plot settings")
    plot.add_argument("--max-cells-plot", type=int, default=3000, help="Max cells per cancer in cell-level clustermap only.")
    plot.add_argument("--dpi", type=int, default=220)
    plot.add_argument(
        "--builtin-infercnvpy-heatmap",
        action="store_true",
        help="Also make inferCNVpy raw window-level chromosome_heatmap; can be memory-heavy.",
    )

    return p.parse_args()


def _discover_auto_datasets(h5ad_dir: Path, recursive: bool = False) -> list[tuple[str, Path]]:
    globber = h5ad_dir.rglob if recursive else h5ad_dir.glob
    files = sorted(globber("*.h5ad"))
    discovered = []
    for p in files:
        stem = p.stem
        # Common names from make_infercnvpy_cnvs.py: AML_infercnvpy.h5ad
        dataset = re.sub(r"_?infercnvpy.*$", "", stem, flags=re.IGNORECASE)
        dataset = dataset or stem
        discovered.append((dataset, p))
    return discovered


def _run_single_h5ad(args: argparse.Namespace) -> None:
    import scanpy as sc

    h5ad = Path(args.h5ad)
    if not h5ad.exists():
        raise FileNotFoundError(h5ad)

    outdir = Path(args.outdir) if args.outdir else h5ad.with_suffix("").parent / f"{h5ad.stem}_cnv_chromosome_clustermaps"
    exclude = None if args.include_sex_chromosomes else ("chrX", "chrY")

    dataset = h5ad.stem.replace("_infercnvpy", "")
    _print(f"[single] reading: {h5ad}")
    with _timed_step("read single h5ad"):
        adata = sc.read_h5ad(h5ad)
        _print(f"[read] loaded cells={adata.n_obs:,}, genes={adata.n_vars:,}, obsm={list(adata.obsm.keys())}")
    _add_dataset_obs_column(
        adata,
        dataset=dataset,
        cancer_col=None if args.cancer_col.lower() == "auto" else args.cancer_col,
    )
    groupby = _ensure_groupby_column(
        adata,
        requested_groupby=args.groupby,
        make_cnv_leiden=args.make_cnv_leiden,
        use_rep=args.use_rep,
        leiden_resolution=args.leiden_resolution,
        neighbors_n=args.neighbors_n,
    )

    export_cnv_chromosome_clustermaps(
        adata,
        outdir=outdir,
        cancer_col=None if args.cancer_col.lower() == "auto" else args.cancer_col,
        groupby=groupby,
        use_rep=args.use_rep,
        window_size=args.window_size,
        step=args.step,
        cnv_type_threshold=args.threshold,
        exclude_chromosomes=exclude,
        max_cells_plot=args.max_cells_plot,
        make_builtin_infercnvpy_heatmap=args.builtin_infercnvpy_heatmap,
        dpi=args.dpi,
        progress=not args.quiet_progress,
    )


def _run_batch(args: argparse.Namespace) -> None:
    import gc
    import scanpy as sc

    h5ad_dir = Path(args.h5ad_dir)
    if not h5ad_dir.exists():
        raise FileNotFoundError(h5ad_dir)

    batch_outdir = Path(args.outdir) if args.outdir else h5ad_dir / "cnv_chromosome_clustermaps"
    batch_outdir.mkdir(parents=True, exist_ok=True)

    exclude = None if args.include_sex_chromosomes else ("chrX", "chrY")
    cancer_col = None if str(args.cancer_col).lower() == "auto" else args.cancer_col

    requested = _split_csv_arg(args.datasets)
    dataset_paths: list[tuple[str, Path]] = []

    if str(args.datasets).strip().lower() in {"auto", "*", "allh5ad", "all-h5ad"}:
        dataset_paths = _discover_auto_datasets(h5ad_dir, recursive=args.recursive)
    else:
        for dataset in requested:
            p = _find_dataset_h5ad(
                h5ad_dir,
                dataset,
                pattern=args.h5ad_pattern,
                recursive=args.recursive,
            )
            if p is None:
                print(f"[skip] {dataset}: no matching h5ad found in {h5ad_dir}")
                continue
            dataset_paths.append((dataset, p))

    if not dataset_paths:
        raise SystemExit("No input h5ad files found. Check --h5ad-dir, --datasets, and --h5ad-pattern.")

    print(f"[batch] h5ad_dir: {h5ad_dir}")
    print(f"[batch] outdir:   {batch_outdir}")
    print(f"[batch] datasets: {[d for d, _ in dataset_paths]}")

    successful = []
    failures = []

    dataset_iter = tqdm(dataset_paths, desc="datasets", unit="dataset", disable=args.quiet_progress)
    for dataset, h5ad in dataset_iter:
        try:
            dataset_iter.set_description(f"dataset {dataset}")
        except Exception:
            pass
        safe_dataset = _safe_name(dataset)
        dataset_outdir = batch_outdir / safe_dataset
        metadata_path = dataset_outdir / "tables" / "export_metadata.json"

        if args.skip_existing and metadata_path.exists():
            print(f"\n[skip] {dataset}: existing export found at {metadata_path}")
            successful.append(dataset)
            continue

        print("\n" + "=" * 96)
        print(f"[dataset] {dataset}")
        print(f"[read] {h5ad}")

        try:
            with _timed_step(f"read h5ad for {dataset}"):
                adata = sc.read_h5ad(h5ad)
                _print(f"[read] loaded cells={adata.n_obs:,}, genes={adata.n_vars:,}, obsm={list(adata.obsm.keys())}")
            _add_dataset_obs_column(adata, dataset=dataset, cancer_col=cancer_col)

            groupby = _ensure_groupby_column(
                adata,
                requested_groupby=args.groupby,
                make_cnv_leiden=args.make_cnv_leiden,
                use_rep=args.use_rep,
                leiden_resolution=args.leiden_resolution,
                neighbors_n=args.neighbors_n,
                cnv_leiden_rep=args.cnv_leiden_rep,
                cnv_svd_components=args.cnv_svd_components,
                progress=not args.quiet_progress,
            )

            export_cnv_chromosome_clustermaps(
                adata,
                outdir=dataset_outdir,
                cancer_col=cancer_col,
                groupby=groupby,
                use_rep=args.use_rep,
                window_size=args.window_size,
                step=args.step,
                cnv_type_threshold=args.threshold,
                exclude_chromosomes=exclude,
                max_cells_plot=args.max_cells_plot,
                make_builtin_infercnvpy_heatmap=args.builtin_infercnvpy_heatmap,
                dpi=args.dpi,
                progress=not args.quiet_progress,
            )

            successful.append(dataset)
            print(f"[done] {dataset}: {dataset_outdir}")

        except Exception as e:
            failures.append((dataset, type(e).__name__, str(e)))
            print(f"[ERROR] {dataset}: {type(e).__name__}: {e}")

        finally:
            try:
                del adata
            except Exception:
                pass
            gc.collect()

    if successful and not args.no_batch_summary:
        print("\n" + "=" * 96)
        print("[batch-summary] making cross-cancer summary plots")
        _make_batch_summary_from_exported_tables(
            batch_outdir=batch_outdir,
            successful=successful,
            dpi=args.dpi,
        )

    print("\n" + "=" * 96)
    print(f"[batch] successful: {successful}")
    if failures:
        print("[batch] failures:")
        for dataset, err_type, msg in failures:
            print(f"  - {dataset}: {err_type}: {msg}")
    print("[done]")


def main() -> None:
    global HEARTBEAT_SECONDS
    args = _parse_args()
    HEARTBEAT_SECONDS = int(args.heartbeat_seconds)
    if args.h5ad:
        _run_single_h5ad(args)
    else:
        _run_batch(args)


if __name__ == "__main__":
    main()
```
