## Supplemental_Code for "scOPE identifies which driver-associated expression programs transfer from bulk tumors to single cells": fix_aml_ensg_symbols_mapped_only.py

```
#!/usr/bin/env python
"""
fix_aml_ensg_symbols_mapped_only.py

Make an AML bulk expression table with ONLY readable gene-symbol columns.

Why this exists:
- The original AML bulk matrix uses Ensembl IDs.
- A GENCODE GTF maps most, but not every, ENSG ID to a current gene symbol.
- Leaving unmapped IDs or ENSG-like GTF placeholder gene_names in the symbolized table makes plots show mixed HGNC/ENSG labels.
- This script drops both unmapped ENSG columns and ENSG-like placeholder gene_names, then writes a clean symbol-only matrix.

Default input/output paths match Andrew's local scOPE tree.
"""

from __future__ import annotations

import argparse
import gzip
import re
from pathlib import Path

import pandas as pd


ENSG_RE = re.compile(r"^(ENSG\d+)(?:\.\d+)?$")


def strip_ensg_version(x: object) -> str:
    s = str(x).strip()
    m = ENSG_RE.match(s)
    return m.group(1) if m else s


def parse_gtf_attrs(attr: str) -> dict[str, str]:
    out = {}
    for part in str(attr).split(";"):
        part = part.strip()
        if not part or " " not in part:
            continue
        k, v = part.split(" ", 1)
        out[k] = v.strip().strip('"')
    return out


def load_gtf_map(gtf_path: Path) -> pd.DataFrame:
    opener = gzip.open if str(gtf_path).endswith(".gz") else open
    rows = []
    with opener(gtf_path, "rt", encoding="utf-8", errors="replace") as fh:
        for line in fh:
            if not line or line.startswith("#"):
                continue
            fields = line.rstrip("\n").split("\t")
            if len(fields) < 9 or fields[2] != "gene":
                continue
            attrs = parse_gtf_attrs(fields[8])
            gene_id = attrs.get("gene_id")
            gene_name = attrs.get("gene_name")
            gene_type = attrs.get("gene_type") or attrs.get("gene_biotype") or ""
            if gene_id and gene_name:
                rows.append({
                    "ensembl_id": strip_ensg_version(gene_id),
                    "gene_symbol": str(gene_name),
                    "gene_type": str(gene_type),
                })
    m = pd.DataFrame(rows).drop_duplicates("ensembl_id")
    if m.empty:
        raise RuntimeError(f"No gene mappings parsed from {gtf_path}")
    return m


def make_unique(names: list[str]) -> list[str]:
    seen = {}
    out = []
    for name in names:
        name = str(name).strip()
        n = seen.get(name, 0)
        seen[name] = n + 1
        out.append(name if n == 0 else f"{name}__{n + 1}")
    return out


def main() -> None:
    default_root = Path(r"C:\Users\ashfo\OneDrive\Desktop\Pathways+Omics\scOPE\scOPE\data")
    ap = argparse.ArgumentParser()
    ap.add_argument(
        "--input",
        default=str(default_root / "scOPE_reviewer_derived" / "AML_bulk_expression_samples_by_genes.tsv.gz"),
        help="Original AML bulk expression matrix, samples x genes.",
    )
    ap.add_argument(
        "--gtf",
        default=str(default_root / "gencode.v44.annotation.gtf.gz"),
        help="GENCODE GTF used to map ENSG IDs to gene symbols.",
    )
    ap.add_argument(
        "--output",
        default=str(default_root / "scOPE_reviewer_derived" / "AML_bulk_expression_samples_by_genes_SYMBOLS_MAPPED_ONLY.tsv.gz"),
        help="Output expression matrix with only mapped gene-symbol columns.",
    )
    ap.add_argument(
        "--unmapped-out",
        default=None,
        help="Optional CSV path for dropped unmapped genes. Default: next to output.",
    )
    ap.add_argument(
        "--collapse-duplicates",
        choices=["none", "mean"],
        default="none",
        help=(
            "How to handle duplicate gene symbols. "
            "'none' keeps duplicates readable as SYMBOL__2; 'mean' averages duplicate symbol columns."
        ),
    )
    args = ap.parse_args()

    inp = Path(args.input)
    gtf = Path(args.gtf)
    out = Path(args.output)
    unmapped_out = Path(args.unmapped_out) if args.unmapped_out else out.with_name(out.name.replace(".tsv.gz", "_dropped_unmapped_genes.csv"))

    print(f"[read] expression: {inp}")
    df = pd.read_csv(inp, sep="\t", index_col=0)
    print(f"[read] matrix shape: {df.shape[0]:,} samples x {df.shape[1]:,} genes")

    print(f"[map] loading GTF: {gtf}")
    gtf_map = load_gtf_map(gtf).set_index("ensembl_id")
    print(f"[map] GTF mappings: {len(gtf_map):,}")

    old_cols = pd.Index(df.columns.astype(str))
    ens_ids = pd.Index([strip_ensg_version(c) for c in old_cols])

    mapped_mask = ens_ids.isin(gtf_map.index)

    # Some GENCODE records technically map, but their gene_name is itself an ENSG-like
    # placeholder rather than a readable HGNC-style symbol. Drop those too so plots
    # are truly symbol-only.
    symbols_all = pd.Series(index=old_cols, dtype="object")
    gene_types_all = pd.Series(index=old_cols, dtype="object")
    mapped_ens = ens_ids[mapped_mask]
    symbols_all.loc[old_cols[mapped_mask]] = gtf_map.loc[mapped_ens, "gene_symbol"].astype(str).to_numpy()
    gene_types_all.loc[old_cols[mapped_mask]] = gtf_map.loc[mapped_ens, "gene_type"].astype(str).to_numpy()

    symbol_is_ensg = symbols_all.fillna("").astype(str).str.match(ENSG_RE)
    readable_mask = mapped_mask & (~symbol_is_ensg.to_numpy())

    n_mapped = int(mapped_mask.sum())
    n_unmapped = int((~mapped_mask).sum())
    n_placeholder = int((mapped_mask & symbol_is_ensg.to_numpy()).sum())
    n_readable = int(readable_mask.sum())
    print(f"[map] GTF-mapped: {n_mapped:,}/{len(old_cols):,}")
    print(f"[map] dropping unmapped: {n_unmapped:,}")
    print(f"[map] dropping ENSG-like GTF gene_name placeholders: {n_placeholder:,}")
    print(f"[map] keeping readable symbols: {n_readable:,}")

    # Save dropped IDs so nothing is hidden.
    dropped_unmapped = pd.DataFrame({
        "old_gene_id": old_cols[~mapped_mask],
        "ensembl_id_no_version": ens_ids[~mapped_mask],
        "gene_symbol_from_gtf": "",
        "gene_type": "",
        "reason": "not_found_in_gtf_gene_records",
    })
    dropped_placeholder = pd.DataFrame({
        "old_gene_id": old_cols[mapped_mask & symbol_is_ensg.to_numpy()],
        "ensembl_id_no_version": ens_ids[mapped_mask & symbol_is_ensg.to_numpy()],
        "gene_symbol_from_gtf": symbols_all.loc[old_cols[mapped_mask & symbol_is_ensg.to_numpy()]].astype(str).to_numpy(),
        "gene_type": gene_types_all.loc[old_cols[mapped_mask & symbol_is_ensg.to_numpy()]].astype(str).to_numpy(),
        "reason": "gtf_gene_name_is_ensg_placeholder",
    })
    dropped = pd.concat([dropped_unmapped, dropped_placeholder], ignore_index=True)
    unmapped_out.parent.mkdir(parents=True, exist_ok=True)
    dropped.to_csv(unmapped_out, index=False)
    print(f"[write] dropped gene list: {unmapped_out}")

    keep_old_cols = old_cols[readable_mask]
    keep_ens_ids = ens_ids[readable_mask]
    symbols = gtf_map.loc[keep_ens_ids, "gene_symbol"].astype(str).to_list()

    clean = df.loc[:, keep_old_cols].copy()

    if args.collapse_duplicates == "mean":
        clean.columns = symbols
        # For log-scale expression matrices, mean is safer than summing duplicate-symbol columns.
        clean = clean.T.groupby(level=0, sort=False).mean().T
        print(f"[dedupe] collapsed duplicate symbols by mean; final genes={clean.shape[1]:,}")
    else:
        clean.columns = make_unique(symbols)
        print(f"[dedupe] kept duplicate symbols with __2 suffixes; final genes={clean.shape[1]:,}")

    # Sanity check: no Ensembl IDs remain as column names.
    still_ensg = [c for c in clean.columns.astype(str) if ENSG_RE.match(c)]
    if still_ensg:
        raise RuntimeError(f"Still found ENSG-like output columns: {still_ensg[:20]}")

    out.parent.mkdir(parents=True, exist_ok=True)
    print(f"[write] clean symbol-only matrix: {out}")
    clean.to_csv(out, sep="\t", compression="infer")

    print("\n[done]")
    print("Update AML expression in your config to:")
    print(str(out))
    print("\nThen delete old AML Fig2 outputs and rerun bulk/interpret:")
    print(r"rmdir /s /q scope_analysis_outputs\fig2_bulk\AML")
    print(r"python drivers\run_all.py --config drivers\config_canonical_v3.json --only bulk,interpret,report")


if __name__ == "__main__":
    main()
```
