## Supplemental_Code for "scOPE identifies which driver-associated expression programs transfer from bulk tumors to single cells": make_infercnvpy_cnvs.py

```
#!/usr/bin/env python
"""
make_infercnvpy_cnvs.py

Run infercnvpy CNV inference on one or more scRNA AnnData objects and export the
tidy CNV call table expected by the scOPE manuscript analysis suite:

    dataset, barcode, method, cnv_score, malignant_call

Default paths are tailored to Andrew's local scOPE tree, but all key paths can be
overridden from the command line.

Examples
--------
# Quick smoke test on 3k cells per cohort:
python make_infercnvpy_cnvs.py --datasets AML --max-cells 3000 --overwrite

# Full run on all configured datasets:
python make_infercnvpy_cnvs.py --datasets AML,BRCA,CRC,GBM,LUAD,PAAD,SKCM --overwrite

# Use less RAM:
python make_infercnvpy_cnvs.py --datasets AML --chunksize 2000 --n-jobs 4 --overwrite

# After it looks good, optionally update the suite CNV table:
python make_infercnvpy_cnvs.py --datasets AML,BRCA,CRC,GBM,LUAD,PAAD,SKCM --update-suite-cnv-csv --overwrite
"""

from __future__ import annotations

import argparse
import gc
import json
import os
import re
import shutil
import sys
import threading
import time
from pathlib import Path
from typing import Optional

import numpy as np
import pandas as pd
import scipy.sparse as sp

try:
    import anndata as ad
except Exception as e:
    raise SystemExit("Missing anndata. Install with: pip install anndata") from e

try:
    import scanpy as sc
except Exception:
    sc = None

try:
    import infercnvpy as cnv
except Exception as e:
    raise SystemExit("Missing infercnvpy. Install with: pip install infercnvpy") from e


DEFAULT_ROOT = Path(r"C:\Users\ashfo\OneDrive\Desktop\Pathways+Omics\scOPE\scOPE\data")

# Prefer expression/scored H5ADs that are already in the user's tree.
# If you regenerate new scored H5ADs after changing SVD settings, pass those with
# --manifest, or edit this dictionary.
DEFAULT_DATASETS = {
    "AML": {
        "path": DEFAULT_ROOT / "scOPE_reviewer_derived" / "AML_scored_with_muttranscript_eval_labels.h5ad",
        "sample_col": "Sample",
        "patient_col": "Sample",
        "patient_strip_regex": r"-D\d+$",
        "celltype_col": "CellType",
        "reference_regex": r"^(?:T|B|NK|CTL)$",
    },
    "BRCA": {
        "path": DEFAULT_ROOT / "models" / "BRCA_scored_z_score_bulk_with_eval_labels.h5ad",
        "sample_col": "sample_id",
        "patient_col": "sample_id",
        "celltype_col": "celltype_major",
        # Strict lymphoid/plasma reference; excludes Cancer Epithelial and Normal Epithelial.
        "reference_regex": r"^(?:T[-_ ]?cells?|B[-_ ]?cells?|NK[-_ ]?cells?|Plasmablasts?|Plasma[-_ ]?cells?)$",
    },
    "CRC": {
        "path": DEFAULT_ROOT / "models" / "CRC_scored_z_score_bulk_with_eval_labels.h5ad",
        "sample_col": "Sample",
        "patient_col": "Patient",
        "celltype_col": "Cell_type",
        # Conservative non-epithelial reference.
        "reference_regex": (
            r"^(?:"
            r"T|T[-_ ]?cells?|CD4[-_ ]?T[-_ ]?cells?|CD8[-_ ]?T[-_ ]?cells?|"
            r"B|B[-_ ]?cells?|NK|NK[-_ ]?cells?|"
            r"Plasma|Plasma[-_ ]?cells?|Plasmablasts?|"
            r"Myeloid|Monocytes?|Macrophages?|Dendritic[-_ ]?cells?|DCs?|"
            r"Mast|Mast[-_ ]?cells?|"
            r"Endothelial|Endothelial[-_ ]?cells?|"
            r"Fibroblasts?|Stromal|Stromal[-_ ]?cells?"
            r")$"
        ),
    },
    "GBM": {
        "path": DEFAULT_ROOT / "models" / "GBM_scored_z_score_bulk_with_eval_labels.h5ad",
        "sample_col": "sample",
        "patient_col": "patient",
        # v10.3.3 FIX (supersedes the earlier "celltype" pointer, which does not exist
        # on the scored GBM h5ad -> _standardize_obs silently fell back to "no usable
        # celltype column; using all-cell reference", i.e. 0 reference cells, which is
        # why GBM previously had to be forced through with --allow-all-cell-reference
        # and produced degenerate calls (~2% malignant vs 10-26% in every other cancer).
        # The scored object instead carries `celltype_coarse` (12 categories) from a
        # generic marker-based classifier with no glioma-specific "malignant" label.
        # Malignant GBM cells were absorbed into non-immune buckets, most heavily
        # "stromal" (9,188/24,131 = 38% of ALL cells) -- almost certainly MES-like
        # malignant glioma cells read as Fibroblast/CAF by marker overlap (a known
        # GBM cell-state confound; Neftel et al. 2019), not real brain stroma.
        # epithelial/erythroid/hspc_prog/unknown are likewise ambiguous/contamination,
        # not a validated non-malignant population.
        "celltype_col": "celltype_coarse",
        # Restricted to the seven immune/vascular categories that ARE reliably
        # non-malignant. This exactly reproduces the dataset's own precomputed
        # `is_reference` obs column: is_reference==True for 100% of lymphoid_t/
        # lymphoid_b/lymphoid_nk/plasma/dendritic/endothelial/myeloid cells and
        # ==False for 100% of stromal/epithelial/erythroid/hspc_prog/unknown cells
        # (verified by groupby(celltype_coarse)["is_reference"].mean() -> exactly
        # 0.0 or 1.0 per category, no partial overlap).
        # Do NOT add stromal/fibroblast back into the GBM reference without marker-
        # level confirmation (e.g. against inferCNV baseline) that a given "stromal"
        # cell is truly non-malignant rather than an MES-like tumor cell.
        "reference_regex": (
            r"^(?:lymphoid_t|lymphoid_b|lymphoid_nk|plasma|dendritic|endothelial|myeloid)$"
        ),
    },
    "LUAD": {
        "path": DEFAULT_ROOT / "models" / "LUAD_scored_z_score_bulk_with_eval_labels_WITH_LUAD_OBS.h5ad",
        "sample_col": "sample",
        "patient_col": "patient",
        "celltype_col": "celltype",
        "reference_regex": (
            r"^(?:"
            r"T|T[-_ ]?cells?|CD4[-_ ]?T[-_ ]?cells?|CD8[-_ ]?T[-_ ]?cells?|"
            r"B|B[-_ ]?cells?|NK|NK[-_ ]?cells?|"
            r"Plasma|Plasma[-_ ]?cells?|Plasmablasts?|"
            r"Myeloid cells|Myeloid|Monocytes?|Macrophages?|Alveolar[-_ ]?Macrophages?|"
            r"Dendritic[-_ ]?cells?|DCs?|"
            r"Mast|Mast[-_ ]?cells?|"
            r"Endothelial cells|Endothelial|Endothelial[-_ ]?cells?|"
            r"Fibroblasts?|Stromal|Stromal[-_ ]?cells?|"
            r"Normal[-_ ]?Epithelial|Epithelial[-_ ]?normal|"
            r"Alveolar|AT1|AT2|Club|Ciliated"
            r")$"
        ),
    },
    "PAAD": {
        "path": DEFAULT_ROOT / "models" / "PAAD_scored_z_score_bulk_with_eval_labels.h5ad",
        "sample_col": "sample",
        "patient_col": "patient",
        "celltype_col": "celltype",
        # PAAD labels are fine-grained and often have suffixes like ".7".
        # Use immune/stromal/endothelial/acinar/endocrine references.
        # Explicitly avoid ductal/cancer clone labels, which may carry tumor CNVs.
        "reference_regex": (
            r"(?i)"
            r"^(?!.*(?:cancer|clone|ductal|malignant|tumou?r|neoplastic))"
            r".*(?:"
            r"t[-_ ]?cells?|cd4|cd8|"
            r"b[-_ ]?cells?|nk|"
            r"plasma|plasmablast|"
            r"myeloid|mono|monocyte|macrophage|dendritic|dc|mast|"
            r"endothelial|"
            r"fibroblast|caf|stellate|stromal|"
            r"acinar|endocrine|islet|alpha|beta|delta"
            r").*$"
        ),
    },
    "SKCM": {
        "path": DEFAULT_ROOT / "models" / "SKCM_scored_z_score_bulk_with_eval_labels.h5ad",
        "sample_col": "samples",
        "patient_col": "patient",
        "celltype_col": "celltype",
        # Conservative non-melanoma reference.
        "reference_regex": (
            r"^(?:"
            r"T|T[-_ ]?cells?|CD4[-_ ]?T[-_ ]?cells?|CD8[-_ ]?T[-_ ]?cells?|"
            r"B|B[-_ ]?cells?|NK|NK[-_ ]?cells?|"
            r"Plasma|Plasma[-_ ]?cells?|Plasmablasts?|"
            r"Myeloid|Monocytes?|Macrophages?|Dendritic[-_ ]?cells?|DCs?|"
            r"Mast|Mast[-_ ]?cells?|"
            r"Endothelial|Endothelial[-_ ]?cells?|"
            r"Fibroblasts?|CAFs?|Stromal|Stromal[-_ ]?cells?"
            r")$"
        ),
    },
}


def _safe_name(x: str) -> str:
    return re.sub(r"[^A-Za-z0-9_.-]+", "_", str(x)).strip("_")


def _heartbeat(stop_event: threading.Event, label: str, interval: int = 60) -> None:
    start = time.time()
    while not stop_event.wait(interval):
        msg = f"[heartbeat] {label}: still running, elapsed={((time.time() - start) / 60):.1f} min"
        try:
            import psutil
            proc = psutil.Process(os.getpid())
            rss_gb = proc.memory_info().rss / (1024**3)
            msg += f", RSS={rss_gb:.2f} GB"
        except Exception:
            pass
        print(msg, flush=True)


def _read_manifest(path: Optional[str]) -> dict:
    if not path:
        return DEFAULT_DATASETS
    p = Path(path)
    if not p.exists():
        raise FileNotFoundError(p)
    data = json.loads(p.read_text())
    out = {}
    for k, v in data.items():
        vv = dict(v)
        vv["path"] = Path(vv["path"])
        out[k] = vv
    return out


def _guess_id_type(var_names: pd.Index) -> tuple[str, Optional[str]]:
    vals = pd.Series(var_names.astype(str)).head(2000)
    ens = vals.str.match(r"^ENSG\d+(?:\.\d+)?$").mean()
    if ens > 0.5:
        return "gene_id", "ensembl_id_no_version"
    return "gene_name", None


def _strip_ens_version(x: object) -> str:
    return re.sub(r"\.\d+$", "", str(x).strip())


def _add_genomic_positions(adata, gtf: Path) -> None:
    if {"chromosome", "start", "end"}.issubset(adata.var.columns):
        ok = (
            adata.var["chromosome"].notna()
            & adata.var["start"].notna()
            & adata.var["end"].notna()
        )
        if ok.sum() > 1000:
            print(f"[gene-pos] existing genomic positions detected for {int(ok.sum()):,} genes")
            return

    gtf_gene_id, adata_gene_id = _guess_id_type(adata.var_names)
    print(f"[gene-pos] annotating from GTF using gtf_gene_id={gtf_gene_id!r}")

    if adata_gene_id:
        adata.var[adata_gene_id] = [_strip_ens_version(x) for x in adata.var_names]
    else:
        adata.var["gene_symbol_for_gtf"] = adata.var_names.astype(str)
        adata_gene_id = "gene_symbol_for_gtf"

    cnv.io.genomic_position_from_gtf(
        str(gtf),
        adata=adata,
        gtf_gene_id=gtf_gene_id,
        adata_gene_id=adata_gene_id,
        inplace=True,
    )


def _force_chr_and_filter(adata, drop_chr_m: bool = True):
    required = {"chromosome", "start", "end"}
    missing = required - set(adata.var.columns)
    if missing:
        raise ValueError(f"adata.var missing required genomic position columns: {missing}")

    chrom = (
        adata.var["chromosome"].astype("string").str.strip()
        .str.replace(r"^chr", "", regex=True, case=False)
        .str.replace(r"\.0$", "", regex=True)
    )
    chrom = chrom.replace({"MT": "M", "Mt": "M", "m": "M", "chrM": "M", "chrMT": "M"})
    canonical = [str(i) for i in range(1, 23)] + ["X", "Y", "M"]
    keep = chrom.isin(canonical)

    adata.var["start"] = pd.to_numeric(adata.var["start"], errors="coerce")
    adata.var["end"] = pd.to_numeric(adata.var["end"], errors="coerce")
    keep &= adata.var["start"].notna() & adata.var["end"].notna()

    if drop_chr_m:
        keep &= chrom.ne("M")

    before = adata.n_vars
    adata = adata[:, keep.to_numpy()].copy()
    chrom = chrom.loc[adata.var_names]
    adata.var["chromosome"] = ("chr" + chrom.astype(str)).to_numpy()
    adata.var["start"] = pd.to_numeric(adata.var["start"], errors="coerce").astype(int)
    adata.var["end"] = pd.to_numeric(adata.var["end"], errors="coerce").astype(int)
    adata.var = adata.var.sort_values(["chromosome", "start", "end"])
    adata = adata[:, adata.var.index].copy()

    print(f"[gene-pos] kept {adata.n_vars:,}/{before:,} genes with canonical genomic positions")
    if adata.n_vars < 3000:
        raise ValueError(
            f"Only {adata.n_vars} genes with genomic positions. This is probably too few; "
            "check whether var_names are symbols vs ENSG and whether the GTF matches."
        )
    return adata


def _standardize_obs(adata, spec: dict, dataset: str):
    obs = adata.obs

    # sample
    sample_col = spec.get("sample_col")
    if sample_col and sample_col in obs.columns:
        adata.obs["sample"] = obs[sample_col].astype(str).to_numpy()
    elif "sample" not in obs.columns:
        adata.obs["sample"] = adata.obs_names.astype(str)

    # patient
    patient_col = spec.get("patient_col")
    if patient_col and patient_col in obs.columns:
        patient = obs[patient_col].astype(str)
    elif "patient" in obs.columns:
        patient = obs["patient"].astype(str)
    else:
        patient = adata.obs["sample"].astype(str)

    strip = spec.get("patient_strip_regex")
    if strip:
        patient = patient.str.replace(strip, "", regex=True)

    adata.obs["patient"] = patient.to_numpy()

    # celltype
    celltype_col = spec.get("celltype_col")
    if celltype_col and celltype_col in obs.columns:
        adata.obs["celltype"] = obs[celltype_col].astype(str).to_numpy()
    elif "celltype" in obs.columns:
        adata.obs["celltype"] = obs["celltype"].astype(str).to_numpy()
    elif "cell_type" in obs.columns:
        adata.obs["celltype"] = obs["cell_type"].astype(str).to_numpy()
    else:
        print(f"[obs] WARNING: {dataset} has no usable celltype column; using all-cell reference.")
        adata.obs["celltype"] = "unknown"

    ref_re = spec.get("reference_regex") or ""
    if ref_re:
        is_ref = adata.obs["celltype"].astype(str).str.contains(ref_re, case=False, regex=True, na=False)
    else:
        is_ref = pd.Series(False, index=adata.obs_names)

    adata.obs["cnv_reference"] = np.where(is_ref.to_numpy(), "reference", "other")
    print(f"[obs] {dataset}: reference cells = {int(is_ref.sum()):,}/{adata.n_obs:,}")
    print("[obs] top celltypes:", adata.obs["celltype"].astype(str).value_counts().head(12).to_dict())

    return int(is_ref.sum())


def _maybe_normalize_log1p(adata, do_it: bool) -> None:
    if not do_it:
        return
    if sc is None:
        raise RuntimeError("scanpy is required for --normalize-log1p")
    print("[preprocess] normalize_total(target_sum=1e4) + log1p")
    sc.pp.normalize_total(adata, target_sum=1e4)
    sc.pp.log1p(adata)


def _cnv_score_from_obsm(adata, key: str) -> np.ndarray:
    """
    infercnvpy may store the CNV matrix under either:
      - adata.obsm[key]
      - adata.obsm["X_" + key]

    For key_added="cnv", the common output key is adata.obsm["X_cnv"].
    """
    candidate_keys = [key, f"X_{key}", "cnv", "X_cnv", "infercnv", "X_infercnv"]
    found_key = None
    for k in candidate_keys:
        if k in adata.obsm:
            found_key = k
            break

    if found_key is None:
        available = list(adata.obsm.keys())
        raise KeyError(
            f"No infercnvpy CNV matrix found. Tried {candidate_keys}. "
            f"Available adata.obsm keys after infercnvpy: {available}"
        )

    print(f"[score] using adata.obsm[{found_key!r}] for CNV score")
    X = adata.obsm[found_key]
    if sp.issparse(X):
        return np.asarray(abs(X).mean(axis=1)).ravel()
    return np.nanmean(np.abs(np.asarray(X)), axis=1)


def run_one(dataset: str, spec: dict, args) -> pd.DataFrame:
    in_path = Path(spec["path"])
    if not in_path.exists():
        raise FileNotFoundError(in_path)

    outdir = Path(args.outdir)
    outdir.mkdir(parents=True, exist_ok=True)
    h5_out = outdir / f"{_safe_name(dataset)}_infercnvpy.h5ad"
    calls_out = outdir / f"cnv_calls_{_safe_name(dataset)}.csv"

    if calls_out.exists() and h5_out.exists() and not args.overwrite:
        print(f"[skip] {dataset}: outputs exist. Use --overwrite to rerun.")
        return pd.read_csv(calls_out)

    print("\n" + "=" * 88)
    print(f"[load] {dataset}: {in_path}")
    adata = ad.read_h5ad(in_path)
    if getattr(adata, "isbacked", False):
        adata = adata.to_memory()
    adata.var_names = adata.var_names.astype(str)

    if args.max_cells and adata.n_obs > args.max_cells:
        rng = np.random.default_rng(args.random_state)
        idx = np.sort(rng.choice(adata.n_obs, size=args.max_cells, replace=False))
        adata = adata[idx].copy()
        print(f"[subset] {dataset}: randomly kept {adata.n_obs:,} cells for smoke test")

    _maybe_normalize_log1p(adata, args.normalize_log1p)
    n_ref = _standardize_obs(adata, spec, dataset)
    _add_genomic_positions(adata, Path(args.gtf))
    adata = _force_chr_and_filter(adata)

    reference_key = "cnv_reference" if n_ref >= args.min_reference_cells else None
    reference_cat = ["reference"] if reference_key else None
    if reference_key is None:
        if not args.allow_all_cell_reference:
            raise ValueError(
                f"{dataset}: only {n_ref} reference cells found. Fix reference_regex/celltype_col "
                "or rerun with --allow-all-cell-reference."
            )
        print(f"[ref] {dataset}: using all-cell average reference")

    print(
        f"[infercnvpy] {dataset}: window_size={args.window_size}, step={args.step}, "
        f"chunksize={args.chunksize}, n_jobs={args.n_jobs}, key_added={args.key_added!r}"
    )

    stop = threading.Event()
    t = threading.Thread(
        target=_heartbeat,
        args=(stop, f"infercnvpy {dataset}", args.heartbeat_seconds),
        daemon=True,
    )
    t.start()

    try:
        cnv.tl.infercnv(
            adata,
            reference_key=reference_key,
            reference_cat=reference_cat,
            lfc_clip=args.lfc_clip,
            window_size=args.window_size,
            step=args.step,
            dynamic_threshold=None if str(args.dynamic_threshold).lower() == "none" else float(args.dynamic_threshold),
            exclude_chromosomes=tuple(args.exclude_chromosomes.split(",")) if args.exclude_chromosomes else (),
            chunksize=args.chunksize,
            n_jobs=args.n_jobs,
            layer=args.layer,
            key_added=args.key_added,
            calculate_gene_values=args.calculate_gene_values,
            inplace=True,
        )
    finally:
        stop.set()
        t.join(timeout=2)

    score = _cnv_score_from_obsm(adata, args.key_added)
    adata.obs["cnv_score"] = score

    if reference_key:
        ref_mask = adata.obs["cnv_reference"].eq("reference").to_numpy()
        threshold = float(np.nanquantile(score[ref_mask], args.malignant_quantile))
        threshold_source = f"reference_q{args.malignant_quantile}"
    else:
        threshold = float(np.nanquantile(score, args.malignant_quantile))
        threshold_source = f"all_cells_q{args.malignant_quantile}"

    malignant = score > threshold
    adata.obs["malignant_call"] = malignant
    adata.obs["cnv_threshold"] = threshold
    adata.obs["cnv_threshold_source"] = threshold_source
    adata.obs["cnv_method"] = "infercnvpy"

    calls = pd.DataFrame({
        "dataset": dataset,
        "barcode": adata.obs_names.astype(str),
        "method": "infercnvpy",
        "cnv_score": score.astype(float),
        "malignant_call": malignant.astype(bool),
        "cnv_threshold": threshold,
        "cnv_threshold_source": threshold_source,
        "sample": adata.obs["sample"].astype(str).to_numpy(),
        "patient": adata.obs["patient"].astype(str).to_numpy(),
        "celltype": adata.obs["celltype"].astype(str).to_numpy(),
        "is_reference": adata.obs["cnv_reference"].eq("reference").to_numpy(),
    })

    print(f"[call] {dataset}: threshold={threshold:.5g} ({threshold_source}); calls={calls['malignant_call'].value_counts().to_dict()}")
    print(f"[write] {calls_out}")
    calls.to_csv(calls_out, index=False)

    print(f"[write] {h5_out}")
    adata.write_h5ad(h5_out)

    del adata
    gc.collect()
    return calls


def main() -> None:
    ap = argparse.ArgumentParser()
    ap.add_argument("--root", default=str(DEFAULT_ROOT), help="scOPE data root")
    ap.add_argument("--gtf", default=str(DEFAULT_ROOT / "gencode.v44.annotation.gtf.gz"))
    ap.add_argument("--outdir", default=str(DEFAULT_ROOT / "scope_robust_real_inputs" / "infercnvpy_new"))
    ap.add_argument("--manifest", default=None, help="Optional JSON dataset manifest overriding defaults")
    ap.add_argument("--datasets", default="AML", help="Comma-separated dataset names, e.g. AML,BRCA,CRC")
    ap.add_argument("--overwrite", action="store_true")
    ap.add_argument("--update-suite-cnv-csv", action="store_true", help="Also copy combined calls to scope_robust_real_inputs/cnv_calls.csv")

    ap.add_argument("--normalize-log1p", action="store_true", help="Use only if input X is raw counts")
    ap.add_argument("--max-cells", type=int, default=0, help="Smoke-test subset size; 0 = all cells")
    ap.add_argument("--min-reference-cells", type=int, default=50)
    ap.add_argument("--allow-all-cell-reference", action="store_true")
    ap.add_argument("--random-state", type=int, default=42)

    ap.add_argument("--window-size", type=int, default=100)
    ap.add_argument("--step", type=int, default=10)
    ap.add_argument("--lfc-clip", type=float, default=3.0)
    ap.add_argument("--dynamic-threshold", default="1.5", help="float or None")
    ap.add_argument("--exclude-chromosomes", default="chrX,chrY")
    ap.add_argument("--chunksize", type=int, default=5000)
    ap.add_argument("--n-jobs", type=int, default=1)
    ap.add_argument("--layer", default=None)
    ap.add_argument("--calculate-gene-values", action="store_true")
    ap.add_argument("--key-added", default="cnv")
    ap.add_argument("--malignant-quantile", type=float, default=0.95)
    ap.add_argument("--heartbeat-seconds", type=int, default=60)

    args = ap.parse_args()

    data_root = Path(args.root)

    manifest = _read_manifest(args.manifest)
    requested = [x.strip() for x in args.datasets.split(",") if x.strip()]
    all_calls = []

    for dataset in requested:
        if dataset not in manifest:
            print(f"[skip] unknown dataset {dataset!r}. Available: {sorted(manifest)}")
            continue
        try:
            calls = run_one(dataset, manifest[dataset], args)
            all_calls.append(calls)
        except Exception as e:
            print(f"[ERROR] {dataset}: {type(e).__name__}: {e}", file=sys.stderr)
            if len(requested) == 1:
                raise

    if all_calls:
        combined = pd.concat(all_calls, ignore_index=True)
        outdir = Path(args.outdir)
        combined_out = outdir / "cnv_calls_infercnvpy_new_all.csv"
        combined.to_csv(combined_out, index=False)
        print(f"\n[write] combined: {combined_out}")

        if args.update_suite_cnv_csv:
            suite_out = data_root / "scope_robust_real_inputs" / "cnv_calls.csv"
            backup = suite_out.with_suffix(".csv.BACKUP_before_infercnvpy_new")
            if suite_out.exists() and not backup.exists():
                shutil.copy2(suite_out, backup)
                print(f"[backup] existing suite CNV calls -> {backup}")
            combined[["dataset", "barcode", "method", "cnv_score", "malignant_call"]].to_csv(suite_out, index=False)
            print(f"[write] suite CNV table: {suite_out}")

    print("[done]")


if __name__ == "__main__":
    main()
```
