## Supplemental_Code for "scOPE identifies which driver-associated expression programs transfer from bulk tumors to single cells": make_sc_patient_status.py

```
#!/usr/bin/env python
"""Create/validate single-cell patient-level mutation-status files.

This is intentionally NOT a bulk-cohort converter.

The output used by the plotting/pancancer code is a tidy CSV:

    patient,mutation,label
    CID3586,TP53,1
    CID3586,NPM1,0

Rules:
  1. Patient/sample IDs are always taken from the scRNA AnnData obs for each
     configured single-cell cohort.
  2. Mutation labels are taken from either:
       a) true per-cell genotype/status columns already present in the scRNA H5AD,
          aggregated to patient level, OR
       b) a manually curated/publication metadata table for that exact scRNA cohort.
  3. Existing patient_status.csv files are only accepted if they overlap the scRNA
     patient IDs. Bulk reviewer/TCGA matrices with unrelated sample IDs are rejected.
  4. If no usable labels exist, the script writes templates for manual curation.

Examples:
  python make_sc_patient_status.py --config drivers/config_canonical_v3.json
  python make_sc_patient_status.py --config drivers/config_canonical_v3.json --overwrite
  python make_sc_patient_status.py --config drivers/config_canonical_v3.json --manual-root C:/path/to/manual_tables --overwrite
"""
from __future__ import annotations

import argparse
import json
import re
import shutil
from dataclasses import dataclass
from pathlib import Path
from typing import Iterable

import numpy as np
import pandas as pd

try:
    import anndata as ad
except Exception as e:  # pragma: no cover
    ad = None
    _ANNDATA_IMPORT_ERROR = e
else:
    _ANNDATA_IMPORT_ERROR = None

try:
    from scope_analysis.data import choose_patient_column as _choose_patient_column
except Exception:  # allow this standalone helper to run before package import works
    _choose_patient_column = None


TRUTH_RE = re.compile(r"(genotype|mutant|mutation|mut_status|status|variant|call|label|truth|snv)", re.I)
PRED_RE = re.compile(r"(mutation_prob|mutation_calib|mutation_adj|mutation_resid|mutation_refpct|_prob|_score|^score|pred|probability|resid|calib|refpct|adj)", re.I)
PATIENT_CANDIDATES = [
    "patient", "Patient", "patient_id", "PatientID", "patientID",
    "donor", "Donor", "donor_id", "case", "Case", "case_id", "CaseID",
    "sample", "Sample", "sample_id", "SampleID", "orig.ident", "orig_ident",
    "individual", "Individual", "subject", "Subject", "library", "sample_name",
]
MUTATION_COL_CANDIDATES = ["mutation", "Mutation", "gene", "Gene", "driver", "Driver", "alteration", "Alteration"]
LABEL_COL_CANDIDATES = ["label", "Label", "status", "Status", "mutated", "Mutated", "is_mutant", "mutation_status", "call", "Call"]
METADATA_LIKE = {
    "patient", "patient_id", "sample", "sample_id", "barcode", "cell", "cell_id",
    "cancer", "cohort", "dataset", "celltype", "cell_type", "timepoint", "day",
    "tumor", "tissue", "orig.ident", "orig_ident", "library", "batch",
}


@dataclass
class CohortAudit:
    cohort: str
    cancer: str
    h5ad: str
    patient_col: str
    n_cells: int
    n_patients: int
    n_drivers: int
    status_source: str
    wrote_patient_status: bool
    status_path: str
    template_path: str
    n_status_rows: int
    n_mut_rows: int
    n_wt_rows: int
    existing_overlap: str
    warning: str


def pth(x: str | Path | None) -> Path | None:
    if not x:
        return None
    return Path(str(x).replace("\\", "/"))


def norm_key(x) -> str:
    return re.sub(r"[^A-Za-z0-9]+", "", str(x).strip()).upper()


def coerce_label(x):
    if pd.isna(x):
        return np.nan
    if isinstance(x, (bool, np.bool_)):
        return int(bool(x))
    if isinstance(x, (int, float, np.integer, np.floating)):
        if not np.isfinite(x):
            return np.nan
        return int(float(x) > 0)
    s = str(x).strip()
    sl = s.lower()
    if sl in {"", "na", "nan", "none", "unknown", "unk", "not_available", "not available"}:
        return np.nan
    if sl in {"1", "true", "t", "yes", "y", "mut", "mutant", "mutated", "pos", "positive", "+", "present"}:
        return 1
    if sl in {"0", "false", "f", "no", "n", "wt", "wildtype", "wild-type", "wild type", "neg", "negative", "-", "absent"}:
        return 0
    # strings like KRAS_G12D / BRAF V600E should count as mutant if they are in a status cell.
    if re.search(r"\b(wt|wild[ -]?type)\b", sl):
        return 0
    if re.search(r"[A-Z][0-9]+[A-Z]|V600E|G12|G13|Q61|R132|R172|frameshift|nonsense|missense", s, re.I):
        return 1
    try:
        return int(float(sl) > 0)
    except Exception:
        return np.nan


def find_first(cols: Iterable[str], candidates: Iterable[str]) -> str | None:
    cols_set = set(cols)
    for c in candidates:
        if c in cols_set:
            return c
    low = {str(c).lower(): c for c in cols}
    for c in candidates:
        if c.lower() in low:
            return low[c.lower()]
    return None


def infer_patient_col(obs: pd.DataFrame, configured: str | None = None) -> str | None:
    """Pick a patient/sample column, rejecting per-cell barcodes.

    This mirrors scope_analysis.data.choose_patient_column so template generation
    cannot accidentally create one row per cell barcode (the LUAD 40k-patient
    failure mode).
    """
    if _choose_patient_column is not None:
        c, diag = _choose_patient_column(obs, configured=configured)
        if c is not None:
            return c
        rejected = diag.get("rejected", []) if isinstance(diag, dict) else []
        if rejected:
            print("[patient-status] rejected candidate patient columns that looked like cell IDs/barcodes:",
                  [(r.get("column"), r.get("n_unique"), round(float(r.get("barcode_like_frac", 0)), 3)) for r in rejected[:5]])
        return None

    # Minimal fallback if the package helper cannot be imported. Prefer columns
    # with repeated labels; reject one-label-per-cell high-cardinality vectors.
    candidates = []
    if configured and str(configured).strip():
        candidates.append(str(configured).strip())
    candidates.extend(PATIENT_CANDIDATES)
    for c in candidates:
        cc = c if c in obs.columns else next((x for x in obs.columns if str(x).lower() == c.lower()), None)
        if cc is None or not obs[cc].notna().any():
            continue
        s = obs[cc].dropna().astype(str).str.strip()
        counts = s.value_counts()
        if counts.empty:
            continue
        unique_frac = counts.size / max(len(s), 1)
        if unique_frac > 0.50 and float(counts.median()) <= 2:
            continue
        return cc
    return None


def read_obs(path: Path) -> pd.DataFrame:
    if ad is None:
        raise RuntimeError(f"Could not import anndata: {_ANNDATA_IMPORT_ERROR}")
    a = ad.read_h5ad(path, backed="r")
    try:
        obs = a.obs.copy()
    finally:
        try:
            a.file.close()
        except Exception:
            pass
    return obs


def drivers_from_config(cfg: dict, cancer: str, obs: pd.DataFrame | None = None) -> list[str]:
    drivers: list[str] = []
    bulk = cfg.get("bulk_cohorts", {}).get(cancer, {})
    mut_path = pth(bulk.get("mutations"))
    if mut_path and mut_path.exists():
        try:
            head = pd.read_csv(mut_path, nrows=2)
            sample_col = bulk.get("sample_col") or head.columns[0]
            drivers.extend([c for c in head.columns if c != sample_col and str(c).strip() not in METADATA_LIKE])
        except Exception:
            pass
    if obs is not None:
        for c in obs.columns:
            m = re.match(r"mutation_(?:prob|calib|adj|resid|refpct)_(.+)$", str(c))
            if m:
                drivers.append(m.group(1))
    out = []
    seen = set()
    for d in drivers:
        d = str(d).strip()
        if not d or d.lower() in METADATA_LIKE:
            continue
        k = d.upper()
        if k not in seen:
            seen.add(k)
            out.append(d)
    return out


def find_genotype_column(obs: pd.DataFrame, driver: str) -> str | None:
    dl = str(driver).lower()
    candidates = []
    for c in obs.columns:
        cs = str(c)
        cl = cs.lower()
        if dl in cl and TRUTH_RE.search(cs) and not PRED_RE.search(cs):
            candidates.append(cs)
    # Prefer columns that include both the driver and strong genotype words.
    for word in ("genotype", "mutation", "mutant", "status", "call", "label", "truth"):
        for c in candidates:
            if word in c.lower():
                return c
    return candidates[0] if candidates else None


def aggregate_per_cell_truth(obs: pd.DataFrame, patient_col: str, drivers: list[str]) -> tuple[pd.DataFrame, pd.DataFrame]:
    rows = []
    detail = []
    patients = obs[patient_col].astype(str)
    for d in drivers:
        col = find_genotype_column(obs, d)
        if not col:
            continue
        vals = obs[col].map(coerce_label)
        if vals.notna().sum() == 0:
            continue
        tmp = pd.DataFrame({"patient": patients, "label_cell": vals})
        for pat, sub in tmp.groupby("patient", observed=False):
            n_obs_truth = int(sub["label_cell"].notna().sum())
            n_mut_cells = int((sub["label_cell"] == 1).sum())
            n_wt_cells = int((sub["label_cell"] == 0).sum())
            if n_obs_truth == 0:
                lab = np.nan
            elif n_mut_cells > 0:
                lab = 1
            elif n_wt_cells > 0:
                lab = 0
            else:
                lab = np.nan
            rows.append({"patient": pat, "mutation": d, "label": lab})
            detail.append({
                "patient": pat, "mutation": d, "genotype_col": col,
                "n_truth_cells": n_obs_truth, "n_mut_cells": n_mut_cells,
                "n_wt_cells": n_wt_cells, "patient_label": lab,
            })
    status = pd.DataFrame(rows)
    if not status.empty:
        status = status.dropna(subset=["label"]).copy()
        status["label"] = status["label"].astype(int)
        status = status.drop_duplicates(["patient", "mutation"], keep="first")
    return status, pd.DataFrame(detail)


def standardize_status_table(path: Path, sc_patients: list[str], drivers: list[str]) -> tuple[pd.DataFrame, str]:
    df = pd.read_csv(path)
    patient_col = find_first(df.columns, ["patient", "Patient", "patient_id", "sample", "Sample", "sample_id", "SampleID", "donor", "case_id"])
    if patient_col is None:
        raise ValueError(f"No patient/sample column found in {path}")
    mut_col = find_first(df.columns, MUTATION_COL_CANDIDATES)
    lab_col = find_first(df.columns, LABEL_COL_CANDIDATES)

    if mut_col and lab_col:
        out = df[[patient_col, mut_col, lab_col]].rename(columns={patient_col: "patient", mut_col: "mutation", lab_col: "label"})
    else:
        # Wide format: patient + one column per driver/gene.
        driver_keys = {d.upper(): d for d in drivers}
        wide_cols = []
        for c in df.columns:
            if c == patient_col:
                continue
            k = str(c).strip().upper()
            if k in driver_keys:
                wide_cols.append(c)
            elif str(c).strip().lower() not in METADATA_LIKE:
                # Allow an external metadata table to include additional drivers too.
                vals = df[c].dropna().head(20).map(coerce_label)
                if vals.notna().any():
                    wide_cols.append(c)
        if not wide_cols:
            raise ValueError(f"No tidy mutation/status columns or wide driver columns found in {path}")
        out = df[[patient_col] + wide_cols].melt(id_vars=[patient_col], var_name="mutation", value_name="label")
        out = out.rename(columns={patient_col: "patient"})

    out["patient"] = out["patient"].astype(str).str.strip()
    out["mutation"] = out["mutation"].astype(str).str.strip()
    out["label"] = out["label"].map(coerce_label)
    out = out.dropna(subset=["patient", "mutation", "label"]).copy()
    out["label"] = out["label"].astype(int)
    out = harmonize_to_sc_patients(out, sc_patients)
    return out[["patient", "mutation", "label"]].drop_duplicates(), f"manual:{path}"


def harmonize_to_sc_patients(status: pd.DataFrame, sc_patients: list[str]) -> pd.DataFrame:
    out = status.copy()
    obs_map = {}
    for p in sc_patients:
        k = norm_key(p)
        if not k:
            continue
        if k not in obs_map:
            obs_map[k] = p
        else:
            obs_map[k] = None
    def map_one(x):
        mapped = obs_map.get(norm_key(x), None)
        return mapped if mapped is not None else str(x)
    out["patient_original"] = out["patient"].astype(str)
    out["patient"] = out["patient"].map(map_one)
    return out


def overlap_summary(status: pd.DataFrame | None, sc_patients: list[str]) -> tuple[int, int, float]:
    if status is None or status.empty:
        return 0, len(sc_patients), 0.0
    sc = set(map(str, sc_patients))
    st = set(status["patient"].astype(str))
    n = len(sc & st)
    frac = n / max(len(sc), 1)
    return n, len(sc), frac


def candidate_manual_files(manual_root: Path | None, cohort: str, cancer: str, configured_status: Path | None) -> list[Path]:
    roots = []
    if manual_root:
        roots.append(manual_root)
    if configured_status:
        roots.append(configured_status.parent)
    names = [
        f"{cohort}_patient_status_manual.csv",
        f"{cohort}_mutation_status_manual.csv",
        f"{cancer}_patient_status_manual.csv",
        f"{cancer}_mutation_status_manual.csv",
        "patient_status_manual.csv",
        "patient_mutation_status_manual.csv",
        "metadata_patient_status.csv",
    ]
    out = []
    for r in roots:
        for n in names:
            p = r / n
            if p.exists() and p not in out:
                out.append(p)
    return out


def write_templates(outdir: Path, cohort: str, patients: list[str], drivers: list[str]) -> tuple[Path, Path]:
    long_path = outdir / f"patient_status_TEMPLATE_{cohort}.csv"
    wide_path = outdir / f"patient_status_wide_TEMPLATE_{cohort}.csv"
    long_rows = [{"patient": p, "mutation": d, "label": "", "notes": "fill 1=mutant, 0=WT, blank=unknown"}
                 for p in patients for d in drivers]
    pd.DataFrame(long_rows).to_csv(long_path, index=False)
    wide = pd.DataFrame({"patient": patients})
    for d in drivers:
        wide[d] = ""
    wide.to_csv(wide_path, index=False)
    return long_path, wide_path


def patient_manifest(obs: pd.DataFrame, patient_col: str, celltype_col: str | None) -> pd.DataFrame:
    base = obs[[patient_col]].copy()
    base[patient_col] = base[patient_col].astype(str)
    rows = []
    for pat, sub in obs.groupby(base[patient_col], observed=False):
        row = {"patient": str(pat), "n_cells": int(len(sub))}
        if celltype_col and celltype_col in sub.columns:
            vc = sub[celltype_col].astype(str).value_counts().head(8)
            row["top_celltypes"] = "; ".join([f"{k}:{v}" for k, v in vc.items()])
        rows.append(row)
    return pd.DataFrame(rows).sort_values("n_cells", ascending=False)


def process_cohort(cohort: str, spec: dict, cfg: dict, args) -> CohortAudit:
    cancer = spec.get("cancer", cohort.split("_")[0])
    h5ad = pth(spec.get("scored_h5ad")) if args.prefer_scored else None
    if not h5ad or not h5ad.exists():
        h5ad = pth(spec.get("raw_h5ad"))
    if not h5ad or not h5ad.exists():
        return CohortAudit(cohort, cancer, "", "", 0, 0, 0, "missing_h5ad", False, "", "", 0, 0, 0, "", "H5AD path not found on this machine")

    obs = read_obs(h5ad)
    patient_col = infer_patient_col(obs, spec.get("sample_col"))
    if patient_col is None:
        return CohortAudit(cohort, cancer, str(h5ad), "", len(obs), 0, 0, "no_patient_col", False, "", "", 0, 0, 0, "", "Could not infer patient/sample column from obs")

    patients = sorted(pd.Series(obs[patient_col]).dropna().astype(str).unique().tolist())
    drivers = drivers_from_config(cfg, cancer, obs)
    outdir = pth(spec.get("patient_status_csv"))
    if outdir is None:
        outdir = Path(args.output_root) / cancer / "patient_status.csv"
    status_path = outdir
    outdir = status_path.parent
    outdir.mkdir(parents=True, exist_ok=True)

    # Always write manifest/templates; they are useful even when labels exist.
    manifest = patient_manifest(obs, patient_col, spec.get("celltype_col"))
    manifest.to_csv(outdir / f"patient_manifest_{cohort}.csv", index=False)
    template_path, wide_template = write_templates(outdir, cohort, patients, drivers)

    warnings = []
    chosen = pd.DataFrame()
    source = "none"

    # 1) Manual/publication exact-scRNA metadata, if user put it next to the output or in --manual-root.
    for mf in candidate_manual_files(pth(args.manual_root), cohort, cancer, status_path):
        try:
            cand, src = standardize_status_table(mf, patients, drivers)
            n_ov, n_sc, frac = overlap_summary(cand, patients)
            if n_ov > 0 and (frac >= args.min_overlap_frac or n_ov >= args.min_overlap_patients):
                chosen, source = cand, src
                break
            warnings.append(f"manual table {mf} rejected: overlap {n_ov}/{n_sc} scRNA patients")
        except Exception as e:
            warnings.append(f"manual table {mf} could not be parsed: {e}")

    # 2) Per-cell truth already inside this same H5AD.
    detail_path = ""
    if chosen.empty:
        truth, detail = aggregate_per_cell_truth(obs, patient_col, drivers)
        if not truth.empty:
            n_ov, n_sc, frac = overlap_summary(truth, patients)
            chosen, source = truth, "aggregated_per_cell_truth_from_h5ad_obs"
            if not detail.empty:
                dp = outdir / f"patient_status_percell_truth_detail_{cohort}.csv"
                detail.to_csv(dp, index=False)
                detail_path = str(dp)

    # 3) Existing patient_status.csv, only if explicitly accepted and it overlaps scRNA IDs.
    existing_overlap = "missing"
    if chosen.empty and args.accept_existing and status_path.exists():
        try:
            ex, src = standardize_status_table(status_path, patients, drivers)
            n_ov, n_sc, frac = overlap_summary(ex, patients)
            existing_overlap = f"{n_ov}/{n_sc} ({frac:.3f})"
            if n_ov > 0 and (frac >= args.min_overlap_frac or n_ov >= args.min_overlap_patients):
                chosen, source = ex, f"accepted_existing:{status_path}"
            else:
                warnings.append(f"existing patient_status rejected: overlap {existing_overlap}")
        except Exception as e:
            existing_overlap = f"parse_failed:{e}"
            warnings.append(f"existing patient_status could not be parsed: {e}")
    elif status_path.exists():
        try:
            ex, _ = standardize_status_table(status_path, patients, drivers)
            n_ov, n_sc, frac = overlap_summary(ex, patients)
            existing_overlap = f"{n_ov}/{n_sc} ({frac:.3f}); not used unless --accept-existing"
            if n_ov == 0:
                warnings.append("existing patient_status has 0 overlap with scRNA patients; likely bulk/unrelated cohort")
        except Exception as e:
            existing_overlap = f"parse_failed:{e}"

    wrote = False
    n_rows = n_mut = n_wt = 0
    if not chosen.empty:
        chosen = chosen.dropna(subset=["label"]).copy()
        chosen["label"] = chosen["label"].astype(int)
        chosen = chosen[chosen["patient"].astype(str).isin(set(map(str, patients)))].copy()
        if not chosen.empty:
            if status_path.exists() and args.overwrite:
                backup = status_path.with_suffix(status_path.suffix + ".bak")
                try:
                    shutil.copy2(status_path, backup)
                except Exception:
                    pass
            if args.overwrite or not status_path.exists():
                chosen[["patient", "mutation", "label"]].to_csv(status_path, index=False)
                wrote = True
            else:
                alt = outdir / f"patient_status_CANDIDATE_{cohort}.csv"
                chosen[["patient", "mutation", "label"]].to_csv(alt, index=False)
                status_path = alt
                wrote = True
                warnings.append("wrote candidate instead of overwriting existing patient_status.csv; rerun with --overwrite to replace")
            n_rows = len(chosen)
            n_mut = int((chosen["label"] == 1).sum())
            n_wt = int((chosen["label"] == 0).sum())
        else:
            warnings.append("candidate labels had no rows after restricting to scRNA patients")
    else:
        warnings.append("no usable exact-scRNA mutation metadata found; fill the generated template manually")

    return CohortAudit(
        cohort=cohort, cancer=cancer, h5ad=str(h5ad), patient_col=patient_col,
        n_cells=len(obs), n_patients=len(patients), n_drivers=len(drivers),
        status_source=source, wrote_patient_status=wrote, status_path=str(status_path),
        template_path=str(template_path), n_status_rows=n_rows, n_mut_rows=n_mut, n_wt_rows=n_wt,
        existing_overlap=existing_overlap, warning=" | ".join(warnings),
    )


def main():
    ap = argparse.ArgumentParser(description=__doc__, formatter_class=argparse.RawTextHelpFormatter)
    ap.add_argument("--config", default="drivers/config_canonical_v3.json")
    ap.add_argument("--manual-root", default="", help="Folder containing publication/manual patient mutation metadata CSVs.")
    ap.add_argument("--output-root", default="sc_patient_status_outputs", help="Fallback output root if a cohort has no patient_status_csv in config.")
    ap.add_argument("--overwrite", action="store_true", help="Replace configured patient_status.csv when a valid exact-scRNA status is found.")
    ap.add_argument("--accept-existing", action="store_true", help="Allow an existing patient_status.csv to be reused if it overlaps scRNA patient IDs.")
    ap.add_argument("--prefer-raw", dest="prefer_scored", action="store_false", help="Use raw_h5ad before scored_h5ad. Default prefers scored_h5ad because truth columns often live there.")
    ap.set_defaults(prefer_scored=True)
    ap.add_argument("--min-overlap-frac", type=float, default=0.25, help="Minimum fraction of scRNA patients covered by manual/existing labels, unless min-overlap-patients is met.")
    ap.add_argument("--min-overlap-patients", type=int, default=2, help="Minimum absolute overlapping patients for manual/existing labels.")
    args = ap.parse_args()

    cfg_path = Path(args.config)
    cfg = json.loads(cfg_path.read_text())
    cohorts = cfg.get("singlecell_cohorts", {})
    if not isinstance(cohorts, dict):
        raise ValueError("Expected config['singlecell_cohorts'] to be a dict")

    audits = []
    for cohort, spec in cohorts.items():
        print(f"\n=== {cohort} ===")
        audit = process_cohort(cohort, spec, cfg, args)
        audits.append(audit.__dict__)
        print(f"patients={audit.n_patients:,} drivers={audit.n_drivers:,} source={audit.status_source}")
        if audit.wrote_patient_status:
            print(f"[write] {audit.status_path} rows={audit.n_status_rows:,} mut={audit.n_mut_rows:,} wt={audit.n_wt_rows:,}")
        else:
            print(f"[template] {audit.template_path}")
        if audit.warning:
            print(f"[warn] {audit.warning}")

    out = pd.DataFrame(audits)
    audit_path = cfg_path.parent / "patient_status_generation_audit.csv"
    out.to_csv(audit_path, index=False)
    print(f"\n[audit] {audit_path}")
    print("\nNext: fill any patient_status_TEMPLATE_*.csv files from the original scRNA publication metadata, then rerun with --manual-root/--overwrite.")


if __name__ == "__main__":
    main()
```
