## Supplemental_Code for "scOPE identifies which driver-associated expression programs transfer from bulk tumors to single cells": patch_bulk_loader_ids.py

```
"""
Patch scope_analysis/scope_analysis/data.py so bulk expression + mutation-label
loading works with your real files.

Fixes:
  1. reads .tsv/.tsv.gz/.txt/.txt.gz with tab separators instead of CSV defaults
  2. strips whitespace from sample/gene IDs
  3. tries conservative sample-ID normalization before declaring no overlap

Run from the suite root:
  cd C:\\Users\\ashfo\\OneDrive\\Desktop\\Pathways+Omics\\scOPE\\scope_analysis
  python patch_bulk_loader_ids.py
"""
from pathlib import Path

DATA_PATH = Path("scope_analysis") / "data.py"

OLD_LOAD_BULK = '''def load_bulk_cohort(
    expression_path: str | Path,
    mutation_path: str | Path,
    sample_col: str = "sample_id",
    genes_as_columns: bool = True,
) -> tuple["ad.AnnData", pd.DataFrame]:
    """Load a bulk cohort as (AnnData, binary-label DataFrame) ready for
    ``BulkPipeline.fit``.

    expression_path: .h5ad (samples x genes) OR a wide CSV (samples x genes).
    mutation_path:   CSV with ``sample_col`` + one 0/1 column per driver.

    The two are inner-joined on sample id; mismatched samples are dropped with a
    printed count so you always know how many tumours actually entered training.
    """
    expression_path = Path(expression_path)
    if expression_path.suffix == ".h5ad":
        adata = ad.read_h5ad(expression_path)
    else:
        df = pd.read_csv(expression_path, index_col=0)
        if not genes_as_columns:
            df = df.T
        adata = ad.AnnData(X=df.values.astype(np.float32))
        adata.obs_names = df.index.astype(str)
        adata.var_names = df.columns.astype(str)

    labels = _read_label_table(mutation_path, sample_col)
    shared = adata.obs_names.intersection(labels.index)
    if len(shared) == 0:
        raise ValueError(
            f"No shared sample ids between {expression_path.name} and "
            f"{Path(mutation_path).name}. Check sample_col and id formatting."
        )
    print(f"[data] {expression_path.name}: {adata.n_obs} samples, "
          f"{len(shared)} with mutation labels, {labels.shape[1]} drivers.")
    adata = adata[shared].copy()
    labels = labels.loc[shared]
    return adata, labels
'''

NEW_LOAD_BULK = '''def load_bulk_cohort(
    expression_path: str | Path,
    mutation_path: str | Path,
    sample_col: str = "sample_id",
    genes_as_columns: bool = True,
) -> tuple["ad.AnnData", pd.DataFrame]:
    """Load a bulk cohort as (AnnData, binary-label DataFrame) ready for
    ``BulkPipeline.fit``.

    Robust version for the local manuscript inputs:
    - .tsv/.tsv.gz/.txt/.txt.gz are read with tab separators
    - sample/gene ids are stripped of whitespace
    - exact join is tried first; conservative normalized join is tried second
    """
    expression_path = Path(expression_path)
    if expression_path.suffix == ".h5ad":
        adata = ad.read_h5ad(expression_path)
        adata.obs_names = adata.obs_names.astype(str).str.strip()
        adata.var_names = adata.var_names.astype(str).str.strip()
    else:
        df = _read_wide_expression_table(expression_path)
        if not genes_as_columns:
            df = df.T
        adata = ad.AnnData(X=df.values.astype(np.float32))
        adata.obs_names = df.index.astype(str)
        adata.var_names = df.columns.astype(str)

    labels = _read_label_table(mutation_path, sample_col)
    adata, labels = _align_bulk_and_labels(
        adata=adata,
        labels=labels,
        expression_name=expression_path.name,
        mutation_name=Path(mutation_path).name,
    )
    print(f"[data] {expression_path.name}: {adata.n_obs} matched samples, "
          f"{adata.n_vars} genes, {labels.shape[1]} drivers.")
    return adata, labels
'''

OLD_READ_LABEL = '''def _read_label_table(path: str | Path, sample_col: str) -> pd.DataFrame:
    """Read a mutation-label CSV and coerce to strict 0/1 ints.

    Accepts 0/1, mutant/WT, yes/no, true/false, pos/neg in any casing.
    """
    df = pd.read_csv(path)
    if sample_col in df.columns:
        df = df.set_index(sample_col)
    elif df.columns[0].lower() in ("sample", "sample_id", "id", "barcode"):
        df = df.set_index(df.columns[0])
    return df.apply(_coerce_binary)
'''

NEW_READ_LABEL = '''def _read_label_table(path: str | Path, sample_col: str) -> pd.DataFrame:
    """Read a mutation-label CSV/TSV and coerce to strict 0/1 ints.

    Accepts 0/1, mutant/WT, yes/no, true/false, pos/neg in any casing.
    """
    path = Path(path)
    sep = _sep_for_table(path)
    df = pd.read_csv(path, sep=sep)
    idx_col = _choose_label_index_column(df, sample_col)
    df = df.set_index(idx_col)
    df.index = df.index.astype(str).str.strip()

    # Keep only putative driver columns; this prevents metadata columns from
    # becoming fake all-zero mutation labels if present.
    metadata_like = {
        "original_sample_id", "sample_id", "sample", "barcode", "patient", "patient_id",
        "case_id", "cancer", "cohort", "dataset"
    }
    keep_cols = [c for c in df.columns if str(c).strip() not in metadata_like]
    df = df[keep_cols]
    return df.apply(_coerce_binary)
'''

HELPERS = r'''

def _sep_for_table(path: str | Path) -> str:
    """Infer separator from filename."""
    name = Path(path).name.lower()
    return "\t" if name.endswith((".tsv", ".tsv.gz", ".txt", ".txt.gz")) else ","


def _read_wide_expression_table(path: str | Path) -> pd.DataFrame:
    """Read wide expression matrix with auto TSV/CSV separator detection."""
    df = pd.read_csv(path, sep=_sep_for_table(path), index_col=0)
    df.index = df.index.astype(str).str.strip()
    df.columns = df.columns.astype(str).str.strip()
    return df


def _normalize_sample_id(x) -> str:
    """Conservative sample-id normalizer for TCGA/BeatAML-style joins."""
    s = str(x).strip()
    s = re.sub(r"^X", "", s)  # R sometimes prefixes sample IDs
    s = s.replace(".", "-").replace("_", "-")
    s = re.sub(r"\s+", "", s)
    s = s.upper()

    # TCGA barcode sample-level matching: TCGA-XX-YYYY-01A-... -> TCGA-XX-YYYY-01
    if s.startswith("TCGA-"):
        parts = s.split("-")
        if len(parts) >= 4:
            sample_type = parts[3][:2]
            s = "-".join(parts[:3] + [sample_type])
    return s


def _make_unique_index(values) -> pd.Index:
    """Make duplicate normalized IDs unique without silently collapsing rows."""
    seen = {}
    out = []
    for v in values:
        v = str(v)
        n = seen.get(v, 0)
        seen[v] = n + 1
        out.append(v if n == 0 else f"{v}__DUP{n}")
    return pd.Index(out)


def _choose_label_index_column(df: pd.DataFrame, requested: str) -> str:
    """Pick a plausible sample-id column for mutation labels."""
    if requested in df.columns:
        return requested
    candidates = [
        "sample_id", "sample", "Sample", "SAMPLE", "id", "ID",
        "barcode", "Barcode", "BARCODE", "patient_id", "patient", "case_id",
        "tumor_sample_barcode", "Tumor_Sample_Barcode", "TumorSampleBarcode",
    ]
    for c in candidates:
        if c in df.columns:
            print(f"[data] sample_col {requested!r} absent; using detected column {c!r}.")
            return c
    print(f"[data] sample_col {requested!r} absent; using first column {df.columns[0]!r}.")
    return df.columns[0]


def _align_bulk_and_labels(
    adata: "ad.AnnData",
    labels: pd.DataFrame,
    expression_name: str,
    mutation_name: str,
) -> tuple["ad.AnnData", pd.DataFrame]:
    """Exact sample-id join first; then normalized join with diagnostics."""
    shared = adata.obs_names.intersection(labels.index)
    if len(shared) > 0:
        return adata[shared].copy(), labels.loc[shared].copy()

    original_expr_ids = pd.Index(adata.obs_names.astype(str))
    original_label_ids = pd.Index(labels.index.astype(str))
    expr_norm = pd.Index([_normalize_sample_id(x) for x in original_expr_ids])
    lab_norm = pd.Index([_normalize_sample_id(x) for x in original_label_ids])

    adata = adata.copy()
    labels = labels.copy()
    adata.obs["original_sample_id"] = original_expr_ids.to_numpy()
    labels["original_sample_id"] = original_label_ids.to_numpy()
    adata.obs_names = _make_unique_index(expr_norm)
    labels.index = _make_unique_index(lab_norm)

    shared = adata.obs_names.intersection(labels.index)
    if len(shared) == 0:
        print("[data] DEBUG expression sample examples:", list(original_expr_ids[:10]))
        print("[data] DEBUG label sample examples:", list(original_label_ids[:10]))
        print("[data] DEBUG normalized expression examples:", list(expr_norm[:10]))
        print("[data] DEBUG normalized label examples:", list(lab_norm[:10]))
        raise ValueError(
            f"No shared sample ids between {expression_name} and {mutation_name}, "
            "even after TSV/CSV detection and conservative ID normalization. "
            "Check sample_col and whether expression rows are samples rather than genes."
        )

    print(
        f"[data] normalized sample-id join rescued {len(shared)} shared samples "
        f"between {expression_name} and {mutation_name}."
    )
    labels = labels.loc[shared].drop(columns=["original_sample_id"], errors="ignore")
    return adata[shared].copy(), labels.copy()
'''


def main():
    if not DATA_PATH.exists():
        raise SystemExit(f"Could not find {DATA_PATH}. Run this from the scope_analysis suite root.")

    text = DATA_PATH.read_text(encoding="utf-8")
    backup = DATA_PATH.with_suffix(".py.BACKUP_before_bulk_id_patch")
    if not backup.exists():
        backup.write_text(text, encoding="utf-8")
        print(f"Backup written: {backup}")

    if "def _sep_for_table" not in text:
        marker = "\n\ndef _coerce_binary"
        if marker not in text:
            raise SystemExit("Could not find insertion point before _coerce_binary().")
        text = text.replace(marker, HELPERS + marker)

    if OLD_LOAD_BULK in text:
        text = text.replace(OLD_LOAD_BULK, NEW_LOAD_BULK)
    elif "def _align_bulk_and_labels" in text and "matched samples" in text:
        print("load_bulk_cohort already appears patched; leaving it as-is.")
    else:
        raise SystemExit("Could not find the original load_bulk_cohort() block. File may already be edited.")

    if OLD_READ_LABEL in text:
        text = text.replace(OLD_READ_LABEL, NEW_READ_LABEL)
    elif "def _choose_label_index_column" in text and "Read a mutation-label CSV/TSV" in text:
        print("_read_label_table already appears patched; leaving it as-is.")
    else:
        raise SystemExit("Could not find the original _read_label_table() block. File may already be edited.")

    DATA_PATH.write_text(text, encoding="utf-8")
    print(f"Patched: {DATA_PATH}")
    print("Next: rerun the bulk stage.")


if __name__ == "__main__":
    main()
```
