## Supplemental_Code for "scOPE identifies which driver-associated expression programs transfer from bulk tumors to single cells": patch_longitudinal_singletons.py

```
from pathlib import Path

run_all = Path("drivers/run_all.py")
longitudinal = Path("scope_analysis/longitudinal.py")

# ---------------------------------------------------------------------
# 1. Fix exact include_samples + include_regex to be OR, not AND
# ---------------------------------------------------------------------
txt = run_all.read_text(encoding="utf-8")

old = '''    analysis_mask = pd.Series(True, index=adata.obs.index)
    if include_samples:
        analysis_mask &= samp.isin(include_samples)
    if include_regex:
        try:
            analysis_mask &= samp.str.contains(include_regex, regex=True, na=False)
        except Exception as e:
            print(f"[filter] {label}: invalid longitudinal_include_samples_regex={include_regex!r}: {e}; ignoring regex.")
'''

new = '''    # Include filters are additive/OR:
    #   longitudinal_include_samples keeps exact named samples
    #   longitudinal_include_samples_regex keeps regex-matched samples
    # This lets you keep the main longitudinal whitelist AND add one-off samples
    # like AML1012-D0 or AML210A-D0.
    if include_samples or include_regex:
        analysis_mask = pd.Series(False, index=adata.obs.index)
        if include_samples:
            analysis_mask |= samp.isin(include_samples)
        if include_regex:
            try:
                analysis_mask |= samp.str.contains(include_regex, regex=True, na=False)
            except Exception as e:
                print(f"[filter] {label}: invalid longitudinal_include_samples_regex={include_regex!r}: {e}; ignoring regex.")
    else:
        analysis_mask = pd.Series(True, index=adata.obs.index)
'''

if old not in txt:
    print("[warn] Could not find exact include filter block in drivers/run_all.py; maybe already patched?")
else:
    txt = txt.replace(old, new, 1)
    run_all.write_text(txt, encoding="utf-8")
    print("[patched] drivers/run_all.py include_samples/regex OR logic")


# ---------------------------------------------------------------------
# 2. Add include_single_timepoint_patients option to longitudinal_trajectory
# ---------------------------------------------------------------------
txt = longitudinal.read_text(encoding="utf-8")

old_sig = '''    reference_mask_col: str = "is_reference",
    malignant_only_col: str | None = None,
    score_mode: str = "configured",
) -> pd.DataFrame:
'''

new_sig = '''    reference_mask_col: str = "is_reference",
    malignant_only_col: str | None = None,
    score_mode: str = "configured",
    include_single_timepoint_patients: bool = False,
) -> pd.DataFrame:
'''

if old_sig not in txt:
    print("[warn] Could not patch longitudinal_trajectory signature; maybe already patched?")
else:
    txt = txt.replace(old_sig, new_sig, 1)

old_skip = '''            if len(tps) < 2:
                continue  # need >=2 timepoints to call it a trajectory
'''

new_skip = '''            if len(tps) < 2 and not include_single_timepoint_patients:
                continue  # default: need >=2 timepoints to call it a trajectory
'''

if old_skip not in txt:
    print("[warn] Could not patch singleton skip; maybe already patched?")
else:
    txt = txt.replace(old_skip, new_skip, 1)

longitudinal.write_text(txt, encoding="utf-8")
print("[patched] scope_analysis/longitudinal.py singleton option")


# ---------------------------------------------------------------------
# 3. Pass the singleton option from the cohort JSON spec in run_all.py
# ---------------------------------------------------------------------
txt = run_all.read_text(encoding="utf-8")

marker = '''            traj = sa.longitudinal.longitudinal_trajectory(adata_long, cfg, drivers=drivers,
                                                           malignant_only_col=malig_col,
                                                           reference_mask_col=long_primary_ref_col,
                                                           score_mode=primary_sc_score_mode)
'''

replacement = '''            include_singletons = bool(spec.get("longitudinal_include_single_timepoint_patients",
                                                spec.get("longitudinal_include_singletons", False)))
            if include_singletons:
                print(f"[longitudinal] {key}: including one-timepoint samples as singleton points; "
                      "trend rho/p will be NaN for singleton patient-driver pairs.")
            traj = sa.longitudinal.longitudinal_trajectory(adata_long, cfg, drivers=drivers,
                                                           malignant_only_col=malig_col,
                                                           reference_mask_col=long_primary_ref_col,
                                                           score_mode=primary_sc_score_mode,
                                                           include_single_timepoint_patients=include_singletons)
'''

if marker not in txt:
    print("[warn] Could not patch primary longitudinal_trajectory call; maybe already patched?")
else:
    txt = txt.replace(marker, replacement, 1)

old_guard = '''                    gtraj = sa.longitudinal.longitudinal_trajectory(
                        adata_long, cfg, drivers=drivers, malignant_only_col=malig_col,
                        reference_mask_col=gref, score_mode=guarded_mode)
'''

new_guard = '''                    gtraj = sa.longitudinal.longitudinal_trajectory(
                        adata_long, cfg, drivers=drivers, malignant_only_col=malig_col,
                        reference_mask_col=gref, score_mode=guarded_mode,
                        include_single_timepoint_patients=include_singletons)
'''

if old_guard not in txt:
    print("[warn] Could not patch guarded longitudinal_trajectory call; maybe already patched?")
else:
    txt = txt.replace(old_guard, new_guard, 1)

run_all.write_text(txt, encoding="utf-8")
print("[patched] drivers/run_all.py singleton option passed into trajectory calls")

print("Done.")
```
