## Supplemental_Code for "scOPE identifies which driver-associated expression programs transfer from bulk tumors to single cells": patch_muttranscript_genotypes.py

```
from pathlib import Path

singlecell = Path("scope_analysis/singlecell.py")
run_all = Path("drivers/run_all.py")

sc_txt = singlecell.read_text(encoding="utf-8")

if "def add_muttranscript_genotype_columns" not in sc_txt:
    insert = r'''

# ---------------------------------------------------------------------------
# van Galen expressed-mutation transcript truth parser
# ---------------------------------------------------------------------------
def _find_obs_col_case_insensitive(obs: pd.DataFrame, names: list[str]) -> str | None:
    """Return the first obs column matching one of `names`, case-insensitively."""
    exact = set(obs.columns)
    for n in names:
        if n in exact:
            return n
    low = {str(c).lower(): c for c in obs.columns}
    for n in names:
        if str(n).lower() in low:
            return low[str(n).lower()]
    return None


def _driver_in_transcript_series(series: pd.Series, driver: str) -> pd.Series:
    """True where a MutTranscript/WtTranscript-style string mentions `driver`."""
    pat = r"(?<![A-Za-z0-9])" + re.escape(str(driver).strip()) + r"(?![A-Za-z0-9])"
    return series.astype(str).str.contains(pat, case=False, regex=True, na=False)


def add_muttranscript_genotype_columns(
    adata,
    drivers: list[str] | None = None,
    mut_col: str | None = None,
    wt_col: str | None = None,
    overwrite: bool = True,
) -> tuple[object, pd.DataFrame]:
    """Parse van Galen-style MutTranscript/WtTranscript obs strings into true
    per-cell genotype columns named <DRIVER>_genotype.

    Semantics:
      * driver mentioned in MutTranscript -> 1 / Mut
      * driver mentioned in WtTranscript  -> 0 / WT
      * neither mentioned                 -> NaN / unknown
      * both mentioned                    -> 1 wins, conservatively
    """
    obs = adata.obs

    mut_col = mut_col if mut_col in obs.columns else _find_obs_col_case_insensitive(
        obs,
        [
            "MutTranscript", "MutTranscripts",
            "mut_transcript", "mut_transcripts",
            "MutatedTranscript", "MutatedTranscripts",
        ],
    )

    wt_col = wt_col if wt_col in obs.columns else _find_obs_col_case_insensitive(
        obs,
        [
            "WtTranscript", "WTTranscript",
            "WtTranscripts", "WTTranscripts",
            "wt_transcript", "wt_transcripts",
            "WildtypeTranscript", "WildTypeTranscript",
        ],
    )

    if mut_col is None and wt_col is None:
        return adata, pd.DataFrame()

    drivers = drivers or scored_drivers(adata)
    mut_s = obs[mut_col].astype(str) if mut_col is not None else pd.Series("", index=obs.index)
    wt_s = obs[wt_col].astype(str) if wt_col is not None else pd.Series("", index=obs.index)

    rows = []

    for d in drivers:
        d = str(d).strip()
        if not d:
            continue

        mut_mask = (
            _driver_in_transcript_series(mut_s, d)
            if mut_col is not None
            else pd.Series(False, index=obs.index)
        )
        wt_mask = (
            _driver_in_transcript_series(wt_s, d)
            if wt_col is not None
            else pd.Series(False, index=obs.index)
        )

        known = mut_mask | wt_mask
        if int(known.sum()) == 0:
            continue

        out = pd.Series(np.nan, index=obs.index, dtype="float")
        out.loc[wt_mask] = 0.0
        out.loc[mut_mask] = 1.0

        out_col = f"{d}_genotype"

        if overwrite or out_col not in obs.columns:
            obs[out_col] = out.values

        rows.append(
            {
                "driver": d,
                "genotype_col": out_col,
                "muttranscript_col": mut_col,
                "wttranscript_col": wt_col,
                "n_mut_cells": int(mut_mask.sum()),
                "n_wt_cells": int(wt_mask.sum()),
                "n_known_cells": int(known.sum()),
                "n_unknown_cells": int(len(obs) - int(known.sum())),
            }
        )

    summary = pd.DataFrame(rows)

    if not summary.empty:
        print(
            f"[singlecell] parsed MutTranscript/WtTranscript into {len(summary)} "
            f"per-cell genotype column(s): "
            f"{summary['driver'].head(12).tolist()}"
            + (" ..." if len(summary) > 12 else "")
        )
    elif mut_col is not None or wt_col is not None:
        print(
            "[singlecell] MutTranscript/WtTranscript columns found, "
            "but no scored driver names were detected inside them."
        )

    return adata, summary
'''

    marker = '\ndef resolve_score_col(adata, cfg: AnalysisConfig, driver: str, score_mode: str = "configured") -> str:\n'
    if marker not in sc_txt:
        raise SystemExit("Could not find insertion marker in scope_analysis/singlecell.py")

    sc_txt = sc_txt.replace(marker, insert + marker)
    singlecell.write_text(sc_txt, encoding="utf-8")
    print("[patched] scope_analysis/singlecell.py")
else:
    print("[skip] scope_analysis/singlecell.py already patched")


ra_txt = run_all.read_text(encoding="utf-8")

if "MutTranscript_genotype_parse_" not in ra_txt:
    marker = "        drivers = sa.data.scored_drivers(adata)\n"

    repl = marker + '''        # van Galen stores per-cell expressed-mutation truth in generic
        # MutTranscript/WtTranscript string columns. Parse those into per-driver
        # <GENE>_genotype columns before Fig3 validation and Fig4 triptychs,
        # otherwise AML falls back to patient-level mutation status.
        try:
            adata, mt_parse = sa.singlecell.add_muttranscript_genotype_columns(
                adata, drivers=drivers, overwrite=True)
            if mt_parse is not None and not mt_parse.empty:
                M.save_table(f"MutTranscript_genotype_parse_{key}", mt_parse)
        except Exception as e:
            print(f"[singlecell] {key}: MutTranscript/WtTranscript genotype parse skipped ({e})")
'''

    if marker not in ra_txt:
        raise SystemExit("Could not find drivers marker in drivers/run_all.py")

    ra_txt = ra_txt.replace(marker, repl, 1)
    run_all.write_text(ra_txt, encoding="utf-8")
    print("[patched] drivers/run_all.py")
else:
    print("[skip] drivers/run_all.py already patched")

print("Done.")
```
