## Supplemental_Code for "scOPE identifies which driver-associated expression programs transfer from bulk tumors to single cells": patch_scopebio_svd_shap_v2.py

```
#!/usr/bin/env python
r"""
patch_scopebio_svd_shap_v2.py

Patch installed/local scope-bio SVDEvaluator.plot_shap so SHAP works with:
  - scope.classification.base._SklearnWrapper
  - scope.classification.models.LogisticMutationClassifier
  - sklearn Pipeline

Run inside your scope_analysis conda/micromamba environment:
  python patch_scopebio_svd_shap_v2.py
r"""

from __future__ import annotations

import inspect
from pathlib import Path

import scope.evaluation.svd_evaluation as svd_eval


NEW_PLOT_SHAP = r'''    def plot_shap(self, output_dir: Path) -> None:
        """SHAP dot + bar summary (requires ``shap``).

        Robust to scOPE classifier wrappers:
        - _SklearnWrapper: apply its StandardScaler to Z, then unwrap ._clf
        - LogisticMutationClassifier: unwrap its underlying sklearn ._model
        - sklearn Pipeline: transform through preprocessing, then unwrap final estimator
        """
        if not _HAS_SHAP:
            log.warning("shap not installed — skipping SHAP plots.")
            return

        clf_final = self._clf
        Z = self.Z_bulk

        # scOPE PerMutationClassifierSet wraps classifiers in _SklearnWrapper
        # when scale_features=True. SHAP cannot parse that wrapper directly.
        # The background matrix must be transformed with the fitted scaler.
        if hasattr(clf_final, "_scaler") and hasattr(clf_final, "_clf"):
            Z = clf_final._scaler.transform(Z)
            clf_final = clf_final._clf

        # sklearn Pipeline support.
        if isinstance(clf_final, Pipeline):
            if len(clf_final.steps) > 1:
                Z = clf_final[:-1].transform(Z)
            clf_final = clf_final[-1]

        # scOPE LogisticMutationClassifier wraps the real sklearn model in ._model.
        # SHAP LinearExplainer expects the actual sklearn estimator.
        if hasattr(clf_final, "_model"):
            clf_final = clf_final._model

        feature_names = [f"SVD{i + 1}" for i in range(self.k)]

        try:
            explainer = _shap_lib.LinearExplainer(clf_final, Z)
            shap_vals = explainer.shap_values(Z)
        except Exception as e:
            log.warning(
                "SHAP failed for %s with %s: %s — skipping SHAP plots.",
                self.mutation, type(e).__name__, e
            )
            return

        if isinstance(shap_vals, list):
            shap_vals = np.array(shap_vals).mean(axis=0)

        _shap_lib.summary_plot(shap_vals, Z, feature_names=feature_names, show=False)
        plt.title(f"SHAP dot summary  ({self.mutation})")
        self._savefig(plt.gcf(), "shap_summary_dot.png", output_dir)

        _shap_lib.summary_plot(
            shap_vals, Z, feature_names=feature_names, plot_type="bar", show=False
        )
        plt.title(f"SHAP bar summary  ({self.mutation})")
        self._savefig(plt.gcf(), "shap_summary_bar.png", output_dir)
'''


def replace_method(text: str, method_name: str, replacement: str) -> str:
    lines = text.splitlines(keepends=True)
    start = None
    for i, line in enumerate(lines):
        if line.startswith(f"    def {method_name}("):
            start = i
            break

    if start is None:
        raise RuntimeError(f"Could not find method {method_name!r}")

    end = None
    for j in range(start + 1, len(lines)):
        if lines[j].startswith("    def "):
            end = j
            break

    if end is None:
        raise RuntimeError(f"Could not find end of method {method_name!r}")

    return "".join(lines[:start]) + replacement + "\n" + "".join(lines[end:])


def main() -> None:
    src = Path(inspect.getfile(svd_eval)).resolve()
    print(f"Patching: {src}")

    text = src.read_text(encoding="utf-8")

    if "Robust to scOPE classifier wrappers" in text:
        print("Already patched.")
        return

    backup = src.with_suffix(src.suffix + ".BACKUP_before_shap_unwrap_patch")
    if not backup.exists():
        backup.write_text(text, encoding="utf-8")
        print(f"Backup written: {backup}")
    else:
        print(f"Backup already exists: {backup}")

    patched = replace_method(text, "plot_shap", NEW_PLOT_SHAP)
    src.write_text(patched, encoding="utf-8")
    print("Done. Restart Python/Jupyter if needed, then rerun interpretation.")


if __name__ == "__main__":
    main()
```
