## Supplemental_Code for "scOPE identifies which driver-associated expression programs transfer from bulk tumors to single cells": patch_shiftresult_nan_return.py

```
#!/usr/bin/env python
"""
patch_shiftresult_nan_return.py

Fixes:
    TypeError: ShiftResult.__init__() takes 9 positional arguments but 10 were given

Cause:
    scope_analysis/stats.py has a degenerate/empty-group return path in
    distribution_shift() that returns 7 NaNs + n_tumour + n_reference.
    But the local ShiftResult class currently has 8 fields total, so that
    empty-path return should provide 6 NaNs + the two counts.

This patch replaces the brittle hard-coded return with a field-count-aware return.
"""

from pathlib import Path

stats_path = Path("scope_analysis") / "stats.py"
if not stats_path.exists():
    raise FileNotFoundError(
        f"Could not find {stats_path}. Run this from the scope_analysis project folder:\n"
        r"  C:\Users\ashfo\OneDrive\Desktop\Pathways+Omics\scOPE\scope_analysis"
    )

text = stats_path.read_text(encoding="utf-8")

old = "return ShiftResult(*([np.nan] * 7), len(s), len(r))"
new = (
    "# Degenerate/empty-group path. Keep this robust to ShiftResult field-count changes.\n"
    "        _n_fields = (\n"
    "            len(getattr(ShiftResult, \"__dataclass_fields__\", {}))\n"
    "            or len(getattr(ShiftResult, \"_fields\", []))\n"
    "            or 8\n"
    "        )\n"
    "        return ShiftResult(*([np.nan] * max(0, _n_fields - 2)), len(s), len(r))"
)

if old not in text:
    print("[warn] Exact old return not found. Looking for nearby ShiftResult NaN returns...")
    import re
    hits = re.findall(r"return ShiftResult\(\*\(\[np\.nan\] \* \d+\), len\(s\), len\(r\)\)", text)
    print("[hits]", hits)
    if not hits:
        raise RuntimeError("No matching ShiftResult NaN return found; nothing patched.")
    for h in hits:
        text = text.replace(h, new)
else:
    text = text.replace(old, new)

backup = stats_path.with_suffix(".py.BACKUP_before_shiftresult_nan_patch")
if not backup.exists():
    backup.write_text(stats_path.read_text(encoding="utf-8"), encoding="utf-8")
    print(f"[backup] {backup}")
else:
    print(f"[backup exists] {backup}")

stats_path.write_text(text, encoding="utf-8")
print(f"[patched] {stats_path}")

# Basic syntax check.
compile(stats_path.read_text(encoding="utf-8"), str(stats_path), "exec")
print("[ok] stats.py compiles")
print("\nNow rerun:")
print("python drivers\\run_all.py --config drivers\\config_canonical_v3.json --only pancancer,cnv,report")
```
