## Supplemental_Code for "scOPE identifies which driver-associated expression programs transfer from bulk tumors to single cells": repair_luad_obs_metadata.py

```
# repair_luad_obs_metadata.py

from pathlib import Path
import pandas as pd
import anndata as ad

ROOT = Path(r"C:\Users\ashfo\OneDrive\Desktop\Pathways+Omics\scOPE\scOPE\data")

ANNOT = ROOT / "LUAD_scRNA" / "GSE131907_cell_annotation.txt.gz"

TARGETS = [
    ROOT / "models" / "LUAD_scored_z_score_bulk_with_eval_labels.h5ad",
    ROOT / "scOPE_reviewer_derived" / "LUAD_sc_expression.h5ad",
]

def repair_one(h5ad_path: Path):
    print("\n" + "=" * 80)
    print("Reading:", h5ad_path)
    a = ad.read_h5ad(h5ad_path)

    print("AnnData:", a.shape)
    print("Current obs columns:", list(a.obs.columns))

    meta = pd.read_csv(ANNOT, sep="\t")
    print("Annotation:", meta.shape)
    print("Annotation columns:", meta.columns.tolist())

    # Most likely match: AnnData obs_names == annotation Index
    meta = meta.copy()
    meta["Index"] = meta["Index"].astype(str)
    meta = meta.set_index("Index", drop=False)

    obs_names = pd.Index(a.obs_names.astype(str))
    overlap = obs_names.intersection(meta.index)
    print(f"Overlap obs_names vs annotation Index: {len(overlap):,}/{a.n_obs:,}")

    if len(overlap) < 0.9 * a.n_obs:
        # fallback: try Barcode_Sample constructed from Barcode + Sample
        meta2 = meta.reset_index(drop=True).copy()
        meta2["Barcode_Sample"] = meta2["Barcode"].astype(str) + "_" + meta2["Sample"].astype(str)
        meta2 = meta2.set_index("Barcode_Sample", drop=False)
        overlap2 = obs_names.intersection(meta2.index)
        print(f"Overlap obs_names vs Barcode_Sample: {len(overlap2):,}/{a.n_obs:,}")

        if len(overlap2) > len(overlap):
            meta = meta2
            overlap = overlap2

    if len(overlap) < 0.9 * a.n_obs:
        print("First AnnData obs_names:", list(obs_names[:10]))
        print("First annotation index:", list(meta.index[:10]))
        raise RuntimeError(
            "Could not confidently match LUAD annotation to AnnData cells. "
            "Need to inspect obs_names format."
        )

    # Reindex annotation to AnnData order.
    meta_aligned = meta.loc[obs_names].copy()

    # Add useful obs fields.
    for col in ["Index", "Barcode", "Sample", "Sample_Origin", "Cell_type", "Cell_type.refined", "Cell_subtype"]:
        if col in meta_aligned.columns:
            a.obs[col] = meta_aligned[col].astype(str).values

    # Add normalized aliases expected by downstream scripts.
    a.obs["sample"] = meta_aligned["Sample"].astype(str).values
    a.obs["patient"] = meta_aligned["Sample"].astype(str).values
    a.obs["celltype"] = meta_aligned["Cell_type.refined"].astype(str).values
    a.obs["cell_type"] = meta_aligned["Cell_type.refined"].astype(str).values
    a.obs["cell_subtype"] = meta_aligned["Cell_subtype"].astype(str).values
    a.obs["sample_origin"] = meta_aligned["Sample_Origin"].astype(str).values

    out = h5ad_path.with_name(h5ad_path.stem + "_WITH_LUAD_OBS.h5ad")
    print("Writing:", out)
    a.write_h5ad(out)

    print("New obs columns:", list(a.obs.columns))
    print("Top celltype values:")
    print(a.obs["celltype"].value_counts().head(20).to_string())

for p in TARGETS:
    if p.exists():
        repair_one(p)
    else:
        print("Missing:", p)
```
