## Supplemental_Code for "scOPE identifies which driver-associated expression programs transfer from bulk tumors to single cells": replace_ensg_with_symbols.py

```
#!/usr/bin/env python
"""
replace_ensg_with_symbols.py

Convert Ensembl gene IDs (ENSG...) to readable HGNC symbols for scOPE analysis files,
using a local GENCODE GTF.

Works on:
  1) AnnData .h5ad files: replaces adata.var_names
  2) wide expression tables: samples x genes, replaces gene columns

It keeps original Ensembl IDs:
  - H5AD: adata.var["ensembl_id_original"]
  - TSV/CSV: writes a separate mapping CSV next to the output

IMPORTANT:
  For modelling, bulk and single-cell gene names must use the same naming convention.
  If AML bulk uses ENSG and AML single-cell uses symbols, convert the AML bulk expression
  table to symbols before running the suite. If both are ENSG, convert both.
"""

from __future__ import annotations

import argparse
import gzip
import re
from pathlib import Path
from typing import Dict, Iterable

import pandas as pd


ENSG_RE = re.compile(r"^(ENSG\d+)(?:\.\d+)?$")


def strip_ensembl_version(x: object) -> str:
    """ENSG000001234.5 -> ENSG000001234; leaves non-ENSG names unchanged."""
    s = str(x).strip()
    m = ENSG_RE.match(s)
    return m.group(1) if m else s


def is_probably_ensg(values: Iterable[object], n_check: int = 2000) -> bool:
    vals = [str(v).strip() for _, v in zip(range(n_check), values)]
    if not vals:
        return False
    n_ensg = sum(bool(ENSG_RE.match(v)) for v in vals)
    return (n_ensg / max(len(vals), 1)) > 0.5


def parse_gtf_attributes(attr: str) -> Dict[str, str]:
    out = {}
    for part in str(attr).split(";"):
        part = part.strip()
        if not part or " " not in part:
            continue
        key, val = part.split(" ", 1)
        out[key] = val.strip().strip('"')
    return out


def load_gencode_gene_map(gtf_path: str | Path) -> pd.DataFrame:
    """
    Return one row per Ensembl gene:
      ensembl_id, gene_symbol, gene_type
    """
    gtf_path = Path(gtf_path)
    opener = gzip.open if gtf_path.name.endswith(".gz") else open

    rows = []
    with opener(gtf_path, "rt", encoding="utf-8", errors="replace") as fh:
        for line in fh:
            if not line or line.startswith("#"):
                continue
            parts = line.rstrip("\n").split("\t")
            if len(parts) < 9:
                continue
            feature = parts[2]
            if feature != "gene":
                continue
            attrs = parse_gtf_attributes(parts[8])
            gene_id = attrs.get("gene_id")
            gene_name = attrs.get("gene_name")
            gene_type = attrs.get("gene_type") or attrs.get("gene_biotype")
            if gene_id and gene_name:
                rows.append(
                    {
                        "ensembl_id": strip_ensembl_version(gene_id),
                        "gene_symbol": gene_name,
                        "gene_type": gene_type or "",
                    }
                )

    gene_map = pd.DataFrame(rows).drop_duplicates("ensembl_id")
    if gene_map.empty:
        raise ValueError(f"No gene rows parsed from {gtf_path}")

    return gene_map


def make_unique(names: Iterable[object]) -> list[str]:
    """
    Make symbols unique for AnnData / matrix columns.
    TP53, TP53 -> TP53, TP53__2
    """
    seen = {}
    out = []
    for x in names:
        s = str(x).strip()
        n = seen.get(s, 0)
        seen[s] = n + 1
        out.append(s if n == 0 else f"{s}__{n + 1}")
    return out


def build_mapping(current_gene_ids: Iterable[object], gene_map: pd.DataFrame) -> pd.DataFrame:
    gm = gene_map.set_index("ensembl_id")
    rows = []
    for old in current_gene_ids:
        old_s = str(old).strip()
        ensg = strip_ensembl_version(old_s)
        if ensg in gm.index:
            symbol = str(gm.loc[ensg, "gene_symbol"])
            gene_type = str(gm.loc[ensg, "gene_type"])
            mapped = True
        else:
            symbol = old_s
            gene_type = ""
            mapped = False

        rows.append(
            {
                "old_gene_id": old_s,
                "ensembl_id": ensg if ensg.startswith("ENSG") else "",
                "gene_symbol": symbol,
                "gene_type": gene_type,
                "mapped": mapped,
            }
        )

    m = pd.DataFrame(rows)
    m["new_gene_name"] = make_unique(m["gene_symbol"])
    return m


def sep_for(path: str | Path) -> str:
    name = str(path).lower()
    return "\t" if name.endswith((".tsv", ".tsv.gz", ".txt", ".txt.gz")) else ","


def convert_h5ad(input_path: Path, output_path: Path, gene_map: pd.DataFrame) -> None:
    import anndata as ad

    print(f"[h5ad] reading: {input_path}")
    adata = ad.read_h5ad(input_path)

    mapping = build_mapping(adata.var_names, gene_map)
    n_mapped = int(mapping["mapped"].sum())
    print(f"[h5ad] mapped {n_mapped:,}/{adata.n_vars:,} var_names to symbols")

    adata.var["ensembl_id_original"] = mapping["old_gene_id"].to_numpy()
    adata.var["gene_symbol"] = mapping["gene_symbol"].to_numpy()
    adata.var["gene_type"] = mapping["gene_type"].to_numpy()
    adata.var_names = pd.Index(mapping["new_gene_name"].astype(str))
    adata.var_names_make_unique()

    output_path.parent.mkdir(parents=True, exist_ok=True)
    print(f"[h5ad] writing: {output_path}")
    adata.write_h5ad(output_path)

    map_path = output_path.with_suffix("").with_name(output_path.with_suffix("").name + "_gene_id_map.csv")
    mapping.to_csv(map_path, index=False)
    print(f"[h5ad] mapping: {map_path}")


def convert_wide_expression(input_path: Path, output_path: Path, gene_map: pd.DataFrame) -> None:
    """
    Convert a wide samples x genes expression table.
    Assumes first column is sample ID and remaining columns are genes.
    """
    sep = sep_for(input_path)
    print(f"[table] reading: {input_path} sep={repr(sep)}")
    df = pd.read_csv(input_path, sep=sep, index_col=0)

    if not is_probably_ensg(df.columns):
        print("[table] WARNING: columns do not look mostly like ENSG IDs.")
        print("[table] First columns:", list(map(str, df.columns[:10])))

    mapping = build_mapping(df.columns, gene_map)
    n_mapped = int(mapping["mapped"].sum())
    print(f"[table] mapped {n_mapped:,}/{df.shape[1]:,} gene columns to symbols")

    df.columns = mapping["new_gene_name"].astype(str).to_list()

    output_path.parent.mkdir(parents=True, exist_ok=True)
    out_sep = sep_for(output_path)
    print(f"[table] writing: {output_path} sep={repr(out_sep)}")
    df.to_csv(output_path, sep=out_sep, compression="infer")

    map_path = output_path.with_suffix("")
    if map_path.name.endswith(".tsv"):
        map_path = map_path.with_suffix("")
    map_path = map_path.with_name(map_path.name + "_gene_id_map.csv")
    mapping.to_csv(map_path, index=False)
    print(f"[table] mapping: {map_path}")


def main() -> None:
    ap = argparse.ArgumentParser()
    ap.add_argument("--gtf", required=True, help="Path to GENCODE GTF, e.g. data/gencode.v44.annotation.gtf.gz")
    ap.add_argument("--input", required=True, help="Input .h5ad, .tsv.gz, .tsv, .csv, or .csv.gz")
    ap.add_argument("--output", required=True, help="Output path")
    ap.add_argument(
        "--type",
        choices=["auto", "h5ad", "wide-expression"],
        default="auto",
        help="auto detects from extension; wide-expression means samples x genes table",
    )
    args = ap.parse_args()

    gtf = Path(args.gtf)
    inp = Path(args.input)
    out = Path(args.output)

    print(f"[map] loading GTF map: {gtf}")
    gene_map = load_gencode_gene_map(gtf)
    print(f"[map] loaded {len(gene_map):,} Ensembl -> symbol mappings")

    ftype = args.type
    if ftype == "auto":
        ftype = "h5ad" if inp.name.endswith(".h5ad") else "wide-expression"

    if ftype == "h5ad":
        convert_h5ad(inp, out, gene_map)
    elif ftype == "wide-expression":
        convert_wide_expression(inp, out, gene_map)
    else:
        raise ValueError(ftype)

    print("[done]")


if __name__ == "__main__":
    main()
```
