## Supplemental_Code for "scOPE identifies which driver-associated expression programs transfer from bulk tumors to single cells": __init__.py

```
"""
scope_analysis
==============
Manuscript-grade analysis utilities for scOPE outputs.

Submodules are lazy-loaded so lightweight imports/tests do not require the full
scope-bio/anndata stack. Driver scripts can still use ``scope_analysis.data`` or
``scope_analysis.plots``; the submodule imports happen on first access.
"""
from __future__ import annotations

import importlib

__version__ = "1.5.1"

from .config import AnalysisConfig

_SUBMODULES = {
    "config", "data", "stats", "bulk", "interpret", "singlecell",
    "longitudinal", "pancancer", "cnv", "robustness", "plots", "report",
    "confidence", "umap_panels", "annotate", "reference", "pathways", "priors", "identifiability",
}


def __getattr__(name: str):
    if name in _SUBMODULES:
        mod = importlib.import_module(f"{__name__}.{name}")
        globals()[name] = mod
        return mod
    raise AttributeError(f"module {__name__!r} has no attribute {name!r}")


__all__ = ["AnalysisConfig", "__version__", *_SUBMODULES]
```
