## Supplemental_Code for "scOPE identifies which driver-associated expression programs transfer from bulk tumors to single cells": annotate.py

```
"""
scope_analysis.annotate
========================
Offline, marker-based coarse cell-type annotation for single-cell objects that lack
a usable cell-type column (so the UMAP context panels and the reference definition
have something sensible to colour by). No network and no model downloads: each cell
is scored against curated canonical marker panels with ``scanpy.tl.score_genes`` and
assigned the highest-scoring lineage (with a low-confidence "Unassigned" fallback).

This is deliberately coarse (immune / stromal / epithelial lineages), which is all
that's needed to (a) give the UMAPs a legend and (b) define a lymphoid/myeloid
reference. For a publication-grade fine annotation, run CellTypist or the dataset's
own labels; this is the graceful fallback when those are absent.
"""
from __future__ import annotations

import numpy as np
import pandas as pd


# Canonical, lineage-level marker panels (human symbols). Intentionally broad.
MARKER_PANELS: dict[str, list[str]] = {
    "T cell":        ["CD3D", "CD3E", "CD3G", "TRAC", "CD2", "CD8A", "CD4", "IL7R"],
    "NK cell":       ["NKG7", "GNLY", "KLRD1", "NCAM1", "KLRF1", "PRF1"],
    "B cell":        ["MS4A1", "CD79A", "CD79B", "CD19", "BANK1"],
    "Plasma":        ["MZB1", "DERL3", "SLAMF7", "XBP1", "PRDM1", "IGHG1"],
    "Myeloid":       ["LYZ", "CD68", "CD14", "FCGR3A", "CSF1R", "ITGAM", "C1QA", "C1QB"],
    "Dendritic":     ["FCER1A", "CLEC9A", "LILRA4", "IRF7", "CD1C"],
    "Erythroid":     ["HBB", "HBA1", "HBA2", "GYPA", "ALAS2"],
    "HSPC/Progenitor": ["CD34", "KIT", "PROM1", "SPINK2", "AVP", "MEIS1", "HLF"],
    "Endothelial":   ["PECAM1", "VWF", "CLDN5", "CDH5", "FLT1"],
    "Fibroblast/CAF": ["COL1A1", "COL1A2", "DCN", "LUM", "PDGFRB", "ACTA2"],
    "Epithelial":    ["EPCAM", "KRT8", "KRT18", "KRT19", "CDH1", "KRT5"],
}

# which coarse lineages are "normal reference" candidates (non-malignant immune/stroma)
DEFAULT_REFERENCE_LINEAGES = ("T cell", "NK cell", "B cell", "Plasma", "Myeloid",
                              "Dendritic", "Endothelial", "Fibroblast/CAF", "Erythroid")


def marker_annotate(adata, out_col: str = "celltype_marker", panels: dict | None = None,
                    min_score: float = 0.0, layer: str | None = None) -> "object":
    """Assign each cell the lineage whose marker panel scores highest.

    Writes `out_col` (the lineage) and `{out_col}_score` (the winning score) to obs.
    Cells whose best score is <= min_score become 'Unassigned'. Returns adata (copy).
    Expects log-normalised expression in .X (or `layer`); if .X looks like raw counts
    a local normalisation+log1p is applied on a copy.
    """
    import scanpy as sc
    a = adata.copy()
    panels = panels or MARKER_PANELS

    X = a.layers[layer] if (layer and layer in a.layers) else a.X
    # heuristic: if values are large non-negative integers, normalise+log
    try:
        mx = float(X.max())
        looks_counts = mx > 50 and np.allclose(np.asarray(X[:10].todense() if hasattr(X, "todense")
                                                          else X[:10]) % 1, 0, atol=1e-6)
    except Exception:
        looks_counts = False
    if looks_counts:
        sc.pp.normalize_total(a, target_sum=1e4)
        sc.pp.log1p(a)

    var_names = set(map(str, a.var_names))
    scores = {}
    used = {}
    for lineage, markers in panels.items():
        present = [g for g in markers if g in var_names]
        if len(present) < 2:
            continue
        sc.tl.score_genes(a, present, score_name=f"_ms_{lineage}", use_raw=False)
        scores[lineage] = a.obs[f"_ms_{lineage}"].values
        used[lineage] = present
    if not scores:
        adata = adata.copy()
        adata.obs[out_col] = "Unassigned"
        adata.obs[f"{out_col}_score"] = np.nan
        print("[annotate] no marker panels had >=2 genes present; all Unassigned.")
        return adata

    S = pd.DataFrame(scores, index=a.obs_names)
    best = S.idxmax(axis=1)
    best_score = S.max(axis=1)
    best[best_score <= min_score] = "Unassigned"

    out = adata.copy()
    out.obs[out_col] = pd.Categorical(best.values)
    out.obs[f"{out_col}_score"] = best_score.values
    vc = out.obs[out_col].value_counts()
    print(f"[annotate] marker-based lineages (n={len(vc)}): "
          + ", ".join(f"{k}={v}" for k, v in vc.items()))
    print(f"[annotate]   panels used: {{"
          + ", ".join(f'{k}:{len(v)}g' for k, v in used.items()) + "}")
    return out


def ensure_celltype(adata, celltype_col: str = "CellType", min_categories: int = 2,
                    fallback_col: str = "celltype_marker"):
    """Return (adata, usable_celltype_col). If the requested cell-type column is
    missing or near-constant, compute a marker-based annotation and use that."""
    ok = (celltype_col in adata.obs.columns and
          adata.obs[celltype_col].astype(str).nunique() >= min_categories)
    if ok:
        return adata, celltype_col
    print(f"[annotate] '{celltype_col}' missing/uninformative; using marker-based annotation.")
    adata = marker_annotate(adata, out_col=fallback_col)
    return adata, fallback_col
```
