## Supplemental_Code for "scOPE identifies which driver-associated expression programs transfer from bulk tumors to single cells": bulk.py

```
"""
scope_analysis.bulk
===================
STAGE 1 of the manuscript story: "Bulk mutation classifiers recover driver-linked
expression programs" (Results 2.2, Fig. 2 / Table 1).

What it does
------------
For every cancer x driver pair it:
  1. trains the real ``scope.BulkPipeline`` (SVD + per-mutation logistic), and
  2. estimates discrimination by **leakage-free** stratified out-of-fold (OOF)
     cross-validation — the entire preprocess -> SVD -> classifier pipeline is
     refit inside each fold, so the latent space never sees held-out tumours, and
  3. wraps every OOF AUROC in a stratified-bootstrap CI and a label-permutation
     p-value (stats.py), then applies one global BH-FDR across all pairs, and
  4. gates each pair as "claim-safe" using your pre-specified rule
     (q<0.05 AND CI lower bound>0.5 AND >=min_pos cases AND >=min_neg controls).

Why OOF rather than scope-bio's ``cross_validate_classifiers``
--------------------------------------------------------------
``scope.evaluation.cross_validate_classifiers`` fixes one latent space computed on
ALL samples and cross-validates only the classifier. That is fine for a quick look
but lets the unsupervised SVD peek at the test fold. For a high-tier submission we
refit the whole pipeline per fold here. The fast path is still available via
``quick_cv_table`` for sanity checks.

Call hierarchy
--------------
    run_all / drivers -> evaluate_all_cancers_bulk
                          -> bulk_oof_predictions  (uses scope.BulkPipeline)
                          -> stats.bootstrap_auroc_ci / permutation_auroc_pvalue
                          -> stats.bh_fdr
    interpret.py      -> fit_bulk_pipeline (re-uses the same fitted object)
"""
from __future__ import annotations

import numpy as np
import pandas as pd
from sklearn.model_selection import StratifiedKFold

from scope import BulkPipeline
from scope.evaluation.metrics import cross_validate_classifiers

from . import stats as st
from .config import AnalysisConfig, cancer_drivers, prior_of


# ---------------------------------------------------------------------------
# Fit a single BulkPipeline on all data (used by interpret.py + final scoring)
# ---------------------------------------------------------------------------
def fit_bulk_pipeline(adata_bulk, labels: pd.DataFrame, cfg: AnalysisConfig) -> BulkPipeline:
    """Fit one scope-bio BulkPipeline on the full cohort with the config's
    primary settings. Returns the fitted object (also holds preprocessor_,
    decomposer_, classifier_set_ for interpretation and single-cell projection)."""
    bp = BulkPipeline(
        norm_method=cfg.norm_method,
        log1p=cfg.log1p,
        decomposition=cfg.decomposition,
        n_components=cfg.n_components,
        decomposition_kwargs=cfg.decomposition_kwargs,
        classifier=cfg.classifier,
        min_positive_frac=cfg.min_positive_frac,
    )
    bp.fit(adata_bulk, labels)
    return bp


# ---------------------------------------------------------------------------
# Leakage-free OOF predictions: refit the whole pipeline inside each fold
# ---------------------------------------------------------------------------
def bulk_auroc_map(adata_bulk, labels: pd.DataFrame, cfg: AnalysisConfig) -> dict:
    """Per-driver out-of-fold bulk AUROC as a {driver: auroc} dict, computed from
    the data without needing the full Fig2 bulk-reporting stage. This is what the
    confidence score's transferability (T) term needs; computing it here makes T
    available even when run_all is invoked with --only singlecell (which otherwise
    leaves the bulk discrimination table empty -> T collapses to its floor)."""
    from sklearn.metrics import roc_auc_score
    oof = bulk_oof_predictions(adata_bulk, labels, cfg)
    out = {}
    for d, sub in oof.groupby("driver"):
        m = sub.dropna(subset=["y_oof"])
        if m["y_true"].nunique() == 2 and len(m) >= 10:
            try:
                out[d] = float(roc_auc_score(m["y_true"], m["y_oof"]))
            except Exception:
                pass
    return out


def bulk_oof_predictions(
    adata_bulk,
    labels: pd.DataFrame,
    cfg: AnalysisConfig,
) -> pd.DataFrame:
    """Return a tidy DataFrame of pooled out-of-fold predictions.

    Columns: ['sample', 'driver', 'y_true', 'y_oof']. One row per (held-out
    sample, driver). The SVD + classifier are refit on the training portion of
    every fold, so y_oof for a sample is always produced by a model that never
    saw that sample.
    """
    skf = StratifiedKFold(n_splits=cfg.cv_folds, shuffle=True,
                          random_state=cfg.random_state)
    drivers = [d for d in labels.columns]
    samples = np.array(adata_bulk.obs_names)

    # Stratify folds on the first driver that has both classes; falls back to a
    # composite key so every fold is reasonably balanced across drivers.
    strat = _composite_stratifier(labels)
    oof = {d: np.full(adata_bulk.n_obs, np.nan) for d in drivers}

    for train_idx, test_idx in skf.split(samples, strat):
        train_samples = samples[train_idx]
        test_samples = samples[test_idx]
        a_tr = adata_bulk[train_samples].copy()
        y_tr = labels.loc[train_samples]
        bp = BulkPipeline(
            norm_method=cfg.norm_method, log1p=cfg.log1p,
            decomposition=cfg.decomposition, n_components=cfg.n_components,
            decomposition_kwargs=cfg.decomposition_kwargs,
            classifier=cfg.classifier, min_positive_frac=cfg.min_positive_frac,
        )
        bp.fit(a_tr, y_tr)
        a_te = adata_bulk[test_samples].copy()
        proba = bp.predict_bulk(a_te)  # columns: mutation_prob_<driver>
        for d in drivers:
            col = f"mutation_prob_{d}"
            if col in proba.columns:
                oof[d][test_idx] = proba[col].values

    rows = []
    for d in drivers:
        for i, s in enumerate(samples):
            rows.append((s, d, int(labels.iloc[i][d]), oof[d][i]))
    return pd.DataFrame(rows, columns=["sample", "driver", "y_true", "y_oof"])


def _composite_stratifier(labels: pd.DataFrame) -> np.ndarray:
    """Build a stratification vector. Prefer the most balanced driver; if none is
    usable, fall back to a hash of the full label row (keeps fold balance sane)."""
    best, best_bal = None, -1.0
    for d in labels.columns:
        f = labels[d].mean()
        bal = min(f, 1 - f)
        if bal > best_bal:
            best, best_bal = d, bal
    if best is not None and best_bal > 0:
        return labels[best].values
    return labels.astype(str).agg("".join, axis=1).values


# ---------------------------------------------------------------------------
# Per-pair metrics + uncertainty + significance (the Table 1 builder)
# ---------------------------------------------------------------------------
def evaluate_all_cancers_bulk(
    cohort_loader,
    cfg: AnalysisConfig,
) -> pd.DataFrame:
    """Run the full bulk evaluation across all configured cancers.

    Parameters
    ----------
    cohort_loader : callable
        ``cohort_loader(cancer) -> (adata_bulk, labels_df)``. You supply this so
        the suite stays agnostic to your exact file layout. See
        ``drivers/run_bulk.py`` for an example built on ``data.load_bulk_cohort``.

    Returns
    -------
    DataFrame (one row per cancer x driver) with AUROC + CI, AUPRC, perm p, the
    pre-specified prior tier, global BH-FDR q, and the claim_safe flag. This is
    Table 1 of the manuscript.
    """
    records = []
    oof_store = {}
    for cancer in cfg.cancers():
        try:
            adata_bulk, labels = cohort_loader(cancer)
        except (FileNotFoundError, ValueError) as e:
            print(f"[bulk] skipping {cancer}: {e}")
            continue
        # restrict to selected drivers that are actually present. Primary mode uses
        # the locked DRIVER_PRIORS list; exploratory mode can evaluate every mutation
        # label column in the file.
        if getattr(cfg, "score_all_label_drivers", False):
            wanted = list(labels.columns)
            mode = "all-label exploratory"
        else:
            wanted = cancer_drivers(cancer) if cfg.target_drivers == "all" else cfg.target_drivers
            mode = "primary/targeted"
        present = [d for d in wanted if d in labels.columns]
        labels = labels[present]
        if not present:
            print(f"[bulk] {cancer}: no selected drivers present, skipping.")
            continue

        print(f"[bulk] === {cancer}: {len(present)} drivers, {adata_bulk.n_obs} samples ({mode}) ===")
        oof = bulk_oof_predictions(adata_bulk, labels, cfg)
        oof_store[cancer] = oof

        for d in present:
            sub = oof[oof.driver == d].dropna(subset=["y_oof"])
            y, p = sub.y_true.values, sub.y_oof.values
            n_pos, n_neg = int(y.sum()), int((1 - y).sum())
            if n_pos < 2 or n_neg < 2:
                print(f"[bulk]   {cancer}/{d}: too few of one class (pos={n_pos}, neg={n_neg}); reporting NaN.")
                records.append(_empty_row(cancer, d, n_pos, n_neg))
                continue
            auc = st.bootstrap_auroc_ci(y, p, n_boot=cfg.n_boot,
                                        random_state=cfg.random_state)
            perm_p = st.permutation_auroc_pvalue(y, p, n_perm=cfg.n_perm,
                                                 random_state=cfg.random_state)
            records.append(dict(
                cancer=cancer, driver=d, prior=prior_of(cancer, d),
                n_pos=n_pos, n_neg=n_neg,
                auroc=auc.auroc, auroc_ci_lo=auc.ci_lo, auroc_ci_hi=auc.ci_hi,
                auprc=st.auprc_safe(y, p),
                auprc_baseline=float(np.mean(y)),  # prevalence baseline for AUPRC
                brier=float(np.mean((p - y) ** 2)),
                perm_p=perm_p,
            ))

    df = pd.DataFrame(records)
    if df.empty:
        return df
    # ONE global FDR correction across every tested pair
    df["fdr_q"] = st.bh_fdr(df["perm_p"].values)
    df["claim_safe"] = (
        (df["fdr_q"] < cfg.fdr_alpha)
        & (df["auroc_ci_lo"] > 0.5)
        & (df["n_pos"] >= cfg.min_pos)
        & (df["n_neg"] >= cfg.min_neg)
    )
    df = df.sort_values(["cancer", "auroc"], ascending=[True, False]).reset_index(drop=True)
    df.attrs["oof_store"] = oof_store  # stash OOF for ROC-curve plotting
    return df


def _empty_row(cancer, d, n_pos, n_neg):
    return dict(cancer=cancer, driver=d, prior=prior_of(cancer, d),
                n_pos=n_pos, n_neg=n_neg, auroc=np.nan, auroc_ci_lo=np.nan,
                auroc_ci_hi=np.nan, auprc=np.nan, auprc_baseline=np.nan,
                brier=np.nan, perm_p=np.nan)


# ---------------------------------------------------------------------------
# Fast sanity path (scope-bio native; latent space fit on all samples)
# ---------------------------------------------------------------------------
def quick_cv_table(adata_bulk, labels: pd.DataFrame, cfg: AnalysisConfig) -> pd.DataFrame:
    """Quick per-fold CV summary using scope-bio's own cross_validate_classifiers.

    Note: this fixes the latent space on all samples (mild optimism vs the
    leakage-free OOF path). Use only for fast iteration, not for the headline."""
    bp = BulkPipeline(norm_method=cfg.norm_method, log1p=cfg.log1p,
                      decomposition=cfg.decomposition, n_components=cfg.n_components,
                      decomposition_kwargs=cfg.decomposition_kwargs,
                      classifier=cfg.classifier, min_positive_frac=cfg.min_positive_frac)
    adata_pp = bp.preprocessor_ if hasattr(bp, "preprocessor_") else None
    bp.fit(adata_bulk, labels)
    Z = bp.transform_bulk(adata_bulk).obsm[bp.obsm_key_]
    import functools
    from scope.classification.models import get_classifier
    factory = functools.partial(get_classifier, cfg.classifier)
    cvres = cross_validate_classifiers(Z, labels, factory, cv=cfg.cv_folds,
                                       random_state=cfg.random_state)
    return cvres.groupby("driver" if "driver" in cvres.columns else "mutation").agg(
        auroc_mean=("auroc", "mean"), auroc_std=("auroc", "std"),
        auprc_mean=("auprc", "mean")).reset_index()
```
