## Supplemental_Code for "scOPE identifies which driver-associated expression programs transfer from bulk tumors to single cells": cnv.py

```
"""
scope_analysis.cnv
==================
EXPRESSION vs COPY-NUMBER COMPLEMENTARITY (Results 2.6, Fig. 6).

The chapter's claim is *complementarity*, not competition: scOPE should agree with
CNV-based malignant-cell calls where broad aneuploidy exists, AND recover signal
where CNV tools are blind (point-mutation-driven, near-diploid tumours). So this
module computes concordance (ARI) but treats DISCORDANCE as the scientifically
interesting output, and adds a continuous score-vs-burden correlation and
per-patient agreement.

It does NOT re-run infercnvpy / CopyKAT / Numbat. It ingests the CNV call tables
you already generate, in the tidy schema:
    columns: [dataset, barcode, method, cnv_score, malignant_call]
and joins them to scOPE scores by barcode using ROBUST matching (the common
failure is a barcode-format mismatch that silently yields zero overlap; here that
is detected, repaired where possible, and reported rather than passed over).

Call hierarchy
--------------
    drivers/run_cnv -> derive_scope_malignant_call
                     -> cnv_concordance            (ARI + confusion, robust join)
                     -> cnv_score_correlation       (continuous: scOPE vs CNV burden)
                     -> per_patient_malignant_fraction
                     -> discordance_analysis         (the complementarity story)
                     -> diploid_patient_standalone   (CNV-blind patients)
"""
from __future__ import annotations

import re

import numpy as np
import pandas as pd
from sklearn.metrics import adjusted_rand_score, confusion_matrix

from . import stats as st
from .config import AnalysisConfig
from .data import scored_drivers, is_usable_patient_series, patient_series_diagnostics


# ---------------------------------------------------------------------------
# Robust barcode matching
# ---------------------------------------------------------------------------
def _normalize_variants(bc: str) -> list[str]:
    """Return a list of normalized keys for a barcode, from strictest to loosest,
    so two formats can be matched even if one carries a sample prefix, a lane
    suffix, or different separators."""
    bc = str(bc)
    variants = [bc]
    # strip trailing -1 / _1 / .1 lane suffix
    stripped = re.sub(r"[-_.]\d+$", "", bc)
    variants.append(stripped)
    # last token after common separators (the raw 16bp-ish barcode)
    for sep in ("_", "-", ":", "#", "."):
        if sep in bc:
            variants.append(bc.split(sep)[-1])
            variants.append(re.sub(r"[-_.]\d+$", "", bc.split(sep)[-1]))
    # uppercase nucleotide core only
    core = re.sub(r"[^ACGTacgt]", "", bc).upper()
    if len(core) >= 12:
        variants.append(core)
    # dedupe preserving order
    seen, out = set(), []
    for v in variants:
        if v and v not in seen:
            seen.add(v); out.append(v)
    return out


def match_barcodes(obs_names, cnv_barcodes) -> tuple[dict, dict]:
    """Find the best obs<->cnv barcode correspondence by trying progressively
    looser normalizations and keeping the level with the most overlap.

    Returns (obs_to_cnv, diagnostics). obs_to_cnv maps an obs_name to the matched
    raw cnv barcode (for a left-join). diagnostics reports overlap counts so a
    near-zero match is visible rather than silent.
    """
    obs_names = list(obs_names)
    cnv_barcodes = list(cnv_barcodes)
    # Pre-index cnv by each of its variant keys -> raw barcode (first wins)
    levels = ["exact", "lane_stripped", "last_token", "nuc_core"]
    # Build per-level cnv lookups
    def level_key(bc, level):
        v = _normalize_variants(bc)
        # map level name to an index heuristically using the variant list
        if level == "exact":
            return v[0]
        if level == "lane_stripped":
            return re.sub(r"[-_.]\d+$", "", str(bc))
        if level == "last_token":
            s = str(bc)
            for sep in ("_", "-", ":", "#", "."):
                if sep in s:
                    return re.sub(r"[-_.]\d+$", "", s.split(sep)[-1])
            return re.sub(r"[-_.]\d+$", "", s)
        if level == "nuc_core":
            return re.sub(r"[^ACGTacgt]", "", str(bc)).upper()
        return str(bc)

    best = {"level": None, "obs_to_cnv": {}, "overlap": 0}
    for level in levels:
        cnv_lookup = {}
        for raw in cnv_barcodes:
            k = level_key(raw, level)
            if k and k not in cnv_lookup:
                cnv_lookup[k] = raw
        mapping = {}
        for o in obs_names:
            k = level_key(o, level)
            if k in cnv_lookup:
                mapping[o] = cnv_lookup[k]
        if len(mapping) > best["overlap"]:
            best = {"level": level, "obs_to_cnv": mapping, "overlap": len(mapping)}
        if best["overlap"] == len(obs_names):
            break
    diagnostics = {
        "n_obs": len(obs_names), "n_cnv": len(cnv_barcodes),
        "match_level": best["level"], "n_matched": best["overlap"],
        "frac_obs_matched": best["overlap"] / max(len(obs_names), 1),
    }
    return best["obs_to_cnv"], diagnostics


# ---------------------------------------------------------------------------
# Derive a scOPE malignant call from per-cell driver probabilities
# ---------------------------------------------------------------------------
def derive_scope_malignant_call(
    adata,
    cfg: AnalysisConfig,
    drivers: list[str] | None = None,
    threshold: float | None = None,
    out_col: str = "scope_malignant",
    reference_mask_col: str | None = "is_reference",
):
    """Call a cell malignant if its MAX RAW driver probability exceeds a threshold
    anchored to the reference upper quantile (not an arbitrary 0.5). Writes a 0/1
    obs column + the continuous max-score, and returns (adata, threshold)."""
    drivers = drivers or scored_drivers(adata)
    cols = [f"{cfg.raw_score_prefix}_{d}" for d in drivers
            if f"{cfg.raw_score_prefix}_{d}" in adata.obs.columns]
    adata = adata.copy()
    max_score = adata.obs[cols].astype(float).max(axis=1).values
    adata.obs["scope_max_driver_score"] = max_score
    if threshold is None:
        if reference_mask_col and reference_mask_col in adata.obs.columns and adata.obs[reference_mask_col].any():
            ref_scores = max_score[adata.obs[reference_mask_col].values]
            threshold = float(np.nanquantile(ref_scores[~np.isnan(ref_scores)], cfg.reference_quantile))
        else:
            threshold = float(np.nanquantile(max_score, 0.5))
    adata.obs[out_col] = (max_score > threshold).astype(int)
    print(f"[cnv] scOPE malignant call: threshold={threshold:.3f} "
          f"({int(adata.obs[out_col].sum())}/{adata.n_obs} cells malignant).")
    return adata, threshold


# ---------------------------------------------------------------------------
# Ingest CNV calls
# ---------------------------------------------------------------------------
def _clean_colname(c) -> str:
    """Normalize a column name for schema detection without changing the original."""
    return re.sub(r"[^a-z0-9]+", "_", str(c).strip().lower()).strip("_")


def _detect_column(df: pd.DataFrame, canonical: str, candidates: tuple[str, ...]) -> str | None:
    """Return the first plausible column for a canonical CNV field."""
    clean = {_clean_colname(c): c for c in df.columns}
    for cand in candidates:
        key = _clean_colname(cand)
        if key in clean:
            return clean[key]
    # fallback token search for messy exported column names
    for c in df.columns:
        cc = _clean_colname(c)
        if canonical == "barcode" and ("barcode" in cc or cc in {"cell", "cellid", "cell_id", "obs", "obs_name", "obs_names"}):
            return c
        if canonical == "malignant_call" and ("malig" in cc or "aneuploid" in cc or cc.endswith("call") or "prediction" in cc):
            return c
        if canonical == "cnv_score" and ("score" in cc or "burden" in cc or "cnv_signal" in cc):
            return c
        if canonical == "dataset" and cc in {"dataset", "cohort", "cancer", "tumor", "tumour"}:
            return c
        if canonical == "method" and cc in {"method", "tool", "caller", "algorithm"}:
            return c
    return None


def _coerce_malignant_call(x: pd.Series) -> pd.Series:
    """Map common CNV call encodings to 0/1, preserving NaN for unknown labels."""
    if pd.api.types.is_bool_dtype(x):
        return x.astype(float)
    if pd.api.types.is_numeric_dtype(x):
        # Accept 0/1 directly; if exported scores slipped into the call column,
        # values >0 are at least a deterministic malignant-ish fallback.
        return (pd.to_numeric(x, errors="coerce") > 0).astype(float)
    mapping = {
        "aneuploid": 1, "malignant": 1, "malig": 1, "malignant_like": 1,
        "malignant like": 1, "tumor": 1, "tumour": 1,
        "cancer": 1, "cancer cell": 1, "malignant cell": 1, "1": 1, "true": 1,
        "t": 1, "yes": 1, "y": 1, "positive": 1, "pos": 1, "cnv": 1,
        "diploid": 0, "normal": 0, "reference": 0, "reference_like": 0,
        "reference like": 0, "non-malignant": 0,
        "nonmalignant": 0, "non malignant": 0, "benign": 0, "0": 0,
        "false": 0, "f": 0, "no": 0, "n": 0, "negative": 0, "neg": 0,
        "not.defined": np.nan, "not defined": np.nan, "unknown": np.nan,
        "nan": np.nan, "na": np.nan, "": np.nan, "none": np.nan,
    }
    return x.astype(str).str.strip().str.lower().map(mapping).astype(float)


def normalize_cnv_calls(df: pd.DataFrame, dataset: str | None = None, source: str | None = None) -> pd.DataFrame:
    """Normalize many real-world CNV exports into the tidy schema used downstream.

    Required downstream columns are [barcode, malignant_call, cnv_score]. Optional
    columns [dataset, method] are preserved/created when possible. This function
    accepts common alternatives such as cell_id/cell/obs_name/Unnamed: 0 for the
    barcode and malignant/aneuploid/call/prediction for the CNV label.
    """
    df = df.copy()
    df.columns = [str(c).strip() for c in df.columns]

    # Detect/rename optional dataset before filtering.
    dataset_col = _detect_column(df, "dataset", ("dataset", "cohort", "cancer", "tumor", "tumour"))
    if dataset_col and dataset_col != "dataset":
        df = df.rename(columns={dataset_col: "dataset"})
    if dataset is not None and "dataset" in df.columns:
        ds = df["dataset"].astype(str).str.strip()
        target = str(dataset).strip()
        exact = ds.str.lower() == target.lower()
        if exact.any():
            df = df[exact].copy()
        else:
            # fallback: substring either direction, e.g. AML_vanGalen table vs AML request
            sub = ds.str.contains(target, case=False, na=False) | pd.Series(
                [target.lower() in str(v).lower() for v in ds], index=ds.index
            )
            if sub.any():
                df = df[sub].copy()

    barcode_col = _detect_column(df, "barcode", (
        "barcode", "barcodes", "cell_barcode", "cell barcode", "cell", "cell_id",
        "cellid", "cell_name", "cellname", "obs_name", "obs_names", "obs",
        "index", "Unnamed: 0", "X", "cell_id_original",
    ))
    if barcode_col is None:
        # Pandas-created RangeIndex is not useful, but a named/nontrivial index is.
        if df.index.name is not None or not isinstance(df.index, pd.RangeIndex):
            df = df.reset_index().rename(columns={df.reset_index().columns[0]: "barcode"})
            barcode_col = "barcode"
    if barcode_col is None:
        raise KeyError(
            "CNV call table has no recognizable barcode/cell-id column. "
            f"Available columns: {list(df.columns)}" + (f" Source: {source}" if source else "")
        )
    if barcode_col != "barcode":
        df = df.rename(columns={barcode_col: "barcode"})

    call_col = _detect_column(df, "malignant_call", (
        "malignant_call", "malignant", "is_malignant", "cnv_malignant",
        "aneuploid", "aneuploid_call", "cnv_call", "call", "prediction",
        "predicted_call", "malignant_status", "cnv_status", "state", "label",
    ))
    if call_col is None:
        # If no discrete call was exported, make a conservative continuous-score
        # threshold below after cnv_score detection.
        df["malignant_call"] = np.nan
    elif call_col != "malignant_call":
        df = df.rename(columns={call_col: "malignant_call"})

    score_col = _detect_column(df, "cnv_score", (
        "cnv_score", "cnv burden", "cnv_burden", "cnv_signal", "score",
        "cnv_score_mean", "malignancy_score", "aneuploidy_score", "cnv_leiden_score",
    ))
    if score_col and score_col != "cnv_score":
        df = df.rename(columns={score_col: "cnv_score"})
    if "cnv_score" not in df.columns:
        df["cnv_score"] = np.nan
    df["cnv_score"] = pd.to_numeric(df["cnv_score"], errors="coerce")

    if "malignant_call" in df.columns:
        coerced = _coerce_malignant_call(df["malignant_call"])
        # Fallback if a call column existed but values were not in the mapping.
        if coerced.notna().sum() == 0 and df["cnv_score"].notna().sum() >= 10 and df["cnv_score"].nunique(dropna=True) > 1:
            thr = float(np.nanquantile(df["cnv_score"].values, 0.95))
            coerced = (df["cnv_score"].astype(float) > thr).astype(float)
            print(f"[cnv] WARNING: could not parse malignant_call values; derived call from cnv_score > 95th pct ({thr:.4g}).")
        df["malignant_call"] = coerced

    method_col = _detect_column(df, "method", ("method", "tool", "caller", "algorithm"))
    if method_col and method_col != "method":
        df = df.rename(columns={method_col: "method"})
    if "method" not in df.columns:
        df["method"] = "cnv"
    if "dataset" not in df.columns and dataset is not None:
        df["dataset"] = dataset

    df["barcode"] = df["barcode"].astype(str).str.strip()
    df = df[df["barcode"].notna() & (df["barcode"].astype(str).str.len() > 0)].copy()
    if df["barcode"].duplicated().any():
        before = len(df)
        # Prefer rows with a parsed malignant call, then with a score.
        df["__has_call__"] = df["malignant_call"].notna().astype(int)
        df["__has_score__"] = df["cnv_score"].notna().astype(int)
        df = (df.sort_values(["barcode", "__has_call__", "__has_score__"], ascending=[True, False, False])
                .drop_duplicates("barcode", keep="first")
                .drop(columns=["__has_call__", "__has_score__"], errors="ignore"))
        print(f"[cnv] WARNING: dropped {before-len(df):,} duplicate CNV barcode rows after schema normalization.")

    keep_first = ["dataset", "barcode", "method", "cnv_score", "malignant_call"]
    rest = [c for c in df.columns if c not in keep_first]
    return df[[c for c in keep_first if c in df.columns] + rest]


def load_cnv_calls(csv_path, dataset: str | None = None) -> pd.DataFrame:
    """Load a CNV-call table and normalize it to
    [dataset, barcode, method, cnv_score, malignant_call].

    The loader is intentionally permissive because infercnvpy/CopyKAT/Numbat and
    ad-hoc merge scripts export different cell-id/call column names. It prints a
    concise schema summary so the selected barcode/call/score columns are visible
    before concordance is computed.
    """
    df = pd.read_csv(csv_path)
    out = normalize_cnv_calls(df, dataset=dataset, source=str(csv_path))
    n_call = int(out["malignant_call"].notna().sum()) if "malignant_call" in out.columns else 0
    print(f"[cnv] loaded {len(out):,} CNV rows from {csv_path}; "
          f"columns normalized to barcode/malignant_call/cnv_score "
          f"({n_call:,} parsed calls).")
    return out


def _merge_scope_cnv(adata, cnv_calls, scope_call_col, score_col=None):
    """Inner-join obs (scope) to cnv_calls via robust barcode matching.
    Returns (merged_df, diagnostics)."""
    if "barcode" not in cnv_calls.columns or "malignant_call" not in cnv_calls.columns:
        cnv_calls = normalize_cnv_calls(cnv_calls)
    obs = adata.obs.copy()
    obs["__obs_bc__"] = obs.index.astype(str)
    obs_to_cnv, diag = match_barcodes(obs["__obs_bc__"].values, cnv_calls["barcode"].astype(str).values)
    obs["__cnv_bc__"] = obs["__obs_bc__"].map(obs_to_cnv)
    take_cols = [c for c in ["barcode", "malignant_call", "cnv_score"] if c in cnv_calls.columns]
    if "cnv_score" not in take_cols:
        cnv_calls = cnv_calls.copy()
        cnv_calls["cnv_score"] = np.nan
        take_cols.append("cnv_score")
    cnv_small = cnv_calls[take_cols].copy()
    cnv_small["barcode"] = cnv_small["barcode"].astype(str)
    keep_obs = [c for c in [scope_call_col, "scope_max_driver_score", score_col, "patient"] if c and c in obs.columns]
    merged = (obs.dropna(subset=["__cnv_bc__"])[["__cnv_bc__"] + keep_obs]
              .merge(cnv_small, left_on="__cnv_bc__", right_on="barcode", how="inner"))
    return merged, diag


# ---------------------------------------------------------------------------
# Concordance: ARI + confusion (robust join, never silently empty)
# ---------------------------------------------------------------------------
def cnv_concordance(adata, cnv_calls: pd.DataFrame, scope_call_col: str = "scope_malignant") -> dict:
    """Adjusted Rand Index + confusion between scOPE and CNV malignant labels on
    matched cells. Reports the barcode-match diagnostics so a format mismatch is
    visible. Returns ari, n, fractions, confusion counts, the merged frame, and
    `match_diagnostics`."""
    merged, diag = _merge_scope_cnv(adata, cnv_calls, scope_call_col)
    merged = merged.dropna(subset=[scope_call_col, "malignant_call"])
    if merged.empty:
        print(f"[cnv] WARNING: 0 cells matched between scored obs and CNV table "
              f"(match level={diag['match_level']}, obs={diag['n_obs']}, cnv={diag['n_cnv']}). "
              f"Check that the CNV 'barcode' column uses the same cell IDs as the .h5ad obs_names.")
        return dict(ari=np.nan, n=0, match_diagnostics=diag, note="no overlapping barcodes")
    a = merged[scope_call_col].astype(int).values
    b = merged["malignant_call"].astype(int).values
    cm = confusion_matrix(b, a, labels=[0, 1])
    print(f"[cnv] matched {len(merged)} cells (level={diag['match_level']}, "
          f"{diag['frac_obs_matched']*100:.0f}% of obs); ARI={adjusted_rand_score(b, a):.3f}")
    return dict(
        ari=float(adjusted_rand_score(b, a)), n=int(len(merged)),
        scope_malignant_frac=float(a.mean()), cnv_malignant_frac=float(b.mean()),
        cnv_normal_scope_normal=int(cm[0, 0]), cnv_normal_scope_malignant=int(cm[0, 1]),
        cnv_malignant_scope_normal=int(cm[1, 0]), cnv_malignant_scope_malignant=int(cm[1, 1]),
        merged=merged, match_diagnostics=diag,
    )


# ---------------------------------------------------------------------------
# Continuous agreement: scOPE max-driver score vs CNV burden score
# ---------------------------------------------------------------------------
def cnv_score_correlation(adata, cnv_calls: pd.DataFrame) -> dict:
    """Spearman correlation between the continuous scOPE max-driver score and the
    continuous CNV burden score across matched cells. A positive correlation is
    complementary evidence that the two malignancy signals co-vary; a weak one
    motivates the discordance analysis."""
    merged, diag = _merge_scope_cnv(adata, cnv_calls, "scope_malignant", score_col=None)
    merged = merged.dropna(subset=["scope_max_driver_score", "cnv_score"])
    if len(merged) < 10 or merged["cnv_score"].nunique() < 3:
        return dict(spearman_rho=np.nan, spearman_p=np.nan, n=len(merged),
                    note="insufficient matched cells or constant cnv_score")
    from scipy.stats import spearmanr
    rho, p = spearmanr(merged["scope_max_driver_score"], merged["cnv_score"])
    return dict(spearman_rho=float(rho), spearman_p=float(p), n=int(len(merged)),
                match_level=diag["match_level"])


# ---------------------------------------------------------------------------
# Per-patient malignant-fraction agreement
# ---------------------------------------------------------------------------
def per_patient_malignant_fraction(adata, cnv_calls: pd.DataFrame,
                                   scope_call_col: str = "scope_malignant",
                                   patient_col: str = "patient",
                                   min_cells_per_patient: int = 2) -> pd.DataFrame:
    """Per patient, the fraction of cells called malignant by scOPE vs by CNV.

    Refuses to aggregate when ``patient_col`` is actually a per-cell barcode column.
    That exact failure mode creates huge tables with one cell per "patient" and
    scatter plots pinned at 0/1, which are not biologically interpretable.
    """
    merged, _ = _merge_scope_cnv(adata, cnv_calls, scope_call_col)
    if merged.empty or patient_col not in merged.columns:
        return pd.DataFrame()
    merged = merged.dropna(subset=[scope_call_col, "malignant_call", patient_col])
    if merged.empty:
        return pd.DataFrame()
    if not is_usable_patient_series(merged[patient_col], colname=patient_col):
        d = patient_series_diagnostics(merged[patient_col], colname=patient_col)
        print(f"[cnv] per-patient malignant fraction skipped: '{patient_col}' looks like cell IDs "
              f"(n_unique={d['n_unique']}, median_cells/id={d['median_cells_per_id']:.1f}, "
              f"barcode_like={d['barcode_like_frac']:.2f}). Repair sample/patient metadata first.")
        return pd.DataFrame()
    g = merged.groupby(patient_col).agg(
        n_cells=("malignant_call", "size"),
        scope_malignant_frac=(scope_call_col, "mean"),
        cnv_malignant_frac=("malignant_call", "mean"),
        mean_cnv_score=("cnv_score", "mean"),
    ).reset_index()
    g = g[g["n_cells"] >= int(min_cells_per_patient)].copy()
    if g.empty:
        print(f"[cnv] per-patient malignant fraction skipped: no patients had >= {min_cells_per_patient} matched cells.")
        return pd.DataFrame()
    return g.sort_values("cnv_malignant_frac", ascending=False)


# ---------------------------------------------------------------------------
# Discordance: the complementarity story
# ---------------------------------------------------------------------------
def discordance_analysis(merged: pd.DataFrame, scope_call_col: str = "scope_malignant") -> pd.DataFrame:
    """Four agreement quadrants with counts, mean CNV score, and mean scOPE
    max-driver score per quadrant. Surfaces scOPE+/CNV- (candidate CNV-blind
    malignant) and scOPE-/CNV+ (aneuploid, off-programme) explicitly."""
    has_score = "scope_max_driver_score" in merged.columns
    q = []
    for sc in (0, 1):
        for cn in (0, 1):
            sub = merged[(merged[scope_call_col] == sc) & (merged["malignant_call"] == cn)]
            name = {(1, 0): "scope+ / CNV- (candidate CNV-blind malignant)",
                    (0, 1): "scope- / CNV+ (aneuploid, off-programme)",
                    (1, 1): "concordant malignant",
                    (0, 0): "concordant normal"}[(sc, cn)]
            row = dict(quadrant=name, scope_malignant=sc, cnv_malignant=cn, n_cells=len(sub),
                       mean_cnv_score=float(sub["cnv_score"].mean()) if len(sub) else np.nan)
            if has_score:
                row["mean_scope_score"] = float(sub["scope_max_driver_score"].mean()) if len(sub) else np.nan
            q.append(row)
    return pd.DataFrame(q)


# ---------------------------------------------------------------------------
# Standalone value where CNV is blind: near-diploid patients
# ---------------------------------------------------------------------------
def diploid_patient_standalone(
    adata, cnv_calls: pd.DataFrame, patient_status: pd.DataFrame, cfg: AnalysisConfig,
    patient_col: str = "patient", diploid_cnv_frac_max: float = 0.10,
) -> pd.DataFrame:
    """In near-diploid patients (CNV calls almost no cells malignant), test whether
    scOPE patient-level scores still separate known-mutant from WT patients — the
    cleanest complementarity claim (signal where CNV is blind)."""
    merged, _ = _merge_scope_cnv(adata, cnv_calls, "scope_malignant")
    if merged.empty or patient_col not in merged.columns:
        return pd.DataFrame()
    merged = merged.dropna(subset=[patient_col])
    if merged.empty or not is_usable_patient_series(merged[patient_col], colname=patient_col):
        print(f"[cnv] near-diploid standalone skipped: '{patient_col}' is absent or barcode-like.")
        return pd.DataFrame()
    pat_cnv_frac = merged.dropna(subset=["malignant_call"]).groupby(patient_col)["malignant_call"].mean()
    diploid_patients = pat_cnv_frac[pat_cnv_frac <= diploid_cnv_frac_max].index.tolist()
    print(f"[cnv] near-diploid patients (CNV malignant frac <= {diploid_cnv_frac_max}): {len(diploid_patients)}")
    if not diploid_patients:
        return pd.DataFrame()
    obs = adata.obs
    rows = []
    for d in scored_drivers(adata):
        sc = cfg.score_col(adata, d)
        cell_df = obs[obs[patient_col].isin(diploid_patients)][[patient_col, sc]]
        agg = st.patient_aggregate(cell_df, patient_col, sc, min_cells=cfg.min_cells_per_patient)
        status = patient_status[patient_status.mutation == d][["patient", "label"]]
        m = agg.merge(status, left_on=patient_col, right_on="patient", how="inner")
        mut, wt = m[m.label == 1][sc].values, m[m.label == 0][sc].values
        if len(mut) < 2 or len(wt) < 2:
            continue
        eff = st.mannwhitney_effect(mut, wt, n_boot=cfg.n_boot, random_state=cfg.random_state)
        rows.append(dict(driver=d, n_diploid_mut=len(mut), n_diploid_wt=len(wt),
                         cliffs_delta=eff.cliffs_delta, mannwhitney_p=eff.p_value,
                         delta_ci_lo=eff.delta_lo, delta_ci_hi=eff.delta_hi))
    out = pd.DataFrame(rows)
    if not out.empty:
        out["fdr_q"] = st.bh_fdr(out["mannwhitney_p"].values)
    return out
```
