## Supplemental_Code for "scOPE identifies which driver-associated expression programs transfer from bulk tumors to single cells": cohort_labels.py

```
"""Canonical human-readable cohort labels for manuscript displays."""

COHORT_DISPLAY_NAMES = {
    "AML_vanGalen": "AML van Galen",
    "BRCA_Wu": "BRCA Wu",
    "CRC_Lee": "CRC Lee",
    "GBM_Neftel": "GBM Neftel",
    "LUAD_Lambrechts": "LUAD Lambrechts",
    "PAAD_Moncada": "PAAD Moncada",
    "SKCM_JerbyArnon": "SKCM Jerby-Arnon",
}

def display_cohort_name(cohort_id: str) -> str:
    return COHORT_DISPLAY_NAMES.get(cohort_id, cohort_id.replace("_", " "))
```
