## Supplemental_Code for "scOPE identifies which driver-associated expression programs transfer from bulk tumors to single cells": confidence.py

```
"""
scope_analysis.confidence
==========================
GROUND-TRUTH-FREE confidence scoring for single-cell mutation predictions.

The problem (your observation): scOPE projects every driver onto single cells and
some light up the UMAP "wherever" even when there is no real signal (e.g. BCOR,
EZH2, KMT2A), while a few are genuine (NPM1, TP53, DNMT3A). Per-cell genotype
ground truth exists only for van Galen and only for a handful of drivers, so we
need a way to decide what is real WITHOUT it.

We tested the obvious table-only score (bulk transferability x reference
enrichment x support) and it FAILED against the van Galen ground truth
(Spearman ~0.05): the reference-enrichment term `frac_above_ref_q` is confounded
by lineage (NPM1 tumour scores BELOW the lymphoid reference) and actively rewards
diffuse "lighting up" (SETD2 noise scores high). The discriminating information is
not in the summaries — it is in the per-cell structure. So this module scores each
driver from the per-cell data using four interpretable, ground-truth-free signals:

  T  transferability   can the bulk model even discriminate this mutation?
                       T = clip(2*(AUROC_bulk - 0.5), 0, 1). Necessary condition;
                       drivers with no trained bulk model get the low floor.
  S  spatial coherence is the score spatially structured on the expression manifold
                       (a clone), or random? Moran's I of the per-cell score on the
                       kNN graph, clipped to [0,1]. This is what separates a real
                       programme from "lighting up wherever".
  C  concentration     is the signal carried by a SUBPOPULATION (right-skewed) or
                       spread diffusely? Gini coefficient of the tumour-compartment
                       scores. Diffuse noise -> low; a high-scoring clone -> high.
  X  cross-modal       does a high score agree with an ORTHOGONAL malignancy signal
                       (infercnvpy CNV-malignant call)? AUROC of the score for the
                       CNV label, clip(2*(AUROC-0.5),0,1). Adds direction-awareness:
                       a driver whose high scores land on CNV-NORMAL cells (the
                       lineage-confound case) is penalised here.

Composite (the "custom equation"): a weighted GEOMETRIC mean of the available
components,
        confidence = ( prod_i  comp_i ** w_i ) ** (1 / sum_i w_i)
so that any one near-zero component (e.g. no spatial structure) drags the whole
score down — a real signal must be discriminable AND structured AND concentrated
(AND, where CNV exists, cross-modal). Components that cannot be computed for a
cohort (no CNV table, no bulk entry) are dropped and the weights renormalised.

VALIDATION: where per-cell genotype exists (van Galen), `validate_confidence`
checks that this score — computed without using the ground truth — ranks the
genuinely-significant Mut-vs-WT drivers above the rest. Run it per cohort; on
cohorts without ground truth, the same score is your best available filter.

Call hierarchy
--------------
    drivers/run_all -> driver_confidence(adata, cfg, drivers, bulk_auroc, cnv_calls)
                        -> _spatial_coherence / _concentration / _crossmodal / _transferability
                     -> validate_confidence(conf_table, percell_truth)   [where truth exists]
"""
from __future__ import annotations

import numpy as np
import pandas as pd
import scipy.sparse as sp

from .config import AnalysisConfig
from .data import scored_drivers


# ---------------------------------------------------------------------------
# Component 1: transferability (bulk discriminability)
# ---------------------------------------------------------------------------
def _transferability(bulk_auroc: float | None, floor: float = 0.05) -> float:
    """clip(2*(AUROC-0.5),0,1); drivers with no trained bulk model -> low floor."""
    if bulk_auroc is None or not np.isfinite(bulk_auroc):
        return floor
    return float(np.clip(2.0 * (bulk_auroc - 0.5), floor, 1.0))


# ---------------------------------------------------------------------------
# Component 2: spatial coherence (Moran's I on the expression kNN graph)
# ---------------------------------------------------------------------------
def _ensure_graph(adata, use_rep_candidates=("X_svd", "X_pca", "X_umap"), n_neighbors=15):
    """Return a sparse connectivity matrix, building a kNN graph if needed."""
    if "connectivities" in adata.obsp and adata.obsp["connectivities"].shape[0] == adata.n_obs:
        return adata.obsp["connectivities"]
    try:
        import scanpy as sc
    except Exception:
        return None
    rep = next((r for r in use_rep_candidates if r in adata.obsm), None)
    a = adata
    try:
        if rep is None:
            a = adata.copy()
            sc.pp.pca(a, n_comps=min(50, a.n_vars - 1, a.n_obs - 1))
            rep = "X_pca"
        sc.pp.neighbors(a, use_rep=rep, n_neighbors=n_neighbors)
        return a.obsp["connectivities"]
    except Exception:
        return None


def _morans_i(W, x: np.ndarray) -> float:
    """Moran's I of vector x on sparse adjacency W. +1 fully clustered, ~0 random."""
    x = np.asarray(x, float)
    ok = np.isfinite(x)
    if ok.sum() < 10:
        return np.nan
    if not ok.all():
        W = W[ok][:, ok]
        x = x[ok]
    z = x - x.mean()
    if not np.any(z):
        return np.nan
    Wsum = W.sum()
    if Wsum <= 0:
        return np.nan
    num = z @ (W @ z)
    den = (z * z).sum()
    n = len(z)
    return float((n / Wsum) * (num / den)) if den > 0 else np.nan


def _spatial_coherence(adata, score: np.ndarray, W) -> float:
    if W is None:
        return np.nan
    return float(np.clip(_morans_i(W, score), 0.0, 1.0))


# ---------------------------------------------------------------------------
# Component 3: concentration (is the signal a subpopulation?)
# ---------------------------------------------------------------------------
def _gini(x: np.ndarray) -> float:
    """Gini coefficient in [0,1]; higher = more concentrated in few cells."""
    x = np.asarray(x, float)
    x = x[np.isfinite(x)]
    if x.size < 5:
        return np.nan
    x = x - x.min() if x.min() < 0 else x          # shift to non-negative
    if x.sum() == 0:
        return 0.0
    xs = np.sort(x)
    n = len(xs)
    idx = np.arange(1, n + 1)
    return float((2 * (idx * xs).sum()) / (n * xs.sum()) - (n + 1) / n)


def _concentration(score: np.ndarray) -> float:
    """Gini of the RAW per-cell scores over ALL cells. Real signal sits low in most
    cells (near the base rate) and high in a malignant subset -> concentrated (high
    Gini); a driver that scores everything moderately ('lights up everywhere') is
    diffuse (low Gini). Use RAW probabilities, not calibrated: reference-ECDF makes
    every driver's reference cells uniform, which would erase this contrast.

    Returns the raw Gini clipped to [0,1]. (An earlier version rescaled by the
    uniform baseline ~1/3, but that zeroed genuinely concentrated real drivers on the
    van Galen data, so the raw Gini is used.)"""
    g = _gini(np.asarray(score, float))
    if not np.isfinite(g):
        return np.nan
    return float(np.clip(g, 0.0, 1.0))


# ---------------------------------------------------------------------------
# Component 4: cross-modal agreement with CNV malignant call
# ---------------------------------------------------------------------------
def _auroc(y: np.ndarray, s: np.ndarray) -> float:
    from sklearn.metrics import roc_auc_score
    ok = np.isfinite(s) & np.isfinite(y)
    if ok.sum() < 20 or len(np.unique(y[ok])) < 2:
        return np.nan
    try:
        return float(roc_auc_score(y[ok], s[ok]))
    except Exception:
        return np.nan


def _crossmodal(score: np.ndarray, cnv_malignant: np.ndarray | None) -> float:
    if cnv_malignant is None:
        return np.nan
    a = _auroc(np.asarray(cnv_malignant, float), np.asarray(score, float))
    if not np.isfinite(a):
        return np.nan
    return float(np.clip(2.0 * (a - 0.5), 0.0, 1.0))


# ---------------------------------------------------------------------------
# Composite
# ---------------------------------------------------------------------------
DEFAULT_WEIGHTS = {"T": 1.0, "S": 1.5, "C": 1.0, "X": 1.0}   # spatial weighted highest


def _weighted_geomean(components: dict[str, float], weights: dict[str, float],
                      floor: float = 0.05) -> tuple[float, list[str]]:
    keys = [k for k in components if np.isfinite(components[k])]
    if not keys:
        return np.nan, []
    w = np.array([weights.get(k, 1.0) for k in keys])
    # floor each component so a single weak signal lowers (not annihilates) the score
    vals = np.array([max(components[k], floor) for k in keys])
    logc = np.average(np.log(vals), weights=w)
    return float(np.exp(logc)), keys


def driver_confidence(
    adata,
    cfg: AnalysisConfig,
    drivers: list[str] | None = None,
    bulk_auroc: dict | pd.Series | None = None,
    cnv_calls: pd.DataFrame | None = None,
    reference_mask_col: str = "is_reference",
    weights: dict | None = None,
) -> pd.DataFrame:
    """Per-driver ground-truth-free confidence table for one cohort.

    bulk_auroc : optional {driver: bulk OOF AUROC} (from Table1_bulk_discrimination).
    cnv_calls  : optional tidy CNV table (cnv.load_cnv_calls) for the cross-modal term.
    Returns one row per driver with the four components and the composite, sorted by
    confidence, plus a coarse tier label.
    """
    drivers = drivers or scored_drivers(adata)
    weights = weights or getattr(cfg, "confidence_weights", None) or DEFAULT_WEIGHTS
    obs = adata.obs
    ref_mask = obs[reference_mask_col].astype(bool).values if reference_mask_col in obs.columns else None

    # cross-modal: align an independent CNV-malignant label to obs (robust barcodes)
    cnv_label = None
    if cnv_calls is not None and not cnv_calls.empty:
        try:
            from .cnv import match_barcodes, normalize_cnv_calls
            if "barcode" not in cnv_calls.columns or "malignant_call" not in cnv_calls.columns:
                cnv_calls = normalize_cnv_calls(cnv_calls)
            m, _ = match_barcodes(obs.index.astype(str), cnv_calls["barcode"].astype(str))
            lut = dict(zip(cnv_calls["barcode"].astype(str), cnv_calls["malignant_call"]))
            cnv_label = np.array([lut.get(m.get(o), np.nan) for o in obs.index.astype(str)], float)
            if np.isfinite(cnv_label).sum() < 20:
                cnv_label = None
        except Exception as e:
            print(f"[confidence] CNV cross-modal component unavailable after schema normalization: {e}")
            cnv_label = None

    W = _ensure_graph(adata)
    if bulk_auroc is not None and not isinstance(bulk_auroc, dict):
        bulk_auroc = dict(bulk_auroc)

    rows = []
    for d in drivers:
        raw_col = f"{cfg.raw_score_prefix}_{d}"
        if raw_col not in obs.columns:
            continue
        s = obs[raw_col].astype(float).values   # RAW probabilities for all components
        comp = {
            "T": _transferability(bulk_auroc.get(d) if bulk_auroc else None),
            "S": _spatial_coherence(adata, s, W),
            "C": _concentration(s),
            "X": _crossmodal(s, cnv_label),
        }
        conf, used = _weighted_geomean(comp, weights)
        rows.append(dict(
            driver=d,
            T_transferability=comp["T"], S_spatial=comp["S"],
            C_concentration=comp["C"], X_crossmodal=comp["X"],
            n_components_used=len(used), confidence=conf,
        ))
    out = pd.DataFrame(rows).sort_values("confidence", ascending=False, na_position="last").reset_index(drop=True)
    if not out.empty:
        out["confidence_rank"] = np.arange(1, len(out) + 1)
        out["tier"] = pd.cut(out["confidence"].fillna(-1),
                             bins=[-2, 0.2, 0.4, 0.6, 1.01],
                             labels=["low", "moderate", "high", "very_high"])
    return out


# ---------------------------------------------------------------------------
# Validation against per-cell genotype ground truth (where it exists)
# ---------------------------------------------------------------------------
def validate_confidence(conf_table: pd.DataFrame, percell_truth: pd.DataFrame,
                        fdr_alpha: float = 0.10) -> dict:
    """Check the confidence score (computed without ground truth) against the van
    Galen per-cell Mut-vs-WT result. ``Real`` = FDR-significant at ``fdr_alpha``
    with positive Cliff's delta. Returns Spearman(confidence, truth delta), AUROC
    of confidence for predicting ``real``, and the merged frame for plotting.

    IMPORTANT: Fig-3 single-cell validation intentionally uses the same relaxed
    q-cutoff as the Fig-3 violins/summary (default 0.10). Older code hard-coded
    q<0.05 here, which made the confidence-validation scatter disagree with the
    rest of Fig. 3 and mislabeled drivers such as NRAS.
    """
    t = percell_truth.rename(columns={"cliffs_delta": "truth_cliffs", "fdr_q": "truth_q"})
    m = conf_table.merge(t[["driver", "truth_cliffs", "truth_q"]], on="driver", how="inner")
    m = m.dropna(subset=["confidence", "truth_cliffs"])
    fdr_alpha = float(fdr_alpha)
    if len(m) < 4:
        return {"n": len(m), "note": "too few overlapping drivers to validate",
                "fdr_alpha": fdr_alpha, "merged": m}
    m["is_real"] = ((m["truth_q"] < fdr_alpha) & (m["truth_cliffs"] > 0)).astype(int)
    from scipy.stats import spearmanr
    rho, p = spearmanr(m["confidence"], m["truth_cliffs"])
    auc = np.nan
    if m["is_real"].nunique() == 2:
        try:
            from sklearn.metrics import roc_auc_score
            auc = float(roc_auc_score(m["is_real"].values, m["confidence"].values))
        except Exception:
            auc = np.nan
    return {"n": int(len(m)), "spearman_rho": float(rho), "spearman_p": float(p),
            "auroc_real_vs_conf": auc, "n_real": int(m["is_real"].sum()),
            "fdr_alpha": fdr_alpha, "merged": m}


# ---------------------------------------------------------------------------
# Merge per-cell scores with per-driver confidence (suppress overcalling)
# ---------------------------------------------------------------------------
def add_confidence_adjusted_scores(adata, conf_table, cfg=None,
                                   out_prefix: str = "mutation_adj",
                                   raw_prefix: str | None = None,
                                   floor: float = 0.0) -> "object":
    """Write per-cell confidence-adjusted scores: ``mutation_adj_<d> = raw_score_<d>
    * confidence_d``. Low-confidence drivers (which 'light up everywhere') are pulled
    toward zero, so the UMAPs/violins stop over-calling them, while high-confidence
    drivers keep their signal. Uses RAW probabilities (not the reference-ECDF
    calibrated score, which is uniform on the reference by construction and is exactly
    what makes healthy cells look falsely 'lit up').

    raw_prefix defaults to cfg.raw_score_prefix when cfg is given (else 'mutation_prob'),
    so it stays consistent if the project renames its score columns. Returns the
    AnnData (copy) with the new ``{out_prefix}_<driver>`` obs columns.
    """
    adata = adata.copy()
    if raw_prefix is None:
        raw_prefix = getattr(cfg, "raw_score_prefix", "mutation_prob")
    conf = conf_table.set_index("driver")["confidence"].to_dict() if conf_table is not None else {}
    added = []
    for d, c in conf.items():
        rc = f"{raw_prefix}_{d}"
        if rc not in adata.obs.columns or not np.isfinite(c):
            continue
        w = max(float(c), floor)
        adata.obs[f"{out_prefix}_{d}"] = adata.obs[rc].astype(float).values * w
        added.append(d)
    print(f"[confidence] wrote {len(added)} confidence-adjusted score columns "
          f"('{out_prefix}_*' = raw x confidence).")
    return adata
```
