## Supplemental_Code for "scOPE identifies which driver-associated expression programs transfer from bulk tumors to single cells": config.py

```
"""
scope_analysis.config
=====================
Single source of truth for: where the data lives, which cancers/drivers are in
play, and — critically — the *a priori* expected fingerprint strength of every
driver BEFORE any metric is computed.

Why the prior table matters
---------------------------
Your Chapter 3 makes a specific methodological promise (Results 2.1, Methods
"Driver selection and prior expectations"): drivers are graded strong /
intermediate / weak *before* performance is seen, so a reviewer cannot accuse you
of reporting only the winners. This module is where that pre-specification is
locked. ``report.py`` reads it back to build the driver-prior table (the chapter's
requested Table next to Fig. 1) and to colour results by expected strength.

The priors below are drawn directly from the biology cited in your Introduction
(NPM1->HOX, IDH1/2->epigenome via 2-HG, BRAF->MAPK, splicing factors->diffuse).
Edit the dictionaries to match your final locked driver list; nothing else in the
suite hard-codes a gene name.

Call hierarchy
--------------
    everything -> AnalysisConfig (paths, knobs)
    bulk/pancancer/report -> DRIVER_PRIORS, cancer_drivers(), prior_of()
"""
from __future__ import annotations

from dataclasses import dataclass, field
from pathlib import Path

# ---------------------------------------------------------------------------
# A-PRIORI DRIVER FINGERPRINT STRENGTH  (pre-specified; do NOT tune to results)
# ---------------------------------------------------------------------------
# tier ∈ {"strong", "intermediate", "weak"}.
# "weak" drivers are deliberately retained as negative/limit controls.
DRIVER_PRIORS: dict[str, dict[str, str]] = {
    # --- AML (BeatAML training; van Galen / Beneyto-Calabuig single-cell) ---
    "AML": {
        "NPM1":  "strong",        # cytoplasmic NPM1 -> characteristic HOX/MEIS1 program
        "IDH1":  "strong",        # neomorphic 2-HG -> broad epigenetic/transcriptional reprogramming
        "IDH2":  "strong",
        "FLT3":  "intermediate",  # FLT3-ITD signalling; signature present but heterogeneous
        "DNMT3A":"intermediate",  # global methylation drift; diffuse expression effect
        "RUNX1": "intermediate",
        "TP53":  "intermediate",
        "NRAS":  "intermediate",  # MAPK axis
        "KRAS":  "intermediate",
        "CEBPA": "intermediate",
        "SRSF2": "weak",          # splicing factor -> diffuse gene-level signature (limit control)
        "U2AF1": "weak",
        "SF3B1": "weak",
        "ASXL1": "weak",
    },
    # --- Colorectal (TCGA COAD/READ; Lee 2020 single-cell) ---
    "CRC": {
        "APC":   "strong",        # WNT activation
        "KRAS":  "strong",        # RAS programme
        "BRAF":  "strong",        # MAPK / serrated, MSI-linked
        "TP53":  "intermediate",
        "PIK3CA":"intermediate",
        "SMAD4": "intermediate",  # TGF-beta axis
        "FBXW7": "weak",
    },
    # --- Breast (TCGA BRCA; Wu 2021 single-cell) ---
    "BRCA": {
        "TP53":  "strong",        # tightly linked to basal-like state
        "PIK3CA":"strong",        # luminal-enriched
        "GATA3": "intermediate",
        "CDH1":  "intermediate",  # lobular
        "MAP3K1":"weak",
    },
    # --- Melanoma (TCGA SKCM; Jerby-Arnon 2018 single-cell) ---
    "SKCM": {
        "BRAF":  "strong",        # V600E -> constitutive MAPK
        "NRAS":  "strong",        # mutually exclusive MAPK class
        "NF1":   "intermediate",  # third MAPK-axis class
        "TP53":  "weak",
    },
    # --- Lung adenocarcinoma (TCGA LUAD; bulk benchmark + exploratory sc) ---
    "LUAD": {
        "EGFR":  "strong",
        "KRAS":  "strong",
        "STK11": "intermediate",
        "KEAP1": "intermediate",  # NRF2/oxidative-stress programme
        "TP53":  "intermediate",
    },
    # --- Glioblastoma (TCGA GBM; bulk benchmark + exploratory sc) ---
    "GBM": {
        "IDH1":  "strong",        # defines G-CIMP / proneural axis
        "EGFR":  "strong",        # classical / RTK
        "PDGFRA":"intermediate",
        "NF1":   "intermediate",  # mesenchymal
        "PTEN":  "weak",
    },
    # --- Pancreatic adenocarcinoma (TCGA PAAD; bulk benchmark + exploratory sc) ---
    "PAAD": {
        "KRAS":  "strong",        # near-universal background driver
        "TP53":  "intermediate",
        "CDKN2A":"intermediate",
        "SMAD4": "intermediate",
    },
}

# Validation tier per cancer: how strong is the single-cell ground truth?
#   "per_cell"        -> expressed-mutation calls at (near) single-cell resolution
#   "patient_level"   -> only patient/sample mutation status; aggregate cells first
#   "exploratory"     -> no single-cell genotype; bulk benchmark + score localisation only
VALIDATION_TIER: dict[str, str] = {
    "AML": "per_cell",          # van Galen expressed-mutation + patient summaries
    "CRC": "patient_level",
    "BRCA": "patient_level",
    "SKCM": "patient_level",
    "LUAD": "exploratory",
    "GBM": "exploratory",
    "PAAD": "exploratory",
}


def cancer_drivers(cancer: str) -> list[str]:
    return list(DRIVER_PRIORS.get(cancer, {}).keys())


def prior_of(cancer: str, driver: str) -> str:
    return DRIVER_PRIORS.get(cancer, {}).get(driver, "unspecified")


def all_cancers() -> list[str]:
    return list(DRIVER_PRIORS.keys())


# ---------------------------------------------------------------------------
# Analysis configuration (paths + every statistical / modelling knob)
# ---------------------------------------------------------------------------
@dataclass
class AnalysisConfig:
    # ---- where things are ----
    data_root: Path = Path("data")
    output_root: Path = Path("scope_analysis_outputs")

    # ---- scope-bio modelling defaults (the manuscript's primary configuration) ----
    norm_method: str = "none"          # set to "cpm" for raw counts; "none" for log2-TPM/RPKM bulk
    log1p: bool = False                # bulk already log-transformed (BeatAML log2(RPKM+1), TCGA log2(norm+1))
    already_log_transformed: bool = True
    decomposition: str = "svd"
    n_components: int = 50
    classifier: str = "logistic"
    alignment_method: str = "z_score_bulk"
    min_positive_frac: float = 0.05
    decomposition_kwargs: dict = field(default_factory=dict)

    # ---- statistical rigor knobs ----
    n_boot: int = 2000                 # bootstrap replicates for CIs
    n_perm: int = 2000                 # label-permutation replicates for nulls
    alpha: float = 0.05
    fdr_alpha: float = 0.10
    min_pos: int = 10                  # claim-safety: >=10 positives
    min_neg: int = 10                  # claim-safety: >=10 negatives
    cv_folds: int = 5
    random_state: int = 42

    # ---- single-cell knobs ----
    min_cells_per_patient: int = 10    # anti-pseudoreplication floor
    reference_quantile: float = 0.95   # healthy-reference threshold for frac_above
    healthy_celltypes: tuple[str, ...] = ("T", "B", "NK", "CTL", "Plasma")

    # ---- scope-bio single-cell preprocessing overrides ----
    # The upstream scope.SingleCellPipeline constructs a SingleCellPreprocessor with
    # raw-droplet defaults unless told otherwise.  That is dangerous when rescoring
    # already-curated h5ads: default QC/doublet/mito filters can remove most cells
    # and change the population used by all downstream plots.  Put kwargs here that
    # should be forwarded to scope.preprocessing.SingleCellPreprocessor, e.g.
    # {"filter_strategy":"none", "already_qc_filtered":true,
    #  "already_normalized":true, "already_log_transformed":true,
    #  "run_doublet_detection":false}.
    singlecell_preprocessing: dict | None = None

    # What to do if projection produces non-finite latent values before classifier
    # prediction.  "zero_fill" replaces NaN/Inf with 0 in bulk-latent space
    # (neutral/mean component value) and reports the count; "median_fill" uses
    # component medians; "raise" fails fast.
    singlecell_projection_nan_policy: str = "zero_fill"

    # ---- single-cell score CALIBRATION ----
    # scope-bio writes mutation_prob_<gene> from predict_proba. For single cells
    # these are genuine probabilities but compress near the base rate (the
    # projected cells land near the latent origin), so a [0,1] plot looks flat.
    # We add a monotonic, reference-anchored calibrated score in [0,1] that is
    # readable AND preserves every rank-based statistic (AUROC, Cliff's, MWU, KS,
    # Spearman are identical under a monotone transform). The raw probability is
    # always retained alongside.
    # Submission-safe default is RAW scores. Reference-ECDF calibration can be useful
    # as an explicitly labelled visualization, but it makes healthy reference cells
    # uniform on [0,1] by construction and can confuse driver/reference diagnostics.
    use_calibrated_scores: bool = False
    calibration_method: str = "reference_ecdf"  # "reference_ecdf" | "minmax" | "rank"
    raw_score_prefix: str = "mutation_prob"
    calibrated_score_prefix: str = "mutation_calib"

    # Longitudinal analyses are deliberately raw by default, even if general plotting
    # calibration is enabled. Treatment trajectories should not be driven by a
    # reference-ECDF transform that redistributes healthy cells across [0,1].
    longitudinal_use_calibrated_scores: bool = False

    # Plot-only scaling. Statistics always use the selected score column unchanged.
    # "auto_robust" keeps absolute raw values in the colorbar but clips color limits
    # to q01-q99 when raw probabilities are compressed (e.g. NPM1 at 0.01-0.04).
    plot_score_scale: str = "auto_robust"  # "auto_robust" | "absolute" | "fixed01"
    plot_robust_lower_q: float = 0.01
    plot_robust_upper_q: float = 0.99
    plot_robust_min_range: float = 0.05
    # Residual score display guards.  Same-cell-state residuals can have tiny numeric
    # ranges for weak/exploratory drivers; without a minimum symmetric scale the UMAPs
    # make numerical dust look like biology.  These are plot-only limits.
    plot_residual_min_abs_vmax: float = 0.05
    plot_residual_abs_vmax: float = 0.25
    heatmap_residual_abs_vmax: float = 0.20
    heatmap_score_scale: str = "global_robust"  # "global_robust" | "driver_robust" | "absolute"
    confidence_display_min: float = 0.35  # drop drivers below this confidence from heatmaps
    # When all-label exploratory scoring is enabled, keep main Fig4 panels from turning
    # into visual confetti by default.  Use "all" for full exploratory atlases.
    plot_driver_set: str = "primary_if_all_labels"  # "primary_if_all_labels" | "primary" | "all" | "confidence"
    # Which cells are greyed in score UMAPs/triptychs:
    #   "reference"      -> only the true reference/healthy baseline (least aggressive; default)
    #   "normal_display" -> all annotated normal/context lineages, e.g. AML non-"-like" cells
    #   "none"           -> color every cell by score
    # Older v9.1 switched automatically to "normal_display" for AML, which was too
    # aggressive and hid a lot of biologically useful context.
    score_umap_grey_mask: str = "reference"

    # ---- reference/cell-state guardrails ----
    # These protect NPM1-like drivers where healthy HSPC/myeloid cells share a
    # developmental programme and can score above AML cells on a global reference
    # comparison. They never change raw scores; they add diagnostic/residual columns.
    enable_reference_guardrails: bool = True
    reference_guardrail_samples: tuple[str, ...] = ("BM1", "BM2", "BM3", "BM4")
    cellstate_col: str = "celltype_coarse"
    cellstate_min_reference_cells: int = 25
    longitudinal_guarded_mode: str = "cellstate_residual"  # "cellstate_residual" | "cellstate_percentile" | "none"

    # ---- driver-selection mode ----
    # Primary/manuscript mode uses the locked DRIVER_PRIORS table. Exploratory mode
    # uses every binary mutation column present in the bulk label matrix. This is
    # useful for broad discovery/rescoring, but tables mark unspecified genes as
    # exploratory rather than pre-specified.
    score_all_label_drivers: bool = False

    # ---- interpretation depth: how many top importance-ranked SVD components and
    #      genes per component to mine/plot for Fig2 SVD interpretation.
    #      interpret_top_components controls the CSV/table depth; plot/heatmap knobs
    #      keep figures readable when the tables are intentionally deep. ----
    interpret_top_components: int = 25
    interpret_genes_per_component: int = 50
    interpret_plot_top_components: int = 6
    interpret_plot_genes_per_component: int = 30
    interpret_heatmap_top_components: int = 10
    interpret_heatmap_genes_per_component: int = 15

    # ---- memory safeguard: cap single cells scored per cohort (0 = no cap) ----
    max_singlecell_cells: int = 0

    # ---- confidence scoring (ground-truth-free signal/noise filter). Component
    #      weights for the weighted geometric mean: T=bulk transferability,
    #      S=spatial coherence (Moran's I), C=concentration (Gini), X=cross-modal CNV. ----
    confidence_weights: dict | None = None

    # ---- malignant-compartment definition (for within-tumour analyses) ----
    # Comparing the whole non-reference compartment to a lineage-mismatched
    # reference (e.g. lymphoid T/B/NK vs myeloid AML blasts) is confounded. When a
    # malignant call is available (CNV or scOPE-derived) the longitudinal / shift
    # analyses can be restricted to malignant cells. This quantile sets the
    # scOPE-derived malignant threshold off the reference distribution.
    malignant_call_quantile: float = 0.95
    restrict_longitudinal_to_malignant: bool = True

    # ---- ablation grids (Methods: alignment / latent-dim / classifier / decomposition) ----
    ablation_alignments: tuple[str, ...] = ("z_score_bulk", "moment_matching", "quantile", "none")
    ablation_n_components: tuple[int, ...] = (1, 5, 10, 25, 50, 75, 100, 150, 250, 500)
    ablation_classifiers: tuple[str, ...] = ("logistic", "random_forest", "gbm")
    ablation_decompositions: tuple[str, ...] = ("svd", "nmf", "pca", "ica")

    target_cancers: tuple[str, ...] | str = "all"
    target_drivers: tuple[str, ...] | str = "all"

    @property
    def score_prefix(self) -> str:
        """Prefix of the obs score column the main analysis should use.

        When ``use_calibrated_scores`` is false, every cfg-aware table/plot resolves
        to raw ``mutation_prob_*`` columns even if stale ``mutation_calib_*`` columns
        are present on a reused h5ad.
        """
        return self.calibrated_score_prefix if self.use_calibrated_scores else self.raw_score_prefix

    def cancers(self) -> list[str]:
        return all_cancers() if self.target_cancers == "all" else list(self.target_cancers)

    def outdir(self, *parts: str) -> Path:
        p = self.output_root.joinpath(*parts)
        p.mkdir(parents=True, exist_ok=True)
        return p

    def figdir(self, *parts: str) -> Path:
        """Organized figure subdirectory, e.g. cfg.figdir('fig3b_longitudinal',
        'AML556'). Keeps the output tree navigable instead of one flat dump."""
        p = self.output_root.joinpath("figures", *parts)
        p.mkdir(parents=True, exist_ok=True)
        return p

    def raw_score_col(self, adata, driver: str) -> str:
        """Raw scope-bio predict_proba column for a driver."""
        return f"{self.raw_score_prefix}_{driver}"

    def calibrated_score_col(self, adata, driver: str) -> str:
        """Reference/rank calibrated score column for a driver."""
        return f"{self.calibrated_score_prefix}_{driver}"

    def score_col(self, adata, driver: str, *, use_calibrated: bool | None = None) -> str:
        """Resolve which obs column to use for a driver's per-cell score.

        Centralizing this prevents stale ``mutation_calib_*`` columns from leaking
        into plots/tables when the JSON sets ``use_calibrated_scores=false``.
        """
        use_cal = self.use_calibrated_scores if use_calibrated is None else bool(use_calibrated)
        calib = self.calibrated_score_col(adata, driver)
        raw = self.raw_score_col(adata, driver)
        if use_cal and calib in adata.obs.columns:
            return calib
        return raw

    def longitudinal_score_col(self, adata, driver: str) -> str:
        """Score column for treatment/timepoint analyses. Raw by default."""
        return self.score_col(adata, driver, use_calibrated=self.longitudinal_use_calibrated_scores)

    def score_kind(self, col: str) -> str:
        if str(col).startswith(f"{self.calibrated_score_prefix}_"):
            return "calibrated_reference_percentile"
        if str(col).startswith(f"{self.raw_score_prefix}_"):
            return "raw_probability"
        if str(col).startswith("mutation_adj_"):
            return "confidence_adjusted_raw"
        if str(col).startswith("mutation_resid_"):
            return "cellstate_residual_raw"
        if str(col).startswith("mutation_refpct_"):
            return "cellstate_reference_percentile"
        return "score"

    def score_label_for_col(self, col: str, driver: str | None = None) -> str:
        gene = driver or str(col).split("_")[-1]
        kind = self.score_kind(col)
        if kind == "calibrated_reference_percentile":
            return f"{gene} reference-percentile score"
        if kind == "raw_probability":
            return f"{gene} raw driver-program score"
        if kind == "confidence_adjusted_raw":
            return f"{gene} confidence-adjusted raw score"
        if kind == "cellstate_residual_raw":
            return f"{gene} same-cell-state residual"
        if kind == "cellstate_reference_percentile":
            return f"{gene} same-cell-state reference percentile"
        return f"{gene} score"
```
