## Supplemental_Code for "scOPE identifies which driver-associated expression programs transfer from bulk tumors to single cells": data.py

```
"""
scope_analysis.data
====================
All disk I/O and metadata wrangling. Two responsibilities:

1. Load the inputs scope-bio needs (bulk AnnData + binary mutation-label table)
   and the single-cell objects you score.
2. Parse van Galen sample identifiers into (patient, timepoint) so the
   longitudinal treatment-response analysis can group cells correctly.

This module deliberately supports BOTH paths:
  (A) train-from-bulk: load raw bulk -> BulkPipeline (real transfer learning), and
  (B) ingest-scored:   load an already-scored .h5ad whose obs already carries
      ``mutation_prob_<GENE>`` columns (the objects you have on disk, e.g.
      ``AML_scored_z_score_bulk_with_eval_labels.h5ad``).
Both feed the identical downstream evaluation, so you are never locked into one.

Call hierarchy
--------------
    bulk.py        -> load_bulk_cohort
    singlecell.py  -> load_scored_singlecell, derive_patient_timepoint, mark_reference_cells
    longitudinal.py-> load_scored_singlecell, derive_patient_timepoint
    pancancer.py   -> load_scored_singlecell, load_patient_status
"""
from __future__ import annotations

import re
from pathlib import Path

import numpy as np
import pandas as pd

try:
    import anndata as ad
except ImportError:  # pragma: no cover - anndata is a hard dep on the analysis box
    ad = None


# ---------------------------------------------------------------------------
# Bulk (for the train-from-bulk path)
# ---------------------------------------------------------------------------
def load_bulk_cohort(
    expression_path: str | Path,
    mutation_path: str | Path,
    sample_col: str = "sample_id",
    genes_as_columns: bool = True,
) -> tuple["ad.AnnData", pd.DataFrame]:
    """Load a bulk cohort as (AnnData, binary-label DataFrame) ready for
    ``BulkPipeline.fit``.

    Robust version for the local manuscript inputs:
    - .tsv/.tsv.gz/.txt/.txt.gz are read with tab separators
    - sample/gene ids are stripped of whitespace
    - exact join is tried first; conservative normalized join is tried second
    """
    expression_path = Path(expression_path)
    if expression_path.suffix == ".h5ad":
        adata = ad.read_h5ad(expression_path)
        adata.obs_names = adata.obs_names.astype(str).str.strip()
        adata.var_names = adata.var_names.astype(str).str.strip()
    else:
        df = _read_wide_expression_table(expression_path)
        if not genes_as_columns:
            df = df.T
        adata = ad.AnnData(X=df.values.astype(np.float32))
        adata.obs_names = df.index.astype(str)
        adata.var_names = df.columns.astype(str)

    labels = _read_label_table(mutation_path, sample_col)
    adata, labels = _align_bulk_and_labels(
        adata=adata,
        labels=labels,
        expression_name=expression_path.name,
        mutation_name=Path(mutation_path).name,
    )
    print(f"[data] {expression_path.name}: {adata.n_obs} matched samples, "
          f"{adata.n_vars} genes, {labels.shape[1]} drivers.")
    return adata, labels


def _read_label_table(path: str | Path, sample_col: str) -> pd.DataFrame:
    """Read a mutation-label CSV/TSV and coerce to strict 0/1 ints.

    Accepts 0/1, mutant/WT, yes/no, true/false, pos/neg in any casing.
    """
    path = Path(path)
    sep = _sep_for_table(path)
    df = pd.read_csv(path, sep=sep)
    idx_col = _choose_label_index_column(df, sample_col)
    df = df.set_index(idx_col)
    df.index = df.index.astype(str).str.strip()

    # Keep only putative driver columns; this prevents metadata columns from
    # becoming fake all-zero mutation labels if present.
    metadata_like = {
        "original_sample_id", "sample_id", "sample", "barcode", "patient", "patient_id",
        "case_id", "cancer", "cohort", "dataset"
    }
    keep_cols = [c for c in df.columns if str(c).strip() not in metadata_like]
    df = df[keep_cols]
    return df.apply(_coerce_binary)


def _sep_for_table(path: str | Path) -> str:
    """Infer separator from filename."""
    name = Path(path).name.lower()
    return "\t" if name.endswith((".tsv", ".tsv.gz", ".txt", ".txt.gz")) else ","


def _read_wide_expression_table(path: str | Path) -> pd.DataFrame:
    """Read wide expression matrix with auto TSV/CSV separator detection.

    Some compressed TCGA/derived matrices on OneDrive occasionally throw a low-level
    C parser read error (``Calling read(nbytes) on source failed``).  Retry with
    pandas' Python engine before giving up so a transient/parser-specific failure
    does not kill the whole run.
    """
    path = Path(path)
    sep = _sep_for_table(path)
    try:
        df = pd.read_csv(path, sep=sep, index_col=0)
    except Exception as e:
        msg = str(e)
        retryable = (
            "Calling read(nbytes) on source failed" in msg
            or "Error tokenizing data" in msg
            or "Try engine='python'" in msg
        )
        if not retryable:
            raise
        print(f"[data] WARNING: C parser failed reading {path.name}; retrying with engine='python' ({e})")
        df = pd.read_csv(path, sep=sep, index_col=0, engine="python")
    df.index = df.index.astype(str).str.strip()
    df.columns = df.columns.astype(str).str.strip()
    return df


def _normalize_sample_id(x) -> str:
    """Conservative sample-id normalizer for TCGA/BeatAML-style joins."""
    s = str(x).strip()
    s = re.sub(r"^X", "", s)  # R sometimes prefixes sample IDs
    s = s.replace(".", "-").replace("_", "-")
    s = re.sub(r"\s+", "", s)
    s = s.upper()

    # TCGA barcode sample-level matching: TCGA-XX-YYYY-01A-... -> TCGA-XX-YYYY-01
    if s.startswith("TCGA-"):
        parts = s.split("-")
        if len(parts) >= 4:
            sample_type = parts[3][:2]
            s = "-".join(parts[:3] + [sample_type])
    return s


def _make_unique_index(values) -> pd.Index:
    """Make duplicate normalized IDs unique without silently collapsing rows."""
    seen = {}
    out = []
    for v in values:
        v = str(v)
        n = seen.get(v, 0)
        seen[v] = n + 1
        out.append(v if n == 0 else f"{v}__DUP{n}")
    return pd.Index(out)


def _choose_label_index_column(df: pd.DataFrame, requested: str) -> str:
    """Pick a plausible sample-id column for mutation labels."""
    if requested in df.columns:
        return requested
    candidates = [
        "sample_id", "sample", "Sample", "SAMPLE", "id", "ID",
        "barcode", "Barcode", "BARCODE", "patient_id", "patient", "case_id",
        "tumor_sample_barcode", "Tumor_Sample_Barcode", "TumorSampleBarcode",
    ]
    for c in candidates:
        if c in df.columns:
            print(f"[data] sample_col {requested!r} absent; using detected column {c!r}.")
            return c
    print(f"[data] sample_col {requested!r} absent; using first column {df.columns[0]!r}.")
    return df.columns[0]


def _align_bulk_and_labels(
    adata: "ad.AnnData",
    labels: pd.DataFrame,
    expression_name: str,
    mutation_name: str,
) -> tuple["ad.AnnData", pd.DataFrame]:
    """Exact sample-id join first; then normalized join with diagnostics."""
    shared = adata.obs_names.intersection(labels.index)
    if len(shared) > 0:
        return adata[shared].copy(), labels.loc[shared].copy()

    original_expr_ids = pd.Index(adata.obs_names.astype(str))
    original_label_ids = pd.Index(labels.index.astype(str))
    expr_norm = pd.Index([_normalize_sample_id(x) for x in original_expr_ids])
    lab_norm = pd.Index([_normalize_sample_id(x) for x in original_label_ids])

    adata = adata.copy()
    labels = labels.copy()
    adata.obs["original_sample_id"] = original_expr_ids.to_numpy()
    labels["original_sample_id"] = original_label_ids.to_numpy()
    adata.obs_names = _make_unique_index(expr_norm)
    labels.index = _make_unique_index(lab_norm)

    shared = adata.obs_names.intersection(labels.index)
    if len(shared) == 0:
        print("[data] DEBUG expression sample examples:", list(original_expr_ids[:10]))
        print("[data] DEBUG label sample examples:", list(original_label_ids[:10]))
        print("[data] DEBUG normalized expression examples:", list(expr_norm[:10]))
        print("[data] DEBUG normalized label examples:", list(lab_norm[:10]))
        raise ValueError(
            f"No shared sample ids between {expression_name} and {mutation_name}, "
            "even after TSV/CSV detection and conservative ID normalization. "
            "Check sample_col and whether expression rows are samples rather than genes."
        )

    print(
        f"[data] normalized sample-id join rescued {len(shared)} shared samples "
        f"between {expression_name} and {mutation_name}."
    )
    labels = labels.loc[shared].drop(columns=["original_sample_id"], errors="ignore")
    return adata[shared].copy(), labels.copy()


def _coerce_binary(col: pd.Series) -> pd.Series:
    if pd.api.types.is_numeric_dtype(col):
        return (col > 0).astype(int)
    mapping = {
        "1": 1, "0": 0, "mut": 1, "mutant": 1, "wt": 0, "wildtype": 0,
        "wild-type": 0, "yes": 1, "no": 0, "true": 1, "false": 0,
        "pos": 1, "positive": 1, "neg": 0, "negative": 0, "present": 1, "absent": 0,
    }
    return col.astype(str).str.strip().str.lower().map(mapping).fillna(0).astype(int)


# ---------------------------------------------------------------------------
# Single-cell (for the ingest-scored path)
# ---------------------------------------------------------------------------
def load_scored_singlecell(h5ad_path: str | Path) -> "ad.AnnData":
    """Load a scored single-cell AnnData and assert it carries scope-bio scores.

    Raises early with a clear message if no ``mutation_prob_*`` columns are
    present, so you never silently analyse an unscored object.
    """
    adata = ad.read_h5ad(h5ad_path)
    prob_cols = [c for c in adata.obs.columns if c.startswith("mutation_prob_")]
    if not prob_cols:
        raise ValueError(
            f"{Path(h5ad_path).name} has no mutation_prob_* columns in obs. "
            "Score it first with SingleCellPipeline.transform (see singlecell.py)."
        )
    print(f"[data] {Path(h5ad_path).name}: {adata.n_obs} cells, "
          f"{len(prob_cols)} scored drivers: "
          f"{[c.replace('mutation_prob_','') for c in prob_cols]}")
    return adata


def scored_drivers(adata: "ad.AnnData") -> list[str]:
    return [c.replace("mutation_prob_", "")
            for c in adata.obs.columns if c.startswith("mutation_prob_")]


# ---------------------------------------------------------------------------
# Robust patient/sample metadata helpers
# ---------------------------------------------------------------------------
_CELL_ID_NAME_RE = re.compile(r"(barcode|cell[_ -]?id|cellid|raw_barcode|obs_name|obs_names|index)$", re.I)
_NUC_BARCODE_RE = re.compile(r"^[ACGTN]{10,}(?:[-_.][0-9]+)?$", re.I)


def _looks_like_cell_barcode_value(x) -> bool:
    """Heuristic for single-cell barcodes, including sample-prefixed barcodes."""
    s = str(x).strip()
    if not s or s.lower() in {"nan", "none", "na"}:
        return False
    parts = re.split(r"[_:#|]", s)
    if any(_NUC_BARCODE_RE.match(p or "") for p in parts):
        return True
    core = re.sub(r"[^ACGTNacgtn]", "", s)
    return len(core) >= 14 and (len(core) / max(len(s), 1)) > 0.75


def patient_series_diagnostics(values, colname: str | None = None) -> dict:
    """Return lightweight diagnostics used to reject cell barcodes as patients."""
    s = pd.Series(values).dropna().astype(str).str.strip()
    s = s[~s.str.lower().isin({"", "nan", "none", "na"})]
    n = int(len(s))
    if n == 0:
        return dict(n=0, n_unique=0, unique_frac=np.nan, median_cells_per_id=0.0,
                    barcode_like_frac=0.0, name_is_cell_id=False, usable=False)
    counts = s.value_counts()
    sample = s.sample(min(len(s), 5000), random_state=0) if len(s) > 5000 else s
    barcode_like_frac = float(sample.map(_looks_like_cell_barcode_value).mean())
    name_is_cell_id = bool(colname and _CELL_ID_NAME_RE.search(str(colname)))
    d = dict(
        n=n,
        n_unique=int(counts.size),
        unique_frac=float(counts.size / max(n, 1)),
        median_cells_per_id=float(counts.median()) if len(counts) else 0.0,
        barcode_like_frac=barcode_like_frac,
        name_is_cell_id=name_is_cell_id,
    )
    d["usable"] = is_usable_patient_series(s, colname=colname, diagnostics=d)
    return d


def is_usable_patient_series(values, colname: str | None = None, diagnostics: dict | None = None) -> bool:
    """True when a vector is plausible patient/sample metadata, not cell barcodes.

    This intentionally rejects the failure mode that produced LUAD plots with
    ~40k "patients": a patient column equal to per-cell barcodes. A real patient or
    sample column should have repeated labels across cells; a barcode column has an
    almost-one-to-one label/cell ratio and often nucleotide-looking values.
    """
    if diagnostics is None:
        s = pd.Series(values).dropna().astype(str).str.strip()
        s = s[~s.str.lower().isin({"", "nan", "none", "na"})]
        n = int(len(s))
        if n == 0:
            return False
        counts = s.value_counts()
        sample = s.sample(min(len(s), 5000), random_state=0) if len(s) > 5000 else s
        diagnostics = dict(
            n=n,
            n_unique=int(counts.size),
            unique_frac=float(counts.size / max(n, 1)),
            median_cells_per_id=float(counts.median()) if len(counts) else 0.0,
            barcode_like_frac=float(sample.map(_looks_like_cell_barcode_value).mean()),
            name_is_cell_id=bool(colname and _CELL_ID_NAME_RE.search(str(colname))),
        )
    n = diagnostics.get("n", 0)
    n_unique = diagnostics.get("n_unique", 0)
    unique_frac = diagnostics.get("unique_frac", 1.0)
    med_cells = diagnostics.get("median_cells_per_id", 0.0)
    barcode_like_frac = diagnostics.get("barcode_like_frac", 0.0)
    name_is_cell_id = diagnostics.get("name_is_cell_id", False)
    if n == 0 or n_unique == 0:
        return False
    # One label per cell (or nearly so) is a cell-id/barcode, not a patient/sample.
    if unique_frac > 0.50 and med_cells <= 2:
        return False
    # A high-cardinality nucleotide-looking column is almost always a barcode.
    if barcode_like_frac >= 0.50 and (unique_frac > 0.20 or med_cells <= 3):
        return False
    if name_is_cell_id and (unique_frac > 0.20 or med_cells <= 3):
        return False
    # Very high cardinality can be valid for sample barcodes in a huge atlas, but not
    # for these patient-level single-cell cohort plots. Keep this permissive enough
    # for dozens of patients while still blocking 40k-cell barcode legends.
    if n_unique > max(250, int(0.10 * n)) and med_cells <= 5:
        return False
    return True


def choose_patient_column(obs: pd.DataFrame, configured: str | None = None,
                          out_patient_col: str = "patient") -> tuple[str | None, dict]:
    """Pick the best existing obs column to represent patients/samples.

    The configured sample column has first priority, then common sample/donor names,
    then an existing patient column. Candidate columns that look like per-cell
    barcodes are rejected with diagnostics.
    """
    candidates: list[str] = []
    if configured and str(configured).strip():
        candidates.append(str(configured).strip())
    candidates.extend([
        "sample", "Sample", "sample_id", "SampleID", "sampleID", "orig.ident", "orig_ident",
        "donor", "Donor", "donor_id", "case", "Case", "case_id", "CaseID",
        out_patient_col, "Patient", "patient_id", "PatientID", "patientID",
        "individual", "Individual", "subject", "Subject", "library", "sample_name",
    ])
    # Preserve order, handle case-insensitive aliases.
    real_cols = {str(c): c for c in obs.columns}
    low_to_col = {str(c).lower(): c for c in obs.columns}
    seen = set()
    rejected = []
    for cand in candidates:
        if cand in seen:
            continue
        seen.add(cand)
        col = real_cols.get(cand, low_to_col.get(cand.lower()))
        if col is None or col not in obs.columns or not obs[col].notna().any():
            continue
        diag = patient_series_diagnostics(obs[col], colname=str(col))
        if diag["usable"]:
            return str(col), {"chosen": str(col), **diag, "rejected": rejected}
        rejected.append({"column": str(col), **diag})
    return None, {"chosen": None, "rejected": rejected}


def derive_patient_timepoint(
    adata: "ad.AnnData",
    sample_col: str = "Sample",
    patient_strip_regex: str = r"-D\d+$",
    timepoint_regex: str = r"-(D\d+)$",
    out_patient_col: str = "patient",
    out_timepoint_col: str = "timepoint",
) -> "ad.AnnData":
    """Split a van-Galen-style sample id (e.g. ``AML556-D31``) into a patient
    (``AML556``) and a timepoint (``D31``).

    Cells whose sample id has no ``-D<n>`` suffix (e.g. normal BM controls like
    ``BM1``) get timepoint ``NaN`` and patient = the sample id unchanged, so they
    remain usable as the healthy reference.

    Returns the AnnData with two new obs columns; original is left untouched
    (operates on a copy).
    """
    adata = adata.copy()
    if sample_col not in adata.obs.columns:
        raise KeyError(f"sample_col '{sample_col}' not in obs. "
                       f"Available: {list(adata.obs.columns)[:20]}")
    samp = adata.obs[sample_col].astype(str)
    adata.obs[out_patient_col] = samp.str.replace(patient_strip_regex, "", regex=True)
    tp = samp.str.extract(timepoint_regex)[0]
    adata.obs[out_timepoint_col] = tp  # NaN where no timepoint suffix
    n_tp = adata.obs[out_timepoint_col].notna().sum()
    print(f"[data] derived patient/timepoint: "
          f"{adata.obs[out_patient_col].nunique()} patients, "
          f"{adata.obs[out_timepoint_col].dropna().nunique()} distinct timepoints, "
          f"{n_tp}/{adata.n_obs} cells carry a timepoint.")
    return adata


def filter_samples(adata: "ad.AnnData", sample_col: str = "Sample",
                   exclude_samples=None, exclude_regex: str | None = None,
                   keep_samples=None) -> "ad.AnnData":
    """Drop cells by sample id. Use to remove contaminating samples that aren't
    representative normals or patients — e.g. van Galen's sorted-progenitor controls
    BM5-34p / BM5-34p38n (CD34+ / CD34+CD38- fractions, which transcriptionally
    resemble blasts and get false-positive mutation calls) and cell lines like
    MUTZ3 / OCI-AML3. exclude_samples: exact ids; exclude_regex: pattern (e.g. '^BM5');
    keep_samples: if given, keep ONLY these (applied after excludes)."""
    if sample_col not in adata.obs.columns or (not exclude_samples and not exclude_regex and not keep_samples):
        return adata
    samp = adata.obs[sample_col].astype(str)
    mask = np.ones(adata.n_obs, dtype=bool)
    if exclude_samples:
        mask &= ~samp.isin([str(s) for s in exclude_samples]).values
    if exclude_regex:
        mask &= ~samp.str.contains(exclude_regex, regex=True, na=False).values
    if keep_samples:
        mask &= samp.isin([str(s) for s in keep_samples]).values
    dropped = int((~mask).sum())
    if dropped:
        kept_samps = sorted(samp[mask].unique().tolist())
        print(f"[data] filter_samples: dropped {dropped}/{adata.n_obs} cells; "
              f"{len(kept_samps)} samples remain.")
    return adata[mask].copy()


def ensure_patient_metadata(
    adata: "ad.AnnData",
    sample_col: str | None = "Sample",
    patient_strip_regex: str = r"-D\d+$",
    timepoint_regex: str = r"-(D\d+)$",
    out_patient_col: str = "patient",
    out_timepoint_col: str = "timepoint",
) -> "ad.AnnData":
    """Create stable patient/timepoint columns without assuming one dataset schema.

    This version explicitly guards against a common LUAD/single-cell failure mode:
    an obs column named ``patient`` that actually contains one cell barcode per row.
    Such columns are rejected and replaced by a real repeated sample/donor column
    when one is available; otherwise patient-level plots are skipped instead of
    silently treating tens of thousands of cells as tens of thousands of patients.
    """
    a = adata.copy()
    obs = a.obs
    configured = str(sample_col).strip() if sample_col is not None and str(sample_col).strip() else None

    # Pick a real repeated sample/patient column. The configured sample_col has
    # priority, but only if it does not look like cell-level barcodes.
    chosen_col, pdiag = choose_patient_column(obs, configured=configured, out_patient_col=out_patient_col)
    existing_ok = (
        out_patient_col in obs.columns
        and obs[out_patient_col].notna().any()
        and is_usable_patient_series(obs[out_patient_col], colname=out_patient_col)
    )

    if chosen_col is not None:
        vals = obs[chosen_col].astype(str)
        # If the chosen column is the configured sample column, derive patient IDs
        # using the cohort regex (AML timepoints: AML556-D31 -> AML556). For solid
        # tumours the default regex is harmless because sample labels usually lack
        # a -D<number> suffix.
        if configured and str(chosen_col).lower() == configured.lower():
            vals = vals.str.replace(patient_strip_regex, "", regex=True)
        # Prefer the configured sample column or any valid replacement over a bad
        # existing patient column. Preserve a valid existing patient only when no
        # better configured column was supplied.
        if (configured and str(chosen_col).lower() == configured.lower()) or not existing_ok or chosen_col != out_patient_col:
            if out_patient_col in obs.columns and not existing_ok and chosen_col != out_patient_col:
                bad = patient_series_diagnostics(obs[out_patient_col], colname=out_patient_col)
                print(f"[data] replacing invalid patient column '{out_patient_col}' "
                      f"(n_unique={bad['n_unique']}, median_cells/id={bad['median_cells_per_id']:.1f}, "
                      f"barcode_like={bad['barcode_like_frac']:.2f}) with obs['{chosen_col}'].")
            elif chosen_col != out_patient_col:
                print(f"[data] patient metadata inferred from obs['{chosen_col}'].")
            obs[out_patient_col] = vals.values
    else:
        # No valid metadata column. If an existing patient column is barcode-like,
        # null it out so downstream patient-level analyses do not create nonsense
        # 1-cell "patients".
        if out_patient_col in obs.columns and obs[out_patient_col].notna().any() and not existing_ok:
            bad = patient_series_diagnostics(obs[out_patient_col], colname=out_patient_col)
            print(f"[data] WARNING: obs['{out_patient_col}'] looks like cell barcodes "
                  f"(n_unique={bad['n_unique']}, median_cells/id={bad['median_cells_per_id']:.1f}); "
                  "clearing it for patient-level analyses. Provide/repair a sample column instead.")
            obs[out_patient_col] = pd.NA

    # Cautious final fallback: common scRNA obs_names are sometimes sample-prefixed.
    if (out_patient_col not in obs.columns or not obs[out_patient_col].notna().any()):
        idx = pd.Series(obs.index.astype(str), index=obs.index)
        prefix = idx.str.extract(r"^([^:_\-]+)[_: -]")[0]
        if prefix.notna().any() and prefix.nunique(dropna=True) > 1 and is_usable_patient_series(prefix, colname="obs_name_prefix"):
            obs[out_patient_col] = prefix.values
            print("[data] patient metadata inferred from obs-name prefixes.")

    # Timepoint detection from the configured sample column when available.
    if configured and configured in obs.columns:
        samp = obs[configured].astype(str)
        tp = samp.str.extract(timepoint_regex)[0]
        if tp.notna().any():
            obs[out_timepoint_col] = tp

    # Optional generic timepoint detection for new longitudinal datasets.
    if out_timepoint_col not in obs.columns or obs[out_timepoint_col].isna().all():
        for c in ["timepoint", "Timepoint", "time_point", "visit", "Visit", "day", "Day"]:
            if c in obs.columns and obs[c].notna().any():
                obs[out_timepoint_col] = obs[c].astype(str)
                break

    if out_patient_col in obs.columns and obs[out_patient_col].notna().any():
        n_pat = obs[out_patient_col].dropna().astype(str).nunique()
        msg = f"[data] patient column '{out_patient_col}': {n_pat} unique values"
        if out_timepoint_col in obs.columns:
            n_tp = obs[out_timepoint_col].dropna().astype(str).nunique()
            msg += f", {n_tp} timepoints"
        if not is_usable_patient_series(obs[out_patient_col], colname=out_patient_col):
            msg += " (WARNING: still looks too high-cardinality for patient-level plots)"
        print(msg + ".")
    else:
        print("[data] WARNING: could not infer a usable patient column for this cohort; patient-level plots will be skipped.")
    return a


def _patient_key(x) -> str:
    """Loose patient/sample key for cross-file matching."""
    return re.sub(r"[^A-Za-z0-9]+", "", str(x).strip()).upper()


# sample-type / timepoint suffix tokens that denote a SAMPLE, not patient identity.
_SAMPLE_SUFFIX = {"N", "T", "P", "M", "R", "B", "BM", "NORMAL", "TUMOR", "TUMOUR",
                  "PRIMARY", "MET", "METASTASIS", "METASTATIC", "RELAPSE", "REL",
                  "BLOOD", "PBMC", "DIAGNOSIS", "DX"}
_SUFFIX_PAT = re.compile(r"^(D|T|P|R|TP|W|C)\d+$", re.I)  # timepoint-like: D0, T1, P2, ...


def _strip_sample_suffix(x) -> str:
    """Drop a SINGLE trailing sample/timepoint token that follows a separator, so
    SMC01-T / SMC01-N -> SMC01 and AML329-D0 -> AML329. Conservative: only acts on
    a separator-delimited trailing token from a known set (never strips letters that
    are fused to the ID, e.g. CID4530N is left untouched)."""
    s = str(x).strip()
    parts = re.split(r"[\-_.\s]+", s)
    if len(parts) >= 2:
        last = parts[-1].upper()
        if last in _SAMPLE_SUFFIX or _SUFFIX_PAT.match(last):
            return "-".join(parts[:-1])
    return s


def _base_key(x) -> str:
    return _patient_key(_strip_sample_suffix(x))


def harmonize_patient_status(patient_status: pd.DataFrame | None, obs_patients) -> pd.DataFrame | None:
    """Map patient_status['patient'] onto the actual obs patient labels when possible.

    Public metadata often differs only by punctuation/case (``CRC-01`` vs ``CRC01``)
    or by a sample/timepoint suffix (``SMC01-T``/``SMC01-N`` vs ``SMC01``;
    ``AML329-D0`` vs ``AML329``). This maps via three conservative tiers — exact,
    punctuation/case-normalized, then sample-suffix-stripped — accepting only
    UNAMBIGUOUS matches (exactly one obs patient), and logs how many mapped at each
    tier plus a few that did not, so a naming mismatch is visible rather than silent.
    The original value is preserved in ``patient_original``.
    """
    if patient_status is None or patient_status.empty or "patient" not in patient_status.columns:
        return patient_status
    obs_vals = pd.Series(obs_patients).dropna().astype(str).unique().tolist()
    obs_set = set(obs_vals)
    out = patient_status.copy()
    out["patient_original"] = out["patient"].astype(str)

    def _build_lut(keyfn):
        d = {}
        for v in obs_vals:
            k = keyfn(v)
            if not k:
                continue
            if k not in d:
                d[k] = v
            elif d[k] != v:
                d[k] = None  # ambiguous -> do not map
        return d
    norm_lut = _build_lut(_patient_key)
    base_lut = _build_lut(_base_key)

    tier_counts = {"exact": 0, "normalized": 0, "suffix_stripped": 0, "unmatched": 0}
    mapped_vals, unmatched = [], []
    for orig in out["patient_original"]:
        if orig in obs_set:
            mapped_vals.append(orig); tier_counts["exact"] += 1; continue
        nk = _patient_key(orig)
        if nk in norm_lut and norm_lut[nk]:
            mapped_vals.append(norm_lut[nk]); tier_counts["normalized"] += 1; continue
        bk = _base_key(orig)
        if bk in base_lut and base_lut[bk]:
            mapped_vals.append(base_lut[bk]); tier_counts["suffix_stripped"] += 1; continue
        mapped_vals.append(orig); tier_counts["unmatched"] += 1
        if orig not in unmatched:
            unmatched.append(orig)
    out["patient"] = mapped_vals
    n_map = tier_counts["exact"] + tier_counts["normalized"] + tier_counts["suffix_stripped"]
    print(f"[data] patient_status match: {n_map}/{len(out)} rows mapped to obs patients "
          f"(exact={tier_counts['exact']}, normalized={tier_counts['normalized']}, "
          f"suffix-stripped={tier_counts['suffix_stripped']}, unmatched={tier_counts['unmatched']}).")
    if unmatched:
        print(f"[data]   unmatched patient IDs (first 6): {unmatched[:6]} "
              f"— if these are real scRNA patients, align the patient_status 'patient' column.")
    return out


def mark_reference_cells(
    adata: "ad.AnnData",
    celltype_col: str = "CellType",
    reference_celltypes: tuple[str, ...] = ("T", "B", "NK"),
    out_col: str = "is_reference",
    sample_col: str = "Sample",
    reference_samples=None,
) -> "ad.AnnData":
    """Flag non-malignant reference cells (the healthy/normal baseline for
    distribution-shift and CNV-reference logic).

    Matching is case-insensitive and exact on the cell-type label. If
    ``reference_samples`` is given, a cell is reference only if it ALSO comes from one
    of those samples — use this to restrict the healthy baseline to clean unsorted
    normal donors (e.g. BM1-BM4) and exclude sorted-progenitor controls (BM5*) that
    would otherwise inflate the reference with blast-like progenitors and drive
    myeloid-driver false positives on the reference.
    """
    adata = adata.copy()
    if celltype_col not in adata.obs.columns:
        print(f"[data] WARNING: celltype_col '{celltype_col}' absent; "
              f"is_reference set all-False.")
        adata.obs[out_col] = False
        return adata
    ct = adata.obs[celltype_col].astype(str)
    ref_lower = {r.lower() for r in reference_celltypes}
    flag = ct.str.lower().isin(ref_lower).values
    if reference_samples and sample_col in adata.obs.columns:
        in_ref_samp = adata.obs[sample_col].astype(str).isin(
            [str(s) for s in reference_samples]).values
        flag = flag & in_ref_samp
        print(f"[data] reference restricted to samples {list(reference_samples)}.")
    adata.obs[out_col] = flag
    print(f"[data] reference cells: {int(flag.sum())}/{adata.n_obs} "
          f"flagged from {reference_celltypes}.")
    return adata


def validate_patient_status_for_obs(
    patient_status: pd.DataFrame | None,
    obs_patients,
    cohort_name: str = "cohort",
    min_overlap_patients: int = 1,
) -> pd.DataFrame | None:
    """Return patient_status only if it overlaps the scRNA patient labels.

    This prevents accidentally using TCGA/reviewer/bulk mutation matrices for
    single-cell UMAP overlays when those sample IDs come from a different cohort.
    A table with zero overlapping patient/sample IDs is treated as unusable and
    downstream plots fall back to prediction-only or unknown-genotype panels.
    """
    if patient_status is None or patient_status.empty or "patient" not in patient_status.columns:
        return patient_status
    obs_vals = pd.Series(obs_patients).dropna().astype(str).unique().tolist()
    if not obs_vals:
        print(f"[data] WARNING {cohort_name}: no obs patients available for patient_status validation; skipping patient_status.")
        return None
    obs_set = set(obs_vals)
    status_patients = set(patient_status["patient"].dropna().astype(str).unique().tolist())
    overlap = obs_set & status_patients
    if len(overlap) < int(min_overlap_patients):
        print(f"[data] WARNING {cohort_name}: patient_status has {len(overlap)}/{len(obs_set)} overlapping scRNA patients; "
              "skipping it because it is probably from a different/bulk cohort.")
        return None
    print(f"[data] {cohort_name}: patient_status overlaps {len(overlap)}/{len(obs_set)} scRNA patients.")
    return patient_status

def load_patient_status(
    csv_path: str | Path,
    patient_col: str = "patient",
    mutation_col: str = "mutation",
    label_col: str = "label",
) -> pd.DataFrame:
    """Load a tidy patient-level mutation-status table.

    Expected columns: patient, mutation, label (label coerced to 0/1).
    Returns a tidy DataFrame; downstream code pivots as needed.
    """
    df = pd.read_csv(csv_path)
    df[label_col] = _coerce_binary(df[label_col])
    return df.rename(columns={patient_col: "patient",
                              mutation_col: "mutation",
                              label_col: "label"})[["patient", "mutation", "label"]]
```
