## Supplemental_Code for "scOPE identifies which driver-associated expression programs transfer from bulk tumors to single cells": identifiability.py

```
"""
scope_analysis.identifiability
==============================
The decisive analysis for the paper: is a scOPE per-driver score detecting the
MUTATION, or the CELL-STATE that the mutation rides on?

A classifier trained on bulk "mutant vs WT" learns the transcriptional correlates of
the mutation, which include real downstream programs but also lineage / blast-fraction /
cell-of-origin. So a per-cell score that lights up on mutant cells could be reading the
genomic event or just "this is a malignant myeloid blast". This module quantifies which,
with three NESTED contrasts that progressively remove confounders:

  1. cross_lineage      Mut malignant cells   vs  normal/reference cells.
                        High AUROC here is necessary but NOT sufficient — a pure
                        malignancy/lineage detector also wins this.
  2. within_malignant   among MALIGNANT cells only:  Mut vs WT.
                        Removes the "is it a blast" confound.
  3. within_lineage     among malignant cells of the SAME coarse lineage:  Mut vs WT.
                        Removes malignancy AND lineage. If the score still separates
                        here, the signal is mutation-specific. If it collapses to ~0.5
                        while (1) stayed high, the score is cell-state-confounded.

Statistics are PATIENT-LEVEL, not pooled across cells (pooling thousands of cells
pseudoreplicates and inflates significance). For each contrast we compute a per-patient
effect size (Cliff's delta, Mut vs WT within that patient ± lineage) and test across
patients with a Wilcoxon signed-rank test — the patient is the unit of inference.

Within the genotyped subset (confirmed Mut vs confirmed WT) AUROC/AUPRC are well defined.
The separate precision@k / hypergeometric test treats the data as positive-unlabeled
(confirmed-Mut = positives, everything else = unlabeled) and never assumes a confirmed
negative — so no accuracy/specificity is ever computed off "no-call" cells.

Clonal architecture is reported per driver because it changes interpretation: a TRUNK
mutation (carried by ~all blasts) makes within-malignant contrast uninformative — there
are essentially no WT malignant cells to compare — so a trunk gene validates the
malignant compartment, not per-cell mutation calling. SUBCLONAL genes are where the
identifiability test has teeth.

Call hierarchy
--------------
    drivers/run_all -> identifiability_report -> {clonality, nested_contrasts,
                                                  pu_enrichment} -> table + figure
"""
from __future__ import annotations

import numpy as np
import pandas as pd

from .config import AnalysisConfig
from .stats import auroc_safe, auprc_safe, bh_fdr

# Clonal-hematopoiesis / commonly-ancestral founders: flagged because they can appear in
# transcriptionally normal HSPCs, so a "mutant" cell is not necessarily malignant.
_CH_FOUNDERS = {"DNMT3A", "TET2", "ASXL1", "JAK2", "TP53", "SF3B1", "SRSF2", "IDH1", "IDH2"}

# minimum sample sizes below which we decline to render a verdict
_MIN_CELLS = 10
_MIN_PATIENTS = 4


def _cliffs(x: np.ndarray, y: np.ndarray) -> float:
    """Cliff's delta in [-1,1]: P(x>y) - P(x<y). Positive => x (Mut) scores higher."""
    x = np.asarray(x, float); y = np.asarray(y, float)
    x = x[np.isfinite(x)]; y = y[np.isfinite(y)]
    if len(x) == 0 or len(y) == 0:
        return np.nan
    # mid-rank formulation, exact with ties, O(n log n)
    allv = np.concatenate([x, y])
    order = allv.argsort()
    ranks = np.empty_like(order, float)
    ranks[order] = np.arange(1, len(allv) + 1)
    # average ranks for ties
    _, inv, cnt = np.unique(allv, return_inverse=True, return_counts=True)
    csum = np.cumsum(cnt)
    start = csum - cnt
    avg = (start + csum + 1) / 2.0
    ranks = avg[inv]
    rx = ranks[:len(x)].sum()
    u = rx - len(x) * (len(x) + 1) / 2.0
    return float(2.0 * u / (len(x) * len(y)) - 1.0)


def classify_clonality(mut_frac_per_patient: list[float], driver: str) -> dict:
    """From the Mut fraction among genotyped MALIGNANT cells in each mutant patient,
    classify clonal architecture. Trunk = mutation in ~all malignant cells."""
    fr = np.array([f for f in mut_frac_per_patient if np.isfinite(f)], float)
    med = float(np.median(fr)) if fr.size else np.nan
    if fr.size == 0:
        cls = "unknown"
    elif med >= 0.80:
        cls = "trunk"
    elif med >= 0.10:
        cls = "subclonal"
    else:
        cls = "rare"
    return {"clonality_class": cls, "mut_frac_malignant_median": med,
            "n_mut_patients": int(fr.size), "ch_founder": driver.upper() in _CH_FOUNDERS}


def _per_patient_effects(df: pd.DataFrame, patient_col: str) -> tuple[np.ndarray, int]:
    """Per-patient Cliff's delta of score between Mut(1) and WT(0) within each group of
    df (already restricted to a contrast). Returns (deltas, n_patients_usable)."""
    deltas = []
    for _, g in df.groupby(patient_col, observed=True):
        xm = g.loc[g["geno"] == 1, "score"].values
        xw = g.loc[g["geno"] == 0, "score"].values
        if len(xm) >= 3 and len(xw) >= 3:
            d = _cliffs(xm, xw)
            if np.isfinite(d):
                deltas.append(d)
    return np.array(deltas, float), len(deltas)


def _patient_level_test(deltas: np.ndarray) -> tuple[float, float]:
    """Wilcoxon signed-rank that per-patient Mut-vs-WT deltas are > 0 (one-sided).
    Returns (median_delta, p). Patient is the unit of inference."""
    if deltas.size < _MIN_PATIENTS:
        return (float(np.median(deltas)) if deltas.size else np.nan), np.nan
    from scipy.stats import wilcoxon
    try:
        # one-sided: deltas greater than 0
        _, p = wilcoxon(deltas, alternative="greater", zero_method="wilcox")
    except Exception:
        p = np.nan
    return float(np.median(deltas)), float(p)


def _pooled_auroc(pos: np.ndarray, neg: np.ndarray) -> float:
    pos = pos[np.isfinite(pos)]; neg = neg[np.isfinite(neg)]
    if len(pos) < 3 or len(neg) < 3:
        return np.nan
    y = np.r_[np.ones(len(pos)), np.zeros(len(neg))]
    s = np.r_[pos, neg]
    return auroc_safe(y, s)


def pu_enrichment(score: np.ndarray, is_confirmed_mut: np.ndarray) -> dict:
    """Positive-unlabeled spike-in: do the top-k scored cells over-represent confirmed
    mutant cells vs base rate? k = number of confirmed mutants. Reports precision@k,
    fold-enrichment over base rate, and a hypergeometric upper-tail p. Never assumes a
    confirmed negative."""
    score = np.asarray(score, float)
    pos = np.asarray(is_confirmed_mut, bool)
    ok = np.isfinite(score)
    score, pos = score[ok], pos[ok]
    n = score.size
    k = int(pos.sum())
    if n == 0 or k == 0 or k == n:
        return {"precision_at_k": np.nan, "fold_enrichment": np.nan, "hypergeom_p": np.nan, "k": k, "n": n}
    topk = np.argsort(-score)[:k]
    hits = int(pos[topk].sum())
    base = k / n
    prec = hits / k
    from scipy.stats import hypergeom
    # P(X >= hits) drawing k from n with k successes
    p = float(hypergeom.sf(hits - 1, n, k, k))
    return {"precision_at_k": prec, "fold_enrichment": (prec / base if base > 0 else np.nan),
            "hypergeom_p": p, "k": k, "n": n}


def identifiability_report(
    adata,
    cfg: AnalysisConfig,
    drivers: list[str] | None = None,
    score_prefix: str | None = None,
    patient_col: str = "patient",
    malignant_col: str = "scope_malignant",
    lineage_col: str = "celltype_coarse",
    reference_col: str = "is_reference",
) -> pd.DataFrame:
    """One row per driver with a per-cell genotype: clonality, the three nested-contrast
    AUROCs, the patient-level within-lineage test, the PU enrichment, and a conservative
    verdict. Requires malignant_col, lineage_col, and per-cell genotype to be present."""
    from .singlecell import find_percell_truth_columns, _binarize_genotype, scored_drivers
    drivers = drivers or scored_drivers(adata)
    score_prefix = score_prefix or getattr(cfg, "raw_score_prefix", "mutation_prob")
    truth = find_percell_truth_columns(adata, drivers)
    obs = adata.obs
    if not truth:
        print("[identifiability] no per-cell genotype columns; skipping.")
        return pd.DataFrame()
    if malignant_col not in obs.columns:
        print(f"[identifiability] '{malignant_col}' missing; run derive_malignant_compartment first.")
        return pd.DataFrame()
    has_line = lineage_col in obs.columns
    has_ref = reference_col in obs.columns
    pat = obs[patient_col].astype(str).values if patient_col in obs.columns else np.array(["p0"] * adata.n_obs)
    malig = obs[malignant_col].astype(bool).values
    ref = obs[reference_col].astype(bool).values if has_ref else ~malig

    rows = []
    for d, tcol in truth.items():
        scol = f"{score_prefix}_{d}"
        if scol not in obs.columns:
            continue
        score = obs[scol].astype(float).values
        geno = _binarize_genotype(obs[tcol]).values.astype(float)
        is_mut = geno == 1
        is_wt = geno == 0
        n_mut, n_wt = int(np.nansum(is_mut)), int(np.nansum(is_wt))

        # clonality from Mut fraction among genotyped malignant cells, per mutant patient
        fr = []
        for p in np.unique(pat):
            pm = (pat == p) & malig & np.isfinite(geno)
            if pm.sum() >= 5 and np.nansum(geno[pm] == 1) > 0:
                fr.append(float(np.nanmean(geno[pm] == 1)))
        clон = classify_clonality(fr, d)

        # contrast 1: Mut-malignant vs reference/normal
        auroc_cross = _pooled_auroc(score[is_mut & malig], score[ref])
        # contrast 2: within malignant, Mut vs WT
        mm, mw = is_mut & malig, is_wt & malig
        auroc_wm = _pooled_auroc(score[mm], score[mw])
        # contrast 3: within malignant AND within lineage, Mut vs WT (pooled descriptive)
        if has_line:
            line = obs[lineage_col].astype(str).values
            pos_l, neg_l = [], []
            for L in np.unique(line):
                sel = malig & (line == L)
                if (is_mut & sel).sum() >= 3 and (is_wt & sel).sum() >= 3:
                    pos_l.append(score[is_mut & sel]); neg_l.append(score[is_wt & sel])
            auroc_wl = (_pooled_auroc(np.concatenate(pos_l), np.concatenate(neg_l))
                        if pos_l else np.nan)
        else:
            auroc_wl = np.nan

        # patient-level test on the within-lineage contrast (the decisive inference)
        if has_line:
            dfc = pd.DataFrame({
                patient_col: pat, "score": score, "geno": geno,
                "_pl": [f"{a}|{b}" for a, b in zip(pat, obs[lineage_col].astype(str).values)],
            })[malig & np.isfinite(geno)]
            # per (patient,lineage) cell groups, then aggregate deltas
            deltas = []
            for _, g in dfc.groupby("_pl", observed=True):
                xm = g.loc[g["geno"] == 1, "score"].values
                xw = g.loc[g["geno"] == 0, "score"].values
                if len(xm) >= 3 and len(xw) >= 3:
                    dd = _cliffs(xm, xw)
                    if np.isfinite(dd):
                        deltas.append(dd)
            deltas = np.array(deltas, float)
        else:
            deltas = np.array([], float)
        med_delta, wilcox_p = _patient_level_test(deltas)

        # PU enrichment among malignant cells (positives = confirmed Mut)
        pu = pu_enrichment(score[malig], is_mut[malig])

        # conservative verdict
        verdict = _verdict(clон["clonality_class"], n_mut, n_wt, deltas.size, auroc_wl, med_delta, wilcox_p)

        rows.append({
            "driver": d, "genotype_col": tcol,
            **clон,
            "n_mut_cells": n_mut, "n_wt_cells": n_wt,
            "auroc_cross_lineage": auroc_cross,
            "auroc_within_malignant": auroc_wm,
            "auroc_within_lineage": auroc_wl,
            "within_lineage_delta_median": med_delta,
            "within_lineage_wilcoxon_p": wilcox_p,
            "n_patient_lineage_groups": int(deltas.size),
            "pu_precision_at_k": pu["precision_at_k"],
            "pu_fold_enrichment": pu["fold_enrichment"],
            "pu_hypergeom_p": pu["hypergeom_p"],
            "identifiability_verdict": verdict,
        })

    out = pd.DataFrame(rows)
    if out.empty:
        return out
    # BH-FDR across drivers on the decisive patient-level test
    p = out["within_lineage_wilcoxon_p"].values
    finite = np.isfinite(p)
    q = np.full(len(p), np.nan)
    if finite.any():
        q[finite] = bh_fdr(p[finite])
    out["within_lineage_fdr_q"] = q
    order = {"mutation-specific": 0, "cell-state-confounded": 1, "trunk: malignancy-proxy": 2,
             "inconclusive": 3, "underpowered": 4}
    out = out.sort_values(by=["identifiability_verdict", "within_lineage_delta_median"],
                          key=lambda s: s.map(order) if s.name == "identifiability_verdict" else -s,
                          ).reset_index(drop=True)
    return out


def _verdict(clonality, n_mut, n_wt, n_groups, auroc_wl, med_delta, wilcox_p) -> str:
    if clonality == "trunk":
        return "trunk: malignancy-proxy"
    if n_mut < _MIN_CELLS or n_wt < _MIN_CELLS or n_groups < _MIN_PATIENTS:
        return "underpowered"
    if np.isfinite(wilcox_p) and wilcox_p < 0.05 and np.isfinite(med_delta) and med_delta > 0.1:
        return "mutation-specific"
    if np.isfinite(auroc_wl) and auroc_wl < 0.58 and np.isfinite(med_delta) and abs(med_delta) <= 0.1:
        return "cell-state-confounded"
    return "inconclusive"


def plot_identifiability(report: pd.DataFrame, outpath, cfg=None):
    """Forest-style panel: per driver, the three nested-contrast AUROCs side by side so
    the drop from cross-lineage to within-lineage (the confounding gap) is obvious.
    Verdict shown as a colour tag."""
    import matplotlib
    matplotlib.use("Agg")
    import matplotlib.pyplot as plt
    from pathlib import Path
    if report is None or report.empty:
        return None
    from .plots import FIGURE_DPI
    df = report.copy()
    n = len(df)
    vcol = {"mutation-specific": "#1b9e77", "cell-state-confounded": "#d73027",
            "trunk: malignancy-proxy": "#7570b3", "inconclusive": "#999999",
            "underpowered": "#cccccc"}
    fig, ax = plt.subplots(figsize=(7.5, 0.55 * n + 1.6))
    ys = np.arange(n)[::-1]
    marks = [("auroc_cross_lineage", "o", "Mut-malig vs normal"),
             ("auroc_within_malignant", "s", "within malignant"),
             ("auroc_within_lineage", "D", "within lineage")]
    shades = ["#bdbdbd", "#6baed6", "#08519c"]
    for (col, mk, lab), sh in zip(marks, shades):
        ax.scatter(df[col], ys, marker=mk, s=46, color=sh, edgecolor="k",
                   linewidth=0.4, label=lab, zorder=3)
    for y, (_, r) in zip(ys, df.iterrows()):
        a, b = r["auroc_cross_lineage"], r["auroc_within_lineage"]
        if np.isfinite(a) and np.isfinite(b):
            ax.plot([min(a, b), max(a, b)], [y, y], color="#cccccc", lw=2, zorder=1)
        ax.add_patch(plt.Rectangle((1.02, y - 0.35), 0.04, 0.7,
                                   color=vcol.get(r["identifiability_verdict"], "#999"),
                                   transform=ax.get_yaxis_transform(), clip_on=False))
    ax.axvline(0.5, ls="--", color="grey", lw=1, label="chance (0.5)")
    ax.set_yticks(ys)
    ax.set_yticklabels([f"{r.driver}  [{r.clonality_class}]" for _, r in df.iterrows()], fontsize=8)
    ax.set_xlim(0.3, 1.06); ax.set_xlabel("apparent AUROC (Mut vs comparator)")
    ax.set_title("Identifiability: does the score survive removing malignancy & lineage?\n"
                 "(collapse toward 0.5 at 'within lineage' = cell-state-confounded)", fontsize=10)
    ax.legend(fontsize=7, loc="lower left", framealpha=0.9)
    fig.tight_layout()
    out = Path(outpath); fig.savefig(out, dpi=FIGURE_DPI, bbox_inches="tight"); plt.close(fig)
    return out
```
