## Supplemental_Code for "scOPE identifies which driver-associated expression programs transfer from bulk tumors to single cells": interpret.py

```
"""
scope_analysis.interpret
=========================
STAGE 1b: "Latent-component interpretation" (Methods 5.x; supports Fig. 2 gene
programs and the per-driver mechanistic read-out).

Thin, honest wrapper around scope-bio's ``SVDEvaluator``. It does NOT re-implement
the SVD interpretation — it drives the real class and adds (a) a multi-driver loop,
(b) a compact "leading gene programme" extraction for the manuscript text, and
(c) a guard that routes you correctly when the decomposition is not SVD (the
evaluator is SVD-only by construction).

This is what produced figures like your "Logistic regression coefficients - NPM1"
and "Aggregated gene scores from SVD + logistic model": SVDEvaluator's
``component_importance``, ``top_genes_per_component`` and ``gene_program_table``.

IMPORTANT (verified against the installed scope-bio 0.2.0):
  - SVDEvaluator requires ``bulk_pipeline.decomposition == 'svd'``.
  - It reads coefficients from a logistic/SVM classifier; for tree/boosting
    classifiers it raises and tells you to use SHAP instead. So run the
    interpretation pass with classifier='logistic' even if an ablation uses GBM.
  - Classifier-weighted importance is |coef| * sigma. (Note: with the default
    scale_by_singular_values=True the latent coordinates carry sigma^2 while this
    importance uses sigma^1 — see HONEST_NOTES in the blueprint.)

Call hierarchy
--------------
    drivers/run_interpret -> interpret_drivers
                              -> scope.evaluation.SVDEvaluator.run_all / individual plots
                              -> leading_gene_programs
"""
from __future__ import annotations

from pathlib import Path

import numpy as np
import pandas as pd

from scope.evaluation import SVDEvaluator

from .config import AnalysisConfig


def _first_attr(obj, *names):
    """Return the first existing non-None attribute from obj."""
    for name in names:
        if hasattr(obj, name):
            val = getattr(obj, name)
            if val is not None:
                return val
    return None


def plot_weighted_scree_improved(ev, outdir, top_k: int | None = None) -> Path:
    """Replacement for scope-bio's weighted scree.

    Fixes two practical issues in scope-bio's default plot:
      1. The colourbar is moved to the bottom so it cannot collide with the
         right-hand cumulative-variance axis.
      2. By default it shows ALL retained SVD components, not just the default
         first ~20 components used by SVDEvaluator.run_all().

    This function intentionally avoids relying on ev.k, because some installed
    scope-bio versions do not expose that attribute. The number of components is
    inferred from the singular-value vector and classifier-importance vector.
    """
    import matplotlib
    matplotlib.use("Agg")
    import matplotlib.pyplot as plt
    from matplotlib.cm import ScalarMappable
    from matplotlib.colors import Normalize
    from pathlib import Path as _P

    sigma = _first_attr(ev, "Sigma", "sigma", "singular_values", "singular_values_")
    importance = _first_attr(ev, "importance", "component_importance", "component_importance_", "weighted_importance")
    mutation = getattr(ev, "mutation", getattr(ev, "driver", "driver"))

    if sigma is None or importance is None:
        keys = sorted([k for k in vars(ev).keys() if not k.startswith("__")])
        raise AttributeError(
            "Could not locate singular values and component importance on SVDEvaluator. "
            f"Available attributes: {keys}"
        )

    sigma = np.asarray(sigma, dtype=float).ravel()
    importance = np.asarray(importance, dtype=float).ravel()
    k_total = int(min(len(sigma), len(importance)))
    if k_total <= 0:
        raise ValueError("No SVD components available for improved scree plot.")
    k = k_total if top_k is None else min(int(top_k), k_total)

    sigma2 = sigma[:k_total] ** 2
    denom = float(np.nansum(sigma2))
    if not np.isfinite(denom) or denom <= 0:
        raise ValueError("Singular values are empty/non-finite; cannot compute explained variance.")
    var_ratio_all = sigma2 / denom
    ev_pct = var_ratio_all[:k] * 100.0
    cum = np.cumsum(var_ratio_all)[:k] * 100.0
    imp = importance[:k]

    finite_imp = imp[np.isfinite(imp)]
    if finite_imp.size == 0:
        finite_imp = np.array([0.0, 1.0])
    vmin, vmax = float(np.nanmin(finite_imp)), float(np.nanmax(finite_imp))
    if not np.isfinite(vmin) or not np.isfinite(vmax) or vmin == vmax:
        vmax = vmin + 1.0

    norm = Normalize(vmin=vmin, vmax=vmax)
    cmap = plt.get_cmap("viridis")
    x = np.arange(1, k + 1)

    # Wider canvas for 100+ components; bottom room for colorbar.
    fig_w = max(10.5, min(28.0, 0.18 * k + 8.5))
    fig, ax1 = plt.subplots(figsize=(fig_w, 6.2), constrained_layout=False)
    fig.subplots_adjust(left=0.08, right=0.90, bottom=0.25, top=0.84)

    ax1.bar(x, ev_pct, color=cmap(norm(imp)), width=0.85, linewidth=0)
    ax1.set_xlabel("SVD component")
    ax1.set_ylabel("Explained variance (%)")
    step = max(1, int(np.ceil(k / 30)))
    ax1.set_xticks(x[::step])
    ax1.set_xlim(0.3, k + 0.7)

    ax2 = ax1.twinx()
    ax2.plot(x, cum, "k.--", lw=1.15, ms=3.2)
    ax2.set_ylabel("Cumulative explained variance (%)", labelpad=10)
    ax2.set_ylim(0, 101)

    sm = ScalarMappable(norm=norm, cmap=cmap)
    sm.set_array([])
    cb = fig.colorbar(sm, ax=[ax1, ax2], orientation="horizontal", fraction=0.055, pad=0.20, aspect=55)
    cb.set_label(f"Classifier importance  |coef|×σ   ({mutation})", fontsize=9)

    ax1.set_title(
        f"Classifier-weighted scree — {mutation} ({k} of {k_total} retained SVD components shown)\n"
        "bar height = explained variance; bar colour = logistic-regression importance",
        fontsize=11,
        pad=12,
    )

    outdir = _P(outdir)
    outdir.mkdir(parents=True, exist_ok=True)
    out = outdir / "weighted_scree.png"
    # Older hotfixes wrote a duplicate named weighted_scree_FIXED.png.
    # Do not create that duplicate anymore, and remove it if it is left over.
    old_duplicate = outdir / "weighted_scree_FIXED.png"
    try:
        old_duplicate.unlink(missing_ok=True)
    except TypeError:  # Python < 3.8 fallback
        if old_duplicate.exists():
            old_duplicate.unlink()
    fig.savefig(out, dpi=300, bbox_inches="tight")
    plt.close(fig)
    return out


def component_signature_scan(
    bulk_pipeline, adata_bulk, drivers: list[str], cfg: AnalysisConfig, cancer: str,
    top_components: int = 25, genes_per_component: int = 20,
) -> pd.DataFrame:
    """Scan MANY top components per driver for gene signatures (deeper than the
    default 10). Returns a tidy table [driver, component_rank, component, importance,
    sigma, abs_coef, direction, gene, loading] across the top-N importance-ranked
    components, and writes it to fig2_bulk/<cancer>/component_signature_scan.csv.
    Use it to mine recurrent programmes across drivers."""
    Z_bulk = bulk_pipeline.transform_bulk(adata_bulk).obsm[bulk_pipeline.obsm_key_]
    trained = set(bulk_pipeline.classifier_set_.classifiers_.keys())
    rows = []
    for d in drivers:
        if d not in trained:
            continue
        ev = SVDEvaluator(bulk_pipeline, Z_bulk, mutation=d)
        ranked = np.argsort(ev.importance)[::-1][:top_components]
        for rank, comp in enumerate(ranked, start=1):
            loadings = ev.V[:, comp]
            order = np.argsort(np.abs(loadings))[::-1][:genes_per_component]
            for gi in order:
                rows.append(dict(
                    driver=d, component_rank=rank, component=int(comp),
                    importance=float(ev.importance[comp]), sigma=float(ev.Sigma[comp]),
                    abs_coef=float(ev.coef_abs[comp]),
                    direction="up" if loadings[gi] > 0 else "down",
                    gene=ev.gene_names[gi], loading=float(loadings[gi]),
                ))
    out = pd.DataFrame(rows)
    if not out.empty:
        path = cfg.outdir("fig2_bulk", cancer) / "component_signature_scan.csv"
        out.to_csv(path, index=False)
        print(f"[interpret] {cancer}: scanned {top_components} components x "
              f"{out.driver.nunique()} drivers -> {path}")
    return out


def _label_vector_for_driver(labels: pd.DataFrame | None, driver: str, fallback_y) -> np.ndarray:
    """Return the 0/1 label vector used for rare-driver CV decisions."""
    if labels is not None:
        # Match robustly while preserving original column spelling.
        d_u = str(driver).strip().upper()
        for col in labels.columns:
            if str(col).strip().upper() == d_u:
                return pd.to_numeric(labels[col], errors="coerce").fillna(0).astype(int).to_numpy()
    return np.asarray(fallback_y, dtype=int).ravel()


def _effective_cv_for_binary_labels(y, requested_cv: int, driver: str, cancer: str) -> tuple[int, dict]:
    """Choose a safe StratifiedKFold n_splits for a binary driver.

    scope-bio's SVDEvaluator defaults to fixed 5-fold CV.  In exploratory
    all-label mode, rare drivers can have <5 positives; fixed 5-fold CV then
    creates one-class folds, returning NaN AUROCs that later crash matplotlib's
    histogram code.  This helper reduces CV to min(requested_cv, n_pos, n_neg),
    or returns 0 to skip CV-dependent plots if either class has <2 samples.
    """
    y = np.asarray(y, dtype=int).ravel()
    n = int(y.size)
    n_pos = int(np.sum(y == 1))
    n_neg = int(np.sum(y == 0))
    min_class = min(n_pos, n_neg)
    requested_cv = int(requested_cv or 5)
    if min_class < 2:
        reason = f"skip_cv_min_class<{2}"
        print(f"[interpret] {cancer}/{driver}: pos={n_pos}, neg={n_neg}; skipping CV-dependent plots ({reason}).")
        return 0, dict(n=n, n_pos=n_pos, n_neg=n_neg, requested_cv=requested_cv,
                       effective_cv=0, cv_reduced=False, cv_skipped=True, cv_reason=reason)
    effective_cv = max(2, min(requested_cv, min_class))
    reduced = effective_cv != requested_cv
    if reduced:
        print(f"[interpret] {cancer}/{driver}: pos={n_pos}, neg={n_neg}; using cv={effective_cv} instead of requested cv={requested_cv} for rare-driver-safe SVD evaluation.")
    return effective_cv, dict(n=n, n_pos=n_pos, n_neg=n_neg, requested_cv=requested_cv,
                              effective_cv=effective_cv, cv_reduced=bool(reduced),
                              cv_skipped=False, cv_reason="adaptive_cv" if reduced else "requested_cv_ok")


def _write_plot_skip_note(outdir: Path, plot_name: str, exc: Exception) -> None:
    """Write a tiny .txt marker when a scope-bio plot is skipped."""
    outdir.mkdir(parents=True, exist_ok=True)
    note = outdir / f"SKIPPED_{plot_name}.txt"
    note.write_text(f"Skipped {plot_name}\nReason: {type(exc).__name__}: {exc}\n")


def _safe_svd_plot_call(ev, outdir: Path, method_name: str, *args, audit: list[dict] | None = None, **kwargs) -> bool:
    """Call one SVDEvaluator plotting/export method without letting rare-label
    numerical issues kill the whole suite."""
    try:
        getattr(ev, method_name)(outdir, *args, **kwargs)
        if audit is not None:
            audit.append(dict(method=method_name, status="ok", reason=""))
        return True
    except Exception as e:
        print(f"[interpret] {getattr(ev, 'mutation', '?')}: {method_name} skipped/failed: {type(e).__name__}: {e}")
        try:
            _write_plot_skip_note(outdir, method_name, e)
        except Exception:
            pass
        if audit is not None:
            audit.append(dict(method=method_name, status="skipped_or_failed", reason=f"{type(e).__name__}: {e}"))
        return False


def _run_svd_evaluator_safe(
    ev,
    outdir: Path,
    cv: int,
    n_permutations: int,
    *,
    table_top_components: int = 25,
    table_genes_per_component: int = 50,
    plot_top_components: int = 6,
    plot_genes_per_component: int = 30,
    heatmap_top_components: int = 10,
    heatmap_genes_per_component: int = 15,
) -> pd.DataFrame:
    """Rare-label-safe replacement for SVDEvaluator.run_all.

    Non-CV plots are still attempted for every trained classifier.  CV-dependent
    plots are skipped if cv==0 and otherwise guarded individually, so an all-NaN
    permutation null or one-class fold cannot crash the run.
    """
    audit: list[dict] = []
    outdir = Path(outdir)

    # Non-CV plots/exports: these should usually be available even for rare labels.
    # Pass depth controls explicitly; scope-bio's defaults are small
    # (top_genes_per_component.png=20 genes, gene_program_table=30 genes).
    _safe_svd_plot_call(ev, outdir, "plot_weighted_scree", audit=audit)
    _safe_svd_plot_call(ev, outdir, "plot_component_importance", top_k=int(table_top_components), audit=audit)
    _safe_svd_plot_call(
        ev, outdir, "plot_gene_loading_heatmap",
        top_components=int(heatmap_top_components),
        top_genes_per_comp=int(heatmap_genes_per_component),
        audit=audit,
    )
    _safe_svd_plot_call(
        ev, outdir, "plot_top_genes_per_component",
        top_components=int(plot_top_components),
        top_genes=int(plot_genes_per_component),
        audit=audit,
    )
    _safe_svd_plot_call(ev, outdir, "plot_latent_scatter", audit=audit)
    _safe_svd_plot_call(ev, outdir, "plot_separation_violins", audit=audit)
    _safe_svd_plot_call(ev, outdir, "plot_component_label_correlation", audit=audit)

    # CV-dependent plots: only run when both classes can support at least 2 folds.
    if cv and int(cv) >= 2:
        _safe_svd_plot_call(ev, outdir, "plot_roc_ablation", cv=int(cv), audit=audit)
        _safe_svd_plot_call(ev, outdir, "plot_permutation_test", n_permutations=int(n_permutations), cv=int(cv), audit=audit)
        _safe_svd_plot_call(ev, outdir, "plot_calibration", cv=int(cv), audit=audit)
    else:
        for method in ["plot_roc_ablation", "plot_permutation_test", "plot_calibration"]:
            msg = RuntimeError("Skipped because effective_cv < 2 for this rare driver")
            print(f"[interpret] {getattr(ev, 'mutation', '?')}: {method} skipped: {msg}")
            _write_plot_skip_note(outdir, method, msg)
            audit.append(dict(method=method, status="skipped", reason=str(msg)))

    # Remaining optional plots.  SHAP/UMAP can legitimately be unavailable or fail
    # depending on optional deps, so keep these guarded too.
    for method in ["plot_gene_biplot", "plot_shap", "plot_umap", "plot_component_crosscorr"]:
        _safe_svd_plot_call(ev, outdir, method, audit=audit)
    _safe_svd_plot_call(
        ev, outdir, "export_gene_program_table",
        top_components=int(table_top_components),
        top_genes=int(table_genes_per_component),
        audit=audit,
    )

    audit_df = pd.DataFrame(audit)
    try:
        audit_df.to_csv(outdir / "svd_evaluator_plot_audit.csv", index=False)
    except Exception:
        pass
    return audit_df

def interpret_drivers(
    bulk_pipeline,
    adata_bulk,
    drivers: list[str],
    cfg: AnalysisConfig,
    cancer: str,
    full_suite: bool = True,
    labels: pd.DataFrame | None = None,
) -> dict[str, pd.DataFrame]:
    """Run SVDEvaluator for each driver and return a dict of gene-program tables.

    full_suite=True calls ``run_all`` (all 15 plots + CSV per driver). Set False
    for a fast pass that only exports the gene-program table and the three plots
    most used in the manuscript (importance, top genes, permutation null).
    """
    if bulk_pipeline.decomposition != "svd":
        raise ValueError(
            f"interpret_drivers needs an SVD pipeline; got "
            f"'{bulk_pipeline.decomposition}'. Re-fit with decomposition='svd' "
            f"for the interpretation pass (ablations can use other methods)."
        )
    Z_bulk = bulk_pipeline.transform_bulk(adata_bulk).obsm[bulk_pipeline.obsm_key_]
    trained = set(bulk_pipeline.classifier_set_.classifiers_.keys())
    out = {}
    cv_audit_rows = []
    for d in drivers:
        if d not in trained:
            print(f"[interpret] {cancer}/{d}: no trained classifier (too rare?), skipping.")
            continue
        ev = SVDEvaluator(bulk_pipeline, Z_bulk, mutation=d)
        outdir = cfg.outdir("fig2_bulk", cancer, f"svd_eval_{d}")
        y_for_cv = _label_vector_for_driver(labels, d, ev.y)
        effective_cv, cv_info = _effective_cv_for_binary_labels(y_for_cv, cfg.cv_folds, d, cancer)
        table_top_components = int(getattr(cfg, "interpret_top_components", 25))
        table_genes_per_component = int(getattr(cfg, "interpret_genes_per_component", 50))
        plot_top_components = int(getattr(cfg, "interpret_plot_top_components", 6))
        plot_genes_per_component = int(getattr(cfg, "interpret_plot_genes_per_component", 30))
        heatmap_top_components = int(getattr(cfg, "interpret_heatmap_top_components", 10))
        heatmap_genes_per_component = int(getattr(cfg, "interpret_heatmap_genes_per_component", 15))
        if full_suite:
            plot_audit = _run_svd_evaluator_safe(
                ev, outdir, cv=effective_cv, n_permutations=cfg.n_perm,
                table_top_components=table_top_components,
                table_genes_per_component=table_genes_per_component,
                plot_top_components=plot_top_components,
                plot_genes_per_component=plot_genes_per_component,
                heatmap_top_components=heatmap_top_components,
                heatmap_genes_per_component=heatmap_genes_per_component,
            )
        else:
            plot_audit_rows = []
            _safe_svd_plot_call(ev, outdir, "plot_component_importance", top_k=table_top_components, audit=plot_audit_rows)
            _safe_svd_plot_call(ev, outdir, "plot_top_genes_per_component", top_components=plot_top_components, top_genes=plot_genes_per_component, audit=plot_audit_rows)
            if effective_cv >= 2:
                _safe_svd_plot_call(ev, outdir, "plot_permutation_test", n_permutations=cfg.n_perm, cv=effective_cv, audit=plot_audit_rows)
            plot_audit = pd.DataFrame(plot_audit_rows)
        cv_record = {**cv_info, "driver": d, "cancer": cancer, "n_permutations": int(cfg.n_perm)}
        cv_audit_rows.append(cv_record)
        try:
            pd.DataFrame([cv_record]).to_csv(outdir / "svd_evaluator_cv_audit.csv", index=False)
        except Exception:
            pass
        # deeper gene-program export + replace the scree with the fixed-colourbar,
        # all-components version (overwrites scope-bio's overlapping one)
        try:
            ev.export_gene_program_table(
                outdir,
                top_components=table_top_components,
                top_genes=table_genes_per_component,
            )
        except Exception as e:
            print(f"[interpret] {cancer}/{d}: gene_program_table export skipped/failed: {type(e).__name__}: {e}")
        try:
            scree = plot_weighted_scree_improved(ev, outdir, top_k=None)
            print(f"[interpret] {cancer}/{d}: overwrote weighted_scree.png with improved all-component scree -> {scree}")
        except Exception as e:
            print(f"[interpret] {cancer}/{d}: improved scree FAILED; leaving scope-bio default plot in place. Reason: {e}")
        # read back the tidy gene-program table for the manuscript
        gp_path = outdir / "gene_program_table.csv"
        if gp_path.exists():
            out[d] = pd.read_csv(gp_path)
        print(f"[interpret] {cancer}/{d}: SVDEvaluator outputs -> {outdir}")
    if cv_audit_rows:
        try:
            audit_path = cfg.outdir("tables") / f"SVD_interpretation_cv_audit_{cancer}.csv"
            pd.DataFrame(cv_audit_rows).to_csv(audit_path, index=False)
            print(f"[interpret] {cancer}: rare-driver CV audit -> {audit_path}")
        except Exception as e:
            print(f"[interpret] {cancer}: failed to write rare-driver CV audit: {e}")
    return out


def leading_gene_programs(
    bulk_pipeline,
    adata_bulk,
    driver: str,
    top_components: int = 3,
    genes_per_component: int = 12,
) -> pd.DataFrame:
    """Compact, text-ready summary: the top mutation-associated components for a
    driver and their leading signed genes.

    Returns columns: [component_rank, component, importance, sigma, abs_coef,
    direction, gene, loading]. Use this to write sentences like "NPM1 is driven by
    component 4 (HOXB-AS3, HOXB9, MEIS1 ...)" directly from the numbers.
    """
    Z_bulk = bulk_pipeline.transform_bulk(adata_bulk).obsm[bulk_pipeline.obsm_key_]
    ev = SVDEvaluator(bulk_pipeline, Z_bulk, mutation=driver)
    ranked = np.argsort(ev.importance)[::-1][:top_components]
    rows = []
    for rank, comp in enumerate(ranked, start=1):
        loadings = ev.V[:, comp]
        order = np.argsort(np.abs(loadings))[::-1][:genes_per_component]
        for gi in order:
            rows.append(dict(
                driver=driver, component_rank=rank, component=int(comp),
                importance=float(ev.importance[comp]),
                sigma=float(ev.Sigma[comp]), abs_coef=float(ev.coef_abs[comp]),
                direction="up" if loadings[gi] > 0 else "down",
                gene=ev.gene_names[gi], loading=float(loadings[gi]),
            ))
    return pd.DataFrame(rows)
```
