## Supplemental_Code for "scOPE identifies which driver-associated expression programs transfer from bulk tumors to single cells": longitudinal.py

```
"""
scope_analysis.longitudinal
============================
THE TREATMENT-RESPONSE STORY (your buried gold).

Your van Galen cohort carries serial timepoints per patient (D0 at diagnosis,
then D15 / D20 / D31 / D37 ... on therapy). Your existing DNMT3A and NPM1 density
plots already show, qualitatively, that a patient's tumour score distribution
sits well above the healthy-BM reference at D0 and then *collapses back* toward
healthy as treatment clears the mutant clone. This module makes that quantitative
and turns it into a falsifiable transfer-learning claim:

    If scOPE scores reflect a real mutation programme (not batch / depth),
    then within a responding patient the per-cell score distribution should move
    monotonically toward the healthy reference over treatment, and the magnitude
    of that move should track the driver's clinical clearance.

It produces, per (patient, driver):
  * a fixed healthy reference distribution (normal cells, pooled once),
  * a distribution-shift statistic at EACH timepoint vs that same reference
    (Cliff's delta, KS, median diff, fraction above the healthy q-quantile),
  * the timepoint TRAJECTORY of those statistics (the thing that should decay),
  * a monotonic-trend test (Spearman of shift vs ordered timepoint), and
  * where per-cell genotype exists, the Mut/WT/unknown split per timepoint
    (your AML556-NRAS / AML328-TP53 violins).

This is a within-patient analysis, so it is immune to the cross-patient
confounding (subtype, composition, batch) that the chapter flags for
patient-level cohorts.

Call hierarchy
--------------
    drivers/run_longitudinal -> build_healthy_reference
                              -> longitudinal_trajectory   (Fig 3/4 trajectory + image 17-22 stats)
                              -> percell_timepoint_split    (image 12/16 violins)
                              -> trajectory_trend_test
"""
from __future__ import annotations

import numpy as np
import pandas as pd

from . import stats as st
from .config import AnalysisConfig
from .data import scored_drivers
from .singlecell import find_percell_truth_columns, _binarize_genotype


# ---------------------------------------------------------------------------
# Fixed healthy reference (pooled once, reused at every timepoint)
# ---------------------------------------------------------------------------
def build_healthy_reference(
    adata,
    driver: str,
    reference_mask_col: str = "is_reference",
    cfg=None,
    score_col: str | None = None,
) -> np.ndarray:
    """Pool scores from flagged normal/reference cells into ONE reference
    distribution for a driver.

    Using a single fixed reference for all timepoints is what makes the
    trajectory interpretable: every timepoint is measured against the same ruler.
    Flag reference cells first with data.mark_reference_cells.
    """
    score_col = score_col or (cfg.longitudinal_score_col(adata, driver) if cfg is not None else f"mutation_prob_{driver}")
    if score_col not in adata.obs.columns:
        raise KeyError(f"score column {score_col!r} missing for {driver}.")
    if reference_mask_col not in adata.obs.columns:
        raise KeyError(f"'{reference_mask_col}' missing; run data.mark_reference_cells.")
    ref = adata.obs.loc[adata.obs[reference_mask_col], score_col].astype(float).values
    ref = ref[~np.isnan(ref)]
    if len(ref) == 0:
        raise ValueError(f"No reference cells with scores for {driver}.")
    return ref


# ---------------------------------------------------------------------------
# Timepoint ordering (D0 < D15 < D31 ...), robust to arbitrary day numbers
# ---------------------------------------------------------------------------
def _order_timepoints(tps: list[str]) -> list[str]:
    """Sort timepoint labels like ['D31','D0','D15'] by their numeric day."""
    def day(t):
        m = "".join(ch for ch in str(t) if ch.isdigit())
        return int(m) if m else 10**9
    return sorted([t for t in tps if pd.notna(t)], key=day)


def _resolve_score_col(adata, cfg: AnalysisConfig | None, driver: str, score_mode: str = "configured") -> str:
    """Resolve score columns for longitudinal analyses.

    score_mode='configured' uses cfg.longitudinal_score_col (raw by default).
    score_mode='raw' forces mutation_prob_*.
    score_mode='cellstate_residual' uses mutation_resid_* if present.
    score_mode='cellstate_percentile' uses mutation_refpct_* if present.
    Missing guarded columns fall back to configured raw score.
    """
    mode = str(score_mode or "configured").lower()
    if mode in {"cellstate_residual", "residual", "guarded"}:
        col = f"mutation_resid_{driver}"
        if col in adata.obs.columns:
            return col
    if mode in {"cellstate_percentile", "percentile", "refpct"}:
        col = f"mutation_refpct_{driver}"
        if col in adata.obs.columns:
            return col
    if mode in {"raw", "raw_probability"}:
        return cfg.score_col(adata, driver, use_calibrated=False) if cfg is not None else f"mutation_prob_{driver}"
    return cfg.longitudinal_score_col(adata, driver) if cfg is not None else f"mutation_prob_{driver}"


def _score_label(cfg: AnalysisConfig | None, score_col: str, driver: str) -> tuple[str, str]:
    if cfg is not None:
        return cfg.score_kind(score_col), cfg.score_label_for_col(score_col, driver)
    return "score", score_col


# ---------------------------------------------------------------------------
# Core: per-patient distribution-shift trajectory over treatment
# ---------------------------------------------------------------------------
def longitudinal_trajectory(
    adata,
    cfg: AnalysisConfig,
    drivers: list[str] | None = None,
    patient_col: str = "patient",
    timepoint_col: str = "timepoint",
    reference_mask_col: str = "is_reference",
    malignant_only_col: str | None = None,
    score_mode: str = "configured",
    include_single_timepoint_patients: bool = False,
) -> pd.DataFrame:
    """For every (patient, driver, timepoint), quantify how far the tumour score
    distribution sits above the fixed healthy reference.

    malignant_only_col : optional bool obs column. If given, the trajectory is
        computed on malignant cells only (recommended when a malignant/normal
        call exists, e.g. from CNV), so the shift reflects the tumour compartment
        rather than changing normal-cell admixture across timepoints.

    Returns tidy rows:
      [patient, driver, timepoint, day, n_cells, median_diff, cliffs_delta,
       ks_statistic, ks_p, frac_above_ref_q, ref_threshold, n_reference]
    sorted by patient, driver, day. Plot cliffs_delta (or frac_above_ref_q)
    against day per patient to see the treatment-response curve.
    """
    drivers = drivers or scored_drivers(adata)
    obs = adata.obs
    rows = []
    has_tp = obs[timepoint_col].notna()
    patients = obs.loc[has_tp, patient_col].unique()

    for driver in drivers:
        score_col = _resolve_score_col(adata, cfg, driver, score_mode=score_mode)
        try:
            ref = build_healthy_reference(adata, driver, reference_mask_col, cfg=cfg, score_col=score_col)
        except ValueError as e:
            print(f"[longitudinal] {driver}: {e}; skipping driver.")
            continue
        for pat in patients:
            sel = (obs[patient_col] == pat) & has_tp
            if malignant_only_col and malignant_only_col in obs.columns:
                sel = sel & obs[malignant_only_col].astype(bool)
            tps = _order_timepoints(obs.loc[sel, timepoint_col].unique().tolist())
            if len(tps) < 2 and not include_single_timepoint_patients:
                continue  # default: need >=2 timepoints to call it a trajectory
            for tp in tps:
                cells = obs.loc[sel & (obs[timepoint_col] == tp), score_col].astype(float).values
                cells = cells[~np.isnan(cells)]
                if len(cells) < cfg.min_cells_per_patient:
                    continue
                sh = st.distribution_shift(cells, ref,
                                           reference_quantile=cfg.reference_quantile)
                score_kind, score_label = _score_label(cfg, score_col, driver)
                ref_median = float(np.nanmedian(ref)) if len(ref) else np.nan
                cell_median = float(np.nanmedian(cells)) if len(cells) else np.nan
                rows.append(dict(
                    patient=pat, driver=driver, timepoint=tp,
                    day=int("".join(c for c in tp if c.isdigit()) or 0),
                    score_col=score_col, score_kind=score_kind, score_label=score_label,
                    score_mode=score_mode, reference_mask_col=reference_mask_col,
                    n_cells=len(cells), median_score=cell_median, reference_median=ref_median,
                    reference_direction=("sample_above_reference" if cell_median > ref_median else
                                         "sample_below_reference" if cell_median < ref_median else "tied"),
                    n_reference=len(ref), median_diff=sh.median_diff,
                    cliffs_delta=sh.cliffs_delta, ks_statistic=sh.ks_statistic,
                    ks_p=sh.ks_p, frac_above_ref_q=sh.frac_above_ref_q,
                    ref_threshold=sh.ref_threshold,
                ))
    out = pd.DataFrame(rows)
    if not out.empty:
        out = out.sort_values(["patient", "driver", "day"]).reset_index(drop=True)
    return out


# ---------------------------------------------------------------------------
# Monotonic-trend test: does the shift decay over treatment?
# ---------------------------------------------------------------------------
def trajectory_trend_test(
    trajectory: pd.DataFrame,
    cfg: AnalysisConfig,
    shift_metric: str = "cliffs_delta",
) -> pd.DataFrame:
    """Per (patient, driver), Spearman-correlate the shift metric against day.

    A clearing/responding patient should show a NEGATIVE trend (shift shrinks as
    day increases). Returns rho, p, n_timepoints, first/last value and their
    delta. One global BH-FDR across all (patient, driver) trajectories.
    """
    from scipy.stats import spearmanr
    rows = []
    for (pat, driver), g in trajectory.groupby(["patient", "driver"]):
        g = g.sort_values("day")
        if g.shape[0] < 3:
            rho, p = np.nan, np.nan  # Spearman needs >=3 points to be meaningful
        else:
            rho, p = spearmanr(g["day"].values, g[shift_metric].values)
        rows.append(dict(
            patient=pat, driver=driver, n_timepoints=g.shape[0],
            shift_metric=shift_metric, spearman_rho=float(rho) if rho==rho else np.nan,
            spearman_p=float(p) if p==p else np.nan,
            first_value=float(g[shift_metric].iloc[0]),
            last_value=float(g[shift_metric].iloc[-1]),
            change=float(g[shift_metric].iloc[-1] - g[shift_metric].iloc[0]),
            direction="decreasing (response-consistent)" if g[shift_metric].iloc[-1] < g[shift_metric].iloc[0]
                       else "increasing/flat",
        ))
    out = pd.DataFrame(rows)
    if not out.empty:
        out["fdr_q"] = st.bh_fdr(out["spearman_p"].values)
    return out


# ---------------------------------------------------------------------------
# Per-cell genotype split by timepoint  -> your AML556-NRAS / AML328-TP53 violins
# ---------------------------------------------------------------------------
def percell_timepoint_split(
    adata,
    patient: str,
    driver: str,
    patient_col: str = "patient",
    timepoint_col: str = "timepoint",
    cfg=None,
    score_mode: str = "configured",
) -> pd.DataFrame:
    """Tidy per-cell scores for one patient/driver, split by timepoint and (if
    available) per-cell genotype label (Mut / WT / Unknown).

    Reproduces and generalises your "NRAS: AML556 - predictions by timepoint" and
    "TP53: AML328 - predictions by timepoint" panels. Returns columns:
    [timepoint, day, score, genotype_label]. If no per-cell genotype column is
    present, genotype_label is 'Unknown' for all cells (still a valid trajectory
    of the score distribution).
    """
    obs = adata.obs
    sub = obs[obs[patient_col] == patient].copy()
    if sub.empty:
        print(f"[longitudinal] patient {patient} not found.")
        return pd.DataFrame()
    score_col = _resolve_score_col(adata, cfg, driver, score_mode=score_mode)
    truth = find_percell_truth_columns(adata, [driver]).get(driver)
    geno = (_binarize_genotype(sub[truth]) if truth is not None
            else pd.Series(np.nan, index=sub.index))
    rows = []
    for idx in sub.index:
        tp = sub.loc[idx, timepoint_col]
        if pd.isna(tp):
            continue
        g = geno.loc[idx]
        label = "Mut" if g == 1 else ("WT" if g == 0 else "Unknown")
        rows.append(dict(timepoint=tp,
                         day=int("".join(c for c in str(tp) if c.isdigit()) or 0),
                         score=float(sub.loc[idx, score_col]),
                         genotype_label=label))
    out = pd.DataFrame(rows)
    if not out.empty:
        out = out.sort_values("day").reset_index(drop=True)
        if cfg is not None:
            out.attrs["score_col"] = score_col
            out.attrs["score_kind"] = cfg.score_kind(score_col)
            out.attrs["score_label"] = cfg.score_label_for_col(score_col, driver)
    return out


# ---------------------------------------------------------------------------
# Density-overlay table  -> your "Healthy (BM) vs AMLxxx.Dn" KDE figures
# ---------------------------------------------------------------------------
def density_overlay_table(
    adata,
    patient: str,
    driver: str,
    cfg: AnalysisConfig,
    patient_col: str = "patient",
    timepoint_col: str = "timepoint",
    reference_mask_col: str = "is_reference",
    score_mode: str = "configured",
) -> dict:
    """Assemble the arrays needed to draw the healthy-vs-tumour KDE overlays at
    each timepoint for one patient/driver, plus the shift stat annotating each.

    Returns {'reference': array, 'timepoints': {tp: {'scores': array,
    'shift': ShiftResult-as-dict}}}. plots.py consumes this directly.
    """
    score_col = _resolve_score_col(adata, cfg, driver, score_mode=score_mode)
    ref = build_healthy_reference(adata, driver, reference_mask_col, cfg=cfg, score_col=score_col)
    obs = adata.obs
    sub = obs[(obs[patient_col] == patient) & obs[timepoint_col].notna()]
    result = {"reference": ref, "driver": driver, "patient": patient,
              "score_col": score_col, "score_kind": cfg.score_kind(score_col),
              "score_label": cfg.score_label_for_col(score_col, driver),
              "timepoints": {}}
    for tp in _order_timepoints(sub[timepoint_col].unique().tolist()):
        s = sub.loc[sub[timepoint_col] == tp, score_col].astype(float).values
        s = s[~np.isnan(s)]
        if len(s) < cfg.min_cells_per_patient:
            continue
        sh = st.distribution_shift(s, ref, reference_quantile=cfg.reference_quantile)
        result["timepoints"][tp] = {"scores": s, "shift": sh.as_dict()}
    return result
```
