## Supplemental_Code for "scOPE identifies which driver-associated expression programs transfer from bulk tumors to single cells": pancancer.py

```
"""
scope_analysis.pancancer
=========================
PAN-CANCER GENERALIZATION (Results 2.5, Fig. 5 / Table 2) + the negative-control
machinery (Methods "Validation safeguards and negative controls").

The chapter is explicit that evidence quality differs by cancer, so this module
separates three tiers and applies the RIGHT test to each:

  TIER 1  per_cell      (AML)            -> singlecell.per_cell_validation
  TIER 2  patient_level (CRC/BRCA/SKCM)  -> patient-level score distribution vs
                                            normal cells + permutation null
  TIER 3  exploratory   (LUAD/GBM/PAAD)  -> score localisation in malignant vs
                                            normal compartments only (no genotype)

For cohorts WITHOUT per-cell genotype (your bulk + single-cell sets with only
patient-level or no mutation truth) the defensible questions are:
  (a) Are scores for a known-mutant patient shifted ABOVE matched normal cells?
      -> normal_vs_tumour_shift (this is the DNMT3A/NPM1 "healthy vs tumour"
         comparison generalised to every cancer).
  (b) Could that shift arise by chance? -> permutation / random-label nulls.

Plus the melanoma BRAF/NRAS mutual-exclusivity check the chapter requests as a
biological sanity test.

Call hierarchy
--------------
    drivers/run_pancancer -> validate_cohort_by_tier
                              -> normal_vs_tumour_shift     (Tier 2/3, image 17-22 generalised)
                              -> permutation_enrichment_null (negative control)
                              -> random_label_control        (negative control)
                              -> mutual_exclusivity_check    (SKCM sanity)
"""
from __future__ import annotations

import numpy as np
import pandas as pd

from . import stats as st
from .config import AnalysisConfig, VALIDATION_TIER, prior_of
from .data import scored_drivers
from .singlecell import resolve_score_col


# ---------------------------------------------------------------------------
# Tier 2/3 workhorse: tumour-compartment scores vs normal-cell scores
# ---------------------------------------------------------------------------
def normal_vs_tumour_shift(
    adata,
    driver: str,
    cfg: AnalysisConfig,
    reference_mask_col: str = "is_reference",
    malignant_mask_col: str | None = None,
    score_mode: str = "configured",
) -> dict:
    """Compare driver scores in the tumour compartment vs normal/reference cells.

    If a malignant mask exists (e.g. CNV-derived), the tumour compartment is those
    cells; otherwise all non-reference cells are treated as the (mixed) tumour
    compartment. Returns a distribution-shift dict: positive cliffs_delta /
    median_diff means the tumour compartment scores above normal, as expected for
    a real fingerprint.

    This is the per-cohort generalisation of your healthy-BM-vs-tumour KDE figures
    to cancers where that is the only available evidence.
    """
    score_col = resolve_score_col(adata, cfg, driver, score_mode=score_mode)
    obs = adata.obs
    if reference_mask_col not in obs.columns:
        raise KeyError(f"'{reference_mask_col}' missing; run data.mark_reference_cells.")
    normal = obs.loc[obs[reference_mask_col], score_col].astype(float).values
    if malignant_mask_col and malignant_mask_col in obs.columns:
        tumour = obs.loc[obs[malignant_mask_col].astype(bool), score_col].astype(float).values
        comp = "malignant(masked)"
    else:
        tumour = obs.loc[~obs[reference_mask_col], score_col].astype(float).values
        comp = "non-reference"
    sh = st.distribution_shift(tumour, normal, reference_quantile=cfg.reference_quantile)
    d = sh.as_dict()
    d.update(driver=driver, score_col=score_col, score_kind=cfg.score_kind(score_col),
             score_mode=score_mode, tumour_compartment=comp,
             n_tumour=len(tumour[~np.isnan(tumour)]),
             n_normal=len(normal[~np.isnan(normal)]))
    return d


# ---------------------------------------------------------------------------
# Negative control 1: permutation enrichment null
# ---------------------------------------------------------------------------
def permutation_enrichment_null(
    adata,
    driver: str,
    cfg: AnalysisConfig,
    reference_mask_col: str = "is_reference",
    score_mode: str = "configured",
) -> dict:
    """Is the tumour-vs-normal score shift larger than under random group labels?

    Permutes the reference/non-reference assignment and recomputes Cliff's delta
    n_perm times. Returns observed delta, permutation p, and the null mean/std.
    A real fingerprint should exceed its own shuffled null.
    """
    score_col = resolve_score_col(adata, cfg, driver, score_mode=score_mode)
    obs = adata.obs
    scores = obs[score_col].astype(float).values
    is_ref = obs[reference_mask_col].astype(bool).values
    keep = ~np.isnan(scores)
    scores, is_ref = scores[keep], is_ref[keep]
    # label: 1 = tumour(non-ref), 0 = normal(ref); test tumour-above-normal
    label = (~is_ref).astype(int)
    res = st.permutation_label_null(scores, label, statistic="cliffs",
                                    n_perm=cfg.n_perm, random_state=cfg.random_state)
    res.update(driver=driver, score_col=score_col, score_kind=cfg.score_kind(score_col),
               score_mode=score_mode, control="permutation_reference_label")
    return res


# ---------------------------------------------------------------------------
# Negative control 2: random-driver-label control (same class balance)
# ---------------------------------------------------------------------------
def random_label_control(
    patient_scores: pd.DataFrame,
    score_col: str,
    label_col: str,
    cfg: AnalysisConfig,
    n_random: int = 200,
) -> dict:
    """Compare the real patient-level mutant-vs-WT effect to effects obtained by
    randomly relabelling patients at the SAME class balance.

    patient_scores : output of stats.patient_aggregate merged with a 0/1 label.
    Returns observed Cliff's delta, the random-label null mean/95th percentile,
    and an empirical p. This guards against the chapter's worry that patient-level
    agreement is driven by subtype/composition rather than the driver.
    """
    s = patient_scores[score_col].values
    y = patient_scores[label_col].values.astype(int)
    n_pos = int(y.sum())
    if n_pos < 2 or (len(y) - n_pos) < 2:
        return dict(observed=np.nan, null_mean=np.nan, null_p=np.nan)
    obs_delta = st._cliffs_point(s[y == 1], s[y == 0])
    rng = np.random.default_rng(cfg.random_state)
    null = np.empty(n_random)
    for i in range(n_random):
        yr = rng.permutation(y)
        null[i] = st._cliffs_point(s[yr == 1], s[yr == 0])
    ge = int(np.sum(np.abs(null) >= abs(obs_delta)))
    return dict(observed=float(obs_delta), null_mean=float(null.mean()),
                null_p95=float(np.percentile(null, 95)),
                null_p=(ge + 1) / (n_random + 1),
                control="random_patient_label_same_balance")


# ---------------------------------------------------------------------------
# Negative control 3: the SAME per-cell statistic under two permutation units
# ---------------------------------------------------------------------------
def unit_matched_permutation_null(
    adata,
    driver: str,
    patient_status: pd.DataFrame,
    cfg: AnalysisConfig,
    patient_col: str = "patient",
    score_mode: str = "configured",
) -> dict:
    """Run ONE per-cell mutant-vs-WT statistic against TWO nulls and report both.

    The per-cell signal is Cliff's delta of the driver score in cells from
    mutant patients vs cells from WT patients (genotype broadcast from
    ``patient_status`` to every cell of the patient — so the mutation label is a
    patient-level property and block permutation is well-defined).

    * ``cell_perm_p``    — observation-level label shuffle (the historical null).
    * ``patient_perm_p`` — patient-block shuffle (fixes the number of mutant
      *patients*; the honest, anti-pseudoreplication null).

    A driver whose cell-level p is tiny but whose patient-level p is large is
    detecting a per-cell association carried by a handful of patients — i.e. the
    effect does not replicate across people. Emitting both columns in one row lets
    the manuscript show that gap directly instead of leaving the cell-level null
    and ``patient_bridge`` to contradict each other across tables.
    """
    score_col = resolve_score_col(adata, cfg, driver, score_mode=score_mode)
    obs = adata.obs
    if patient_col not in obs.columns:
        raise KeyError(f"patient_col '{patient_col}' missing.")
    status_d = patient_status[patient_status.mutation == driver][["patient", "label"]]
    lab_map = dict(zip(status_d["patient"], status_d["label"]))
    cell_pat = obs[patient_col].astype(object).values
    cell_lab = np.array([lab_map.get(p, np.nan) for p in cell_pat], dtype=float)
    score = obs[score_col].astype(float).values
    keep = ~np.isnan(score) & np.isin(cell_lab, [0.0, 1.0])
    score, lab, pats = score[keep], cell_lab[keep].astype(int), cell_pat[keep]
    n_mut_pat = int(np.unique(pats[lab == 1]).size)
    n_wt_pat = int(np.unique(pats[lab == 0]).size)
    base = dict(driver=driver, score_col=score_col,
                score_kind=cfg.score_kind(score_col), score_mode=score_mode,
                n_cells=int(keep.sum()), n_mut_patients=n_mut_pat,
                n_wt_patients=n_wt_pat,
                control="unit_matched(cell_vs_patient)")
    if n_mut_pat < 2 or n_wt_pat < 2 or lab.sum() == 0 or (lab == 0).sum() == 0:
        base.update(observed=np.nan, cell_perm_p=np.nan, patient_perm_p=np.nan,
                    power_flag="insufficient_patients")
        return base
    cell = st.permutation_label_null(score, lab, statistic="cliffs",
                                     n_perm=cfg.n_perm, random_state=cfg.random_state)
    pat = st.permutation_label_null(score, lab, statistic="cliffs",
                                    n_perm=cfg.n_perm, random_state=cfg.random_state,
                                    groups=pats)
    base.update(observed=cell["observed"],
                cell_perm_p=cell["perm_p"], cell_null_mean=cell["null_mean"],
                patient_perm_p=pat["perm_p"], patient_null_mean=pat["null_mean"],
                power_flag=("ok" if min(n_mut_pat, n_wt_pat) >= 5 else "underpowered"))
    return base


# ---------------------------------------------------------------------------
# SKCM sanity: BRAF / NRAS mutual exclusivity in score space
# ---------------------------------------------------------------------------
def mutual_exclusivity_check(
    adata,
    driver_a: str = "BRAF",
    driver_b: str = "NRAS",
    patient_col: str = "patient",
    cfg: AnalysisConfig | None = None,
) -> dict:
    """Melanoma sanity check: BRAF and NRAS are near-mutually-exclusive drivers,
    so their patient-level scores should be NEGATIVELY associated.

    Aggregates both driver scores to patient level and reports the Spearman
    correlation. A negative rho is the expected, biology-consistent result and is
    a clean reviewer-facing sanity test that the scores carry driver-specific
    (not generic-tumour) signal.
    """
    from scipy.stats import spearmanr
    cfg = cfg or AnalysisConfig()
    ca, cb = cfg.score_col(adata, driver_a), cfg.score_col(adata, driver_b)
    obs = adata.obs
    if ca not in obs.columns or cb not in obs.columns:
        return dict(driver_a=driver_a, driver_b=driver_b, rho=np.nan, p=np.nan,
                    note="one or both drivers not scored")
    agg = obs.groupby(patient_col)[[ca, cb]].median().dropna()
    if agg.shape[0] < 3:
        return dict(driver_a=driver_a, driver_b=driver_b, rho=np.nan, p=np.nan,
                    note="too few patients")
    rho, p = spearmanr(agg[ca], agg[cb])
    return dict(driver_a=driver_a, driver_b=driver_b, n_patients=agg.shape[0],
                spearman_rho=float(rho), spearman_p=float(p),
                expected="negative (mutual exclusivity)",
                consistent=bool(rho < 0))


# ---------------------------------------------------------------------------
# Orchestrator: apply the right test to each cohort by its tier
# ---------------------------------------------------------------------------
def validate_cohort_by_tier(
    adata,
    cancer: str,
    cfg: AnalysisConfig,
    patient_status: pd.DataFrame | None = None,
    drivers: list[str] | None = None,
    reference_mask_col: str = "is_reference",
    malignant_mask_col: str | None = None,
    score_mode: str = "configured",
) -> dict[str, pd.DataFrame]:
    """Run the tier-appropriate validation for one scored single-cell cohort.

    Returns a dict of result tables keyed by analysis name. Always computes the
    tumour-vs-normal shift and its permutation null (valid for every tier); adds
    the patient bridge when a patient_status table is supplied; adds the
    mutual-exclusivity check for SKCM.
    """
    drivers = drivers or scored_drivers(adata)
    tier = VALIDATION_TIER.get(cancer, "exploratory")
    print(f"[pancancer] {cancer}: tier='{tier}', {len(drivers)} drivers.")
    results: dict[str, pd.DataFrame] = {}

    # --- tumour vs normal shift + permutation null (all tiers) ---
    shift_rows, null_rows = [], []
    for d in drivers:
        try:
            sh = normal_vs_tumour_shift(adata, d, cfg, reference_mask_col, malignant_mask_col, score_mode=score_mode)
            sh.update(cancer=cancer, prior=prior_of(cancer, d), tier=tier)
            shift_rows.append(sh)
            nl = permutation_enrichment_null(adata, d, cfg, reference_mask_col, score_mode=score_mode)
            nl.update(cancer=cancer)
            null_rows.append(nl)
        except (KeyError, ValueError) as e:
            print(f"[pancancer]   {cancer}/{d}: {e}")
    shift_df = pd.DataFrame(shift_rows)
    null_df = pd.DataFrame(null_rows)
    if not null_df.empty:
        null_df["fdr_q"] = st.bh_fdr(null_df["perm_p"].values)
    results["normal_vs_tumour_shift"] = shift_df
    results["permutation_null"] = null_df

    # --- patient bridge (Tier 2 when status provided) ---
    if patient_status is not None and "patient" in adata.obs.columns:
        from .singlecell import patient_bridge
        results["patient_bridge"] = patient_bridge(
            adata, patient_status, cfg, drivers=drivers, score_mode=score_mode)

    # --- SKCM mutual-exclusivity sanity ---
    if cancer == "SKCM":
        results["mutual_exclusivity"] = pd.DataFrame(
            [mutual_exclusivity_check(adata, "BRAF", "NRAS", cfg=cfg)])

    return results
```
