## Supplemental_Code for "scOPE identifies which driver-associated expression programs transfer from bulk tumors to single cells": pathways.py

```
"""
scope_analysis.pathways
=======================
Pathway / gene-set over-representation analysis (ORA) on the gene programs scOPE
learns for each driver.

scOPE already exports, per driver, the leading signed genes of the mutation-
associated SVD components (`tables/genes_<cancer>_<driver>.csv` and
`fig2_bulk/<cancer>/component_signature_scan.csv`). This module tests those gene
lists for enrichment of curated gene sets, so a prediction ("these cells look NPM1-
mutant") can be tied to a biological programme ("HOXA/MEIS1 stemness", etc.).

Two backends, chosen automatically:
  * OFFLINE (default, no network): a hypergeometric ORA against a local `.gmt`
    file (e.g. MSigDB Hallmark `h.all.*.symbols.gmt`, or KEGG/Reactome `.gmt`).
  * ONLINE: gseapy.enrichr, if gseapy is installed and `use_gseapy=True`.

The offline path has no dependency beyond scipy and is fully reproducible, which is
what you want for a manuscript. Download a `.gmt` once from MSigDB and point to it.

Call hierarchy
--------------
    drivers/make_pathway_analysis -> enrich_driver_programs
                                      -> read_gmt / ora_hypergeometric
                                      -> plot_pathway_bar
"""
from __future__ import annotations

from pathlib import Path

import numpy as np
import pandas as pd


# ---------------------------------------------------------------------------
# gene-set IO + hypergeometric ORA
# ---------------------------------------------------------------------------
def read_gmt(path: str | Path) -> dict[str, set]:
    """Parse a .gmt (name<TAB>desc<TAB>gene1<TAB>gene2...). Returns {name: {genes}}."""
    sets = {}
    with open(path) as fh:
        for line in fh:
            parts = line.rstrip("\n").split("\t")
            if len(parts) < 3:
                continue
            name, _desc, *genes = parts
            g = {x.strip().upper() for x in genes if x.strip()}
            if g:
                sets[name] = g
    return sets


def ora_hypergeometric(genes, gene_sets: dict[str, set], background=None,
                       min_set_size: int = 5, max_set_size: int = 1000) -> pd.DataFrame:
    """Over-representation analysis by the hypergeometric test.

    genes      : query gene list (e.g. a driver's leading genes), symbols.
    gene_sets  : {term: {genes}} from read_gmt.
    background : universe of testable genes (e.g. all bulk gene symbols). If None,
                 the union of all gene-set members is used (a reasonable default,
                 but supplying your assayed-gene universe is more correct).
    Returns a table sorted by FDR: [term, overlap, set_size, n_query, n_background,
    p_value, fdr, odds_ratio, genes].
    """
    from scipy.stats import hypergeom
    query = {str(g).strip().upper() for g in genes if str(g).strip()}
    if background is None:
        background = set().union(*gene_sets.values()) if gene_sets else set()
    else:
        background = {str(g).strip().upper() for g in background}
    query &= background
    N = len(background)
    n = len(query)
    if N == 0 or n == 0:
        return pd.DataFrame()

    rows = []
    for term, gs in gene_sets.items():
        gs = gs & background
        K = len(gs)
        if K < min_set_size or K > max_set_size:
            continue
        hit = query & gs
        k = len(hit)
        if k == 0:
            continue
        # P(X >= k) for X ~ Hypergeom(N, K, n)
        p = hypergeom.sf(k - 1, N, K, n)
        exp = n * K / N
        odds = (k / exp) if exp > 0 else np.nan
        rows.append(dict(term=term, overlap=k, set_size=K, n_query=n, n_background=N,
                         p_value=float(p), fold_enrichment=float(odds),
                         genes=",".join(sorted(hit))))
    out = pd.DataFrame(rows)
    if out.empty:
        return out
    out = out.sort_values("p_value").reset_index(drop=True)
    # Benjamini-Hochberg FDR
    m = len(out)
    out["fdr"] = (out["p_value"] * m / (np.arange(1, m + 1))).clip(upper=1.0)
    out["fdr"] = out["fdr"][::-1].cummin()[::-1]
    return out


# ---------------------------------------------------------------------------
# driver programmes -> enrichment
# ---------------------------------------------------------------------------
def _leading_genes_from_table(df: pd.DataFrame, direction: str | None, top_n: int) -> list[str]:
    """Pull top genes from a genes_<cancer>_<driver>.csv / component_signature_scan
    table. Prefers ranking by |loading|; optionally filter by up/down direction."""
    d = df.copy()
    if direction in ("up", "down") and "direction" in d.columns:
        d = d[d["direction"] == direction]
    if "loading" in d.columns:
        d = d.reindex(d["loading"].abs().sort_values(ascending=False).index)
    return d["gene"].astype(str).head(top_n).tolist()


def enrich_driver_programs(genes_table: pd.DataFrame, gmt_path: str | Path,
                           background=None, top_n: int = 100, by_direction: bool = True,
                           use_gseapy: bool = False, organism: str = "human") -> pd.DataFrame:
    """Run ORA for one driver's gene programme. Returns a tidy enrichment table with a
    'direction' column ('combined'/'up'/'down'). Falls back to hypergeometric ORA
    against gmt_path unless use_gseapy=True and gseapy is importable."""
    blocks = []
    directions = [None] if not by_direction else [None, "up", "down"]
    if use_gseapy:
        try:
            import gseapy as gp
            for direction in directions:
                gl = _leading_genes_from_table(genes_table, direction, top_n)
                if len(gl) < 5:
                    continue
                enr = gp.enrichr(gene_list=gl, gene_sets=str(gmt_path), organism=organism,
                                 outdir=None, no_plot=True)
                t = enr.results.rename(columns={"Term": "term", "Adjusted P-value": "fdr",
                                                "P-value": "p_value", "Genes": "genes"})
                t["direction"] = direction or "combined"
                blocks.append(t)
            if blocks:
                return pd.concat(blocks, ignore_index=True)
        except Exception as e:
            print(f"[pathways] gseapy unavailable/failed ({e}); using offline hypergeometric ORA.")

    gene_sets = read_gmt(gmt_path)
    for direction in directions:
        gl = _leading_genes_from_table(genes_table, direction, top_n)
        if len(gl) < 5:
            continue
        t = ora_hypergeometric(gl, gene_sets, background=background)
        if not t.empty:
            t["direction"] = direction or "combined"
            blocks.append(t)
    return pd.concat(blocks, ignore_index=True) if blocks else pd.DataFrame()


def plot_pathway_bar(enrich_table: pd.DataFrame, outpath, title: str = "",
                     top_n: int = 15, direction: str = "combined", dpi: int = 200):
    """Horizontal bar of the top enriched terms (-log10 FDR) for one direction."""
    import matplotlib
    matplotlib.use("Agg")
    import matplotlib.pyplot as plt
    t = enrich_table[enrich_table.get("direction", "combined") == direction].copy()
    if t.empty:
        return None
    t = t.sort_values("fdr").head(top_n).iloc[::-1]
    y = np.arange(len(t))
    val = -np.log10(t["fdr"].clip(lower=1e-300))
    fig, ax = plt.subplots(figsize=(8.5, max(2.4, 0.42 * len(t))))
    ax.barh(y, val, color="#4575b4", alpha=0.9)
    ax.set_yticks(y); ax.set_yticklabels([s[:48] for s in t["term"]], fontsize=8)
    ax.axvline(-np.log10(0.05), color="grey", ls="--", lw=1)
    ax.set_xlabel("-log10 FDR"); ax.set_title(title, fontsize=11, loc="left")
    for sp in ("top", "right"):
        ax.spines[sp].set_visible(False)
    fig.tight_layout()
    out = Path(outpath); fig.savefig(out, dpi=dpi, bbox_inches="tight"); plt.close(fig)
    return out
```
