## Supplemental_Code for "scOPE identifies which driver-associated expression programs transfer from bulk tumors to single cells": plots.py

```
"""
scope_analysis.plots
====================
Manuscript figure builders. Every function returns the saved file path(s) and is
keyed to a specific chapter figure, so the figure list and the code never drift.

Where scope-bio already has the right plot, this module CALLS it rather than
re-implementing (UMAP probability overlays, scree, per-cluster heatmap). The
custom builders here are the panels scope-bio does not provide: forest plots with
CIs coloured by prior tier, ROC overlays, expressed-mutation annotation violins, the
treatment-response trajectory, and the healthy-vs-tumour density overlays.

Style: one consistent look (white background, no top/right spines, colour-blind-
safe palette). Prior tiers are coloured strong=teal, intermediate=amber,
weak=grey so the strong-vs-subtle gradient reads at a glance.

Call hierarchy
--------------
    drivers/* and report.py -> the figure functions below
    several functions       -> scope.visualization.{compute_umap,
                               plot_mutation_probabilities, plot_mutation_heatmap,
                               plot_scree}
"""
from __future__ import annotations

from pathlib import Path

import matplotlib
matplotlib.use("Agg")
import matplotlib.pyplot as plt
from matplotlib.lines import Line2D
import numpy as np
import pandas as pd
from scipy.stats import gaussian_kde, mannwhitneyu

from .stats import auroc_safe, bh_fdr, cliffs_delta

# Manuscript figure quality: a single knob for crisp output everywhere.
FIGURE_DPI = 400

# Significance cutoff (BH-FDR q) for the Fig3 single-cell plots only. Loosened to 0.10
# given the low power of the per-cell PU analysis; other figures keep the 0.05 default.
FIG3_FDR_ALPHA = 0.10

# High-contrast palette for boolean / 2-3 category panels (e.g. is_reference),
# so True/False are never blue-on-blue and read clearly in print.
_BOOL_COLORS = {
    "true": "#d62728", "reference": "#d62728", "yes": "#d62728", "1": "#d62728",
    "false": "#9aa0a6", "non-reference": "#9aa0a6", "no": "#9aa0a6", "0": "#9aa0a6",
}
_TWO_CAT_FALLBACK = ["#4C72B0", "#DD8452", "#55A868"]  # distinct, colourblind-friendly

_TIER_COLOR = {"strong": "#1b9e77", "intermediate": "#d95f02",
               "weak": "#7570b3", "unspecified": "#999999"}
_PALETTE = {"WT": "#4575b4", "Mut": "#d73027", "Unknown": "#bdbdbd",
            "Healthy (reference)": "#4575b4", "Tumour": "#fc8d59"}


def _axis_label(driver: str, score_label: str | None, fallback: str = "score") -> str:
    """Avoid labels like 'NPM1 NPM1 same-cell-state residual'."""
    lab = str(score_label or fallback)
    if driver and lab.strip().lower().startswith(str(driver).lower()):
        return lab
    return f"{driver} {lab}" if driver else lab


def _finite_stat(x):
    try:
        if x is None or pd.isna(x):
            return None
        v = float(x)
        return v if np.isfinite(v) else None
    except Exception:
        return None


def _format_driver_stats(stats: dict | None) -> str:
    """Compact annotation for available driver confidence / ground-truth stats."""
    if not stats:
        return ""
    bits = []
    conf = _finite_stat(stats.get("confidence", stats.get("driver_confidence")))
    if conf is not None:
        bits.append(f"conf={conf:.2f}")
    p = _finite_stat(stats.get("mannwhitney_p", stats.get("p", stats.get("p_value"))))
    if p is not None:
        bits.append(f"MWU p={p:.1e}")
    q = _finite_stat(stats.get("fdr_q", stats.get("q")))
    if q is not None:
        bits.append(f"q={q:.1e}")
    delta = _finite_stat(stats.get("cliffs_delta"))
    if delta is not None:
        bits.append(f"δ={delta:.2f}")
    auc = _finite_stat(stats.get("apparent_auroc", stats.get("auroc")))
    if auc is not None:
        bits.append(f"AUC={auc:.2f}")
    return " · ".join(bits)


def _timepoint_sort_key(t) -> int:
    return int("".join(c for c in str(t) if c.isdigit()) or 0)


def _clean_scores(values) -> np.ndarray:
    arr = np.asarray(values, dtype=float)
    return arr[np.isfinite(arr)]


def _sig_stars(p: float | None) -> str:
    """Star labels from an adjusted q-value when available, else raw p-value."""
    p = _finite_stat(p)
    if p is None:
        return ""
    if p < 1e-4:
        return "****"
    if p < 1e-3:
        return "***"
    if p < 1e-2:
        return "**"
    if p < 0.05:
        return "*"
    return "ns"


def _safe_mannwhitney(x: np.ndarray, y: np.ndarray) -> tuple[float, float]:
    try:
        res = mannwhitneyu(x, y, alternative="two-sided")
        return float(res.statistic), float(res.pvalue)
    except Exception:
        return np.nan, np.nan


def _add_sig_bracket(ax, x1: float, x2: float, y: float, h: float, label: str,
                     fontsize: int = 7, lw: float = 0.8) -> None:
    ax.plot([x1, x1, x2, x2], [y, y + h, y + h, y], color="black", lw=lw, clip_on=False)
    ax.text((x1 + x2) / 2.0, y + h, label, ha="center", va="bottom",
            fontsize=fontsize, clip_on=False)


def _natural_label_order(labels) -> list[str]:
    """Stable visual/statistical order for genotype labels, with fallbacks."""
    preferred = ["WT", "Mut", "Unknown"]
    seen = [str(x) for x in labels if pd.notna(x)]
    out = [x for x in preferred if x in set(seen)]
    out.extend(sorted([x for x in set(seen) if x not in set(out)]))
    return out


def timepoint_genotype_violin_stats(
    split: pd.DataFrame,
    min_n: int = 3,
    comparison_mode: str = "full",
) -> pd.DataFrame:
    """Mann-Whitney comparisons for per-patient longitudinal violin panels.

    Default ``comparison_mode='full'`` emits the two comparison families that are
    useful for mixed WT/Mut/Unknown longitudinal panels:

      1. ``within_timepoint_label_pairwise`` — all pairwise label/status
         contrasts within each timepoint, e.g. WT vs Mut, WT vs Unknown,
         Mut vs Unknown.
      2. ``between_timepoint_same_label_pairwise`` — all pairwise timepoint
         contrasts within each label/status, e.g. D0 vs D15 and D0 vs D31
         among WT cells only, then again among Mut cells, then Unknown.

    Positive ``cliffs_delta_b_vs_a`` and ``median_diff_b_minus_a`` always mean
    group B tends to have higher scores than group A. For between-timepoint
    comparisons, group B is the later timepoint.

    Supported modes:
      * ``full`` / ``auto`` / ``status_and_timepoint``: both families above.
      * ``within_timepoint`` / ``genotype``: label/status contrasts only.
      * ``timepoint`` / ``longitudinal``: all pairwise timepoint contrasts
        within each label/status.
      * ``adjacent``: adjacent timepoint contrasts only within each label/status.
    """
    if split is None or split.empty or "score" not in split.columns:
        return pd.DataFrame()
    needed = {"timepoint", "genotype_label", "score"}
    if not needed.issubset(split.columns):
        return pd.DataFrame()

    work = split.copy()
    work["genotype_label"] = work["genotype_label"].astype(str)
    tps = sorted(work["timepoint"].dropna().unique().tolist(), key=_timepoint_sort_key)
    labels_present = _natural_label_order(work["genotype_label"].dropna().unique().tolist())

    def vals(tp, lab):
        return _clean_scores(work.loc[(work["timepoint"] == tp) &
                                      (work["genotype_label"] == str(lab)), "score"].values)

    rows = []
    mode = str(comparison_mode or "full").lower()
    do_within = mode in {
        "auto", "full", "status_and_timepoint", "all", "genotype", "status",
        "within_timepoint", "within_timepoint_genotype", "within_timepoint_label",
        "within_timepoint_label_pairwise",
    }
    do_between_all = mode in {
        "auto", "full", "status_and_timepoint", "all", "timepoint", "longitudinal",
        "between_timepoint", "between_timepoint_same_label", "between_timepoint_same_label_pairwise",
    }
    do_between_adjacent = mode in {"adjacent", "adjacent_timepoint", "adjacent_timepoint_same_label"}

    # 1) Status/genotype contrasts within each timepoint.
    if do_within and len(labels_present) >= 2:
        from itertools import combinations
        for tp in tps:
            for lab_a, lab_b in combinations(labels_present, 2):
                a = vals(tp, lab_a)
                b = vals(tp, lab_b)
                if len(a) < min_n or len(b) < min_n:
                    continue
                U, pval = _safe_mannwhitney(a, b)
                rows.append(dict(
                    comparison_family="status_within_timepoint",
                    comparison_type="within_timepoint_label_pairwise",
                    timepoint=tp,
                    timepoint_a=tp,
                    timepoint_b=tp,
                    label=f"{lab_b}_vs_{lab_a}",
                    genotype_label_a=lab_a,
                    genotype_label_b=lab_b,
                    group_a=lab_a,
                    group_b=lab_b,
                    n_a=len(a),
                    n_b=len(b),
                    median_a=float(np.nanmedian(a)),
                    median_b=float(np.nanmedian(b)),
                    median_diff_b_minus_a=float(np.nanmedian(b) - np.nanmedian(a)),
                    cliffs_delta_b_vs_a=float(cliffs_delta(b, a, n_boot=0)),
                    mannwhitney_u=U,
                    p_value=pval,
                ))

    # 2) Longitudinal contrasts within the same status/genotype label.
    if (do_between_all or do_between_adjacent) and len(tps) >= 2:
        from itertools import combinations
        if do_between_adjacent and not do_between_all:
            tp_pairs = list(zip(tps[:-1], tps[1:]))
        else:
            tp_pairs = list(combinations(tps, 2))
        for lab in labels_present:
            for tp_a, tp_b in tp_pairs:
                a = vals(tp_a, lab)
                b = vals(tp_b, lab)
                if len(a) < min_n or len(b) < min_n:
                    continue
                U, pval = _safe_mannwhitney(a, b)
                rows.append(dict(
                    comparison_family="timepoint_within_status",
                    comparison_type="between_timepoint_same_label_pairwise",
                    timepoint=f"{tp_a}_vs_{tp_b}",
                    timepoint_a=tp_a,
                    timepoint_b=tp_b,
                    label=lab,
                    genotype_label_a=lab,
                    genotype_label_b=lab,
                    group_a=f"{tp_a}:{lab}",
                    group_b=f"{tp_b}:{lab}",
                    n_a=len(a),
                    n_b=len(b),
                    median_a=float(np.nanmedian(a)),
                    median_b=float(np.nanmedian(b)),
                    median_diff_b_minus_a=float(np.nanmedian(b) - np.nanmedian(a)),
                    cliffs_delta_b_vs_a=float(cliffs_delta(b, a, n_boot=0)),
                    mannwhitney_u=U,
                    p_value=pval,
                ))

    out = pd.DataFrame(rows)
    if not out.empty:
        out["fdr_q"] = bh_fdr(out["p_value"].values)
        out["significance"] = [_sig_stars(q) for q in out["fdr_q"].values]
        out["test"] = "Mann-Whitney U, two-sided; BH-FDR across all comparisons in this violin panel"
    return out

def _style(ax):
    ax.spines[["top", "right"]].set_visible(False)
    return ax


def _score_kind_from_col(col: str, cfg=None) -> str:
    if cfg is not None and hasattr(cfg, "score_kind"):
        return cfg.score_kind(col)
    col = str(col)
    if "mutation_calib_" in col:
        return "calibrated_reference_percentile"
    if "mutation_prob_" in col:
        return "raw_probability"
    if "mutation_adj_" in col:
        return "confidence_adjusted_raw"
    if "mutation_resid_" in col:
        return "cellstate_residual_raw"
    if "mutation_refpct_" in col:
        return "cellstate_reference_percentile"
    return "score"


def _score_label(driver: str, col: str, cfg=None) -> str:
    if cfg is not None and hasattr(cfg, "score_label_for_col"):
        return cfg.score_label_for_col(col, driver)
    kind = _score_kind_from_col(col, cfg)
    if kind == "calibrated_reference_percentile":
        return f"{driver} reference-percentile score"
    if kind == "raw_probability":
        return f"{driver} raw driver-program score"
    if kind == "confidence_adjusted_raw":
        return f"{driver} confidence-adjusted raw score"
    if kind == "cellstate_residual_raw":
        return f"{driver} same-cell-state residual"
    if kind == "cellstate_reference_percentile":
        return f"{driver} same-cell-state reference percentile"
    return f"{driver} score"


def _select_score_col(adata, driver: str, cfg=None, score_mode: str = "configured") -> str:
    mode = str(score_mode or "configured").lower()
    if mode in {"cellstate_residual", "residual", "guarded"}:
        c = f"mutation_resid_{driver}"
        if c in adata.obs.columns:
            return c
    if mode in {"cellstate_percentile", "percentile", "refpct"}:
        c = f"mutation_refpct_{driver}"
        if c in adata.obs.columns:
            return c
    if mode in {"raw", "raw_probability"}:
        return cfg.score_col(adata, driver, use_calibrated=False) if cfg is not None else f"mutation_prob_{driver}"
    return cfg.score_col(adata, driver) if cfg is not None else f"mutation_prob_{driver}"


def _display_score_col(adata, driver: str, cfg=None, score_mode: str = "configured",
                       prefer_guarded_for_reference_high: bool = False) -> str:
    """Resolve the display score column.

    For main UMAP probability panels we often want a safer default: if a driver is
    flagged as reference-high (healthy reference systematically above tumour, as in
    NPM1/HSPC overlap), use the same-cell-state guarded score for display when the
    caller asked for configured/main plots. Explicit raw/guarded score_mode values
    always win.
    """
    mode = str(score_mode or "configured").lower()
    explicit = mode in {"cellstate_residual", "residual", "guarded",
                        "cellstate_percentile", "percentile", "refpct",
                        "raw", "raw_probability"}
    if (not explicit) and prefer_guarded_for_reference_high:
        ref_high = set((adata.uns.get("reference_high_drivers", []) or []))
        if driver in ref_high:
            guarded_mode = getattr(cfg, "longitudinal_guarded_mode", "cellstate_residual") if cfg is not None else "cellstate_residual"
            gcol = _select_score_col(adata, driver, cfg=cfg, score_mode=guarded_mode)
            if gcol in adata.obs.columns:
                return gcol
    return _select_score_col(adata, driver, cfg=cfg, score_mode=score_mode)


def _robust_limits(values, cfg=None, col: str | None = None):
    """Return plot-only color limits plus a short scale note.

    Same-cell-state residuals need special handling: weak/exploratory drivers often
    have microscopic numeric ranges.  Per-panel q01-q99 scaling makes that noise look
    bright.  For residual columns we therefore use a symmetric color scale around
    zero with a configurable minimum span.  Statistics are never affected.
    """
    x = np.asarray(values, dtype=float)
    x = x[np.isfinite(x)]
    if x.size == 0:
        return 0.0, 1.0, ""
    kind = _score_kind_from_col(col or "", cfg)
    if kind == "cellstate_residual_raw":
        uq = float(getattr(cfg, "plot_robust_upper_q", 0.99)) if cfg is not None else 0.99
        min_abs = float(getattr(cfg, "plot_residual_min_abs_vmax", 0.05)) if cfg is not None else 0.05
        max_abs = float(getattr(cfg, "plot_residual_abs_vmax", 0.25)) if cfg is not None else 0.25
        try:
            vmax = float(np.nanquantile(np.abs(x), uq))
        except Exception:
            vmax = float(np.nanmax(np.abs(x)))
        if not np.isfinite(vmax) or vmax <= 0:
            vmax = min_abs
        vmax = min(max(vmax, min_abs), max_abs)
        return -vmax, vmax, f"centered ±{vmax:g} color scale"
    if kind in {"calibrated_reference_percentile", "cellstate_reference_percentile"}:
        return 0.0, 1.0, ""
    mode = getattr(cfg, "plot_score_scale", "auto_robust") if cfg is not None else "auto_robust"
    mode_l = str(mode).lower()
    if mode_l in {"fixed01", "probability", "0_1"}:
        return 0.0, 1.0, "fixed 0-1 scale"
    if mode_l in {"absolute", "none", "fixed"}:
        lo, hi = float(np.nanmin(x)), float(np.nanmax(x))
        if hi <= lo: hi = lo + 1e-9
        return lo, hi, ""
    lq = float(getattr(cfg, "plot_robust_lower_q", 0.01)) if cfg is not None else 0.01
    uq = float(getattr(cfg, "plot_robust_upper_q", 0.99)) if cfg is not None else 0.99
    min_range = float(getattr(cfg, "plot_robust_min_range", 0.05)) if cfg is not None else 0.05
    lo, hi = np.nanquantile(x, [lq, uq])
    if not np.isfinite(lo) or not np.isfinite(hi) or hi <= lo:
        lo, hi = float(np.nanmin(x)), float(np.nanmax(x))
    if hi <= lo:
        hi = lo + 1e-9
    full_range = float(np.nanmax(x) - np.nanmin(x))
    compressed = (full_range < min_range) or (float(np.nanmax(x)) < 0.15 and kind == "raw_probability")
    if compressed:
        return float(lo), float(hi), f"q{int(lq*100):02d}-q{int(uq*100):02d} display scale"
    lo, hi = float(np.nanmin(x)), float(np.nanmax(x))
    if kind == "raw_probability" and lo >= 0 and hi <= 1:
        return 0.0, 1.0, ""
    return lo, hi, ""


def _robust_rescale_column(values, lower_q=0.01, upper_q=0.99):
    x = np.asarray(values, dtype=float)
    out = np.full_like(x, np.nan, dtype=float)
    finite = np.isfinite(x)
    if not finite.any():
        return out
    lo, hi = np.nanquantile(x[finite], [lower_q, upper_q])
    if not np.isfinite(lo) or not np.isfinite(hi) or hi <= lo:
        lo, hi = np.nanmin(x[finite]), np.nanmax(x[finite])
    if hi <= lo:
        out[finite] = 0.5
    else:
        out[finite] = np.clip((x[finite] - lo) / (hi - lo), 0, 1)
    return out


# ===========================================================================
# FIGURE 2 / TABLE 1 — bulk discrimination
# ===========================================================================
def forest_bulk_auroc(bulk_table: pd.DataFrame, outpath: str | Path,
                      title: str = "Bulk cross-validated AUROC (95% CI)") -> Path:
    """Forest plot of OOF AUROC with bootstrap CIs, one row per cancer x driver,
    coloured by pre-specified prior tier, sorted within cancer. Reference line at
    0.5; claim-unsafe pairs drawn hollow. This is the Fig. 2 / Table 1 visual."""
    df = bulk_table.dropna(subset=["auroc"]).copy()
    df = df.sort_values(["cancer", "auroc"], ascending=[True, True]).reset_index(drop=True)
    n = len(df)
    fig, ax = plt.subplots(figsize=(7.5, max(3, 0.32 * n)))
    for i, r in df.iterrows():
        color = _TIER_COLOR.get(r["prior"], "#999")
        safe = bool(r.get("claim_safe", False))
        ax.plot([r["auroc_ci_lo"], r["auroc_ci_hi"]], [i, i], color=color, lw=2, zorder=1)
        ax.scatter([r["auroc"]], [i], s=46, color=color if safe else "white",
                   edgecolor=color, linewidth=1.6, zorder=2)
    ax.axvline(0.5, ls="--", color="grey", lw=1)
    ax.set_yticks(range(n))
    ax.set_yticklabels([f"{r.cancer} · {r.driver}" for r in df.itertuples()], fontsize=8)
    ax.set_xlabel("AUROC (out-of-fold)"); ax.set_xlim(0.3, 1.02)
    ax.set_title(title)
    handles = [plt.Line2D([0], [0], marker="o", color="w", markerfacecolor=c,
                          markeredgecolor=c, label=t, markersize=8)
               for t, c in _TIER_COLOR.items() if t != "unspecified"]
    handles.append(plt.Line2D([0], [0], marker="o", color="w", markerfacecolor="white",
                              markeredgecolor="black", label="claim-unsafe (hollow)", markersize=8))
    ax.legend(handles=handles, title="prior fingerprint", fontsize=7,
              title_fontsize=8, loc="lower right", frameon=False)
    _style(ax); fig.tight_layout()
    out = Path(outpath); fig.savefig(out, dpi=FIGURE_DPI, bbox_inches="tight"); plt.close(fig)
    return out


def roc_overlay(oof_store: dict, cancer: str, drivers: list[str],
                outpath: str | Path) -> Path:
    """Overlay OOF ROC curves for several drivers of one cancer (image 7 style).
    Reads the OOF predictions stashed on the bulk table (bulk_table.attrs)."""
    from sklearn.metrics import roc_curve
    oof = oof_store[cancer]
    fig, ax = plt.subplots(figsize=(5.2, 5))
    for d in drivers:
        sub = oof[oof.driver == d].dropna(subset=["y_oof"])
        y, p = sub.y_true.values, sub.y_oof.values
        if len(np.unique(y)) < 2:
            continue
        fpr, tpr, _ = roc_curve(y, p)
        ax.plot(fpr, tpr, lw=2, label=f"{d} (AUC={auroc_safe(y,p):.3f})")
    ax.plot([0, 1], [0, 1], ls="--", color="grey", lw=1)
    ax.set_xlabel("False positive rate"); ax.set_ylabel("True positive rate")
    ax.set_title(f"Bulk OOF ROC — {cancer}"); ax.legend(fontsize=8, frameon=False, loc="lower right")
    _style(ax); fig.tight_layout()
    out = Path(outpath); fig.savefig(out, dpi=FIGURE_DPI, bbox_inches="tight"); plt.close(fig)
    return out


# ===========================================================================
# FIGURE 3 — single-cell validation
# ===========================================================================
def percell_violin(cell_long: pd.DataFrame, driver: str, outpath: str | Path,
                   p_annot: float | None = None, q_annot: float | None = None,
                   score_label: str = "score", reference_scores=None,
                   describe: str = "calibrated", show_reference: bool = False,
                   confidence: float | None = None) -> Path:
    """Per-cell score violin split by genotype (Mut vs WT), with optional healthy
    reference and per-group medians + counts.

    show_reference defaults to False: under reference-ECDF calibration the reference
    is uniform on [0,1] by construction, so its violin is an uninformative blob
    spanning the whole axis. The WT-vs-Mut contrast is the actual signal. Pass
    show_reference=True to restore the reference context violin.
    """
    sub = cell_long[cell_long.driver == driver]
    if sub.empty:
        return None
    groups, data, colors = [], [], []
    if show_reference and reference_scores is not None and len(reference_scores) > 1:
        groups.append("Reference"); data.append(np.asarray(reference_scores, float))
        colors.append(_PALETTE["Healthy (reference)"])
    for g in ["WT", "Mut"]:
        if g in sub.genotype_label.unique():
            vals = sub[sub.genotype_label == g]["score"].values
            vals = vals[~np.isnan(vals)]
            if len(vals):
                groups.append(g); data.append(vals); colors.append(_PALETTE[g])
    if not data:
        return None
    fig, ax = plt.subplots(figsize=(1.5 * len(groups) + 2, 4.6))
    parts = ax.violinplot(data, showmedians=False, showextrema=False)
    for pc, c in zip(parts["bodies"], colors):
        pc.set_facecolor(c); pc.set_alpha(0.65)
    for i, (vals, c) in enumerate(zip(data, colors), start=1):
        med = np.median(vals)
        ax.hlines(med, i - 0.28, i + 0.28, color="black", lw=1.6, zorder=5)
        # light jittered points for small groups (shows real cells, not just KDE)
        if len(vals) <= 600:
            xj = np.random.default_rng(0).normal(i, 0.05, len(vals))
            ax.scatter(xj, vals, s=5, color=c, alpha=0.35, edgecolor="none", zorder=3)
    ax.set_xticks(range(1, len(groups) + 1))
    ax.set_xticklabels([f"{g}\n(n={len(d)})" for g, d in zip(groups, data)], fontsize=8)
    if score_label and score_label != "score":
        ylab = score_label
    else:
        ylab = (f"{driver} reference-percentile score\n(vs healthy)"
                if str(describe).startswith("calibrated")
                else f"{driver} raw driver-program score")
    ax.set_ylabel(ylab, fontsize=9)
    # auto-scale with a little headroom; clamp to [0,1] when within it
    allv = np.concatenate(data)
    lo, hi = np.nanmin(allv), np.nanmax(allv)
    pad = 0.05 * (hi - lo + 1e-9)
    ax.set_ylim(max(-0.02, lo - pad), min(1.02, hi + pad) if hi <= 1 else hi + pad)
    title = f"{driver}: direct-label score enrichment"
    if p_annot is not None:
        title += f"\nMann–Whitney p = {p_annot:.2e}"
        if q_annot is not None and np.isfinite(q_annot):
            title += f"; FDR q = {q_annot:.2e}"
            if q_annot >= FIG3_FDR_ALPHA and p_annot < 0.05:
                title += f" (nominal; not FDR<{FIG3_FDR_ALPHA:g})"
    if confidence is not None and np.isfinite(confidence):
        title += f"   ·   confidence = {confidence:.2f}"
    ax.set_title(title, fontsize=10)
    _style(ax); fig.tight_layout()
    out = Path(outpath); fig.savefig(out, dpi=FIGURE_DPI, bbox_inches="tight"); plt.close(fig)
    return out


def patient_bridge_strip(bridge_table: pd.DataFrame, adata, driver: str,
                         patient_status: pd.DataFrame, outpath: str | Path,
                         patient_col: str = "patient") -> Path:
    """Patient-level aggregated scores, mutant vs WT patients, as a strip+box.
    The unit is the patient (anti-pseudoreplication), matching the chapter."""
    from .stats import patient_aggregate
    sc = f"mutation_prob_{driver}"
    agg = patient_aggregate(adata.obs[[patient_col, sc]].copy(), patient_col, sc)
    status = patient_status[patient_status.mutation == driver][["patient", "label"]]
    m = agg.merge(status, left_on=patient_col, right_on="patient", how="inner")
    if m.empty:
        return None
    fig, ax = plt.subplots(figsize=(4, 4.4))
    for lab, name, color in [(0, "WT", _PALETTE["WT"]), (1, "Mut", _PALETTE["Mut"])]:
        vals = m[m.label == lab][sc].values
        x = np.random.default_rng(0).normal(lab, 0.06, len(vals))
        ax.scatter(x, vals, color=color, alpha=0.8, s=40, edgecolor="k", linewidth=0.3)
        if len(vals):
            ax.hlines(np.median(vals), lab - 0.2, lab + 0.2, color=color, lw=2)
    ax.set_xticks([0, 1]); ax.set_xticklabels(["WT", "Mut"])
    ax.set_ylabel(f"patient median P({driver})")
    ax.set_title(f"{driver}: patient-level bridge", fontsize=10)
    _style(ax); fig.tight_layout()
    out = Path(outpath); fig.savefig(out, dpi=FIGURE_DPI, bbox_inches="tight"); plt.close(fig)
    return out


# ===========================================================================
# FIGURE 3/4 — TREATMENT-RESPONSE (the timepoint gold)
# ===========================================================================
def treatment_response_trajectory(trajectory: pd.DataFrame, driver: str,
                                  outpath: str | Path,
                                  metric: str = "cliffs_delta") -> Path:
    """One line per patient over treatment.

    For raw/global reference trajectories, `cliffs_delta` answers whether the sample
    is above the selected healthy reference. For NPM1-like reference-high drivers,
    guarded residual trajectories should usually be plotted with `median_diff` so
    zero means matched healthy reference.
    """
    sub = trajectory[trajectory.driver == driver]
    if sub.empty or metric not in sub.columns:
        return None
    fig, ax = plt.subplots(figsize=(6.4, 4.7))
    cmap = plt.get_cmap("tab10")
    for i, (pat, g) in enumerate(sub.groupby("patient")):
        g = g.sort_values("day")
        ax.plot(g["day"], g[metric], marker="o", color=cmap(i % 10), label=pat, lw=1.8)
    ax.axhline(0, ls="--", color="grey", lw=1)
    ax.set_xlabel("Treatment day")
    score_label = str(sub.get("score_label", pd.Series(["score"])).dropna().iloc[0]) if "score_label" in sub else "score"
    score_mode = str(sub.get("score_mode", pd.Series([""])).dropna().iloc[0]) if "score_mode" in sub else ""
    ref_col = str(sub.get("reference_mask_col", pd.Series(["healthy reference"])).dropna().iloc[0]) if "reference_mask_col" in sub else "healthy reference"
    if metric == "cliffs_delta":
        ylab = f"Cliff's δ vs {ref_col}"
    elif metric == "median_diff":
        ylab = f"Median difference vs {ref_col}"
    elif metric == "median_score":
        ylab = f"Median {score_label}"
    else:
        ylab = metric
    ax.set_ylabel(ylab)
    ref_high = False
    if "reference_direction" in sub.columns:
        d0 = sub[sub["day"] == sub["day"].min()]
        ref_high = (d0.get("reference_direction", pd.Series([], dtype=str)).astype(str) == "sample_below_reference").mean() > 0.5 if not d0.empty else False
    title = f"{driver}: treatment trajectory"
    if score_mode:
        title += f" ({score_mode})"
    if ref_high and metric == "cliffs_delta":
        title += "\nreference-high at baseline: use guarded residual panel for interpretation"
    ax.set_title(title, fontsize=10)
    ax.legend(fontsize=7, frameon=False, ncol=2, title="patient", title_fontsize=8)
    _style(ax); fig.tight_layout()
    out = Path(outpath); fig.savefig(out, dpi=FIGURE_DPI, bbox_inches="tight"); plt.close(fig)
    return out


def density_overlay(overlay: dict, outpath: str | Path,
                    score_label: str | None = None,
                    driver_stats: dict | None = None) -> Path:
    """Healthy-vs-tumour KDE overlays across timepoints for one patient/driver.

    For guarded residual/percentile overlays the annotation should reflect the
    guarded score rather than a raw Cliff's-δ story, so we dynamically choose a
    more interpretable subtitle from the score kind."""
    score_label = score_label or overlay.get("score_label", "score")
    driver_stats = driver_stats or overlay.get("driver_stats")
    stats_note = _format_driver_stats(driver_stats)
    score_kind = str(overlay.get("score_kind", ""))
    tps = list(overlay["timepoints"].keys())
    if not tps:
        return None
    ref = np.asarray(overlay["reference"], float)
    # global x-range across reference + all timepoints
    allv = [ref] + [np.asarray(overlay["timepoints"][tp]["scores"], float) for tp in tps]
    allv = np.concatenate([a[~np.isnan(a)] for a in allv if len(a)])
    lo, hi = np.nanmin(allv), np.nanmax(allv)
    if hi <= lo:
        hi = lo + 1e-3
    pad = 0.04 * (hi - lo)
    grid = np.linspace(lo - pad, hi + pad, 256)
    fig, axes = plt.subplots(1, len(tps), figsize=(3.2 * len(tps), 3.4), squeeze=False, sharex=True)
    for j, tp in enumerate(tps):
        ax = axes[0][j]
        s = np.asarray(overlay["timepoints"][tp]["scores"], float)
        sh = overlay["timepoints"][tp]["shift"]
        for arr, name in [(ref, "Healthy (reference)"), (s, "Tumour")]:
            arr = arr[~np.isnan(arr)]
            if len(arr) > 1 and np.std(arr) > 1e-9:
                kde = gaussian_kde(arr)
                ax.fill_between(grid, kde(grid), alpha=0.5, color=_PALETTE[name], label=name)
            elif len(arr) >= 1:
                ax.axvline(np.mean(arr), color=_PALETTE[name], lw=2, alpha=0.6, label=name)
        if score_kind in {"cellstate_residual_raw", "cellstate_reference_percentile"}:
            ttl = (f"{overlay['patient']} · {tp}\nΔmed={sh['median_diff']:.3g}, "
                   f"med={np.nanmedian(s):.3g}")
        else:
            ttl = (f"{overlay['patient']} · {tp}\nδ={sh['cliffs_delta']:.2f}, "
                   f"frac>q={sh['frac_above_ref_q']:.2f}")
        ax.set_title(ttl, fontsize=8)
        ax.set_xlabel(_axis_label(overlay.get('driver', ''), score_label), fontsize=8)
        ax.set_ylabel("density" if j == 0 else "")
        if j == 0:
            ax.legend(fontsize=7, frameon=False)
        _style(ax)
    if stats_note:
        fig.suptitle(stats_note, fontsize=9, y=1.03)
    fig.tight_layout()
    out = Path(outpath); fig.savefig(out, dpi=FIGURE_DPI, bbox_inches="tight"); plt.close(fig)
    return out


def timepoint_genotype_violin(
    split: pd.DataFrame,
    patient: str,
    driver: str,
    outpath: str | Path,
    score_label: str | None = None,
    driver_stats: dict | None = None,
    annotate_significance: bool = True,
    annotate_ns: bool = False,
    min_n_sig: int = 3,
    comparison_mode: str = "full",
    significance_stats: pd.DataFrame | None = None,
    stats_outpath: str | Path | None = None,
) -> Path:
    """Per-cell scores by timepoint, split by genotype (Mut/WT/Unknown).

    Adds Mann-Whitney significance brackets automatically. By default this tests
    both classes of comparisons that matter in mixed WT/Mut/Unknown panels:
      * all pairwise genotype/status contrasts within each timepoint;
      * all pairwise longitudinal timepoint contrasts within each genotype/status.
    The bracket stars use BH-FDR q-values computed across the panel's emitted
    comparisons. Non-significant brackets are hidden by default to avoid turning
    dense panels into spaghetti.
    """
    if split.empty:
        return None
    tps = sorted(split.timepoint.unique(), key=_timepoint_sort_key)
    labels_present = [g for g in ["WT", "Mut", "Unknown"] if g in split.genotype_label.unique()]
    if not labels_present:
        labels_present = sorted(split.genotype_label.dropna().astype(str).unique().tolist())

    fig, ax = plt.subplots(figsize=(1.6 * len(tps) + 2, 4.8))
    width = 0.8 / max(1, len(labels_present))
    group_positions: dict[tuple[str, str], float] = {}
    group_values: dict[tuple[str, str], np.ndarray] = {}

    for li, lab in enumerate(labels_present):
        positions, data = [], []
        for ti, tp in enumerate(tps):
            vals = _clean_scores(split[(split.timepoint == tp) &
                                       (split.genotype_label.astype(str) == str(lab))]["score"].values)
            if len(vals) >= 3:
                pos = ti + li * width - 0.4 + width / 2
                positions.append(pos)
                data.append(vals)
                group_positions[(str(tp), str(lab))] = pos
                group_values[(str(tp), str(lab))] = vals
        if data:
            parts = ax.violinplot(data, positions=positions, widths=width * 0.9,
                                  showmedians=True, showextrema=False)
            for pc in parts["bodies"]:
                pc.set_facecolor(_PALETTE.get(lab, "#999999")); pc.set_alpha(0.7)

    ax.set_xticks(range(len(tps))); ax.set_xticklabels(tps)
    score_label = score_label or split.attrs.get("score_label", f"{driver} score")
    ax.set_xlabel("Timepoint"); ax.set_ylabel(score_label, fontsize=9)
    _allv = _clean_scores(split["score"].values) if not split.empty else np.array([0, 1])
    if len(_allv) == 0:
        _allv = np.array([0, 1])
    _lo, _hi = np.nanmin(_allv), np.nanmax(_allv)
    if not np.isfinite(_lo) or not np.isfinite(_hi):
        _lo, _hi = 0.0, 1.0
    if _hi <= _lo:
        _hi = _lo + 1e-3
    _yr = _hi - _lo
    _pad = 0.06 * (_yr + 1e-9)
    ymin = max(-0.02, _lo - _pad) if _hi <= 1 and _lo >= 0 else _lo - _pad
    ymax = _hi + _pad

    sig_df = significance_stats
    if annotate_significance:
        if sig_df is None:
            sig_df = timepoint_genotype_violin_stats(
                split, min_n=min_n_sig, comparison_mode=comparison_mode)
        if stats_outpath is not None and sig_df is not None and not sig_df.empty:
            Path(stats_outpath).parent.mkdir(parents=True, exist_ok=True)
            sig_df.to_csv(stats_outpath, index=False)

        if sig_df is not None and not sig_df.empty:
            h = 0.025 * (_yr + 1e-9)
            step = 0.075 * (_yr + 1e-9)
            y0 = _hi + _pad

            # Map each statistical row back onto the actual violin x-positions.
            # We stack brackets on the lowest non-overlapping y-level, so dense
            # mixed panels remain readable instead of exploding upward one row at
            # a time.
            bracket_rows = []
            for _, row in sig_df.iterrows():
                label = row.get("significance") or _sig_stars(row.get("fdr_q", row.get("p_value")))
                if not label or (label == "ns" and not annotate_ns):
                    continue
                ctype = str(row.get("comparison_type", ""))
                x1 = x2 = None
                if ctype in {"within_timepoint_genotype", "within_timepoint_label_pairwise"}:
                    tp = str(row.get("timepoint"))
                    ga = str(row.get("genotype_label_a", row.get("group_a", "WT")))
                    gb = str(row.get("genotype_label_b", row.get("group_b", "Mut")))
                    x1 = group_positions.get((tp, ga))
                    x2 = group_positions.get((tp, gb))
                elif ctype in {"adjacent_timepoint_same_label", "between_timepoint_same_label_pairwise"}:
                    lab = str(row.get("label", "Unknown"))
                    x1 = group_positions.get((str(row.get("timepoint_a")), lab))
                    x2 = group_positions.get((str(row.get("timepoint_b")), lab))
                if x1 is None or x2 is None:
                    continue
                xa, xb = sorted([float(x1), float(x2)])
                q = _finite_stat(row.get("fdr_q", row.get("p_value")))
                span = xb - xa
                fam = str(row.get("comparison_family", ""))
                bracket_rows.append((span, 0 if fam == "status_within_timepoint" else 1, q if q is not None else np.inf, xa, xb, label))

            bracket_rows.sort(key=lambda z: (z[0], z[1], z[2]))
            levels: list[list[tuple[float, float]]] = []
            level_count = 0
            for _, _, _, xa, xb, label in bracket_rows:
                # Treat touching/near-touching intervals as overlapping; brackets
                # need a little horizontal breathing room around the vertical caps.
                placed = False
                for lev, intervals in enumerate(levels):
                    if all((xb + 0.04) < a or (xa - 0.04) > b for a, b in intervals):
                        intervals.append((xa, xb))
                        _add_sig_bracket(ax, xa, xb, y0 + lev * step, h, label)
                        level_count = max(level_count, lev + 1)
                        placed = True
                        break
                if not placed:
                    levels.append([(xa, xb)])
                    lev = len(levels) - 1
                    _add_sig_bracket(ax, xa, xb, y0 + lev * step, h, label)
                    level_count = max(level_count, lev + 1)
            if level_count:
                ymax = max(ymax, y0 + level_count * step + h + 0.02 * (_yr + 1e-9))

    ax.set_ylim(ymin, ymax)
    stats_note = _format_driver_stats(driver_stats or split.attrs.get("driver_stats"))
    title = f"{driver}: {patient} — driver-program scores by timepoint"
    if stats_note:
        title += f"\n{stats_note}"
    ax.set_title(title, fontsize=10)
    handles = [plt.Rectangle((0, 0), 1, 1, color=_PALETTE.get(l, "#999999"), alpha=0.7) for l in labels_present]
    ax.legend(handles, labels_present, title="Label", fontsize=8, frameon=False)
    if annotate_significance:
        ax.text(0.995, 0.01, "MWU; BH-FDR per panel; sig only", transform=ax.transAxes,
                ha="right", va="bottom", fontsize=6.5, color="#555555")
    _style(ax); fig.tight_layout()
    out = Path(outpath); fig.savefig(out, dpi=FIGURE_DPI, bbox_inches="tight"); plt.close(fig)
    return out

# ===========================================================================
# FIGURE 4 — subclonal maps (delegate to scope-bio viz)
# ===========================================================================
def _ensure_embedding(adata, latent_keys=("X_umap", "X_svd", "X_pca")):
    """Return (adata, key) with a 2-D embedding in obsm['X_umap'], computing one if
    needed. Tries, in order: an existing X_umap; UMAP from an existing latent
    (X_svd/X_pca); else PCA->UMAP from .X via scanpy. Self-contained so it works on
    scored objects that may not carry a latent embedding."""
    import numpy as np
    if "X_umap" in adata.obsm and adata.obsm["X_umap"].shape[1] >= 2:
        return adata, "X_umap"
    adata = adata.copy()
    try:
        import scanpy as sc
    except Exception:
        sc = None
    # UMAP from existing latent
    for k in ("X_svd", "X_pca"):
        if k in adata.obsm and sc is not None:
            try:
                sc.pp.neighbors(adata, use_rep=k, n_neighbors=15)
                sc.tl.umap(adata)
                return adata, "X_umap"
            except Exception:
                pass
    # Fresh PCA -> UMAP from .X
    if sc is not None:
        try:
            ad2 = adata.copy()
            sc.pp.pca(ad2, n_comps=min(50, ad2.n_vars - 1, ad2.n_obs - 1))
            sc.pp.neighbors(ad2, use_rep="X_pca", n_neighbors=15)
            sc.tl.umap(ad2)
            adata.obsm["X_umap"] = ad2.obsm["X_umap"]
            return adata, "X_umap"
        except Exception as e:
            print(f"[plots] UMAP computation failed ({e}); falling back to first 2 latent dims.")
    # last resort: first 2 dims of any latent
    for k in ("X_svd", "X_pca"):
        if k in adata.obsm:
            adata.obsm["X_umap"] = np.asarray(adata.obsm[k])[:, :2]
            return adata, "X_umap"
    raise KeyError("No embedding available and UMAP could not be computed.")


def umap_probability_panels(adata, drivers: list[str], outpath: str | Path,
                            cfg=None, max_panels: int = 12, recompute_umap: bool = False,
                            point_size: float | None = None,
                            confidence: dict | pd.Series | None = None,
                            panel_size: float = 4.3,
                            score_mode: str = "configured",
                            reference_mask_col: str = "is_reference",
                            grey_reference: bool = True,
                            prefer_guarded_for_reference_high: bool = True) -> Path:
    """Per-cell score overlays on a UMAP, one panel per driver.
    Self-contained (computes a UMAP if absent). Dots are small and adaptive so dense
    embeddings don't smear; each panel is annotated with the driver's confidence
    score (if provided) so "lighting up" is read against whether the signal is real.
    Panels are ordered by confidence (most trustworthy first) when confidence given."""
    if recompute_umap and "X_umap" in adata.obsm:
        del adata.obsm["X_umap"]
    adata, key = _ensure_embedding(adata)
    emb = np.asarray(adata.obsm[key])
    drivers = [d for d in drivers if _display_score_col(adata, d, cfg, score_mode=score_mode,
                                                        prefer_guarded_for_reference_high=prefer_guarded_for_reference_high) in adata.obs.columns]
    conf = dict(confidence) if confidence is not None else {}
    if conf:
        drivers = sorted(drivers, key=lambda d: conf.get(d, -1), reverse=True)
    drivers = drivers[:max_panels]
    if not drivers:
        return None
    # adaptive point size: smaller for more cells, so dense UMAPs stay legible
    if point_size is None:
        point_size = float(np.clip(9000.0 / max(adata.n_obs, 1), 0.6, 4.0))
    ncol = min(4, len(drivers)); nrow = int(np.ceil(len(drivers) / ncol))
    fig, axes = plt.subplots(nrow, ncol, figsize=(panel_size * ncol, panel_size * nrow),
                             squeeze=False)
    ref_mask = None
    if grey_reference and reference_mask_col in adata.obs.columns:
        ref_mask = adata.obs[reference_mask_col].astype(bool).values
    flagged = set((adata.uns.get("reference_high_drivers", []) or []))
    for i, d in enumerate(drivers):
        ax = axes[i // ncol][i % ncol]
        col = _display_score_col(adata, d, cfg, score_mode=score_mode,
                                 prefer_guarded_for_reference_high=prefer_guarded_for_reference_high)
        c = adata.obs[col].astype(float).values
        # Display scale should be informed by the claim-bearing compartment, not by
        # greyed reference cells that are intentionally de-emphasised.
        c_scale = c[(~ref_mask) & np.isfinite(c)] if ref_mask is not None and (~ref_mask).any() else c
        vmin, vmax, scale_note = _robust_limits(c_scale, cfg=cfg, col=col)
        if ref_mask is not None and ref_mask.any():
            ax.scatter(emb[ref_mask, 0], emb[ref_mask, 1], c="#e8e8e8", s=point_size,
                       edgecolor="none", rasterized=True, linewidths=0, zorder=1)
            idx = np.where(~ref_mask)[0]
            order = idx[np.argsort(np.nan_to_num(c[idx], nan=-np.inf))]
        else:
            order = np.argsort(np.nan_to_num(c, nan=-np.inf))  # high-score cells on top
        cmap = "coolwarm" if _score_kind_from_col(col, cfg) == "cellstate_residual_raw" else "magma"
        scat = ax.scatter(emb[order, 0], emb[order, 1], c=c[order], s=point_size,
                          cmap=cmap, vmin=vmin, vmax=vmax,
                          edgecolor="none", rasterized=True, linewidths=0, zorder=3)
        title = d
        if d in flagged and col.startswith(("mutation_resid_", "mutation_refpct_")) and score_mode == "configured":
            title += " (guarded display)"
        if scale_note:
            title += f"\n{scale_note}"
        if d in conf and np.isfinite(conf[d]):
            title = f"{title}   conf={conf[d]:.2f}"
        ax.set_title(title, fontsize=11); ax.set_xticks([]); ax.set_yticks([])
        for sp in ax.spines.values():
            sp.set_visible(False)
        cb = fig.colorbar(scat, ax=ax, fraction=0.046, pad=0.04)
        cb.ax.tick_params(labelsize=7)
        lbl = _score_label(d, col, cfg)
        if ref_mask is not None and ref_mask.any():
            lbl += " (reference greyed)"
        cb.set_label(lbl, fontsize=7)
    for j in range(len(drivers), nrow * ncol):
        axes[j // ncol][j % ncol].axis("off")
    mode_label = str(score_mode).replace("_", " ") if score_mode != "configured" else ("reference-percentile" if (cfg is not None and cfg.use_calibrated_scores) else "raw probability")
    sub = f"Per-cell {mode_label} score (UMAP)"
    if grey_reference and reference_mask_col in adata.obs.columns and adata.obs[reference_mask_col].astype(bool).any():
        sub += " — reference cells greyed"
    if prefer_guarded_for_reference_high and score_mode == "configured":
        sub += "; reference-high drivers auto-switch to guarded display"
    if conf:
        sub += "  —  ordered by confidence; conf<0.2 ≈ likely noise"
    fig.suptitle(sub, fontsize=12, y=1.002)
    fig.tight_layout()
    out = Path(outpath); fig.savefig(out, dpi=FIGURE_DPI, bbox_inches="tight"); plt.close(fig)
    return out


def _distinct_colors(n: int):
    """A long, perceptually distinct qualitative palette (>20 categories).
    Returns uniform RGBA 4-tuples — matplotlib's qualitative colormaps mix RGB
    3-tuples and RGBA 4-tuples, and concatenating them made np.array(...) ragged
    ('inhomogeneous shape'), which crashed categorical UMAP panels (e.g. PAAD)."""
    from matplotlib.colors import to_rgba
    base = []
    for name in ("tab20", "tab20b", "tab20c", "Set3", "Set2", "Dark2"):
        base.extend(list(plt.get_cmap(name).colors))
    if n <= len(base):
        chosen = base[:n]
    else:
        extra = plt.get_cmap("hsv")(np.linspace(0, 1, n - len(base), endpoint=False))
        chosen = base + [tuple(c) for c in extra]
    return [tuple(to_rgba(c)) for c in chosen]   # uniform 4-tuples


def _categorical_umap(ax, emb, vals, title, s, max_legend=24, seed=0):
    """One categorical UMAP panel: shuffled z-order (no category sits on top),
    distinct colours, and a multi-column legend that shows the most frequent
    categories with a '+N more' marker instead of silently dropping the legend.
    Boolean / 2-3 category panels (e.g. is_reference) get a high-contrast palette so
    True/False never render blue-on-blue."""
    vals = pd.Series(vals).astype(str).values
    counts = pd.Series(vals).value_counts()
    cats = list(counts.index)                       # frequency-ordered
    if len(cats) <= 3:
        # boolean-like: map by meaning where possible, else a distinct fallback trio
        colors = {}
        for i, c in enumerate(cats):
            key = str(c).strip().lower()
            colors[c] = _BOOL_COLORS.get(key, _TWO_CAT_FALLBACK[i % len(_TWO_CAT_FALLBACK)])
    else:
        colors = {c: col for c, col in zip(cats, _distinct_colors(len(cats)))}
    cvec = np.array([colors[v] for v in vals])
    order = np.random.default_rng(seed).permutation(len(emb))  # kill overplot bias
    ax.scatter(emb[order, 0], emb[order, 1], c=cvec[order], s=s, edgecolor="none",
               rasterized=True, linewidths=0)
    ax.set_title(f"{title}  (n={len(cats)})", fontsize=11)
    ax.set_xticks([]); ax.set_yticks([])
    for sp in ax.spines.values():
        sp.set_edgecolor("#cccccc")
    show = cats[:max_legend]
    handles = [Line2D([0], [0], marker="o", color="w", markerfacecolor=colors[c],
                      markersize=6, label=(c if len(c) <= 26 else c[:25] + "…")) for c in show]
    if len(cats) > max_legend:
        handles.append(Line2D([0], [0], marker="o", color="w", markerfacecolor="#bbbbbb",
                              markersize=6, label=f"+{len(cats)-max_legend} more"))
    ncol = 1 if len(handles) <= 14 else 2
    ax.legend(handles=handles, fontsize=6.5, loc="center left", bbox_to_anchor=(1.0, 0.5),
              frameon=False, handletextpad=0.25, labelspacing=0.25, ncol=ncol,
              columnspacing=0.8, borderaxespad=0.0)


def umap_context_panel(adata, outpath: str | Path, celltype_col: str = "CellType",
                       reference_mask_col: str = "is_reference",
                       extra_cols: list[str] | None = None, cfg=None,
                       annotate_if_missing: bool = True) -> Path:
    """UMAP context panels (cell type · reference flag · patient). High quality:
    adaptive point size, shuffled z-order so no patient/lineage sits on top, distinct
    colours for many categories, and legends that never get silently dropped. If the
    cell-type column is missing/uninformative and annotate_if_missing, a marker-based
    coarse annotation is computed so the panel always has a meaningful legend."""
    adata, key = _ensure_embedding(adata)
    ct_col = celltype_col
    if annotate_if_missing:
        try:
            from .annotate import ensure_celltype
            adata, ct_col = ensure_celltype(adata, celltype_col)
        except Exception as e:
            print(f"[plots] marker annotation skipped ({e}).")
    emb = np.asarray(adata.obsm[key])
    s = float(np.clip(6000.0 / max(adata.n_obs, 1), 0.35, 3.0))

    panels = []
    if ct_col in adata.obs.columns:
        panels.append((ct_col, adata.obs[ct_col].values))
    if reference_mask_col in adata.obs.columns:
        _ref_title = {"is_normal_display": "normal (greyed on score maps)",
                      "is_reference": "is_reference"}.get(reference_mask_col, reference_mask_col)
        panels.append((_ref_title, adata.obs[reference_mask_col].astype(str).values))
    for c in (extra_cols or ["patient"]):
        if c in adata.obs.columns and c != ct_col:
            if str(c).lower() == "patient":
                try:
                    from .data import is_usable_patient_series, patient_series_diagnostics
                    if not is_usable_patient_series(adata.obs[c], colname=c):
                        d = patient_series_diagnostics(adata.obs[c], colname=c)
                        print(f"[plots] skipping patient UMAP panel: '{c}' looks like cell IDs "
                              f"(n_unique={d['n_unique']}, median_cells/id={d['median_cells_per_id']:.1f}).")
                        continue
                except Exception:
                    pass
            panels.append((c, adata.obs[c].values))
    if not panels:
        return None
    ncol = len(panels)
    fig, axes = plt.subplots(1, ncol, figsize=(5.6 * ncol, 4.8), squeeze=False)
    for i, (title, vals) in enumerate(panels):
        _categorical_umap(axes[0][i], emb, vals, title, s, seed=i)
    fig.tight_layout()
    out = Path(outpath); fig.savefig(out, dpi=FIGURE_DPI, bbox_inches="tight"); plt.close(fig)
    return out


def percell_summary_heatmap(percell_table: pd.DataFrame, outpath: str | Path) -> Path:
    """Compact Fig-3 summary across ALL genotyped drivers: apparent AUROC and
    Cliff's delta per driver, sorted, with significance stars. One figure that
    captures the whole single-cell validation instead of many separate violins."""
    df = percell_table.dropna(subset=["apparent_auroc"]).copy()
    if df.empty:
        return None
    df = df.sort_values("cliffs_delta")
    n = len(df)
    fig, ax = plt.subplots(figsize=(5.5, max(3, 0.34 * n)))
    colors = ["#1b9e77" if v > 0 else "#d73027" for v in df["cliffs_delta"]]
    ax.barh(range(n), df["cliffs_delta"], color=colors, alpha=0.85)
    for i, (_, r) in enumerate(df.iterrows()):
        q = r.get("fdr_q", np.nan)
        star = "***" if q < 1e-3 else "**" if q < 1e-2 else "*" if q < FIG3_FDR_ALPHA else ""
        ax.text(r["cliffs_delta"] + (0.01 if r["cliffs_delta"] >= 0 else -0.01), i,
                f"{star}  AUC={r['apparent_auroc']:.2f} (n={int(r['n_mut'])}/{int(r['n_wt'])})",
                va="center", ha="left" if r["cliffs_delta"] >= 0 else "right", fontsize=7)
    ax.axvline(0, color="grey", lw=1)
    ax.set_yticks(range(n)); ax.set_yticklabels(df["driver"], fontsize=8)
    ax.set_xlabel("Cliff's δ  (Mut vs WT, per-cell; positive = mutant scores higher)")
    ax.set_title("Single-cell Mut-vs-WT enrichment by driver\n(* q<0.10, ** q<0.01, *** q<0.001)", fontsize=10)
    lim = max(0.1, np.abs(df["cliffs_delta"]).max() * 1.5)
    ax.set_xlim(-lim, lim)
    _style(ax); fig.tight_layout()
    out = Path(outpath); fig.savefig(out, dpi=FIGURE_DPI, bbox_inches="tight"); plt.close(fig)
    return out


def cluster_probability_heatmap(adata, drivers: list[str], cluster_key: str,
                                outpath: str | Path, cfg=None,
                                score_mode: str = "configured",
                                confidence: dict | pd.Series | None = None,
                                min_confidence: float | None = None,
                                cluster: bool = True) -> Path:
    """Mean score per cell-type/cluster heatmap (drivers x clusters).

    To cut noise, drivers below ``min_confidence`` are dropped (keeps the panel to the
    trustworthy signal). When ``cluster`` is True, rows and columns are hierarchically
    ordered (a clustermap) so co-varying drivers/clusters sit together and patterns
    are visible instead of an arbitrary alphabetical grid. The displayed matrix is
    per-driver robust-scaled; group means come from the raw selected score columns.
    """
    if cluster_key not in adata.obs.columns:
        return None
    conf = dict(confidence) if confidence is not None else {}
    if min_confidence is not None and conf:
        kept = [d for d in drivers if conf.get(d, np.inf) >= min_confidence]
        if kept:
            drivers = kept
    cols = {d: _select_score_col(adata, d, cfg, score_mode=score_mode) for d in drivers}
    cols = {d: c for d, c in cols.items() if c in adata.obs.columns}
    if not cols:
        return None
    M = (adata.obs.assign(**{d: adata.obs[c].astype(float) for d, c in cols.items()})
         .groupby(cluster_key)[list(cols.keys())].mean())
    display = M.copy()
    scale_note = ""
    hmode = str(getattr(cfg, "heatmap_score_scale", "global_robust") if cfg is not None else "global_robust").lower()
    kinds = [_score_kind_from_col(c, cfg) for c in cols.values()]
    residual_heatmap = bool(kinds) and all(k == "cellstate_residual_raw" for k in kinds)
    cmap = "viridis"
    vmin, vmax = None, None
    if hmode == "driver_robust":
        lq = float(getattr(cfg, "plot_robust_lower_q", 0.01)) if cfg is not None else 0.01
        uq = float(getattr(cfg, "plot_robust_upper_q", 0.99)) if cfg is not None else 0.99
        for d in display.columns:
            display[d] = _robust_rescale_column(display[d].values, lq, uq)
        vmin, vmax = 0.0, 1.0
        scale_note = " (per-driver robust display; exploratory)"
    elif residual_heatmap:
        vals = display.values.astype(float)
        vals = vals[np.isfinite(vals)]
        min_abs = float(getattr(cfg, "plot_residual_min_abs_vmax", 0.05)) if cfg is not None else 0.05
        max_abs = float(getattr(cfg, "heatmap_residual_abs_vmax", 0.20)) if cfg is not None else 0.20
        if vals.size:
            if hmode == "absolute":
                lim = float(np.nanmax(np.abs(vals)))
            else:
                uq = float(getattr(cfg, "plot_robust_upper_q", 0.99)) if cfg is not None else 0.99
                lim = float(np.nanquantile(np.abs(vals), uq))
            if not np.isfinite(lim) or lim <= 0:
                lim = min_abs
            lim = min(max(lim, min_abs), max_abs)
        else:
            lim = min_abs
        vmin, vmax = -lim, lim
        cmap = "coolwarm"
        scale_note = f" (global centered ±{lim:g}; comparable)"
    elif hmode == "global_robust":
        vals = display.values.astype(float)
        vals = vals[np.isfinite(vals)]
        if vals.size:
            lq = float(getattr(cfg, "plot_robust_lower_q", 0.01)) if cfg is not None else 0.01
            uq = float(getattr(cfg, "plot_robust_upper_q", 0.99)) if cfg is not None else 0.99
            vmin, vmax = np.nanquantile(vals, [lq, uq])
            if not np.isfinite(vmin) or not np.isfinite(vmax) or vmax <= vmin:
                vmin, vmax = float(np.nanmin(vals)), float(np.nanmax(vals))
            scale_note = f" (global q{int(lq*100):02d}-q{int(uq*100):02d}; comparable)"
    else:
        vals = display.values.astype(float)
        vals = vals[np.isfinite(vals)]
        if vals.size:
            vmin, vmax = float(np.nanmin(vals)), float(np.nanmax(vals))
        scale_note = " (absolute scale)"

    row_labels = list(M.index)
    col_labels = list(cols.keys())
    Dv = display.values
    # hierarchical ordering (clustermap) when there is enough to cluster
    if cluster and Dv.shape[0] >= 3 and Dv.shape[1] >= 3:
        try:
            from scipy.cluster.hierarchy import linkage, leaves_list
            from scipy.spatial.distance import pdist
            clean = np.nan_to_num(Dv, nan=0.0)
            r_order = leaves_list(linkage(pdist(clean, "euclidean"), "average"))
            c_order = leaves_list(linkage(pdist(clean.T, "euclidean"), "average"))
            Dv = clean[np.ix_(r_order, c_order)]
            row_labels = [row_labels[i] for i in r_order]
            col_labels = [col_labels[i] for i in c_order]
            scale_note += " · clustered"
        except Exception as e:
            print(f"[plots] heatmap clustering skipped ({e}).")

    fig, ax = plt.subplots(figsize=(0.5 * len(col_labels) + 3, 0.4 * len(row_labels) + 2))
    im = ax.imshow(Dv, aspect="auto", cmap=cmap, vmin=vmin, vmax=vmax)
    ax.set_xticks(range(len(col_labels))); ax.set_xticklabels(col_labels, rotation=90, fontsize=7)
    ax.set_yticks(range(len(row_labels))); ax.set_yticklabels(row_labels, fontsize=7)
    mode = str(score_mode).replace("_", " ") if score_mode != "configured" else ("reference-percentile" if (cfg is not None and cfg.use_calibrated_scores) else "raw probability")
    thr_note = f"  [conf≥{min_confidence}]" if (min_confidence is not None and conf) else ""
    ax.set_title(f"Mean {mode} score by {cluster_key}{scale_note}{thr_note}", fontsize=10)
    fig.colorbar(im, ax=ax, fraction=0.046, pad=0.04)
    fig.tight_layout()
    out = Path(outpath); fig.savefig(out, dpi=FIGURE_DPI, bbox_inches="tight"); plt.close(fig)
    return out


# ===========================================================================
# FIGURE 6 — CNV concordance / discordance
# ===========================================================================
def cnv_discordance_bar(discordance: pd.DataFrame, outpath: str | Path,
                        ari: float | None = None) -> Path:
    """Bar chart of the four scOPE-vs-CNV agreement quadrants (Fig. 6)."""
    fig, ax = plt.subplots(figsize=(6.5, 4))
    colors = ["#4575b4", "#fc8d59", "#fc8d59", "#bdbdbd"]
    ax.bar(range(len(discordance)), discordance["n_cells"], color=colors)
    ax.set_xticks(range(len(discordance)))
    ax.set_xticklabels(discordance["quadrant"], rotation=25, ha="right", fontsize=7)
    ax.set_ylabel("cells")
    ax.set_title(f"scOPE vs CNV agreement" + (f" (ARI={ari:.3f})" if ari is not None else ""),
                 fontsize=10)
    _style(ax); fig.tight_layout()
    out = Path(outpath); fig.savefig(out, dpi=FIGURE_DPI, bbox_inches="tight"); plt.close(fig)
    return out


def cnv_confusion_heatmap(conc: dict, outpath: str | Path) -> Path:
    """2x2 confusion heatmap of scOPE vs CNV malignant calls with counts + ARI."""
    if conc.get("n", 0) == 0:
        return None
    M = np.array([[conc["cnv_normal_scope_normal"], conc["cnv_normal_scope_malignant"]],
                  [conc["cnv_malignant_scope_normal"], conc["cnv_malignant_scope_malignant"]]])
    fig, ax = plt.subplots(figsize=(4.2, 3.8))
    im = ax.imshow(M, cmap="Blues")
    for i in range(2):
        for j in range(2):
            ax.text(j, i, f"{M[i, j]:,}", ha="center", va="center",
                    color="white" if M[i, j] > M.max() / 2 else "black", fontsize=12)
    ax.set_xticks([0, 1]); ax.set_xticklabels(["scOPE normal", "scOPE malignant"], fontsize=8)
    ax.set_yticks([0, 1]); ax.set_yticklabels(["CNV normal", "CNV malignant"], fontsize=8)
    ax.set_title(f"scOPE vs CNV calls (ARI={conc['ari']:.3f}, n={conc['n']:,})", fontsize=10)
    fig.colorbar(im, ax=ax, fraction=0.046, pad=0.04)
    fig.tight_layout()
    out = Path(outpath); fig.savefig(out, dpi=FIGURE_DPI, bbox_inches="tight"); plt.close(fig)
    return out


def cnv_malignant_fraction_scatter(frac_table: pd.DataFrame, outpath: str | Path) -> Path:
    """Per-patient scOPE vs CNV malignant fraction scatter (agreement at the
    patient level), with the identity line and near-diploid region shaded."""
    if frac_table.empty:
        return None
    fig, ax = plt.subplots(figsize=(4.8, 4.6))
    # near-diploid is defined by CNV fraction < 10%, which is the X axis -> a VERTICAL
    # band. (Previously this was a horizontal axhspan on the scOPE axis, which put the
    # 'CNV<10%' shading on the wrong axis and made the axes look mislabeled.)
    ax.axvspan(0, 0.10, color="#fff3cd", alpha=0.6, zorder=0, label="near-diploid (CNV<10%)")
    ax.plot([0, 1], [0, 1], ls="--", color="grey", lw=1)
    sizes = 20 + 80 * (frac_table["n_cells"] / frac_table["n_cells"].max())
    ax.scatter(frac_table["cnv_malignant_frac"], frac_table["scope_malignant_frac"],
               s=sizes, alpha=0.75, color="#2c7fb8", edgecolor="k", linewidth=0.4)
    metric_cols = {"n_cells", "scope_malignant_frac", "cnv_malignant_frac", "mean_cnv_score"}
    patient_label_col = next((c for c in frac_table.columns if c not in metric_cols), frac_table.columns[0])
    if frac_table.shape[0] <= 80:
        for _, r in frac_table.iterrows():
            ax.annotate(str(r[patient_label_col]),
                        (r["cnv_malignant_frac"], r["scope_malignant_frac"]),
                        xytext=(3, 3), textcoords="offset points",
                        fontsize=7 if frac_table.shape[0] <= 30 else 5.5,
                        alpha=0.85, clip_on=True)
    else:
        ax.text(0.02, 0.98, f"{frac_table.shape[0]} patients (labels suppressed)",
                transform=ax.transAxes, va="top", ha="left", fontsize=7, color="0.35")
    ax.set_xlabel("CNV malignant fraction  (per patient)")
    ax.set_ylabel("scOPE malignant fraction  (per patient)")
    ax.set_xlim(-0.02, 1.02); ax.set_ylim(-0.02, 1.02)
    ax.set_title("Per-patient malignant fraction: scOPE vs CNV", fontsize=10)
    ax.legend(fontsize=7, frameon=False, loc="lower right")
    _style(ax); fig.tight_layout()
    out = Path(outpath); fig.savefig(out, dpi=FIGURE_DPI, bbox_inches="tight"); plt.close(fig)
    return out


def cnv_score_scatter(merged: pd.DataFrame, outpath: str | Path, rho: float | None = None) -> Path:
    """Continuous scOPE max-driver score vs CNV burden score, coloured by the
    discordance quadrant, to show where the two malignancy signals agree/disagree."""
    if merged is None or merged.empty or "scope_max_driver_score" not in merged.columns:
        return None
    sub = merged.dropna(subset=["scope_max_driver_score", "cnv_score"])
    if sub.empty:
        return None
    fig, ax = plt.subplots(figsize=(5, 4.6))
    sc = sub["scope_malignant"].astype(int).values if "scope_malignant" in sub else np.zeros(len(sub), int)
    cn = sub["malignant_call"].astype(int).values
    quad = np.where((sc == 1) & (cn == 1), "both",
            np.where((sc == 1) & (cn == 0), "scope+ / CNV-",
            np.where((sc == 0) & (cn == 1), "scope- / CNV+", "neither")))
    cmap = {"both": "#7570b3", "scope+ / CNV-": "#1b9e77", "scope- / CNV+": "#d95f02", "neither": "#bdbdbd"}
    for q, c in cmap.items():
        m = quad == q
        if m.any():
            ax.scatter(sub["cnv_score"].values[m], sub["scope_max_driver_score"].values[m],
                       s=6, alpha=0.4, color=c, label=q, edgecolor="none", rasterized=True)
    ax.set_xlabel("CNV burden score"); ax.set_ylabel("scOPE max-driver score")
    title = "scOPE score vs CNV burden (per cell)"
    if rho is not None:
        title += f"\nSpearman ρ = {rho:.2f}"
    ax.set_title(title, fontsize=10)
    ax.legend(fontsize=6, frameon=False, markerscale=2, loc="best")
    _style(ax); fig.tight_layout()
    out = Path(outpath); fig.savefig(out, dpi=FIGURE_DPI, bbox_inches="tight"); plt.close(fig)
    return out


def reference_direction_bar(diag_table: pd.DataFrame, outpath: str | Path) -> Path:
    """Diagnostic: per-driver tumour-vs-reference vs Mut-vs-WT direction. Bars show
    the two Cliff's deltas side by side; drivers where they disagree (lineage
    confound) are flagged. Makes the reference-lineage issue legible at a glance."""
    df = diag_table.dropna(subset=["mut_vs_wt_cliffs"]).copy()
    if df.empty:
        return None
    df = df.sort_values("mut_vs_wt_cliffs")
    n = len(df); y = np.arange(n); h = 0.4
    fig, ax = plt.subplots(figsize=(6, max(3, 0.4 * n)))
    ax.barh(y + h/2, df["mut_vs_wt_cliffs"], height=h, color="#1b9e77", label="Mut vs WT (per-cell)")
    ax.barh(y - h/2, df["tumour_vs_ref_cliffs"], height=h, color="#fc8d59", label="Tumour vs reference")
    for i, (_, r) in enumerate(df.iterrows()):
        if r.get("lineage_confound_flag", False):
            ax.text(0, i, "  ⚠ disagree", va="center", ha="left", fontsize=7, color="firebrick")
    ax.axvline(0, color="grey", lw=1)
    ax.set_yticks(y); ax.set_yticklabels(df["driver"], fontsize=8)
    ax.set_xlabel("Cliff's δ"); ax.set_title("Direction check: Mut-vs-WT vs Tumour-vs-reference\n"
                                             "(disagreement ⇒ reference-lineage confound)", fontsize=10)
    ax.legend(fontsize=7, frameon=False, loc="lower right")
    _style(ax); fig.tight_layout()
    out = Path(outpath); fig.savefig(out, dpi=FIGURE_DPI, bbox_inches="tight"); plt.close(fig)
    return out


def confidence_bar(conf_table: pd.DataFrame, outpath: str | Path, title_suffix: str = "") -> Path:
    """Ranked per-driver confidence with the component breakdown (T/S/C/X) shown as a
    small stacked strip, so you can see WHY each driver scored as it did. Drivers are
    sorted by composite confidence; a guide line marks the low-confidence cutoff."""
    df = conf_table.dropna(subset=["confidence"]).sort_values("confidence").copy()
    if df.empty:
        return None
    n = len(df)
    fig, (axc, axb) = plt.subplots(1, 2, figsize=(11, max(3, 0.34 * n)),
                                   gridspec_kw={"width_ratios": [2.4, 2]})
    # left: composite confidence bars, coloured by tier
    tier_col = {"very_high": "#1a9850", "high": "#91cf60", "moderate": "#fee08b", "low": "#d73027"}
    colors = [tier_col.get(str(t), "#999999") for t in df.get("tier", ["low"] * n)]
    axc.barh(range(n), df["confidence"], color=colors, alpha=0.9)
    axc.axvline(0.2, color="grey", ls="--", lw=1)
    axc.set_yticks(range(n)); axc.set_yticklabels(df["driver"], fontsize=8)
    axc.set_xlim(0, 1); axc.set_xlabel("confidence  (ground-truth-free)")
    axc.set_title("Composite confidence" + title_suffix, fontsize=11, loc="left")
    _style(axc)
    # right: component heatmap T/S/C/X
    comp_cols = ["T_transferability", "S_spatial", "C_concentration", "X_crossmodal"]
    present = [c for c in comp_cols if c in df.columns]
    M = df[present].astype(float).values
    im = axb.imshow(M, aspect="auto", cmap="viridis", vmin=0, vmax=1)
    axb.set_yticks(range(n)); axb.set_yticklabels([""] * n)
    axb.set_xticks(range(len(present)))
    axb.set_xticklabels([c.split("_")[0] for c in present])
    axb.set_title("Components  (T=transfer S=spatial C=concentr. X=cross-modal)", fontsize=9, loc="left")
    fig.colorbar(im, ax=axb, fraction=0.046, pad=0.04)
    fig.tight_layout()
    out = Path(outpath); fig.savefig(out, dpi=FIGURE_DPI, bbox_inches="tight"); plt.close(fig)
    return out


def confidence_vs_truth(validation: dict, outpath: str | Path) -> Path:
    """Validation scatter: ground-truth-free confidence (x) vs the van Galen per-cell
    Mut-vs-WT effect (y), where genotype exists. Points coloured by whether the driver
    is a 'real' (FDR-significant, positive) signal. Confirms the score ranks the real
    drivers high WITHOUT having seen the ground truth."""
    m = validation.get("merged")
    if m is None or m.empty or "is_real" not in m.columns:
        return None
    fig, ax = plt.subplots(figsize=(5.6, 4.8))
    fdr_alpha = float(validation.get("fdr_alpha", FIG3_FDR_ALPHA))
    for is_real, col, lab in [(1, "#1a9850", f"real (FDR q<{fdr_alpha:g}, Mut>WT)"),
                              (0, "#d73027", f"not FDR-significant Mut>WT at q<{fdr_alpha:g}")]:
        sub = m[m["is_real"] == is_real]
        ax.scatter(sub["confidence"], sub["truth_cliffs"], c=col, s=60,
                   edgecolor="k", linewidth=0.4, label=lab, zorder=3)
    for _, r in m.iterrows():
        ax.annotate(r["driver"], (r["confidence"], r["truth_cliffs"]),
                    fontsize=7, alpha=0.75, xytext=(3, 2), textcoords="offset points")
    ax.axhline(0, color="grey", lw=1); ax.axvline(0.2, color="grey", ls="--", lw=1)
    rho = validation.get("spearman_rho"); auc = validation.get("auroc_real_vs_conf")
    sub = f"Spearman ρ={rho:.2f}" if rho is not None else ""
    if auc is not None and np.isfinite(auc):
        sub += f"   ·   AUROC(real | confidence)={auc:.2f}"
    ax.set_xlabel("ground-truth-free confidence"); ax.set_ylabel("per-cell Mut-vs-WT Cliff's δ (truth)")
    ax.set_title("Confidence compared with van Galen direct labels\n" + sub +
                 f"\nReal label uses FDR q<{fdr_alpha:g} and positive Cliff's δ", fontsize=10)
    ax.legend(fontsize=8, frameon=False, loc="lower right")
    _style(ax); fig.tight_layout()
    out = Path(outpath); fig.savefig(out, dpi=FIGURE_DPI, bbox_inches="tight"); plt.close(fig)
    return out
```
