## Supplemental_Code for "scOPE identifies which driver-associated expression programs transfer from bulk tumors to single cells": priors.py

```
"""
scope_analysis.priors
=====================
Cancer-type driver-mutation PRIORS, used only when a cohort has no per-patient
genotype for a scored driver (e.g. BRCA's patient_status carries only ERBB2 while the
scored drivers are PIK3CA/TP53/GATA3/...; GBM and PAAD patient_status are empty).

IMPORTANT — what this is and is NOT:
  * This is a population-level prior: "how often is gene X mutated in cancer type Y",
    from well-established large cohorts (TCGA / COSMIC / cBioPortal, approximate and
    rounded). It tells you which mutations to *expect* in a cancer type.
  * It is NOT a per-patient genotype. It must never be used to claim that a specific
    patient or cell carries a mutation. In figures it is shown as a clearly-labelled
    expectation ("expected in BRCA ~moderate"), and the genotype panel cells stay
    'unknown' (grey) — we do not fabricate Mut/WT calls from a prevalence.

Frequencies are coarse on purpose (tiers + an approximate %), so nobody over-reads a
spuriously precise number. Use `prevalence_tier()` for the qualitative label.

Call hierarchy
--------------
    umap_panels.driver_triptych_grid -> expected_label  (genotype panel annotation)
    drivers/run_all                  -> expected_drivers_table  (Expected_drivers_*.csv)
"""
from __future__ import annotations

import re
import pandas as pd

# Approximate mutation prevalence (fraction of patients) from large public cohorts.
# Rounded; meant as a prior, not a precise estimate. Amplification-driven events
# (e.g. ERBB2 in BRCA, EGFR in GBM) are included where they are the canonical driver.
CANCER_DRIVER_PRIORS: dict[str, dict[str, float]] = {
    "AML": {  # van Galen / TCGA-LAML driver landscape
        "NPM1": 0.30, "FLT3": 0.30, "DNMT3A": 0.25, "IDH2": 0.12, "IDH1": 0.08,
        "NRAS": 0.10, "RUNX1": 0.10, "TET2": 0.10, "TP53": 0.08, "CEBPA": 0.10,
        "WT1": 0.08, "KIT": 0.05, "PTPN11": 0.05, "KRAS": 0.05, "ASXL1": 0.08,
        "STAG2": 0.06, "SRSF2": 0.06, "U2AF1": 0.04, "SMC1A": 0.02, "SMC3": 0.02,
        "RAD21": 0.03, "BCOR": 0.05, "SETD2": 0.03, "PHF6": 0.03, "EZH2": 0.03,
        "KMT2A": 0.05, "GATA2": 0.03, "SF3B1": 0.04, "ETV6": 0.03,
    },
    "BRCA": {  # TCGA-BRCA
        "PIK3CA": 0.35, "TP53": 0.30, "GATA3": 0.10, "CDH1": 0.10, "MAP3K1": 0.08,
        "ERBB2": 0.15, "PTEN": 0.05, "AKT1": 0.03, "MAP2K4": 0.03,
    },
    "CRC": {  # TCGA-COADREAD
        "APC": 0.75, "TP53": 0.55, "KRAS": 0.40, "PIK3CA": 0.18, "SMAD4": 0.10,
        "FBXW7": 0.10, "BRAF": 0.10, "NRAS": 0.05, "CTNNB1": 0.05, "TCF7L2": 0.05,
        "ARID1A": 0.08, "SOX9": 0.05, "FAT4": 0.05, "RNF43": 0.08, "PTEN": 0.05,
    },
    "GBM": {  # TCGA-GBM (primary)
        "EGFR": 0.40, "TP53": 0.30, "PTEN": 0.30, "NF1": 0.10, "PIK3CA": 0.10,
        "RB1": 0.08, "IDH1": 0.05, "PDGFRA": 0.10, "ATRX": 0.07,
    },
    "LUAD": {  # TCGA-LUAD
        "TP53": 0.45, "KRAS": 0.30, "EGFR": 0.15, "STK11": 0.15, "KEAP1": 0.15,
        "NF1": 0.10, "BRAF": 0.07, "PIK3CA": 0.07, "MET": 0.05,
    },
    "PAAD": {  # TCGA-PAAD
        "KRAS": 0.90, "TP53": 0.70, "CDKN2A": 0.50, "SMAD4": 0.25, "ARID1A": 0.06,
    },
    "SKCM": {  # TCGA-SKCM (cutaneous)
        "BRAF": 0.50, "NRAS": 0.25, "NF1": 0.15, "TP53": 0.15, "CDKN2A": 0.15,
        "PTEN": 0.10, "KIT": 0.03,
    },
}

# Cohort-key prefix -> cancer type (cohort keys look like 'BRCA_Wu', 'AML_vanGalen').
_KEY_TO_CANCER = {
    "AML": "AML", "BRCA": "BRCA", "CRC": "CRC", "COAD": "CRC", "READ": "CRC",
    "GBM": "GBM", "LUAD": "LUAD", "NSCLC": "LUAD", "PAAD": "PAAD", "PDAC": "PAAD",
    "SKCM": "SKCM", "MEL": "SKCM",
}


def cancer_type_for_cohort(cohort_key: str, explicit: str | None = None) -> str | None:
    """Resolve a cancer type from an explicit config value or the cohort key prefix."""
    if explicit:
        return str(explicit).upper()
    if not cohort_key:
        return None
    head = re.split(r"[_\-\s]", str(cohort_key))[0].upper()
    return _KEY_TO_CANCER.get(head)


def prevalence_tier(freq: float | None) -> str:
    if freq is None:
        return "not characteristic"
    if freq >= 0.30:
        return "frequent"
    if freq >= 0.10:
        return "moderate"
    if freq > 0.0:
        return "rare"
    return "not characteristic"


def driver_prior(cancer_type: str | None, driver: str) -> float | None:
    """Approximate prevalence of `driver` in `cancer_type`, or None if not characteristic."""
    if not cancer_type:
        return None
    return CANCER_DRIVER_PRIORS.get(str(cancer_type).upper(), {}).get(str(driver).upper())


def expected_label(cancer_type: str | None, driver: str) -> str:
    """Short human-readable expectation for the triptych genotype-panel annotation when
    no per-patient genotype exists. Never asserts a per-patient call."""
    freq = driver_prior(cancer_type, driver)
    ct = (cancer_type or "this cancer").upper()
    if freq is None:
        return f"no per-patient genotype · not a characteristic {ct} driver (prior)"
    return f"no per-patient genotype · {prevalence_tier(freq)} in {ct} ~{int(round(freq*100))}% (cancer-type prior)"


def expected_drivers_table(cancer_type: str | None, drivers: list[str]) -> pd.DataFrame:
    """One row per scored driver: its cancer-type prior tier/prevalence. Saved as
    Expected_drivers_<cohort>.csv so the expectation is explicit and auditable."""
    rows = []
    for d in drivers:
        f = driver_prior(cancer_type, d)
        rows.append({
            "driver": d,
            "cancer_type": (cancer_type or "").upper(),
            "expected": f is not None,
            "prior_tier": prevalence_tier(f),
            "approx_prevalence": (round(f, 3) if f is not None else 0.0),
            "source": "literature/TCGA prior (population-level, not per-patient)",
        })
    return pd.DataFrame(rows).sort_values("approx_prevalence", ascending=False).reset_index(drop=True)
```
