## Supplemental_Code for "scOPE identifies which driver-associated expression programs transfer from bulk tumors to single cells": reference.py

```
"""
scope_analysis.reference
========================
Reference-set and cell-state guardrails for transferred mutation scores.

Why this exists
---------------
Some AML drivers (especially NPM1) look like developmental / myeloid-progenitor
programmes rather than clean tumour-vs-healthy markers. A global healthy reference
can therefore sit *above* AML cells even when Mut > WT within AML. This module
makes that failure mode explicit and provides safer same-cell-state residual scores.
"""
from __future__ import annotations

import re
import numpy as np
import pandas as pd


LYMPHOID_STATES = {"lymphoid_t", "lymphoid_b", "lymphoid_nk", "plasma"}
MYELOID_STATES = {"myeloid", "dendritic"}
HSPC_STATES = {"hspc_prog"}
ERY_STATES = {"erythroid"}


def _norm(x) -> str:
    return re.sub(r"[^a-z0-9]+", "", str(x).strip().lower())


def coarse_state_one(x) -> str:
    """Map heterogeneous labels to coarse lineage states.

    Handles van Galen labels (HSC-like, Prog-like, Mono-like, GMP-like, cDC-like),
    marker fallback labels (T cell, Myeloid, HSPC/Progenitor), and common variants.
    """
    s0 = str(x).strip()
    s = _norm(s0)
    if not s or s in {"nan", "none", "unknown", "unassigned"}:
        return "unknown"
    if s in {"t", "tcell", "tcells", "cd4t", "cd8t"} or "ctl" in s or s.startswith("cd3"):
        return "lymphoid_t"
    if s in {"nk", "nkcell", "nkcells"} or "natural" in s:
        return "lymphoid_nk"
    if s in {"b", "bcell", "bcells", "prob", "probt"} or s.startswith("prob"):
        return "lymphoid_b"
    if "plasma" in s:
        return "plasma"
    if "ery" in s or "eryth" in s:
        return "erythroid"
    if "dendritic" in s or s in {"cdc", "cdclike", "pdc", "pdclike", "dc"}:
        return "dendritic"
    if "mono" in s or "myeloid" in s or s in {"monocyte", "macrophage", "mac", "promono", "promonolike"}:
        return "myeloid"
    if s in {"hsc", "hsclike", "prog", "proglike", "gmp", "gmplike", "hspc", "hspcprogenitor", "progenitor", "stem"}:
        return "hspc_prog"
    if "endothelial" in s or s in {"endo"}:
        return "endothelial"
    if "fibro" in s or "caf" in s or "strom" in s:
        return "stromal"
    if "epi" in s or "krt" in s:
        return "epithelial"
    # fallbacks for exact van Galen labels that normalize oddly
    low = s0.lower()
    if any(k in low for k in ["hsc", "prog", "gmp"]):
        return "hspc_prog"
    if any(k in low for k in ["mono", "myeloid"]):
        return "myeloid"
    return "other"


def ensure_coarse_cellstate(adata, celltype_col: str = "CellType", out_col: str = "celltype_coarse"):
    """Write a coarse cell-state column to adata.obs and return (adata, out_col)."""
    a = adata.copy()
    src = celltype_col if celltype_col in a.obs.columns else None
    if src is None:
        try:
            from .annotate import ensure_celltype
            a, src = ensure_celltype(a, celltype_col)
        except Exception:
            src = None
    if src is None or src not in a.obs.columns:
        a.obs[out_col] = "unknown"
    else:
        a.obs[out_col] = pd.Categorical([coarse_state_one(v) for v in a.obs[src].astype(str).values])
    return a, out_col


def _sample_mask(adata, sample_col: str = "Sample", reference_samples=None):
    if reference_samples and sample_col in adata.obs.columns:
        return adata.obs[sample_col].astype(str).isin([str(x) for x in reference_samples]).values
    # if no sample whitelist was provided, use existing is_reference as a conservative fallback
    if "is_reference" in adata.obs.columns:
        return adata.obs["is_reference"].astype(bool).values
    return np.zeros(adata.n_obs, dtype=bool)


def add_reference_sets(
    adata,
    celltype_col: str = "CellType",
    sample_col: str = "Sample",
    reference_samples=None,
    out_prefix: str = "is_reference",
):
    """Add multiple named reference masks for sensitivity diagnostics.

    The BM1-BM4 versions are activated by passing reference_samples=["BM1",...].
    If no reference_samples are supplied, the functions fall back to the existing
    `is_reference` mask as the candidate healthy pool.
    """
    a, coarse_col = ensure_coarse_cellstate(adata, celltype_col=celltype_col)
    candidate = _sample_mask(a, sample_col=sample_col, reference_samples=reference_samples)
    state = a.obs[coarse_col].astype(str)
    lymph = state.isin(LYMPHOID_STATES).values
    myprog = state.isin(MYELOID_STATES | HSPC_STATES | ERY_STATES).values
    hspc = state.isin(HSPC_STATES).values
    # names preserve earlier patch contract; if not BM1-BM4, still useful as the configured healthy pool
    suffix = "BM1_BM4" if reference_samples else "configured"
    masks = {
        f"{out_prefix}_lymphoid_{suffix}": candidate & lymph,
        f"{out_prefix}_all_healthy_{suffix}": candidate,
        f"{out_prefix}_myeloid_prog_{suffix}": candidate & myprog,
        f"{out_prefix}_hspc_prog_{suffix}": candidate & hspc,
    }
    for k, v in masks.items():
        a.obs[k] = v
    rows = []
    for name, mask in masks.items():
        vc = state[mask].value_counts()
        rows.append({
            "reference_set": name,
            "n_cells": int(mask.sum()),
            "n_states": int(vc.shape[0]),
            "top_states": "; ".join(f"{idx}:{int(val)}" for idx, val in vc.head(8).items()),
            "sample_col": sample_col,
            "reference_samples": ",".join(map(str, reference_samples or [])),
        })
    return a, pd.DataFrame(rows), list(masks.keys())


def add_normal_display_mask(
    adata,
    celltype_col: str = "CellType",
    malignant_label_regex: str | None = None,
    reference_mask_col: str = "is_reference",
    out_col: str = "is_normal_display",
):
    """Flag cells that should be GREYED on score UMAPs as non-malignant ("normal"),
    comprehensively — including normal immune/stromal cells that live *inside patient
    samples*, which a sample-restricted reference (e.g. BM1-BM4 only) misses.

    Why this exists: for AML/van Galen the healthy *baseline* is correctly restricted
    to clean normal marrow (BM1-BM4), but that makes ``is_reference`` cover only BM1-BM4
    cells, so the large normal T/NK/myeloid populations within the AML patient samples
    are never greyed and a myeloid-leaning score (NPM1) lights them up. Solid tumours
    don't hit this because their normal compartment is immune/stromal with no sample
    restriction.

    Resolution:
      * If ``malignant_label_regex`` is given (van Galen: ``-like$``), the malignant
        compartment is the dataset's own annotation (cells whose label matches), and
        every other annotated cell is greyed. This is label-based and reliable.
      * Otherwise falls back to ``is_reference`` so other cohorts are unchanged.

    Writes ``out_col`` (True = normal, grey it) and, when a regex is used,
    ``is_malignant_label``. Returns (adata, info).
    """
    a = adata.copy()
    info = {"method": None, "n_normal": 0, "n_malignant": 0}
    if malignant_label_regex and celltype_col in a.obs.columns:
        lab = a.obs[celltype_col].astype(str)
        is_mal = lab.str.contains(malignant_label_regex, regex=True, case=False, na=False)
        known = ~lab.str.strip().str.lower().isin(["nan", "none", "unknown", "", "?"])
        a.obs["is_malignant_label"] = is_mal.values
        a.obs[out_col] = (known & ~is_mal).values
        info.update(method=f"label_regex:{malignant_label_regex}",
                    n_normal=int((known & ~is_mal).sum()), n_malignant=int(is_mal.sum()))
        print(f"[reference] normal-display mask via '{malignant_label_regex}': "
              f"{info['n_malignant']} malignant-labelled, {info['n_normal']} greyed as normal.")
    else:
        # General solid-tumour / non-AML fallback: grey out expected normal
        # immune/stromal compartments even when no clean reference sample exists.
        # This makes score UMAPs interpretable for public cohorts that only have
        # patient samples: epithelial/HSPC-like tumour compartments remain in the
        # foreground, while lymphoid/myeloid/endothelial/stromal cells are shown as
        # normal context rather than competing score claims.
        normal = np.zeros(a.n_obs, dtype=bool)
        method_bits = []
        if reference_mask_col in a.obs.columns:
            normal |= a.obs[reference_mask_col].astype(bool).values
            method_bits.append(reference_mask_col)
        if celltype_col in a.obs.columns:
            coarse = pd.Series([coarse_state_one(v) for v in a.obs[celltype_col].astype(str).values], index=a.obs.index)
            normal_states = (LYMPHOID_STATES | MYELOID_STATES | {"plasma", "dendritic", "endothelial", "stromal", "erythroid"})
            normal |= coarse.isin(normal_states).values
            method_bits.append("coarse_normal_lineages")
        a.obs[out_col] = normal
        info.update(method="+".join(method_bits) if method_bits else "none", n_normal=int(normal.sum()))
        if normal.any():
            print(f"[reference] normal-display mask via {info['method']}: {info['n_normal']} greyed as expected normal/context cells.")
    return a, info


def _ecdf_percentile(values, ref):
    """Fraction of same-cell-state reference scores <= each value.

    This is intentionally simple/non-parametric and is used by
    add_cellstate_guarded_scores to create mutation_refpct_<GENE>.
    """
    values = np.asarray(values, dtype=float)
    out = np.full(len(values), np.nan, dtype=float)
    ref = np.asarray(ref, dtype=float)
    ref = ref[np.isfinite(ref)]
    if ref.size == 0:
        return out
    r = np.sort(ref)
    finite = np.isfinite(values)
    out[finite] = np.searchsorted(r, values[finite], side="right") / float(len(r))
    return np.clip(out, 0.0, 1.0)


def add_cellstate_guarded_scores(
    adata,
    cfg,
    drivers: list[str] | None = None,
    reference_mask_col: str | None = None,
    cellstate_col: str = "celltype_coarse",
    min_reference_per_state: int = 25,
):
    """Add same-cell-state residual and percentile score columns.

    Columns written per driver:
      mutation_resid_<GENE>   = raw score - median(raw score of healthy cells in same coarse state)
      mutation_refpct_<GENE>  = ECDF percentile vs healthy cells in same coarse state

    If a state has too few reference cells, the global reference distribution is used
    as a fallback and the per-driver/state fallback is recorded in the returned table.
    """
    a = adata.copy()
    drivers = drivers or [c.replace("mutation_prob_", "") for c in a.obs.columns if c.startswith("mutation_prob_")]
    if cellstate_col not in a.obs.columns:
        a, cellstate_col = ensure_coarse_cellstate(a, out_col=cellstate_col)
    if reference_mask_col is None:
        preferred = ["is_reference_all_healthy_BM1_BM4", "is_reference_all_healthy_configured", "is_reference"]
        reference_mask_col = next((c for c in preferred if c in a.obs.columns and a.obs[c].astype(bool).sum() > 0), None)
    if reference_mask_col is None or reference_mask_col not in a.obs.columns:
        print("[reference] no usable reference mask for cell-state guarded scores; skipped.")
        return a, pd.DataFrame()
    ref_mask = a.obs[reference_mask_col].astype(bool).values
    state = a.obs[cellstate_col].astype(str).values
    rows = []
    for d in drivers:
        raw_col = cfg.score_col(a, d, use_calibrated=False) if hasattr(cfg, "score_col") else f"mutation_prob_{d}"
        if raw_col not in a.obs.columns:
            continue
        x = a.obs[raw_col].astype(float).values
        global_ref = x[ref_mask & np.isfinite(x)]
        if global_ref.size == 0:
            continue
        resid = np.full(len(x), np.nan, dtype=float)
        pct = np.full(len(x), np.nan, dtype=float)
        for st in pd.unique(state):
            m_state = state == st
            ref_state = x[ref_mask & m_state & np.isfinite(x)]
            fallback = False
            if ref_state.size < min_reference_per_state:
                ref_state = global_ref
                fallback = True
            med = float(np.nanmedian(ref_state)) if ref_state.size else np.nan
            vals = x[m_state]
            resid[m_state] = vals - med
            pct[m_state] = _ecdf_percentile(vals, ref_state)
            rows.append({
                "driver": d,
                "cellstate": st,
                "reference_mask_col": reference_mask_col,
                "n_cells_state": int(m_state.sum()),
                "n_reference_state": int((ref_mask & m_state).sum()),
                "n_reference_used": int(ref_state.size),
                "used_global_fallback": bool(fallback),
                "reference_median_raw": med,
                "raw_col": raw_col,
                "resid_col": f"mutation_resid_{d}",
                "pct_col": f"mutation_refpct_{d}",
            })
        a.obs[f"mutation_resid_{d}"] = resid
        a.obs[f"mutation_refpct_{d}"] = pct
    return a, pd.DataFrame(rows)


def reference_direction_diagnostics(
    adata,
    cfg,
    drivers: list[str] | None = None,
    reference_mask_col: str = "is_reference",
    malignant_mask_col: str | None = None,
) -> pd.DataFrame:
    """Diagnose when global healthy reference is higher than tumour cells.

    This catches the NPM1-like case: Mut > WT may be true, while Tumour vs global
    healthy reference is negative because healthy HSPC/myeloid cells share the program.
    """
    from . import stats as st
    from .singlecell import find_percell_truth_columns, _binarize_genotype
    drivers = drivers or [c.replace("mutation_prob_", "") for c in adata.obs.columns if c.startswith("mutation_prob_")]
    if reference_mask_col not in adata.obs.columns:
        return pd.DataFrame()
    ref_mask = adata.obs[reference_mask_col].astype(bool).values
    if malignant_mask_col and malignant_mask_col in adata.obs.columns:
        sample_mask = adata.obs[malignant_mask_col].astype(bool).values
    else:
        sample_mask = ~ref_mask
    truth_cols = find_percell_truth_columns(adata, drivers)
    rows = []
    for d in drivers:
        col = cfg.score_col(adata, d, use_calibrated=False) if hasattr(cfg, "score_col") else f"mutation_prob_{d}"
        if col not in adata.obs.columns:
            continue
        x = adata.obs[col].astype(float).values
        ref = x[ref_mask & np.isfinite(x)]
        tum = x[sample_mask & np.isfinite(x)]
        if ref.size and tum.size:
            sh = st.distribution_shift(tum, ref, reference_quantile=getattr(cfg, "reference_quantile", 0.95))
            tumour_vs_ref = sh.cliffs_delta
            median_tumour = float(np.nanmedian(tum))
            median_ref = float(np.nanmedian(ref))
        else:
            sh = None
            tumour_vs_ref = np.nan
            median_tumour = np.nan
            median_ref = np.nan
        mut_vs_wt = np.nan
        n_mut = n_wt = 0
        if d in truth_cols:
            geno = _binarize_genotype(adata.obs[truth_cols[d]]).values
            keep = np.isfinite(geno) & np.isfinite(x)
            mut = x[keep & (geno == 1)]
            wt = x[keep & (geno == 0)]
            n_mut, n_wt = int(len(mut)), int(len(wt))
            if n_mut >= 2 and n_wt >= 2:
                mut_vs_wt = float(st.cliffs_delta(mut, wt, n_boot=0))
        flag = bool(np.isfinite(tumour_vs_ref) and tumour_vs_ref < -0.2)
        disagree = bool(flag and np.isfinite(mut_vs_wt) and mut_vs_wt > 0)
        interp = "reference_high_cellstate_confounded" if disagree else ("reference_higher_than_sample" if flag else "sample_not_below_reference")
        rows.append({
            "driver": d,
            "score_col": col,
            "reference_mask_col": reference_mask_col,
            "n_reference": int(ref.size),
            "n_sample": int(tum.size),
            "median_reference": median_ref,
            "median_sample": median_tumour,
            "tumour_vs_ref_cliffs": tumour_vs_ref,
            "mut_vs_wt_cliffs": mut_vs_wt,
            "n_mut_cells": n_mut,
            "n_wt_cells": n_wt,
            "reference_high_flag": flag,
            "lineage_confound_flag": disagree,
            "recommended_primary_singlecell_metric": "cellstate_residual_or_mut_vs_wt" if flag else "global_reference_shift_ok",
            "interpretation": interp,
        })
    return pd.DataFrame(rows)
```
