## Supplemental_Code for "scOPE identifies which driver-associated expression programs transfer from bulk tumors to single cells": report.py

```
"""
scope_analysis.report
======================
Assembles every table and figure into a manuscript-ready manifest and writes the
driver-prior table the chapter asks for next to Fig. 1.

Nothing here computes science; it collects, formats, and writes, so the final
"what goes where" mapping lives in one auditable place.

Call hierarchy
--------------
    drivers/run_all -> build_driver_prior_table
                     -> save_table / register_figure
                     -> write_manifest
"""
from __future__ import annotations

import json
from dataclasses import dataclass, field
from pathlib import Path

import pandas as pd

from .config import AnalysisConfig, DRIVER_PRIORS, VALIDATION_TIER


# Maps each chapter figure to the suite function(s) that produce it.
FIGURE_MAP = {
    "Fig1_overview":        "(schematic — not data-generated)",
    "Fig2_Table1_bulk":     "bulk.evaluate_all_cancers_bulk + plots.forest_bulk_auroc + interpret.interpret_drivers",
    "Fig3_aml_singlecell":  "singlecell.per_cell_validation + singlecell.patient_bridge + plots.percell_violin",
    "Fig3b_treatment":      "longitudinal.longitudinal_trajectory + plots.treatment_response_trajectory + plots.density_overlay",
    "Fig4_subclonal":       "plots.umap_probability_panels + plots.cluster_probability_heatmap",
    "Fig5_Table2_pancancer":"pancancer.validate_cohort_by_tier + pancancer.normal_vs_tumour_shift",
    "Fig6_cnv":             "cnv.cnv_concordance + cnv.discordance_analysis + plots.cnv_discordance_bar",
    "RobustnessTable":      "robustness.* + pancancer permutation/random-label nulls",
}


@dataclass
class Manifest:
    cfg: AnalysisConfig
    tables: dict[str, str] = field(default_factory=dict)
    figures: dict[str, list[str]] = field(default_factory=dict)

    def save_table(self, name: str, df: pd.DataFrame, subdir: str = "tables") -> Path:
        out = self.cfg.outdir(subdir) / f"{name}.csv"
        df.to_csv(out, index=False)
        self.tables[name] = str(out)
        print(f"[report] table  -> {out}")
        return out

    def register_figure(self, figure_key: str, path) -> None:
        if path is None:
            return
        self.figures.setdefault(figure_key, []).append(str(path))
        print(f"[report] figure -> [{figure_key}] {path}")

    def write_manifest(self) -> Path:
        out = self.cfg.outdir() / "MANIFEST.json"
        payload = {
            "suite": "scope_analysis",
            "config": {k: (str(v) if isinstance(v, Path) else v)
                       for k, v in self.cfg.__dict__.items()},
            "figure_map": FIGURE_MAP,
            "tables": self.tables,
            "figures": self.figures,
        }
        out.write_text(json.dumps(payload, indent=2, default=str))
        print(f"[report] MANIFEST -> {out}")
        return out


def build_driver_prior_table(cfg: AnalysisConfig) -> pd.DataFrame:
    """The chapter's pre-specified driver-prior table (Table next to Fig. 1):
    cancer, driver, prior fingerprint strength, validation tier. Bulk positive/
    negative counts are filled in later by merging with the bulk results table."""
    rows = []
    for cancer, drivers in DRIVER_PRIORS.items():
        for d, tier in drivers.items():
            rows.append(dict(cancer=cancer, driver=d, prior_fingerprint=tier,
                             validation_tier=VALIDATION_TIER.get(cancer, "exploratory")))
    return pd.DataFrame(rows)


def merge_priors_with_results(prior_table: pd.DataFrame,
                              bulk_table: pd.DataFrame) -> pd.DataFrame:
    """Join the pre-specified prior table with realised bulk counts/AUROC so the
    final Table 1 shows prior tier alongside n_pos/n_neg and performance."""
    cols = ["cancer", "driver", "n_pos", "n_neg", "auroc", "auroc_ci_lo",
            "auroc_ci_hi", "fdr_q", "claim_safe"]
    have = [c for c in cols if c in bulk_table.columns]
    return prior_table.merge(bulk_table[have], on=["cancer", "driver"], how="left")
```
