## Supplemental_Code for "scOPE identifies which driver-associated expression programs transfer from bulk tumors to single cells": robustness.py

```
"""
scope_analysis.robustness
==========================
ROBUSTNESS TABLE (Methods "Validation safeguards and negative controls";
supplement to every results figure).

The chapter asks for sensitivity to: latent dimension, classifier family,
decomposition method, and the bulk->sc alignment method, plus calibration
before/after scaling. This module provides each, leaning on scope-bio's own
sweep/grid-search utilities so the ablations exercise the real pipeline.

Call hierarchy
--------------
    drivers/run_robustness -> latent_dim_ablation   (scope.utils.sweep_n_components)
                            -> method_grid_ablation   (scope.utils.grid_search_pipeline)
                            -> alignment_ablation      (re-score sc under each aligner)
                            -> calibration_table       (Brier + ECE before/after)
"""
from __future__ import annotations

import numpy as np
import pandas as pd

from scope import BulkPipeline, SingleCellPipeline
from scope.utils import sweep_n_components, grid_search_pipeline, summarise_grid_search

from . import stats as st
from .config import AnalysisConfig


def _sanitize_bulk_for_decomposition(adata_bulk):
    """Return a copy of the bulk AnnData safe for SVD/decomposition: finite values
    only, and zero-variance genes removed. The crash this prevents is real — when a
    gene has zero variance, scope-bio's scaler divides by a zero std and emits NaN,
    which then breaks randomized_svd ('Input contains NaN'). Constant genes carry no
    information for the latent space, so dropping them is correct, not a workaround."""
    import scipy.sparse as sp
    a = adata_bulk.copy()
    X = a.X
    if sp.issparse(X):
        X = X.toarray()
    X = np.asarray(X, dtype=float)
    X[~np.isfinite(X)] = 0.0
    var = X.var(axis=0)
    keep = np.asarray(var > 1e-12).ravel()
    n_drop = int((~keep).sum())
    if n_drop:
        print(f"[robustness] dropping {n_drop} zero-variance/non-finite genes before ablation "
              f"({int(keep.sum())} retained).")
    a = a[:, keep].copy()
    a.X = X[:, keep]
    return a


def _clean_labels_for_ablation(labels: pd.DataFrame, adata_bulk) -> pd.DataFrame:
    """Numeric 0/1 labels aligned to adata obs, with NaNs handled explicitly."""
    y = labels.copy()
    if not y.index.equals(adata_bulk.obs_names):
        common = [x for x in adata_bulk.obs_names if x in y.index]
        if common:
            y = y.loc[common]
        else:
            y.index = adata_bulk.obs_names[:len(y)]
    y = y.apply(pd.to_numeric, errors="coerce")
    # Mutation matrices mean absent/not-called when missing in these derived files;
    # fill NaN as 0 for robustness rather than letting sklearn/scope crash.
    y = y.fillna(0).astype(int)
    keep_cols = []
    for c in y.columns:
        vals = y[c].values
        if np.unique(vals).size == 2 and vals.sum() >= 2 and (len(vals) - vals.sum()) >= 2:
            keep_cols.append(c)
    return y[keep_cols]


def latent_dim_ablation(adata_bulk, labels: pd.DataFrame, cfg: AnalysisConfig) -> pd.DataFrame:
    """Leakage-free latent-size ablation using this suite's hardened bulk OOF path.

    The previous implementation delegated to ``scope.utils.sweep_n_components``;
    in current scope-bio that helper defaults to log1p=True and can re-introduce
    NaNs even after matrix sanitization. Here we reuse ``bulk_oof_predictions`` so
    the ablation honors the same cfg.log1p/norm/decomposition settings as the main
    manuscript model, sanitizes both X and labels, clamps invalid k values, and
    skips only the failing k/cancer instead of killing the run.
    """
    from dataclasses import replace
    from sklearn.metrics import roc_auc_score, average_precision_score
    from . import bulk as bulk_eval

    a = _sanitize_bulk_for_decomposition(adata_bulk)
    labels2 = _clean_labels_for_ablation(labels, a)
    if labels2.empty:
        print("[robustness] no binary mutation labels with >=2 positives/negatives; skipping latent_dim_ablation.")
        return pd.DataFrame()
    # align sanitized AnnData to labels if needed
    common = [x for x in a.obs_names if x in labels2.index]
    if common and len(common) != a.n_obs:
        a = a[common].copy()
        labels2 = labels2.loc[common]
    n_samples, n_genes = a.n_obs, a.n_vars
    kmax = max(min(n_samples - 2, n_genes - 1), 2)
    ks = sorted({int(k) for k in cfg.ablation_n_components if 2 <= int(k) <= kmax})
    if not ks:
        print("[robustness] no valid n_components for this cohort; skipping latent_dim_ablation.")
        return pd.DataFrame()

    rows = []
    # Keep folds feasible for rare mutation labels.
    min_class = min(int(min(labels2[c].sum(), len(labels2) - labels2[c].sum())) for c in labels2.columns)
    cv = max(2, min(int(cfg.cv_folds), int(min_class))) if min_class >= 2 else 2
    for k in ks:
        try:
            cfg_k = replace(cfg, n_components=k, cv_folds=cv, n_boot=100, n_perm=100)
            oof = bulk_eval.bulk_oof_predictions(a, labels2, cfg_k)
            for d, sub in oof.dropna(subset=["y_oof"]).groupby("driver"):
                y = sub["y_true"].astype(int).values
                p = sub["y_oof"].astype(float).values
                finite = np.isfinite(p)
                y, p = y[finite], p[finite]
                if len(y) < 4 or np.unique(y).size < 2:
                    continue
                rows.append({
                    "n_components": k,
                    "driver": d,
                    "auroc_mean": float(roc_auc_score(y, p)),
                    "auroc_std": np.nan,
                    "auprc_mean": float(average_precision_score(y, p)),
                    "n_samples": int(len(y)),
                    "cv_folds": int(cv),
                    "implementation": "suite_bulk_oof_hardened",
                })
        except Exception as e:
            print(f"[robustness] latent_dim_ablation: n_components={k} failed ({e}); skipping that k.")
    return pd.DataFrame(rows)

def method_grid_ablation(adata_bulk, labels: pd.DataFrame, cfg: AnalysisConfig) -> pd.DataFrame:
    """Grid over decomposition x classifier via scope-bio's grid_search_pipeline,
    summarised to mean +/- std AUROC per combination. Demonstrates the result is
    not an artefact of one modelling choice."""
    grid = {"decomposition": list(cfg.ablation_decompositions),
            "classifier": list(cfg.ablation_classifiers),
            "n_components": [cfg.n_components]}
    res = grid_search_pipeline(adata_bulk, labels, grid, cv=cfg.cv_folds)
    if res.empty:
        return res
    return summarise_grid_search(res, metric="auroc")


def alignment_ablation(
    bulk_pipeline,
    adata_bulk_pp,
    adata_sc_raw,
    cfg: AnalysisConfig,
    drivers: list[str],
    per_cell_truth: dict[str, str] | None = None,
) -> pd.DataFrame:
    """Re-score single cells under each alignment method and compare per-cell
    apparent AUROC (where genotype exists) or score spread.

    The chapter flags alignment as a key sensitivity. For each method in
    cfg.ablation_alignments this re-runs SingleCellPipeline and reports the
    per-driver apparent AUROC, so you can show the conclusion is stable across
    z-score / moment-matching / quantile alignment.
    """
    from .singlecell import _binarize_genotype
    rows = []
    for method in cfg.ablation_alignments:
        sc = SingleCellPipeline(bulk_pipeline=bulk_pipeline, alignment_method=method)
        sc.fit(adata_bulk_pp, adata_sc_raw)
        scored = sc.transform(adata_sc_raw)
        for d in drivers:
            col = f"mutation_prob_{d}"
            if col not in scored.obs.columns:
                continue
            score = scored.obs[col].astype(float).values
            if per_cell_truth and d in per_cell_truth and per_cell_truth[d] in scored.obs.columns:
                geno = _binarize_genotype(scored.obs[per_cell_truth[d]]).values
                keep = ~np.isnan(geno)
                auc = st.auroc_safe(geno[keep].astype(int), score[keep])
                rows.append(dict(alignment=method, driver=d, apparent_auroc=auc,
                                 metric="apparent_auroc"))
            else:
                rows.append(dict(alignment=method, driver=d,
                                 score_iqr=float(np.subtract(*np.percentile(score, [75, 25]))),
                                 metric="score_iqr"))
    return pd.DataFrame(rows)


def calibration_table(
    y_true: np.ndarray,
    y_prob: np.ndarray,
    n_bins: int = 10,
) -> dict:
    """Brier score + expected calibration error (ECE, equal-width bins) and an
    isotonic-recalibrated Brier/ECE, so you can report calibration before/after.

    scope-bio's SVDEvaluator draws the reliability curve; this gives the numbers
    for the robustness table.
    """
    from sklearn.isotonic import IsotonicRegression
    y_true = np.asarray(y_true).astype(int); y_prob = np.asarray(y_prob, dtype=float)

    def ece(p):
        bins = np.linspace(0, 1, n_bins + 1)
        idx = np.clip(np.digitize(p, bins) - 1, 0, n_bins - 1)
        e = 0.0
        for b in range(n_bins):
            m = idx == b
            if m.sum() == 0:
                continue
            e += m.mean() * abs(p[m].mean() - y_true[m].mean())
        return float(e)

    brier = float(np.mean((y_prob - y_true) ** 2))
    iso = IsotonicRegression(out_of_bounds="clip").fit(y_prob, y_true)
    p_cal = iso.predict(y_prob)
    return dict(brier=brier, ece=ece(y_prob),
                brier_isotonic=float(np.mean((p_cal - y_true) ** 2)),
                ece_isotonic=ece(p_cal), n=len(y_true))
```
