## Supplemental_Code for "scOPE identifies which driver-associated expression programs transfer from bulk tumors to single cells": singlecell.py

```
"""
scope_analysis.singlecell
==========================
STAGE 2: "Single-cell AML validation tests per-cell mutation-score enrichment"
(Results 2.3, Fig. 3).

Two entry points, one downstream evaluation:

  (A) project_singlecell(bulk_pipeline, adata_bulk_pp, adata_sc_raw, cfg)
        -> real transfer learning via scope.SingleCellPipeline; writes
           mutation_prob_<GENE> into obs. Use when you want to (re)score.

  (B) you already have a scored .h5ad on disk -> load via data.load_scored_singlecell
        and go straight to evaluation.

Evaluation respects the chapter's epistemics:
  * Per-cell metrics are computed ONLY where a genuine per-cell genotype column
    exists (van Galen expressed-mutation: Mut / WT / unknown). They are reported
    as POSITIVE-UNLABELED ("apparent") AUROC/AUPRC — reported-positive enrichment,
    not truth-vs-truth — because an unreported cell is not a confirmed wild type.
  * Cross-patient inference runs on patient-aggregated scores (patient = unit of
    replication), never on pooled cells.

Call hierarchy
--------------
    drivers/run_singlecell -> per_cell_validation        (Fig 3 violins, image 5/13)
                            -> patient_bridge             (Fig 3 per-patient enrichment)
                            -> project_singlecell         (scope.SingleCellPipeline)
"""
from __future__ import annotations

from contextlib import contextmanager
import inspect
import re

import numpy as np
import pandas as pd

from . import stats as st
from .config import AnalysisConfig
from .data import scored_drivers


# ---------------------------------------------------------------------------
# (A) Real transfer-learning projection (scope-bio)
# ---------------------------------------------------------------------------
def _clean_preprocessor_kwargs(kwargs: dict | None) -> dict:
    """Drop comments/empty values before passing kwargs into scope-bio."""
    out = {}
    for k, v in dict(kwargs or {}).items():
        if str(k).startswith("_") or v is None:
            continue
        out[str(k)] = v
    return out


@contextmanager
def _patched_scope_preprocessor(preprocess_kwargs: dict | None):
    """Temporarily make scope.SingleCellPipeline use configured sc-preprocess args.

    scope-bio constructs ``SingleCellPreprocessor`` internally.  The analysis suite
    therefore cannot simply pass an already-built preprocessor object.  This scoped
    monkeypatch is intentionally narrow: while the SingleCellPipeline is being
    constructed/fitted/transformed, any internal ``SingleCellPreprocessor()`` call
    receives the current run's explicit kwargs.  Afterward the original class is
    restored.
    """
    kwargs = _clean_preprocessor_kwargs(preprocess_kwargs)
    if not kwargs:
        yield
        return

    patched = []
    base_cls = None
    try:
        import scope.preprocessing as _scope_preprocessing
        base_cls = getattr(_scope_preprocessing, "SingleCellPreprocessor", None)
    except Exception:
        _scope_preprocessing = None
    try:
        import scope.pipeline.sc_pipeline as _scope_sc_pipeline
        base_cls = getattr(_scope_sc_pipeline, "SingleCellPreprocessor", base_cls)
    except Exception:
        _scope_sc_pipeline = None

    if base_cls is None:
        print("[singlecell-preprocess] WARNING: could not locate scope SingleCellPreprocessor; "
              "configured preprocessing overrides were not applied.")
        yield
        return

    class ConfiguredSingleCellPreprocessor(base_cls):
        def __init__(self, *args, **init_kwargs):
            merged = dict(init_kwargs)
            merged.update(kwargs)
            try:
                super().__init__(*args, **merged)
            except TypeError as e:
                # Be forgiving across scope-bio versions: keep only kwargs accepted
                # by this installed SingleCellPreprocessor.
                try:
                    sig = inspect.signature(base_cls.__init__)
                    allowed = {k: v for k, v in merged.items() if k in sig.parameters}
                    dropped = sorted(set(merged) - set(allowed))
                    if dropped:
                        print(f"[singlecell-preprocess] WARNING: installed scope ignored unsupported "
                              f"preprocess kwarg(s): {dropped}")
                    super().__init__(*args, **allowed)
                except Exception:
                    raise e

    for mod in (_scope_preprocessing, _scope_sc_pipeline):
        if mod is not None and hasattr(mod, "SingleCellPreprocessor"):
            old = getattr(mod, "SingleCellPreprocessor")
            setattr(mod, "SingleCellPreprocessor", ConfiguredSingleCellPreprocessor)
            patched.append((mod, old))

    print(f"[singlecell-preprocess] scope SingleCellPreprocessor overrides: {kwargs}")
    try:
        yield
    finally:
        for mod, old in reversed(patched):
            setattr(mod, "SingleCellPreprocessor", old)


@contextmanager
def _finite_classifier_projection_guard(bulk_pipeline, nan_policy: str = "zero_fill"):
    """Guard classifier prediction against NaN/Inf latent projections.

    The upstream transform computes Z_sc and immediately calls the classifier set.
    If the installed scope-bio preprocessing/alignment creates any non-finite latent
    values, sklearn raises deep inside LogisticRegression.  This guard reports the
    issue at the useful boundary and applies a conservative fill so the run can
    finish and the audit table can expose that it happened.
    """
    cset = getattr(bulk_pipeline, "classifier_set_", None)
    orig = getattr(cset, "predict_proba", None)
    if cset is None or orig is None:
        yield
        return

    policy = str(nan_policy or "zero_fill").lower().strip()

    def guarded_predict_proba(Z, *args, **kwargs):
        arr = np.asarray(Z)
        bad = ~np.isfinite(arr)
        if bad.any():
            n_bad = int(bad.sum())
            n_rows = int(bad.any(axis=1).sum()) if arr.ndim == 2 else int(n_bad)
            print(f"[singlecell-project] WARNING: projected latent matrix contains "
                  f"{n_bad:,} non-finite value(s) across {n_rows:,} cell(s); "
                  f"nan_policy={policy!r}.")
            if policy in {"raise", "fail", "error"}:
                raise ValueError("Projected single-cell latent matrix contains NaN/Inf values.")
            z = np.asarray(Z, dtype=float).copy()
            if policy in {"median", "median_fill", "col_median", "column_median"} and z.ndim == 2:
                finite_z = np.where(np.isfinite(z), z, np.nan)
                med = np.nanmedian(finite_z, axis=0)
                med = np.where(np.isfinite(med), med, 0.0)
                rr, cc = np.where(~np.isfinite(z))
                z[rr, cc] = med[cc]
            else:
                # In the bulk latent space, 0 is the centered/neutral component value.
                z = np.nan_to_num(z, nan=0.0, posinf=0.0, neginf=0.0)
            return orig(z, *args, **kwargs)
        return orig(Z, *args, **kwargs)

    setattr(cset, "predict_proba", guarded_predict_proba)
    try:
        yield
    finally:
        try:
            setattr(cset, "predict_proba", orig)
        except Exception:
            pass


def _sanitize_projected_latents(adata, nan_policy: str = "zero_fill"):
    """Replace NaN/Inf in latent obsm matrices so later UMAP/plots don't crash."""
    policy = str(nan_policy or "zero_fill").lower().strip()
    if policy in {"raise", "fail", "error"}:
        return adata
    for key in list(getattr(adata, "obsm", {}).keys()):
        if not (str(key).lower().startswith("x_svd") or "latent" in str(key).lower() or "scope" in str(key).lower()):
            continue
        try:
            arr = np.asarray(adata.obsm[key])
            bad = ~np.isfinite(arr)
            if not bad.any():
                continue
            z = np.asarray(arr, dtype=float).copy()
            if policy in {"median", "median_fill", "col_median", "column_median"} and z.ndim == 2:
                med = np.nanmedian(np.where(np.isfinite(z), z, np.nan), axis=0)
                med = np.where(np.isfinite(med), med, 0.0)
                rr, cc = np.where(~np.isfinite(z))
                z[rr, cc] = med[cc]
            else:
                z = np.nan_to_num(z, nan=0.0, posinf=0.0, neginf=0.0)
            adata.obsm[key] = z
            print(f"[singlecell-project] sanitized {int(bad.sum()):,} non-finite value(s) in adata.obsm[{key!r}].")
        except Exception as e:
            print(f"[singlecell-project] WARNING: could not sanitize obsm[{key!r}] ({e})")
    return adata


def project_singlecell(
    bulk_pipeline,
    adata_bulk_pp,
    adata_sc_raw,
    cfg: AnalysisConfig,
    preprocess_kwargs: dict | None = None,
    nan_policy: str | None = None,
):
    """Project raw single cells into the bulk latent space and infer per-cell
    mutation probabilities. Thin wrapper over scope.SingleCellPipeline so the
    transfer step is the real, published projection (Z_sc = A'_sc V).

    adata_bulk_pp must be the *preprocessed* bulk object
    (``bulk_pipeline.preprocessor_.transform(adata_bulk)``), per scope-bio's API.
    Returns the scored single-cell AnnData (mutation_prob_<GENE> in obs).

    ``preprocess_kwargs`` are forwarded to the upstream scope-bio
    SingleCellPreprocessor.  For already-curated/scored h5ads, use e.g.
    filter_strategy='none', already_qc_filtered=True, already_normalized=True,
    already_log_transformed=True, run_doublet_detection=False so rescoring uses the
    same cells/matrix instead of re-QCing away most of the object.
    """
    if preprocess_kwargs is None:
        preprocess_kwargs = getattr(cfg, "singlecell_preprocessing", None)
    nan_policy = nan_policy or getattr(cfg, "singlecell_projection_nan_policy", "zero_fill")

    with _patched_scope_preprocessor(preprocess_kwargs), _finite_classifier_projection_guard(bulk_pipeline, nan_policy):
        from scope import SingleCellPipeline  # deferred: keeps scope-free imports/tests working
        sc = SingleCellPipeline(
            bulk_pipeline=bulk_pipeline,
            alignment_method=cfg.alignment_method,
        )
        sc.fit(adata_bulk_pp, adata_sc_raw)
        out = sc.transform(adata_sc_raw)
    return _sanitize_projected_latents(out, nan_policy)


# ---------------------------------------------------------------------------
# Per-cell genotype columns: detect real truth, reject prediction look-alikes
# ---------------------------------------------------------------------------
_TRUTH_RE = re.compile(r"(genotype|mutant|mutation|status|variant|call|label)", re.I)
_PRED_RE = re.compile(r"(mutation_prob|_prob|_score|^score|pred|probability)", re.I)
# scOPE's derived SCORE columns all live in the 'mutation_<kind>_<gene>' namespace.
# They must never be read as genotype (the bug that let mutation_resid_*/refpct_*/adj_*
# pass the truth filter and be picked up as ground truth depending on column order).
_SCORE_NAMESPACE = re.compile(
    r"^mutation_(prob|calib|calibrated|resid|residual|refpct|pct|adj|adjusted|"
    r"gated|gatedadj|score|conf|confidence)(?:_|$)", re.I)

# Columns created by this analysis suite.  Keeping this centralized matters because
# stale derived columns (mutation_adj_*, mutation_resid_*, mutation_refpct_*, old
# mutation_calib_*) can otherwise be silently reused by plotting code even after the
# raw mutation_prob_* values have been recomputed.
_SCORE_COLUMN_RE = re.compile(
    r"^mutation_(prob|calib|calibrated|resid|residual|refpct|pct|adj|adjusted|"
    r"gated|gatedadj|score|conf|confidence)(?:_|$)", re.I)
_RAW_SCORE_COLUMN_RE = re.compile(r"^mutation_prob(?:_|$)", re.I)


def scope_score_columns(adata, include_raw: bool = True, include_derived: bool = True) -> list[str]:
    """Return obs columns created by scOPE/scope_analysis scoring.

    Parameters
    ----------
    include_raw
        Include raw transferred probabilities (mutation_prob_*). Use True before a
        forced rescore so old probabilities cannot leak into the new object.
    include_derived
        Include run-dependent derived scores (calibrated, confidence-adjusted,
        residual/percentile guardrail scores, etc.). Use True at the start of every
        run because these depend on the current config/reference/masking choices.
    """
    cols = []
    for c in list(getattr(adata, "obs", pd.DataFrame()).columns):
        cs = str(c)
        is_score = bool(_SCORE_COLUMN_RE.search(cs))
        is_raw = bool(_RAW_SCORE_COLUMN_RE.search(cs))
        if is_score and ((include_raw and is_raw) or (include_derived and not is_raw)):
            cols.append(c)
    return cols


def clear_scope_score_columns(adata, include_raw: bool = False, include_derived: bool = True,
                              *, label: str = "singlecell"):
    """Drop stale score columns from ``adata.obs`` and report what was removed.

    This deliberately leaves true genotype/status columns alone. It only targets
    the reserved score namespace used by the pipeline. Returns ``(adata, cols)`` so
    callers can include the count in audit tables.
    """
    cols = scope_score_columns(adata, include_raw=include_raw, include_derived=include_derived)
    if cols:
        adata.obs.drop(columns=cols, inplace=True, errors="ignore")
        shown = [str(c) for c in cols[:8]]
        more = "" if len(cols) <= 8 else f" ... +{len(cols) - 8} more"
        print(f"[singlecell] {label}: dropped {len(cols)} stale score column(s): {shown}{more}")
    return adata, cols


def _is_discrete_genotype(series: pd.Series) -> bool:
    """True if values look like a discrete genotype (binarizes to {0,1,nan},
    <=12 distinct raw values), not a continuous score."""
    s = pd.Series(series)
    nun = s.dropna().nunique()
    if nun == 0 or nun > 12:
        return False
    try:
        b = _binarize_genotype(s)
        vals = set(np.unique(b.values[~pd.isna(b.values)]))
        return vals.issubset({0.0, 1.0}) and len(vals) >= 1
    except Exception:
        return True


def find_percell_truth_columns(adata, drivers: list[str]) -> dict[str, str]:
    """Map driver -> obs column holding genuine per-cell genotype, if present.

    A column qualifies only if it (a) mentions the driver gene, (b) matches a
    truth-like keyword, (c) is NOT a prediction look-alike, (d) is NOT in the scOPE
    score namespace, and (e) has discrete genotype-like values. (d)+(e) close the
    loophole that let derived score columns (mutation_resid_*/refpct_*/adj_*) be
    mistaken for ground truth — scores can never be read as truth.
    """
    found = {}
    for d in drivers:
        dl = d.lower()
        cands = [c for c in adata.obs.columns
                 if dl in str(c).lower() and _TRUTH_RE.search(c)
                 and not _PRED_RE.search(c) and not _SCORE_NAMESPACE.search(str(c).lower())]
        cands.sort(key=lambda c: (0 if re.search(r"genotype|variant", str(c), re.I) else 1, len(str(c))))
        for c in cands:
            if _is_discrete_genotype(adata.obs[c]):
                found[d] = c
                break
    return found


def _binarize_genotype(series: pd.Series) -> pd.Series:
    """Map a per-cell genotype column to {1: mutant, 0: WT/reference, NaN: unknown}.

    Handles string ('Mut'/'WT'), integer (1/0), float ('1.0'/'0.0'), and boolean
    (True/False) encodings — numeric/boolean columns are common when genotype is
    stored as a 0/1 indicator, and missing them silently drops the driver from the
    per-cell validation. Anything not clearly mutant or WT becomes NaN (unknown) and
    is excluded from apparent-AUROC, consistent with positive-unlabeled handling.
    """
    s = series.astype(str).str.strip().str.lower()
    mut = s.isin(["mut", "mutant", "1", "1.0", "alt", "positive", "pos", "variant", "true"])
    wt = s.isin(["wt", "wildtype", "wild-type", "0", "0.0", "ref", "reference",
                 "negative", "neg", "false"])
    out = pd.Series(np.nan, index=series.index)
    out[mut] = 1.0
    out[wt] = 0.0
    return out


# ---------------------------------------------------------------------------
# van Galen expressed-mutation transcript truth parser
# ---------------------------------------------------------------------------
def _find_obs_col_case_insensitive(obs: pd.DataFrame, names: list[str]) -> str | None:
    """Return the first obs column matching one of `names`, case-insensitively."""
    exact = set(obs.columns)
    for n in names:
        if n in exact:
            return n
    low = {str(c).lower(): c for c in obs.columns}
    for n in names:
        if str(n).lower() in low:
            return low[str(n).lower()]
    return None


def _driver_in_transcript_series(series: pd.Series, driver: str) -> pd.Series:
    """True where a MutTranscript/WtTranscript-style string mentions `driver`."""
    pat = r"(?<![A-Za-z0-9])" + re.escape(str(driver).strip()) + r"(?![A-Za-z0-9])"
    return series.astype(str).str.contains(pat, case=False, regex=True, na=False)


def add_muttranscript_genotype_columns(
    adata,
    drivers: list[str] | None = None,
    mut_col: str | None = None,
    wt_col: str | None = None,
    overwrite: bool = True,
) -> tuple[object, pd.DataFrame]:
    """Parse van Galen-style MutTranscript/WtTranscript obs strings into true
    per-cell genotype columns named <DRIVER>_genotype.

    Semantics:
      * driver mentioned in MutTranscript -> 1 / Mut
      * driver mentioned in WtTranscript  -> 0 / WT
      * neither mentioned                 -> NaN / unknown
      * both mentioned                    -> 1 wins, conservatively
    """
    obs = adata.obs

    mut_col = mut_col if mut_col in obs.columns else _find_obs_col_case_insensitive(
        obs,
        [
            "MutTranscript", "MutTranscripts",
            "mut_transcript", "mut_transcripts",
            "MutatedTranscript", "MutatedTranscripts",
        ],
    )

    wt_col = wt_col if wt_col in obs.columns else _find_obs_col_case_insensitive(
        obs,
        [
            "WtTranscript", "WTTranscript",
            "WtTranscripts", "WTTranscripts",
            "wt_transcript", "wt_transcripts",
            "WildtypeTranscript", "WildTypeTranscript",
        ],
    )

    if mut_col is None and wt_col is None:
        return adata, pd.DataFrame()

    drivers = drivers or scored_drivers(adata)
    mut_s = obs[mut_col].astype(str) if mut_col is not None else pd.Series("", index=obs.index)
    wt_s = obs[wt_col].astype(str) if wt_col is not None else pd.Series("", index=obs.index)

    rows = []

    for d in drivers:
        d = str(d).strip()
        if not d:
            continue

        mut_mask = (
            _driver_in_transcript_series(mut_s, d)
            if mut_col is not None
            else pd.Series(False, index=obs.index)
        )
        wt_mask = (
            _driver_in_transcript_series(wt_s, d)
            if wt_col is not None
            else pd.Series(False, index=obs.index)
        )

        known = mut_mask | wt_mask
        if int(known.sum()) == 0:
            continue

        out = pd.Series(np.nan, index=obs.index, dtype="float")
        out.loc[wt_mask] = 0.0
        out.loc[mut_mask] = 1.0

        out_col = f"{d}_genotype"

        if overwrite or out_col not in obs.columns:
            obs[out_col] = out.values

        rows.append(
            {
                "driver": d,
                "genotype_col": out_col,
                "muttranscript_col": mut_col,
                "wttranscript_col": wt_col,
                "n_mut_cells": int(mut_mask.sum()),
                "n_wt_cells": int(wt_mask.sum()),
                "n_known_cells": int(known.sum()),
                "n_unknown_cells": int(len(obs) - int(known.sum())),
            }
        )

    summary = pd.DataFrame(rows)

    if not summary.empty:
        print(
            f"[singlecell] parsed MutTranscript/WtTranscript into {len(summary)} "
            f"per-cell genotype column(s): "
            f"{summary['driver'].head(12).tolist()}"
            + (" ..." if len(summary) > 12 else "")
        )
    elif mut_col is not None or wt_col is not None:
        print(
            "[singlecell] MutTranscript/WtTranscript columns found, "
            "but no scored driver names were detected inside them."
        )

    return adata, summary

def resolve_score_col(adata, cfg: AnalysisConfig, driver: str, score_mode: str = "configured") -> str:
    """Resolve the score column for claim-bearing single-cell statistics.

    score_mode="configured" keeps the old cfg.score_col behavior.
    score_mode="cellstate_residual" / "guarded" uses mutation_resid_<GENE> when present.
    score_mode="cellstate_percentile" / "refpct" uses mutation_refpct_<GENE> when present.
    Missing guarded columns fall back to cfg.score_col so a cohort without guardrails
    still runs instead of crashing.
    """
    mode = str(score_mode or "configured").lower()
    if mode in {"cellstate_residual", "residual"}:
        c = f"mutation_resid_{driver}"
        if c in adata.obs.columns:
            return c
    if mode in {"cellstate_percentile", "percentile", "refpct"}:
        c = f"mutation_refpct_{driver}"
        if c in adata.obs.columns:
            return c
    if mode in {"guarded", "guarded_if_available"}:
        preferred = getattr(cfg, "longitudinal_guarded_mode", "cellstate_residual")
        first = f"mutation_refpct_{driver}" if str(preferred).lower() in {"cellstate_percentile", "percentile", "refpct"} else f"mutation_resid_{driver}"
        second = f"mutation_resid_{driver}" if first.startswith("mutation_refpct_") else f"mutation_refpct_{driver}"
        for c in (first, second):
            if c in adata.obs.columns:
                return c
    if mode in {"raw", "raw_probability"}:
        return cfg.score_col(adata, driver, use_calibrated=False)
    return cfg.score_col(adata, driver)


# ---------------------------------------------------------------------------
# Per-cell validation (apparent / positive-unlabeled)  -> Fig 3 violins, image 5
# ---------------------------------------------------------------------------
def per_cell_validation(
    adata,
    cfg: AnalysisConfig,
    drivers: list[str] | None = None,
    score_mode: str = "configured",
) -> tuple[pd.DataFrame, pd.DataFrame]:
    """Where per-cell genotype exists, compare scores between reported-mutant and
    WT/reference cells.

    score_mode : "configured" uses the cfg score column (raw/calibrated). Any other
        value (e.g. "cellstate_residual") prefers the matching guarded column
        (mutation_resid_*/mutation_refpct_*) when present, so Fig 3 can be shown
        guarded against normal-cell contamination. Falls back to the cfg column.

    Returns
    -------
    summary : DataFrame  (one row per driver with a usable truth column)
        apparent AUROC + CI, Mann-Whitney p, Cliff's delta + CI, n_mut, n_wt.
        These are labelled apparent/PU metrics, NOT truth-vs-truth.
    cell_long : DataFrame  (tidy per-cell rows for plotting the violins)
        columns: [driver, score, genotype_label] for cells with known genotype.
    """
    drivers = drivers or scored_drivers(adata)
    truth_cols = find_percell_truth_columns(adata, drivers)
    if not truth_cols:
        print("[singlecell] no per-cell genotype columns found; "
              "use patient_bridge() instead for this cohort.")
        return pd.DataFrame(), pd.DataFrame()

    rows, cell_rows = [], []
    for d, col in truth_cols.items():
        score_col = resolve_score_col(adata, cfg, d, score_mode=score_mode)
        score = adata.obs[score_col].astype(float).values
        geno = _binarize_genotype(adata.obs[col]).values
        keep = ~np.isnan(geno)
        y, p = geno[keep].astype(int), score[keep]
        n_mut, n_wt = int(y.sum()), int((1 - y).sum())
        for yi, pi in zip(y, p):
            cell_rows.append(dict(driver=d, score=pi, score_col=score_col,
                                  score_kind=cfg.score_kind(score_col),
                                  genotype_label="Mut" if yi == 1 else "WT"))
        if n_mut < 2 or n_wt < 2:
            print(f"[singlecell]   {d}: too few of one class (mut={n_mut}, wt={n_wt}).")
            continue
        auc = st.bootstrap_auroc_ci(y, p, n_boot=cfg.n_boot, random_state=cfg.random_state)
        eff = st.mannwhitney_effect(p[y == 1], p[y == 0], n_boot=cfg.n_boot,
                                    random_state=cfg.random_state)
        rows.append(dict(
            driver=d, truth_col=col, score_col=score_col, score_kind=cfg.score_kind(score_col),
            score_mode=score_mode,
            n_mut=n_mut, n_wt=n_wt,
            apparent_auroc=auc.auroc, auroc_ci_lo=auc.ci_lo, auroc_ci_hi=auc.ci_hi,
            mannwhitney_p=eff.p_value, cliffs_delta=eff.cliffs_delta,
            delta_ci_lo=eff.delta_lo, delta_ci_hi=eff.delta_hi,
            metric_type="apparent_positive_unlabeled",
        ))
    summary = pd.DataFrame(rows)
    if not summary.empty:
        summary["fdr_q"] = st.bh_fdr(summary["mannwhitney_p"].values)
    return summary, pd.DataFrame(cell_rows)


# ---------------------------------------------------------------------------
# Patient-level bridge  -> Fig 3 per-patient enrichment (anti-pseudoreplication)
# ---------------------------------------------------------------------------
def patient_bridge(
    adata,
    patient_status: pd.DataFrame,
    cfg: AnalysisConfig,
    patient_col: str = "patient",
    drivers: list[str] | None = None,
    score_mode: str = "configured",
) -> pd.DataFrame:
    """Aggregate per-cell scores to patient x driver and test mutant vs WT
    patients. The patient is the unit of replication.

    patient_status : tidy [patient, mutation, label] (data.load_patient_status).

    Returns one row per driver: number of mutant/WT patients, Mann-Whitney p on
    patient-level aggregated scores, Cliff's delta + CI, and a global FDR q.
    """
    drivers = drivers or scored_drivers(adata)
    if patient_col not in adata.obs.columns:
        raise KeyError(f"patient_col '{patient_col}' missing; run "
                       f"data.derive_patient_timepoint first.")
    rows = []
    for d in drivers:
        score_col = resolve_score_col(adata, cfg, d, score_mode=score_mode)
        cell_df = adata.obs[[patient_col, score_col]].copy()
        agg = st.patient_aggregate(cell_df, patient_col, score_col,
                                   min_cells=cfg.min_cells_per_patient)
        status_d = patient_status[patient_status.mutation == d][["patient", "label"]]
        merged = agg.merge(status_d, left_on=patient_col, right_on="patient", how="inner")
        if merged.empty:
            continue
        mut = merged[merged.label == 1][score_col].values
        wt = merged[merged.label == 0][score_col].values
        if len(mut) < 2 or len(wt) < 2:
            print(f"[singlecell]   patient bridge {d}: mutant patients={len(mut)}, "
                  f"WT patients={len(wt)} (need >=2 each).")
            continue
        eff = st.mannwhitney_effect(mut, wt, n_boot=cfg.n_boot, random_state=cfg.random_state)
        rows.append(dict(
            driver=d, score_col=score_col, score_kind=cfg.score_kind(score_col),
            score_mode=score_mode,
            n_mut_patients=len(mut), n_wt_patients=len(wt),
            median_mut=float(np.median(mut)), median_wt=float(np.median(wt)),
            mannwhitney_p=eff.p_value, cliffs_delta=eff.cliffs_delta,
            delta_ci_lo=eff.delta_lo, delta_ci_hi=eff.delta_hi,
        ))
    out = pd.DataFrame(rows)
    if not out.empty:
        out["fdr_q"] = st.bh_fdr(out["mannwhitney_p"].values)
    return out


# ---------------------------------------------------------------------------
# Score CALIBRATION  ->  readable [0,1] scores that preserve all rank statistics
# ---------------------------------------------------------------------------
def _reference_ecdf_monotone(x: np.ndarray, ref: np.ndarray) -> np.ndarray:
    """Strictly-monotone, reference-anchored map of raw scores to [0,1].

    Within the reference range it is a linearly-interpolated reference ECDF (so a
    value of 0.9 ≈ 'above ~90% of healthy cells'); the lower/upper tails are
    extended linearly to the global score range so cells below/above all reference
    still get distinct, ordered values reaching 0 and 1. Because the mapping is
    strictly increasing in the raw score, the rank ordering of cells is preserved
    EXACTLY — hence AUROC, Cliff's delta, Mann-Whitney and KS are identical to the
    raw-score values; only the spread (readability) changes.
    """
    x = np.asarray(x, float)
    if ref.size == 0:
        order = x.argsort().argsort()
        return order / max(len(x) - 1, 1)
    n = ref.size
    # midpoint-convention CDF knots at each (unique) reference value
    xs_u, idx = np.unique(ref, return_index=True)
    ys_u = np.array([np.mean(((np.arange(n) + 0.5) / n)[ref == v]) for v in xs_u])
    gmin, gmax = float(np.nanmin(x)), float(np.nanmax(x))
    lo_x = min(gmin, xs_u[0]); hi_x = max(gmax, xs_u[-1])
    knots_x = np.concatenate([[lo_x], xs_u, [hi_x]])
    knots_y = np.concatenate([[0.0], ys_u, [1.0]])
    # enforce strictly increasing x for interpolation (dedupe, averaging y)
    kx, ky = [], []
    for xv, yv in zip(knots_x, knots_y):
        if kx and xv <= kx[-1]:
            xv = kx[-1] + 1e-12
        kx.append(xv); ky.append(yv)
    return np.clip(np.interp(x, kx, ky), 0.0, 1.0)


def calibrate_scores(
    adata,
    cfg: AnalysisConfig,
    drivers: list[str] | None = None,
    reference_mask_col: str = "is_reference",
):
    """Add a monotonic, reference-anchored calibrated score in [0,1] for each
    driver, written to obs as ``{cfg.calibrated_score_prefix}_<gene>``.

    Why: scope-bio's ``mutation_prob_<gene>`` are real predict_proba values, but
    for single cells they pile up near the base rate (projected cells sit near the
    latent origin), so plotting on a 0-1 axis looks like a flat smear even when the
    Mut-vs-WT difference is hugely significant. Calibration spreads the scores
    across [0,1] WITHOUT changing any rank-based statistic (AUROC, Cliff's delta,
    Mann-Whitney, KS, Spearman are all invariant under a strictly monotone map),
    so significance is identical; only readability improves.

    Methods (``cfg.calibration_method``)
      * ``"reference_ecdf"`` (default, recommended): calib_i = fraction of healthy
        reference cells whose raw score <= cell i's raw score. Interpretable as the
        cell's percentile against the healthy null ("how abnormal vs normal"):
        healthy cells are ~uniform with median 0.5, tumour cells carrying the
        programme are pushed toward 1. Ties handled by 'right' side.
      * ``"minmax"``: (x - min)/(max - min) per driver over all cells.
      * ``"rank"``: global percentile rank over all cells.

    The raw probability column is never overwritten. Returns the (copied) AnnData.
    """
    drivers = drivers or scored_drivers(adata)
    adata = adata.copy()
    method = cfg.calibration_method
    has_ref = reference_mask_col in adata.obs.columns and adata.obs[reference_mask_col].any()
    if method == "reference_ecdf" and not has_ref:
        print("[singlecell] calibrate_scores: no reference cells flagged; "
              "falling back to 'rank' calibration for this cohort.")
        method = "rank"

    for d in drivers:
        raw_col = f"{cfg.raw_score_prefix}_{d}"
        if raw_col not in adata.obs.columns:
            continue
        x = adata.obs[raw_col].astype(float).values
        out = np.full(len(x), np.nan)
        finite = ~np.isnan(x)
        xf = x[finite]
        if xf.size == 0:
            adata.obs[f"{cfg.calibrated_score_prefix}_{d}"] = out
            continue
        if method == "reference_ecdf":
            ref = adata.obs.loc[adata.obs[reference_mask_col], raw_col].astype(float).values
            ref = np.sort(ref[~np.isnan(ref)])
            out[finite] = _reference_ecdf_monotone(xf, ref)
        elif method == "minmax":
            lo, hi = np.nanmin(xf), np.nanmax(xf)
            out[finite] = (xf - lo) / (hi - lo) if hi > lo else 0.5
        else:  # rank
            order = xf.argsort().argsort()
            out[finite] = order / max(len(xf) - 1, 1)
        adata.obs[f"{cfg.calibrated_score_prefix}_{d}"] = out
    print(f"[singlecell] calibrated {len([d for d in drivers if f'{cfg.raw_score_prefix}_{d}' in adata.obs.columns])} "
          f"drivers via '{method}' -> '{cfg.calibrated_score_prefix}_*' (monotone; ranks/significance unchanged).")
    return adata


# ---------------------------------------------------------------------------
# Direction / lineage-confound diagnostic
# ---------------------------------------------------------------------------
def reference_direction_diagnostic(
    adata,
    cfg: AnalysisConfig,
    drivers: list[str] | None = None,
    reference_mask_col: str = "is_reference",
    score_mode: str = "configured",
) -> pd.DataFrame:
    """Per driver, report whether the non-reference (tumour) compartment scores
    ABOVE or BELOW the reference compartment, and whether that agrees with the
    per-cell Mut-vs-WT direction where genotype exists.

    This surfaces lineage confounding: if the reference cells are a different
    lineage than the malignant cells (e.g. lymphoid T/B/NK reference vs myeloid
    AML blasts), a driver's tumour-vs-reference shift can invert even though
    Mut-vs-WT among genotyped cells is correct. A row where
    ``tumour_vs_ref_sign`` and ``mut_vs_wt_sign`` disagree is a flag that the
    tumour-vs-reference framing is unreliable for that driver.
    """
    drivers = drivers or scored_drivers(adata)
    truth = find_percell_truth_columns(adata, drivers)
    obs = adata.obs
    is_ref = obs[reference_mask_col].astype(bool).values if reference_mask_col in obs.columns else None
    rows = []
    for d in drivers:
        col = resolve_score_col(adata, cfg, d, score_mode=score_mode)
        s = obs[col].astype(float).values
        ref_sign = mut_sign = np.nan
        if is_ref is not None and is_ref.any() and (~is_ref).any():
            tum, ref = s[~is_ref], s[is_ref]
            tum, ref = tum[~np.isnan(tum)], ref[~np.isnan(ref)]
            if len(tum) and len(ref):
                ref_sign = st._cliffs_point(tum, ref)
        if d in truth:
            g = _binarize_genotype(obs[truth[d]]).values
            k = ~np.isnan(g)
            if k.sum() and (g[k] == 1).any() and (g[k] == 0).any():
                mut_sign = st._cliffs_point(s[k][g[k] == 1], s[k][g[k] == 0])
        agree = (np.sign(ref_sign) == np.sign(mut_sign)) if (ref_sign == ref_sign and mut_sign == mut_sign) else np.nan
        rows.append(dict(
            driver=d,
            score_col=col,
            score_kind=cfg.score_kind(col),
            score_mode=score_mode,
            tumour_vs_ref_cliffs=ref_sign,
            tumour_vs_ref_sign=("above" if ref_sign > 0 else "below") if ref_sign == ref_sign else "n/a",
            mut_vs_wt_cliffs=mut_sign,
            mut_vs_wt_sign=("mut>wt" if mut_sign > 0 else "mut<wt") if mut_sign == mut_sign else "n/a",
            directions_agree=agree,
            lineage_confound_flag=(agree == False),  # noqa: E712 (want explicit False, not NaN)
        ))
    return pd.DataFrame(rows)


# ---------------------------------------------------------------------------
# Malignant-compartment call (for within-tumour longitudinal / shift analyses)
# ---------------------------------------------------------------------------
def derive_malignant_compartment(
    adata,
    cfg: AnalysisConfig,
    drivers: list[str] | None = None,
    reference_mask_col: str = "is_reference",
    out_col: str = "scope_malignant",
):
    """Flag cells as malignant if their max raw driver probability exceeds the
    reference upper quantile (anchored to what normal cells look like, not an
    arbitrary 0.5). Writes a bool obs column and returns (adata, threshold).

    Used to restrict the longitudinal trajectory to the malignant compartment so
    the treatment-response signal is not diluted by changing normal-cell
    admixture across timepoints (and is robust to lineage-mismatched references).
    """
    drivers = drivers or scored_drivers(adata)
    cols = [f"{cfg.raw_score_prefix}_{d}" for d in drivers
            if f"{cfg.raw_score_prefix}_{d}" in adata.obs.columns]
    adata = adata.copy()
    mx = adata.obs[cols].astype(float).max(axis=1).values
    if reference_mask_col in adata.obs.columns and adata.obs[reference_mask_col].any():
        ref = mx[adata.obs[reference_mask_col].values]
        thr = float(np.nanquantile(ref, cfg.malignant_call_quantile))
    else:
        thr = float(np.nanquantile(mx, 0.5))
    adata.obs[out_col] = (mx > thr)
    print(f"[singlecell] malignant compartment: {int(adata.obs[out_col].sum())}/{adata.n_obs} "
          f"cells (max-driver raw score > {thr:.3g}).")
    return adata, thr
```
