## Supplemental_Code for "scOPE identifies which driver-associated expression programs transfer from bulk tumors to single cells": stats.py

```
"""
scope_analysis.stats
====================
Rigorous, self-contained statistical primitives used everywhere downstream.

Design philosophy
-----------------
``scope-bio`` gives you point estimates (AUROC / AUPRC / Brier via
``scope.evaluation.metrics``). For a high-tier manuscript you also need
*uncertainty* and *significance* around every one of those point estimates, plus
effect sizes for distribution comparisons and a discipline against
pseudoreplication. That is what this module adds. None of it duplicates
``scope-bio`` logic — it wraps and extends it.

Every function here is deterministic given ``random_state`` and is unit-tested in
``tests/test_stats.py`` against analytically known answers.

Call hierarchy (who calls what)
--------------------------------
    bulk.py        -> bootstrap_auroc_ci, permutation_auroc_pvalue, delong_test
    singlecell.py  -> bootstrap_auroc_ci, mannwhitney_effect, patient_aggregate
    longitudinal.py-> distribution_shift, cliffs_delta
    pancancer.py   -> distribution_shift, permutation_label_null, bh_fdr
    (all)          -> bh_fdr  (applied ONCE, globally, at the end of a stage)

Nothing in this module imports scope-bio, anndata, or any plotting library, so it
stays fast and trivially testable.
"""
from __future__ import annotations

from dataclasses import dataclass, asdict
from typing import Sequence

import numpy as np
import pandas as pd
from scipy import stats as _sps
from sklearn.metrics import roc_auc_score, average_precision_score

__all__ = [
    "AUROCResult",
    "EffectResult",
    "ShiftResult",
    "auroc_safe",
    "auprc_safe",
    "bootstrap_auroc_ci",
    "delong_auc_variance",
    "delong_test",
    "permutation_auroc_pvalue",
    "permutation_label_null",
    "bh_fdr",
    "cliffs_delta",
    "mannwhitney_effect",
    "distribution_shift",
    "patient_aggregate",
]


# ---------------------------------------------------------------------------
# Result containers (so downstream code reads as tidy rows, never loose tuples)
# ---------------------------------------------------------------------------
@dataclass
class AUROCResult:
    auroc: float
    ci_lo: float
    ci_hi: float
    n_pos: int
    n_neg: int
    perm_p: float = np.nan      # filled in by permutation_auroc_pvalue if requested
    method: str = "bootstrap"

    def as_dict(self) -> dict:
        return asdict(self)


@dataclass
class EffectResult:
    statistic: float            # Mann-Whitney U
    p_value: float
    cliffs_delta: float
    delta_lo: float
    delta_hi: float
    n_x: int
    n_y: int

    def as_dict(self) -> dict:
        return asdict(self)


@dataclass
class ShiftResult:
    """Distribution comparison of a tumour sample vs a normal/healthy reference."""
    median_diff: float          # median(sample) - median(reference)
    cliffs_delta: float         # >0 means sample shifted ABOVE reference
    ks_statistic: float
    ks_p: float
    frac_above_ref_q: float     # fraction of sample cells above the reference q-quantile
    ref_threshold: float        # the reference quantile value used
    n_sample: int
    n_reference: int

    def as_dict(self) -> dict:
        return asdict(self)


# ---------------------------------------------------------------------------
# Safe metric wrappers (degenerate label vectors -> NaN, never an exception)
# ---------------------------------------------------------------------------
def auroc_safe(y: np.ndarray, p: np.ndarray) -> float:
    y = np.asarray(y); p = np.asarray(p)
    if len(np.unique(y)) < 2:
        return np.nan
    return float(roc_auc_score(y, p))


def auprc_safe(y: np.ndarray, p: np.ndarray) -> float:
    y = np.asarray(y); p = np.asarray(p)
    if len(np.unique(y)) < 2:
        return np.nan
    return float(average_precision_score(y, p))


# ---------------------------------------------------------------------------
# Stratified bootstrap CI for AUROC (cases & controls resampled separately)
# ---------------------------------------------------------------------------
def bootstrap_auroc_ci(
    y: Sequence[int],
    p: Sequence[float],
    n_boot: int = 2000,
    alpha: float = 0.05,
    stratified: bool = True,
    random_state: int = 42,
) -> AUROCResult:
    """Percentile bootstrap CI for AUROC.

    Stratified resampling preserves the case/control ratio in every replicate,
    which is the correct procedure for a fixed-prevalence discrimination metric.
    """
    y = np.asarray(y).astype(int)
    p = np.asarray(p, dtype=float)
    n_pos = int(y.sum())
    n_neg = int((1 - y).sum())
    point = auroc_safe(y, p)
    if np.isnan(point) or n_pos < 2 or n_neg < 2:
        return AUROCResult(point, np.nan, np.nan, n_pos, n_neg)

    rng = np.random.default_rng(random_state)
    pos_idx = np.where(y == 1)[0]
    neg_idx = np.where(y == 0)[0]
    boots = np.empty(n_boot)
    for b in range(n_boot):
        if stratified:
            bi = np.concatenate([
                rng.choice(pos_idx, size=n_pos, replace=True),
                rng.choice(neg_idx, size=n_neg, replace=True),
            ])
        else:
            bi = rng.choice(len(y), size=len(y), replace=True)
        yb, pb = y[bi], p[bi]
        boots[b] = auroc_safe(yb, pb)
    boots = boots[~np.isnan(boots)]
    lo = float(np.percentile(boots, 100 * alpha / 2))
    hi = float(np.percentile(boots, 100 * (1 - alpha / 2)))
    return AUROCResult(point, lo, hi, n_pos, n_neg)


# ---------------------------------------------------------------------------
# Fast DeLong (Sun & Xu 2014 midrank) — variance of one AUC and paired test
# ---------------------------------------------------------------------------
def _compute_midrank(x: np.ndarray) -> np.ndarray:
    J = np.argsort(x)
    z = x[J]
    N = len(x)
    t = np.zeros(N, dtype=float)
    i = 0
    while i < N:
        j = i
        while j < N and z[j] == z[i]:
            j += 1
        t[i:j] = 0.5 * (i + j - 1) + 1
        i = j
    out = np.empty(N, dtype=float)
    out[J] = t
    return out


def _fast_delong(predictions_sorted_transposed: np.ndarray, label_1_count: int):
    """Core fast DeLong; returns (aucs, covariance-matrix)."""
    m = label_1_count
    n = predictions_sorted_transposed.shape[1] - m
    positive = predictions_sorted_transposed[:, :m]
    negative = predictions_sorted_transposed[:, m:]
    k = predictions_sorted_transposed.shape[0]

    tx = np.empty([k, m]); ty = np.empty([k, n]); tz = np.empty([k, m + n])
    for r in range(k):
        tx[r, :] = _compute_midrank(positive[r, :])
        ty[r, :] = _compute_midrank(negative[r, :])
        tz[r, :] = _compute_midrank(predictions_sorted_transposed[r, :])
    aucs = tz[:, :m].sum(axis=1) / m / n - (m + 1.0) / 2.0 / n
    v01 = (tz[:, :m] - tx[:, :]) / n
    v10 = 1.0 - (tz[:, m:] - ty[:, :]) / m
    sx = np.cov(v01)
    sy = np.cov(v10)
    delongcov = sx / m + sy / n
    return aucs, np.atleast_2d(delongcov)


def _delong_prep(y: np.ndarray, *probs: np.ndarray):
    order = (-y).argsort(kind="mergesort")  # positives first
    label_1_count = int(y.sum())
    preds = np.vstack(probs)[:, order]
    return preds, label_1_count


def delong_auc_variance(y: Sequence[int], p: Sequence[float]) -> tuple[float, float]:
    """Return (AUC, variance) for a single predictor via DeLong."""
    y = np.asarray(y).astype(int)
    p = np.asarray(p, dtype=float)
    preds, m = _delong_prep(y, p)
    aucs, cov = _fast_delong(preds, m)
    return float(aucs[0]), float(cov[0, 0])


def delong_test(
    y: Sequence[int],
    p1: Sequence[float],
    p2: Sequence[float],
) -> dict:
    """Paired DeLong test that two AUCs (same samples) differ.

    Use this to compare scOPE against a baseline scored on the *identical*
    samples (single best marker gene, full-feature logistic, passenger control).
    Returns AUCs, their difference, a two-sided p-value, and the z statistic.
    """
    y = np.asarray(y).astype(int)
    p1 = np.asarray(p1, dtype=float)
    p2 = np.asarray(p2, dtype=float)
    if len(np.unique(y)) < 2:
        return dict(auc1=np.nan, auc2=np.nan, diff=np.nan, z=np.nan, p_value=np.nan)
    preds, m = _delong_prep(y, p1, p2)
    aucs, cov = _fast_delong(preds, m)
    var = cov[0, 0] + cov[1, 1] - 2 * cov[0, 1]
    diff = aucs[0] - aucs[1]
    if var <= 0:
        z = 0.0 if diff == 0 else np.sign(diff) * np.inf
        pval = 1.0 if diff == 0 else 0.0
    else:
        z = diff / np.sqrt(var)
        pval = 2.0 * _sps.norm.sf(abs(z))
    return dict(auc1=float(aucs[0]), auc2=float(aucs[1]), diff=float(diff),
                z=float(z), p_value=float(pval))


# ---------------------------------------------------------------------------
# Label-permutation null for AUROC (one-sided: AUROC > 0.5)
# ---------------------------------------------------------------------------
def permutation_auroc_pvalue(
    y: Sequence[int],
    p: Sequence[float],
    n_perm: int = 2000,
    random_state: int = 42,
) -> float:
    """Empirical one-sided p-value: P(AUROC_perm >= AUROC_obs) under label shuffle.

    Uses the (B + 1) correction so the p-value can never be exactly zero.
    """
    y = np.asarray(y).astype(int)
    p = np.asarray(p, dtype=float)
    obs = auroc_safe(y, p)
    if np.isnan(obs):
        return np.nan
    rng = np.random.default_rng(random_state)
    ge = 0
    for _ in range(n_perm):
        yp = rng.permutation(y)
        if auroc_safe(yp, p) >= obs:
            ge += 1
    return (ge + 1) / (n_perm + 1)


def permutation_label_null(
    score: Sequence[float],
    label: Sequence[int],
    statistic: str = "auroc",
    n_perm: int = 2000,
    random_state: int = 42,
    groups: Sequence | None = None,
) -> dict:
    """General label-permutation null for an arbitrary 2-group statistic.

    statistic: 'auroc' | 'cliffs' | 'median_diff'. Returns observed value,
    one-sided p (obs >= null), and the null mean/std for reporting.

    groups : optional per-observation grouping key (e.g. patient id). When the
    label is a GROUP-LEVEL property — constant within each group, as a per-patient
    mutation status is — pass ``groups`` to permute the label at the GROUP level:
    each group keeps its block of observations and the group->label assignment is
    shuffled. This is the anti-pseudoreplication null. It fixes the number of
    positive *patients*, not the number of positive *cells*, so a per-cell
    statistic that is really driven by one or two patients no longer reads as
    significant — which is exactly the cell-vs-patient discrepancy that otherwise
    shows up between this null and ``patient_bridge``. The returned dict carries
    ``n_perm_unit`` ('observation' or 'group') so the manuscript can state the
    unit of permutation explicitly. Default (groups=None) is the original
    observation-level shuffle and is bit-for-bit unchanged.

    Raises ValueError if ``groups`` is given but the label is not constant within
    every group (a within-group-varying label, e.g. a tumour/reference compartment
    flag, is not a group-level property and block permutation would be ill-defined
    — aggregate first or use a within-group restricted design instead).
    """
    score = np.asarray(score, dtype=float)
    label = np.asarray(label).astype(int)

    def stat(s, lab):
        if statistic == "auroc":
            return auroc_safe(lab, s)
        a, b = s[lab == 1], s[lab == 0]
        if len(a) == 0 or len(b) == 0:
            return np.nan
        if statistic == "cliffs":
            return _cliffs_point(a, b)
        if statistic == "median_diff":
            return float(np.median(a) - np.median(b))
        raise ValueError(statistic)

    rng = np.random.default_rng(random_state)

    if groups is None:
        unit = "observation"
        def draw_labels():
            return rng.permutation(label)
    else:
        unit = "group"
        groups = np.asarray(groups)
        uniq = np.unique(groups)
        gidx = [np.where(groups == g)[0] for g in uniq]
        for ix in gidx:
            if np.unique(label[ix]).size > 1:
                raise ValueError(
                    "permutation_label_null(groups=...) requires the label to be "
                    "constant within each group (a group-level property such as a "
                    "per-patient mutation status); a group with mixed labels was found.")
        glabel = np.array([label[ix[0]] for ix in gidx])
        def draw_labels():
            out = np.empty_like(label)
            for ix, lab in zip(gidx, rng.permutation(glabel)):
                out[ix] = lab
            return out

    obs = stat(score, label)
    if np.isnan(obs):
        return dict(observed=np.nan, perm_p=np.nan, null_mean=np.nan,
                    null_std=np.nan, n_perm_unit=unit)
    null = np.empty(n_perm)
    for i in range(n_perm):
        null[i] = stat(score, draw_labels())
    null = null[~np.isnan(null)]
    ge = int(np.sum(null >= obs))
    return dict(observed=float(obs), perm_p=(ge + 1) / (len(null) + 1),
                null_mean=float(null.mean()), null_std=float(null.std()),
                n_perm_unit=unit)


# ---------------------------------------------------------------------------
# Benjamini-Hochberg FDR (vectorised, NaN-safe)
# ---------------------------------------------------------------------------
def bh_fdr(pvals: Sequence[float]) -> np.ndarray:
    """Benjamini-Hochberg q-values. NaNs pass through as NaN.

    Apply this ONCE per analysis stage, across every test in that stage, so the
    manuscript's multiple-testing correction is honest and global rather than
    per-figure.
    """
    p = np.asarray(pvals, dtype=float)
    out = np.full_like(p, np.nan)
    mask = ~np.isnan(p)
    pm = p[mask]
    n = len(pm)
    if n == 0:
        return out
    order = np.argsort(pm)
    ranked = pm[order]
    q = ranked * n / (np.arange(n) + 1)
    q = np.minimum.accumulate(q[::-1])[::-1]
    q = np.clip(q, 0, 1)
    filled = np.empty(n)
    filled[order] = q
    out[mask] = filled
    return out


# ---------------------------------------------------------------------------
# Cliff's delta (non-parametric effect size) + bootstrap CI
# ---------------------------------------------------------------------------
def _cliffs_point(x: np.ndarray, y: np.ndarray) -> float:
    x = np.asarray(x, dtype=float); y = np.asarray(y, dtype=float)
    # delta = (#(x>y) - #(x<y)) / (nx*ny), computed via rank trick for O(n log n)
    nx, ny = len(x), len(y)
    if nx == 0 or ny == 0:
        return np.nan
    combined = np.concatenate([x, y])
    ranks = _sps.rankdata(combined)
    rx = ranks[:nx].sum()
    # Mann-Whitney U for x
    U = rx - nx * (nx + 1) / 2.0
    return float(2.0 * U / (nx * ny) - 1.0)


def cliffs_delta(
    x: Sequence[float],
    y: Sequence[float],
    n_boot: int = 0,
    random_state: int = 42,
) -> EffectResult | float:
    """Cliff's delta between groups x and y.

    delta > 0  ->  x tends to exceed y.
    |delta|:  <0.147 negligible, <0.33 small, <0.474 medium, else large
    (Romano et al. thresholds). If n_boot>0, returns an EffectResult with a CI;
    otherwise returns the bare float (used inside permutation nulls).
    """
    point = _cliffs_point(np.asarray(x), np.asarray(y))
    if n_boot <= 0:
        return point
    rng = np.random.default_rng(random_state)
    x = np.asarray(x, dtype=float); y = np.asarray(y, dtype=float)
    boots = np.empty(n_boot)
    for b in range(n_boot):
        xb = rng.choice(x, len(x), replace=True)
        yb = rng.choice(y, len(y), replace=True)
        boots[b] = _cliffs_point(xb, yb)
    lo, hi = np.percentile(boots, [2.5, 97.5])
    U, p = _mwu(x, y)
    return EffectResult(float(U), float(p), float(point), float(lo), float(hi),
                        len(x), len(y))


def _mwu(x, y):
    try:
        res = _sps.mannwhitneyu(x, y, alternative="two-sided")
        return res.statistic, res.pvalue
    except ValueError:
        return np.nan, np.nan


def mannwhitney_effect(
    x: Sequence[float],
    y: Sequence[float],
    n_boot: int = 2000,
    random_state: int = 42,
) -> EffectResult:
    """Mann-Whitney U test + Cliff's delta with bootstrap CI in one call."""
    res = cliffs_delta(x, y, n_boot=n_boot, random_state=random_state)
    if isinstance(res, EffectResult):
        return res
    U, p = _mwu(x, y)
    return EffectResult(float(U), float(p), float(res), np.nan, np.nan, len(x), len(y))


# ---------------------------------------------------------------------------
# Distribution shift vs a healthy/normal reference (the timepoint workhorse)
# ---------------------------------------------------------------------------
def distribution_shift(
    sample_scores: Sequence[float],
    reference_scores: Sequence[float],
    reference_quantile: float = 0.95,
) -> ShiftResult:
    """Quantify how far a tumour sample's score distribution sits ABOVE a
    healthy/normal reference distribution.

    This is the quantitative backbone of the DNMT3A / NPM1 "healthy BM vs tumour"
    density plots, and of the longitudinal treatment-response trajectory: call it
    per (patient, timepoint, driver) with the same fixed reference each time and
    watch ``cliffs_delta`` / ``frac_above_ref_q`` decay as treatment clears the
    mutant clone.
    """
    s = np.asarray(sample_scores, dtype=float)
    r = np.asarray(reference_scores, dtype=float)
    s = s[~np.isnan(s)]; r = r[~np.isnan(r)]
    if len(s) == 0 or len(r) == 0:
        # Degenerate/empty-group path. Keep this robust to ShiftResult field-count changes.
        _n_fields = (
            len(getattr(ShiftResult, "__dataclass_fields__", {}))
            or len(getattr(ShiftResult, "_fields", []))
            or 8
        )
        return ShiftResult(*([np.nan] * max(0, _n_fields - 2)), len(s), len(r))
    thr = float(np.quantile(r, reference_quantile))
    try:
        ks_stat, ks_p = _sps.ks_2samp(s, r)
    except ValueError:
        ks_stat, ks_p = np.nan, np.nan
    return ShiftResult(
        median_diff=float(np.median(s) - np.median(r)),
        cliffs_delta=_cliffs_point(s, r),
        ks_statistic=float(ks_stat),
        ks_p=float(ks_p),
        frac_above_ref_q=float(np.mean(s > thr)),
        ref_threshold=thr,
        n_sample=len(s),
        n_reference=len(r),
    )


# ---------------------------------------------------------------------------
# Patient-level aggregation (the anti-pseudoreplication discipline)
# ---------------------------------------------------------------------------
def patient_aggregate(
    cell_df: pd.DataFrame,
    patient_col: str,
    score_col: str,
    agg: str = "median",
    min_cells: int = 10,
) -> pd.DataFrame:
    """Collapse per-cell scores to one value per patient.

    Thousands of cells from one patient are NOT independent biological
    replicates; treating them as such inflates significance by orders of
    magnitude. Every cross-patient inference in this suite runs on the output of
    this function, with the patient as the unit of replication.

    Returns columns: [patient_col, score_col, 'n_cells'].
    """
    grp = cell_df.groupby(patient_col)[score_col]
    agg_fn = {"median": grp.median, "mean": grp.mean}[agg]
    out = agg_fn().to_frame(score_col)
    out["n_cells"] = grp.size()
    out = out[out["n_cells"] >= min_cells].reset_index()
    return out


def concordance_kappa(call_a, call_b, n_boot: int = 2000, seed: int = 0) -> dict:
    """Agreement between two binary malignancy calls (e.g. scOPE vs infercnvpy CNV)
    BEYOND chance: Cohen's kappa with a bootstrap CI, plus sensitivity/specificity of
    A treating B as reference, and the 2x2 counts. Reporting kappa (not raw % agree)
    is the right move for a reviewer because it corrects for the high base-rate
    agreement you get when most cells are non-malignant.
    """
    a = np.asarray(call_a).astype(float); b = np.asarray(call_b).astype(float)
    ok = np.isfinite(a) & np.isfinite(b)
    a, b = a[ok].astype(int), b[ok].astype(int)
    n = len(a)
    if n < 10 or len(np.unique(np.r_[a, b])) < 2:
        return {"n": int(n), "kappa": np.nan, "note": "insufficient data/variation"}

    def _kappa(a, b):
        po = np.mean(a == b)
        pa1 = np.mean(a); pb1 = np.mean(b)
        pe = pa1 * pb1 + (1 - pa1) * (1 - pb1)
        return (po - pe) / (1 - pe) if pe < 1 else np.nan

    k = _kappa(a, b)
    tp = int(np.sum((a == 1) & (b == 1))); tn = int(np.sum((a == 0) & (b == 0)))
    fp = int(np.sum((a == 1) & (b == 0))); fn = int(np.sum((a == 0) & (b == 1)))
    sens = tp / (tp + fn) if (tp + fn) else np.nan          # A recall vs B
    spec = tn / (tn + fp) if (tn + fp) else np.nan
    rng = np.random.default_rng(seed)
    boot = []
    for _ in range(n_boot):
        idx = rng.integers(0, n, n)
        boot.append(_kappa(a[idx], b[idx]))
    boot = np.array([x for x in boot if np.isfinite(x)])
    lo, hi = (np.percentile(boot, [2.5, 97.5]) if len(boot) else (np.nan, np.nan))
    return {"n": int(n), "kappa": float(k), "kappa_lo": float(lo), "kappa_hi": float(hi),
            "percent_agree": float(np.mean(a == b)),
            "sensitivity_a_vs_b": float(sens), "specificity_a_vs_b": float(spec),
            "tp": tp, "tn": tn, "fp": fp, "fn": fn}
```
