## Supplemental_Code for "scOPE identifies which driver-associated expression programs transfer from bulk tumors to single cells": umap_panels.py

```
"""
scope_analysis.umap_panels
===========================
High-quality multi-panel UMAP figures for single-cell cohorts.

For each driver this renders a row of three UMAP panels that all share the SAME
embedding, each with its own legend / colourbar:

    [ patient ]   [ ground-truth mutation (red = Mut) ]   [ driver-program score ]

so you can read, at a glance: which patient a region belongs to, where the cells
that REALLY carry the mutation sit (where genotype exists), and whether the scOPE
prediction lights up the same place. Driver-level confidence is annotated on the
score panel. Small, rasterised points keep dense embeddings legible.

Also provides a per-cohort context strip (patient / cell type / malignant
compartment) and a pure prediction-score grid for drivers without genotype.

Nothing here re-projects or re-scores; it only reads obs the pipeline already wrote
(mutation_calib_* / mutation_prob_*, a expressed-mutation annotation column where present,
patient, cell type, scope_malignant). Use the standalone driver
`drivers/make_umap_panels.py` to render from a scored h5ad without a full run.
"""
from __future__ import annotations

from pathlib import Path
import re

import numpy as np
import pandas as pd
import matplotlib
matplotlib.use("Agg")
import matplotlib.pyplot as plt
from matplotlib.lines import Line2D


# ---------------------------------------------------------------------------
# helpers
# ---------------------------------------------------------------------------
_TRUTH_RE = re.compile(r"(genotype|mutant|mutation|status|variant|call|label)", re.I)
_PRED_RE = re.compile(r"(mutation_prob|mutation_calib|_prob|_score|^score|pred|probability)", re.I)
# scOPE's own derived SCORE columns all live in the 'mutation_<kind>_<gene>' namespace.
# These must never be mistaken for a expressed-mutation annotation (the bug that made the triptych
# binarize a continuous residual/percentile score as if it were truth, and bypass the
# patient-level mutation panel entirely).
_SCORE_NAMESPACE = re.compile(
    r"^mutation_(prob|calib|calibrated|resid|residual|refpct|pct|adj|adjusted|"
    r"gated|gatedadj|score|conf|confidence)\b", re.I)

_BOOL_COLORS = {
    "true": "#d73027", "1": "#d73027", "mut": "#d73027", "mutant": "#d73027", "malignant": "#d73027",
    "false": "#4daf4a", "0": "#4daf4a", "wt": "#4daf4a", "wild-type": "#4daf4a", "non-malignant": "#4daf4a",
    "unknown": "#bdbdbd", "nan": "#bdbdbd", "none": "#bdbdbd",
}
_TWO_CAT_FALLBACK = ["#d73027", "#4daf4a", "#984ea3"]


def _looks_like_genotype(series: pd.Series) -> bool:
    """True if a column's values look like a discrete genotype call, not a continuous
    score. A genuine genotype binarizes to {0,1,nan} with both classes present and has
    only a handful of distinct raw values; a score has many distinct floats in (0,1)."""
    s = pd.Series(series)
    nun = s.dropna().nunique()
    if nun == 0 or nun > 12:           # continuous score -> many distinct values
        return False
    try:
        from .singlecell import _binarize_genotype
        b = _binarize_genotype(s)
        vals = set(np.unique(b.values[~pd.isna(b.values)]))
        return vals.issubset({0.0, 1.0}) and len({0.0, 1.0} & vals) >= 1
    except Exception:
        return True


def find_genotype_column(adata, driver: str) -> str | None:
    """obs column holding a genuine expressed-mutation annotation for `driver` (not a score).

    Excludes the scOPE score namespace (mutation_prob/calib/resid/refpct/adj/gated/...)
    and any column whose values look continuous, so derived scores are never treated as
    ground truth. Returns None when only scores exist — which lets the triptych fall
    back to the patient-level mutation panel."""
    cand = []
    dl = driver.lower()
    for c in adata.obs.columns:
        cl = str(c).lower()
        if dl not in cl:
            continue
        if not _TRUTH_RE.search(c) or _PRED_RE.search(c) or _SCORE_NAMESPACE.search(cl):
            continue
        cand.append(c)
    # prefer the most genotype-looking name, then verify values are discrete
    cand.sort(key=lambda c: (0 if re.search(r"genotype|variant", str(c), re.I) else 1, len(str(c))))
    for c in cand:
        if _looks_like_genotype(adata.obs[c]):
            return c
    return None


def binarize_genotype(series: pd.Series) -> np.ndarray:
    """-> array in {1 mutant, 0 WT, nan unknown}. Delegates to the canonical
    singlecell._binarize_genotype so the UMAP panels and the Fig-3 validation always
    agree on what counts as Mut/WT (string, int, float, and boolean encodings)."""
    from .singlecell import _binarize_genotype
    return np.asarray(_binarize_genotype(series).values, dtype=float)


def patient_level_genotype(adata, patient_status: pd.DataFrame | None, driver: str,
                           patient_col: str = "patient") -> np.ndarray | None:
    """Return per-cell labels from a tidy patient-status table.

    Used for non-AML cohorts where the middle UMAP panel should be patient-level
    mutation status, not fake expressed-mutation annotation. Values are {1 Mut-patient,
    0 WT-patient, NaN unknown/unmatched}.
    """
    if patient_status is None or patient_col not in adata.obs.columns:
        return None
    try:
        from .data import is_usable_patient_series
        if not is_usable_patient_series(adata.obs[patient_col], colname=patient_col):
            return None
    except Exception:
        pass
    if not {"patient", "mutation", "label"}.issubset(patient_status.columns):
        return None
    status = patient_status[patient_status["mutation"].astype(str).str.upper() == str(driver).upper()]
    if status.empty:
        return None
    labels = pd.to_numeric(status["label"], errors="coerce")
    # A patient may have several samples (e.g. CRC SMC01-T / SMC01-N) that harmonized to
    # the same obs patient. Aggregate to one call per patient: mutant if ANY sample is
    # mutant, else WT if any sample is WT, else unknown. (scRNA cells are the tumour,
    # so a positive call in any tumour sample means the patient's tumour carries it.)
    agg = (pd.DataFrame({"patient": status["patient"].astype(str), "label": labels})
           .groupby("patient")["label"]
           .agg(lambda s: 1.0 if (s == 1).any() else (0.0 if (s == 0).any() else np.nan)))
    lut = agg.to_dict()
    obs_pat = adata.obs[patient_col].astype(str)
    vals = obs_pat.map(lut).astype(float)
    # Loose punctuation/case-insensitive + sample-suffix fallback fills only still-
    # unmatched cells, so exact matches are preserved but SMC01 still joins SMC01-T.
    import re as _re
    def _key(x): return _re.sub(r"[^A-Za-z0-9]+", "", str(x).strip()).upper()
    try:
        from .data import _base_key as _bk
    except Exception:
        def _bk(x): return _key(x)
    lut2 = {_key(k): v for k, v in lut.items()}
    lut3 = {_bk(k): v for k, v in lut.items()}
    loose = obs_pat.map(lambda x: lut2.get(_key(x), lut3.get(_bk(x), np.nan)))
    vals = vals.where(vals.notna(), loose)
    if vals.notna().any():
        return vals.astype(float).values
    return None


def triptych_genotype_audit(adata, drivers, patient_status: pd.DataFrame | None = None,
                            patient_col: str = "patient") -> pd.DataFrame:
    """Audit exactly what the triptych middle panel will use for each driver.

    This is intentionally generated before rendering so silent failures are obvious:
    source is one of ``per-cell``, ``patient``, or ``prior/none``; counts are computed
    from the actual per-cell vector that will be plotted. It also catches the old bug
    where a barcode-like patient column prevented patient-level mutation overlays.
    """
    rows = []
    obs_pat = adata.obs[patient_col].astype(str) if patient_col in adata.obs.columns else None
    patient_usable = False
    if obs_pat is not None:
        try:
            from .data import is_usable_patient_series
            patient_usable = bool(is_usable_patient_series(obs_pat, colname=patient_col))
        except Exception:
            patient_usable = True
    status_has_cols = patient_status is not None and {"patient", "mutation", "label"}.issubset(patient_status.columns)
    for d in drivers:
        gcol = find_genotype_column(adata, d)
        pgeno = patient_level_genotype(adata, patient_status, d, patient_col=patient_col)
        if gcol is not None:
            geno = binarize_genotype(adata.obs[gcol])
            source = "per-cell"
        elif pgeno is not None:
            geno = pgeno
            source = "patient"
        else:
            geno = np.full(adata.n_obs, np.nan)
            source = "prior/none"
        finite = np.isfinite(geno)
        mut = int(np.sum(geno == 1))
        wt = int(np.sum(geno == 0))
        unk = int(np.sum(~finite))
        status_rows = 0
        status_patients = 0
        status_overlap_patients = 0
        status_mut_patients = 0
        status_wt_patients = 0
        if status_has_cols:
            ss = patient_status[patient_status["mutation"].astype(str).str.upper() == str(d).upper()].copy()
            status_rows = int(len(ss))
            if not ss.empty:
                status_patients = int(ss["patient"].astype(str).nunique())
                lab = pd.to_numeric(ss["label"], errors="coerce")
                status_mut_patients = int(ss.loc[lab == 1, "patient"].astype(str).nunique())
                status_wt_patients = int(ss.loc[lab == 0, "patient"].astype(str).nunique())
                if obs_pat is not None:
                    status_overlap_patients = int(len(set(ss["patient"].astype(str)) & set(obs_pat.dropna().astype(str).unique())))
        rows.append({
            "driver": d,
            "source": source,
            "genotype_col": gcol,
            "patient_col": patient_col if patient_col in adata.obs.columns else None,
            "patient_col_usable": bool(patient_usable),
            "patient_status_rows": status_rows,
            "patient_status_patients": status_patients,
            "patient_status_overlap_patients": status_overlap_patients,
            "patient_status_mut_patients": status_mut_patients,
            "patient_status_wt_patients": status_wt_patients,
            "n_mut_cells_plotted": mut,
            "n_wt_cells_plotted": wt,
            "n_unknown_unmatched_cells": unk,
            "n_cells": int(adata.n_obs),
        })
    return pd.DataFrame(rows)


def _embedding(adata, prefer=("X_umap", "X_umap_hvg3000", "X_umap_hvg2000")):
    for k in prefer:
        if k in adata.obsm:
            return np.asarray(adata.obsm[k])[:, :2]
    for k in adata.obsm:
        if k.lower().startswith("x_umap"):
            return np.asarray(adata.obsm[k])[:, :2]
    # last resort: compute a quick one
    try:
        import scanpy as sc
        a = adata.copy()
        rep = next((r for r in ("X_svd", "X_pca") if r in a.obsm), None)
        if rep is None:
            sc.pp.pca(a, n_comps=min(50, a.n_vars - 1, a.n_obs - 1)); rep = "X_pca"
        sc.pp.neighbors(a, use_rep=rep); sc.tl.umap(a)
        adata.obsm["X_umap"] = a.obsm["X_umap"]
        return np.asarray(a.obsm["X_umap"])[:, :2]
    except Exception:
        raise ValueError("No UMAP embedding found and none could be computed.")


def _point_size(n: int) -> float:
    return float(np.clip(6000.0 / max(n, 1), 0.35, 3.0))


def _score_kind_from_col(col: str, cfg=None) -> str:
    if cfg is not None and hasattr(cfg, "score_kind"):
        return cfg.score_kind(col)
    col = str(col)
    if "mutation_calib_" in col:
        return "calibrated_reference_percentile"
    if "mutation_prob_" in col:
        return "raw_probability"
    if "mutation_adj_" in col:
        return "confidence_adjusted_raw"
    if "mutation_resid_" in col:
        return "cellstate_residual_raw"
    if "mutation_refpct_" in col:
        return "cellstate_reference_percentile"
    return "score"


def _robust_limits(values, cfg=None, col: str | None = None):
    """Return plot-only color limits plus a short scale note.

    Same-cell-state residuals need special handling: weak/exploratory drivers often
    have microscopic numeric ranges.  Per-panel q01-q99 scaling makes that noise look
    bright.  For residual columns we therefore use a symmetric color scale around
    zero with a configurable minimum span.  Statistics are never affected.
    """
    x = np.asarray(values, dtype=float)
    x = x[np.isfinite(x)]
    if x.size == 0:
        return 0.0, 1.0, ""
    kind = _score_kind_from_col(col or "", cfg)
    if kind == "cellstate_residual_raw":
        uq = float(getattr(cfg, "plot_robust_upper_q", 0.99)) if cfg is not None else 0.99
        min_abs = float(getattr(cfg, "plot_residual_min_abs_vmax", 0.05)) if cfg is not None else 0.05
        max_abs = float(getattr(cfg, "plot_residual_abs_vmax", 0.25)) if cfg is not None else 0.25
        try:
            vmax = float(np.nanquantile(np.abs(x), uq))
        except Exception:
            vmax = float(np.nanmax(np.abs(x)))
        if not np.isfinite(vmax) or vmax <= 0:
            vmax = min_abs
        vmax = min(max(vmax, min_abs), max_abs)
        return -vmax, vmax, f"centered ±{vmax:g} color scale"
    if kind in {"calibrated_reference_percentile", "cellstate_reference_percentile"}:
        return 0.0, 1.0, ""
    mode = getattr(cfg, "plot_score_scale", "auto_robust") if cfg is not None else "auto_robust"
    mode_l = str(mode).lower()
    if mode_l in {"fixed01", "probability", "0_1"}:
        return 0.0, 1.0, "fixed 0-1 scale"
    if mode_l in {"absolute", "none", "fixed"}:
        lo, hi = float(np.nanmin(x)), float(np.nanmax(x))
        if hi <= lo: hi = lo + 1e-9
        return lo, hi, ""
    lq = float(getattr(cfg, "plot_robust_lower_q", 0.01)) if cfg is not None else 0.01
    uq = float(getattr(cfg, "plot_robust_upper_q", 0.99)) if cfg is not None else 0.99
    min_range = float(getattr(cfg, "plot_robust_min_range", 0.05)) if cfg is not None else 0.05
    lo, hi = np.nanquantile(x, [lq, uq])
    if not np.isfinite(lo) or not np.isfinite(hi) or hi <= lo:
        lo, hi = float(np.nanmin(x)), float(np.nanmax(x))
    if hi <= lo:
        hi = lo + 1e-9
    full_range = float(np.nanmax(x) - np.nanmin(x))
    compressed = (full_range < min_range) or (float(np.nanmax(x)) < 0.15 and kind == "raw_probability")
    if compressed:
        return float(lo), float(hi), f"q{int(lq*100):02d}-q{int(uq*100):02d} display scale"
    lo, hi = float(np.nanmin(x)), float(np.nanmax(x))
    if kind == "raw_probability" and lo >= 0 and hi <= 1:
        return 0.0, 1.0, ""
    return lo, hi, ""


def _score_col(adata, driver, cfg=None, prefer=("adj", "raw", "calib")):
    """Resolve which per-cell score column to plot.

    Default preference is ADJUSTED (raw x confidence) -> RAW probability -> calibrated.
    RAW/adjusted are preferred over the reference-ECDF *calibrated* score on purpose:
    calibration transforms each cell by the reference distribution's own CDF, which
    forces the reference cells to spread ~uniformly over [0,1] (so ~half of them exceed
    0.5 *by construction*) and makes healthy cells look falsely 'lit up'. Raw scores
    don't have that artifact, so the panel reflects model output rather than the
    calibration transform.
    """
    name = {"adj": f"mutation_adj_{driver}", "raw": f"mutation_prob_{driver}",
            "calib": f"mutation_calib_{driver}", "resid": f"mutation_resid_{driver}",
            "refpct": f"mutation_refpct_{driver}", "guarded": f"mutation_resid_{driver}"}
    for kind in prefer:
        c = name[kind]
        if c in adata.obs.columns:
            return c, kind
    if cfg is not None:
        try:
            c = cfg.score_col(adata, driver)
            return c, cfg.score_kind(c)
        except Exception:
            pass
    return None, None


def _blank(ax):
    ax.set_xticks([]); ax.set_yticks([])
    for sp in ax.spines.values():
        sp.set_visible(False)


# ---------------------------------------------------------------------------
# single panels
# ---------------------------------------------------------------------------
def _panel_patient(ax, emb, labels, s, max_legend=18):
    labels = pd.Series(labels).astype(str).values
    uniq = pd.Index(pd.unique(labels))
    cmap = plt.get_cmap("tab20")
    colors = {u: cmap(i % 20) for i, u in enumerate(uniq)}
    cvec = np.array([colors[u] for u in labels])
    order = np.random.default_rng(0).permutation(len(emb))   # avoid one patient on top
    ax.scatter(emb[order, 0], emb[order, 1], c=cvec[order], s=s, edgecolor="none",
               rasterized=True, linewidths=0)
    _blank(ax); ax.set_title(f"patient  (n={len(uniq)})", fontsize=11)
    show = list(uniq[:max_legend])
    handles = [Line2D([0], [0], marker="o", color="w", markerfacecolor=colors[u],
                      markersize=6, label=str(u)) for u in show]
    if len(uniq) > max_legend:
        handles.append(Line2D([0], [0], marker="o", color="w", markerfacecolor="grey",
                              markersize=6, label=f"+{len(uniq)-max_legend} more"))
    ax.legend(handles=handles, fontsize=6, loc="center left", bbox_to_anchor=(1.0, 0.5),
              frameon=False, handletextpad=0.2, labelspacing=0.25)


def _panel_genotype(ax, emb, geno, driver, s, label_kind="expressed-mutation annotation"):
    n_mut = int(np.nansum(geno == 1)); n_wt = int(np.nansum(geno == 0))
    unknown = ~np.isfinite(geno)
    # background: unknown (very light), WT (light blue), Mut (red) on top
    ax.scatter(emb[unknown, 0], emb[unknown, 1], c="#eeeeee", s=s, edgecolor="none",
               rasterized=True, linewidths=0)
    wt = geno == 0
    ax.scatter(emb[wt, 0], emb[wt, 1], c="#9ecae1", s=s, edgecolor="none",
               rasterized=True, linewidths=0)
    mut = geno == 1
    ax.scatter(emb[mut, 0], emb[mut, 1], c="#d73027", s=max(s * 2.2, 3.0),
               edgecolor="none", rasterized=True, linewidths=0, zorder=5)
    _blank(ax); ax.set_title(f"{driver}  {label_kind}", fontsize=11)
    handles = [Line2D([0], [0], marker="o", color="w", markerfacecolor="#d73027",
                      markersize=7, label=f"Mut (n={n_mut})"),
               Line2D([0], [0], marker="o", color="w", markerfacecolor="#9ecae1",
                      markersize=6, label=f"WT (n={n_wt})"),
               Line2D([0], [0], marker="o", color="w", markerfacecolor="#eeeeee",
                      markersize=6, label=f"unknown/unmatched (n={int(unknown.sum())})")]
    ax.legend(handles=handles, fontsize=7, loc="center left", bbox_to_anchor=(1.0, 0.5),
              frameon=False, handletextpad=0.2, labelspacing=0.3)


def _panel_score(ax, emb, score, driver, s, confidence=None, cmap="magma",
                 ref_mask=None, score_label="raw score", cfg=None, score_col=None):
    """Score panel. If ref_mask is given, reference cells are drawn as light-grey
    background (the model is out-of-distribution there and the panel should make no
    claim), and the colour scale + draw order are computed on the NON-reference
    (malignant-compartment) cells, so genuine Mut-vs-WT contrast isn't compressed by
    lit-up reference cells."""
    score = np.asarray(score, dtype=float)
    if ref_mask is None:
        ref_mask = np.zeros(len(score), dtype=bool)
    ref_mask = np.asarray(ref_mask, dtype=bool)
    fg = ~ref_mask
    # background: reference cells, no score claim
    if ref_mask.any():
        ax.scatter(emb[ref_mask, 0], emb[ref_mask, 1], c="#e8e8e8", s=s,
                   edgecolor="none", rasterized=True, linewidths=0, zorder=1)
    fg_score = score[fg]
    vmin, vmax, scale_note = _robust_limits(fg_score, cfg=cfg, col=score_col)
    idx = np.where(fg)[0]
    order = idx[np.argsort(np.nan_to_num(score[idx], nan=-1))]   # high on top
    if _score_kind_from_col(score_col or "", cfg) == "cellstate_residual_raw":
        cmap = "coolwarm"
    scat = ax.scatter(emb[order, 0], emb[order, 1], c=score[order], s=s, cmap=cmap,
                      vmin=vmin, vmax=vmax, edgecolor="none", rasterized=True,
                      linewidths=0, zorder=3)
    _blank(ax)
    title = f"{driver}  driver-program score"
    if scale_note:
        title += f"\n{scale_note}"
    if confidence is not None and np.isfinite(confidence):
        title += f"   (conf={confidence:.2f})"
    ax.set_title(title, fontsize=11)
    cb = plt.colorbar(scat, ax=ax, fraction=0.046, pad=0.04)
    cb.ax.tick_params(labelsize=7)
    lbl = score_label + ("  (reference greyed)" if ref_mask.any() else "")
    cb.set_label(lbl, fontsize=8)


def _panel_categorical(ax, emb, labels, s, title, max_legend=14):
    labels = pd.Series(labels).astype(str).values
    counts = pd.Series(labels).value_counts()
    uniq = pd.Index(list(counts.index))
    if len(uniq) <= 3:
        colors = {}
        for i, u in enumerate(uniq):
            key = str(u).strip().lower()
            colors[u] = _BOOL_COLORS.get(key, _TWO_CAT_FALLBACK[i % len(_TWO_CAT_FALLBACK)])
    else:
        cmap = plt.get_cmap("tab20")
        colors = {u: cmap(i % 20) for i, u in enumerate(uniq)}
    cvec = np.array([colors[u] for u in labels])
    order = np.random.default_rng(1).permutation(len(emb))
    ax.scatter(emb[order, 0], emb[order, 1], c=cvec[order], s=s, edgecolor="none",
               rasterized=True, linewidths=0)
    _blank(ax); ax.set_title(title, fontsize=11)
    show = list(uniq[:max_legend])
    handles = [Line2D([0], [0], marker="o", color="w", markerfacecolor=colors[u],
                      markersize=6, label=str(u)) for u in show]
    if len(uniq) > max_legend:
        handles.append(Line2D([0], [0], marker="o", color="w", markerfacecolor="grey",
                              markersize=6, label=f"+{len(uniq)-max_legend} more"))
    ax.legend(handles=handles, fontsize=6.5, loc="center left", bbox_to_anchor=(1.0, 0.5),
              frameon=False, handletextpad=0.2, labelspacing=0.25)


# ---------------------------------------------------------------------------
# public figures
# ---------------------------------------------------------------------------
def driver_triptych_grid(adata, drivers, outpath, cfg=None, confidence=None,
                         patient_col="patient", only_genotyped=True, max_rows=14,
                         dpi=300, mask_reference=True, reference_col="is_reference",
                         score_prefer=("adj", "raw", "calib"),
                         patient_status: pd.DataFrame | None = None,
                         cancer_type: str | None = None) -> Path | None:
    """One row per driver: [patient | mutation annotation/status | driver-program score].
    By default the score panel plots RAW/adjusted (not reference-calibrated) scores and
    greys out reference cells, so it shows where the model actually fires among the
    malignant compartment rather than the reference-uniform calibration artifact.
    Rows are restricted to drivers with a expressed-mutation annotation column by default
    (set only_genotyped=False to include all scored drivers, genotype panel blank)."""
    emb = _embedding(adata)
    s = _point_size(adata.n_obs)
    conf = dict(confidence) if confidence is not None else {}
    use_patient_panel = False
    if patient_col in adata.obs.columns:
        try:
            from .data import is_usable_patient_series
            use_patient_panel = is_usable_patient_series(adata.obs[patient_col], colname=patient_col)
        except Exception:
            use_patient_panel = True
    pat = adata.obs[patient_col].values if use_patient_panel else np.array(["patient metadata unavailable"] * adata.n_obs)
    ref_mask = (adata.obs[reference_col].astype(bool).values
                if (mask_reference and reference_col in adata.obs.columns) else None)
    _label = {"adj": "adjusted score (raw x conf)", "raw": "raw probability",
              "calib": "calibrated/reference-percentile score",
              "resid": "same-cell-state residual",
              "refpct": "same-cell-state reference percentile",
              "guarded": "same-cell-state residual",
              "raw_probability": "raw probability",
              "calibrated_reference_percentile": "calibrated/reference-percentile score",
              "confidence_adjusted_raw": "adjusted score (raw x conf)",
              "cellstate_residual_raw": "same-cell-state residual",
              "cellstate_reference_percentile": "same-cell-state reference percentile"}

    rows = []
    for d in drivers:
        gcol = find_genotype_column(adata, d)
        pgeno = patient_level_genotype(adata, patient_status, d, patient_col=patient_col)
        scol, kind = _score_col(adata, d, cfg, prefer=score_prefer)
        if scol is None:
            continue
        if only_genotyped and gcol is None and pgeno is None:
            continue
        rows.append((d, gcol, pgeno, scol, kind))
    if not rows:
        return None
    rows = rows if max_rows is None else rows[:max_rows]
    nrow = len(rows)
    fig, axes = plt.subplots(nrow, 3, figsize=(15.5, 4.3 * nrow), squeeze=False)
    for i, (d, gcol, pgeno, scol, kind) in enumerate(rows):
        _panel_patient(axes[i][0], emb, pat, s)
        if gcol:
            geno = binarize_genotype(adata.obs[gcol])
            label_kind = "expressed-mutation annotation"
        elif pgeno is not None:
            geno = pgeno
            label_kind = "patient mutation status"
        else:
            geno = np.full(adata.n_obs, np.nan)
            # No per-patient call: annotate with the cancer-type prior (which mutations
            # to EXPECT in this cancer type). Cells stay 'unknown' — we never fabricate
            # a per-patient Mut/WT call from a population prevalence.
            try:
                from . import priors
                label_kind = priors.expected_label(cancer_type, d)
            except Exception:
                label_kind = "mutation status (no per-patient genotype)"
        _panel_genotype(axes[i][1], emb, geno, d, s, label_kind=label_kind)
        _panel_score(axes[i][2], emb, adata.obs[scol].astype(float).values, d, s,
                     confidence=conf.get(d), ref_mask=ref_mask,
                     score_label=_label.get(kind, "score"), cfg=cfg, score_col=scol)
    fig.suptitle("UMAP per driver — patient · mutation status · driver-program score "
                 "(per-cell where available; otherwise patient-level; reference/context greyed)", fontsize=13, y=1.0)
    fig.tight_layout()
    out = Path(outpath); fig.savefig(out, dpi=dpi, bbox_inches="tight"); plt.close(fig)
    return out


def context_strip(adata, outpath, patient_col="patient", celltype_col="CellType",
                  malignant_col="scope_malignant", dpi=300) -> Path | None:
    """A single context row: [patient | cell type | malignant compartment]."""
    emb = _embedding(adata)
    s = _point_size(adata.n_obs)
    panels = []
    if patient_col in adata.obs.columns:
        try:
            from .data import is_usable_patient_series
            if is_usable_patient_series(adata.obs[patient_col], colname=patient_col):
                panels.append(("patient", adata.obs[patient_col].values, "cat"))
            else:
                print(f"[umap_panels] skipping context patient strip: '{patient_col}' looks like cell IDs/barcodes.")
        except Exception:
            panels.append(("patient", adata.obs[patient_col].values, "cat"))
    if celltype_col in adata.obs.columns:
        panels.append(("cell type", adata.obs[celltype_col].values, "cat"))
    if malignant_col in adata.obs.columns:
        panels.append(("malignant compartment", adata.obs[malignant_col].astype(str).values, "cat"))
    if not panels:
        return None
    fig, axes = plt.subplots(1, len(panels), figsize=(5.2 * len(panels), 4.6), squeeze=False)
    for j, (title, lab, _) in enumerate(panels):
        _panel_categorical(axes[0][j], emb, lab, s, title)
    fig.tight_layout()
    out = Path(outpath); fig.savefig(out, dpi=dpi, bbox_inches="tight"); plt.close(fig)
    return out
```
