## Supplementary material for "scOPE identifies which driver-associated expression programs transfer from bulk tumors to single cells": scOPE_supplement

#### Abbreviations

AML, acute myeloid leukemia; ARI, adjusted Rand index; AUPRC, area under the precision–recall curve; AUROC, area under the receiver operating characteristic curve; BRCA, breast invasive carcinoma; CNV, copy-number variation; CRC, colorectal cancer; FDR, false-discovery rate; GBM, glioblastoma; LUAD, lung adenocarcinoma; PAAD, pancreatic adenocarcinoma; scRNA-seq, single-cell RNA sequencing; SHAP, SHapley Additive exPlanations; SKCM, skin cutaneous melanoma; SVD, singular value decomposition; TCGA, The Cancer Genome Atlas; UMAP, uniform manifold approximation and projection.

#### Organization of the Supplementary Information

The supplementary material extends the main-text analyses without reproducing complete main-text panels. Supplementary Figures S1–S3 provide bulk-model diagnostics that are not shown in main Figure 2; Supplementary Figure S4 reports confidence profiles for the six cancers not displayed in main Figure 3A; Supplementary Figure S5 shows the nine AML direct-label distributions not displayed in main Figure 3D–F; and Supplementary Figure S6 contains non-*NPM1* longitudinal trajectories together with the additional *NPM1* samples omitted from main Figure 5. Supplementary Figure S7 retains the complete AML and CRC genotype-localization atlases because the full driver-by-driver context is itself the purpose of the figure, even though selected rows appear in main Figure 4. Supplementary Figure S8 provides cellular-state context and score atlases for CRC, GBM, LUAD, PAAD, and SKCM, complementing the AML and BRCA examples in main Figure 6. Supplementary Figure S9 provides CNV panels omitted from main Figure 7, and Supplementary Figure S10 summarizes controls and guardrail audits generated by the canonical analysis.

Supplementary Tables S1–S10 are provided in two parallel forms: readable typeset versions are included in Section 3 below for direct consultation, and the complete machine-readable CSV files remain the authoritative data release. The much larger set of per-driver calibration, permutation, component, loading, SHAP, longitudinal density, violin, UMAP, and audit files is retained in `Supplemental_Code.zip` rather than reproduced indiscriminately in the PDF.

### 1 Additional Supplementary Methods

#### 1.1 Scope of the extended bulk-model audit

Main Figure 2 reports the complete pooled out-of-fold AUROC forest plot, representative pooled out-of-fold ROC curves, and the selected AML *NPM1* component example. Supplementary Figure S1 therefore focuses on information not visible in that figure: improvement in AUPRC above the prevalence baseline, the relationship between positive-class size and AUROC uncertainty, Brier-score distributions, claim-safety retention, and selected calibration and label-permutation diagnostics. The calibration and permutation panels were generated by a separate driver-specific interpretation evaluator operating on the full-cohort fitted latent representation; they are diagnostic rather than claim-bearing, and their displayed AUROCs can differ slightly from the leakage-free pooled out-of-fold estimates in main Figure 2 and Supplementary Table S1. The three model examples were selected before figure assembly to illustrate distinct audit conditions: AML *NPM1* as a well-powered high-performing model, GBM *IDH1* as a high-performing but rare-positive model, and AML *SMC1A* as a high-point-estimate model that remained claim-unsafe. The full AML cohort contained 10 *SMC1A*-positive tumors, but only four positives were represented among valid pooled out-of-fold predictions because one fold did not produce a trained driver classifier. All 158 claim-bearing model-level values are provided in Supplementary Table S1.

#### 1.2 Latent-dimension sensitivity and component interpretation

The canonical analysis used 30 truncated-SVD components. The archived sensitivity grid evaluated 5, 10, 25, 50, 75, 100, 150, and, where permitted by cohort dimensions, 250 and 500 components. Because 30 was not an explicit member of this secondary grid, Supplementary Figure S2 reports the cancer-level trajectories across the full grid and directly summarizes the 25-to-50-component change that brackets the canonical setting. These sensitivity models were refit and evaluated within the corresponding bulk cohort and were not used to select a different value after inspection.

Supplementary Figure S3 extends the component-ablation analysis beyond the AML *NPM1* example shown in main Figure 2D. It reports ablation curves for the leading BRCA, CRC, GBM, LUAD, PAAD, and SKCM models and the signed gene loadings associated with the highest-ranked AML *NPM1* components. These plots are interpretability diagnostics rather than independent evidence of driver specificity.

#### 1.3 Confidence and direct-label atlas selection

Main Figure 3 contains the complete AML confidence profile, confidence-versus-truth comparison, all-driver effect-size summary, direction diagnostic, and the selected *NPM1*, *TP53*, and *DNMT3A* violins. Supplementary Figure S4 therefore contains only the six additional cancer-level confidence profiles. Supplementary Figure S5 contains the remaining nine AML driver distributions and a separate coverage summary showing how strongly the number of truth-evaluable cells differed among drivers. The full numerical direct-label audit, including apparent AUROC, confidence intervals, Cliff's $\delta$, and FDR, is provided in Supplementary Table S3.

#### 1.4 Longitudinal atlas selection

The main figure displays the complete *NPM1* effect-size trajectory and five selected patient-level distributions. Supplementary Figure S6 reports the corresponding trajectory summaries for every other evaluated driver and adds *NPM1* distributions for AML1012, AML328, AML371, AML420B, AML707B, AML916, and AML921A. The complete per-patient, per-driver violin and density output set remains available in `Supplemental_Code.zip`; reproducing every generated panel in the PDF would add hundreds of nearly identical pages without adding a distinct inferential result. Numerical first-to-last changes and trend statistics are provided in Supplementary Table S7.

#### 1.5 Cellular-state atlas selection

Main Figure 6 provides detailed AML and BRCA examples. Supplementary Figure S8 therefore focuses on CRC, GBM, LUAD, PAAD, and SKCM. For each cohort, the expression-derived UMAP context page shows patient or sample identity, source cell-type annotation, and the malignant/reference compartment used by the analysis. A second page displays every retained driver score map ordered by the ground-truth-free confidence score. The associated driver-by-cell-type heatmap is shown beside the context panels. Score color limits are visualization scales; all tests used the untruncated values recorded in the analysis tables.

##### 1.6 CNV panels and near-diploid analyses

Main Figure 7 already shows continuous score–CNV concordance for all seven cohorts, BRCA, CRC, and PAAD confusion matrices, six cohort-level discordance counts, and AML and BRCA patient fractions. Supplementary Figure S9 therefore adds the omitted AML, GBM, LUAD, and SKCM confusion matrices and the omitted CRC, GBM, LUAD, PAAD, and SKCM patient-fraction plots. The complete continuous and thresholded results are provided in Supplementary Table S5; near-diploid patient-level tests are provided in Supplementary Table S6.

##### 1.7 Controls and guardrail audits

Supplementary Figure S10 summarizes controls that were present in the canonical output. Bulk label-permutation results were available for all 158 attempted driver–cancer models. Solid-tumor pan-cancer analyses included observation-level permutations of the reference/non-reference labels, allowing the observed directed Cliff's $\delta$ to be compared with an empirical null. The reference-cell residualization audit recorded whether each driver–state combination used a same-state reference median or the prespecified cohort-wide fallback when fewer than 25 same-state reference cells were available. A separate direction audit flagged drivers for which the healthy-reference compartment had unexpectedly high scores or for which available mutation-label and tumor/reference directions were inconsistent.

The code archive also contains a patient-block permutation implementation and broader decomposition, classifier, and alignment ablation utilities. These utilities were not represented as completed claim-bearing results in the canonical output and are therefore not presented as if they had been run. This distinction avoids converting available code paths into retrospective evidence.

##### 1.8 Supplemental Code and provenance

The complete manuscript analysis layer is supplied as `Supplemental_Code.zip`. It contains `scope_analysis_v10.3.2.2`, the canonical configuration `drivers/config_canonical_v3.json`, the forced-rescore command, model and score audits, generated tables, figure-generation scripts, and the machine-readable output manifest. Public software installation is pinned to `scope-bio==0.2.0`. Supplementary Table S8 maps each main-text panel to its source artifact, score column, subset, statistical unit, analysis version, configuration hash, and random seed.

##### 1.9 Source-data provenance and cohort attribution

Supplementary Table S2 is the source-data index for both the bulk and single-cell analyses. It reports source publications and DOIs, repository/project accessions, access conditions, analysis-input descriptions, modeled or retained cohort sizes, and the reference annotations used in the single-cell analyses. The table distinguishes public processed deposits from raw human data that require controlled access or were not released directly by the source study. For the previously derived BeatAML and TCGA bulk matrices, project-level identifiers are reported because the original portal-specific download manifests and file UUIDs were not preserved in the frozen analysis tree; the exact derived matrix filenames remain recorded in the canonical configuration.

#### 2 Supplementary Figures

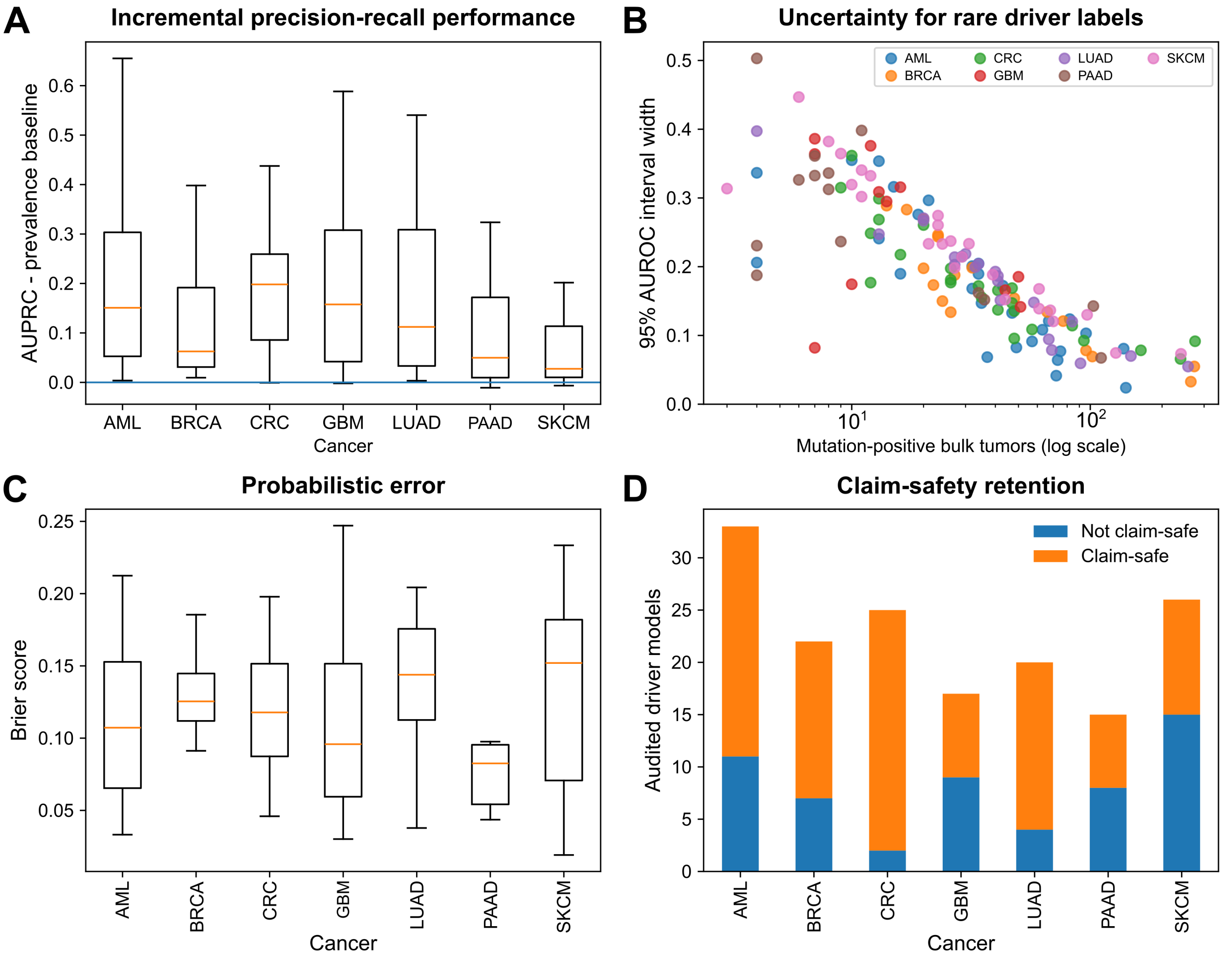


Supplementary Figure S1A–D. Bulk-model performance beyond the main-text AUROC display. (A) AUPRC improvement above the driver-prevalence baseline for all audited models, grouped by cancer. (B) Width of the 95% AUROC bootstrap interval versus the number of mutation-positive bulk tumors. (C) Brier-score distributions across cancers. (D) Counts of claim-safe and non-claim-safe evaluations; non-claim-safe and failed evaluations were retained rather than removed after inspection.

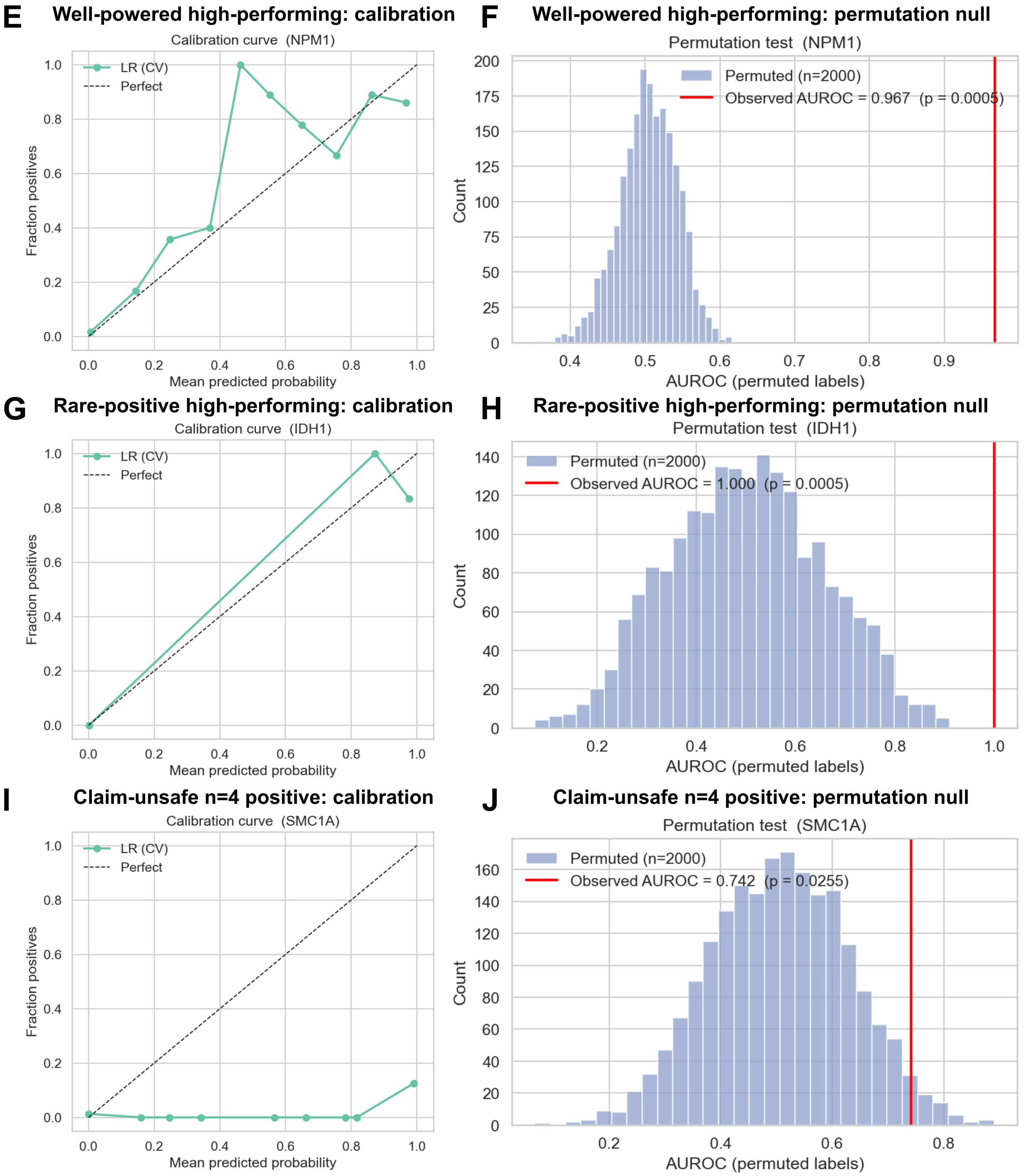


Supplementary Figure S1E–J. Representative calibration and bulk-label permutation diagnostics from the separate driver-specific interpretation evaluator. Rows show AML *NPM1*, GBM *IDH1*, and AML *SMC1A*, respectively; the left panel of each row is the calibration curve and the right panel is the observed AUROC relative to its label-permutation null. These diagnostic AUROCs can differ from the leakage-free pooled out-of-fold values used for claim-bearing inference. The examples represent a well-powered high-performing model, a rare-positive high-performing model, and a claim-unsafe high-point-estimate model. For *SMC1A*, the full fitted cohort contained 10 positives, whereas only four positives were represented among valid pooled out-of-fold predictions. Complete claim-bearing numerical results are in Supplementary Table S1.

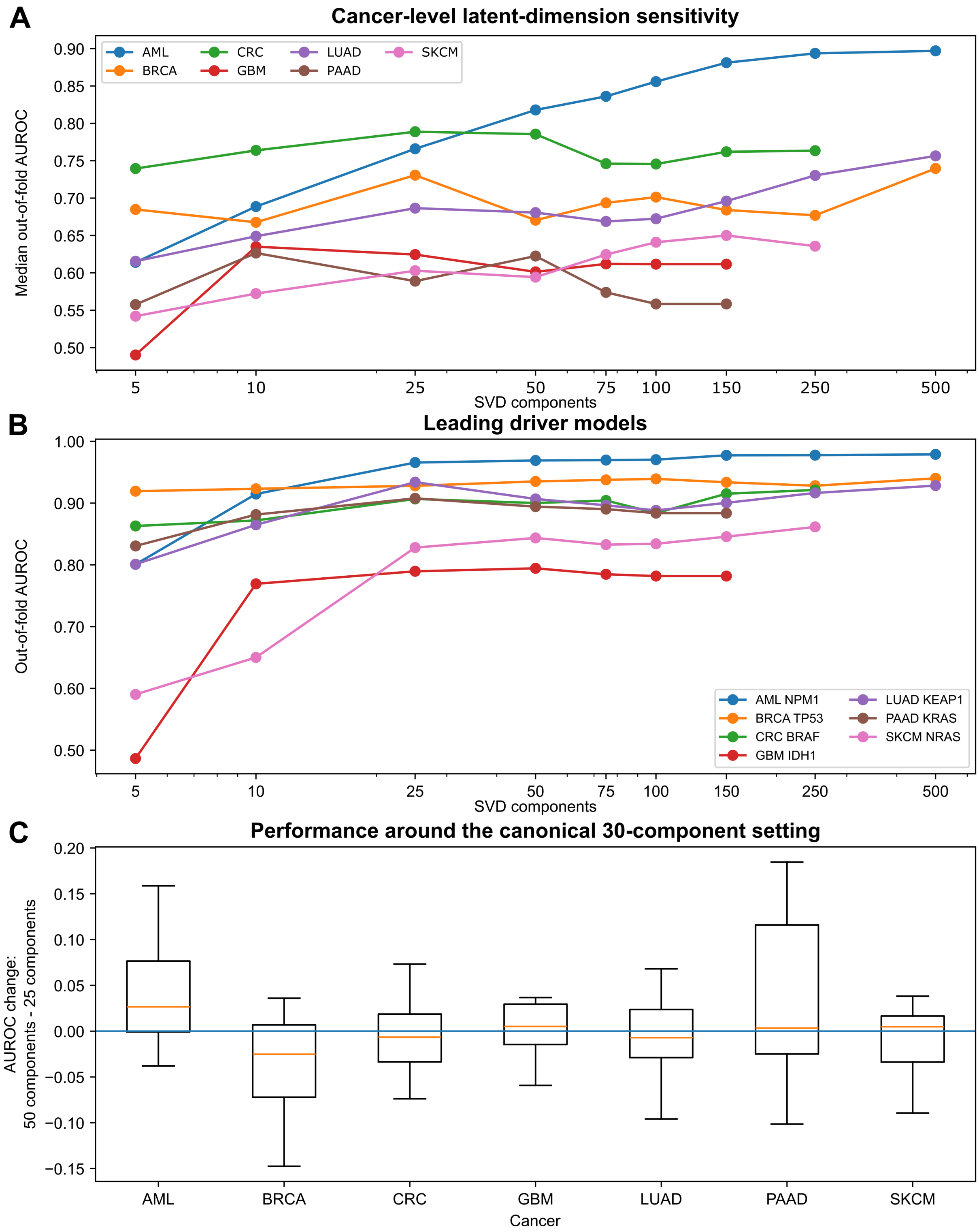


Supplementary Figure S2. Latent-dimension sensitivity. (A) Median out-of-fold AUROC across drivers as a function of the number of SVD components for each cancer. (B) Sensitivity curves for the leading model from each cancer. (C) Per-driver AUROC change between 25 and 50 components, the two sensitivity-grid values bracketing the canonical 30-component model. The sensitivity grid was diagnostic and was not used for post hoc model selection.

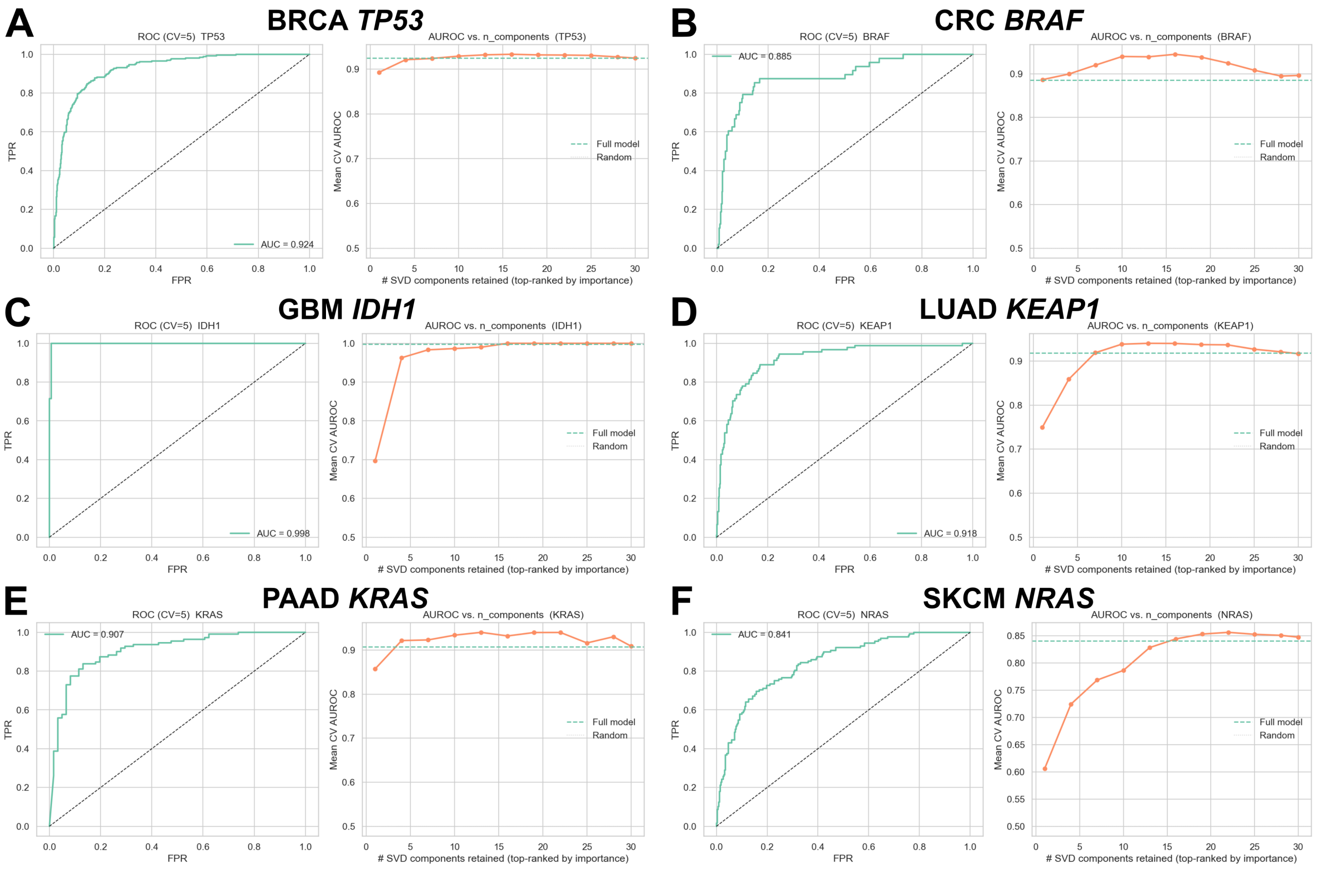


Supplementary Figure S3A. Component-ablation diagnostics for leading non-AML bulk models. Curves show cross-validated performance after retaining increasing numbers of classifier-ranked latent components for BRCA *TP53*, CRC *BRAF*, GBM *IDH1*, LUAD *KEAP1*, PAAD *KRAS*, and SKCM *NRAS*. The corresponding AML *NPM1* ablation is shown in main Figure 2D.

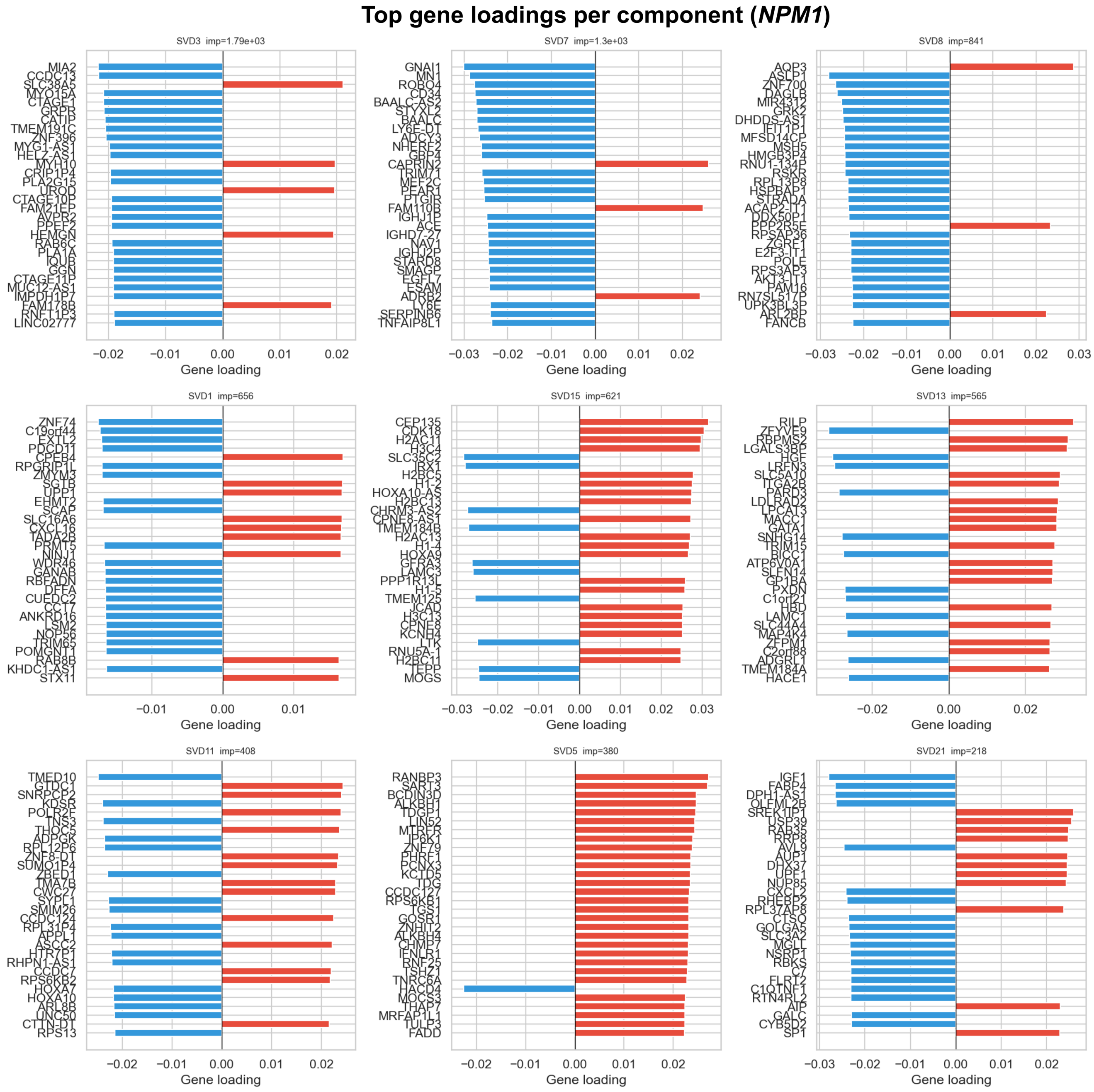


Supplementary Figure S3B. Signed gene loadings for the highest-ranked AML *NPM1* latent components. Positive and negative bars show the genes with the largest signed loadings within each component; component labels report their model-importance ranking. These loadings describe the transferred multicomponent expression program and are not interpreted as a causal mutation signature.

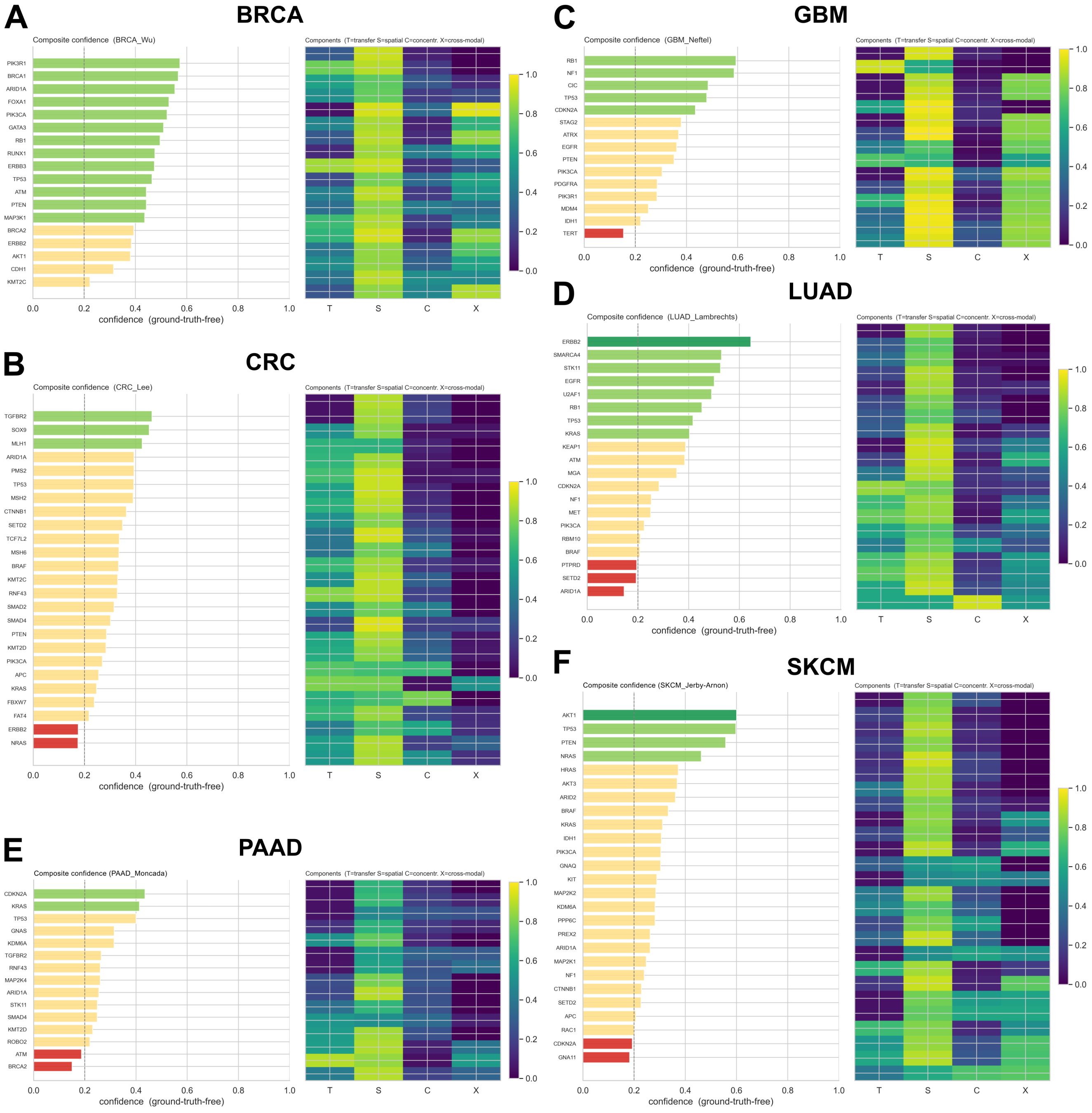


Supplementary Figure S4. Ground-truth-free confidence profiles for the six additional cancers. Composite confidence and its transferability (T), spatial-coherence (S), concentration (C), and CNV-axis agreement (X) components are shown for BRCA, CRC, GBM, LUAD, PAAD, and SKCM. The AML profile and its direct-label validation are shown in main Figure 3.

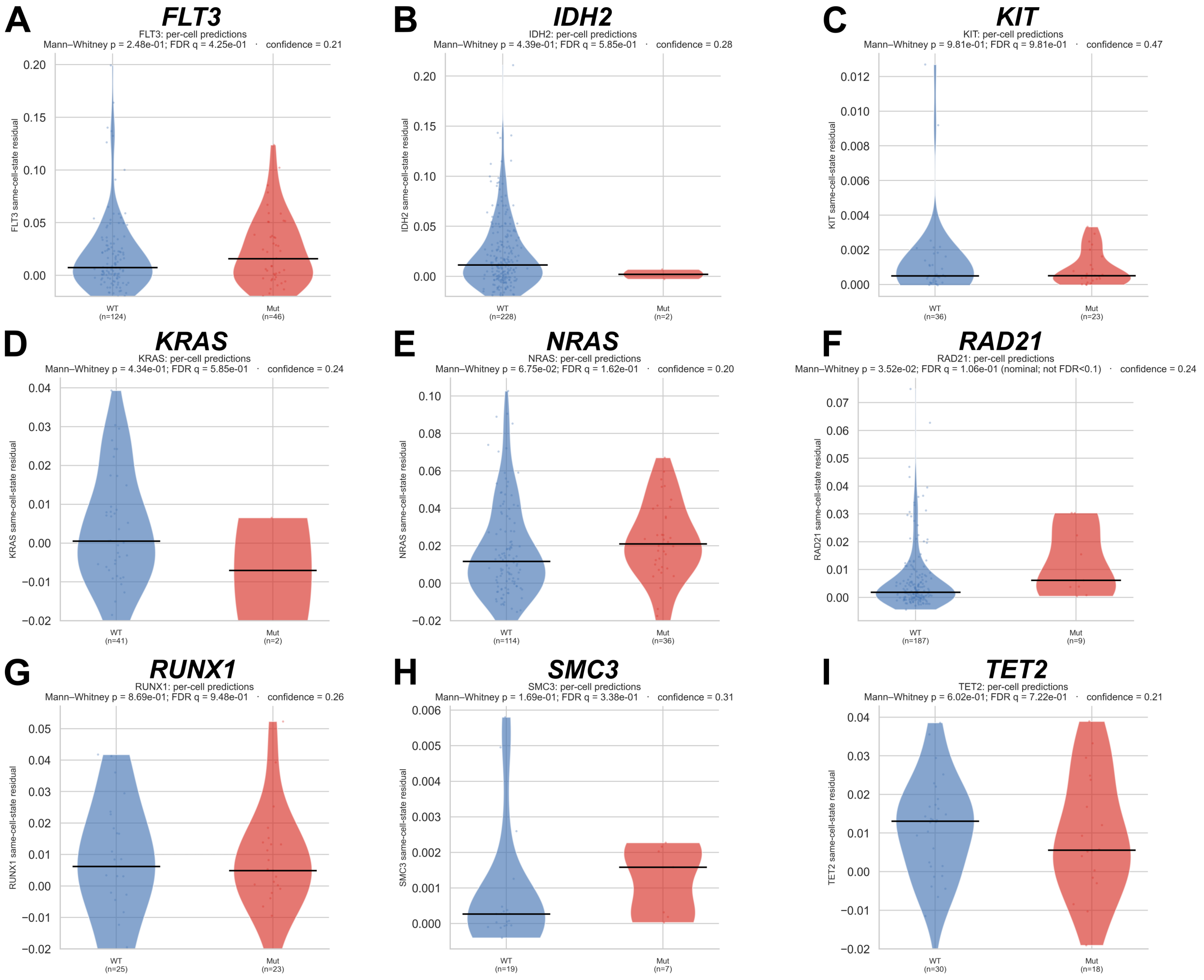


Supplementary Figure S5A. Direct AML mutation-label distributions for the nine drivers not displayed as violins in main Figure 3. Cell-state-residual scores are compared between reported-mutant cells and explicit wild-type-transcript comparison cells, with mutually exclusive transcript evidence required in both groups. Sample sizes, nominal Mann–Whitney values, FDR values, and confidence scores are shown in the source panels. These are positive-unlabeled cellular enrichment analyses rather than complete genomic genotype calls.

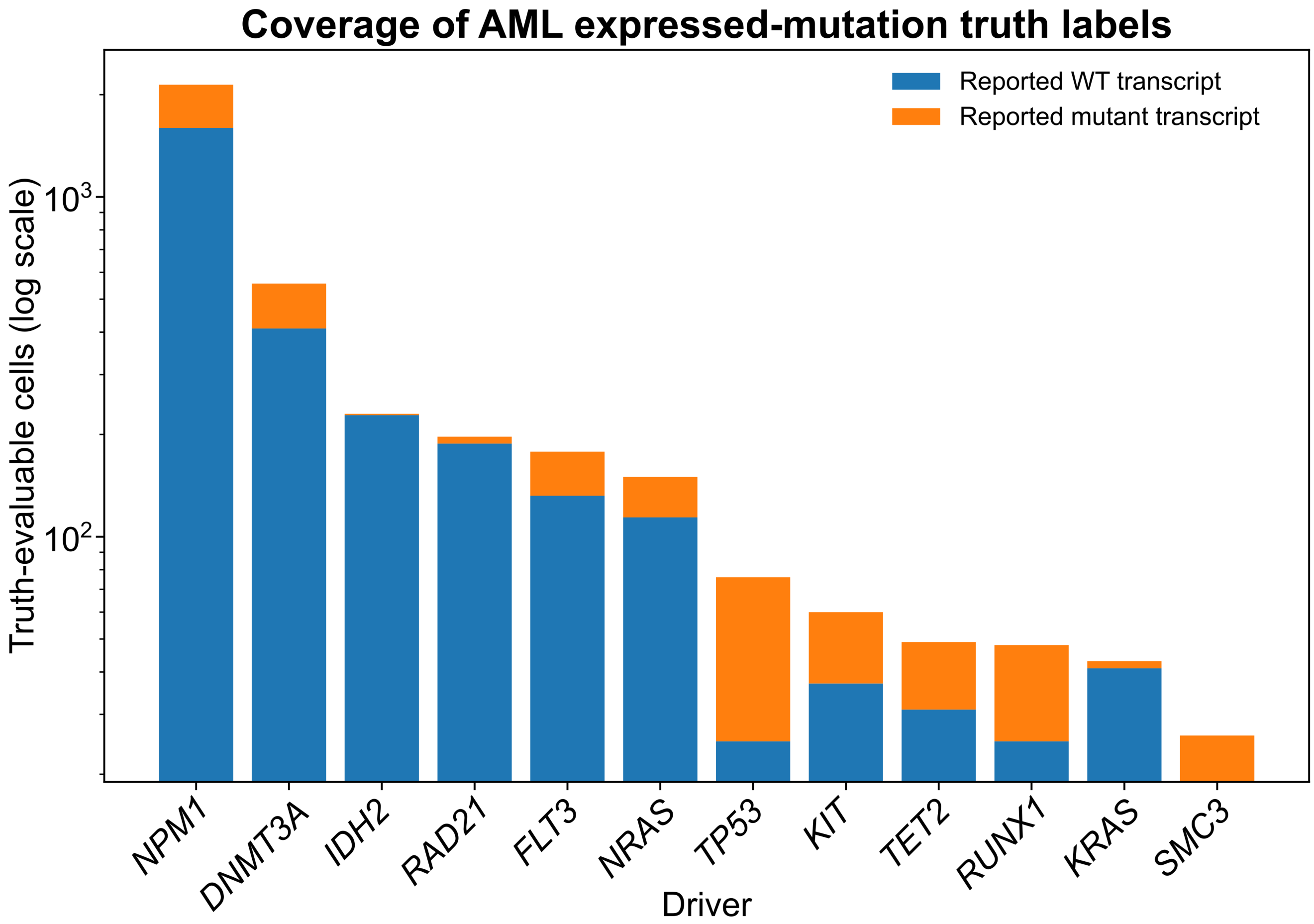


Supplementary Figure S5B. Coverage of AML expressed-mutation truth labels. Stacked counts show cells carrying mutually exclusive reported wild-type or mutant transcript evidence for each of the 12 drivers included in the direct-label analysis. The logarithmic axis emphasizes the strong driver-to-driver difference in truth-label availability.

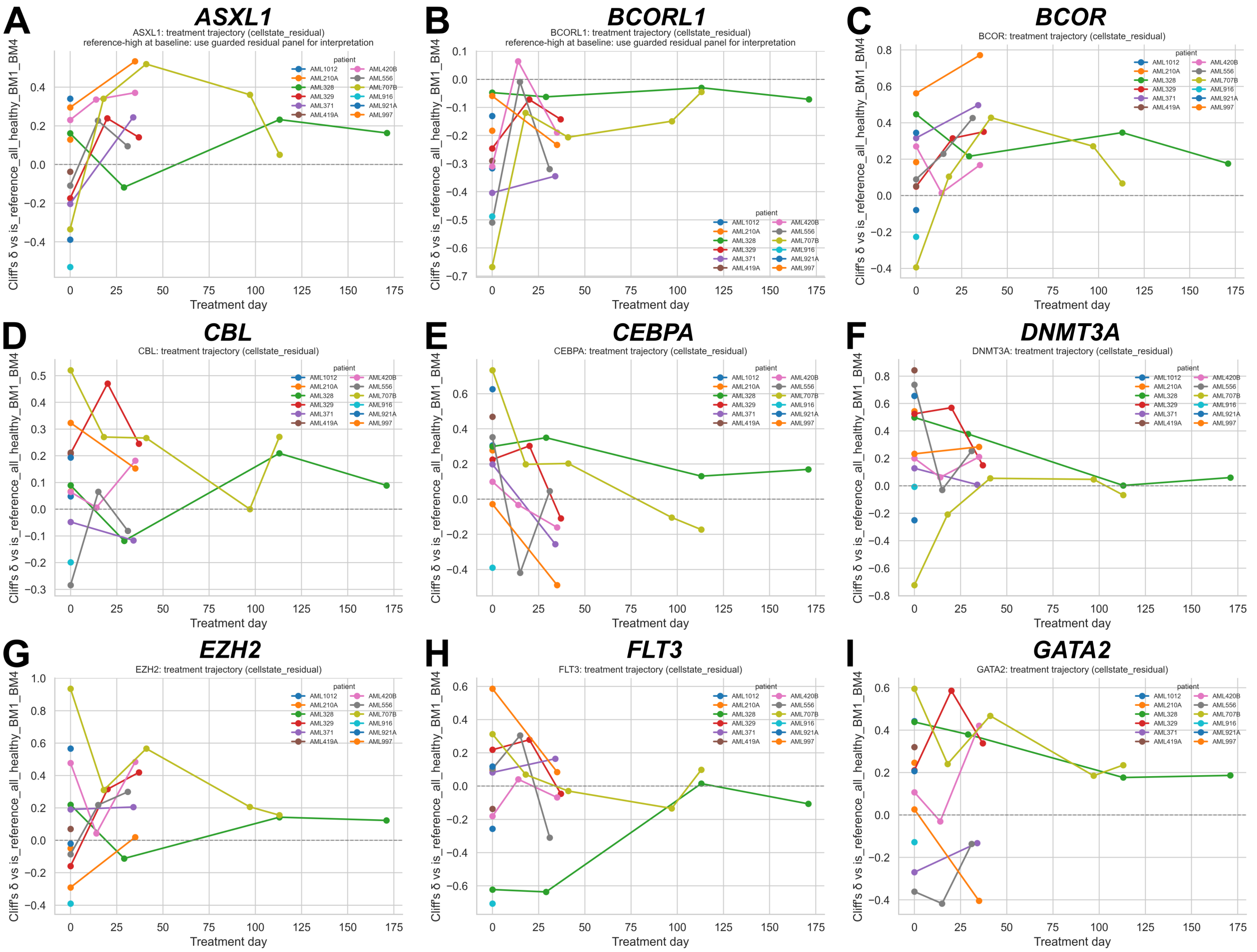


Supplementary Figure S6A, page 1. Longitudinal effect-size trajectories for non-*NPM1* AML driver programs. Each panel shows Cliff's $\delta$ relative to the fixed healthy bone-marrow reference across treatment day for every retained patient. Drivers without direct-label support are interpreted as exploratory state dynamics.

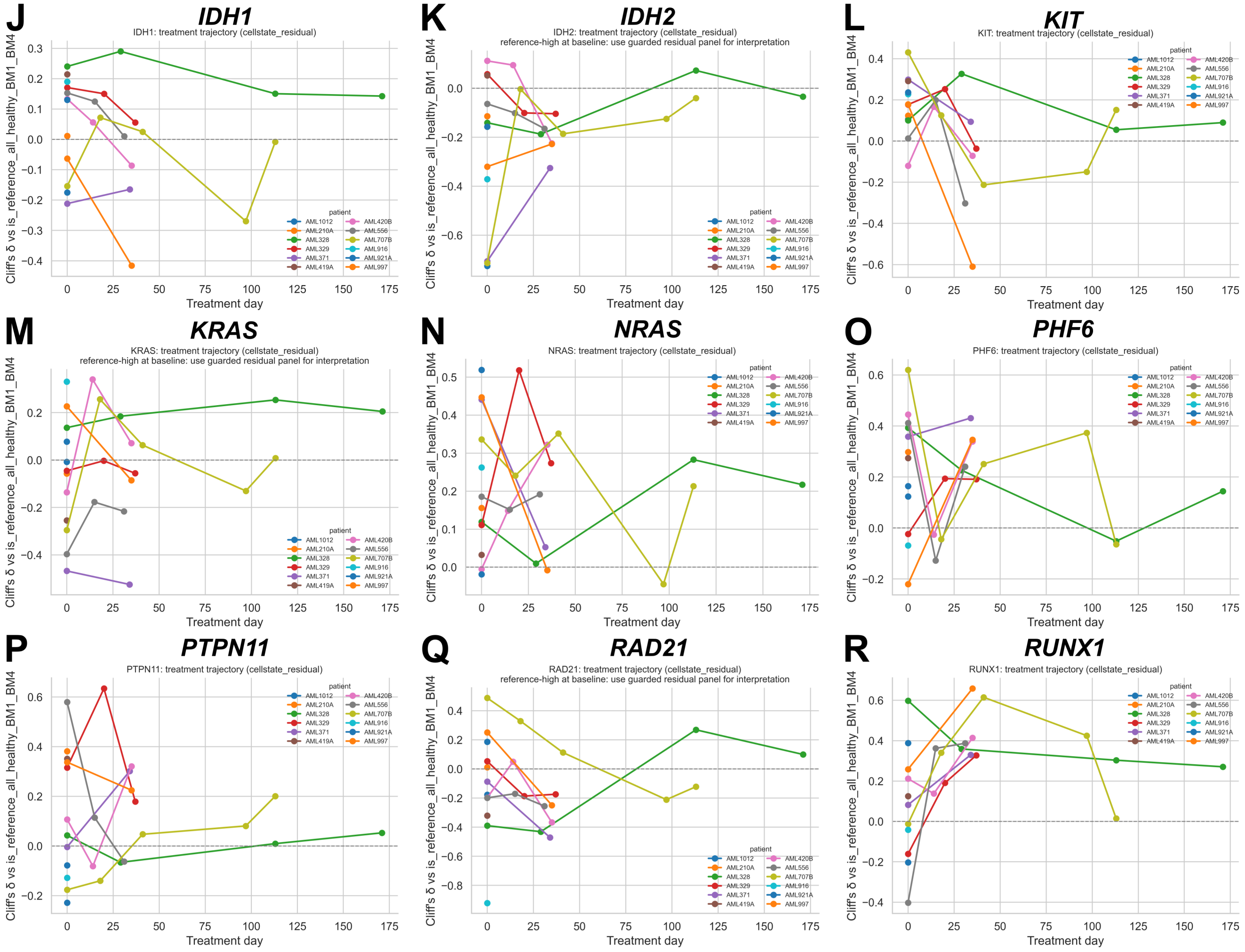


Supplementary Figure S6A, page 2. Continuation of the non-*NPM1* longitudinal trajectory atlas.

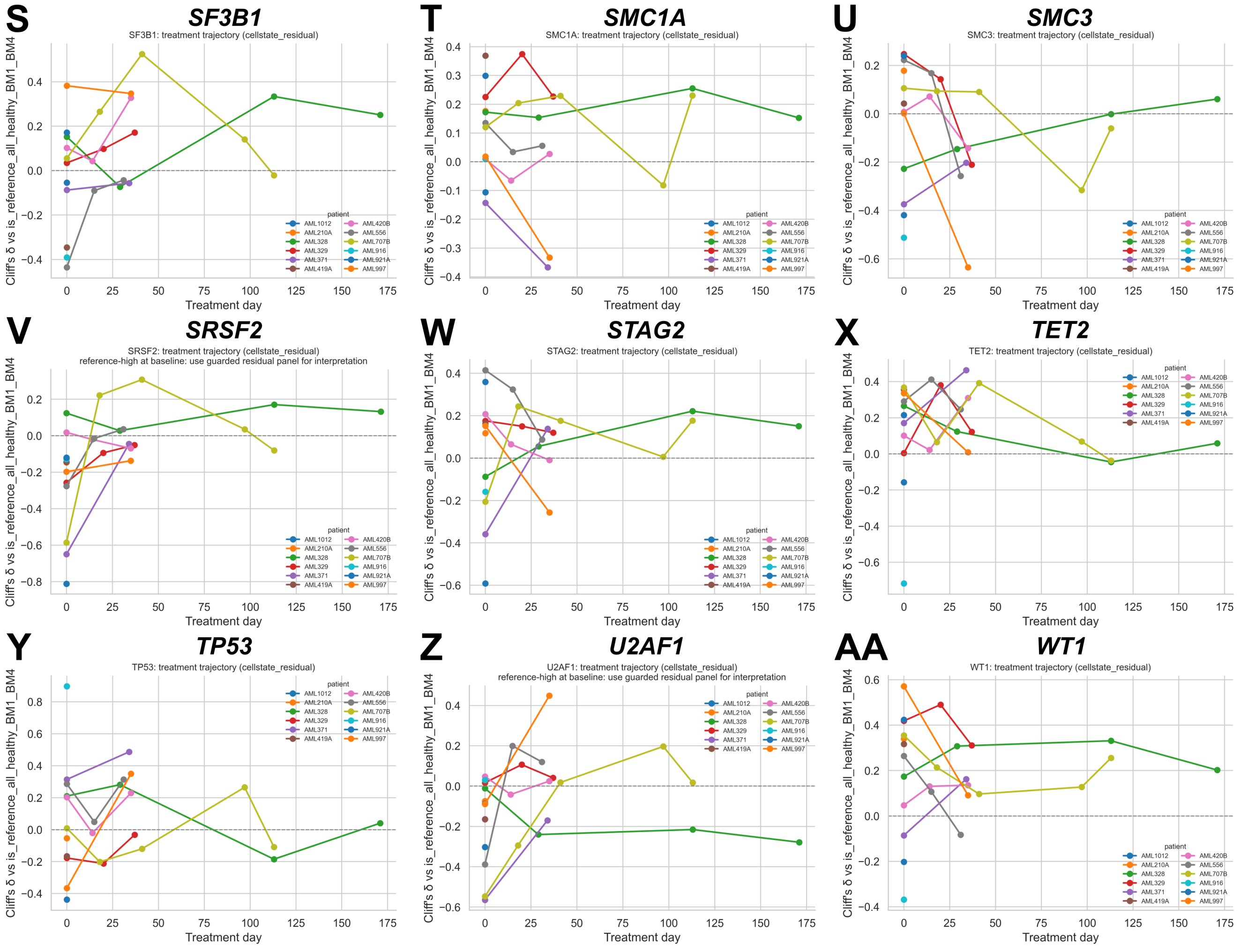


Supplementary Figure S6A, page 3. Continuation of the non-*NPM1* longitudinal trajectory atlas.

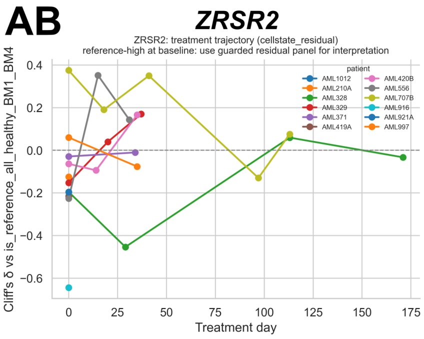


Supplementary Figure S6A, page 4. Continuation of the non-*NPM1* longitudinal trajectory atlas.

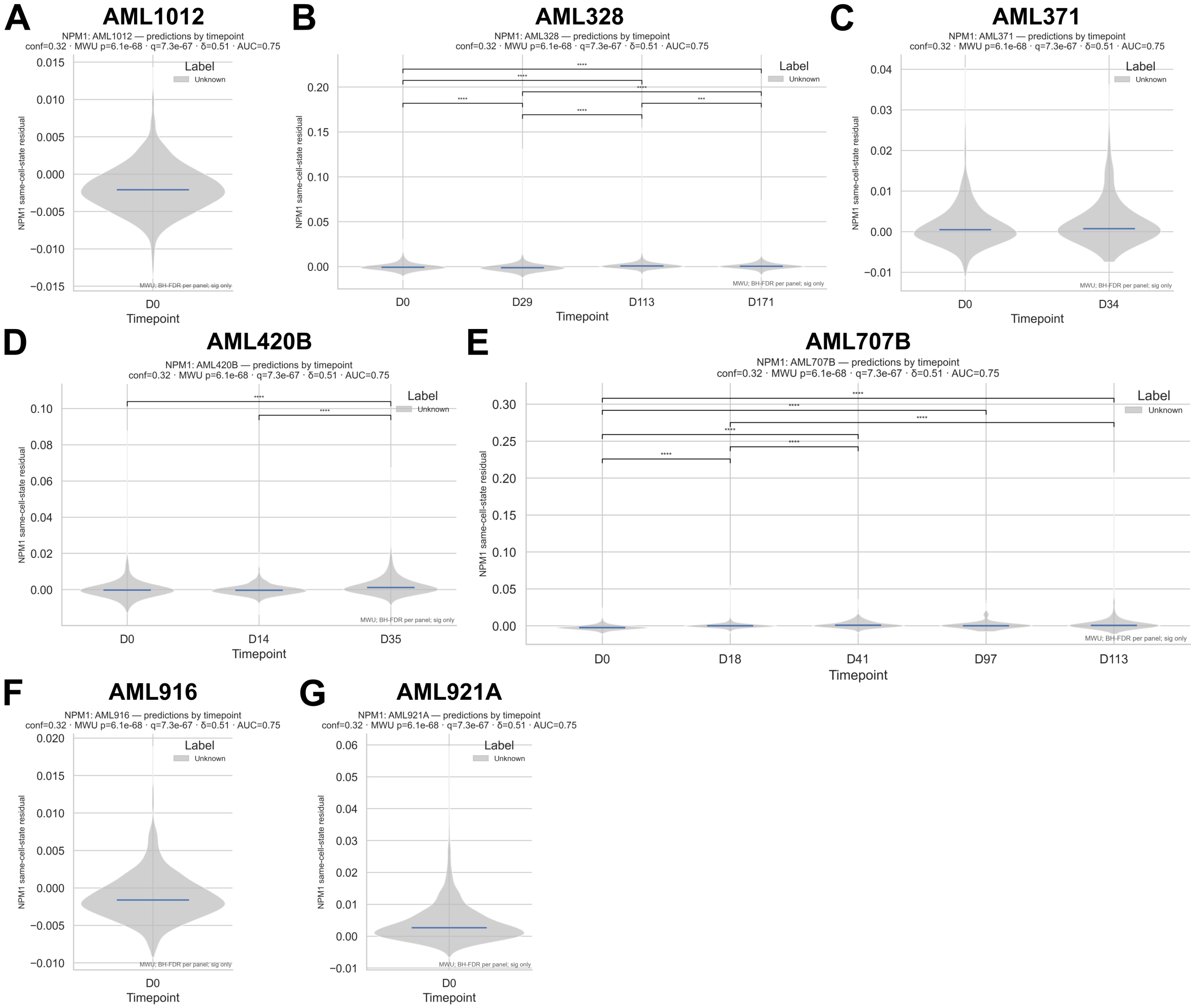


Supplementary Figure S6B. Additional AML *NPM1* patient-level score distributions not displayed in main Figure 5. Panels show AML1012, AML328, AML371, AML420B, AML707B, AML916, and AML921A. Pairwise annotations are descriptive within-patient comparisons and do not establish clonal identity.

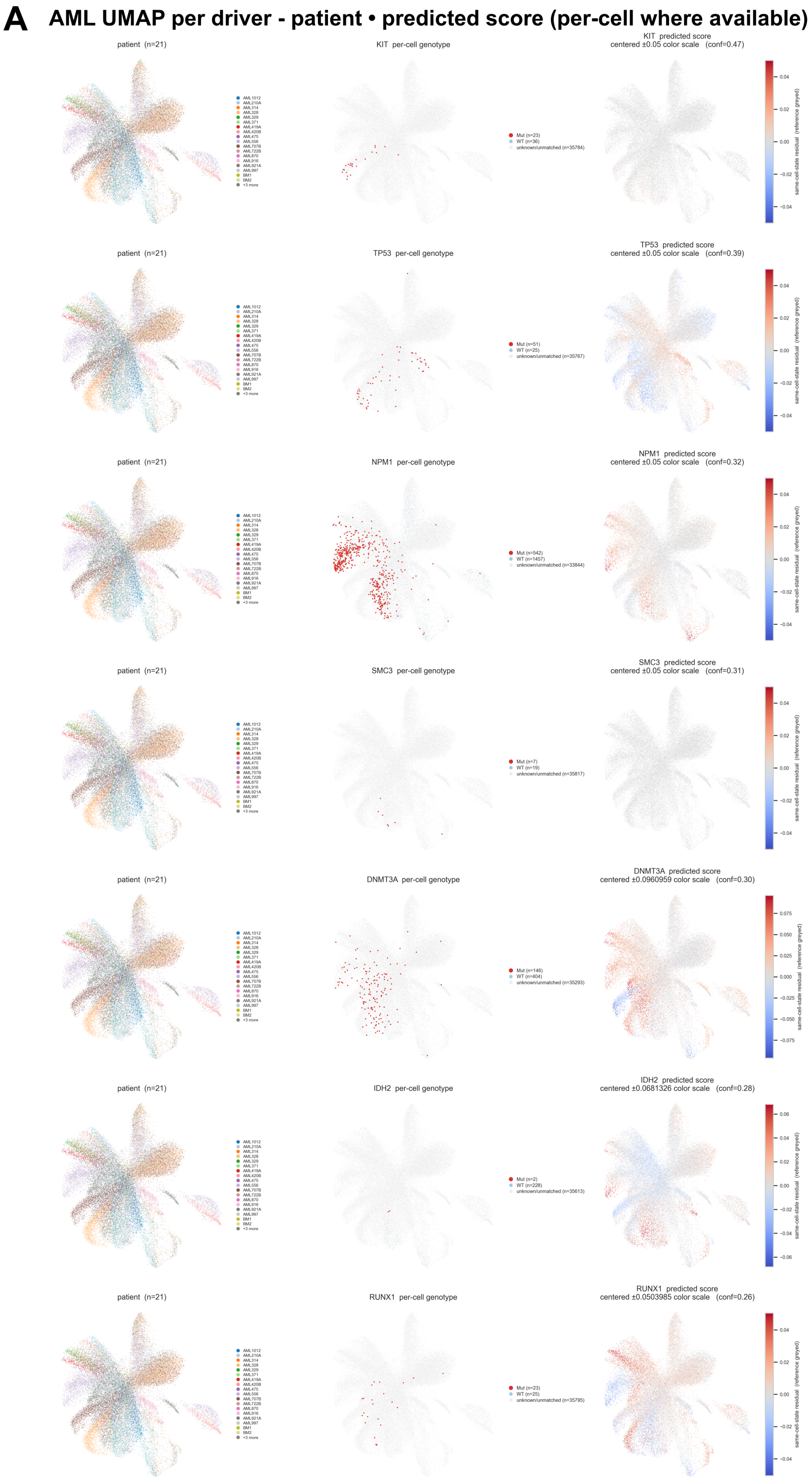


Supplementary Figure S7A. Complete AML genotype-localization atlas. Patient identity, guarded direct per-cell expressed-mutation annotation, and the matched cell-state-residual score are displayed on common expression-derived UMAP coordinates for every evaluable driver. Mutant and comparison labels require mutually exclusive transcript evidence; unlabeled cells are not assumed to be wild type.

**B**

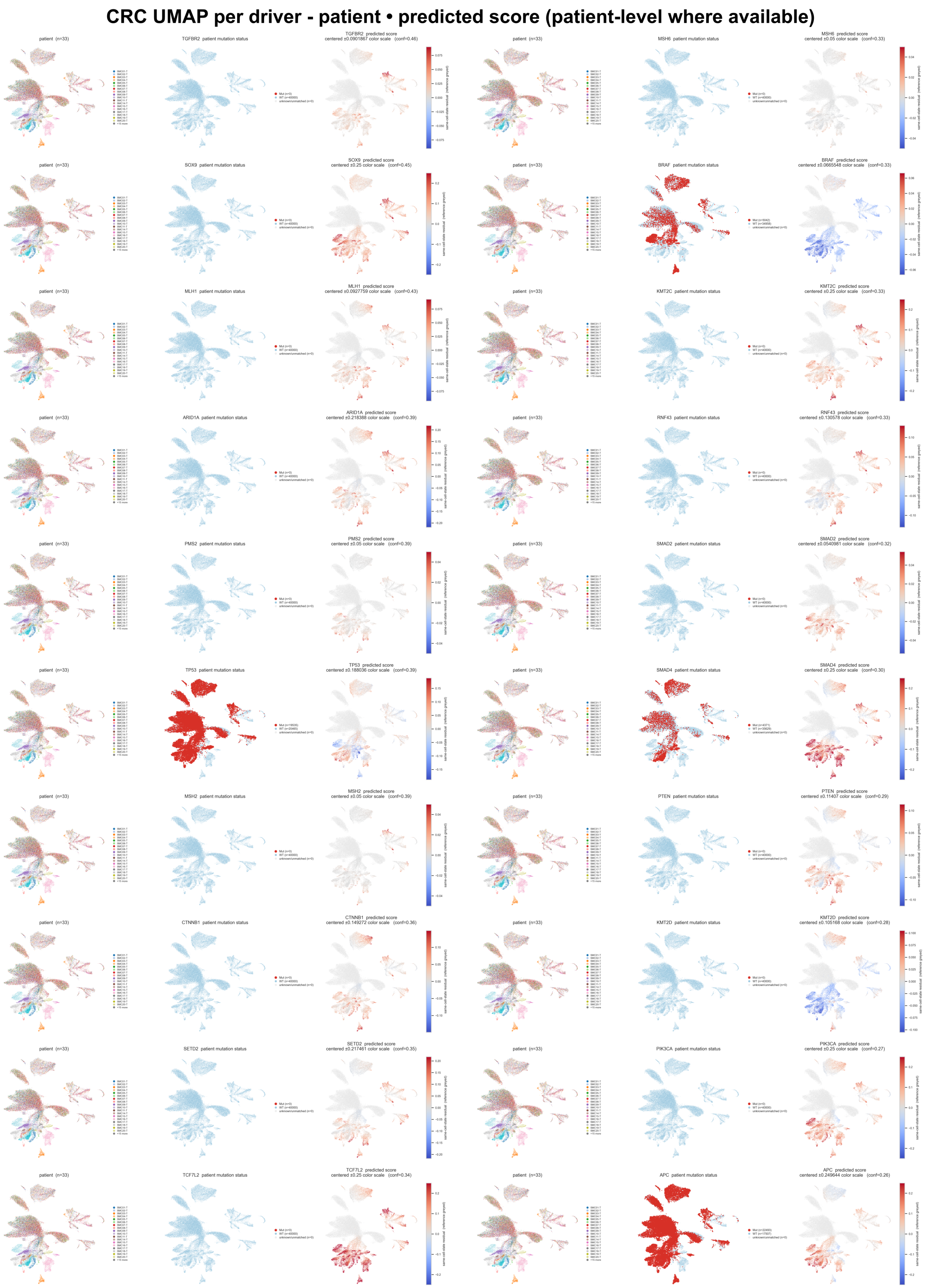


Supplementary Figure S7B. Complete CRC genotype-localization atlas. Patient identity, patient-level genotype context, and the matched cell-state-residual score are displayed on common expression-derived UMAP coordinates. Genotype colors are assigned from tumor-level metadata and are not interpreted as per-cell truth.

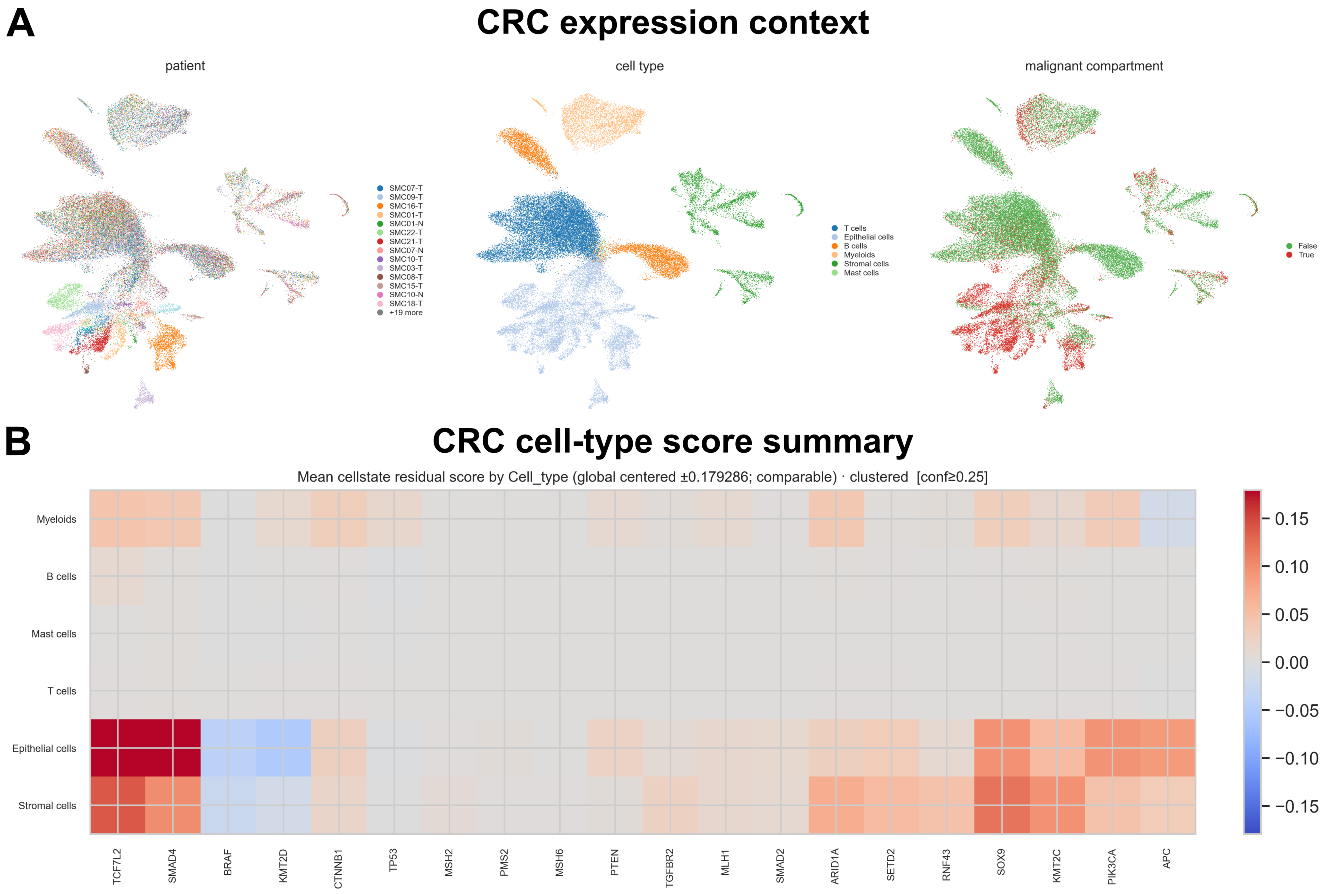


Supplementary Figure S8A. CRC cellular-state context. Expression-derived UMAPs show patient identity, source cell type, and malignant/reference compartment; the heatmap summarizes mean cell-state-residual score by source cell type and driver.

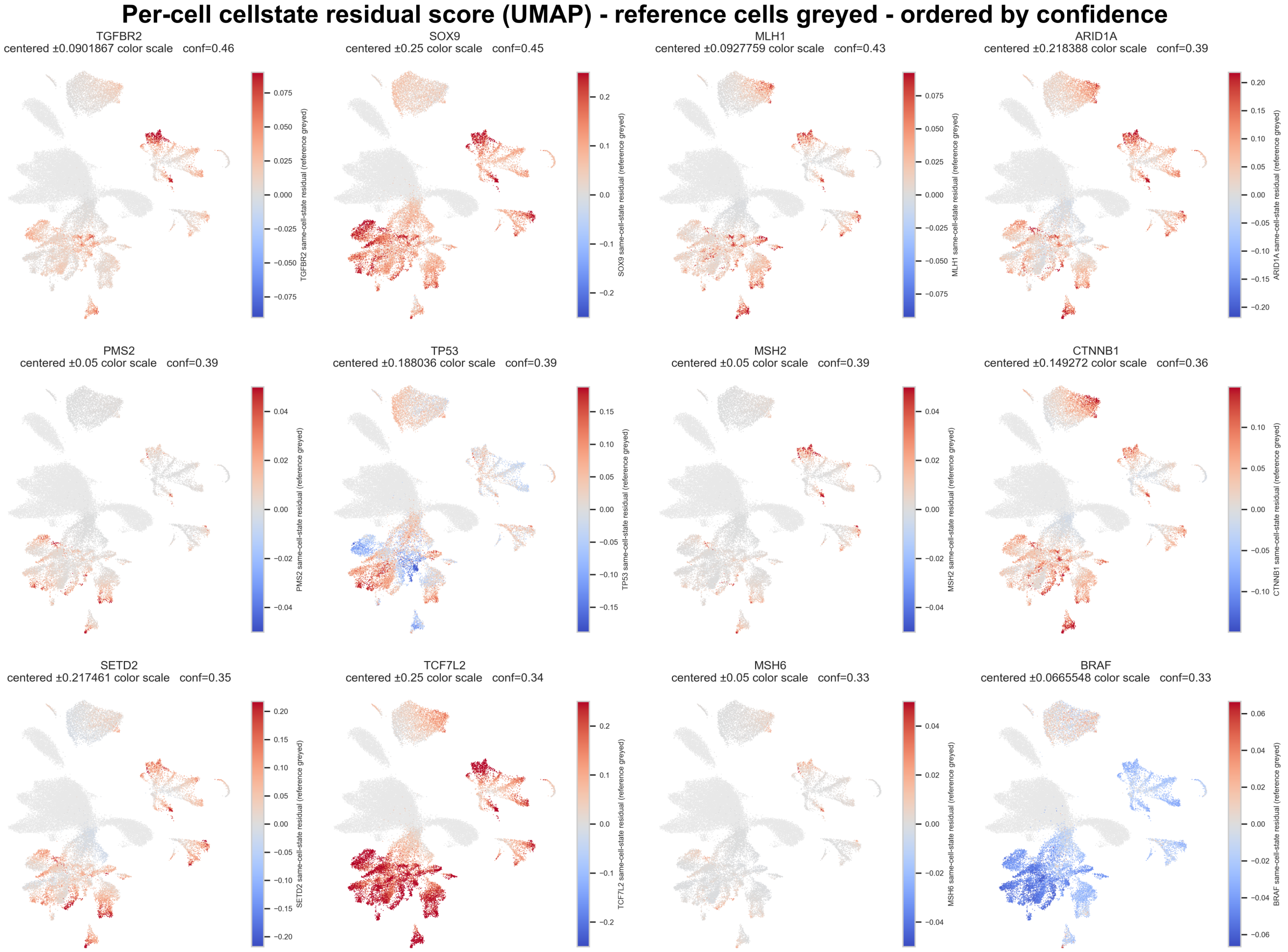


Supplementary Figure S8B. Complete CRC driver-score atlas. Cell-state-residual scores are displayed on the shared expression-derived UMAP and ordered by confidence. Reference cells are greyed.

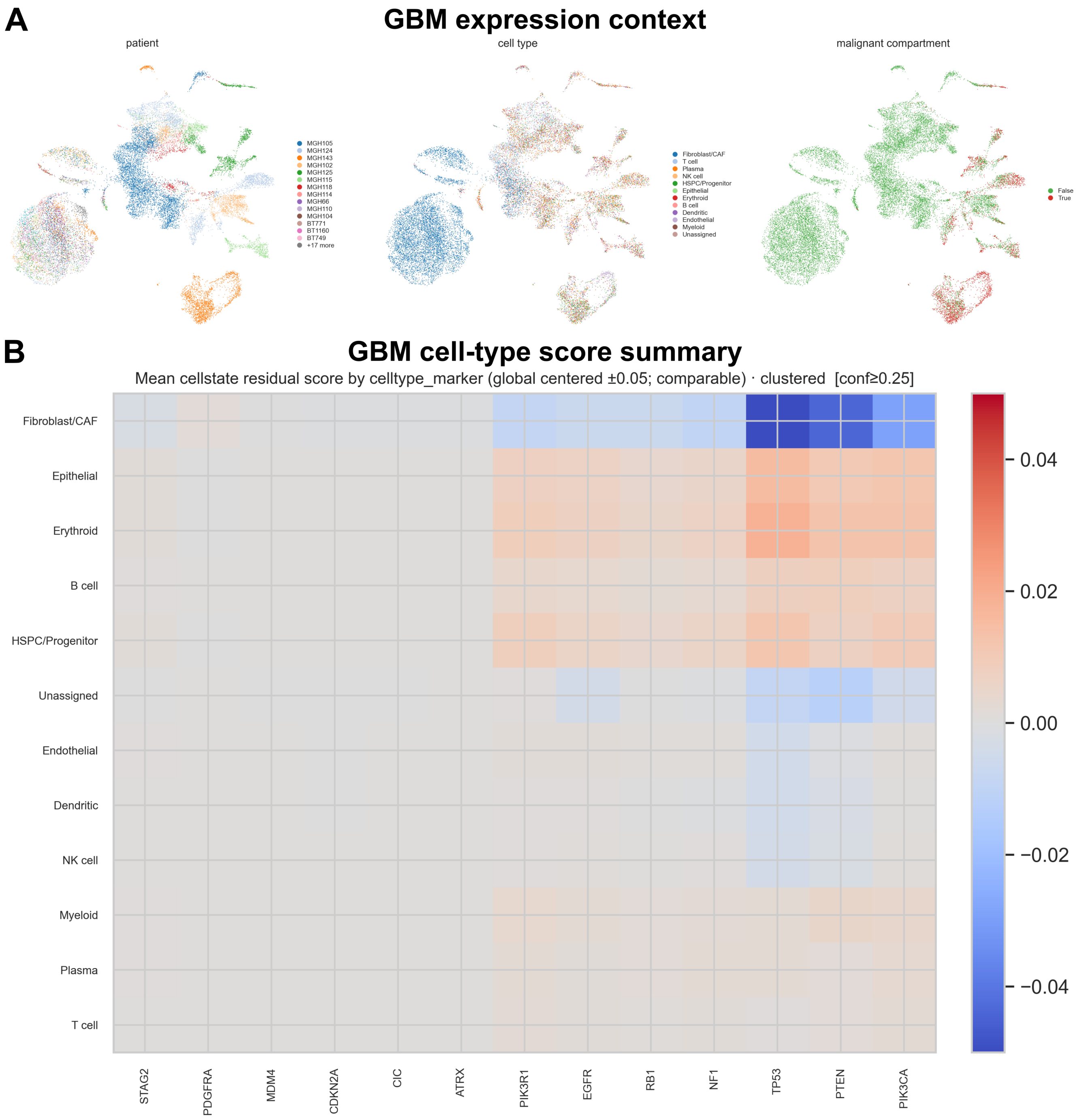


Supplementary Figure S8C. GBM cellular-state context and driver-by-cell-type score summary.

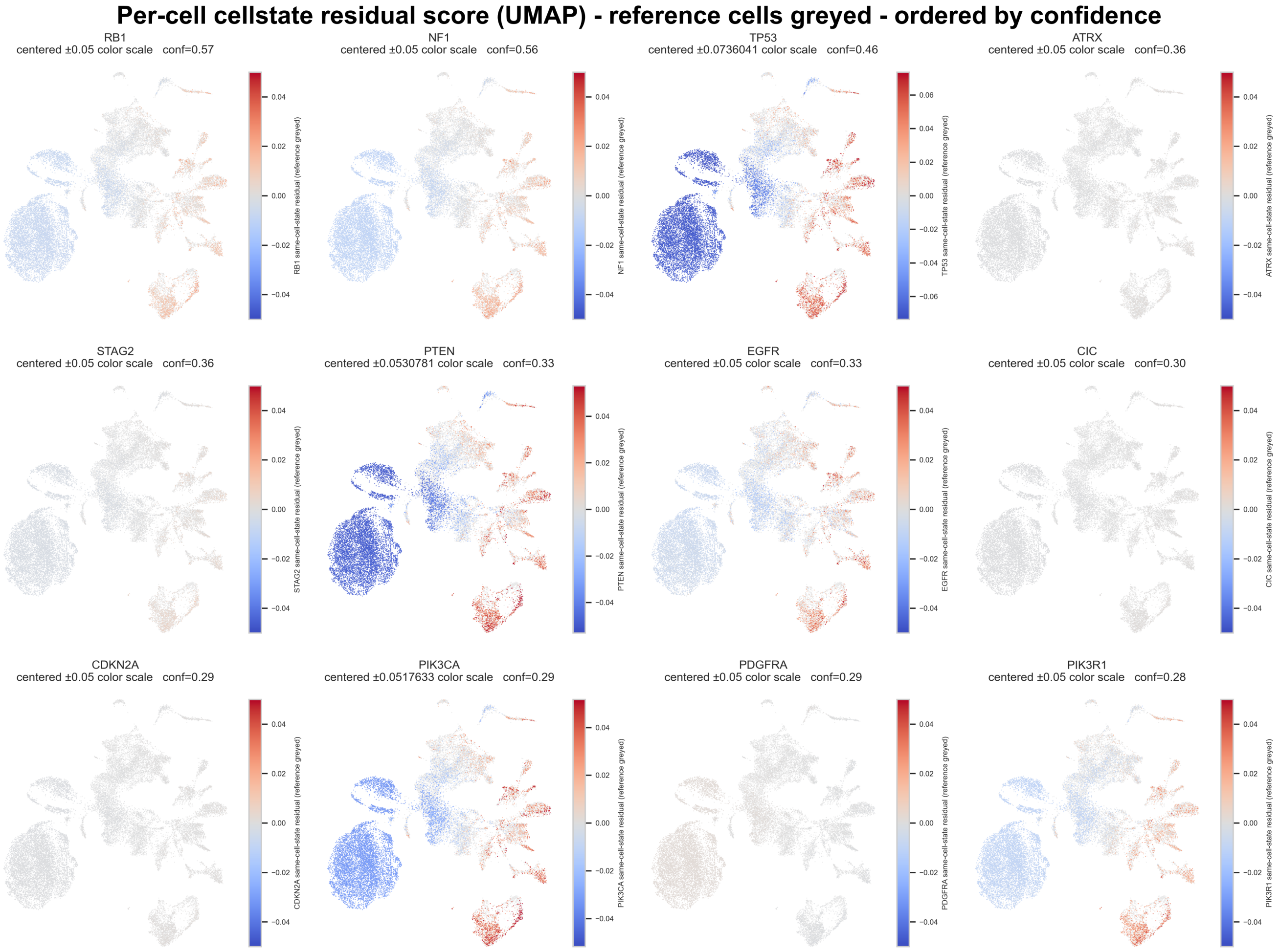


Supplementary Figure S8D. Complete GBM driver-score atlas.

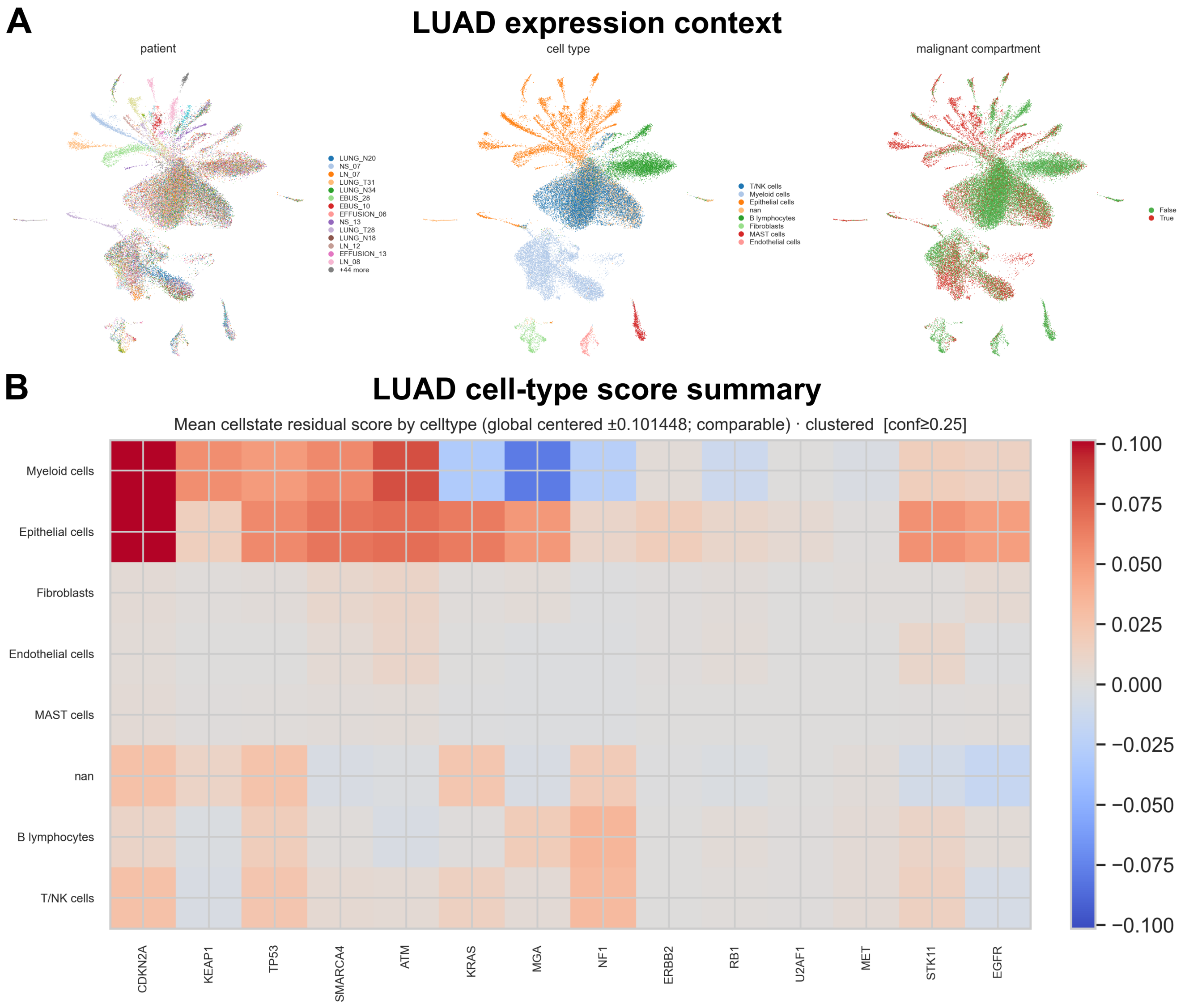


Supplementary Figure S8E. LUAD cellular-state context and driver-by-cell-type score summary.

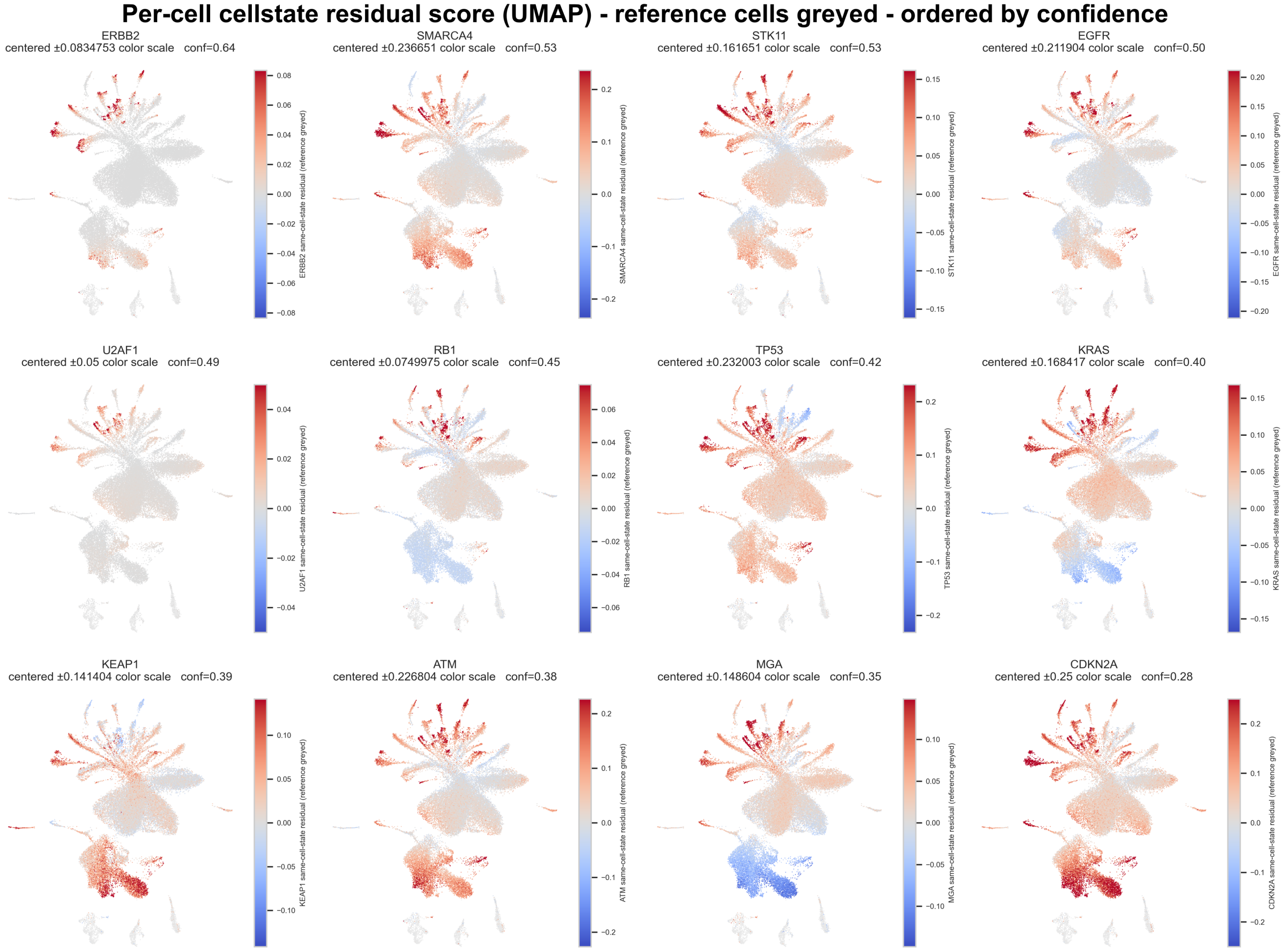


Supplementary Figure S8F. Complete LUAD driver-score atlas.

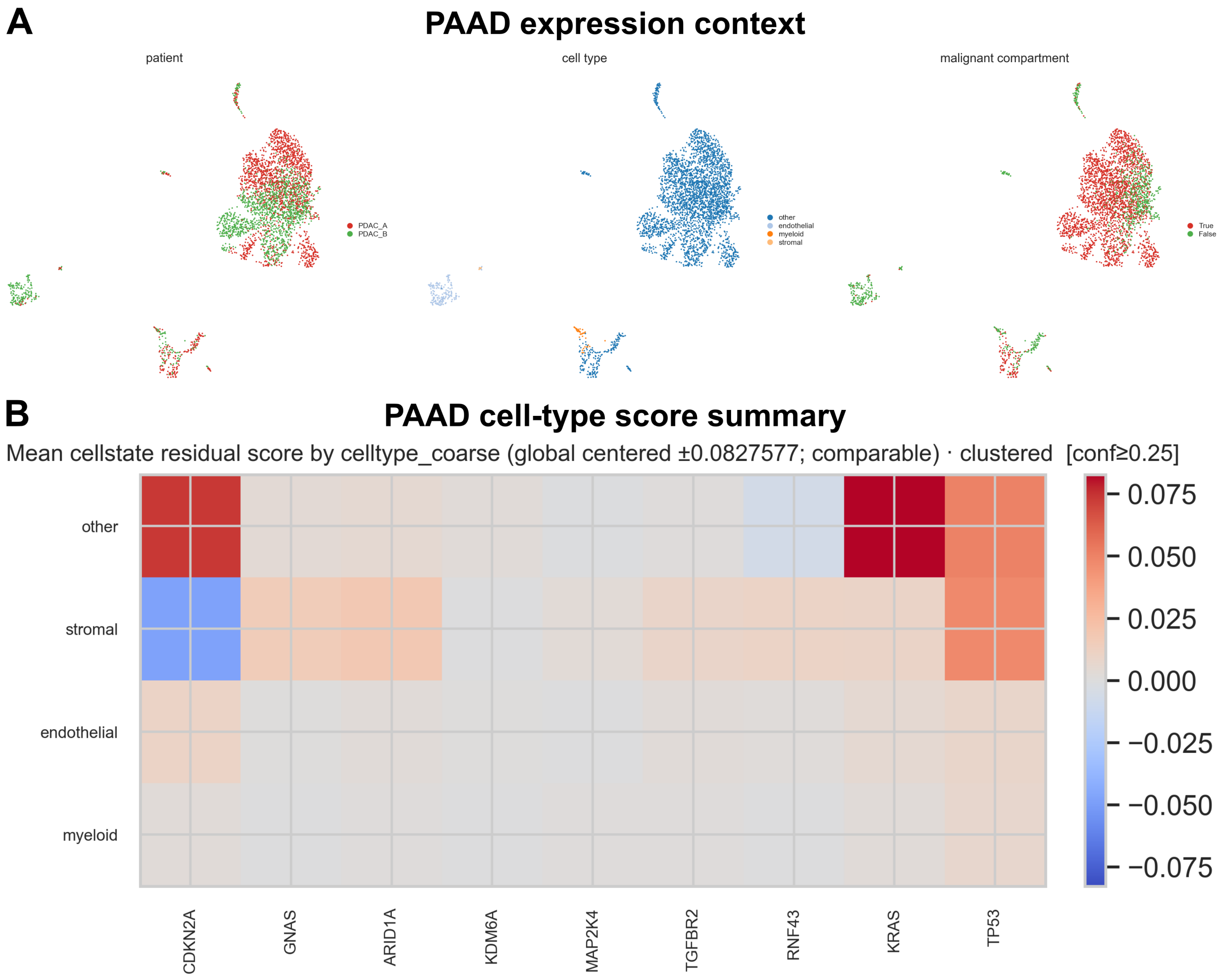


Supplementary Figure S8G. PAAD cellular-state context and driver-by-cell-type score summary.

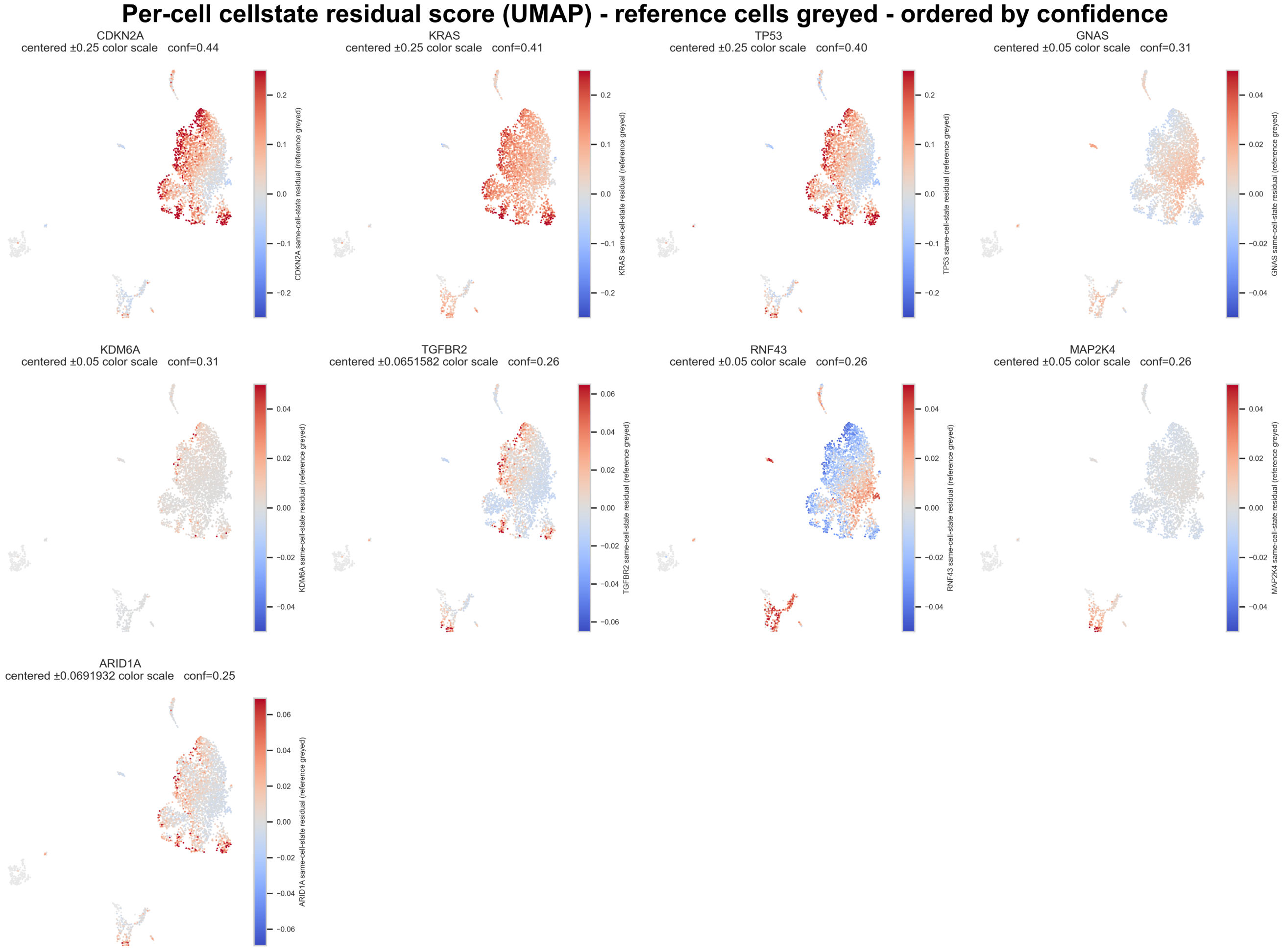


Supplementary Figure S8H. Complete PAAD driver-score atlas.

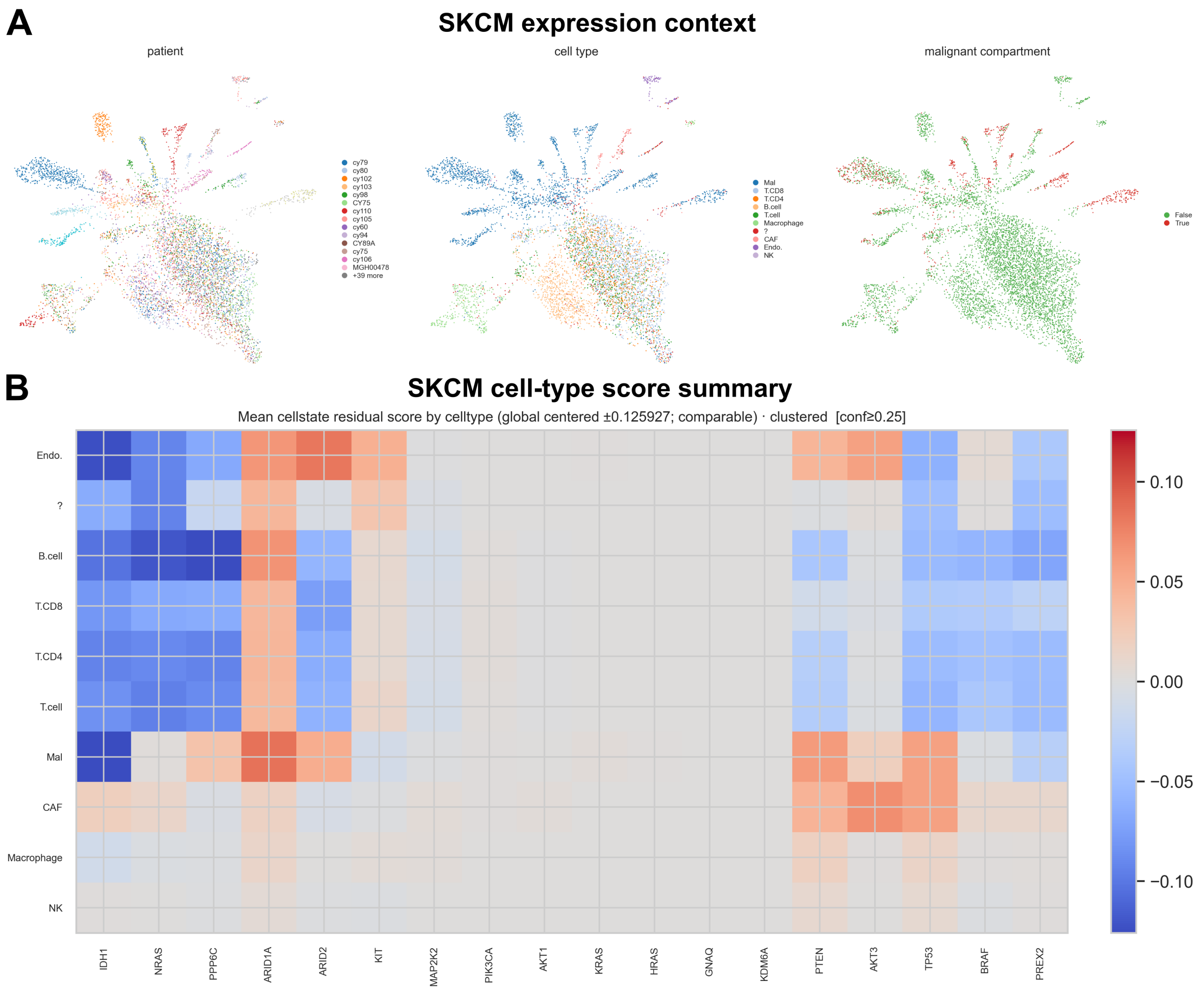


Supplementary Figure S8I. SKCM cellular-state context and driver-by-cell-type score summary.

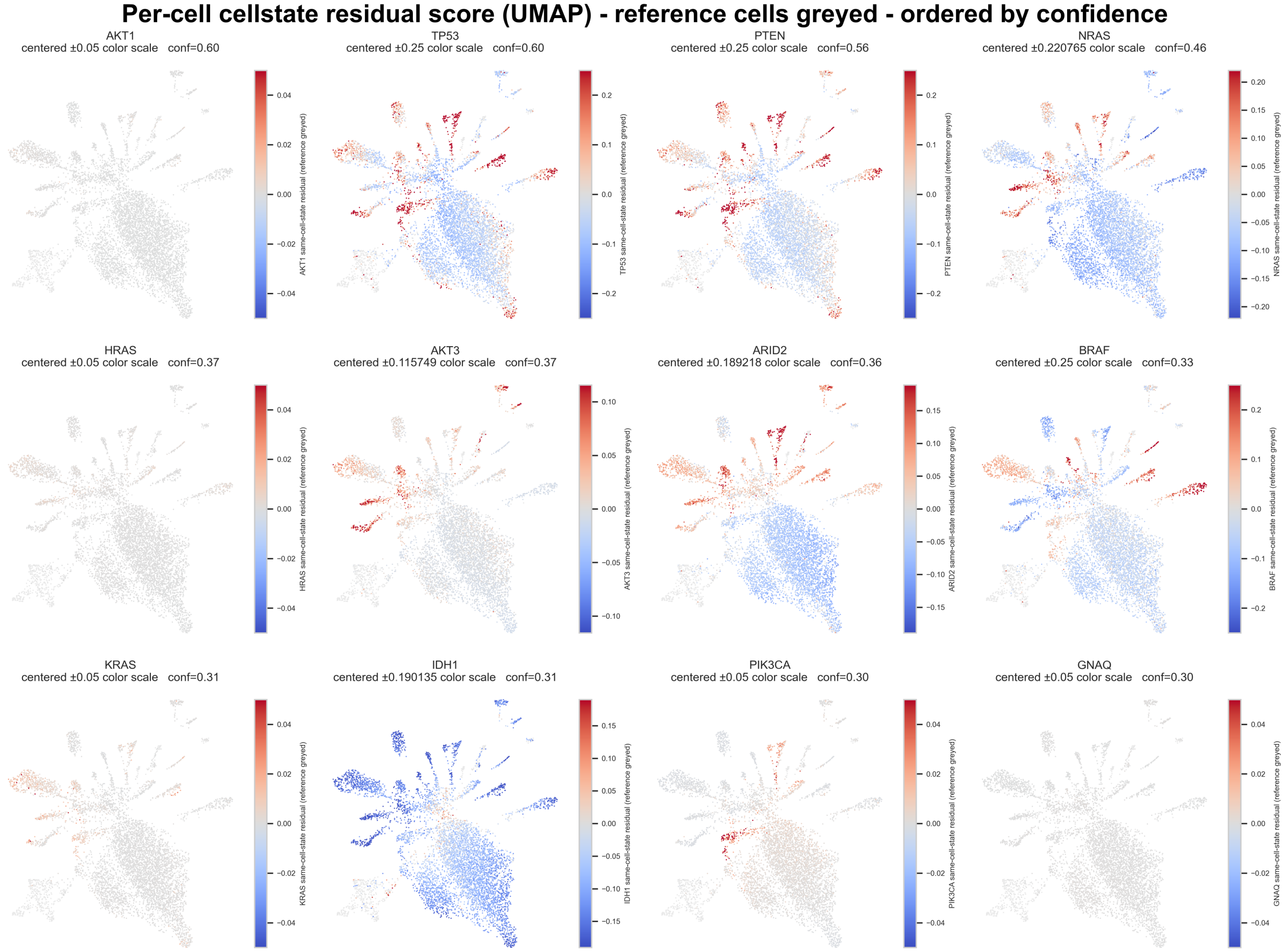


Supplementary Figure S8J. Complete SKCM driver-score atlas.

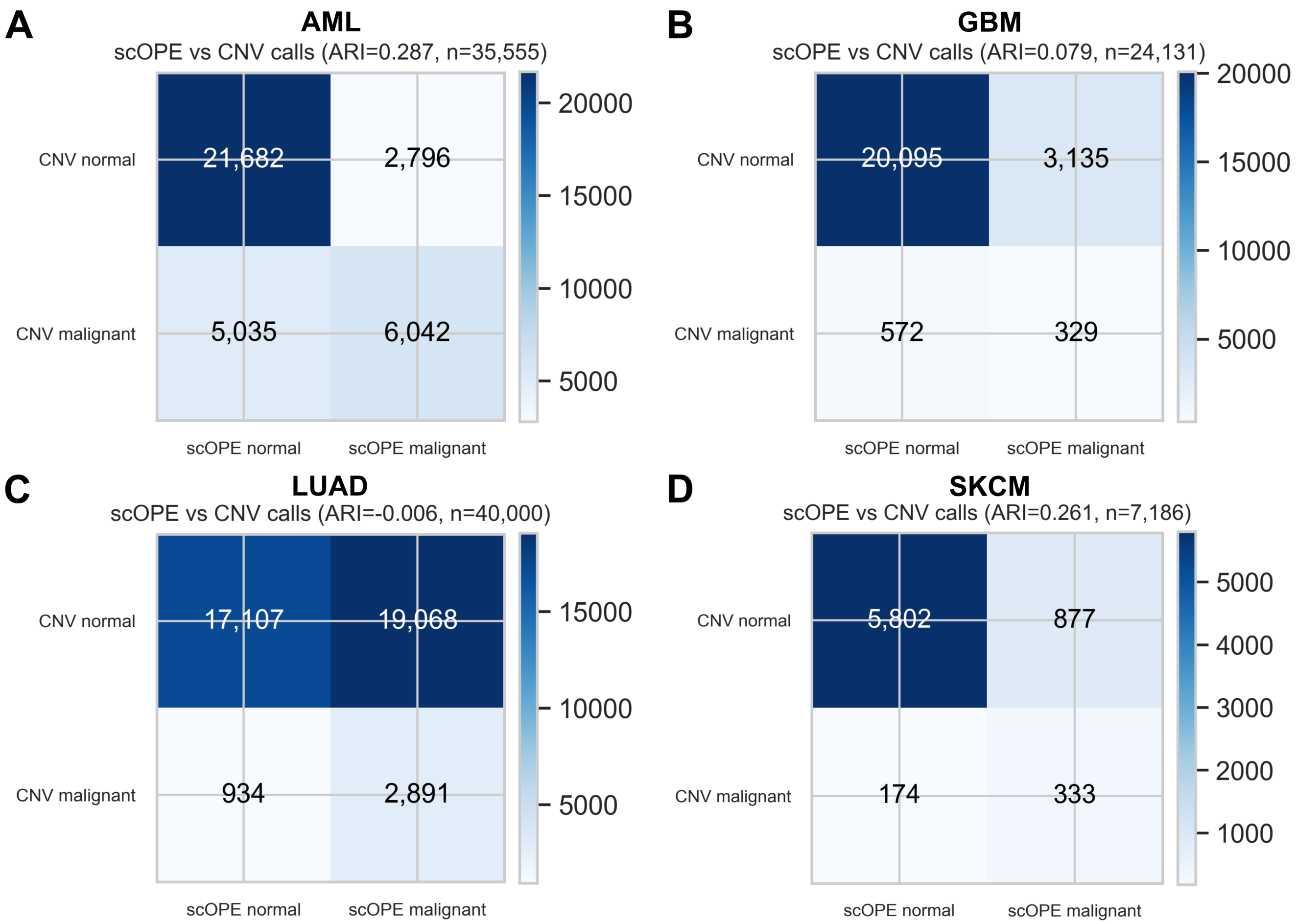


Supplementary Figure S9A. Thresholded scOPE–CNV agreement for cohorts whose confusion matrices are not shown individually in main Figure 7. Panels show AML, GBM, LUAD, and SKCM; adjusted Rand index and matched cell counts are reported in each source panel.

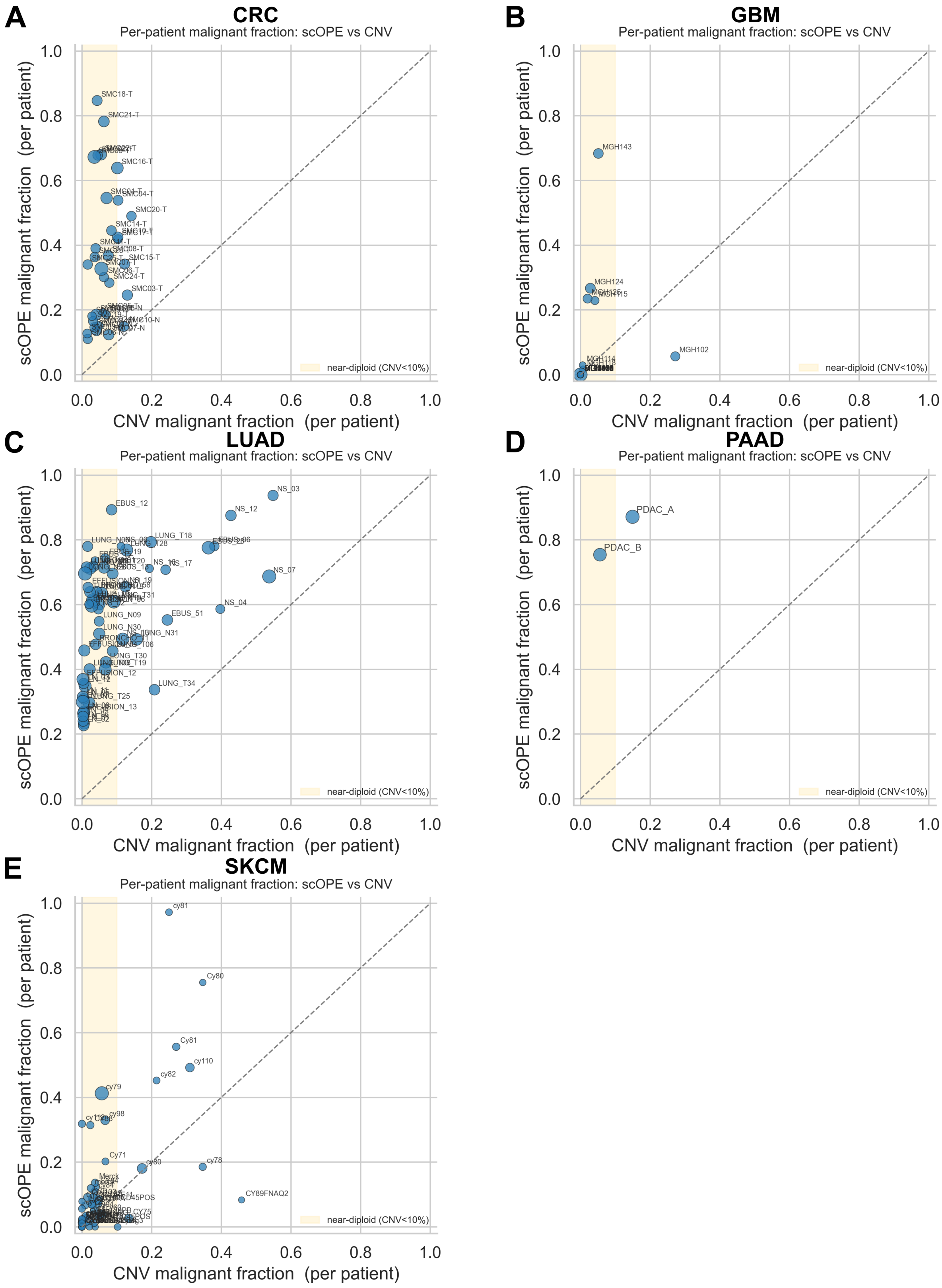


Supplementary Figure S9B. Patient-level malignant fractions for cohorts omitted from main Figure 7E. Per-patient fractions called malignant by scOPE and inferred CNV are shown for CRC, GBM, LUAD, PAAD, and SKCM.

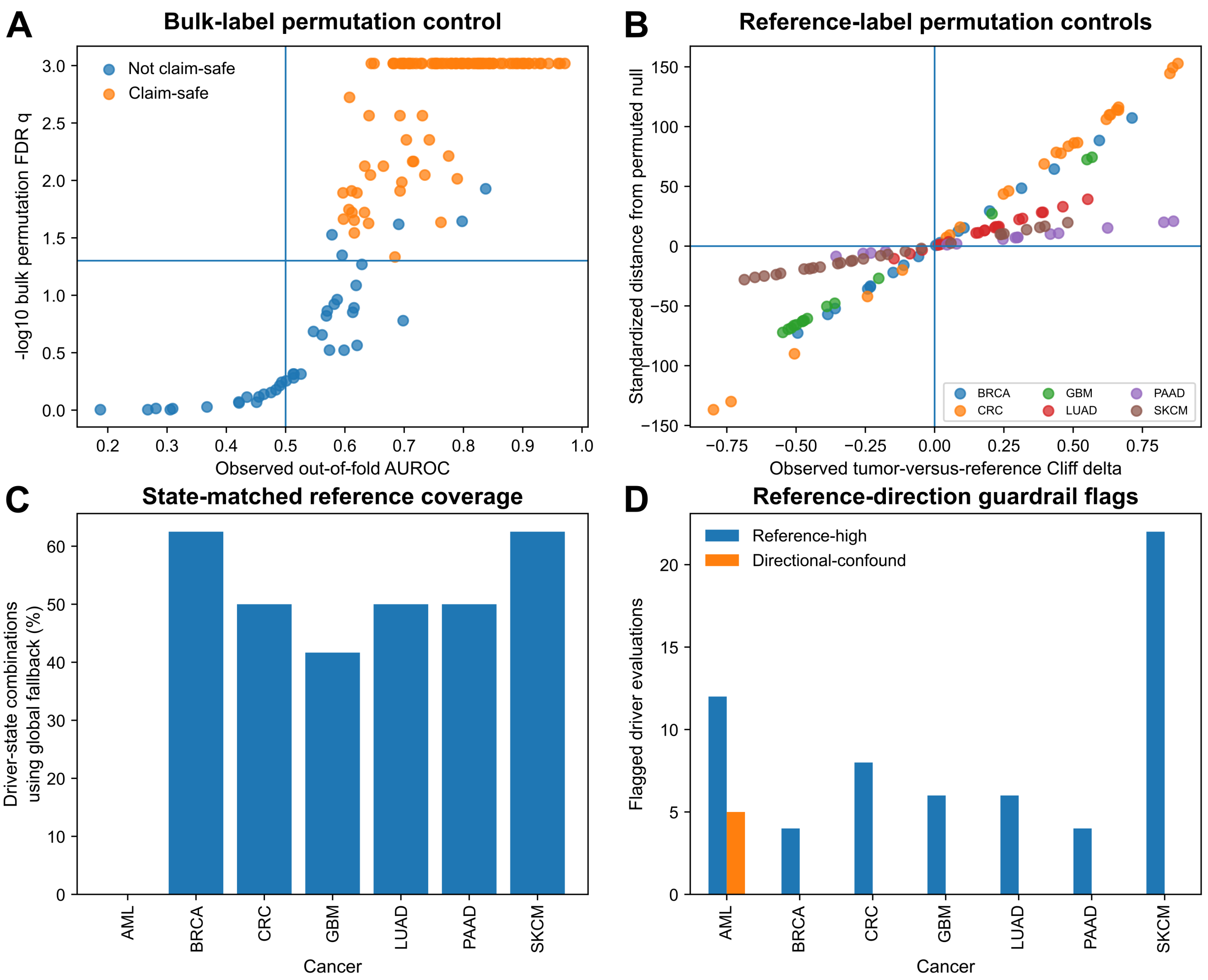


Supplementary Figure S10. Negative controls and reference guardrails present in the canonical output. (A) Observed bulk AUROC versus empirical label-permutation FDR across all 158 attempted models. (B) Observed tumor-versus-reference Cliff's $\delta$ versus its standardized distance from the reference-label permutation null in the six solid-tumor cohorts. (C) Fraction of driver–cell-state combinations that required the prespecified cohort-wide reference fallback because fewer than 25 same-state reference cells were available. (D) Counts of reference-high and directional-confound flags from the reference-direction audit. Patient-block permutations were implemented in the codebase but are not shown because they were not part of the completed canonical result set.

### 3 Supplementary Tables and Data Files

The scOPE Supplementary Tables are provided in two parallel forms: the complete, machine-readable CSV files remain the authoritative data release, and readable typeset versions are included below for direct consultation within the dissertation. The typeset layouts combine closely related fields or use record-style formatting where a single wide grid would require impractically small text; no analytical rows are intentionally filtered. The complete LaTeX table environments are embedded directly in this document so it compiles without separate generated-table fragments. The accompanying CSV files remain the authoritative machine-readable release and preserve full numerical precision.

**Supplementary Table S1. Complete bulk-model audit across 158 driver–cancer models.** AUROC values are pooled out-of-fold estimates with bootstrap intervals. AUPRC is shown with the positive-class prevalence baseline. Permutation and FDR values are from the completed bulk-label permutation analysis. The separate CSV retains every source field at full numerical precision.

| Cancer | Driver | Prior | $n_+$ | $n_-$ | AUROC (95% CI) | AUPRC (base) | Brier | Perm. $p$ | FDR $q$ | Claim-safe |
|---|---|---|---|---|---|---|---|---|---|---|
| AML | *NPM1* | strong | 141 | 510 | 0.971 (0.958–0.982) | 0.872 (0.217) | 0.053 | $5.00 \times 10^{-4}$ | $9.59 \times 10^{-4}$ | Yes |
| AML | *SRSF2* | weak | 72 | 579 | 0.960 (0.937–0.979) | 0.816 (0.111) | 0.047 | $5.00 \times 10^{-4}$ | $9.59 \times 10^{-4}$ | Yes |
| AML | *CEBPA* | intermediate | 37 | 614 | 0.929 (0.893–0.961) | 0.608 (0.057) | 0.054 | $5.00 \times 10^{-4}$ | $9.59 \times 10^{-4}$ | Yes |
| AML | *IDH2* | strong | 73 | 578 | 0.910 (0.876–0.941) | 0.529 (0.112) | 0.088 | $5.00 \times 10^{-4}$ | $9.59 \times 10^{-4}$ | Yes |
| AML | *TP53* | intermediate | 57 | 594 | 0.901 (0.851–0.943) | 0.672 (0.088) | 0.074 | $5.00 \times 10^{-4}$ | $9.59 \times 10^{-4}$ | Yes |
| AML | *IDH1* | strong | 49 | 602 | 0.898 (0.852–0.934) | 0.404 (0.075) | 0.080 | $5.00 \times 10^{-4}$ | $9.59 \times 10^{-4}$ | Yes |
| AML | *RUNX1* | intermediate | 75 | 576 | 0.892 (0.850–0.927) | 0.527 (0.115) | 0.103 | $5.00 \times 10^{-4}$ | $9.59 \times 10^{-4}$ | Yes |
| AML | *KIT* | unspecified | 16 | 635 | 0.844 (0.738–0.928) | 0.153 (0.025) | 0.065 | $5.00 \times 10^{-4}$ | $9.59 \times 10^{-4}$ | Yes |
| AML | *SMC1A* | unspecified | 4 | 517 | 0.838 (0.648–0.985) | 0.285 (0.008) | 0.047 | 0.007 | 0.012 | No |
| AML | *ASXL1* | weak | 63 | 588 | 0.833 (0.776–0.884) | 0.400 (0.097) | 0.137 | $5.00 \times 10^{-4}$ | $9.59 \times 10^{-4}$ | Yes |
| AML | *BCORL1* | unspecified | 13 | 638 | 0.808 (0.679–0.920) | 0.155 (0.020) | 0.041 | $5.00 \times 10^{-4}$ | $9.59 \times 10^{-4}$ | Yes |
| AML | *SF3B1* | weak | 35 | 616 | 0.807 (0.730–0.878) | 0.298 (0.054) | 0.107 | $5.00 \times 10^{-4}$ | $9.59 \times 10^{-4}$ | Yes |
| AML | *DNMT3A* | intermediate | 138 | 513 | 0.797 (0.756–0.837) | 0.515 (0.212) | 0.170 | $5.00 \times 10^{-4}$ | $9.59 \times 10^{-4}$ | Yes |
| AML | *GATA2* | unspecified | 29 | 622 | 0.787 (0.672–0.887) | 0.300 (0.045) | 0.107 | $5.00 \times 10^{-4}$ | $9.59 \times 10^{-4}$ | Yes |
| AML | *WT1* | unspecified | 47 | 604 | 0.779 (0.714–0.846) | 0.223 (0.072) | 0.141 | $5.00 \times 10^{-4}$ | $9.59 \times 10^{-4}$ | Yes |
| AML | *STAG2* | unspecified | 43 | 608 | 0.772 (0.681–0.853) | 0.310 (0.066) | 0.113 | $5.00 \times 10^{-4}$ | $9.59 \times 10^{-4}$ | Yes |
| AML | *TET2* | unspecified | 82 | 569 | 0.728 (0.667–0.791) | 0.349 (0.126) | 0.170 | $5.00 \times 10^{-4}$ | $9.59 \times 10^{-4}$ | Yes |
| AML | *FLT3* | intermediate | 67 | 584 | 0.709 (0.648–0.769) | 0.208 (0.103) | 0.195 | $5.00 \times 10^{-4}$ | $9.59 \times 10^{-4}$ | Yes |
| AML | *KRAS* | intermediate | 34 | 617 | 0.698 (0.603–0.792) | 0.145 (0.052) | 0.156 | $5.00 \times 10^{-4}$ | $9.59 \times 10^{-4}$ | Yes |
| AML | *PTPN11* | unspecified | 32 | 619 | 0.693 (0.605–0.773) | 0.101 (0.049) | 0.153 | $5.00 \times 10^{-4}$ | $9.59 \times 10^{-4}$ | Yes |
| AML | *SMC3* | unspecified | 10 | 641 | 0.691 (0.497–0.852) | 0.051 (0.015) | 0.056 | 0.017 | 0.024 | No |
| AML | *NRAS* | intermediate | 96 | 555 | 0.684 (0.632–0.736) | 0.254 (0.147) | 0.212 | $5.00 \times 10^{-4}$ | $9.59 \times 10^{-4}$ | Yes |
| AML | *BCOR* | unspecified | 34 | 617 | 0.634 (0.527–0.732) | 0.083 (0.052) | 0.162 | 0.004 | 0.008 | Yes |
| AML | *RAD21* | unspecified | 15 | 636 | 0.629 (0.469–0.786) | 0.085 (0.023) | 0.092 | 0.041 | 0.054 | No |
| AML | *U2AF1* | weak | 32 | 619 | 0.620 (0.519–0.720) | 0.102 (0.049) | 0.153 | 0.008 | 0.013 | Yes |
| AML | *ZRSR2* | unspecified | 4 | 517 | 0.599 (0.486–0.692) | 0.011 (0.008) | 0.033 | 0.256 | 0.301 | No |
| AML | *PHF6* | unspecified | 21 | 630 | 0.587 (0.434–0.731) | 0.073 (0.032) | 0.110 | 0.084 | 0.109 | No |
| AML | *EZH2* | unspecified | 19 | 632 | 0.514 (0.374–0.650) | 0.037 (0.029) | 0.163 | 0.421 | 0.485 | No |
| AML | *CBL* | unspecified | 13 | 638 | 0.455 (0.279–0.633) | 0.057 (0.020) | 0.118 | 0.713 | 0.767 | No |
| AML | *ETV6* | unspecified | 0 | 131 | – (——) | – (–) | – | – | – | No |
| AML | *KDM6A* | unspecified | 0 | 0 | – (——) | – (–) | – | – | – | No |
| AML | *SETD2* | unspecified | 0 | 0 | – (——) | – (–) | – | – | – | No |
| AML | *KMT2A* | unspecified | 0 | 0 | – (——) | – (–) | – | – | – | No |
| BRCA | *TP53* | strong | 264 | 639 | 0.931 (0.914–0.946) | 0.834 (0.292) | 0.099 | $5.00 \times 10^{-4}$ | $9.59 \times 10^{-4}$ | Yes |
| BRCA | *CDH1* | intermediate | 102 | 801 | 0.889 (0.852–0.921) | 0.511 (0.113) | 0.122 | $5.00 \times 10^{-4}$ | $9.59 \times 10^{-4}$ | Yes |
| BRCA | *GATA3* | intermediate | 96 | 807 | 0.844 (0.803–0.881) | 0.459 (0.106) | 0.139 | $5.00 \times 10^{-4}$ | $9.59 \times 10^{-4}$ | Yes |
| BRCA | *PIK3CA* | strong | 273 | 630 | 0.827 (0.800–0.855) | 0.632 (0.302) | 0.171 | $5.00 \times 10^{-4}$ | $9.59 \times 10^{-4}$ | Yes |
| BRCA | *AKT1* | unspecified | 24 | 879 | 0.787 (0.707–0.857) | 0.083 (0.027) | 0.098 | $5.00 \times 10^{-4}$ | $9.59 \times 10^{-4}$ | Yes |
| BRCA | *ERBB2* | unspecified | 23 | 880 | 0.751 (0.622–0.866) | 0.199 (0.025) | 0.111 | $5.00 \times 10^{-4}$ | $9.59 \times 10^{-4}$ | Yes |
| BRCA | *ARID1A* | unspecified | 26 | 877 | 0.747 (0.681–0.815) | 0.059 (0.029) | 0.121 | $5.00 \times 10^{-4}$ | $9.59 \times 10^{-4}$ | Yes |
| BRCA | *MAP3K1* | weak | 66 | 837 | 0.730 (0.662–0.796) | 0.271 (0.073) | 0.171 | $5.00 \times 10^{-4}$ | $9.59 \times 10^{-4}$ | Yes |
| BRCA | *RUNX1* | unspecified | 32 | 871 | 0.720 (0.616–0.815) | 0.114 (0.035) | 0.145 | $5.00 \times 10^{-4}$ | $9.59 \times 10^{-4}$ | Yes |
| BRCA | *FOXA1* | unspecified | 27 | 876 | 0.699 (0.599–0.787) | 0.062 (0.030) | 0.143 | $5.00 \times 10^{-4}$ | $9.59 \times 10^{-4}$ | Yes |
| BRCA | *BRCA1* | unspecified | 22 | 881 | 0.693 (0.600–0.773) | 0.043 (0.024) | 0.116 | 0.001 | 0.003 | Yes |
| BRCA | *RB1* | unspecified | 20 | 883 | 0.682 (0.581–0.778) | 0.136 (0.022) | 0.110 | $5.00 \times 10^{-4}$ | $9.59 \times 10^{-4}$ | Yes |
| BRCA | *KMT2C* | unspecified | 77 | 826 | 0.645 (0.584–0.705) | 0.154 (0.085) | 0.209 | $5.00 \times 10^{-4}$ | $9.59 \times 10^{-4}$ | Yes |
| BRCA | *PTEN* | unspecified | 48 | 855 | 0.641 (0.561–0.715) | 0.100 (0.053) | 0.185 | 0.001 | 0.003 | Yes |

Continued on next page

*Supplementary Table S1 continued*

| Cancer | Driver | Prior | $n_+$ | $n_-$ | AUROC (95% CI) | AUPRC (base) | Brier | Perm. $p$ | FDR $q$ | Claim-safe |
|---|---|---|---|---|---|---|---|---|---|---|
| BRCA | *PIK3R1* | unspecified | 23 | 880 | 0.633 (0.504–0.750) | 0.067 (0.025) | 0.129 | 0.013 | 0.019 | Yes |
| BRCA | *ERBB3* | unspecified | 14 | 889 | 0.619 (0.466–0.754) | 0.025 (0.016) | 0.091 | 0.063 | 0.082 | No |
| BRCA | *ATM* | unspecified | 17 | 886 | 0.561 (0.423–0.706) | 0.036 (0.019) | 0.114 | 0.184 | 0.221 | No |
| BRCA | *BRCA2* | unspecified | 20 | 883 | 0.513 (0.377–0.646) | 0.051 (0.022) | 0.129 | 0.424 | 0.485 | No |
| BRCA | *KRAS* | unspecified | 0 | 0 | – (——) | – (–) | – | – | – | No |
| BRCA | *BRAF* | unspecified | 0 | 0 | – (——) | – (–) | – | – | – | No |
| BRCA | *PALB2* | unspecified | 0 | 0 | – (——) | – (–) | – | – | – | No |
| BRCA | *CCND1* | unspecified | 0 | 0 | – (——) | – (–) | – | – | – | No |
| CRC | *BRAF* | strong | 48 | 360 | 0.911 (0.856–0.951) | 0.684 (0.118) | 0.080 | $5.00 \times 10^{-4}$ | $9.59 \times 10^{-4}$ | Yes |
| CRC | *TP53* | intermediate | 239 | 169 | 0.903 (0.868–0.934) | 0.898 (0.586) | 0.118 | $5.00 \times 10^{-4}$ | $9.59 \times 10^{-4}$ | Yes |
| CRC | *MSH2* | unspecified | 12 | 396 | 0.867 (0.772–0.949) | 0.181 (0.029) | 0.046 | $5.00 \times 10^{-4}$ | $9.59 \times 10^{-4}$ | Yes |
| CRC | *APC* | strong | 275 | 133 | 0.850 (0.802–0.893) | 0.872 (0.674) | 0.137 | $5.00 \times 10^{-4}$ | $9.59 \times 10^{-4}$ | Yes |
| CRC | *RNF43* | unspecified | 35 | 373 | 0.838 (0.755–0.910) | 0.473 (0.086) | 0.104 | $5.00 \times 10^{-4}$ | $9.59 \times 10^{-4}$ | Yes |
| CRC | *ARID1A* | unspecified | 41 | 367 | 0.833 (0.762–0.899) | 0.340 (0.100) | 0.119 | $5.00 \times 10^{-4}$ | $9.59 \times 10^{-4}$ | Yes |
| CRC | *KRAS* | strong | 163 | 245 | 0.829 (0.789–0.867) | 0.708 (0.400) | 0.169 | $5.00 \times 10^{-4}$ | $9.59 \times 10^{-4}$ | Yes |
| CRC | *FBXW7* | weak | 57 | 351 | 0.824 (0.768–0.876) | 0.467 (0.140) | 0.139 | $5.00 \times 10^{-4}$ | $9.59 \times 10^{-4}$ | Yes |
| CRC | *KMT2D* | unspecified | 47 | 361 | 0.820 (0.733–0.902) | 0.553 (0.115) | 0.113 | $5.00 \times 10^{-4}$ | $9.59 \times 10^{-4}$ | Yes |
| CRC | *PMS2* | unspecified | 9 | 399 | 0.816 (0.641–0.956) | 0.220 (0.022) | 0.046 | $5.00 \times 10^{-4}$ | $9.59 \times 10^{-4}$ | Yes |
| CRC | *CTNNB1* | unspecified | 26 | 382 | 0.809 (0.715–0.895) | 0.323 (0.064) | 0.087 | $5.00 \times 10^{-4}$ | $9.59 \times 10^{-4}$ | Yes |
| CRC | *SETD2* | unspecified | 26 | 382 | 0.798 (0.695–0.893) | 0.317 (0.064) | 0.104 | $5.00 \times 10^{-4}$ | $9.59 \times 10^{-4}$ | Yes |
| CRC | *PIK3CA* | intermediate | 94 | 314 | 0.788 (0.741–0.834) | 0.481 (0.230) | 0.183 | $5.00 \times 10^{-4}$ | $9.59 \times 10^{-4}$ | Yes |
| CRC | *SOX9* | unspecified | 48 | 360 | 0.781 (0.709–0.844) | 0.333 (0.118) | 0.152 | $5.00 \times 10^{-4}$ | $9.59 \times 10^{-4}$ | Yes |
| CRC | *TGFBR2* | unspecified | 13 | 395 | 0.766 (0.628–0.897) | 0.160 (0.032) | 0.067 | $5.00 \times 10^{-4}$ | $9.59 \times 10^{-4}$ | Yes |
| CRC | *KMT2C* | unspecified | 34 | 374 | 0.755 (0.664–0.836) | 0.257 (0.083) | 0.136 | $5.00 \times 10^{-4}$ | $9.59 \times 10^{-4}$ | Yes |
| CRC | *PTEN* | unspecified | 26 | 382 | 0.754 (0.659–0.836) | 0.149 (0.064) | 0.136 | $5.00 \times 10^{-4}$ | $9.59 \times 10^{-4}$ | Yes |
| CRC | *FAT4* | unspecified | 84 | 324 | 0.753 (0.696–0.810) | 0.420 (0.206) | 0.187 | $5.00 \times 10^{-4}$ | $9.59 \times 10^{-4}$ | Yes |
| CRC | *MSH6* | unspecified | 12 | 396 | 0.742 (0.612–0.860) | 0.101 (0.029) | 0.059 | 0.002 | 0.004 | Yes |
| CRC | *MLH1* | unspecified | 16 | 392 | 0.714 (0.597–0.815) | 0.076 (0.039) | 0.092 | 0.004 | 0.007 | Yes |
| CRC | *SMAD4* | intermediate | 47 | 361 | 0.696 (0.620–0.767) | 0.202 (0.115) | 0.193 | $5.00 \times 10^{-4}$ | $9.59 \times 10^{-4}$ | Yes |
| CRC | *SMAD2* | unspecified | 13 | 395 | 0.693 (0.536–0.835) | 0.104 (0.032) | 0.058 | 0.008 | 0.012 | Yes |
| CRC | *TCF7L2* | unspecified | 41 | 367 | 0.611 (0.528–0.694) | 0.143 (0.100) | 0.198 | 0.008 | 0.012 | Yes |
| CRC | *NRAS* | unspecified | 20 | 388 | 0.501 (0.363–0.624) | 0.056 (0.049) | 0.154 | 0.494 | 0.556 | No |
| CRC | *ERBB2* | unspecified | 10 | 398 | 0.421 (0.246–0.608) | 0.024 (0.025) | 0.104 | 0.802 | 0.850 | No |
| GBM | *IDH1* | strong | 7 | 148 | 0.963 (0.914–0.996) | 0.633 (0.045) | 0.030 | $5.00 \times 10^{-4}$ | $9.59 \times 10^{-4}$ | Yes |
| GBM | *ATRX* | unspecified | 10 | 145 | 0.862 (0.769–0.943) | 0.384 (0.065) | 0.053 | $5.00 \times 10^{-4}$ | $9.59 \times 10^{-4}$ | Yes |
| GBM | *TP53* | unspecified | 51 | 104 | 0.829 (0.753–0.894) | 0.747 (0.329) | 0.163 | $5.00 \times 10^{-4}$ | $9.59 \times 10^{-4}$ | Yes |
| GBM | *PDGFRA* | intermediate | 7 | 148 | 0.790 (0.584–0.948) | 0.203 (0.045) | 0.047 | 0.006 | 0.010 | Yes |
| GBM | *RB1* | unspecified | 13 | 142 | 0.731 (0.572–0.880) | 0.380 (0.084) | 0.091 | 0.001 | 0.003 | Yes |
| GBM | *EGFR* | strong | 44 | 111 | 0.715 (0.629–0.795) | 0.439 (0.284) | 0.218 | $5.00 \times 10^{-4}$ | $9.59 \times 10^{-4}$ | Yes |
| GBM | *NF1* | intermediate | 16 | 139 | 0.696 (0.528–0.844) | 0.298 (0.103) | 0.096 | 0.006 | 0.010 | Yes |
| GBM | *PTEN* | weak | 50 | 105 | 0.612 (0.515–0.701) | 0.399 (0.323) | 0.247 | 0.013 | 0.019 | Yes |
| GBM | *STAG2* | unspecified | 7 | 148 | 0.514 (0.318–0.704) | 0.053 (0.045) | 0.066 | 0.461 | 0.524 | No |
| GBM | *PIK3R1* | unspecified | 12 | 143 | 0.463 (0.276–0.652) | 0.080 (0.077) | 0.140 | 0.671 | 0.728 | No |
| GBM | *PIK3CA* | unspecified | 14 | 141 | 0.421 (0.280–0.574) | 0.088 (0.090) | 0.134 | 0.830 | 0.866 | No |
| GBM | *IDH2* | unspecified | 0 | 0 | – (——) | – (–) | – | – | – | No |
| GBM | *CDKN2A* | unspecified | 1 | 123 | – (——) | – (–) | – | – | – | No |
| GBM | *CIC* | unspecified | 1 | 123 | – (——) | – (–) | – | – | – | No |
| GBM | *TERT* | unspecified | 0 | 93 | – (——) | – (–) | – | – | – | No |
| GBM | *MDM2* | unspecified | 0 | 0 | – (——) | – (–) | – | – | – | No |
| GBM | *MDM4* | unspecified | 0 | 124 | – (——) | – (–) | – | – | – | No |
| LUAD | *KEAP1* | intermediate | 91 | 476 | 0.920 (0.888–0.948) | 0.701 (0.160) | 0.093 | $5.00 \times 10^{-4}$ | $9.59 \times 10^{-4}$ | Yes |

Continued on next page

*Supplementary Table S1 continued*

| **Cancer** | **Driver** | **Prior** | $n_+$ | $n_-$ | **AUROC (95% CI)** | **AUPRC (base)** | **Brier** | **Perm.** $p$ | **FDR** $q$ | **Claim-safe** |
|---|---|---|---|---|---|---|---|---|---|---|
| LUAD | *TP53* | intermediate | 257 | 310 | 0.883 (0.854–0.909) | 0.828 (0.453) | 0.139 | $5.00 \times 10^{-4}$ | $9.59 \times 10^{-4}$ | Yes |
| LUAD | *STK11* | intermediate | 69 | 498 | 0.878 (0.836–0.915) | 0.468 (0.122) | 0.119 | $5.00 \times 10^{-4}$ | $9.59 \times 10^{-4}$ | Yes |
| LUAD | *EGFR* | strong | 67 | 500 | 0.850 (0.800–0.895) | 0.494 (0.118) | 0.125 | $5.00 \times 10^{-4}$ | $9.59 \times 10^{-4}$ | Yes |
| LUAD | *KRAS* | strong | 148 | 419 | 0.845 (0.810–0.880) | 0.623 (0.261) | 0.151 | $5.00 \times 10^{-4}$ | $9.59 \times 10^{-4}$ | Yes |
| LUAD | *ERBB2* | unspecified | 4 | 450 | 0.798 (0.595–0.992) | 0.305 (0.009) | 0.038 | 0.016 | 0.023 | No |
| LUAD | *SMARCA4* | unspecified | 42 | 525 | 0.797 (0.718–0.869) | 0.323 (0.074) | 0.124 | $5.00 \times 10^{-4}$ | $9.59 \times 10^{-4}$ | Yes |
| LUAD | *RB1* | unspecified | 27 | 540 | 0.796 (0.686–0.889) | 0.318 (0.048) | 0.082 | $5.00 \times 10^{-4}$ | $9.59 \times 10^{-4}$ | Yes |
| LUAD | *U2AF1* | unspecified | 13 | 554 | 0.716 (0.584–0.831) | 0.068 (0.023) | 0.075 | 0.004 | 0.007 | Yes |
| LUAD | *ATM* | unspecified | 41 | 526 | 0.700 (0.606–0.793) | 0.220 (0.072) | 0.149 | $5.00 \times 10^{-4}$ | $9.59 \times 10^{-4}$ | Yes |
| LUAD | *PTPRD* | unspecified | 84 | 483 | 0.682 (0.618–0.739) | 0.273 (0.148) | 0.202 | $5.00 \times 10^{-4}$ | $9.59 \times 10^{-4}$ | Yes |
| LUAD | *NF1* | unspecified | 58 | 509 | 0.649 (0.574–0.722) | 0.177 (0.102) | 0.204 | $5.00 \times 10^{-4}$ | $9.59 \times 10^{-4}$ | Yes |
| LUAD | *SETD2* | unspecified | 33 | 534 | 0.643 (0.537–0.736) | 0.157 (0.058) | 0.157 | 0.005 | 0.009 | Yes |
| LUAD | *MET* | unspecified | 20 | 547 | 0.640 (0.503–0.770) | 0.070 (0.035) | 0.089 | 0.017 | 0.024 | Yes |
| LUAD | *PIK3CA* | unspecified | 27 | 540 | 0.616 (0.508–0.722) | 0.075 (0.048) | 0.131 | 0.015 | 0.022 | Yes |
| LUAD | *BRAF* | unspecified | 41 | 526 | 0.607 (0.513–0.693) | 0.102 (0.072) | 0.194 | 0.012 | 0.018 | Yes |
| LUAD | *MGA* | unspecified | 40 | 527 | 0.598 (0.502–0.695) | 0.168 (0.071) | 0.190 | 0.015 | 0.022 | Yes |
| LUAD | *RBM10* | unspecified | 34 | 533 | 0.547 (0.445–0.650) | 0.073 (0.060) | 0.174 | 0.170 | 0.207 | No |
| LUAD | *ARID1A* | unspecified | 30 | 537 | 0.484 (0.375–0.593) | 0.056 (0.053) | 0.181 | 0.605 | 0.666 | No |
| LUAD | *CDKN2A* | unspecified | 20 | 547 | 0.475 (0.340–0.611) | 0.042 (0.035) | 0.158 | 0.643 | 0.703 | No |
| PAAD | *KRAS* | strong | 111 | 61 | 0.944 (0.907–0.975) | 0.969 (0.645) | 0.086 | $5.00 \times 10^{-4}$ | $9.59 \times 10^{-4}$ | Yes |
| PAAD | *TP53* | intermediate | 103 | 69 | 0.779 (0.705–0.848) | 0.816 (0.599) | 0.189 | $5.00 \times 10^{-4}$ | $9.59 \times 10^{-4}$ | Yes |
| PAAD | *GNAS* | unspecified | 9 | 163 | 0.775 (0.646–0.883) | 0.133 (0.052) | 0.079 | 0.003 | 0.006 | Yes |
| PAAD | *KDM6A* | unspecified | 6 | 166 | 0.762 (0.592–0.919) | 0.245 (0.035) | 0.044 | 0.016 | 0.023 | Yes |
| PAAD | *SMAD4* | intermediate | 36 | 136 | 0.756 (0.677–0.829) | 0.402 (0.209) | 0.177 | $5.00 \times 10^{-4}$ | $9.59 \times 10^{-4}$ | Yes |
| PAAD | *CDKN2A* | intermediate | 34 | 138 | 0.710 (0.625–0.787) | 0.307 (0.198) | 0.201 | $5.00 \times 10^{-4}$ | $9.59 \times 10^{-4}$ | Yes |
| PAAD | *TGFBR2* | unspecified | 8 | 164 | 0.684 (0.507–0.843) | 0.089 (0.047) | 0.088 | 0.035 | 0.046 | Yes |
| PAAD | *MAP2K4* | unspecified | 4 | 168 | 0.621 (0.369–0.872) | 0.069 (0.023) | 0.053 | 0.229 | 0.273 | No |
| PAAD | *RNF43* | unspecified | 11 | 161 | 0.613 (0.406–0.804) | 0.118 (0.064) | 0.098 | 0.113 | 0.141 | No |
| PAAD | *KMT2D* | unspecified | 7 | 165 | 0.574 (0.390–0.752) | 0.060 (0.041) | 0.070 | 0.254 | 0.300 | No |
| PAAD | *ARID1A* | unspecified | 8 | 164 | 0.526 (0.370–0.682) | 0.053 (0.047) | 0.089 | 0.421 | 0.485 | No |
| PAAD | *ROBO2* | unspecified | 4 | 168 | 0.281 (0.193–0.381) | 0.020 (0.023) | 0.051 | 0.941 | 0.968 | No |
| PAAD | *ATM* | unspecified | 7 | 165 | 0.268 (0.108–0.441) | 0.030 (0.041) | 0.058 | 0.988 | 0.992 | No |
| PAAD | *STK11* | unspecified | 4 | 168 | 0.188 (0.088–0.318) | 0.017 (0.023) | 0.044 | 0.985 | 0.992 | No |
| PAAD | *BRCA2* | unspecified | 0 | 138 | – (——) | – (–) | – | – | – | No |
| SKCM | *NRAS* | strong | 128 | 338 | 0.843 (0.803–0.877) | 0.651 (0.275) | 0.153 | $5.00 \times 10^{-4}$ | $9.59 \times 10^{-4}$ | Yes |
| SKCM | *BRAF* | strong | 240 | 226 | 0.824 (0.787–0.860) | 0.838 (0.515) | 0.171 | $5.00 \times 10^{-4}$ | $9.59 \times 10^{-4}$ | Yes |
| SKCM | *TP53* | weak | 70 | 396 | 0.801 (0.736–0.857) | 0.504 (0.150) | 0.141 | $5.00 \times 10^{-4}$ | $9.59 \times 10^{-4}$ | Yes |
| SKCM | *PTEN* | unspecified | 44 | 422 | 0.762 (0.680–0.832) | 0.296 (0.094) | 0.143 | $5.00 \times 10^{-4}$ | $9.59 \times 10^{-4}$ | Yes |
| SKCM | *GNAQ* | unspecified | 10 | 456 | 0.735 (0.561–0.880) | 0.076 (0.021) | 0.039 | 0.005 | 0.009 | Yes |
| SKCM | *NF1* | intermediate | 68 | 398 | 0.707 (0.636–0.773) | 0.302 (0.146) | 0.190 | $5.00 \times 10^{-4}$ | $9.59 \times 10^{-4}$ | Yes |
| SKCM | *KIT* | unspecified | 21 | 445 | 0.704 (0.583–0.816) | 0.105 (0.045) | 0.111 | 0.002 | 0.004 | Yes |
| SKCM | *AKT1* | unspecified | 3 | 369 | 0.698 (0.523–0.837) | 0.018 (0.008) | 0.019 | 0.136 | 0.166 | No |
| SKCM | *IDH1* | unspecified | 23 | 443 | 0.665 (0.528–0.788) | 0.163 (0.049) | 0.107 | 0.004 | 0.008 | Yes |
| SKCM | *MAP2K1* | unspecified | 27 | 439 | 0.616 (0.517–0.715) | 0.082 (0.058) | 0.152 | 0.021 | 0.029 | Yes |
| SKCM | *PIK3CA* | unspecified | 11 | 455 | 0.615 (0.464–0.765) | 0.047 (0.024) | 0.051 | 0.101 | 0.129 | No |
| SKCM | *PREX2* | unspecified | 97 | 369 | 0.608 (0.543–0.673) | 0.276 (0.208) | 0.233 | $10.00 \times 10^{-4}$ | 0.002 | Yes |
| SKCM | *CDKN2A* | unspecified | 61 | 405 | 0.597 (0.527–0.666) | 0.179 (0.131) | 0.214 | 0.008 | 0.013 | Yes |
| SKCM | *APC* | unspecified | 39 | 427 | 0.595 (0.498–0.687) | 0.138 (0.084) | 0.196 | 0.033 | 0.045 | No |
| SKCM | *CTNNB1* | unspecified | 24 | 442 | 0.582 (0.461–0.694) | 0.079 (0.052) | 0.156 | 0.093 | 0.120 | No |
| SKCM | *ARID2* | unspecified | 61 | 405 | 0.578 (0.494–0.661) | 0.277 (0.131) | 0.208 | 0.022 | 0.030 | No |
| SKCM | *RAC1* | unspecified | 29 | 437 | 0.570 (0.460–0.674) | 0.084 (0.062) | 0.182 | 0.109 | 0.137 | No |

Continued on next page

*Supplementary Table S1 continued*

| **Cancer** | **Driver** | **Prior** | $n_+$ | $n_-$ | **AUROC (95% CI)** | **AUPRC (base)** | **Brier** | **Perm.** $p$ | **FDR** $q$ | **Claim-safe** |
|---|---|---|---|---|---|---|---|---|---|---|
| SKCM | *SETD2* | unspecified | 26 | 440 | 0.569 (0.452–0.689) | 0.078 (0.056) | 0.171 | 0.122 | 0.151 | No |
| SKCM | *KRAS* | unspecified | 11 | 455 | 0.494 (0.325–0.665) | 0.035 (0.024) | 0.071 | 0.511 | 0.572 | No |
| SKCM | *ARID1A* | unspecified | 23 | 443 | 0.490 (0.351–0.625) | 0.054 (0.049) | 0.171 | 0.552 | 0.613 | No |
| SKCM | *PPP6C* | unspecified | 31 | 435 | 0.451 (0.333–0.566) | 0.063 (0.067) | 0.208 | 0.812 | 0.854 | No |
| SKCM | *AKT3* | unspecified | 8 | 458 | 0.435 (0.241–0.623) | 0.017 (0.017) | 0.067 | 0.722 | 0.770 | No |
| SKCM | *MAP2K2* | unspecified | 9 | 457 | 0.367 (0.196–0.561) | 0.017 (0.019) | 0.066 | 0.906 | 0.939 | No |
| SKCM | *HRAS* | unspecified | 6 | 460 | 0.309 (0.105–0.552) | 0.011 (0.013) | 0.026 | 0.950 | 0.970 | No |
| SKCM | *GNA11* | unspecified | 12 | 454 | 0.305 (0.147–0.480) | 0.019 (0.026) | 0.115 | 0.992 | 0.992 | No |
| SKCM | *KDM6A* | unspecified | 1 | 279 | – (——) | – (–) | – | – | – | No |

**Supplementary Table S2. Data sources, accessions, and cohort characteristics.** Data sources, accessions, access conditions, analysis inputs, cohort sizes, and single-cell reference annotations. Each cohort is presented as a readable record rather than an impractically wide grid. The separate CSV preserves the normalized field structure for programmatic use.

| Domain / cohort | Source, access, analysis input, and retained cohort details |
|---|---|
| **bulk / AML / Beat-AML** | **Publication:** Functional genomic landscape of acute myeloid leukaemia; Integrative analysis of drug response and clinical outcome in acute myeloid leukemia<br>**DOI:** `10.1038/s41586-018-0623-z;10.1016/j.ccell.2022.07.002`<br>**Repository:** NCI Genomic Data Commons / dbGaP / Vizome<br>**Accession/project:** `BEATAML1.0-COHORT;phs001657`<br>**Access:** Processed/open resources are available through GDC/Vizome; controlled sequence-level data require authorized access.<br>**Analysis input:** Previously derived normalized/log-transformed bulk RNA-seq expression matrix and somatic mutation-label matrix; exact local filenames are recorded in config_canonical_v3.json.<br>**Modeled/retained:** 651<br>**Provenance:** Project-level source identifier; original portal-specific download manifest/file UUID was not retained. |
| **bulk / BRCA / TCGA-BRCA** | **Publication:** Comprehensive molecular portraits of human breast tumours<br>**DOI:** `10.1038/nature11412`<br>**Repository:** NCI Genomic Data Commons<br>**Accession/project:** `TCGA-BRCA`<br>**Access:** Open processed data and controlled raw human sequencing data.<br>**Analysis input:** Previously derived normalized/log-transformed bulk RNA-seq expression matrix and somatic mutation-label matrix; exact local filenames are recorded in config_canonical_v3.json.<br>**Modeled/retained:** 903<br>**Provenance:** Project-level source identifier; original portal-specific download manifest/file UUID was not retained. |
| **bulk / CRC / TCGA-COAD/TCGA-READ** | **Publication:** Comprehensive molecular characterization of human colon and rectal cancer<br>**DOI:** `10.1038/nature11252`<br>**Repository:** NCI Genomic Data Commons<br>**Accession/project:** `TCGA-COAD;TCGA-READ`<br>**Access:** Open processed data and controlled raw human sequencing data.<br>**Analysis input:** Previously derived normalized/log-transformed bulk RNA-seq expression matrix and somatic mutation-label matrix; exact local filenames are recorded in config_canonical_v3.json.<br>**Modeled/retained:** 408<br>**Provenance:** Project-level source identifier; original portal-specific download manifest/file UUID was not retained. |
| **bulk / GBM / TCGA-GBM** | **Publication:** Integrated genomic analysis identifies clinically relevant subtypes of glioblastoma characterized by abnormalities in PDGFRA, IDH1, EGFR, and NF1<br>**DOI:** `10.1016/j.ccr.2009.12.020`<br>**Repository:** NCI Genomic Data Commons<br>**Accession/project:** `TCGA-GBM`<br>**Access:** Open processed data and controlled raw human sequencing data.<br>**Analysis input:** Previously derived normalized/log-transformed bulk RNA-seq expression matrix and somatic mutation-label matrix; exact local filenames are recorded in config_canonical_v3.json.<br>**Modeled/retained:** 155<br>**Provenance:** Project-level source identifier; original portal-specific download manifest/file UUID was not retained. |

Continued on next page

| Domain / cohort | Source, access, analysis input, and retained cohort details |
|---|---|
| **bulk / LUAD / TCGA-LUAD** | **Publication:** Comprehensive molecular profiling of lung adenocarcinoma<br>**DOI:** `10.1038/nature13385`<br>**Repository:** NCI Genomic Data Commons<br>**Accession/project:** `TCGA-LUAD`<br>**Access:** Open processed data and controlled raw human sequencing data.<br>**Analysis input:** Previously derived normalized/log-transformed bulk RNA-seq expression matrix and somatic mutation-label matrix; exact local filenames are recorded in config_canonical_v3.json.<br>**Modeled/retained:** 567<br>**Provenance:** Project-level source identifier; original portal-specific download manifest/file UUID was not retained. |
| **bulk / PAAD / TCGA-PAAD** | **Publication:** Integrated genomic characterization of pancreatic ductal adenocarcinoma<br>**DOI:** `10.1016/j.ccell.2017.07.007`<br>**Repository:** NCI Genomic Data Commons<br>**Accession/project:** `TCGA-PAAD`<br>**Access:** Open processed data and controlled raw human sequencing data.<br>**Analysis input:** Previously derived normalized/log-transformed bulk RNA-seq expression matrix and somatic mutation-label matrix; exact local filenames are recorded in config_canonical_v3.json.<br>**Modeled/retained:** 172<br>**Provenance:** Project-level source identifier; original portal-specific download manifest/file UUID was not retained. |
| **bulk / SKCM / TCGA-SKCM** | **Publication:** Genomic classification of cutaneous melanoma<br>**DOI:** `10.1016/j.cell.2015.05.044`<br>**Repository:** NCI Genomic Data Commons<br>**Accession/project:** `TCGA-SKCM`<br>**Access:** Open processed data and controlled raw human sequencing data.<br>**Analysis input:** Previously derived normalized/log-transformed bulk RNA-seq expression matrix and somatic mutation-label matrix; exact local filenames are recorded in config_canonical_v3.json.<br>**Modeled/retained:** 466<br>**Provenance:** Project-level source identifier; original portal-specific download manifest/file UUID was not retained. |
| **single-cell / AML / van Galen** | **Publication:** Single-Cell RNA-Seq Reveals AML Hierarchies Relevant to Disease Progression and Immunity<br>**DOI:** `10.1016/j.cell.2019.01.031`<br>**Repository:** NCBI GEO / SRA<br>**Accession/project:** `GSE116256;PRJNA477870;SRP183188`<br>**Access:** Processed data are public in GEO; raw reads are available in SRA.<br>**Analysis input:** Processed, quality-controlled, normalized, log-transformed AnnData object; exact local filename is recorded in config_canonical_v3.json.<br>**Modeled/retained:** 35,843<br>**CNV-matched cells:** 35,555<br>**Reference cells:** 5,946<br>**Reference annotation:** `CellType`<br>**Reference categories:** Prog (1,076); T (697); ProMono (610); earlyEry (601); Mono (556); GMP (522); HSC (493); cDC (272); CTL (257); lateEry (254); ProB (188); NK (152); B (113); pDC (90); Plasma (65); GMP-like (0); HSC-like (0); Mono-like (0); ProMono-like (0); Prog-like (0); cDC-like (0)<br>**Provenance:** Source attribution verified against repository accession, sample count, and source publication. |

Continued on next page

| Domain / cohort | Source, access, analysis input, and retained cohort details |
|---|---|
| **single-cell / BRCA / Wu** | **Publication:** A single-cell and spatially resolved atlas of human breast cancers<br>**DOI:** `10.1038/s41588-021-00911-1`<br>**Repository:** NCBI GEO / EGA<br>**Accession/project:** `GSE176078;EGAS00001005173`<br>**Access:** Processed data are public in GEO; raw human data require controlled EGA access.<br>**Analysis input:** Processed, quality-controlled, normalized, log-transformed AnnData object; exact local filename is recorded in config_canonical_v3.json.<br>**Modeled/retained:** 39,998<br>**CNV-matched cells:** 39,998<br>**Reference cells:** 9,529<br>**Reference annotation:** `celltype_major`<br>**Reference categories:** Myeloid (3,841); Endothelial (3,103); CAFs (2,585); B-cells (0); Cancer Epithelial (0); Normal Epithelial (0); PVL (0); Plasmablasts (0); T-cells (0)<br>**Provenance:** Source attribution verified against repository accession, sample count, and source publication. |
| **single-cell / CRC / Lee** | **Publication:** Lineage-dependent gene expression programs influence the immune landscape of colorectal cancer<br>**DOI:** `10.1038/s41588-020-0636-z`<br>**Repository:** NCBI GEO<br>**Accession/project:** `GSE132465;PRJNA548146`<br>**Access:** Processed matrices and annotations are public in GEO; raw reads were not provided in the GEO record because of patient-privacy constraints.<br>**Analysis input:** Processed, quality-controlled, normalized, log-transformed AnnData object; exact local filename is recorded in config_canonical_v3.json.<br>**Modeled/retained:** 40,000<br>**CNV-matched cells:** 40,000<br>**Reference cells:** 20,435<br>**Reference annotation:** `Cell_type`<br>**Reference categories:** T cells (14,523); B cells (5,789); Mast cells (123); Epithelial cells (0); Myeloids (0); Stromal cells (0)<br>**Provenance:** Source attribution verified against repository accession, sample count, and source publication. |
| **single-cell / GBM / Neftel** | **Publication:** An Integrative Model of Cellular States, Plasticity, and Genetics for Glioblastoma<br>**DOI:** `10.1016/j.cell.2019.06.024`<br>**Repository:** NCBI GEO<br>**Accession/project:** `GSE131928;PRJNA545332`<br>**Access:** Processed data and metadata are public in GEO; raw reads were not provided in the GEO record.<br>**Analysis input:** Processed, quality-controlled, normalized, log-transformed AnnData object; exact local filename is recorded in config_canonical_v3.json.<br>**Modeled/retained:** 24,131<br>**CNV-matched cells:** 24,131<br>**Reference cells:** 10,680<br>**Reference annotation:** `celltype`<br>**Reference categories:** T cell (2,560); Plasma (2,047); NK cell (1,730); B cell (1,203); Dendritic (1,151); Endothelial (1,113); Myeloid (876); Epithelial (0); Erythroid (0); Fibroblast/CAF (0); HSPC/Progenitor (0); Unassigned (0)<br>**Provenance:** Source attribution verified against repository accession, sample count, and source publication. |

Continued on next page

*Supplementary Table S2 continued*

| Domain / cohort | Source, access, analysis input, and retained cohort details |
|---|---|
| **single-cell / LUAD / Lambrechts** | **Publication:** Phenotype molding of stromal cells in the lung tumor microenvironment<br>**DOI:** `10.1038/s41591-018-0096-5`<br>**Repository:** EMBL-EBI BioStudies/ArrayExpress<br>**Accession/project:** `E-MTAB-6149`<br>**Access:** The source study record and deposited files are public in BioStudies/ArrayExpress.<br>**Analysis input:** Processed, quality-controlled, normalized, log-transformed AnnData object; exact local filename is recorded in config_canonical_v3.json.<br>**Modeled/retained:** 40,000<br>**CNV-matched cells:** 40,000<br>**Reference cells:** 1,783<br>**Reference annotation:** `celltype`<br>**Reference categories:** Fibroblasts (700); MAST cells (666); Endothelial cells (417); B lymphocytes (0); Epithelial cells (0); Myeloid cells (0); T/NK cells (0); nan (0)<br>**Provenance:** Source attribution verified against repository accession, sample count, and source publication. |
| **single-cell / PAAD / Moncada** | **Publication:** Integrating microarray-based spatial transcriptomics and single-cell RNA-seq reveals tissue architecture in pancreatic ductal adenocarcinomas<br>**DOI:** `10.1038/s41587-019-0392-8`<br>**Repository:** NCBI GEO / SRA<br>**Accession/project:** `GSE111672;PRJNA437847;SRP134863`<br>**Access:** Processed data are public in GEO; raw reads are available in SRA.<br>**Analysis input:** Processed, quality-controlled, normalized, log-transformed AnnData object; exact local filename is recorded in config_canonical_v3.json.<br>**Modeled/retained:** 3,659<br>**CNV-matched cells:** 3,659<br>**Reference cells:** 208<br>**Reference annotation:** `celltype`<br>**Reference categories:** endothelial (170); myeloid (38); other (0); stromal (0)<br>**Provenance:** Source attribution verified against repository accession, sample count, and source publication. |
| **single-cell / SKCM / Jerby-Arnon** | **Publication:** A cancer cell program promotes T cell exclusion and resistance to checkpoint blockade<br>**DOI:** `10.1016/j.cell.2018.09.006`<br>**Repository:** NCBI GEO / dbGaP<br>**Accession/project:** `GSE115978;PRJNA476644`<br>**Access:** Processed counts, TPM values, and annotations are public in GEO; the GEO record states that raw reads are to be made available through dbGaP and does not provide them directly.<br>**Analysis input:** Processed, quality-controlled, normalized, log-transformed AnnData object; exact local filename is recorded in config_canonical_v3.json.<br>**Modeled/retained:** 7,186<br>**CNV-matched cells:** 7,186<br>**Reference cells:** 618<br>**Reference annotation:** `celltype`<br>**Reference categories:** Macrophage (420); CAF (106); NK (92); ? (0); B.cell (0); Endo. (0); Mal (0); T.CD4 (0); T.CD8 (0); T.cell (0)<br>**Provenance:** Source attribution verified against repository accession, sample count, and source publication. |

**Supplementary Table S3. AML per-cell expressed-mutation truth validation.** AML expressed-mutation direct-label validation for all 12 evaluable drivers. AUROCs are apparent positive–unlabeled discrimination values; intervals, Mann–Whitney tests, Cliff's $\delta$, and FDR values are reported from the canonical analysis. The final columns identify the exact score and truth fields.

| Driver | $n_{\text{mut}}$ | $n_{\text{WT}}$ | AUROC (95% CI) | MW $p$ | Cliff's $\delta$ (95% CI) | FDR $q$ | Score column / mode | Truth / metric type |
|---|---|---|---|---|---|---|---|---|
| *FLT3* | 46 | 124 | 0.558 (0.456–0.660) | 0.248 | 0.116 (-0.088–0.320) | 0.425 | `mutation_resid_FLT3` cellstate__residual__raw; cell-state__residual | `FLT3_genotype` apparent__positive__unlabeled |
| *NPM1* | 542 | 1,457 | 0.753 (0.730–0.776) | $6.11 \times 10^{-68}$ | 0.506 (0.460–0.551) | $7.34 \times 10^{-67}$ | `mutation_resid_NPM1` cellstate__residual__raw; cell-state__residual | `NPM1_genotype` apparent__positive__unlabeled |
| *DNMT3A* | 146 | 404 | 0.714 (0.669–0.761) | $1.56 \times 10^{-14}$ | 0.429 (0.338–0.523) | $9.39 \times 10^{-14}$ | `mutation_resid_DNMT3A` cellstate__residual__raw; cell-state__residual | `DNMT3A_genotype` apparent__positive__unlabeled |
| *IDH2* | 2 | 228 | 0.329 (0.197–0.469) | 0.439 | -0.342 (-0.605—0.061) | 0.585 | `mutation_resid_IDH2` cellstate__residual__raw; cell-state__residual | `IDH2_genotype` apparent__positive__unlabeled |
| *TET2* | 18 | 30 | 0.454 (0.276–0.632) | 0.602 | -0.093 (-0.448–0.263) | 0.722 | `mutation_resid_TET2` cellstate__residual__raw; cell-state__residual | `TET2_genotype` apparent__positive__unlabeled |
| *RUNX1* | 23 | 25 | 0.485 (0.322–0.649) | 0.869 | -0.030 (-0.357–0.297) | 0.948 | `mutation_resid_RUNX1` cellstate__residual__raw; cell-state__residual | `RUNX1_genotype` apparent__positive__unlabeled |
| *NRAS* | 36 | 114 | 0.601 (0.500–0.700) | 0.067 | 0.203 (-0.000–0.399) | 0.162 | `mutation_resid_NRAS` cellstate__residual__raw; cell-state__residual | `NRAS_genotype` apparent__positive__unlabeled |
| *KRAS* | 2 | 41 | 0.317 (0.000–0.683) | 0.434 | -0.366 (-1.000–0.366) | 0.585 | `mutation_resid_KRAS` cellstate__residual__raw; cell-state__residual | `KRAS_genotype` apparent__positive__unlabeled |
| *RAD21* | 9 | 187 | 0.708 (0.549–0.850) | 0.035 | 0.417 (0.098–0.699) | 0.106 | `mutation_resid_RAD21` cellstate__residual__raw; cell-state__residual | `RAD21_genotype` apparent__positive__unlabeled |
| *SMC3* | 7 | 19 | 0.684 (0.451–0.880) | 0.169 | 0.368 (-0.098–0.759) | 0.338 | `mutation_resid_SMC3` cellstate__residual__raw; cell-state__residual | `SMC3_genotype` apparent__positive__unlabeled |
| *TP53* | 51 | 25 | 0.743 (0.629–0.859) | $6.35 \times 10^{-4}$ | 0.485 (0.258–0.718) | 0.003 | `mutation_resid_TP53` cellstate__residual__raw; cell-state__residual | `TP53_genotype` apparent__positive__unlabeled |
| *KIT* | 23 | 36 | 0.502 (0.353–0.655) | 0.981 | 0.005 (-0.295–0.309) | 0.981 | `mutation_resid_KIT` cellstate__residual__raw; cell-state__residual | `KIT_genotype` apparent__positive__unlabeled |

**Supplementary Table S4. Patient-level genotype bridge.** Patient-level genotype bridge using median cell-state-residual scores. Mutant and comparison patient counts, group medians, Mann–Whitney tests, Cliff's $\delta$ intervals, FDR, and the exact score fields are shown. FDR was controlled within each cohort's evaluable driver family.

| Cancer | Cohort | Driver | $n_{\text{mut}}$ | $n_{\text{comp}}$ | Median mut / comp | Cliffś $\delta$ (95% CI) | MW $p$ | FDR $q$ | Score column / kind / mode |
|---|---|---|---|---|---|---|---|---|---|
| AML | van Galen | *FLT3* | 2 | 5 | 0.0022 / -0.0012 | 0.400 (-0.600–1.000) | 0.571 | 0.714 | `mutation_resid_FLT3` cellstate_residual_raw; cellstate_residual |
| AML | van Galen | *NPM1* | 5 | 5 | 0.0050 / -0.0000 | 1.000 (1.000–1.000) | 0.008 | 0.040 | `mutation_resid_NPM1` cellstate_residual_raw; cellstate_residual |
| AML | van Galen | *DNMT3A* | 6 | 6 | 0.0160 / 0.0016 | 0.778 (0.333–1.000) | 0.026 | 0.060 | `mutation_resid_DNMT3A` cellstate_residual_raw; cellstate_residual |
| AML | van Galen | *NRAS* | 3 | 5 | 0.0099 / -0.0003 | 1.000 (1.000–1.000) | 0.036 | 0.060 | `mutation_resid_NRAS` cellstate_residual_raw; cellstate_residual |
| AML | van Galen | *TP53* | 3 | 5 | 0.0010 / 0.0018 | -0.067 (-1.000–1.000) | 1.000 | 1.000 | `mutation_resid_TP53` cellstate_residual_raw; cellstate_residual |
| CRC | Lee | *APC* | 17 | 16 | 0.0085 / 0.0013 | 0.404 (0.007–0.772) | 0.050 | 0.124 | `mutation_resid_APC` cellstate_residual_raw; cellstate_residual |
| CRC | Lee | *KRAS* | 10 | 23 | 0.0079 / 0.0059 | 0.087 (-0.357–0.513) | 0.710 | 0.710 | `mutation_resid_KRAS` cellstate_residual_raw; cellstate_residual |
| CRC | Lee | *BRAF* | 4 | 29 | -0.0110 / -0.0052 | -0.293 (-0.845–0.311) | 0.377 | 0.472 | `mutation_resid_BRAF` cellstate_residual_raw; cellstate_residual |
| CRC | Lee | *SMAD4* | 4 | 29 | 0.0410 / 0.0120 | 0.397 (-0.224–0.862) | 0.224 | 0.373 | `mutation_resid_SMAD4` cellstate_residual_raw; cellstate_residual |
| CRC | Lee | *TP53* | 14 | 19 | 0.0072 / -0.0057 | 0.571 (0.180–0.887) | 0.006 | 0.030 | `mutation_resid_TP53` cellstate_residual_raw; cellstate_residual |
| BRCA | Wu | *ERBB2* | 5 | 21 | -0.0013 / -0.0027 | 0.200 (-0.371–0.714) | 0.527 | 0.527 | `mutation_resid_ERBB2` cellstate_residual_raw; cellstate_residual |

**Supplementary Table S5. scOPE–CNV concordance and discordance.** Continuous and thresholded concordance between scOPE and inferred CNV burden. Each agreement category reports the cell count followed by mean scOPE score and mean inferred-CNV burden. CN denotes concordant normal and CM denotes concordant malignant.

| Cancer | Cohort | Cells | Spearman $\rho$ ($p$) | ARI | CN: $n$; mean scOPE / CNV | CNV-only: $n$; mean scOPE / CNV | scOPE-only: $n$; mean scOPE / CNV | CM: $n$; mean scOPE / CNV |
|---|---|---|---|---|---|---|---|---|
| AML | AML_vanGalen | 35,555 | 0.559 (0) | 0.287 | 21,682; 0.368 / 0.006 | 5,035; 0.374 / 0.034 | 2,796; 0.403 / 0.010 | 6,042; 0.420 / 0.046 |
| BRCA | BRCA_Wu | 39,998 | 0.761 (0) | 0.477 | 25,164; 0.297 / 0.008 | 4,875; 0.323 / 0.032 | 1,122; 0.403 / 0.017 | 8,837; 0.478 / 0.042 |
| CRC | CRC_Lee | 40,000 | 0.065 ($8.04 \times 10^{-39}$) | 0.017 | 22,996; 0.586 / 0.015 | 1,207; 0.580 / 0.030 | 14,332; 0.662 / 0.016 | 1,465; 0.688 / 0.031 |
| GBM | GBM_Neftel | 24,131 | 0.787 (0) | 0.079 | 20,095; 0.376 / 0.008 | 572; 0.412 / 0.030 | 3,135; 0.442 / 0.018 | 329; 0.445 / 0.031 |
| LUAD | LUAD_Lambrechts | 40,000 | 0.321 (0) | -0.006 | 17,107; 0.413 / 0.014 | 934; 0.397 / 0.033 | 19,068; 0.469 / 0.016 | 2,891; 0.538 / 0.035 |
| PAAD | PAAD_Moncada | 3,659 | 0.598 (0) | -0.075 | 658; 0.683 / 0.009 | 16; 0.670 / 0.036 | 2,616; 0.770 / 0.014 | 369; 0.850 / 0.043 |
| SKCM | SKCM_JerbyArnon | 7,186 | 0.268 ($2.56 \times 10^{-118}$) | 0.261 | 5,802; 0.491 / 0.018 | 174; 0.497 / 0.044 | 877; 0.638 / 0.024 | 333; 0.712 / 0.042 |

**Supplementary Table S6. Near-diploid patient-level validation.** Patient-level driver comparisons after restricting to patients with inferred-CNV malignant fraction below 0.10. Counts refer to near-diploid mutant and comparison patients; effect sizes, intervals, Mann–Whitney tests, and cohort-family FDR are shown.

| **Cancer** | **Cohort** | **Driver** | $n_{\mathrm{mut}}$ | $n_{\mathrm{comp}}$ | **Cliff's** $\delta$ **(95% CI)** | **MW** $p$ | **FDR** $q$ |
|---|---|---|---|---|---|---|---|
| AML | `AML_vanGalen` | *NPM1* | 2 | 3 | 1.000 (1.000–1.000) | 0.200 | 0.200 |
| AML | `AML_vanGalen` | *DNMT3A* | 2 | 4 | 1.000 (1.000–1.000) | 0.133 | 0.200 |
| AML | `AML_vanGalen` | *NRAS* | 2 | 3 | 1.000 (1.000–1.000) | 0.200 | 0.200 |
| BRCA | `BRCA_Wu` | *ERBB2* | 2 | 13 | -0.154 (-0.692–0.385) | 0.800 | 0.800 |
| CRC | `CRC_Lee` | *APC* | 14 | 11 | 0.623 (0.181–1.000) | 0.009 | 0.019 |
| CRC | `CRC_Lee` | *KRAS* | 8 | 17 | 0.176 (-0.294–0.662) | 0.511 | 0.511 |
| CRC | `CRC_Lee` | *SMAD4* | 3 | 22 | 0.303 (-0.455–0.909) | 0.446 | 0.511 |
| CRC | `CRC_Lee` | *TP53* | 12 | 13 | 0.654 (0.231–1.000) | 0.006 | 0.019 |

**Supplementary Table S7. AML longitudinal driver trends.** Complete AML patient-by-driver longitudinal summary. First and last effect values, change and direction, Spearman trend statistics, trend FDR, driver confidence, direct-label support metrics, and truth-evaluable cell counts are shown. Blank trend fields indicate that the available number of time points did not support the corresponding statistic.

| Patient | Driver | TP | First / last | Change / direction | $\rho$ / $p$ | Trend $q$ | Confidence | Truth $\delta$ / AUROC | Truth $p$ / $q$ | Truth cells mut / WT | Metric |
|---|---|---|---|---|---|---|---|---|---|---|---|
| AML1012 | *ASXL1* | 1 | -0.388 / -0.388 | 0.000; increasing/flat | – / – | – | 0.209 | – / – | – / – | – / – | cliffs_delta |
| AML1012 | *BCOR* | 1 | -0.079 / -0.079 | 0.000; increasing/flat | – / – | – | 0.177 | – / – | – / – | – / – | cliffs_delta |
| AML1012 | *BCORL1* | 1 | -0.316 / -0.316 | 0.000; increasing/flat | – / – | – | 0.281 | – / – | – / – | – / – | cliffs_delta |
| AML1012 | *CBL* | 1 | 0.048 / 0.048 | 0.000; increasing/flat | – / – | – | 0.163 | – / – | – / – | – / – | cliffs_delta |
| AML1012 | *CEBPA* | 1 | 0.306 / 0.306 | 0.000; increasing/flat | – / – | – | 0.370 | – / – | – / – | – / – | cliffs_delta |
| AML1012 | *DNMT3A* | 1 | -0.250 / -0.250 | 0.000; increasing/flat | – / – | – | 0.298 | 0.429 / 0.714 | $1.56 \times 10^{-14}$ / $9.39 \times 10^{-14}$ | 146 / 404 | cliffs_delta |
| AML1012 | *EZH2* | 1 | 0.566 / 0.566 | 0.000; increasing/flat | – / – | – | 0.130 | – / – | – / – | – / – | cliffs_delta |
| AML1012 | *FLT3* | 1 | -0.256 / -0.256 | 0.000; increasing/flat | – / – | – | 0.207 | 0.116 / 0.558 | 0.248 / 0.425 | 46 / 124 | cliffs_delta |
| AML1012 | *GATA2* | 1 | 0.443 / 0.443 | 0.000; increasing/flat | – / – | – | 0.244 | – / – | – / – | – / – | cliffs_delta |
| AML1012 | *IDH1* | 1 | -0.175 / -0.175 | 0.000; increasing/flat | – / – | – | 0.449 | – / – | – / – | – / – | cliffs_delta |
| AML1012 | *IDH2* | 1 | -0.726 / -0.726 | 0.000; increasing/flat | – / – | – | 0.284 | -0.342 / 0.329 | 0.439 / 0.585 | 2 / 228 | cliffs_delta |
| AML1012 | *KIT* | 1 | 0.106 / 0.106 | 0.000; increasing/flat | – / – | – | 0.471 | 0.005 / 0.502 | 0.981 / 0.981 | 23 / 36 | cliffs_delta |
| AML1012 | *KRAS* | 1 | -0.008 / -0.008 | 0.000; increasing/flat | – / – | – | 0.240 | -0.366 / 0.317 | 0.434 / 0.585 | 2 / 41 | cliffs_delta |
| AML1012 | *NPM1* | 1 | -0.362 / -0.362 | 0.000; increasing/flat | – / – | – | 0.318 | 0.506 / 0.753 | $6.11 \times 10^{-68}$ / $7.34 \times 10^{-67}$ | 542 / 1,457 | cliffs_delta |
| AML1012 | *NRAS* | 1 | 0.519 / 0.519 | 0.000; increasing/flat | – / – | – | 0.199 | 0.203 / 0.601 | 0.067 / 0.162 | 36 / 114 | cliffs_delta |
| AML1012 | *PHF6* | 1 | 0.164 / 0.164 | 0.000; increasing/flat | – / – | – | 0.251 | – / – | – / – | – / – | cliffs_delta |
| AML1012 | *PTPN11* | 1 | -0.228 / -0.228 | 0.000; increasing/flat | – / – | – | 0.212 | – / – | – / – | – / – | cliffs_delta |
| AML1012 | *RAD21* | 1 | -0.178 / -0.178 | 0.000; increasing/flat | – / – | – | 0.237 | 0.417 / 0.708 | 0.035 / 0.106 | 9 / 187 | cliffs_delta |
| AML1012 | *RUNX1* | 1 | -0.203 / -0.203 | 0.000; increasing/flat | – / – | – | 0.256 | -0.030 / 0.485 | 0.869 / 0.948 | 23 / 25 | cliffs_delta |
| AML1012 | *SF3B1* | 1 | -0.054 / -0.054 | 0.000; increasing/flat | – / – | – | 0.264 | – / – | – / – | – / – | cliffs_delta |
| AML1012 | *SMC1A* | 1 | -0.106 / -0.106 | 0.000; increasing/flat | – / – | – | 0.312 | – / – | – / – | – / – | cliffs_delta |
| AML1012 | *SMC3* | 1 | -0.419 / -0.419 | 0.000; increasing/flat | – / – | – | 0.311 | 0.368 / 0.684 | 0.169 / 0.338 | 7 / 19 | cliffs_delta |
| AML1012 | *SRSF2* | 1 | -0.812 / -0.812 | 0.000; increasing/flat | – / – | – | 0.341 | – / – | – / – | – / – | cliffs_delta |
| AML1012 | *STAG2* | 1 | -0.591 / -0.591 | 0.000; increasing/flat | – / – | – | 0.235 | – / – | – / – | – / – | cliffs_delta |
| AML1012 | *TET2* | 1 | -0.157 / -0.157 | 0.000; increasing/flat | – / – | – | 0.214 | -0.093 / 0.454 | 0.602 / 0.722 | 18 / 30 | cliffs_delta |
| AML1012 | *TP53* | 1 | 0.204 / 0.204 | 0.000; increasing/flat | – / – | – | 0.395 | 0.485 / 0.743 | $6.35 \times 10^{-4}$ / 0.003 | 51 / 25 | cliffs_delta |
| AML1012 | *U2AF1* | 1 | -0.303 / -0.303 | 0.000; increasing/flat | – / – | – | 0.166 | – / – | – / – | – / – | cliffs_delta |
| AML1012 | *WT1* | 1 | -0.202 / -0.202 | 0.000; increasing/flat | – / – | – | 0.244 | – / – | – / – | – / – | cliffs_delta |
| AML1012 | *ZRSR2* | 1 | -0.214 / -0.214 | 0.000; increasing/flat | – / – | – | 0.219 | – / – | – / – | – / – | cliffs_delta |
| AML210A | *ASXL1* | 1 | 0.129 / 0.129 | 0.000; increasing/flat | – / – | – | 0.209 | – / – | – / – | – / – | cliffs_delta |
| AML210A | *BCOR* | 1 | 0.184 / 0.184 | 0.000; increasing/flat | – / – | – | 0.177 | – / – | – / – | – / – | cliffs_delta |
| AML210A | *BCORL1* | 1 | -0.183 / -0.183 | 0.000; increasing/flat | – / – | – | 0.281 | – / – | – / – | – / – | cliffs_delta |
| AML210A | *CBL* | 1 | 0.066 / 0.066 | 0.000; increasing/flat | – / – | – | 0.163 | – / – | – / – | – / – | cliffs_delta |
| AML210A | *CEBPA* | 1 | 0.278 / 0.278 | 0.000; increasing/flat | – / – | – | 0.370 | – / – | – / – | – / – | cliffs_delta |
| AML210A | *DNMT3A* | 1 | 0.544 / 0.544 | 0.000; increasing/flat | – / – | – | 0.298 | 0.429 / 0.714 | $1.56 \times 10^{-14}$ / $9.39 \times 10^{-14}$ | 146 / 404 | cliffs_delta |
| AML210A | *EZH2* | 1 | -0.049 / -0.049 | 0.000; increasing/flat | – / – | – | 0.130 | – / – | – / – | – / – | cliffs_delta |
| AML210A | *FLT3* | 1 | 0.096 / 0.096 | 0.000; increasing/flat | – / – | – | 0.207 | 0.116 / 0.558 | 0.248 / 0.425 | 46 / 124 | cliffs_delta |
| AML210A | *GATA2* | 1 | 0.246 / 0.246 | 0.000; increasing/flat | – / – | – | 0.244 | – / – | – / – | – / – | cliffs_delta |
| AML210A | *IDH1* | 1 | 0.012 / 0.012 | 0.000; increasing/flat | – / – | – | 0.449 | – / – | – / – | – / – | cliffs_delta |
| AML210A | *IDH2* | 1 | -0.114 / -0.114 | 0.000; increasing/flat | – / – | – | 0.284 | -0.342 / 0.329 | 0.439 / 0.585 | 2 / 228 | cliffs_delta |
| AML210A | *KIT* | 1 | 0.123 / 0.123 | 0.000; increasing/flat | – / – | – | 0.471 | 0.005 / 0.502 | 0.981 / 0.981 | 23 / 36 | cliffs_delta |
| AML210A | *KRAS* | 1 | -0.057 / -0.057 | 0.000; increasing/flat | – / – | – | 0.240 | -0.366 / 0.317 | 0.434 / 0.585 | 2 / 41 | cliffs_delta |
| AML210A | *NPM1* | 1 | 0.523 / 0.523 | 0.000; increasing/flat | – / – | – | 0.318 | 0.506 / 0.753 | $6.11 \times 10^{-68}$ / $7.34 \times 10^{-67}$ | 542 / 1,457 | cliffs_delta |
| AML210A | *NRAS* | 1 | 0.156 / 0.156 | 0.000; increasing/flat | – / – | – | 0.199 | 0.203 / 0.601 | 0.067 / 0.162 | 36 / 114 | cliffs_delta |
| AML210A | *PHF6* | 1 | 0.298 / 0.298 | 0.000; increasing/flat | – / – | – | 0.251 | – / – | – / – | – / – | cliffs_delta |
| AML210A | *PTPN11* | 1 | 0.382 / 0.382 | 0.000; increasing/flat | – / – | – | 0.212 | – / – | – / – | – / – | cliffs_delta |
| AML210A | *RAD21* | 1 | 0.011 / 0.011 | 0.000; increasing/flat | – / – | – | 0.237 | 0.417 / 0.708 | 0.035 / 0.106 | 9 / 187 | cliffs_delta |
| AML210A | *RUNX1* | 1 | 0.082 / 0.082 | 0.000; increasing/flat | – / – | – | 0.256 | -0.030 / 0.485 | 0.869 / 0.948 | 23 / 25 | cliffs_delta |
| AML210A | *SF3B1* | 1 | 0.035 / 0.035 | 0.000; increasing/flat | – / – | – | 0.264 | – / – | – / – | – / – | cliffs_delta |
| AML210A | *SMC1A* | 1 | 0.176 / 0.176 | 0.000; increasing/flat | – / – | – | 0.312 | – / – | – / – | – / – | cliffs_delta |
| AML210A | *SMC3* | 1 | 0.178 / 0.178 | 0.000; increasing/flat | – / – | – | 0.311 | 0.368 / 0.684 | 0.169 / 0.338 | 7 / 19 | cliffs_delta |

Continued on next page

*Supplementary Table S7 continued*

| Patient | Driver | TP | First / last | Change / direction | $\rho$ / $p$ | Trend $q$ | Confidence | Truth $\delta$ / AUROC | Truth $p$ / $q$ | Truth cells mut / WT | Metric |
|---|---|---|---|---|---|---|---|---|---|---|---|
| AML210A | *SRSF2* | 1 | -0.146 / -0.146 | 0.000; increasing/flat | – / – | – | 0.341 | – / – | – / – | – / – | cliffs_delta |
| AML210A | *STAG2* | 1 | 0.118 / 0.118 | 0.000; increasing/flat | – / – | – | 0.235 | – / – | – / – | – / – | cliffs_delta |
| AML210A | *TET2* | 1 | 0.356 / 0.356 | 0.000; increasing/flat | – / – | – | 0.214 | -0.093 / 0.454 | 0.602 / 0.722 | 18 / 30 | cliffs_delta |
| AML210A | *TP53* | 1 | -0.054 / -0.054 | 0.000; increasing/flat | – / – | – | 0.395 | 0.485 / 0.743 | $6.35 \times 10^{-4}$ / 0.003 | 51 / 25 | cliffs_delta |
| AML210A | *U2AF1* | 1 | -0.074 / -0.074 | 0.000; increasing/flat | – / – | – | 0.166 | – / – | – / – | – / – | cliffs_delta |
| AML210A | *WT1* | 1 | 0.341 / 0.341 | 0.000; increasing/flat | – / – | – | 0.244 | – / – | – / – | – / – | cliffs_delta |
| AML210A | *ZRSR2* | 1 | -0.125 / -0.125 | 0.000; increasing/flat | – / – | – | 0.219 | – / – | – / – | – / – | cliffs_delta |
| AML328 | *ASXL1* | 4 | 0.161 / 0.163 | 0.002; increasing/flat | 0.600 / 0.400 | 0.690 | 0.209 | – / – | – / – | – / – | cliffs_delta |
| AML328 | *BCOR* | 4 | 0.447 / 0.176 | -0.271; decreasing (response-consistent) | -0.800 / 0.200 | 0.569 | 0.177 | – / – | – / – | – / – | cliffs_delta |
| AML328 | *BCORL1* | 4 | -0.047 / -0.071 | -0.024; decreasing (response-consistent) | -0.400 / 0.600 | 0.690 | 0.281 | – / – | – / – | – / – | cliffs_delta |
| AML328 | *CBL* | 4 | 0.089 / 0.089 | -0.000; decreasing (response-consistent) | 0.000 / 1.000 | 1.000 | 0.163 | – / – | – / – | – / – | cliffs_delta |
| AML328 | *CEBPA* | 4 | 0.299 / 0.169 | -0.130; decreasing (response-consistent) | -0.600 / 0.400 | 0.690 | 0.370 | – / – | – / – | – / – | cliffs_delta |
| AML328 | *DNMT3A* | 4 | 0.499 / 0.060 | -0.438; decreasing (response-consistent) | -0.800 / 0.200 | 0.569 | 0.298 | 0.429 / 0.714 | $1.56 \times 10^{-14}$ / $9.39 \times 10^{-14}$ | 146 / 404 | cliffs_delta |
| AML328 | *EZH2* | 4 | 0.220 / 0.123 | -0.097; decreasing (response-consistent) | -0.400 / 0.600 | 0.690 | 0.130 | – / – | – / – | – / – | cliffs_delta |
| AML328 | *FLT3* | 4 | -0.622 / -0.107 | 0.515; increasing/flat | 0.600 / 0.400 | 0.690 | 0.207 | 0.116 / 0.558 | 0.248 / 0.425 | 46 / 124 | cliffs_delta |
| AML328 | *GATA2* | 4 | 0.438 / 0.187 | -0.252; decreasing (response-consistent) | -0.800 / 0.200 | 0.569 | 0.244 | – / – | – / – | – / – | cliffs_delta |
| AML328 | *IDH1* | 4 | 0.240 / 0.143 | -0.098; decreasing (response-consistent) | -0.800 / 0.200 | 0.569 | 0.449 | – / – | – / – | – / – | cliffs_delta |
| AML328 | *IDH2* | 4 | -0.141 / -0.034 | 0.107; increasing/flat | 0.600 / 0.400 | 0.690 | 0.284 | -0.342 / 0.329 | 0.439 / 0.585 | 2 / 228 | cliffs_delta |
| AML328 | *KIT* | 4 | 0.101 / 0.090 | -0.011; decreasing (response-consistent) | -0.600 / 0.400 | 0.690 | 0.471 | 0.005 / 0.502 | 0.981 / 0.981 | 23 / 36 | cliffs_delta |
| AML328 | *KRAS* | 4 | 0.137 / 0.205 | 0.068; increasing/flat | 0.800 / 0.200 | 0.569 | 0.240 | -0.366 / 0.317 | 0.434 / 0.585 | 2 / 41 | cliffs_delta |
| AML328 | *NPM1* | 4 | -0.127 / 0.062 | 0.188; increasing/flat | 0.600 / 0.400 | 0.690 | 0.318 | 0.506 / 0.753 | $6.11 \times 10^{-68}$ / $7.34 \times 10^{-67}$ | 542 / 1,457 | cliffs_delta |
| AML328 | *NRAS* | 4 | 0.119 / 0.217 | 0.098; increasing/flat | 0.600 / 0.400 | 0.690 | 0.199 | 0.203 / 0.601 | 0.067 / 0.162 | 36 / 114 | cliffs_delta |
| AML328 | *PHF6* | 4 | 0.393 / 0.145 | -0.249; decreasing (response-consistent) | -0.800 / 0.200 | 0.569 | 0.251 | – / – | – / – | – / – | cliffs_delta |
| AML328 | *PTPN11* | 4 | 0.043 / 0.053 | 0.010; increasing/flat | 0.400 / 0.600 | 0.690 | 0.212 | – / – | – / – | – / – | cliffs_delta |
| AML328 | *RAD21* | 4 | -0.390 / 0.100 | 0.489; increasing/flat | 0.600 / 0.400 | 0.690 | 0.237 | 0.417 / 0.708 | 0.035 / 0.106 | 9 / 187 | cliffs_delta |
| AML328 | *RUNX1* | 4 | 0.598 / 0.271 | -0.327; decreasing (response-consistent) | -1.000 / 0 | 0 | 0.256 | -0.030 / 0.485 | 0.869 / 0.948 | 23 / 25 | cliffs_delta |
| AML328 | *SF3B1* | 4 | 0.152 / 0.251 | 0.099; increasing/flat | 0.600 / 0.400 | 0.690 | 0.264 | – / – | – / – | – / – | cliffs_delta |
| AML328 | *SMC1A* | 4 | 0.173 / 0.153 | -0.020; decreasing (response-consistent) | -0.400 / 0.600 | 0.690 | 0.312 | – / – | – / – | – / – | cliffs_delta |
| AML328 | *SMC3* | 4 | -0.227 / 0.061 | 0.288; increasing/flat | 1.000 / 0 | 0 | 0.311 | 0.368 / 0.684 | 0.169 / 0.338 | 7 / 19 | cliffs_delta |
| AML328 | *SRSF2* | 4 | 0.124 / 0.132 | 0.009; increasing/flat | 0.600 / 0.400 | 0.690 | 0.341 | – / – | – / – | – / – | cliffs_delta |
| AML328 | *STAG2* | 4 | -0.088 / 0.151 | 0.239; increasing/flat | 0.800 / 0.200 | 0.569 | 0.235 | – / – | – / – | – / – | cliffs_delta |
| AML328 | *TET2* | 4 | 0.266 / 0.059 | -0.207; decreasing (response-consistent) | -0.800 / 0.200 | 0.569 | 0.214 | -0.093 / 0.454 | 0.602 / 0.722 | 18 / 30 | cliffs_delta |
| AML328 | *TP53* | 4 | 0.210 / 0.041 | -0.169; decreasing (response-consistent) | -0.600 / 0.400 | 0.690 | 0.395 | 0.485 / 0.743 | $6.35 \times 10^{-4}$ / 0.003 | 51 / 25 | cliffs_delta |
| AML328 | *U2AF1* | 4 | -0.011 / -0.279 | -0.268; decreasing (response-consistent) | -0.800 / 0.200 | 0.569 | 0.166 | – / – | – / – | – / – | cliffs_delta |
| AML328 | *WT1* | 4 | 0.174 / 0.202 | 0.029; increasing/flat | 0.400 / 0.600 | 0.690 | 0.244 | – / – | – / – | – / – | cliffs_delta |
| AML328 | *ZRSR2* | 4 | -0.195 / -0.033 | 0.162; increasing/flat | 0.600 / 0.400 | 0.690 | 0.219 | – / – | – / – | – / – | cliffs_delta |
| AML329 | *ASXL1* | 3 | -0.175 / 0.141 | 0.315; increasing/flat | 0.500 / 0.667 | 0.690 | 0.209 | – / – | – / – | – / – | cliffs_delta |
| AML329 | *BCOR* | 3 | 0.049 / 0.350 | 0.301; increasing/flat | 1.000 / 0 | 0 | 0.177 | – / – | – / – | – / – | cliffs_delta |
| AML329 | *BCORL1* | 3 | -0.246 / -0.142 | 0.104; increasing/flat | 0.500 / 0.667 | 0.690 | 0.281 | – / – | – / – | – / – | cliffs_delta |
| AML329 | *CBL* | 3 | 0.209 / 0.245 | 0.036; increasing/flat | 0.500 / 0.667 | 0.690 | 0.163 | – / – | – / – | – / – | cliffs_delta |
| AML329 | *CEBPA* | 3 | 0.226 / -0.110 | -0.335; decreasing (response-consistent) | -0.500 / 0.667 | 0.690 | 0.370 | – / – | – / – | – / – | cliffs_delta |
| AML329 | *DNMT3A* | 3 | 0.525 / 0.150 | -0.375; decreasing (response-consistent) | -0.500 / 0.667 | 0.690 | 0.298 | 0.429 / 0.714 | $1.56 \times 10^{-14}$ / $9.39 \times 10^{-14}$ | 146 / 404 | cliffs_delta |
| AML329 | *EZH2* | 3 | -0.159 / 0.419 | 0.578; increasing/flat | 1.000 / 0 | 0 | 0.130 | – / – | – / – | – / – | cliffs_delta |

Continued on next page

*Supplementary Table S7 continued*

| Patient | Driver | TP | First / last | Change / direction | $\rho$ / $p$ | Trend $q$ | Confidence | Truth $\delta$ / AUROC | Truth $p$ / $q$ | Truth cells mut / WT | Metric |
|---|---|---|---|---|---|---|---|---|---|---|---|
| AML329 | *FLT3* | 3 | 0.218 / -0.046 | -0.265; decreasing (response-consistent) | -0.500 / 0.667 | 0.690 | 0.207 | 0.116 / 0.558 | 0.248 / 0.425 | 46 / 124 | cliffs_delta |
| AML329 | *GATA2* | 3 | 0.211 / 0.338 | 0.127; increasing/flat | 0.500 / 0.667 | 0.690 | 0.244 | – / – | – / – | – / – | cliffs_delta |
| AML329 | *IDH1* | 3 | 0.171 / 0.056 | -0.115; decreasing (response-consistent) | -1.000 / 0 | 0 | 0.449 | – / – | – / – | – / – | cliffs_delta |
| AML329 | *IDH2* | 3 | 0.058 / -0.104 | -0.162; decreasing (response-consistent) | -1.000 / 0 | 0 | 0.284 | -0.342 / 0.329 | 0.439 / 0.585 | 2 / 228 | cliffs_delta |
| AML329 | *KIT* | 3 | 0.177 / -0.037 | -0.214; decreasing (response-consistent) | -0.500 / 0.667 | 0.690 | 0.471 | 0.005 / 0.502 | 0.981 / 0.981 | 23 / 36 | cliffs_delta |
| AML329 | *KRAS* | 3 | -0.046 / -0.056 | -0.010; decreasing (response-consistent) | -0.500 / 0.667 | 0.690 | 0.240 | -0.366 / 0.317 | 0.434 / 0.585 | 2 / 41 | cliffs_delta |
| AML329 | *NPM1* | 3 | 0.551 / 0.180 | -0.371; decreasing (response-consistent) | -0.500 / 0.667 | 0.690 | 0.318 | 0.506 / 0.753 | $6.11 \times 10^{-68}$ / $7.34 \times 10^{-67}$ | 542 / 1,457 | cliffs_delta |
| AML329 | *NRAS* | 3 | 0.111 / 0.274 | 0.163; increasing/flat | 0.500 / 0.667 | 0.690 | 0.199 | 0.203 / 0.601 | 0.067 / 0.162 | 36 / 114 | cliffs_delta |
| AML329 | *PHF6* | 3 | -0.024 / 0.191 | 0.215; increasing/flat | 0.500 / 0.667 | 0.690 | 0.251 | – / – | – / – | – / – | cliffs_delta |
| AML329 | *PTPN11* | 3 | 0.315 / 0.179 | -0.136; decreasing (response-consistent) | -0.500 / 0.667 | 0.690 | 0.212 | – / – | – / – | – / – | cliffs_delta |
| AML329 | *RAD21* | 3 | 0.053 / -0.174 | -0.227; decreasing (response-consistent) | -0.500 / 0.667 | 0.690 | 0.237 | 0.417 / 0.708 | 0.035 / 0.106 | 9 / 187 | cliffs_delta |
| AML329 | *RUNX1* | 3 | -0.161 / 0.327 | 0.488; increasing/flat | 1.000 / 0 | 0 | 0.256 | -0.030 / 0.485 | 0.869 / 0.948 | 23 / 25 | cliffs_delta |
| AML329 | *SF3B1* | 3 | 0.035 / 0.171 | 0.136; increasing/flat | 1.000 / 0 | 0 | 0.264 | – / – | – / – | – / – | cliffs_delta |
| AML329 | *SMC1A* | 3 | 0.225 / 0.227 | 0.002; increasing/flat | 0.500 / 0.667 | 0.690 | 0.312 | – / – | – / – | – / – | cliffs_delta |
| AML329 | *SMC3* | 3 | 0.247 / -0.210 | -0.457; decreasing (response-consistent) | -1.000 / 0 | 0 | 0.311 | 0.368 / 0.684 | 0.169 / 0.338 | 7 / 19 | cliffs_delta |
| AML329 | *SRSF2* | 3 | -0.257 / -0.051 | 0.206; increasing/flat | 1.000 / 0 | 0 | 0.341 | – / – | – / – | – / – | cliffs_delta |
| AML329 | *STAG2* | 3 | 0.176 / 0.120 | -0.056; decreasing (response-consistent) | -1.000 / 0 | 0 | 0.235 | – / – | – / – | – / – | cliffs_delta |
| AML329 | *TET2* | 3 | 0.004 / 0.122 | 0.118; increasing/flat | 0.500 / 0.667 | 0.690 | 0.214 | -0.093 / 0.454 | 0.602 / 0.722 | 18 / 30 | cliffs_delta |
| AML329 | *TP53* | 3 | -0.177 / -0.032 | 0.145; increasing/flat | 0.500 / 0.667 | 0.690 | 0.395 | 0.485 / 0.743 | $6.35 \times 10^{-4}$ / 0.003 | 51 / 25 | cliffs_delta |
| AML329 | *U2AF1* | 3 | 0.017 / 0.041 | 0.024; increasing/flat | 0.500 / 0.667 | 0.690 | 0.166 | – / – | – / – | – / – | cliffs_delta |
| AML329 | *WT1* | 3 | 0.419 / 0.311 | -0.108; decreasing (response-consistent) | -0.500 / 0.667 | 0.690 | 0.244 | – / – | – / – | – / – | cliffs_delta |
| AML329 | *ZRSR2* | 3 | -0.152 / 0.170 | 0.322; increasing/flat | 1.000 / 0 | 0 | 0.219 | – / – | – / – | – / – | cliffs_delta |
| AML371 | *ASXL1* | 2 | -0.203 / 0.244 | 0.448; increasing/flat | – / – | – | 0.209 | – / – | – / – | – / – | cliffs_delta |
| AML371 | *BCOR* | 2 | 0.316 / 0.497 | 0.181; increasing/flat | – / – | – | 0.177 | – / – | – / – | – / – | cliffs_delta |
| AML371 | *BCORL1* | 2 | -0.404 / -0.345 | 0.060; increasing/flat | – / – | – | 0.281 | – / – | – / – | – / – | cliffs_delta |
| AML371 | *CBL* | 2 | -0.048 / -0.117 | -0.069; decreasing (response-consistent) | – / – | – | 0.163 | – / – | – / – | – / – | cliffs_delta |
| AML371 | *CEBPA* | 2 | 0.198 / -0.256 | -0.454; decreasing (response-consistent) | – / – | – | 0.370 | – / – | – / – | – / – | cliffs_delta |
| AML371 | *DNMT3A* | 2 | 0.128 / 0.009 | -0.120; decreasing (response-consistent) | – / – | – | 0.298 | 0.429 / 0.714 | $1.56 \times 10^{-14}$ / $9.39 \times 10^{-14}$ | 146 / 404 | cliffs_delta |
| AML371 | *EZH2* | 2 | 0.191 / 0.206 | 0.015; increasing/flat | – / – | – | 0.130 | – / – | – / – | – / – | cliffs_delta |
| AML371 | *FLT3* | 2 | 0.082 / 0.165 | 0.083; increasing/flat | – / – | – | 0.207 | 0.116 / 0.558 | 0.248 / 0.425 | 46 / 124 | cliffs_delta |
| AML371 | *GATA2* | 2 | -0.270 / -0.132 | 0.138; increasing/flat | – / – | – | 0.244 | – / – | – / – | – / – | cliffs_delta |
| AML371 | *IDH1* | 2 | -0.211 / -0.165 | 0.047; increasing/flat | – / – | – | 0.449 | – / – | – / – | – / – | cliffs_delta |
| AML371 | *IDH2* | 2 | -0.707 / -0.326 | 0.381; increasing/flat | – / – | – | 0.284 | -0.342 / 0.329 | 0.439 / 0.585 | 2 / 228 | cliffs_delta |
| AML371 | *KIT* | 2 | 0.299 / 0.095 | -0.205; decreasing (response-consistent) | – / – | – | 0.471 | 0.005 / 0.502 | 0.981 / 0.981 | 23 / 36 | cliffs_delta |
| AML371 | *KRAS* | 2 | -0.468 / -0.525 | -0.057; decreasing (response-consistent) | – / – | – | 0.240 | -0.366 / 0.317 | 0.434 / 0.585 | 2 / 41 | cliffs_delta |
| AML371 | *NPM1* | 2 | 0.072 / 0.157 | 0.084; increasing/flat | – / – | – | 0.318 | 0.506 / 0.753 | $6.11 \times 10^{-68}$ / $7.34 \times 10^{-67}$ | 542 / 1,457 | cliffs_delta |
| AML371 | *NRAS* | 2 | 0.441 / 0.053 | -0.388; decreasing (response-consistent) | – / – | – | 0.199 | 0.203 / 0.601 | 0.067 / 0.162 | 36 / 114 | cliffs_delta |
| AML371 | *PHF6* | 2 | 0.358 / 0.431 | 0.073; increasing/flat | – / – | – | 0.251 | – / – | – / – | – / – | cliffs_delta |
| AML371 | *PTPN11* | 2 | -0.004 / 0.301 | 0.306; increasing/flat | – / – | – | 0.212 | – / – | – / – | – / – | cliffs_delta |
| AML371 | *RAD21* | 2 | -0.086 / -0.471 | -0.385; decreasing (response-consistent) | – / – | – | 0.237 | 0.417 / 0.708 | 0.035 / 0.106 | 9 / 187 | cliffs_delta |
| AML371 | *RUNX1* | 2 | 0.083 / 0.330 | 0.247; increasing/flat | – / – | – | 0.256 | -0.030 / 0.485 | 0.869 / 0.948 | 23 / 25 | cliffs_delta |
| AML371 | *SF3B1* | 2 | -0.088 / -0.056 | 0.031; increasing/flat | – / – | – | 0.264 | – / – | – / – | – / – | cliffs_delta |

Continued on next page

*Supplementary Table S7 continued*

| Patient | Driver | TP | First / last | Change / direction | $\rho$ / $p$ | Trend $q$ | Confidence | Truth $\delta$ / AUROC | Truth $p$ / $q$ | Truth cells mut / WT | Metric |
|---|---|---|---|---|---|---|---|---|---|---|---|
| AML371 | *SMC1A* | 2 | -0.143 / -0.367 | -0.224; decreasing (response-consistent) | – / – | – | 0.312 | – / – | – / – | – / – | cliffs_delta |
| AML371 | *SMC3* | 2 | -0.374 / -0.203 | 0.172; increasing/flat | – / – | – | 0.311 | 0.368 / 0.684 | 0.169 / 0.338 | 7 / 19 | cliffs_delta |
| AML371 | *SRSF2* | 2 | -0.650 / -0.045 | 0.605; increasing/flat | – / – | – | 0.341 | – / – | – / – | – / – | cliffs_delta |
| AML371 | *STAG2* | 2 | -0.359 / 0.139 | 0.497; increasing/flat | – / – | – | 0.235 | – / – | – / – | – / – | cliffs_delta |
| AML371 | *TET2* | 2 | 0.171 / 0.463 | 0.293; increasing/flat | – / – | – | 0.214 | -0.093 / 0.454 | 0.602 / 0.722 | 18 / 30 | cliffs_delta |
| AML371 | *TP53* | 2 | 0.314 / 0.487 | 0.173; increasing/flat | – / – | – | 0.395 | 0.485 / 0.743 | $6.35 \times 10^{-4}$ / 0.003 | 51 / 25 | cliffs_delta |
| AML371 | *U2AF1* | 2 | -0.565 / -0.170 | 0.395; increasing/flat | – / – | – | 0.166 | – / – | – / – | – / – | cliffs_delta |
| AML371 | *WT1* | 2 | -0.086 / 0.163 | 0.248; increasing/flat | – / – | – | 0.244 | – / – | – / – | – / – | cliffs_delta |
| AML371 | *ZRSR2* | 2 | -0.029 / -0.010 | 0.019; increasing/flat | – / – | – | 0.219 | – / – | – / – | – / – | cliffs_delta |
| AML419A | *ASXL1* | 1 | -0.038 / -0.038 | 0.000; increasing/flat | – / – | – | 0.209 | – / – | – / – | – / – | cliffs_delta |
| AML419A | *BCOR* | 1 | 0.052 / 0.052 | 0.000; increasing/flat | – / – | – | 0.177 | – / – | – / – | – / – | cliffs_delta |
| AML419A | *BCORL1* | 1 | -0.290 / -0.290 | 0.000; increasing/flat | – / – | – | 0.281 | – / – | – / – | – / – | cliffs_delta |
| AML419A | *CBL* | 1 | 0.212 / 0.212 | 0.000; increasing/flat | – / – | – | 0.163 | – / – | – / – | – / – | cliffs_delta |
| AML419A | *CEBPA* | 1 | 0.468 / 0.468 | 0.000; increasing/flat | – / – | – | 0.370 | – / – | – / – | – / – | cliffs_delta |
| AML419A | *DNMT3A* | 1 | 0.843 / 0.843 | 0.000; increasing/flat | – / – | – | 0.298 | 0.429 / 0.714 | $1.56 \times 10^{-14}$ / $9.39 \times 10^{-14}$ | 146 / 404 | cliffs_delta |
| AML419A | *EZH2* | 1 | 0.070 / 0.070 | 0.000; increasing/flat | – / – | – | 0.130 | – / – | – / – | – / – | cliffs_delta |
| AML419A | *FLT3* | 1 | -0.137 / -0.137 | 0.000; increasing/flat | – / – | – | 0.207 | 0.116 / 0.558 | 0.248 / 0.425 | 46 / 124 | cliffs_delta |
| AML419A | *GATA2* | 1 | 0.320 / 0.320 | 0.000; increasing/flat | – / – | – | 0.244 | – / – | – / – | – / – | cliffs_delta |
| AML419A | *IDH1* | 1 | 0.215 / 0.215 | 0.000; increasing/flat | – / – | – | 0.449 | – / – | – / – | – / – | cliffs_delta |
| AML419A | *IDH2* | 1 | 0.052 / 0.052 | 0.000; increasing/flat | – / – | – | 0.284 | -0.342 / 0.329 | 0.439 / 0.585 | 2 / 228 | cliffs_delta |
| AML419A | *KIT* | 1 | 0.292 / 0.292 | 0.000; increasing/flat | – / – | – | 0.471 | 0.005 / 0.502 | 0.981 / 0.981 | 23 / 36 | cliffs_delta |
| AML419A | *KRAS* | 1 | -0.255 / -0.255 | 0.000; increasing/flat | – / – | – | 0.240 | -0.366 / 0.317 | 0.434 / 0.585 | 2 / 41 | cliffs_delta |
| AML419A | *NPM1* | 1 | 0.695 / 0.695 | 0.000; increasing/flat | – / – | – | 0.318 | 0.506 / 0.753 | $6.11 \times 10^{-68}$ / $7.34 \times 10^{-67}$ | 542 / 1,457 | cliffs_delta |
| AML419A | *NRAS* | 1 | 0.032 / 0.032 | 0.000; increasing/flat | – / – | – | 0.199 | 0.203 / 0.601 | 0.067 / 0.162 | 36 / 114 | cliffs_delta |
| AML419A | *PHF6* | 1 | 0.274 / 0.274 | 0.000; increasing/flat | – / – | – | 0.251 | – / – | – / – | – / – | cliffs_delta |
| AML419A | *PTPN11* | 1 | 0.351 / 0.351 | 0.000; increasing/flat | – / – | – | 0.212 | – / – | – / – | – / – | cliffs_delta |
| AML419A | *RAD21* | 1 | -0.320 / -0.320 | 0.000; increasing/flat | – / – | – | 0.237 | 0.417 / 0.708 | 0.035 / 0.106 | 9 / 187 | cliffs_delta |
| AML419A | *RUNX1* | 1 | 0.125 / 0.125 | 0.000; increasing/flat | – / – | – | 0.256 | -0.030 / 0.485 | 0.869 / 0.948 | 23 / 25 | cliffs_delta |
| AML419A | *SF3B1* | 1 | -0.346 / -0.346 | 0.000; increasing/flat | – / – | – | 0.264 | – / – | – / – | – / – | cliffs_delta |
| AML419A | *SMC1A* | 1 | 0.369 / 0.369 | 0.000; increasing/flat | – / – | – | 0.312 | – / – | – / – | – / – | cliffs_delta |
| AML419A | *SMC3* | 1 | 0.042 / 0.042 | 0.000; increasing/flat | – / – | – | 0.311 | 0.368 / 0.684 | 0.169 / 0.338 | 7 / 19 | cliffs_delta |
| AML419A | *SRSF2* | 1 | -0.145 / -0.145 | 0.000; increasing/flat | – / – | – | 0.341 | – / – | – / – | – / – | cliffs_delta |
| AML419A | *STAG2* | 1 | 0.163 / 0.163 | 0.000; increasing/flat | – / – | – | 0.235 | – / – | – / – | – / – | cliffs_delta |
| AML419A | *TET2* | 1 | 0.355 / 0.355 | 0.000; increasing/flat | – / – | – | 0.214 | -0.093 / 0.454 | 0.602 / 0.722 | 18 / 30 | cliffs_delta |
| AML419A | *TP53* | 1 | -0.166 / -0.166 | 0.000; increasing/flat | – / – | – | 0.395 | 0.485 / 0.743 | $6.35 \times 10^{-4}$ / 0.003 | 51 / 25 | cliffs_delta |
| AML419A | *U2AF1* | 1 | -0.164 / -0.164 | 0.000; increasing/flat | – / – | – | 0.166 | – / – | – / – | – / – | cliffs_delta |
| AML419A | *WT1* | 1 | 0.317 / 0.317 | 0.000; increasing/flat | – / – | – | 0.244 | – / – | – / – | – / – | cliffs_delta |
| AML419A | *ZRSR2* | 1 | -0.217 / -0.217 | 0.000; increasing/flat | – / – | – | 0.219 | – / – | – / – | – / – | cliffs_delta |
| AML420B | *ASXL1* | 3 | 0.231 / 0.371 | 0.141; increasing/flat | 1.000 / 0 | 0 | 0.209 | – / – | – / – | – / – | cliffs_delta |
| AML420B | *BCOR* | 3 | 0.271 / 0.168 | -0.103; decreasing (response-consistent) | -0.500 / 0.667 | 0.690 | 0.177 | – / – | – / – | – / – | cliffs_delta |
| AML420B | *BCORL1* | 3 | -0.310 / -0.189 | 0.120; increasing/flat | 0.500 / 0.667 | 0.690 | 0.281 | – / – | – / – | – / – | cliffs_delta |
| AML420B | *CBL* | 3 | 0.065 / 0.182 | 0.117; increasing/flat | 0.500 / 0.667 | 0.690 | 0.163 | – / – | – / – | – / – | cliffs_delta |
| AML420B | *CEBPA* | 3 | 0.100 / -0.161 | -0.261; decreasing (response-consistent) | -1.000 / 0 | 0 | 0.370 | – / – | – / – | – / – | cliffs_delta |
| AML420B | *DNMT3A* | 3 | 0.201 / 0.211 | 0.010; increasing/flat | 0.500 / 0.667 | 0.690 | 0.298 | 0.429 / 0.714 | $1.56 \times 10^{-14}$ / $9.39 \times 10^{-14}$ | 146 / 404 | cliffs_delta |
| AML420B | *EZH2* | 3 | 0.477 / 0.484 | 0.006; increasing/flat | 0.500 / 0.667 | 0.690 | 0.130 | – / – | – / – | – / – | cliffs_delta |
| AML420B | *FLT3* | 3 | -0.181 / -0.068 | 0.113; increasing/flat | 0.500 / 0.667 | 0.690 | 0.207 | 0.116 / 0.558 | 0.248 / 0.425 | 46 / 124 | cliffs_delta |
| AML420B | *GATA2* | 3 | 0.107 / 0.421 | 0.313; increasing/flat | 0.500 / 0.667 | 0.690 | 0.244 | – / – | – / – | – / – | cliffs_delta |
| AML420B | *IDH1* | 3 | 0.133 / -0.086 | -0.219; decreasing (response-consistent) | -1.000 / 0 | 0 | 0.449 | – / – | – / – | – / – | cliffs_delta |
| AML420B | *IDH2* | 3 | 0.112 / -0.225 | -0.337; decreasing (response-consistent) | -1.000 / 0 | 0 | 0.284 | -0.342 / 0.329 | 0.439 / 0.585 | 2 / 228 | cliffs_delta |
| AML420B | *KIT* | 3 | -0.120 / -0.071 | 0.048; increasing/flat | 0.500 / 0.667 | 0.690 | 0.471 | 0.005 / 0.502 | 0.981 / 0.981 | 23 / 36 | cliffs_delta |
| AML420B | *KRAS* | 3 | -0.136 / 0.071 | 0.207; increasing/flat | 0.500 / 0.667 | 0.690 | 0.240 | -0.366 / 0.317 | 0.434 / 0.585 | 2 / 41 | cliffs_delta |
| AML420B | *NPM1* | 3 | -0.066 / 0.180 | 0.246; increasing/flat | 0.500 / 0.667 | 0.690 | 0.318 | 0.506 / 0.753 | $6.11 \times 10^{-68}$ / $7.34 \times 10^{-67}$ | 542 / 1,457 | cliffs_delta |

Continued on next page

*Supplementary Table S7 continued*

| Patient | Driver | TP | First / last | Change / direction | $\rho$ / $p$ | Trend $q$ | Confidence | Truth $\delta$ / AUROC | Truth $p$ / $q$ | Truth cells mut / WT | Metric |
|---|---|---|---|---|---|---|---|---|---|---|---|
| AML420B | *NRAS* | 3 | -0.005 / 0.322 | 0.327; increasing/flat | 1.000 / 0 | 0 | 0.199 | 0.203 / 0.601 | 0.067 / 0.162 | 36 / 114 | cliffs_delta |
| AML420B | *PHF6* | 3 | 0.445 / 0.339 | -0.106; decreasing (response-consistent) | -0.500 / 0.667 | 0.690 | 0.251 | – / – | – / – | – / – | cliffs_delta |
| AML420B | *PTPN11* | 3 | 0.107 / 0.321 | 0.214; increasing/flat | 0.500 / 0.667 | 0.690 | 0.212 | – / – | – / – | – / – | cliffs_delta |
| AML420B | *RAD21* | 3 | -0.198 / -0.368 | -0.170; decreasing (response-consistent) | -0.500 / 0.667 | 0.690 | 0.237 | 0.417 / 0.708 | 0.035 / 0.106 | 9 / 187 | cliffs_delta |
| AML420B | *RUNX1* | 3 | 0.212 / 0.414 | 0.203; increasing/flat | 0.500 / 0.667 | 0.690 | 0.256 | -0.030 / 0.485 | 0.869 / 0.948 | 23 / 25 | cliffs_delta |
| AML420B | *SF3B1* | 3 | 0.103 / 0.328 | 0.226; increasing/flat | 0.500 / 0.667 | 0.690 | 0.264 | – / – | – / – | – / – | cliffs_delta |
| AML420B | *SMC1A* | 3 | 0.009 / 0.027 | 0.018; increasing/flat | 0.500 / 0.667 | 0.690 | 0.312 | – / – | – / – | – / – | cliffs_delta |
| AML420B | *SMC3* | 3 | 0.008 / -0.142 | -0.150; decreasing (response-consistent) | -0.500 / 0.667 | 0.690 | 0.311 | 0.368 / 0.684 | 0.169 / 0.338 | 7 / 19 | cliffs_delta |
| AML420B | *SRSF2* | 3 | 0.018 / -0.069 | -0.088; decreasing (response-consistent) | -1.000 / 0 | 0 | 0.341 | – / – | – / – | – / – | cliffs_delta |
| AML420B | *STAG2* | 3 | 0.208 / -0.010 | -0.217; decreasing (response-consistent) | -1.000 / 0 | 0 | 0.235 | – / – | – / – | – / – | cliffs_delta |
| AML420B | *TET2* | 3 | 0.101 / 0.310 | 0.209; increasing/flat | 0.500 / 0.667 | 0.690 | 0.214 | -0.093 / 0.454 | 0.602 / 0.722 | 18 / 30 | cliffs_delta |
| AML420B | *TP53* | 3 | 0.203 / 0.229 | 0.026; increasing/flat | 0.500 / 0.667 | 0.690 | 0.395 | 0.485 / 0.743 | $6.35 \times 10^{-4}$ / 0.003 | 51 / 25 | cliffs_delta |
| AML420B | *U2AF1* | 3 | 0.048 / 0.025 | -0.023; decreasing (response-consistent) | -0.500 / 0.667 | 0.690 | 0.166 | – / – | – / – | – / – | cliffs_delta |
| AML420B | *WT1* | 3 | 0.047 / 0.136 | 0.089; increasing/flat | 1.000 / 0 | 0 | 0.244 | – / – | – / – | – / – | cliffs_delta |
| AML420B | *ZRSR2* | 3 | -0.064 / 0.166 | 0.229; increasing/flat | 0.500 / 0.667 | 0.690 | 0.219 | – / – | – / – | – / – | cliffs_delta |
| AML556 | *ASXL1* | 3 | -0.109 / 0.095 | 0.204; increasing/flat | 0.500 / 0.667 | 0.690 | 0.209 | – / – | – / – | – / – | cliffs_delta |
| AML556 | *BCOR* | 3 | 0.089 / 0.426 | 0.337; increasing/flat | 1.000 / 0 | 0 | 0.177 | – / – | – / – | – / – | cliffs_delta |
| AML556 | *BCORL1* | 3 | -0.510 / -0.319 | 0.190; increasing/flat | 0.500 / 0.667 | 0.690 | 0.281 | – / – | – / – | – / – | cliffs_delta |
| AML556 | *CBL* | 3 | -0.284 / -0.081 | 0.203; increasing/flat | 0.500 / 0.667 | 0.690 | 0.163 | – / – | – / – | – / – | cliffs_delta |
| AML556 | *CEBPA* | 3 | 0.353 / 0.046 | -0.307; decreasing (response-consistent) | -0.500 / 0.667 | 0.690 | 0.370 | – / – | – / – | – / – | cliffs_delta |
| AML556 | *DNMT3A* | 3 | 0.737 / 0.255 | -0.482; decreasing (response-consistent) | -0.500 / 0.667 | 0.690 | 0.298 | 0.429 / 0.714 | $1.56 \times 10^{-14}$ / $9.39 \times 10^{-14}$ | 146 / 404 | cliffs_delta |
| AML556 | *EZH2* | 3 | -0.086 / 0.299 | 0.385; increasing/flat | 1.000 / 0 | 0 | 0.130 | – / – | – / – | – / – | cliffs_delta |
| AML556 | *FLT3* | 3 | 0.104 / -0.310 | -0.414; decreasing (response-consistent) | -0.500 / 0.667 | 0.690 | 0.207 | 0.116 / 0.558 | 0.248 / 0.425 | 46 / 124 | cliffs_delta |
| AML556 | *GATA2* | 3 | -0.361 / -0.137 | 0.224; increasing/flat | 0.500 / 0.667 | 0.690 | 0.244 | – / – | – / – | – / – | cliffs_delta |
| AML556 | *IDH1* | 3 | 0.153 / 0.010 | -0.143; decreasing (response-consistent) | -1.000 / 0 | 0 | 0.449 | – / – | – / – | – / – | cliffs_delta |
| AML556 | *IDH2* | 3 | -0.064 / -0.165 | -0.101; decreasing (response-consistent) | -1.000 / 0 | 0 | 0.284 | -0.342 / 0.329 | 0.439 / 0.585 | 2 / 228 | cliffs_delta |
| AML556 | *KIT* | 3 | 0.013 / -0.302 | -0.315; decreasing (response-consistent) | -0.500 / 0.667 | 0.690 | 0.471 | 0.005 / 0.502 | 0.981 / 0.981 | 23 / 36 | cliffs_delta |
| AML556 | *KRAS* | 3 | -0.397 / -0.216 | 0.180; increasing/flat | 0.500 / 0.667 | 0.690 | 0.240 | -0.366 / 0.317 | 0.434 / 0.585 | 2 / 41 | cliffs_delta |
| AML556 | *NPM1* | 3 | 0.634 / 0.104 | -0.529; decreasing (response-consistent) | -1.000 / 0 | 0 | 0.318 | 0.506 / 0.753 | $6.11 \times 10^{-68}$ / $7.34 \times 10^{-67}$ | 542 / 1,457 | cliffs_delta |
| AML556 | *NRAS* | 3 | 0.185 / 0.192 | 0.006; increasing/flat | 0.500 / 0.667 | 0.690 | 0.199 | 0.203 / 0.601 | 0.067 / 0.162 | 36 / 114 | cliffs_delta |
| AML556 | *PHF6* | 3 | 0.412 / 0.241 | -0.171; decreasing (response-consistent) | -0.500 / 0.667 | 0.690 | 0.251 | – / – | – / – | – / – | cliffs_delta |
| AML556 | *PTPN11* | 3 | 0.580 / -0.063 | -0.642; decreasing (response-consistent) | -1.000 / 0 | 0 | 0.212 | – / – | – / – | – / – | cliffs_delta |
| AML556 | *RAD21* | 3 | -0.198 / -0.255 | -0.057; decreasing (response-consistent) | -0.500 / 0.667 | 0.690 | 0.237 | 0.417 / 0.708 | 0.035 / 0.106 | 9 / 187 | cliffs_delta |
| AML556 | *RUNX1* | 3 | -0.402 / 0.387 | 0.790; increasing/flat | 1.000 / 0 | 0 | 0.256 | -0.030 / 0.485 | 0.869 / 0.948 | 23 / 25 | cliffs_delta |
| AML556 | *SF3B1* | 3 | -0.435 / -0.043 | 0.393; increasing/flat | 1.000 / 0 | 0 | 0.264 | – / – | – / – | – / – | cliffs_delta |
| AML556 | *SMC1A* | 3 | 0.135 / 0.055 | -0.079; decreasing (response-consistent) | -0.500 / 0.667 | 0.690 | 0.312 | – / – | – / – | – / – | cliffs_delta |
| AML556 | *SMC3* | 3 | 0.223 / -0.257 | -0.480; decreasing (response-consistent) | -1.000 / 0 | 0 | 0.311 | 0.368 / 0.684 | 0.169 / 0.338 | 7 / 19 | cliffs_delta |
| AML556 | *SRSF2* | 3 | -0.277 / 0.036 | 0.313; increasing/flat | 1.000 / 0 | 0 | 0.341 | – / – | – / – | – / – | cliffs_delta |
| AML556 | *STAG2* | 3 | 0.415 / 0.088 | -0.327; decreasing (response-consistent) | -1.000 / 0 | 0 | 0.235 | – / – | – / – | – / – | cliffs_delta |
| AML556 | *TET2* | 3 | 0.291 / 0.247 | -0.044; decreasing (response-consistent) | -0.500 / 0.667 | 0.690 | 0.214 | -0.093 / 0.454 | 0.602 / 0.722 | 18 / 30 | cliffs_delta |
| AML556 | *TP53* | 3 | 0.286 / 0.313 | 0.028; increasing/flat | 0.500 / 0.667 | 0.690 | 0.395 | 0.485 / 0.743 | $6.35 \times 10^{-4}$ / 0.003 | 51 / 25 | cliffs_delta |
| AML556 | *U2AF1* | 3 | -0.388 / 0.120 | 0.508; increasing/flat | 0.500 / 0.667 | 0.690 | 0.166 | – / – | – / – | – / – | cliffs_delta |

Continued on next page

*Supplementary Table S7 continued*

| Patient | Driver | TP | First / last | Change / direction | $\rho$ / $p$ | Trend $q$ | Confidence | Truth $\delta$ / AUROC | Truth $p$ / $q$ | Truth cells mut / WT | Metric |
|---|---|---|---|---|---|---|---|---|---|---|---|
| AML556 | *WT1* | 3 | 0.264 / -0.083 | -0.347; decreasing (response-consistent) | -1.000 / 0 | 0 | 0.244 | – / – | – / – | – / – | cliffs_delta |
| AML556 | *ZRSR2* | 3 | -0.226 / 0.143 | 0.369; increasing/flat | 0.500 / 0.667 | 0.690 | 0.219 | – / – | – / – | – / – | cliffs_delta |
| AML707B | *ASXL1* | 5 | -0.335 / 0.050 | 0.385; increasing/flat | 0.300 / 0.624 | 0.690 | 0.209 | – / – | – / – | – / – | cliffs_delta |
| AML707B | *BCOR* | 5 | -0.394 / 0.067 | 0.461; increasing/flat | 0.300 / 0.624 | 0.690 | 0.177 | – / – | – / – | – / – | cliffs_delta |
| AML707B | *BCORL1* | 5 | -0.668 / -0.045 | 0.623; increasing/flat | 0.700 / 0.188 | 0.569 | 0.281 | – / – | – / – | – / – | cliffs_delta |
| AML707B | *CBL* | 5 | 0.520 / 0.271 | -0.249; decreasing (response-consistent) | -0.400 / 0.505 | 0.690 | 0.163 | – / – | – / – | – / – | cliffs_delta |
| AML707B | *CEBPA* | 5 | 0.733 / -0.173 | -0.906; decreasing (response-consistent) | -0.900 / 0.037 | 0.147 | 0.370 | – / – | – / – | – / – | cliffs_delta |
| AML707B | *DNMT3A* | 5 | -0.723 / -0.067 | 0.656; increasing/flat | 0.600 / 0.285 | 0.690 | 0.298 | 0.429 / 0.714 | $1.56 \times 10^{-14}$ / $9.39 \times 10^{-14}$ | 146 / 404 | cliffs_delta |
| AML707B | *EZH2* | 5 | 0.935 / 0.155 | -0.780; decreasing (response-consistent) | -0.900 / 0.037 | 0.147 | 0.130 | – / – | – / – | – / – | cliffs_delta |
| AML707B | *FLT3* | 5 | 0.313 / 0.098 | -0.215; decreasing (response-consistent) | -0.400 / 0.505 | 0.690 | 0.207 | 0.116 / 0.558 | 0.248 / 0.425 | 46 / 124 | cliffs_delta |
| AML707B | *GATA2* | 5 | 0.595 / 0.235 | -0.360; decreasing (response-consistent) | -0.800 / 0.104 | 0.387 | 0.244 | – / – | – / – | – / – | cliffs_delta |
| AML707B | *IDH1* | 5 | -0.154 / -0.008 | 0.146; increasing/flat | -0.200 / 0.747 | 0.763 | 0.449 | – / – | – / – | – / – | cliffs_delta |
| AML707B | *IDH2* | 5 | -0.714 / -0.040 | 0.674; increasing/flat | 0.400 / 0.505 | 0.690 | 0.284 | -0.342 / 0.329 | 0.439 / 0.585 | 2 / 228 | cliffs_delta |
| AML707B | *KIT* | 5 | 0.431 / 0.152 | -0.280; decreasing (response-consistent) | -0.300 / 0.624 | 0.690 | 0.471 | 0.005 / 0.502 | 0.981 / 0.981 | 23 / 36 | cliffs_delta |
| AML707B | *KRAS* | 5 | -0.296 / 0.008 | 0.304; increasing/flat | 0.100 / 0.873 | 0.879 | 0.240 | -0.366 / 0.317 | 0.434 / 0.585 | 2 / 41 | cliffs_delta |
| AML707B | *NPM1* | 5 | -0.396 / 0.149 | 0.545; increasing/flat | 0.700 / 0.188 | 0.569 | 0.318 | 0.506 / 0.753 | $6.11 \times 10^{-68}$ / $7.34 \times 10^{-67}$ | 542 / 1,457 | cliffs_delta |
| AML707B | *NRAS* | 5 | 0.336 / 0.213 | -0.123; decreasing (response-consistent) | -0.600 / 0.285 | 0.690 | 0.199 | 0.203 / 0.601 | 0.067 / 0.162 | 36 / 114 | cliffs_delta |
| AML707B | *PHF6* | 5 | 0.620 / -0.064 | -0.684; decreasing (response-consistent) | -0.600 / 0.285 | 0.690 | 0.251 | – / – | – / – | – / – | cliffs_delta |
| AML707B | *PTPN11* | 5 | -0.176 / 0.201 | 0.377; increasing/flat | 1.000 / $1.40 \times 10^{-24}$ | $6.17 \times 10^{-24}$ | 0.212 | – / – | – / – | – / – | cliffs_delta |
| AML707B | *RAD21* | 5 | 0.488 / -0.122 | -0.610; decreasing (response-consistent) | -0.900 / 0.037 | 0.147 | 0.237 | 0.417 / 0.708 | 0.035 / 0.106 | 9 / 187 | cliffs_delta |
| AML707B | *RUNX1* | 5 | -0.012 / 0.015 | 0.026; increasing/flat | 0.300 / 0.624 | 0.690 | 0.256 | -0.030 / 0.485 | 0.869 / 0.948 | 23 / 25 | cliffs_delta |
| AML707B | *SF3B1* | 5 | 0.055 / -0.021 | -0.076; decreasing (response-consistent) | -0.300 / 0.624 | 0.690 | 0.264 | – / – | – / – | – / – | cliffs_delta |
| AML707B | *SMC1A* | 5 | 0.120 / 0.230 | 0.110; increasing/flat | 0.400 / 0.505 | 0.690 | 0.312 | – / – | – / – | – / – | cliffs_delta |
| AML707B | *SMC3* | 5 | 0.106 / -0.060 | -0.166; decreasing (response-consistent) | -0.900 / 0.037 | 0.147 | 0.311 | 0.368 / 0.684 | 0.169 / 0.338 | 7 / 19 | cliffs_delta |
| AML707B | *SRSF2* | 5 | -0.586 / -0.081 | 0.506; increasing/flat | 0.100 / 0.873 | 0.879 | 0.341 | – / – | – / – | – / – | cliffs_delta |
| AML707B | *STAG2* | 5 | -0.206 / 0.177 | 0.383; increasing/flat | 0.300 / 0.624 | 0.690 | 0.235 | – / – | – / – | – / – | cliffs_delta |
| AML707B | *TET2* | 5 | 0.368 / -0.037 | -0.405; decreasing (response-consistent) | -0.500 / 0.391 | 0.690 | 0.214 | -0.093 / 0.454 | 0.602 / 0.722 | 18 / 30 | cliffs_delta |
| AML707B | *TP53* | 5 | 0.009 / -0.109 | -0.117; decreasing (response-consistent) | 0.200 / 0.747 | 0.763 | 0.395 | 0.485 / 0.743 | $6.35 \times 10^{-4}$ / 0.003 | 51 / 25 | cliffs_delta |
| AML707B | *U2AF1* | 5 | -0.547 / 0.017 | 0.564; increasing/flat | 0.700 / 0.188 | 0.569 | 0.166 | – / – | – / – | – / – | cliffs_delta |
| AML707B | *WT1* | 5 | 0.355 / 0.255 | -0.100; decreasing (response-consistent) | -0.300 / 0.624 | 0.690 | 0.244 | – / – | – / – | – / – | cliffs_delta |
| AML707B | *ZRSR2* | 5 | 0.375 / 0.076 | -0.299; decreasing (response-consistent) | -0.800 / 0.104 | 0.387 | 0.219 | – / – | – / – | – / – | cliffs_delta |
| AML916 | *ASXL1* | 1 | -0.530 / -0.530 | 0.000; increasing/flat | – / – | – | 0.209 | – / – | – / – | – / – | cliffs_delta |
| AML916 | *BCOR* | 1 | -0.226 / -0.226 | 0.000; increasing/flat | – / – | – | 0.177 | – / – | – / – | – / – | cliffs_delta |
| AML916 | *BCORL1* | 1 | -0.488 / -0.488 | 0.000; increasing/flat | – / – | – | 0.281 | – / – | – / – | – / – | cliffs_delta |
| AML916 | *CBL* | 1 | -0.199 / -0.199 | 0.000; increasing/flat | – / – | – | 0.163 | – / – | – / – | – / – | cliffs_delta |
| AML916 | *CEBPA* | 1 | -0.390 / -0.390 | 0.000; increasing/flat | – / – | – | 0.370 | – / – | – / – | – / – | cliffs_delta |
| AML916 | *DNMT3A* | 1 | -0.007 / -0.007 | 0.000; increasing/flat | – / – | – | 0.298 | 0.429 / 0.714 | $1.56 \times 10^{-14}$ / $9.39 \times 10^{-14}$ | 146 / 404 | cliffs_delta |
| AML916 | *EZH2* | 1 | -0.390 / -0.390 | 0.000; increasing/flat | – / – | – | 0.130 | – / – | – / – | – / – | cliffs_delta |
| AML916 | *FLT3* | 1 | -0.706 / -0.706 | 0.000; increasing/flat | – / – | – | 0.207 | 0.116 / 0.558 | 0.248 / 0.425 | 46 / 124 | cliffs_delta |
| AML916 | *GATA2* | 1 | -0.128 / -0.128 | 0.000; increasing/flat | – / – | – | 0.244 | – / – | – / – | – / – | cliffs_delta |
| AML916 | *IDH1* | 1 | 0.190 / 0.190 | 0.000; increasing/flat | – / – | – | 0.449 | – / – | – / – | – / – | cliffs_delta |
| AML916 | *IDH2* | 1 | -0.371 / -0.371 | 0.000; increasing/flat | – / – | – | 0.284 | -0.342 / 0.329 | 0.439 / 0.585 | 2 / 228 | cliffs_delta |
| AML916 | *KIT* | 1 | 0.228 / 0.228 | 0.000; increasing/flat | – / – | – | 0.471 | 0.005 / 0.502 | 0.981 / 0.981 | 23 / 36 | cliffs_delta |

Continued on next page

*Supplementary Table S7 continued*

| Patient | Driver | TP | First / last | Change / direction | $\rho$ / $p$ | Trend $q$ | Confidence | Truth $\delta$ / AUROC | Truth $p$ / $q$ | Truth cells mut / WT | Metric |
|---|---|---|---|---|---|---|---|---|---|---|---|
| AML916 | *KRAS* | 1 | 0.331 / 0.331 | 0.000; increasing/flat | – / – | – | 0.240 | -0.366 / 0.317 | 0.434 / 0.585 | 2 / 41 | cliffs_delta |
| AML916 | *NPM1* | 1 | -0.269 / -0.269 | 0.000; increasing/flat | – / – | – | 0.318 | 0.506 / 0.753 | $6.11 \times 10^{-68}$ / $7.34 \times 10^{-67}$ | 542 / 1,457 | cliffs_delta |
| AML916 | *NRAS* | 1 | 0.262 / 0.262 | 0.000; increasing/flat | – / – | – | 0.199 | 0.203 / 0.601 | 0.067 / 0.162 | 36 / 114 | cliffs_delta |
| AML916 | *PHF6* | 1 | -0.069 / -0.069 | 0.000; increasing/flat | – / – | – | 0.251 | – / – | – / – | – / – | cliffs_delta |
| AML916 | *PTPN11* | 1 | -0.128 / -0.128 | 0.000; increasing/flat | – / – | – | 0.212 | – / – | – / – | – / – | cliffs_delta |
| AML916 | *RAD21* | 1 | -0.921 / -0.921 | 0.000; increasing/flat | – / – | – | 0.237 | 0.417 / 0.708 | 0.035 / 0.106 | 9 / 187 | cliffs_delta |
| AML916 | *RUNX1* | 1 | -0.041 / -0.041 | 0.000; increasing/flat | – / – | – | 0.256 | -0.030 / 0.485 | 0.869 / 0.948 | 23 / 25 | cliffs_delta |
| AML916 | *SF3B1* | 1 | -0.391 / -0.391 | 0.000; increasing/flat | – / – | – | 0.264 | – / – | – / – | – / – | cliffs_delta |
| AML916 | *SMC1A* | 1 | 0.010 / 0.010 | 0.000; increasing/flat | – / – | – | 0.312 | – / – | – / – | – / – | cliffs_delta |
| AML916 | *SMC3* | 1 | -0.512 / -0.512 | 0.000; increasing/flat | – / – | – | 0.311 | 0.368 / 0.684 | 0.169 / 0.338 | 7 / 19 | cliffs_delta |
| AML916 | *SRSF2* | 1 | -0.128 / -0.128 | 0.000; increasing/flat | – / – | – | 0.341 | – / – | – / – | – / – | cliffs_delta |
| AML916 | *STAG2* | 1 | -0.159 / -0.159 | 0.000; increasing/flat | – / – | – | 0.235 | – / – | – / – | – / – | cliffs_delta |
| AML916 | *TET2* | 1 | -0.717 / -0.717 | 0.000; increasing/flat | – / – | – | 0.214 | -0.093 / 0.454 | 0.602 / 0.722 | 18 / 30 | cliffs_delta |
| AML916 | *TP53* | 1 | 0.897 / 0.897 | 0.000; increasing/flat | – / – | – | 0.395 | 0.485 / 0.743 | $6.35 \times 10^{-4}$ / 0.003 | 51 / 25 | cliffs_delta |
| AML916 | *U2AF1* | 1 | 0.031 / 0.031 | 0.000; increasing/flat | – / – | – | 0.166 | – / – | – / – | – / – | cliffs_delta |
| AML916 | *WT1* | 1 | -0.368 / -0.368 | 0.000; increasing/flat | – / – | – | 0.244 | – / – | – / – | – / – | cliffs_delta |
| AML916 | *ZRSR2* | 1 | -0.644 / -0.644 | 0.000; increasing/flat | – / – | – | 0.219 | – / – | – / – | – / – | cliffs_delta |
| AML921A | *ASXL1* | 1 | 0.340 / 0.340 | 0.000; increasing/flat | – / – | – | 0.209 | – / – | – / – | – / – | cliffs_delta |
| AML921A | *BCOR* | 1 | 0.346 / 0.346 | 0.000; increasing/flat | – / – | – | 0.177 | – / – | – / – | – / – | cliffs_delta |
| AML921A | *BCORL1* | 1 | -0.131 / -0.131 | 0.000; increasing/flat | – / – | – | 0.281 | – / – | – / – | – / – | cliffs_delta |
| AML921A | *CBL* | 1 | 0.193 / 0.193 | 0.000; increasing/flat | – / – | – | 0.163 | – / – | – / – | – / – | cliffs_delta |
| AML921A | *CEBPA* | 1 | 0.626 / 0.626 | 0.000; increasing/flat | – / – | – | 0.370 | – / – | – / – | – / – | cliffs_delta |
| AML921A | *DNMT3A* | 1 | 0.656 / 0.656 | 0.000; increasing/flat | – / – | – | 0.298 | 0.429 / 0.714 | $1.56 \times 10^{-14}$ / $9.39 \times 10^{-14}$ | 146 / 404 | cliffs_delta |
| AML921A | *EZH2* | 1 | -0.020 / -0.020 | 0.000; increasing/flat | – / – | – | 0.130 | – / – | – / – | – / – | cliffs_delta |
| AML921A | *FLT3* | 1 | 0.118 / 0.118 | 0.000; increasing/flat | – / – | – | 0.207 | 0.116 / 0.558 | 0.248 / 0.425 | 46 / 124 | cliffs_delta |
| AML921A | *GATA2* | 1 | 0.207 / 0.207 | 0.000; increasing/flat | – / – | – | 0.244 | – / – | – / – | – / – | cliffs_delta |
| AML921A | *IDH1* | 1 | 0.131 / 0.131 | 0.000; increasing/flat | – / – | – | 0.449 | – / – | – / – | – / – | cliffs_delta |
| AML921A | *IDH2* | 1 | -0.157 / -0.157 | 0.000; increasing/flat | – / – | – | 0.284 | -0.342 / 0.329 | 0.439 / 0.585 | 2 / 228 | cliffs_delta |
| AML921A | *KIT* | 1 | 0.237 / 0.237 | 0.000; increasing/flat | – / – | – | 0.471 | 0.005 / 0.502 | 0.981 / 0.981 | 23 / 36 | cliffs_delta |
| AML921A | *KRAS* | 1 | 0.077 / 0.077 | 0.000; increasing/flat | – / – | – | 0.240 | -0.366 / 0.317 | 0.434 / 0.585 | 2 / 41 | cliffs_delta |
| AML921A | *NPM1* | 1 | 0.384 / 0.384 | 0.000; increasing/flat | – / – | – | 0.318 | 0.506 / 0.753 | $6.11 \times 10^{-68}$ / $7.34 \times 10^{-67}$ | 542 / 1,457 | cliffs_delta |
| AML921A | *NRAS* | 1 | -0.019 / -0.019 | 0.000; increasing/flat | – / – | – | 0.199 | 0.203 / 0.601 | 0.067 / 0.162 | 36 / 114 | cliffs_delta |
| AML921A | *PHF6* | 1 | 0.124 / 0.124 | 0.000; increasing/flat | – / – | – | 0.251 | – / – | – / – | – / – | cliffs_delta |
| AML921A | *PTPN11* | 1 | -0.078 / -0.078 | 0.000; increasing/flat | – / – | – | 0.212 | – / – | – / – | – / – | cliffs_delta |
| AML921A | *RAD21* | 1 | 0.187 / 0.187 | 0.000; increasing/flat | – / – | – | 0.237 | 0.417 / 0.708 | 0.035 / 0.106 | 9 / 187 | cliffs_delta |
| AML921A | *RUNX1* | 1 | 0.388 / 0.388 | 0.000; increasing/flat | – / – | – | 0.256 | -0.030 / 0.485 | 0.869 / 0.948 | 23 / 25 | cliffs_delta |
| AML921A | *SF3B1* | 1 | 0.171 / 0.171 | 0.000; increasing/flat | – / – | – | 0.264 | – / – | – / – | – / – | cliffs_delta |
| AML921A | *SMC1A* | 1 | 0.299 / 0.299 | 0.000; increasing/flat | – / – | – | 0.312 | – / – | – / – | – / – | cliffs_delta |
| AML921A | *SMC3* | 1 | 0.239 / 0.239 | 0.000; increasing/flat | – / – | – | 0.311 | 0.368 / 0.684 | 0.169 / 0.338 | 7 / 19 | cliffs_delta |
| AML921A | *SRSF2* | 1 | -0.120 / -0.120 | 0.000; increasing/flat | – / – | – | 0.341 | – / – | – / – | – / – | cliffs_delta |
| AML921A | *STAG2* | 1 | 0.359 / 0.359 | 0.000; increasing/flat | – / – | – | 0.235 | – / – | – / – | – / – | cliffs_delta |
| AML921A | *TET2* | 1 | 0.215 / 0.215 | 0.000; increasing/flat | – / – | – | 0.214 | -0.093 / 0.454 | 0.602 / 0.722 | 18 / 30 | cliffs_delta |
| AML921A | *TP53* | 1 | -0.437 / -0.437 | 0.000; increasing/flat | – / – | – | 0.395 | 0.485 / 0.743 | $6.35 \times 10^{-4}$ / 0.003 | 51 / 25 | cliffs_delta |
| AML921A | *U2AF1* | 1 | -0.087 / -0.087 | 0.000; increasing/flat | – / – | – | 0.166 | – / – | – / – | – / – | cliffs_delta |
| AML921A | *WT1* | 1 | 0.424 / 0.424 | 0.000; increasing/flat | – / – | – | 0.244 | – / – | – / – | – / – | cliffs_delta |
| AML921A | *ZRSR2* | 1 | -0.196 / -0.196 | 0.000; increasing/flat | – / – | – | 0.219 | – / – | – / – | – / – | cliffs_delta |
| AML997 | *ASXL1* | 2 | 0.295 / 0.534 | 0.238; increasing/flat | – / – | – | 0.209 | – / – | – / – | – / – | cliffs_delta |
| AML997 | *BCOR* | 2 | 0.562 / 0.772 | 0.210; increasing/flat | – / – | – | 0.177 | – / – | – / – | – / – | cliffs_delta |
| AML997 | *BCORL1* | 2 | -0.059 / -0.233 | -0.174; decreasing (response-consistent) | – / – | – | 0.281 | – / – | – / – | – / – | cliffs_delta |
| AML997 | *CBL* | 2 | 0.323 / 0.153 | -0.171; decreasing (response-consistent) | – / – | – | 0.163 | – / – | – / – | – / – | cliffs_delta |
| AML997 | *CEBPA* | 2 | -0.028 / -0.489 | -0.461; decreasing (response-consistent) | – / – | – | 0.370 | – / – | – / – | – / – | cliffs_delta |
| AML997 | *DNMT3A* | 2 | 0.233 / 0.285 | 0.051; increasing/flat | – / – | – | 0.298 | 0.429 / 0.714 | $1.56 \times 10^{-14}$ / $9.39 \times 10^{-14}$ | 146 / 404 | cliffs_delta |
| AML997 | *EZH2* | 2 | -0.291 / 0.020 | 0.310; increasing/flat | – / – | – | 0.130 | – / – | – / – | – / – | cliffs_delta |

Continued on next page

*Supplementary Table S7 continued*

| Patient | Driver | TP | First / last | Change / direction | $\rho$ / $p$ | Trend $q$ | Confidence | Truth $\delta$ / AUROC | Truth $p$ / $q$ | Truth cells mut / WT | Metric |
|---|---|---|---|---|---|---|---|---|---|---|---|
| AML997 | *FLT3* | 2 | 0.586 / 0.085 | -0.501; decreasing (response-consistent) | – / – | – | 0.207 | 0.116 / 0.558 | 0.248 / 0.425 | 46 / 124 | cliffs_delta |
| AML997 | *GATA2* | 2 | 0.027 / -0.405 | -0.431; decreasing (response-consistent) | – / – | – | 0.244 | – / – | – / – | – / – | cliffs_delta |
| AML997 | *IDH1* | 2 | -0.063 / -0.416 | -0.353; decreasing (response-consistent) | – / – | – | 0.449 | – / – | – / – | – / – | cliffs_delta |
| AML997 | *IDH2* | 2 | -0.320 / -0.228 | 0.093; increasing/flat | – / – | – | 0.284 | -0.342 / 0.329 | 0.439 / 0.585 | 2 / 228 | cliffs_delta |
| AML997 | *KIT* | 2 | 0.179 / -0.610 | -0.788; decreasing (response-consistent) | – / – | – | 0.471 | 0.005 / 0.502 | 0.981 / 0.981 | 23 / 36 | cliffs_delta |
| AML997 | *KRAS* | 2 | 0.227 / -0.086 | -0.312; decreasing (response-consistent) | – / – | – | 0.240 | -0.366 / 0.317 | 0.434 / 0.585 | 2 / 41 | cliffs_delta |
| AML997 | *NPM1* | 2 | 0.706 / -0.134 | -0.840; decreasing (response-consistent) | – / – | – | 0.318 | 0.506 / 0.753 | $6.11 \times 10^{-68}$ / $7.34 \times 10^{-67}$ | 542 / 1,457 | cliffs_delta |
| AML997 | *NRAS* | 2 | 0.447 / -0.008 | -0.455; decreasing (response-consistent) | – / – | – | 0.199 | 0.203 / 0.601 | 0.067 / 0.162 | 36 / 114 | cliffs_delta |
| AML997 | *PHF6* | 2 | -0.220 / 0.346 | 0.566; increasing/flat | – / – | – | 0.251 | – / – | – / – | – / – | cliffs_delta |
| AML997 | *PTPN11* | 2 | 0.338 / 0.225 | -0.113; decreasing (response-consistent) | – / – | – | 0.212 | – / – | – / – | – / – | cliffs_delta |
| AML997 | *RAD21* | 2 | 0.251 / -0.249 | -0.500; decreasing (response-consistent) | – / – | – | 0.237 | 0.417 / 0.708 | 0.035 / 0.106 | 9 / 187 | cliffs_delta |
| AML997 | *RUNX1* | 2 | 0.259 / 0.659 | 0.400; increasing/flat | – / – | – | 0.256 | -0.030 / 0.485 | 0.869 / 0.948 | 23 / 25 | cliffs_delta |
| AML997 | *SF3B1* | 2 | 0.382 / 0.347 | -0.035; decreasing (response-consistent) | – / – | – | 0.264 | – / – | – / – | – / – | cliffs_delta |
| AML997 | *SMC1A* | 2 | 0.018 / -0.333 | -0.352; decreasing (response-consistent) | – / – | – | 0.312 | – / – | – / – | – / – | cliffs_delta |
| AML997 | *SMC3* | 2 | 0.002 / -0.635 | -0.637; decreasing (response-consistent) | – / – | – | 0.311 | 0.368 / 0.684 | 0.169 / 0.338 | 7 / 19 | cliffs_delta |
| AML997 | *SRSF2* | 2 | -0.197 / -0.137 | 0.060; increasing/flat | – / – | – | 0.341 | – / – | – / – | – / – | cliffs_delta |
| AML997 | *STAG2* | 2 | 0.152 / -0.256 | -0.409; decreasing (response-consistent) | – / – | – | 0.235 | – / – | – / – | – / – | cliffs_delta |
| AML997 | *TET2* | 2 | 0.336 / 0.009 | -0.328; decreasing (response-consistent) | – / – | – | 0.214 | -0.093 / 0.454 | 0.602 / 0.722 | 18 / 30 | cliffs_delta |
| AML997 | *TP53* | 2 | -0.366 / 0.350 | 0.716; increasing/flat | – / – | – | 0.395 | 0.485 / 0.743 | $6.35 \times 10^{-4}$ / 0.003 | 51 / 25 | cliffs_delta |
| AML997 | *U2AF1* | 2 | -0.089 / 0.448 | 0.537; increasing/flat | – / – | – | 0.166 | – / – | – / – | – / – | cliffs_delta |
| AML997 | *WT1* | 2 | 0.571 / 0.091 | -0.481; decreasing (response-consistent) | – / – | – | 0.244 | – / – | – / – | – / – | cliffs_delta |
| AML997 | *ZRSR2* | 2 | 0.060 / -0.076 | -0.136; decreasing (response-consistent) | – / – | – | 0.219 | – / – | – / – | – / – | cliffs_delta |

**Supplementary Table S8. Panel-level provenance.** Panel-level provenance linking manuscript items to their source artifacts, analysis version, configuration, model object, alignment, component count, score field, subset, statistical unit, test or metric, correction family, random seed, and canonical status.

| Manuscript item / panel | Complete provenance record |
|---|---|
| **Figure 1 / A-C** | **Purpose:** Workflow schematic<br>**Source artifact:** `figures/Fig_1_overview/Figure_1_overview.png`<br>**Analysis version:** v10.3.2.2<br>**Config file:** `drivers/config_canonical_v3.json`<br>**Config hash:** `295f7b08b4e769f84eca625b76dbfad56b6bd025b1acd5cec1e5528b fc3dfcf2`<br>**Model ID:** fresh v10.3.2.2 forced-rescore model/output object<br>**Cancer:** all<br>**Driver:** all<br>**Alignment:** moment_matching<br>**Components:** 30<br>**Score column:** not applicable<br>**Score kind:** schematic<br>**Cell subset:** not applicable<br>**Statistical unit:** schematic<br>**Test/metric:** not applicable<br>**Multiple-testing family:** not applicable<br>**Random seed:** 67<br>**Status:** canonical |
| **Figure 2 / A-D** | **Purpose:** Bulk discrimination and latent interpretation<br>**Source artifact:** `Table1_bulk_discrimination.csv;SVD_interpretation_cv_audit_ *.csv;genes_<cancer>_<driver>.csv`<br>**Analysis version:** v10.3.2.2<br>**Config file:** `drivers/config_canonical_v3.json`<br>**Config hash:** `295f7b08b4e769f84eca625b76dbfad56b6bd025b1acd5cec1e5528b fc3dfcf2`<br>**Model ID:** fresh cohort-specific bulk fit from canonical run<br>**Cancer:** all<br>**Driver:** all<br>**Alignment:** not applicable to bulk<br>**Components:** 30<br>**Score column:** out-of-fold probability<br>**Score kind:** bulk probability<br>**Cell subset:** bulk tumors<br>**Statistical unit:** tumor<br>**Test/metric:** AUROC, AUPRC, Brier, bootstrap CI, label permutation<br>**Multiple-testing family:** all tested cancer-driver models<br>**Random seed:** 67<br>**Status:** canonical<br>**Notes:** Bulk models refit in this run. |

Continued on next page

| Manuscript item / panel | Complete provenance record |
|---|---|
| **Figure 3 / A-B** | **Purpose:** Ground-truth-free confidence<br>**Source artifact:** `Fig3_confidence_AML_vanGalen.csv;Fig3_confidence_validation_AML_vanGalen.csv`<br>**Analysis version:** v10.3.2.2<br>**Config file:** `drivers/config_canonical_v3.json`<br>**Config hash:** `295f7b08b4e769f84eca625b76dbfad56b6bd025b1acd5cec1e5528bfc3dfcf2`<br>**Model ID:** fresh v10.3.2.2 forced-rescore model/output object<br>**Cancer:** AML<br>**Driver:** truth-evaluable drivers<br>**Alignment:** moment_matching<br>**Components:** 30<br>**Score column:** mutation_prob_*<br>**Score kind:** raw probability<br>**Cell subset:** all retained cells<br>**Statistical unit:** driver<br>**Test/metric:** weighted geometric confidence; Spearman; AUROC<br>**Multiple-testing family:** AML truth-evaluable drivers<br>**Random seed:** 67<br>**Status:** canonical<br>**Notes:** Confidence components use raw scores. |
| **Figure 3 / C-G** | **Purpose:** AML direct per-cell mutation-label validation<br>**Source artifact:** `Fig3_percell_AML_vanGalen.csv;MutTranscript_genotype_parse_AML_vanGalen.csv`<br>**Analysis version:** v10.3.2.2<br>**Config file:** `drivers/config_canonical_v3.json`<br>**Config hash:** `295f7b08b4e769f84eca625b76dbfad56b6bd025b1acd5cec1e5528bfc3dfcf2`<br>**Model ID:** fresh v10.3.2.2 forced-rescore model/output object<br>**Cancer:** AML<br>**Driver:** NPM1, TP53, DNMT3A and all evaluable<br>**Alignment:** moment_matching<br>**Components:** 30<br>**Score column:** mutation_resid_*<br>**Score kind:** cell-state residual raw<br>**Cell subset:** cells with noncontradictory MutTranscripts/WtTranscripts labels<br>**Statistical unit:** cell<br>**Test/metric:** apparent AUROC, Mann-Whitney, Cliff delta<br>**Multiple-testing family:** AML evaluable drivers<br>**Random seed:** 67<br>**Status:** canonical<br>**Notes:** Raw scores and derived residuals recomputed in v10.3.2.2. |

Continued on next page

| Manuscript item / panel | Complete provenance record |
|---|---|
| **Figure 4 / A-J** | **Purpose:** Matched genotype and score localization<br>**Source artifact:** `Fig4_triptych_genotype_audit_AML_vanGalen.csv;Fig4_triptych_genotype_audit_CRC_Lee.csv`<br>**Analysis version:** v10.3.2.2<br>**Config file:** `drivers/config_canonical_v3.json`<br>**Config hash:** `295f7b08b4e769f84eca625b76dbfad56b6bd025b1acd5cec1e5528bfc3dfcf2`<br>**Model ID:** fresh v10.3.2.2 forced-rescore model/output object<br>**Cancer:** AML, CRC<br>**Driver:** displayed drivers<br>**Alignment:** moment_matching<br>**Components:** 30<br>**Score column:** mutation_resid_*<br>**Score kind:** cell-state residual raw<br>**Cell subset:** retained cells on shared expression UMAP<br>**Statistical unit:** cell<br>**Test/metric:** descriptive localization<br>**Multiple-testing family:** not applicable<br>**Random seed:** 67<br>**Status:** canonical<br>**Notes:** AML context is per-cell; CRC context is patient-level. |
| **Figure 5 / A-F** | **Purpose:** Longitudinal AML dynamics<br>**Source artifact:** `Fig3b_trajectory_AML_vanGalen.csv;Fig3b_tpviolin_significance_*.csv`<br>**Analysis version:** v10.3.2.2<br>**Config file:** `drivers/config_canonical_v3.json`<br>**Config hash:** `295f7b08b4e769f84eca625b76dbfad56b6bd025b1acd5cec1e5528bfc3dfcf2`<br>**Model ID:** fresh v10.3.2.2 forced-rescore model/output object<br>**Cancer:** AML<br>**Driver:** NPM1 and displayed controls<br>**Alignment:** moment_matching<br>**Components:** 30<br>**Score column:** mutation_resid_*<br>**Score kind:** cell-state residual raw<br>**Cell subset:** prespecified AML patient/sample filters plus fixed BM reference<br>**Statistical unit:** patient-time point and cell<br>**Test/metric:** Cliff delta trajectory; Mann-Whitney within panel<br>**Multiple-testing family:** within displayed panel/driver families<br>**Random seed:** 67<br>**Status:** canonical<br>**Notes:** AML870 excluded; singleton diagnosis panels are cross-sectional. |

Continued on next page

| Manuscript item / panel | Complete provenance record |
|---|---|
| **Figure 6 / A-G** | **Purpose:** Cell-state localization atlas<br>**Source artifact:** `Fig4_cluster_heatmap_*.csvandcohortUMAPexports`<br>**Analysis version:** v10.3.2.2<br>**Config file:** `drivers/config_canonical_v3.json`<br>**Config hash:** `295f7b08b4e769f84eca625b76dbfad56b6bd025b1acd5cec1e5528bfc3dfcf2`<br>**Model ID:** fresh v10.3.2.2 forced-rescore model/output object<br>**Cancer:** AML, BRCA<br>**Driver:** displayed drivers<br>**Alignment:** moment_matching<br>**Components:** 30<br>**Score column:** mutation_resid_*<br>**Score kind:** cell-state residual raw<br>**Cell subset:** retained cells and source/coarse annotations<br>**Statistical unit:** cell and cell type<br>**Test/metric:** mean residual score; descriptive UMAP localization<br>**Multiple-testing family:** not applicable<br>**Random seed:** 67<br>**Status:** canonical<br>**Notes:** Fresh HVG UMAP; 3,000 HVGs, 50 PCs, 15 neighbors, min_dist 0.5. |
| **Figure 7 / A-E** | **Purpose:** CNV concordance and discordance<br>**Source artifact:** `Fig6_discordance_*.csv;Fig6_score_correlation_*.csv;Fig6_patient_fraction_*.csv`<br>**Analysis version:** v10.3.2.2<br>**Config file:** `drivers/config_canonical_v3.json`<br>**Config hash:** `295f7b08b4e769f84eca625b76dbfad56b6bd025b1acd5cec1e5528bfc3dfcf2`<br>**Model ID:** fresh v10.3.2.2 forced-rescore model/output object<br>**Cancer:** all<br>**Driver:** locked primary-driver set<br>**Alignment:** moment_matching<br>**Components:** 30<br>**Score column:** scope_max_driver_score = max(mutation_prob_*)<br>**Score kind:** unadjusted raw maximum<br>**Cell subset:** CNV-matched cells/patients<br>**Statistical unit:** cell and patient<br>**Test/metric:** Spearman, ARI, quadrant counts, malignant fractions<br>**Multiple-testing family:** cohort-specific<br>**Random seed:** 67<br>**Status:** canonical<br>**Notes:** No confidence adjustment or residual score used; scOPE reference threshold q=0.8; CNV reference threshold q=0.95. |

**Supplementary Table S9. Latent-dimension sensitivity.** Cancer-, driver-, and component-count-specific cross-validated AUROC and AUPRC values from the archived latent-dimension sensitivity grid. The canonical model used 30 components; the sensitivity grid evaluated the component counts listed here.

| Cancer | $k$ | Driver | AUROC mean | AUROC SD | AUPRC mean | Samples | CV folds | Implementation |
|---|---|---|---|---|---|---|---|---|
| AML | 5 | *ASXL1* | 0.635 | – | 0.191 | 651 | 2 | suite_bulk_oof_hardened |
| AML | 5 | *BCOR* | 0.668 | – | 0.107 | 651 | 2 | suite_bulk_oof_hardened |
| AML | 5 | *BCORL1* | 0.455 | – | 0.019 | 651 | 2 | suite_bulk_oof_hardened |
| AML | 5 | *CBL* | 0.490 | – | 0.026 | 651 | 2 | suite_bulk_oof_hardened |
| AML | 5 | *CEBPA* | 0.664 | – | 0.091 | 651 | 2 | suite_bulk_oof_hardened |
| AML | 5 | *DNMT3A* | 0.628 | – | 0.291 | 651 | 2 | suite_bulk_oof_hardened |
| AML | 5 | *ETV6* | 0.772 | – | 0.013 | 326 | 2 | suite_bulk_oof_hardened |
| AML | 5 | *EZH2* | 0.386 | – | 0.030 | 651 | 2 | suite_bulk_oof_hardened |
| AML | 5 | *FLT3* | 0.674 | – | 0.205 | 651 | 2 | suite_bulk_oof_hardened |
| AML | 5 | *GATA2* | 0.703 | – | 0.127 | 651 | 2 | suite_bulk_oof_hardened |
| AML | 5 | *IDH1* | 0.612 | – | 0.108 | 651 | 2 | suite_bulk_oof_hardened |
| AML | 5 | *IDH2* | 0.545 | – | 0.127 | 651 | 2 | suite_bulk_oof_hardened |
| AML | 5 | *KIT* | 0.414 | – | 0.021 | 651 | 2 | suite_bulk_oof_hardened |
| AML | 5 | *KRAS* | 0.683 | – | 0.143 | 651 | 2 | suite_bulk_oof_hardened |
| AML | 5 | *NPM1* | 0.801 | – | 0.458 | 651 | 2 | suite_bulk_oof_hardened |
| AML | 5 | *NRAS* | 0.616 | – | 0.195 | 651 | 2 | suite_bulk_oof_hardened |
| AML | 5 | *PHF6* | 0.562 | – | 0.039 | 651 | 2 | suite_bulk_oof_hardened |
| AML | 5 | *PTPN11* | 0.584 | – | 0.075 | 651 | 2 | suite_bulk_oof_hardened |
| AML | 5 | *RAD21* | 0.520 | – | 0.026 | 651 | 2 | suite_bulk_oof_hardened |
| AML | 5 | *RUNX1* | 0.668 | – | 0.171 | 651 | 2 | suite_bulk_oof_hardened |
| AML | 5 | *SF3B1* | 0.535 | – | 0.055 | 651 | 2 | suite_bulk_oof_hardened |
| AML | 5 | *SMC1A* | 0.799 | – | 0.023 | 326 | 2 | suite_bulk_oof_hardened |
| AML | 5 | *SMC3* | 0.816 | – | 0.062 | 651 | 2 | suite_bulk_oof_hardened |
| AML | 5 | *SRSF2* | 0.645 | – | 0.169 | 651 | 2 | suite_bulk_oof_hardened |
| AML | 5 | *STAG2* | 0.499 | – | 0.067 | 651 | 2 | suite_bulk_oof_hardened |
| AML | 5 | *TET2* | 0.604 | – | 0.171 | 651 | 2 | suite_bulk_oof_hardened |
| AML | 5 | *TP53* | 0.724 | – | 0.166 | 651 | 2 | suite_bulk_oof_hardened |
| AML | 5 | *U2AF1* | 0.396 | – | 0.039 | 651 | 2 | suite_bulk_oof_hardened |
| AML | 5 | *WT1* | 0.568 | – | 0.085 | 651 | 2 | suite_bulk_oof_hardened |
| AML | 5 | *ZRSR2* | 0.542 | – | 0.007 | 326 | 2 | suite_bulk_oof_hardened |
| AML | 10 | *ASXL1* | 0.723 | – | 0.219 | 651 | 2 | suite_bulk_oof_hardened |
| AML | 10 | *BCOR* | 0.691 | – | 0.109 | 651 | 2 | suite_bulk_oof_hardened |
| AML | 10 | *BCORL1* | 0.708 | – | 0.048 | 651 | 2 | suite_bulk_oof_hardened |
| AML | 10 | *CBL* | 0.452 | – | 0.021 | 651 | 2 | suite_bulk_oof_hardened |
| AML | 10 | *CEBPA* | 0.733 | – | 0.321 | 651 | 2 | suite_bulk_oof_hardened |
| AML | 10 | *DNMT3A* | 0.680 | – | 0.350 | 651 | 2 | suite_bulk_oof_hardened |
| AML | 10 | *ETV6* | 0.877 | – | 0.024 | 326 | 2 | suite_bulk_oof_hardened |
| AML | 10 | *EZH2* | 0.427 | – | 0.026 | 651 | 2 | suite_bulk_oof_hardened |
| AML | 10 | *FLT3* | 0.662 | – | 0.224 | 651 | 2 | suite_bulk_oof_hardened |
| AML | 10 | *GATA2* | 0.724 | – | 0.222 | 651 | 2 | suite_bulk_oof_hardened |
| AML | 10 | *IDH1* | 0.694 | – | 0.130 | 651 | 2 | suite_bulk_oof_hardened |
| AML | 10 | *IDH2* | 0.703 | – | 0.232 | 651 | 2 | suite_bulk_oof_hardened |
| AML | 10 | *KIT* | 0.627 | – | 0.045 | 651 | 2 | suite_bulk_oof_hardened |
| AML | 10 | *KRAS* | 0.723 | – | 0.157 | 651 | 2 | suite_bulk_oof_hardened |
| AML | 10 | *NPM1* | 0.915 | – | 0.736 | 651 | 2 | suite_bulk_oof_hardened |
| AML | 10 | *NRAS* | 0.668 | – | 0.236 | 651 | 2 | suite_bulk_oof_hardened |
| AML | 10 | *PHF6* | 0.494 | – | 0.035 | 651 | 2 | suite_bulk_oof_hardened |
| AML | 10 | *PTPN11* | 0.601 | – | 0.084 | 651 | 2 | suite_bulk_oof_hardened |
| AML | 10 | *RAD21* | 0.631 | – | 0.039 | 651 | 2 | suite_bulk_oof_hardened |
| AML | 10 | *RUNX1* | 0.835 | – | 0.440 | 651 | 2 | suite_bulk_oof_hardened |
| AML | 10 | *SF3B1* | 0.686 | – | 0.101 | 651 | 2 | suite_bulk_oof_hardened |
| AML | 10 | *SMC1A* | 0.755 | – | 0.032 | 326 | 2 | suite_bulk_oof_hardened |
| AML | 10 | *SMC3* | 0.754 | – | 0.074 | 651 | 2 | suite_bulk_oof_hardened |
| AML | 10 | *SRSF2* | 0.711 | – | 0.250 | 651 | 2 | suite_bulk_oof_hardened |
| AML | 10 | *STAG2* | 0.587 | – | 0.085 | 651 | 2 | suite_bulk_oof_hardened |
| AML | 10 | *TET2* | 0.660 | – | 0.256 | 651 | 2 | suite_bulk_oof_hardened |
| AML | 10 | *TP53* | 0.829 | – | 0.454 | 651 | 2 | suite_bulk_oof_hardened |
| AML | 10 | *U2AF1* | 0.543 | – | 0.061 | 651 | 2 | suite_bulk_oof_hardened |
| AML | 10 | *WT1* | 0.615 | – | 0.119 | 651 | 2 | suite_bulk_oof_hardened |
| AML | 10 | *ZRSR2* | 0.535 | – | 0.007 | 326 | 2 | suite_bulk_oof_hardened |
| AML | 25 | *ASXL1* | 0.807 | – | 0.369 | 651 | 2 | suite_bulk_oof_hardened |
| AML | 25 | *BCOR* | 0.651 | – | 0.107 | 651 | 2 | suite_bulk_oof_hardened |
| AML | 25 | *BCORL1* | 0.771 | – | 0.112 | 651 | 2 | suite_bulk_oof_hardened |
| AML | 25 | *CBL* | 0.480 | – | 0.026 | 651 | 2 | suite_bulk_oof_hardened |
| AML | 25 | *CEBPA* | 0.913 | – | 0.612 | 651 | 2 | suite_bulk_oof_hardened |
| AML | 25 | *DNMT3A* | 0.786 | – | 0.517 | 651 | 2 | suite_bulk_oof_hardened |
| AML | 25 | *ETV6* | 0.969 | – | 0.091 | 326 | 2 | suite_bulk_oof_hardened |
| AML | 25 | *EZH2* | 0.522 | – | 0.035 | 651 | 2 | suite_bulk_oof_hardened |
| AML | 25 | *FLT3* | 0.681 | – | 0.195 | 651 | 2 | suite_bulk_oof_hardened |

Continued on next page

*Supplementary Table S9 continued*

| Cancer | $k$ | Driver | AUROC mean | AUROC SD | AUPRC mean | Samples | CV folds | Implementation |
|---|---|---|---|---|---|---|---|---|
| AML | 25 | *GATA2* | 0.727 | – | 0.274 | 651 | 2 | suite_bulk_oof_hardened |
| AML | 25 | *IDH1* | 0.898 | – | 0.363 | 651 | 2 | suite_bulk_oof_hardened |
| AML | 25 | *IDH2* | 0.862 | – | 0.479 | 651 | 2 | suite_bulk_oof_hardened |
| AML | 25 | *KIT* | 0.802 | – | 0.134 | 651 | 2 | suite_bulk_oof_hardened |
| AML | 25 | *KRAS* | 0.677 | – | 0.185 | 651 | 2 | suite_bulk_oof_hardened |
| AML | 25 | *NPM1* | 0.966 | – | 0.859 | 651 | 2 | suite_bulk_oof_hardened |
| AML | 25 | *NRAS* | 0.680 | – | 0.245 | 651 | 2 | suite_bulk_oof_hardened |
| AML | 25 | *PHF6* | 0.599 | – | 0.056 | 651 | 2 | suite_bulk_oof_hardened |
| AML | 25 | *PTPN11* | 0.695 | – | 0.111 | 651 | 2 | suite_bulk_oof_hardened |
| AML | 25 | *RAD21* | 0.588 | – | 0.030 | 651 | 2 | suite_bulk_oof_hardened |
| AML | 25 | *RUNX1* | 0.888 | – | 0.543 | 651 | 2 | suite_bulk_oof_hardened |
| AML | 25 | *SF3B1* | 0.761 | – | 0.267 | 651 | 2 | suite_bulk_oof_hardened |
| AML | 25 | *SMC1A* | 0.969 | – | 0.545 | 326 | 2 | suite_bulk_oof_hardened |
| AML | 25 | *SMC3* | 0.770 | – | 0.110 | 651 | 2 | suite_bulk_oof_hardened |
| AML | 25 | *SRSF2* | 0.944 | – | 0.702 | 651 | 2 | suite_bulk_oof_hardened |
| AML | 25 | *STAG2* | 0.768 | – | 0.257 | 651 | 2 | suite_bulk_oof_hardened |
| AML | 25 | *TET2* | 0.764 | – | 0.372 | 651 | 2 | suite_bulk_oof_hardened |
| AML | 25 | *TP53* | 0.860 | – | 0.579 | 651 | 2 | suite_bulk_oof_hardened |
| AML | 25 | *U2AF1* | 0.619 | – | 0.085 | 651 | 2 | suite_bulk_oof_hardened |
| AML | 25 | *WT1* | 0.677 | – | 0.155 | 651 | 2 | suite_bulk_oof_hardened |
| AML | 25 | *ZRSR2* | 0.640 | – | 0.008 | 326 | 2 | suite_bulk_oof_hardened |
| AML | 50 | *ASXL1* | 0.858 | – | 0.415 | 651 | 2 | suite_bulk_oof_hardened |
| AML | 50 | *BCOR* | 0.720 | – | 0.201 | 651 | 2 | suite_bulk_oof_hardened |
| AML | 50 | *BCORL1* | 0.852 | – | 0.280 | 651 | 2 | suite_bulk_oof_hardened |
| AML | 50 | *CBL* | 0.478 | – | 0.109 | 651 | 2 | suite_bulk_oof_hardened |
| AML | 50 | *CEBPA* | 0.875 | – | 0.598 | 651 | 2 | suite_bulk_oof_hardened |
| AML | 50 | *DNMT3A* | 0.775 | – | 0.542 | 651 | 2 | suite_bulk_oof_hardened |
| AML | 50 | *ETV6* | 0.978 | – | 0.125 | 326 | 2 | suite_bulk_oof_hardened |
| AML | 50 | *EZH2* | 0.623 | – | 0.169 | 651 | 2 | suite_bulk_oof_hardened |
| AML | 50 | *FLT3* | 0.712 | – | 0.242 | 651 | 2 | suite_bulk_oof_hardened |
| AML | 50 | *GATA2* | 0.806 | – | 0.444 | 651 | 2 | suite_bulk_oof_hardened |
| AML | 50 | *IDH1* | 0.881 | – | 0.395 | 651 | 2 | suite_bulk_oof_hardened |
| AML | 50 | *IDH2* | 0.860 | – | 0.457 | 651 | 2 | suite_bulk_oof_hardened |
| AML | 50 | *KIT* | 0.856 | – | 0.170 | 651 | 2 | suite_bulk_oof_hardened |
| AML | 50 | *KRAS* | 0.700 | – | 0.129 | 651 | 2 | suite_bulk_oof_hardened |
| AML | 50 | *NPM1* | 0.969 | – | 0.862 | 651 | 2 | suite_bulk_oof_hardened |
| AML | 50 | *NRAS* | 0.686 | – | 0.247 | 651 | 2 | suite_bulk_oof_hardened |
| AML | 50 | *PHF6* | 0.709 | – | 0.113 | 651 | 2 | suite_bulk_oof_hardened |
| AML | 50 | *PTPN11* | 0.715 | – | 0.120 | 651 | 2 | suite_bulk_oof_hardened |
| AML | 50 | *RAD21* | 0.669 | – | 0.067 | 651 | 2 | suite_bulk_oof_hardened |
| AML | 50 | *RUNX1* | 0.907 | – | 0.572 | 651 | 2 | suite_bulk_oof_hardened |
| AML | 50 | *SF3B1* | 0.920 | – | 0.669 | 651 | 2 | suite_bulk_oof_hardened |
| AML | 50 | *SMC1A* | 0.980 | – | 0.567 | 326 | 2 | suite_bulk_oof_hardened |
| AML | 50 | *SMC3* | 0.799 | – | 0.076 | 651 | 2 | suite_bulk_oof_hardened |
| AML | 50 | *SRSF2* | 0.994 | – | 0.940 | 651 | 2 | suite_bulk_oof_hardened |
| AML | 50 | *STAG2* | 0.829 | – | 0.398 | 651 | 2 | suite_bulk_oof_hardened |
| AML | 50 | *TET2* | 0.738 | – | 0.333 | 651 | 2 | suite_bulk_oof_hardened |
| AML | 50 | *TP53* | 0.853 | – | 0.604 | 651 | 2 | suite_bulk_oof_hardened |
| AML | 50 | *U2AF1* | 0.869 | – | 0.434 | 651 | 2 | suite_bulk_oof_hardened |
| AML | 50 | *WT1* | 0.757 | – | 0.255 | 651 | 2 | suite_bulk_oof_hardened |
| AML | 50 | *ZRSR2* | 0.523 | – | 0.006 | 326 | 2 | suite_bulk_oof_hardened |
| AML | 75 | *ASXL1* | 0.871 | – | 0.474 | 651 | 2 | suite_bulk_oof_hardened |
| AML | 75 | *BCOR* | 0.819 | – | 0.335 | 651 | 2 | suite_bulk_oof_hardened |
| AML | 75 | *BCORL1* | 0.845 | – | 0.335 | 651 | 2 | suite_bulk_oof_hardened |
| AML | 75 | *CBL* | 0.515 | – | 0.180 | 651 | 2 | suite_bulk_oof_hardened |
| AML | 75 | *CEBPA* | 0.891 | – | 0.611 | 651 | 2 | suite_bulk_oof_hardened |
| AML | 75 | *DNMT3A* | 0.791 | – | 0.576 | 651 | 2 | suite_bulk_oof_hardened |
| AML | 75 | *ETV6* | 0.945 | – | 0.053 | 326 | 2 | suite_bulk_oof_hardened |
| AML | 75 | *EZH2* | 0.679 | – | 0.167 | 651 | 2 | suite_bulk_oof_hardened |
| AML | 75 | *FLT3* | 0.720 | – | 0.260 | 651 | 2 | suite_bulk_oof_hardened |
| AML | 75 | *GATA2* | 0.870 | – | 0.493 | 651 | 2 | suite_bulk_oof_hardened |
| AML | 75 | *IDH1* | 0.882 | – | 0.437 | 651 | 2 | suite_bulk_oof_hardened |
| AML | 75 | *IDH2* | 0.889 | – | 0.471 | 651 | 2 | suite_bulk_oof_hardened |
| AML | 75 | *KIT* | 0.842 | – | 0.275 | 651 | 2 | suite_bulk_oof_hardened |
| AML | 75 | *KRAS* | 0.756 | – | 0.159 | 651 | 2 | suite_bulk_oof_hardened |
| AML | 75 | *NPM1* | 0.970 | – | 0.893 | 651 | 2 | suite_bulk_oof_hardened |
| AML | 75 | *NRAS* | 0.706 | – | 0.256 | 651 | 2 | suite_bulk_oof_hardened |
| AML | 75 | *PHF6* | 0.665 | – | 0.081 | 651 | 2 | suite_bulk_oof_hardened |
| AML | 75 | *PTPN11* | 0.762 | – | 0.167 | 651 | 2 | suite_bulk_oof_hardened |
| AML | 75 | *RAD21* | 0.722 | – | 0.066 | 651 | 2 | suite_bulk_oof_hardened |
| AML | 75 | *RUNX1* | 0.899 | – | 0.562 | 651 | 2 | suite_bulk_oof_hardened |
| AML | 75 | *SF3B1* | 0.964 | – | 0.783 | 651 | 2 | suite_bulk_oof_hardened |
| AML | 75 | *SMC1A* | 0.983 | – | 0.333 | 326 | 2 | suite_bulk_oof_hardened |

Continued on next page

*Supplementary Table S9 continued*

| Cancer | $k$ | Driver | AUROC mean | AUROC SD | AUPRC mean | Samples | CV folds | Implementation |
|---|---|---|---|---|---|---|---|---|
| AML | 75 | *SMC3* | 0.808 | – | 0.061 | 651 | 2 | suite_bulk_oof_hardened |
| AML | 75 | *SRSF2* | 0.996 | – | 0.950 | 651 | 2 | suite_bulk_oof_hardened |
| AML | 75 | *STAG2* | 0.831 | – | 0.486 | 651 | 2 | suite_bulk_oof_hardened |
| AML | 75 | *TET2* | 0.741 | – | 0.392 | 651 | 2 | suite_bulk_oof_hardened |
| AML | 75 | *TP53* | 0.841 | – | 0.593 | 651 | 2 | suite_bulk_oof_hardened |
| AML | 75 | *U2AF1* | 0.945 | – | 0.723 | 651 | 2 | suite_bulk_oof_hardened |
| AML | 75 | *WT1* | 0.801 | – | 0.291 | 651 | 2 | suite_bulk_oof_hardened |
| AML | 75 | *ZRSR2* | 0.572 | – | 0.007 | 326 | 2 | suite_bulk_oof_hardened |
| AML | 100 | *ASXL1* | 0.861 | – | 0.476 | 651 | 2 | suite_bulk_oof_hardened |
| AML | 100 | *BCOR* | 0.862 | – | 0.415 | 651 | 2 | suite_bulk_oof_hardened |
| AML | 100 | *BCORL1* | 0.873 | – | 0.412 | 651 | 2 | suite_bulk_oof_hardened |
| AML | 100 | *CBL* | 0.526 | – | 0.308 | 651 | 2 | suite_bulk_oof_hardened |
| AML | 100 | *CEBPA* | 0.912 | – | 0.653 | 651 | 2 | suite_bulk_oof_hardened |
| AML | 100 | *DNMT3A* | 0.795 | – | 0.600 | 651 | 2 | suite_bulk_oof_hardened |
| AML | 100 | *ETV6* | 0.908 | – | 0.032 | 326 | 2 | suite_bulk_oof_hardened |
| AML | 100 | *EZH2* | 0.732 | – | 0.228 | 651 | 2 | suite_bulk_oof_hardened |
| AML | 100 | *FLT3* | 0.709 | – | 0.245 | 651 | 2 | suite_bulk_oof_hardened |
| AML | 100 | *GATA2* | 0.882 | – | 0.485 | 651 | 2 | suite_bulk_oof_hardened |
| AML | 100 | *IDH1* | 0.899 | – | 0.469 | 651 | 2 | suite_bulk_oof_hardened |
| AML | 100 | *IDH2* | 0.886 | – | 0.478 | 651 | 2 | suite_bulk_oof_hardened |
| AML | 100 | *KIT* | 0.847 | – | 0.379 | 651 | 2 | suite_bulk_oof_hardened |
| AML | 100 | *KRAS* | 0.761 | – | 0.169 | 651 | 2 | suite_bulk_oof_hardened |
| AML | 100 | *NPM1* | 0.970 | – | 0.884 | 651 | 2 | suite_bulk_oof_hardened |
| AML | 100 | *NRAS* | 0.687 | – | 0.273 | 651 | 2 | suite_bulk_oof_hardened |
| AML | 100 | *PHF6* | 0.732 | – | 0.126 | 651 | 2 | suite_bulk_oof_hardened |
| AML | 100 | *PTPN11* | 0.736 | – | 0.166 | 651 | 2 | suite_bulk_oof_hardened |
| AML | 100 | *RAD21* | 0.734 | – | 0.072 | 651 | 2 | suite_bulk_oof_hardened |
| AML | 100 | *RUNX1* | 0.908 | – | 0.559 | 651 | 2 | suite_bulk_oof_hardened |
| AML | 100 | *SF3B1* | 0.982 | – | 0.832 | 651 | 2 | suite_bulk_oof_hardened |
| AML | 100 | *SMC1A* | 0.944 | – | 0.277 | 326 | 2 | suite_bulk_oof_hardened |
| AML | 100 | *SMC3* | 0.784 | – | 0.071 | 651 | 2 | suite_bulk_oof_hardened |
| AML | 100 | *SRSF2* | 0.996 | – | 0.945 | 651 | 2 | suite_bulk_oof_hardened |
| AML | 100 | *STAG2* | 0.851 | – | 0.557 | 651 | 2 | suite_bulk_oof_hardened |
| AML | 100 | *TET2* | 0.758 | – | 0.421 | 651 | 2 | suite_bulk_oof_hardened |
| AML | 100 | *TP53* | 0.873 | – | 0.672 | 651 | 2 | suite_bulk_oof_hardened |
| AML | 100 | *U2AF1* | 0.968 | – | 0.860 | 651 | 2 | suite_bulk_oof_hardened |
| AML | 100 | *WT1* | 0.803 | – | 0.290 | 651 | 2 | suite_bulk_oof_hardened |
| AML | 100 | *ZRSR2* | 0.698 | – | 0.010 | 326 | 2 | suite_bulk_oof_hardened |
| AML | 150 | *ASXL1* | 0.896 | – | 0.565 | 651 | 2 | suite_bulk_oof_hardened |
| AML | 150 | *BCOR* | 0.907 | – | 0.491 | 651 | 2 | suite_bulk_oof_hardened |
| AML | 150 | *BCORL1* | 0.880 | – | 0.443 | 651 | 2 | suite_bulk_oof_hardened |
| AML | 150 | *CBL* | 0.552 | – | 0.355 | 651 | 2 | suite_bulk_oof_hardened |
| AML | 150 | *CEBPA* | 0.949 | – | 0.695 | 651 | 2 | suite_bulk_oof_hardened |
| AML | 150 | *DNMT3A* | 0.801 | – | 0.619 | 651 | 2 | suite_bulk_oof_hardened |
| AML | 150 | *ETV6* | 0.929 | – | 0.042 | 326 | 2 | suite_bulk_oof_hardened |
| AML | 150 | *EZH2* | 0.691 | – | 0.288 | 651 | 2 | suite_bulk_oof_hardened |
| AML | 150 | *FLT3* | 0.717 | – | 0.294 | 651 | 2 | suite_bulk_oof_hardened |
| AML | 150 | *GATA2* | 0.848 | – | 0.445 | 651 | 2 | suite_bulk_oof_hardened |
| AML | 150 | *IDH1* | 0.919 | – | 0.537 | 651 | 2 | suite_bulk_oof_hardened |
| AML | 150 | *IDH2* | 0.898 | – | 0.557 | 651 | 2 | suite_bulk_oof_hardened |
| AML | 150 | *KIT* | 0.882 | – | 0.423 | 651 | 2 | suite_bulk_oof_hardened |
| AML | 150 | *KRAS* | 0.794 | – | 0.235 | 651 | 2 | suite_bulk_oof_hardened |
| AML | 150 | *NPM1* | 0.977 | – | 0.909 | 651 | 2 | suite_bulk_oof_hardened |
| AML | 150 | *NRAS* | 0.690 | – | 0.315 | 651 | 2 | suite_bulk_oof_hardened |
| AML | 150 | *PHF6* | 0.840 | – | 0.147 | 651 | 2 | suite_bulk_oof_hardened |
| AML | 150 | *PTPN11* | 0.741 | – | 0.194 | 651 | 2 | suite_bulk_oof_hardened |
| AML | 150 | *RAD21* | 0.750 | – | 0.082 | 651 | 2 | suite_bulk_oof_hardened |
| AML | 150 | *RUNX1* | 0.924 | – | 0.622 | 651 | 2 | suite_bulk_oof_hardened |
| AML | 150 | *SF3B1* | 0.995 | – | 0.911 | 651 | 2 | suite_bulk_oof_hardened |
| AML | 150 | *SMC1A* | 0.949 | – | 0.197 | 326 | 2 | suite_bulk_oof_hardened |
| AML | 150 | *SMC3* | 0.829 | – | 0.061 | 651 | 2 | suite_bulk_oof_hardened |
| AML | 150 | *SRSF2* | 0.996 | – | 0.947 | 651 | 2 | suite_bulk_oof_hardened |
| AML | 150 | *STAG2* | 0.870 | – | 0.657 | 651 | 2 | suite_bulk_oof_hardened |
| AML | 150 | *TET2* | 0.803 | – | 0.460 | 651 | 2 | suite_bulk_oof_hardened |
| AML | 150 | *TP53* | 0.889 | – | 0.683 | 651 | 2 | suite_bulk_oof_hardened |
| AML | 150 | *U2AF1* | 0.979 | – | 0.918 | 651 | 2 | suite_bulk_oof_hardened |
| AML | 150 | *WT1* | 0.814 | – | 0.393 | 651 | 2 | suite_bulk_oof_hardened |
| AML | 150 | *ZRSR2* | 0.938 | – | 0.048 | 326 | 2 | suite_bulk_oof_hardened |
| AML | 250 | *ASXL1* | 0.898 | – | 0.583 | 651 | 2 | suite_bulk_oof_hardened |
| AML | 250 | *BCOR* | 0.938 | – | 0.634 | 651 | 2 | suite_bulk_oof_hardened |
| AML | 250 | *BCORL1* | 0.892 | – | 0.446 | 651 | 2 | suite_bulk_oof_hardened |
| AML | 250 | *CBL* | 0.592 | – | 0.400 | 651 | 2 | suite_bulk_oof_hardened |
| AML | 250 | *CEBPA* | 0.953 | – | 0.721 | 651 | 2 | suite_bulk_oof_hardened |

Continued on next page

*Supplementary Table S9 continued*

| Cancer | $k$ | Driver | AUROC mean | AUROC SD | AUPRC mean | Samples | CV folds | Implementation |
|---|---|---|---|---|---|---|---|---|
| AML | 250 | *DNMT3A* | 0.836 | – | 0.680 | 651 | 2 | suite_bulk_oof_hardened |
| AML | 250 | *ETV6* | 0.855 | – | 0.021 | 326 | 2 | suite_bulk_oof_hardened |
| AML | 250 | *EZH2* | 0.746 | – | 0.303 | 651 | 2 | suite_bulk_oof_hardened |
| AML | 250 | *FLT3* | 0.751 | – | 0.293 | 651 | 2 | suite_bulk_oof_hardened |
| AML | 250 | *GATA2* | 0.854 | – | 0.470 | 651 | 2 | suite_bulk_oof_hardened |
| AML | 250 | *IDH1* | 0.927 | – | 0.557 | 651 | 2 | suite_bulk_oof_hardened |
| AML | 250 | *IDH2* | 0.895 | – | 0.560 | 651 | 2 | suite_bulk_oof_hardened |
| AML | 250 | *KIT* | 0.905 | – | 0.436 | 651 | 2 | suite_bulk_oof_hardened |
| AML | 250 | *KRAS* | 0.772 | – | 0.276 | 651 | 2 | suite_bulk_oof_hardened |
| AML | 250 | *NPM1* | 0.978 | – | 0.897 | 651 | 2 | suite_bulk_oof_hardened |
| AML | 250 | *NRAS* | 0.728 | – | 0.320 | 651 | 2 | suite_bulk_oof_hardened |
| AML | 250 | *PHF6* | 0.906 | – | 0.262 | 651 | 2 | suite_bulk_oof_hardened |
| AML | 250 | *PTPN11* | 0.793 | – | 0.245 | 651 | 2 | suite_bulk_oof_hardened |
| AML | 250 | *RAD21* | 0.796 | – | 0.097 | 651 | 2 | suite_bulk_oof_hardened |
| AML | 250 | *RUNX1* | 0.941 | – | 0.681 | 651 | 2 | suite_bulk_oof_hardened |
| AML | 250 | *SF3B1* | 0.997 | – | 0.950 | 651 | 2 | suite_bulk_oof_hardened |
| AML | 250 | *SMC1A* | 0.968 | – | 0.125 | 326 | 2 | suite_bulk_oof_hardened |
| AML | 250 | *SMC3* | 0.822 | – | 0.054 | 651 | 2 | suite_bulk_oof_hardened |
| AML | 250 | *SRSF2* | 0.997 | – | 0.955 | 651 | 2 | suite_bulk_oof_hardened |
| AML | 250 | *STAG2* | 0.889 | – | 0.684 | 651 | 2 | suite_bulk_oof_hardened |
| AML | 250 | *TET2* | 0.839 | – | 0.542 | 651 | 2 | suite_bulk_oof_hardened |
| AML | 250 | *TP53* | 0.919 | – | 0.734 | 651 | 2 | suite_bulk_oof_hardened |
| AML | 250 | *U2AF1* | 0.979 | – | 0.933 | 651 | 2 | suite_bulk_oof_hardened |
| AML | 250 | *WT1* | 0.815 | – | 0.440 | 651 | 2 | suite_bulk_oof_hardened |
| AML | 250 | *ZRSR2* | 0.938 | – | 0.048 | 326 | 2 | suite_bulk_oof_hardened |
| AML | 500 | *ASXL1* | 0.896 | – | 0.603 | 651 | 2 | suite_bulk_oof_hardened |
| AML | 500 | *BCOR* | 0.934 | – | 0.641 | 651 | 2 | suite_bulk_oof_hardened |
| AML | 500 | *BCORL1* | 0.894 | – | 0.506 | 651 | 2 | suite_bulk_oof_hardened |
| AML | 500 | *CBL* | 0.605 | – | 0.403 | 651 | 2 | suite_bulk_oof_hardened |
| AML | 500 | *CEBPA* | 0.950 | – | 0.734 | 651 | 2 | suite_bulk_oof_hardened |
| AML | 500 | *DNMT3A* | 0.832 | – | 0.688 | 651 | 2 | suite_bulk_oof_hardened |
| AML | 500 | *ETV6* | 0.846 | – | 0.020 | 326 | 2 | suite_bulk_oof_hardened |
| AML | 500 | *EZH2* | 0.758 | – | 0.312 | 651 | 2 | suite_bulk_oof_hardened |
| AML | 500 | *FLT3* | 0.756 | – | 0.311 | 651 | 2 | suite_bulk_oof_hardened |
| AML | 500 | *GATA2* | 0.870 | – | 0.500 | 651 | 2 | suite_bulk_oof_hardened |
| AML | 500 | *IDH1* | 0.924 | – | 0.539 | 651 | 2 | suite_bulk_oof_hardened |
| AML | 500 | *IDH2* | 0.908 | – | 0.579 | 651 | 2 | suite_bulk_oof_hardened |
| AML | 500 | *KIT* | 0.898 | – | 0.456 | 651 | 2 | suite_bulk_oof_hardened |
| AML | 500 | *KRAS* | 0.754 | – | 0.235 | 651 | 2 | suite_bulk_oof_hardened |
| AML | 500 | *NPM1* | 0.979 | – | 0.896 | 651 | 2 | suite_bulk_oof_hardened |
| AML | 500 | *NRAS* | 0.749 | – | 0.339 | 651 | 2 | suite_bulk_oof_hardened |
| AML | 500 | *PHF6* | 0.907 | – | 0.230 | 651 | 2 | suite_bulk_oof_hardened |
| AML | 500 | *PTPN11* | 0.814 | – | 0.234 | 651 | 2 | suite_bulk_oof_hardened |
| AML | 500 | *RAD21* | 0.757 | – | 0.096 | 651 | 2 | suite_bulk_oof_hardened |
| AML | 500 | *RUNX1* | 0.941 | – | 0.691 | 651 | 2 | suite_bulk_oof_hardened |
| AML | 500 | *SF3B1* | 0.997 | – | 0.948 | 651 | 2 | suite_bulk_oof_hardened |
| AML | 500 | *SMC1A* | 0.954 | – | 0.110 | 326 | 2 | suite_bulk_oof_hardened |
| AML | 500 | *SMC3* | 0.819 | – | 0.070 | 651 | 2 | suite_bulk_oof_hardened |
| AML | 500 | *SRSF2* | 0.997 | – | 0.952 | 651 | 2 | suite_bulk_oof_hardened |
| AML | 500 | *STAG2* | 0.898 | – | 0.691 | 651 | 2 | suite_bulk_oof_hardened |
| AML | 500 | *TET2* | 0.849 | – | 0.582 | 651 | 2 | suite_bulk_oof_hardened |
| AML | 500 | *TP53* | 0.914 | – | 0.727 | 651 | 2 | suite_bulk_oof_hardened |
| AML | 500 | *U2AF1* | 0.981 | – | 0.939 | 651 | 2 | suite_bulk_oof_hardened |
| AML | 500 | *WT1* | 0.853 | – | 0.495 | 651 | 2 | suite_bulk_oof_hardened |
| AML | 500 | *ZRSR2* | 0.914 | – | 0.034 | 326 | 2 | suite_bulk_oof_hardened |
| BRCA | 5 | *AKT1* | 0.752 | – | 0.062 | 903 | 5 | suite_bulk_oof_hardened |
| BRCA | 5 | *ARID1A* | 0.621 | – | 0.040 | 903 | 5 | suite_bulk_oof_hardened |
| BRCA | 5 | *ATM* | 0.586 | – | 0.024 | 903 | 5 | suite_bulk_oof_hardened |
| BRCA | 5 | *BRCA1* | 0.721 | – | 0.052 | 903 | 5 | suite_bulk_oof_hardened |
| BRCA | 5 | *BRCA2* | 0.623 | – | 0.037 | 903 | 5 | suite_bulk_oof_hardened |
| BRCA | 5 | *CDH1* | 0.826 | – | 0.369 | 903 | 5 | suite_bulk_oof_hardened |
| BRCA | 5 | *ERBB2* | 0.680 | – | 0.045 | 903 | 5 | suite_bulk_oof_hardened |
| BRCA | 5 | *ERBB3* | 0.732 | – | 0.113 | 903 | 5 | suite_bulk_oof_hardened |
| BRCA | 5 | *FOXA1* | 0.638 | – | 0.043 | 903 | 5 | suite_bulk_oof_hardened |
| BRCA | 5 | *GATA3* | 0.737 | – | 0.182 | 903 | 5 | suite_bulk_oof_hardened |
| BRCA | 5 | *KMT2C* | 0.635 | – | 0.132 | 903 | 5 | suite_bulk_oof_hardened |
| BRCA | 5 | *MAP3K1* | 0.690 | – | 0.139 | 903 | 5 | suite_bulk_oof_hardened |
| BRCA | 5 | *PIK3CA* | 0.764 | – | 0.550 | 903 | 5 | suite_bulk_oof_hardened |
| BRCA | 5 | *PIK3R1* | 0.626 | – | 0.038 | 903 | 5 | suite_bulk_oof_hardened |
| BRCA | 5 | *PTEN* | 0.592 | – | 0.072 | 903 | 5 | suite_bulk_oof_hardened |
| BRCA | 5 | *RB1* | 0.694 | – | 0.068 | 903 | 5 | suite_bulk_oof_hardened |
| BRCA | 5 | *RUNX1* | 0.565 | – | 0.041 | 903 | 5 | suite_bulk_oof_hardened |
| BRCA | 5 | *TP53* | 0.919 | – | 0.804 | 903 | 5 | suite_bulk_oof_hardened |

Continued on next page

*Supplementary Table S9 continued*

| Cancer | $k$ | Driver | AUROC mean | AUROC SD | AUPRC mean | Samples | CV folds | Implementation |
|---|---|---|---|---|---|---|---|---|
| BRCA | 10 | *AKT1* | 0.799 | – | 0.079 | 903 | 5 | suite_bulk_oof_hardened |
| BRCA | 10 | *ARID1A* | 0.654 | – | 0.050 | 903 | 5 | suite_bulk_oof_hardened |
| BRCA | 10 | *ATM* | 0.610 | – | 0.033 | 903 | 5 | suite_bulk_oof_hardened |
| BRCA | 10 | *BRCA1* | 0.759 | – | 0.061 | 903 | 5 | suite_bulk_oof_hardened |
| BRCA | 10 | *BRCA2* | 0.551 | – | 0.034 | 903 | 5 | suite_bulk_oof_hardened |
| BRCA | 10 | *CDH1* | 0.853 | – | 0.391 | 903 | 5 | suite_bulk_oof_hardened |
| BRCA | 10 | *ERBB2* | 0.733 | – | 0.107 | 903 | 5 | suite_bulk_oof_hardened |
| BRCA | 10 | *ERBB3* | 0.650 | – | 0.040 | 903 | 5 | suite_bulk_oof_hardened |
| BRCA | 10 | *FOXA1* | 0.652 | – | 0.049 | 903 | 5 | suite_bulk_oof_hardened |
| BRCA | 10 | *GATA3* | 0.728 | – | 0.172 | 903 | 5 | suite_bulk_oof_hardened |
| BRCA | 10 | *KMT2C* | 0.651 | – | 0.135 | 903 | 5 | suite_bulk_oof_hardened |
| BRCA | 10 | *MAP3K1* | 0.675 | – | 0.129 | 903 | 5 | suite_bulk_oof_hardened |
| BRCA | 10 | *PIK3CA* | 0.816 | – | 0.602 | 903 | 5 | suite_bulk_oof_hardened |
| BRCA | 10 | *PIK3R1* | 0.638 | – | 0.070 | 903 | 5 | suite_bulk_oof_hardened |
| BRCA | 10 | *PTEN* | 0.630 | – | 0.112 | 903 | 5 | suite_bulk_oof_hardened |
| BRCA | 10 | *RB1* | 0.740 | – | 0.063 | 903 | 5 | suite_bulk_oof_hardened |
| BRCA | 10 | *RUNX1* | 0.660 | – | 0.057 | 903 | 5 | suite_bulk_oof_hardened |
| BRCA | 10 | *TP53* | 0.923 | – | 0.811 | 903 | 5 | suite_bulk_oof_hardened |
| BRCA | 25 | *AKT1* | 0.795 | – | 0.098 | 903 | 5 | suite_bulk_oof_hardened |
| BRCA | 25 | *ARID1A* | 0.737 | – | 0.058 | 903 | 5 | suite_bulk_oof_hardened |
| BRCA | 25 | *ATM* | 0.644 | – | 0.033 | 903 | 5 | suite_bulk_oof_hardened |
| BRCA | 25 | *BRCA1* | 0.728 | – | 0.049 | 903 | 5 | suite_bulk_oof_hardened |
| BRCA | 25 | *BRCA2* | 0.591 | – | 0.037 | 903 | 5 | suite_bulk_oof_hardened |
| BRCA | 25 | *CDH1* | 0.890 | – | 0.498 | 903 | 5 | suite_bulk_oof_hardened |
| BRCA | 25 | *ERBB2* | 0.761 | – | 0.168 | 903 | 5 | suite_bulk_oof_hardened |
| BRCA | 25 | *ERBB3* | 0.650 | – | 0.027 | 903 | 5 | suite_bulk_oof_hardened |
| BRCA | 25 | *FOXA1* | 0.724 | – | 0.096 | 903 | 5 | suite_bulk_oof_hardened |
| BRCA | 25 | *GATA3* | 0.846 | – | 0.447 | 903 | 5 | suite_bulk_oof_hardened |
| BRCA | 25 | *KMT2C* | 0.666 | – | 0.145 | 903 | 5 | suite_bulk_oof_hardened |
| BRCA | 25 | *MAP3K1* | 0.734 | – | 0.262 | 903 | 5 | suite_bulk_oof_hardened |
| BRCA | 25 | *PIK3CA* | 0.826 | – | 0.625 | 903 | 5 | suite_bulk_oof_hardened |
| BRCA | 25 | *PIK3R1* | 0.607 | – | 0.045 | 903 | 5 | suite_bulk_oof_hardened |
| BRCA | 25 | *PTEN* | 0.651 | – | 0.100 | 903 | 5 | suite_bulk_oof_hardened |
| BRCA | 25 | *RB1* | 0.734 | – | 0.110 | 903 | 5 | suite_bulk_oof_hardened |
| BRCA | 25 | *RUNX1* | 0.726 | – | 0.140 | 903 | 5 | suite_bulk_oof_hardened |
| BRCA | 25 | *TP53* | 0.928 | – | 0.832 | 903 | 5 | suite_bulk_oof_hardened |
| BRCA | 50 | *AKT1* | 0.802 | – | 0.099 | 903 | 5 | suite_bulk_oof_hardened |
| BRCA | 50 | *ARID1A* | 0.705 | – | 0.056 | 903 | 5 | suite_bulk_oof_hardened |
| BRCA | 50 | *ATM* | 0.588 | – | 0.060 | 903 | 5 | suite_bulk_oof_hardened |
| BRCA | 50 | *BRCA1* | 0.650 | – | 0.045 | 903 | 5 | suite_bulk_oof_hardened |
| BRCA | 50 | *BRCA2* | 0.449 | – | 0.046 | 903 | 5 | suite_bulk_oof_hardened |
| BRCA | 50 | *CDH1* | 0.876 | – | 0.491 | 903 | 5 | suite_bulk_oof_hardened |
| BRCA | 50 | *ERBB2* | 0.750 | – | 0.218 | 903 | 5 | suite_bulk_oof_hardened |
| BRCA | 50 | *ERBB3* | 0.544 | – | 0.018 | 903 | 5 | suite_bulk_oof_hardened |
| BRCA | 50 | *FOXA1* | 0.639 | – | 0.067 | 903 | 5 | suite_bulk_oof_hardened |
| BRCA | 50 | *GATA3* | 0.864 | – | 0.534 | 903 | 5 | suite_bulk_oof_hardened |
| BRCA | 50 | *KMT2C* | 0.628 | – | 0.159 | 903 | 5 | suite_bulk_oof_hardened |
| BRCA | 50 | *MAP3K1* | 0.715 | – | 0.212 | 903 | 5 | suite_bulk_oof_hardened |
| BRCA | 50 | *PIK3CA* | 0.833 | – | 0.633 | 903 | 5 | suite_bulk_oof_hardened |
| BRCA | 50 | *PIK3R1* | 0.643 | – | 0.043 | 903 | 5 | suite_bulk_oof_hardened |
| BRCA | 50 | *PTEN* | 0.658 | – | 0.089 | 903 | 5 | suite_bulk_oof_hardened |
| BRCA | 50 | *RB1* | 0.586 | – | 0.064 | 903 | 5 | suite_bulk_oof_hardened |
| BRCA | 50 | *RUNX1* | 0.683 | – | 0.108 | 903 | 5 | suite_bulk_oof_hardened |
| BRCA | 50 | *TP53* | 0.935 | – | 0.828 | 903 | 5 | suite_bulk_oof_hardened |
| BRCA | 75 | *AKT1* | 0.790 | – | 0.105 | 903 | 5 | suite_bulk_oof_hardened |
| BRCA | 75 | *ARID1A* | 0.708 | – | 0.060 | 903 | 5 | suite_bulk_oof_hardened |
| BRCA | 75 | *ATM* | 0.545 | – | 0.048 | 903 | 5 | suite_bulk_oof_hardened |
| BRCA | 75 | *BRCA1* | 0.603 | – | 0.038 | 903 | 5 | suite_bulk_oof_hardened |
| BRCA | 75 | *BRCA2* | 0.462 | – | 0.087 | 903 | 5 | suite_bulk_oof_hardened |
| BRCA | 75 | *CDH1* | 0.873 | – | 0.463 | 903 | 5 | suite_bulk_oof_hardened |
| BRCA | 75 | *ERBB2* | 0.814 | – | 0.241 | 903 | 5 | suite_bulk_oof_hardened |
| BRCA | 75 | *ERBB3* | 0.667 | – | 0.028 | 903 | 5 | suite_bulk_oof_hardened |
| BRCA | 75 | *FOXA1* | 0.707 | – | 0.076 | 903 | 5 | suite_bulk_oof_hardened |
| BRCA | 75 | *GATA3* | 0.898 | – | 0.573 | 903 | 5 | suite_bulk_oof_hardened |
| BRCA | 75 | *KMT2C* | 0.640 | – | 0.177 | 903 | 5 | suite_bulk_oof_hardened |
| BRCA | 75 | *MAP3K1* | 0.739 | – | 0.246 | 903 | 5 | suite_bulk_oof_hardened |
| BRCA | 75 | *PIK3CA* | 0.830 | – | 0.643 | 903 | 5 | suite_bulk_oof_hardened |
| BRCA | 75 | *PIK3R1* | 0.604 | – | 0.037 | 903 | 5 | suite_bulk_oof_hardened |
| BRCA | 75 | *PTEN* | 0.583 | – | 0.072 | 903 | 5 | suite_bulk_oof_hardened |
| BRCA | 75 | *RB1* | 0.617 | – | 0.071 | 903 | 5 | suite_bulk_oof_hardened |
| BRCA | 75 | *RUNX1* | 0.680 | – | 0.103 | 903 | 5 | suite_bulk_oof_hardened |
| BRCA | 75 | *TP53* | 0.938 | – | 0.851 | 903 | 5 | suite_bulk_oof_hardened |

Continued on next page

| Cancer | *k* | Driver | AUROC mean | AUROC SD | AUPRC mean | Samples | CV folds | Implementation |
|---|---|---|---|---|---|---|---|---|
| BRCA | 100 | *AKT1* | 0.767 | – | 0.174 | 903 | 5 | suite_bulk_oof_hardened |
| BRCA | 100 | *ARID1A* | 0.722 | – | 0.075 | 903 | 5 | suite_bulk_oof_hardened |
| BRCA | 100 | *ATM* | 0.521 | – | 0.028 | 903 | 5 | suite_bulk_oof_hardened |
| BRCA | 100 | *BRCA1* | 0.571 | – | 0.075 | 903 | 5 | suite_bulk_oof_hardened |
| BRCA | 100 | *BRCA2* | 0.463 | – | 0.080 | 903 | 5 | suite_bulk_oof_hardened |
| BRCA | 100 | *CDH1* | 0.871 | – | 0.454 | 903 | 5 | suite_bulk_oof_hardened |
| BRCA | 100 | *ERBB2* | 0.871 | – | 0.310 | 903 | 5 | suite_bulk_oof_hardened |
| BRCA | 100 | *ERBB3* | 0.681 | – | 0.031 | 903 | 5 | suite_bulk_oof_hardened |
| BRCA | 100 | *FOXA1* | 0.739 | – | 0.094 | 903 | 5 | suite_bulk_oof_hardened |
| BRCA | 100 | *GATA3* | 0.898 | – | 0.610 | 903 | 5 | suite_bulk_oof_hardened |
| BRCA | 100 | *KMT2C* | 0.647 | – | 0.173 | 903 | 5 | suite_bulk_oof_hardened |
| BRCA | 100 | *MAP3K1* | 0.725 | – | 0.287 | 903 | 5 | suite_bulk_oof_hardened |
| BRCA | 100 | *PIK3CA* | 0.822 | – | 0.636 | 903 | 5 | suite_bulk_oof_hardened |
| BRCA | 100 | *PIK3R1* | 0.605 | – | 0.043 | 903 | 5 | suite_bulk_oof_hardened |
| BRCA | 100 | *PTEN* | 0.534 | – | 0.066 | 903 | 5 | suite_bulk_oof_hardened |
| BRCA | 100 | *RB1* | 0.585 | – | 0.065 | 903 | 5 | suite_bulk_oof_hardened |
| BRCA | 100 | *RUNX1* | 0.679 | – | 0.121 | 903 | 5 | suite_bulk_oof_hardened |
| BRCA | 100 | *TP53* | 0.939 | – | 0.859 | 903 | 5 | suite_bulk_oof_hardened |
| BRCA | 150 | *AKT1* | 0.797 | – | 0.194 | 903 | 5 | suite_bulk_oof_hardened |
| BRCA | 150 | *ARID1A* | 0.679 | – | 0.060 | 903 | 5 | suite_bulk_oof_hardened |
| BRCA | 150 | *ATM* | 0.556 | – | 0.031 | 903 | 5 | suite_bulk_oof_hardened |
| BRCA | 150 | *BRCA1* | 0.558 | – | 0.050 | 903 | 5 | suite_bulk_oof_hardened |
| BRCA | 150 | *BRCA2* | 0.570 | – | 0.034 | 903 | 5 | suite_bulk_oof_hardened |
| BRCA | 150 | *CDH1* | 0.866 | – | 0.439 | 903 | 5 | suite_bulk_oof_hardened |
| BRCA | 150 | *ERBB2* | 0.886 | – | 0.276 | 903 | 5 | suite_bulk_oof_hardened |
| BRCA | 150 | *ERBB3* | 0.643 | – | 0.031 | 903 | 5 | suite_bulk_oof_hardened |
| BRCA | 150 | *FOXA1* | 0.785 | – | 0.131 | 903 | 5 | suite_bulk_oof_hardened |
| BRCA | 150 | *GATA3* | 0.897 | – | 0.645 | 903 | 5 | suite_bulk_oof_hardened |
| BRCA | 150 | *KMT2C* | 0.651 | – | 0.185 | 903 | 5 | suite_bulk_oof_hardened |
| BRCA | 150 | *MAP3K1* | 0.713 | – | 0.341 | 903 | 5 | suite_bulk_oof_hardened |
| BRCA | 150 | *PIK3CA* | 0.824 | – | 0.643 | 903 | 5 | suite_bulk_oof_hardened |
| BRCA | 150 | *PIK3R1* | 0.593 | – | 0.040 | 903 | 5 | suite_bulk_oof_hardened |
| BRCA | 150 | *PTEN* | 0.527 | – | 0.084 | 903 | 5 | suite_bulk_oof_hardened |
| BRCA | 150 | *RB1* | 0.537 | – | 0.085 | 903 | 5 | suite_bulk_oof_hardened |
| BRCA | 150 | *RUNX1* | 0.689 | – | 0.110 | 903 | 5 | suite_bulk_oof_hardened |
| BRCA | 150 | *TP53* | 0.934 | – | 0.841 | 903 | 5 | suite_bulk_oof_hardened |
| BRCA | 250 | *AKT1* | 0.804 | – | 0.157 | 903 | 5 | suite_bulk_oof_hardened |
| BRCA | 250 | *ARID1A* | 0.632 | – | 0.047 | 903 | 5 | suite_bulk_oof_hardened |
| BRCA | 250 | *ATM* | 0.616 | – | 0.041 | 903 | 5 | suite_bulk_oof_hardened |
| BRCA | 250 | *BRCA1* | 0.514 | – | 0.028 | 903 | 5 | suite_bulk_oof_hardened |
| BRCA | 250 | *BRCA2* | 0.590 | – | 0.052 | 903 | 5 | suite_bulk_oof_hardened |
| BRCA | 250 | *CDH1* | 0.861 | – | 0.447 | 903 | 5 | suite_bulk_oof_hardened |
| BRCA | 250 | *ERBB2* | 0.882 | – | 0.195 | 903 | 5 | suite_bulk_oof_hardened |
| BRCA | 250 | *ERBB3* | 0.607 | – | 0.021 | 903 | 5 | suite_bulk_oof_hardened |
| BRCA | 250 | *FOXA1* | 0.805 | – | 0.221 | 903 | 5 | suite_bulk_oof_hardened |
| BRCA | 250 | *GATA3* | 0.921 | – | 0.669 | 903 | 5 | suite_bulk_oof_hardened |
| BRCA | 250 | *KMT2C* | 0.649 | – | 0.207 | 903 | 5 | suite_bulk_oof_hardened |
| BRCA | 250 | *MAP3K1* | 0.768 | – | 0.413 | 903 | 5 | suite_bulk_oof_hardened |
| BRCA | 250 | *PIK3CA* | 0.813 | – | 0.622 | 903 | 5 | suite_bulk_oof_hardened |
| BRCA | 250 | *PIK3R1* | 0.559 | – | 0.033 | 903 | 5 | suite_bulk_oof_hardened |
| BRCA | 250 | *PTEN* | 0.591 | – | 0.095 | 903 | 5 | suite_bulk_oof_hardened |
| BRCA | 250 | *RB1* | 0.587 | – | 0.085 | 903 | 5 | suite_bulk_oof_hardened |
| BRCA | 250 | *RUNX1* | 0.705 | – | 0.082 | 903 | 5 | suite_bulk_oof_hardened |
| BRCA | 250 | *TP53* | 0.928 | – | 0.824 | 903 | 5 | suite_bulk_oof_hardened |
| BRCA | 500 | *AKT1* | 0.860 | – | 0.165 | 903 | 5 | suite_bulk_oof_hardened |
| BRCA | 500 | *ARID1A* | 0.676 | – | 0.057 | 903 | 5 | suite_bulk_oof_hardened |
| BRCA | 500 | *ATM* | 0.674 | – | 0.032 | 903 | 5 | suite_bulk_oof_hardened |
| BRCA | 500 | *BRCA1* | 0.595 | – | 0.031 | 903 | 5 | suite_bulk_oof_hardened |
| BRCA | 500 | *BRCA2* | 0.627 | – | 0.062 | 903 | 5 | suite_bulk_oof_hardened |
| BRCA | 500 | *CDH1* | 0.877 | – | 0.456 | 903 | 5 | suite_bulk_oof_hardened |
| BRCA | 500 | *ERBB2* | 0.828 | – | 0.167 | 903 | 5 | suite_bulk_oof_hardened |
| BRCA | 500 | *ERBB3* | 0.745 | – | 0.060 | 903 | 5 | suite_bulk_oof_hardened |
| BRCA | 500 | *FOXA1* | 0.863 | – | 0.291 | 903 | 5 | suite_bulk_oof_hardened |
| BRCA | 500 | *GATA3* | 0.936 | – | 0.717 | 903 | 5 | suite_bulk_oof_hardened |
| BRCA | 500 | *KMT2C* | 0.680 | – | 0.184 | 903 | 5 | suite_bulk_oof_hardened |
| BRCA | 500 | *MAP3K1* | 0.769 | – | 0.460 | 903 | 5 | suite_bulk_oof_hardened |
| BRCA | 500 | *PIK3CA* | 0.808 | – | 0.634 | 903 | 5 | suite_bulk_oof_hardened |
| BRCA | 500 | *PIK3R1* | 0.567 | – | 0.030 | 903 | 5 | suite_bulk_oof_hardened |
| BRCA | 500 | *PTEN* | 0.584 | – | 0.080 | 903 | 5 | suite_bulk_oof_hardened |
| BRCA | 500 | *RB1* | 0.631 | – | 0.051 | 903 | 5 | suite_bulk_oof_hardened |
| BRCA | 500 | *RUNX1* | 0.735 | – | 0.105 | 903 | 5 | suite_bulk_oof_hardened |
| BRCA | 500 | *TP53* | 0.940 | – | 0.854 | 903 | 5 | suite_bulk_oof_hardened |

Continued on next page

*Supplementary Table S9 continued*

| Cancer | $k$ | Driver | AUROC mean | AUROC SD | AUPRC mean | Samples | CV folds | Implementation |
|---|---|---|---|---|---|---|---|---|
| CRC | 5 | *APC* | 0.882 | – | 0.922 | 408 | 5 | suite_bulk_oof_hardened |
| CRC | 5 | *ARID1A* | 0.823 | – | 0.423 | 408 | 5 | suite_bulk_oof_hardened |
| CRC | 5 | *BRAF* | 0.863 | – | 0.546 | 408 | 5 | suite_bulk_oof_hardened |
| CRC | 5 | *CTNNB1* | 0.740 | – | 0.178 | 408 | 5 | suite_bulk_oof_hardened |
| CRC | 5 | *ERBB2* | 0.537 | – | 0.044 | 408 | 5 | suite_bulk_oof_hardened |
| CRC | 5 | *FAT4* | 0.731 | – | 0.444 | 408 | 5 | suite_bulk_oof_hardened |
| CRC | 5 | *FBXW7* | 0.754 | – | 0.350 | 408 | 5 | suite_bulk_oof_hardened |
| CRC | 5 | *KMT2C* | 0.707 | – | 0.235 | 408 | 5 | suite_bulk_oof_hardened |
| CRC | 5 | *KMT2D* | 0.850 | – | 0.559 | 408 | 5 | suite_bulk_oof_hardened |
| CRC | 5 | *KRAS* | 0.619 | – | 0.453 | 408 | 5 | suite_bulk_oof_hardened |
| CRC | 5 | *MLH1* | 0.811 | – | 0.128 | 408 | 5 | suite_bulk_oof_hardened |
| CRC | 5 | *MSH2* | 0.772 | – | 0.129 | 408 | 5 | suite_bulk_oof_hardened |
| CRC | 5 | *MSH6* | 0.863 | – | 0.129 | 408 | 5 | suite_bulk_oof_hardened |
| CRC | 5 | *NRAS* | 0.610 | – | 0.067 | 408 | 5 | suite_bulk_oof_hardened |
| CRC | 5 | *PIK3CA* | 0.722 | – | 0.373 | 408 | 5 | suite_bulk_oof_hardened |
| CRC | 5 | *PMS2* | 0.845 | – | 0.125 | 408 | 5 | suite_bulk_oof_hardened |
| CRC | 5 | *PTEN* | 0.737 | – | 0.243 | 408 | 5 | suite_bulk_oof_hardened |
| CRC | 5 | *RNF43* | 0.912 | – | 0.568 | 408 | 5 | suite_bulk_oof_hardened |
| CRC | 5 | *SETD2* | 0.819 | – | 0.304 | 408 | 5 | suite_bulk_oof_hardened |
| CRC | 5 | *SMAD2* | 0.557 | – | 0.038 | 408 | 5 | suite_bulk_oof_hardened |
| CRC | 5 | *SMAD4* | 0.570 | – | 0.142 | 408 | 5 | suite_bulk_oof_hardened |
| CRC | 5 | *SOX9* | 0.671 | – | 0.172 | 408 | 5 | suite_bulk_oof_hardened |
| CRC | 5 | *TCF7L2* | 0.566 | – | 0.118 | 408 | 5 | suite_bulk_oof_hardened |
| CRC | 5 | *TGFBR2* | 0.722 | – | 0.163 | 408 | 5 | suite_bulk_oof_hardened |
| CRC | 5 | *TP53* | 0.838 | – | 0.853 | 408 | 5 | suite_bulk_oof_hardened |
| CRC | 10 | *APC* | 0.882 | – | 0.924 | 408 | 5 | suite_bulk_oof_hardened |
| CRC | 10 | *ARID1A* | 0.844 | – | 0.393 | 408 | 5 | suite_bulk_oof_hardened |
| CRC | 10 | *BRAF* | 0.872 | – | 0.594 | 408 | 5 | suite_bulk_oof_hardened |
| CRC | 10 | *CTNNB1* | 0.772 | – | 0.215 | 408 | 5 | suite_bulk_oof_hardened |
| CRC | 10 | *ERBB2* | 0.500 | – | 0.034 | 408 | 5 | suite_bulk_oof_hardened |
| CRC | 10 | *FAT4* | 0.731 | – | 0.414 | 408 | 5 | suite_bulk_oof_hardened |
| CRC | 10 | *FBXW7* | 0.762 | – | 0.395 | 408 | 5 | suite_bulk_oof_hardened |
| CRC | 10 | *KMT2C* | 0.764 | – | 0.269 | 408 | 5 | suite_bulk_oof_hardened |
| CRC | 10 | *KMT2D* | 0.874 | – | 0.584 | 408 | 5 | suite_bulk_oof_hardened |
| CRC | 10 | *KRAS* | 0.765 | – | 0.655 | 408 | 5 | suite_bulk_oof_hardened |
| CRC | 10 | *MLH1* | 0.763 | – | 0.111 | 408 | 5 | suite_bulk_oof_hardened |
| CRC | 10 | *MSH2* | 0.780 | – | 0.180 | 408 | 5 | suite_bulk_oof_hardened |
| CRC | 10 | *MSH6* | 0.867 | – | 0.149 | 408 | 5 | suite_bulk_oof_hardened |
| CRC | 10 | *NRAS* | 0.595 | – | 0.067 | 408 | 5 | suite_bulk_oof_hardened |
| CRC | 10 | *PIK3CA* | 0.737 | – | 0.395 | 408 | 5 | suite_bulk_oof_hardened |
| CRC | 10 | *PMS2* | 0.827 | – | 0.098 | 408 | 5 | suite_bulk_oof_hardened |
| CRC | 10 | *PTEN* | 0.760 | – | 0.229 | 408 | 5 | suite_bulk_oof_hardened |
| CRC | 10 | *RNF43* | 0.878 | – | 0.473 | 408 | 5 | suite_bulk_oof_hardened |
| CRC | 10 | *SETD2* | 0.794 | – | 0.268 | 408 | 5 | suite_bulk_oof_hardened |
| CRC | 10 | *SMAD2* | 0.565 | – | 0.041 | 408 | 5 | suite_bulk_oof_hardened |
| CRC | 10 | *SMAD4* | 0.590 | – | 0.146 | 408 | 5 | suite_bulk_oof_hardened |
| CRC | 10 | *SOX9* | 0.724 | – | 0.258 | 408 | 5 | suite_bulk_oof_hardened |
| CRC | 10 | *TCF7L2* | 0.601 | – | 0.147 | 408 | 5 | suite_bulk_oof_hardened |
| CRC | 10 | *TGFBR2* | 0.655 | – | 0.068 | 408 | 5 | suite_bulk_oof_hardened |
| CRC | 10 | *TP53* | 0.889 | – | 0.902 | 408 | 5 | suite_bulk_oof_hardened |
| CRC | 25 | *APC* | 0.851 | – | 0.874 | 408 | 5 | suite_bulk_oof_hardened |
| CRC | 25 | *ARID1A* | 0.845 | – | 0.343 | 408 | 5 | suite_bulk_oof_hardened |
| CRC | 25 | *BRAF* | 0.907 | – | 0.668 | 408 | 5 | suite_bulk_oof_hardened |
| CRC | 25 | *CTNNB1* | 0.810 | – | 0.314 | 408 | 5 | suite_bulk_oof_hardened |
| CRC | 25 | *ERBB2* | 0.433 | – | 0.024 | 408 | 5 | suite_bulk_oof_hardened |
| CRC | 25 | *FAT4* | 0.735 | – | 0.417 | 408 | 5 | suite_bulk_oof_hardened |
| CRC | 25 | *FBXW7* | 0.821 | – | 0.501 | 408 | 5 | suite_bulk_oof_hardened |
| CRC | 25 | *KMT2C* | 0.764 | – | 0.233 | 408 | 5 | suite_bulk_oof_hardened |
| CRC | 25 | *KMT2D* | 0.832 | – | 0.513 | 408 | 5 | suite_bulk_oof_hardened |
| CRC | 25 | *KRAS* | 0.818 | – | 0.712 | 408 | 5 | suite_bulk_oof_hardened |
| CRC | 25 | *MLH1* | 0.722 | – | 0.077 | 408 | 5 | suite_bulk_oof_hardened |
| CRC | 25 | *MSH2* | 0.801 | – | 0.165 | 408 | 5 | suite_bulk_oof_hardened |
| CRC | 25 | *MSH6* | 0.785 | – | 0.116 | 408 | 5 | suite_bulk_oof_hardened |
| CRC | 25 | *NRAS* | 0.538 | – | 0.057 | 408 | 5 | suite_bulk_oof_hardened |
| CRC | 25 | *PIK3CA* | 0.789 | – | 0.486 | 408 | 5 | suite_bulk_oof_hardened |
| CRC | 25 | *PMS2* | 0.835 | – | 0.158 | 408 | 5 | suite_bulk_oof_hardened |
| CRC | 25 | *PTEN* | 0.767 | – | 0.178 | 408 | 5 | suite_bulk_oof_hardened |
| CRC | 25 | *RNF43* | 0.863 | – | 0.523 | 408 | 5 | suite_bulk_oof_hardened |
| CRC | 25 | *SETD2* | 0.819 | – | 0.320 | 408 | 5 | suite_bulk_oof_hardened |
| CRC | 25 | *SMAD2* | 0.704 | – | 0.072 | 408 | 5 | suite_bulk_oof_hardened |
| CRC | 25 | *SMAD4* | 0.689 | – | 0.219 | 408 | 5 | suite_bulk_oof_hardened |
| CRC | 25 | *SOX9* | 0.771 | – | 0.344 | 408 | 5 | suite_bulk_oof_hardened |
| CRC | 25 | *TCF7L2* | 0.599 | – | 0.158 | 408 | 5 | suite_bulk_oof_hardened |

Continued on next page

*Supplementary Table S9 continued*

| Cancer | $k$ | Driver | AUROC mean | AUROC SD | AUPRC mean | Samples | CV folds | Implementation |
|---|---|---|---|---|---|---|---|---|
| CRC | 25 | *TGFBR2* | 0.777 | – | 0.128 | 408 | 5 | suite_bulk_oof_hardened |
| CRC | 25 | *TP53* | 0.900 | – | 0.890 | 408 | 5 | suite_bulk_oof_hardened |
| CRC | 50 | *APC* | 0.844 | – | 0.875 | 408 | 5 | suite_bulk_oof_hardened |
| CRC | 50 | *ARID1A* | 0.805 | – | 0.343 | 408 | 5 | suite_bulk_oof_hardened |
| CRC | 50 | *BRAF* | 0.900 | – | 0.648 | 408 | 5 | suite_bulk_oof_hardened |
| CRC | 50 | *CTNNB1* | 0.786 | – | 0.283 | 408 | 5 | suite_bulk_oof_hardened |
| CRC | 50 | *ERBB2* | 0.453 | – | 0.030 | 408 | 5 | suite_bulk_oof_hardened |
| CRC | 50 | *FAT4* | 0.744 | – | 0.400 | 408 | 5 | suite_bulk_oof_hardened |
| CRC | 50 | *FBXW7* | 0.835 | – | 0.527 | 408 | 5 | suite_bulk_oof_hardened |
| CRC | 50 | *KMT2C* | 0.789 | – | 0.239 | 408 | 5 | suite_bulk_oof_hardened |
| CRC | 50 | *KMT2D* | 0.808 | – | 0.515 | 408 | 5 | suite_bulk_oof_hardened |
| CRC | 50 | *KRAS* | 0.849 | – | 0.734 | 408 | 5 | suite_bulk_oof_hardened |
| CRC | 50 | *MLH1* | 0.741 | – | 0.097 | 408 | 5 | suite_bulk_oof_hardened |
| CRC | 50 | *MSH2* | 0.823 | – | 0.114 | 408 | 5 | suite_bulk_oof_hardened |
| CRC | 50 | *MSH6* | 0.719 | – | 0.106 | 408 | 5 | suite_bulk_oof_hardened |
| CRC | 50 | *NRAS* | 0.611 | – | 0.068 | 408 | 5 | suite_bulk_oof_hardened |
| CRC | 50 | *PIK3CA* | 0.775 | – | 0.468 | 408 | 5 | suite_bulk_oof_hardened |
| CRC | 50 | *PMS2* | 0.862 | – | 0.126 | 408 | 5 | suite_bulk_oof_hardened |
| CRC | 50 | *PTEN* | 0.694 | – | 0.116 | 408 | 5 | suite_bulk_oof_hardened |
| CRC | 50 | *RNF43* | 0.826 | – | 0.477 | 408 | 5 | suite_bulk_oof_hardened |
| CRC | 50 | *SETD2* | 0.771 | – | 0.255 | 408 | 5 | suite_bulk_oof_hardened |
| CRC | 50 | *SMAD2* | 0.671 | – | 0.056 | 408 | 5 | suite_bulk_oof_hardened |
| CRC | 50 | *SMAD4* | 0.673 | – | 0.188 | 408 | 5 | suite_bulk_oof_hardened |
| CRC | 50 | *SOX9* | 0.787 | – | 0.367 | 408 | 5 | suite_bulk_oof_hardened |
| CRC | 50 | *TCF7L2* | 0.613 | – | 0.163 | 408 | 5 | suite_bulk_oof_hardened |
| CRC | 50 | *TGFBR2* | 0.730 | – | 0.096 | 408 | 5 | suite_bulk_oof_hardened |
| CRC | 50 | *TP53* | 0.904 | – | 0.902 | 408 | 5 | suite_bulk_oof_hardened |
| CRC | 75 | *APC* | 0.844 | – | 0.875 | 408 | 5 | suite_bulk_oof_hardened |
| CRC | 75 | *ARID1A* | 0.799 | – | 0.344 | 408 | 5 | suite_bulk_oof_hardened |
| CRC | 75 | *BRAF* | 0.904 | – | 0.647 | 408 | 5 | suite_bulk_oof_hardened |
| CRC | 75 | *CTNNB1* | 0.741 | – | 0.224 | 408 | 5 | suite_bulk_oof_hardened |
| CRC | 75 | *ERBB2* | 0.452 | – | 0.025 | 408 | 5 | suite_bulk_oof_hardened |
| CRC | 75 | *FAT4* | 0.717 | – | 0.388 | 408 | 5 | suite_bulk_oof_hardened |
| CRC | 75 | *FBXW7* | 0.825 | – | 0.568 | 408 | 5 | suite_bulk_oof_hardened |
| CRC | 75 | *KMT2C* | 0.777 | – | 0.247 | 408 | 5 | suite_bulk_oof_hardened |
| CRC | 75 | *KMT2D* | 0.820 | – | 0.524 | 408 | 5 | suite_bulk_oof_hardened |
| CRC | 75 | *KRAS* | 0.867 | – | 0.781 | 408 | 5 | suite_bulk_oof_hardened |
| CRC | 75 | *MLH1* | 0.743 | – | 0.157 | 408 | 5 | suite_bulk_oof_hardened |
| CRC | 75 | *MSH2* | 0.770 | – | 0.183 | 408 | 5 | suite_bulk_oof_hardened |
| CRC | 75 | *MSH6* | 0.739 | – | 0.121 | 408 | 5 | suite_bulk_oof_hardened |
| CRC | 75 | *NRAS* | 0.608 | – | 0.067 | 408 | 5 | suite_bulk_oof_hardened |
| CRC | 75 | *PIK3CA* | 0.746 | – | 0.446 | 408 | 5 | suite_bulk_oof_hardened |
| CRC | 75 | *PMS2* | 0.830 | – | 0.101 | 408 | 5 | suite_bulk_oof_hardened |
| CRC | 75 | *PTEN* | 0.716 | – | 0.124 | 408 | 5 | suite_bulk_oof_hardened |
| CRC | 75 | *RNF43* | 0.837 | – | 0.527 | 408 | 5 | suite_bulk_oof_hardened |
| CRC | 75 | *SETD2* | 0.734 | – | 0.188 | 408 | 5 | suite_bulk_oof_hardened |
| CRC | 75 | *SMAD2* | 0.674 | – | 0.057 | 408 | 5 | suite_bulk_oof_hardened |
| CRC | 75 | *SMAD4* | 0.666 | – | 0.205 | 408 | 5 | suite_bulk_oof_hardened |
| CRC | 75 | *SOX9* | 0.782 | – | 0.379 | 408 | 5 | suite_bulk_oof_hardened |
| CRC | 75 | *TCF7L2* | 0.631 | – | 0.169 | 408 | 5 | suite_bulk_oof_hardened |
| CRC | 75 | *TGFBR2* | 0.696 | – | 0.078 | 408 | 5 | suite_bulk_oof_hardened |
| CRC | 75 | *TP53* | 0.913 | – | 0.910 | 408 | 5 | suite_bulk_oof_hardened |
| CRC | 100 | *APC* | 0.825 | – | 0.867 | 408 | 5 | suite_bulk_oof_hardened |
| CRC | 100 | *ARID1A* | 0.768 | – | 0.272 | 408 | 5 | suite_bulk_oof_hardened |
| CRC | 100 | *BRAF* | 0.884 | – | 0.664 | 408 | 5 | suite_bulk_oof_hardened |
| CRC | 100 | *CTNNB1* | 0.701 | – | 0.208 | 408 | 5 | suite_bulk_oof_hardened |
| CRC | 100 | *ERBB2* | 0.372 | – | 0.022 | 408 | 5 | suite_bulk_oof_hardened |
| CRC | 100 | *FAT4* | 0.698 | – | 0.362 | 408 | 5 | suite_bulk_oof_hardened |
| CRC | 100 | *FBXW7* | 0.834 | – | 0.578 | 408 | 5 | suite_bulk_oof_hardened |
| CRC | 100 | *KMT2C* | 0.793 | – | 0.297 | 408 | 5 | suite_bulk_oof_hardened |
| CRC | 100 | *KMT2D* | 0.834 | – | 0.533 | 408 | 5 | suite_bulk_oof_hardened |
| CRC | 100 | *KRAS* | 0.853 | – | 0.760 | 408 | 5 | suite_bulk_oof_hardened |
| CRC | 100 | *MLH1* | 0.734 | – | 0.090 | 408 | 5 | suite_bulk_oof_hardened |
| CRC | 100 | *MSH2* | 0.834 | – | 0.146 | 408 | 5 | suite_bulk_oof_hardened |
| CRC | 100 | *MSH6* | 0.746 | – | 0.109 | 408 | 5 | suite_bulk_oof_hardened |
| CRC | 100 | *NRAS* | 0.604 | – | 0.065 | 408 | 5 | suite_bulk_oof_hardened |
| CRC | 100 | *PIK3CA* | 0.743 | – | 0.435 | 408 | 5 | suite_bulk_oof_hardened |
| CRC | 100 | *PMS2* | 0.766 | – | 0.071 | 408 | 5 | suite_bulk_oof_hardened |
| CRC | 100 | *PTEN* | 0.691 | – | 0.136 | 408 | 5 | suite_bulk_oof_hardened |
| CRC | 100 | *RNF43* | 0.847 | – | 0.475 | 408 | 5 | suite_bulk_oof_hardened |
| CRC | 100 | *SETD2* | 0.734 | – | 0.254 | 408 | 5 | suite_bulk_oof_hardened |
| CRC | 100 | *SMAD2* | 0.595 | – | 0.044 | 408 | 5 | suite_bulk_oof_hardened |
| CRC | 100 | *SMAD4* | 0.673 | – | 0.221 | 408 | 5 | suite_bulk_oof_hardened |

Continued on next page

*Supplementary Table S9 continued*

| Cancer | $k$ | Driver | AUROC mean | AUROC SD | AUPRC mean | Samples | CV folds | Implementation |
|---|---|---|---|---|---|---|---|---|
| CRC | 100 | *SOX9* | 0.798 | – | 0.369 | 408 | 5 | `suite_bulk_oof_hardened` |
| CRC | 100 | *TCF7L2* | 0.649 | – | 0.150 | 408 | 5 | `suite_bulk_oof_hardened` |
| CRC | 100 | *TGFBR2* | 0.705 | – | 0.083 | 408 | 5 | `suite_bulk_oof_hardened` |
| CRC | 100 | *TP53* | 0.912 | – | 0.917 | 408 | 5 | `suite_bulk_oof_hardened` |
| CRC | 150 | *APC* | 0.842 | – | 0.867 | 408 | 5 | `suite_bulk_oof_hardened` |
| CRC | 150 | *ARID1A* | 0.802 | – | 0.304 | 408 | 5 | `suite_bulk_oof_hardened` |
| CRC | 150 | *BRAF* | 0.915 | – | 0.744 | 408 | 5 | `suite_bulk_oof_hardened` |
| CRC | 150 | *CTNNB1* | 0.762 | – | 0.272 | 408 | 5 | `suite_bulk_oof_hardened` |
| CRC | 150 | *ERBB2* | 0.319 | – | 0.019 | 408 | 5 | `suite_bulk_oof_hardened` |
| CRC | 150 | *FAT4* | 0.689 | – | 0.356 | 408 | 5 | `suite_bulk_oof_hardened` |
| CRC | 150 | *FBXW7* | 0.863 | – | 0.637 | 408 | 5 | `suite_bulk_oof_hardened` |
| CRC | 150 | *KMT2C* | 0.800 | – | 0.311 | 408 | 5 | `suite_bulk_oof_hardened` |
| CRC | 150 | *KMT2D* | 0.865 | – | 0.555 | 408 | 5 | `suite_bulk_oof_hardened` |
| CRC | 150 | *KRAS* | 0.862 | – | 0.789 | 408 | 5 | `suite_bulk_oof_hardened` |
| CRC | 150 | *MLH1* | 0.750 | – | 0.108 | 408 | 5 | `suite_bulk_oof_hardened` |
| CRC | 150 | *MSH2* | 0.823 | – | 0.133 | 408 | 5 | `suite_bulk_oof_hardened` |
| CRC | 150 | *MSH6* | 0.795 | – | 0.159 | 408 | 5 | `suite_bulk_oof_hardened` |
| CRC | 150 | *NRAS* | 0.573 | – | 0.070 | 408 | 5 | `suite_bulk_oof_hardened` |
| CRC | 150 | *PIK3CA* | 0.739 | – | 0.463 | 408 | 5 | `suite_bulk_oof_hardened` |
| CRC | 150 | *PMS2* | 0.712 | – | 0.061 | 408 | 5 | `suite_bulk_oof_hardened` |
| CRC | 150 | *PTEN* | 0.657 | – | 0.175 | 408 | 5 | `suite_bulk_oof_hardened` |
| CRC | 150 | *RNF43* | 0.859 | – | 0.464 | 408 | 5 | `suite_bulk_oof_hardened` |
| CRC | 150 | *SETD2* | 0.743 | – | 0.206 | 408 | 5 | `suite_bulk_oof_hardened` |
| CRC | 150 | *SMAD2* | 0.625 | – | 0.058 | 408 | 5 | `suite_bulk_oof_hardened` |
| CRC | 150 | *SMAD4* | 0.670 | – | 0.256 | 408 | 5 | `suite_bulk_oof_hardened` |
| CRC | 150 | *SOX9* | 0.818 | – | 0.467 | 408 | 5 | `suite_bulk_oof_hardened` |
| CRC | 150 | *TCF7L2* | 0.645 | – | 0.169 | 408 | 5 | `suite_bulk_oof_hardened` |
| CRC | 150 | *TGFBR2* | 0.703 | – | 0.068 | 408 | 5 | `suite_bulk_oof_hardened` |
| CRC | 150 | *TP53* | 0.919 | – | 0.917 | 408 | 5 | `suite_bulk_oof_hardened` |
| CRC | 250 | *APC* | 0.866 | – | 0.881 | 408 | 5 | `suite_bulk_oof_hardened` |
| CRC | 250 | *ARID1A* | 0.831 | – | 0.350 | 408 | 5 | `suite_bulk_oof_hardened` |
| CRC | 250 | *BRAF* | 0.921 | – | 0.790 | 408 | 5 | `suite_bulk_oof_hardened` |
| CRC | 250 | *CTNNB1* | 0.758 | – | 0.243 | 408 | 5 | `suite_bulk_oof_hardened` |
| CRC | 250 | *ERBB2* | 0.382 | – | 0.023 | 408 | 5 | `suite_bulk_oof_hardened` |
| CRC | 250 | *FAT4* | 0.690 | – | 0.402 | 408 | 5 | `suite_bulk_oof_hardened` |
| CRC | 250 | *FBXW7* | 0.899 | – | 0.654 | 408 | 5 | `suite_bulk_oof_hardened` |
| CRC | 250 | *KMT2C* | 0.812 | – | 0.296 | 408 | 5 | `suite_bulk_oof_hardened` |
| CRC | 250 | *KMT2D* | 0.869 | – | 0.575 | 408 | 5 | `suite_bulk_oof_hardened` |
| CRC | 250 | *KRAS* | 0.887 | – | 0.817 | 408 | 5 | `suite_bulk_oof_hardened` |
| CRC | 250 | *MLH1* | 0.753 | – | 0.130 | 408 | 5 | `suite_bulk_oof_hardened` |
| CRC | 250 | *MSH2* | 0.860 | – | 0.189 | 408 | 5 | `suite_bulk_oof_hardened` |
| CRC | 250 | *MSH6* | 0.763 | – | 0.120 | 408 | 5 | `suite_bulk_oof_hardened` |
| CRC | 250 | *NRAS* | 0.586 | – | 0.069 | 408 | 5 | `suite_bulk_oof_hardened` |
| CRC | 250 | *PIK3CA* | 0.760 | – | 0.476 | 408 | 5 | `suite_bulk_oof_hardened` |
| CRC | 250 | *PMS2* | 0.726 | – | 0.096 | 408 | 5 | `suite_bulk_oof_hardened` |
| CRC | 250 | *PTEN* | 0.689 | – | 0.176 | 408 | 5 | `suite_bulk_oof_hardened` |
| CRC | 250 | *RNF43* | 0.865 | – | 0.448 | 408 | 5 | `suite_bulk_oof_hardened` |
| CRC | 250 | *SETD2* | 0.777 | – | 0.245 | 408 | 5 | `suite_bulk_oof_hardened` |
| CRC | 250 | *SMAD2* | 0.680 | – | 0.079 | 408 | 5 | `suite_bulk_oof_hardened` |
| CRC | 250 | *SMAD4* | 0.710 | – | 0.247 | 408 | 5 | `suite_bulk_oof_hardened` |
| CRC | 250 | *SOX9* | 0.814 | – | 0.480 | 408 | 5 | `suite_bulk_oof_hardened` |
| CRC | 250 | *TCF7L2* | 0.657 | – | 0.165 | 408 | 5 | `suite_bulk_oof_hardened` |
| CRC | 250 | *TGFBR2* | 0.725 | – | 0.085 | 408 | 5 | `suite_bulk_oof_hardened` |
| CRC | 250 | *TP53* | 0.930 | – | 0.929 | 408 | 5 | `suite_bulk_oof_hardened` |
| GBM | 5 | *ATRX* | 0.473 | – | 0.187 | 155 | 2 | `suite_bulk_oof_hardened` |
| GBM | 5 | *CDKN2A* | 0.471 | – | 0.031 | 155 | 2 | `suite_bulk_oof_hardened` |
| GBM | 5 | *CIC* | 0.263 | – | 0.017 | 155 | 2 | `suite_bulk_oof_hardened` |
| GBM | 5 | *EGFR* | 0.569 | – | 0.359 | 155 | 2 | `suite_bulk_oof_hardened` |
| GBM | 5 | *IDH1* | 0.486 | – | 0.058 | 155 | 2 | `suite_bulk_oof_hardened` |
| GBM | 5 | *NF1* | 0.716 | – | 0.237 | 155 | 2 | `suite_bulk_oof_hardened` |
| GBM | 5 | *PDGFRA* | 0.665 | – | 0.117 | 155 | 2 | `suite_bulk_oof_hardened` |
| GBM | 5 | *PIK3CA* | 0.494 | – | 0.099 | 155 | 2 | `suite_bulk_oof_hardened` |
| GBM | 5 | *PIK3R1* | 0.407 | – | 0.074 | 155 | 2 | `suite_bulk_oof_hardened` |
| GBM | 5 | *PTEN* | 0.625 | – | 0.428 | 155 | 2 | `suite_bulk_oof_hardened` |
| GBM | 5 | *RB1* | 0.572 | – | 0.175 | 155 | 2 | `suite_bulk_oof_hardened` |
| GBM | 5 | *STAG2* | 0.314 | – | 0.036 | 155 | 2 | `suite_bulk_oof_hardened` |
| GBM | 5 | *TERT* | 0.301 | – | 0.013 | 155 | 2 | `suite_bulk_oof_hardened` |
| GBM | 5 | *TP53* | 0.669 | – | 0.508 | 155 | 2 | `suite_bulk_oof_hardened` |
| GBM | 10 | *ATRX* | 0.837 | – | 0.341 | 155 | 2 | `suite_bulk_oof_hardened` |
| GBM | 10 | *CDKN2A* | 0.401 | – | 0.022 | 155 | 2 | `suite_bulk_oof_hardened` |
| GBM | 10 | *CIC* | 0.625 | – | 0.033 | 155 | 2 | `suite_bulk_oof_hardened` |
| GBM | 10 | *EGFR* | 0.640 | – | 0.460 | 155 | 2 | `suite_bulk_oof_hardened` |

Continued on next page

| Cancer | *k* | Driver | AUROC mean | AUROC SD | AUPRC mean | Samples | CV folds | Implementation |
|---|---|---|---|---|---|---|---|---|
| GBM | 10 | *IDH1* | 0.769 | – | 0.228 | 155 | 2 | suite_bulk_oof_hardened |
| GBM | 10 | *NF1* | 0.698 | – | 0.216 | 155 | 2 | suite_bulk_oof_hardened |
| GBM | 10 | *PDGFRA* | 0.659 | – | 0.106 | 155 | 2 | suite_bulk_oof_hardened |
| GBM | 10 | *PIK3CA* | 0.456 | – | 0.107 | 155 | 2 | suite_bulk_oof_hardened |
| GBM | 10 | *PIK3R1* | 0.336 | – | 0.059 | 155 | 2 | suite_bulk_oof_hardened |
| GBM | 10 | *PTEN* | 0.638 | – | 0.466 | 155 | 2 | suite_bulk_oof_hardened |
| GBM | 10 | *RB1* | 0.632 | – | 0.176 | 155 | 2 | suite_bulk_oof_hardened |
| GBM | 10 | *STAG2* | 0.332 | – | 0.037 | 155 | 2 | suite_bulk_oof_hardened |
| GBM | 10 | *TERT* | 0.422 | – | 0.016 | 155 | 2 | suite_bulk_oof_hardened |
| GBM | 10 | *TP53* | 0.775 | – | 0.654 | 155 | 2 | suite_bulk_oof_hardened |
| GBM | 25 | *ATRX* | 0.803 | – | 0.295 | 155 | 2 | suite_bulk_oof_hardened |
| GBM | 25 | *CDKN2A* | 0.524 | – | 0.029 | 155 | 2 | suite_bulk_oof_hardened |
| GBM | 25 | *CIC* | 0.618 | – | 0.032 | 155 | 2 | suite_bulk_oof_hardened |
| GBM | 25 | *EGFR* | 0.691 | – | 0.506 | 155 | 2 | suite_bulk_oof_hardened |
| GBM | 25 | *IDH1* | 0.790 | – | 0.247 | 155 | 2 | suite_bulk_oof_hardened |
| GBM | 25 | *NF1* | 0.594 | – | 0.200 | 155 | 2 | suite_bulk_oof_hardened |
| GBM | 25 | *PDGFRA* | 0.778 | – | 0.216 | 155 | 2 | suite_bulk_oof_hardened |
| GBM | 25 | *PIK3CA* | 0.469 | – | 0.085 | 155 | 2 | suite_bulk_oof_hardened |
| GBM | 25 | *PIK3R1* | 0.428 | – | 0.075 | 155 | 2 | suite_bulk_oof_hardened |
| GBM | 25 | *PTEN* | 0.640 | – | 0.453 | 155 | 2 | suite_bulk_oof_hardened |
| GBM | 25 | *RB1* | 0.631 | – | 0.194 | 155 | 2 | suite_bulk_oof_hardened |
| GBM | 25 | *STAG2* | 0.379 | – | 0.041 | 155 | 2 | suite_bulk_oof_hardened |
| GBM | 25 | *TERT* | 0.431 | – | 0.019 | 155 | 2 | suite_bulk_oof_hardened |
| GBM | 25 | *TP53* | 0.801 | – | 0.731 | 155 | 2 | suite_bulk_oof_hardened |
| GBM | 50 | *ATRX* | 0.808 | – | 0.295 | 155 | 2 | suite_bulk_oof_hardened |
| GBM | 50 | *CDKN2A* | 0.504 | – | 0.028 | 155 | 2 | suite_bulk_oof_hardened |
| GBM | 50 | *CIC* | 0.559 | – | 0.028 | 155 | 2 | suite_bulk_oof_hardened |
| GBM | 50 | *EGFR* | 0.696 | – | 0.517 | 155 | 2 | suite_bulk_oof_hardened |
| GBM | 50 | *IDH1* | 0.794 | – | 0.249 | 155 | 2 | suite_bulk_oof_hardened |
| GBM | 50 | *NF1* | 0.601 | – | 0.170 | 155 | 2 | suite_bulk_oof_hardened |
| GBM | 50 | *PDGFRA* | 0.779 | – | 0.208 | 155 | 2 | suite_bulk_oof_hardened |
| GBM | 50 | *PIK3CA* | 0.601 | – | 0.129 | 155 | 2 | suite_bulk_oof_hardened |
| GBM | 50 | *PIK3R1* | 0.580 | – | 0.121 | 155 | 2 | suite_bulk_oof_hardened |
| GBM | 50 | *PTEN* | 0.602 | – | 0.414 | 155 | 2 | suite_bulk_oof_hardened |
| GBM | 50 | *RB1* | 0.742 | – | 0.286 | 155 | 2 | suite_bulk_oof_hardened |
| GBM | 50 | *STAG2* | 0.416 | – | 0.045 | 155 | 2 | suite_bulk_oof_hardened |
| GBM | 50 | *TERT* | 0.118 | – | 0.011 | 155 | 2 | suite_bulk_oof_hardened |
| GBM | 50 | *TP53* | 0.809 | – | 0.696 | 155 | 2 | suite_bulk_oof_hardened |
| GBM | 75 | *ATRX* | 0.806 | – | 0.296 | 155 | 2 | suite_bulk_oof_hardened |
| GBM | 75 | *CDKN2A* | 0.439 | – | 0.024 | 155 | 2 | suite_bulk_oof_hardened |
| GBM | 75 | *CIC* | 0.526 | – | 0.027 | 155 | 2 | suite_bulk_oof_hardened |
| GBM | 75 | *EGFR* | 0.715 | – | 0.524 | 155 | 2 | suite_bulk_oof_hardened |
| GBM | 75 | *IDH1* | 0.785 | – | 0.250 | 155 | 2 | suite_bulk_oof_hardened |
| GBM | 75 | *NF1* | 0.619 | – | 0.159 | 155 | 2 | suite_bulk_oof_hardened |
| GBM | 75 | *PDGFRA* | 0.775 | – | 0.196 | 155 | 2 | suite_bulk_oof_hardened |
| GBM | 75 | *PIK3CA* | 0.605 | – | 0.122 | 155 | 2 | suite_bulk_oof_hardened |
| GBM | 75 | *PIK3R1* | 0.509 | – | 0.105 | 155 | 2 | suite_bulk_oof_hardened |
| GBM | 75 | *PTEN* | 0.590 | – | 0.390 | 155 | 2 | suite_bulk_oof_hardened |
| GBM | 75 | *RB1* | 0.710 | – | 0.293 | 155 | 2 | suite_bulk_oof_hardened |
| GBM | 75 | *STAG2* | 0.482 | – | 0.059 | 155 | 2 | suite_bulk_oof_hardened |
| GBM | 75 | *TERT* | 0.056 | – | 0.010 | 155 | 2 | suite_bulk_oof_hardened |
| GBM | 75 | *TP53* | 0.830 | – | 0.744 | 155 | 2 | suite_bulk_oof_hardened |
| GBM | 100 | *ATRX* | 0.805 | – | 0.295 | 155 | 2 | suite_bulk_oof_hardened |
| GBM | 100 | *CDKN2A* | 0.428 | – | 0.024 | 155 | 2 | suite_bulk_oof_hardened |
| GBM | 100 | *CIC* | 0.529 | – | 0.027 | 155 | 2 | suite_bulk_oof_hardened |
| GBM | 100 | *EGFR* | 0.712 | – | 0.537 | 155 | 2 | suite_bulk_oof_hardened |
| GBM | 100 | *IDH1* | 0.782 | – | 0.248 | 155 | 2 | suite_bulk_oof_hardened |
| GBM | 100 | *NF1* | 0.621 | – | 0.160 | 155 | 2 | suite_bulk_oof_hardened |
| GBM | 100 | *PDGFRA* | 0.775 | – | 0.200 | 155 | 2 | suite_bulk_oof_hardened |
| GBM | 100 | *PIK3CA* | 0.602 | – | 0.122 | 155 | 2 | suite_bulk_oof_hardened |
| GBM | 100 | *PIK3R1* | 0.498 | – | 0.102 | 155 | 2 | suite_bulk_oof_hardened |
| GBM | 100 | *PTEN* | 0.588 | – | 0.388 | 155 | 2 | suite_bulk_oof_hardened |
| GBM | 100 | *RB1* | 0.710 | – | 0.293 | 155 | 2 | suite_bulk_oof_hardened |
| GBM | 100 | *STAG2* | 0.479 | – | 0.059 | 155 | 2 | suite_bulk_oof_hardened |
| GBM | 100 | *TERT* | 0.052 | – | 0.010 | 155 | 2 | suite_bulk_oof_hardened |
| GBM | 100 | *TP53* | 0.831 | – | 0.745 | 155 | 2 | suite_bulk_oof_hardened |
| GBM | 150 | *ATRX* | 0.805 | – | 0.295 | 155 | 2 | suite_bulk_oof_hardened |
| GBM | 150 | *CDKN2A* | 0.428 | – | 0.024 | 155 | 2 | suite_bulk_oof_hardened |
| GBM | 150 | *CIC* | 0.529 | – | 0.027 | 155 | 2 | suite_bulk_oof_hardened |
| GBM | 150 | *EGFR* | 0.712 | – | 0.537 | 155 | 2 | suite_bulk_oof_hardened |
| GBM | 150 | *IDH1* | 0.782 | – | 0.248 | 155 | 2 | suite_bulk_oof_hardened |
| GBM | 150 | *NF1* | 0.621 | – | 0.160 | 155 | 2 | suite_bulk_oof_hardened |

Continued on next page

*Supplementary Table S9 continued*

| Cancer | $k$ | Driver | AUROC mean | AUROC SD | AUPRC mean | Samples | CV folds | Implementation |
|---|---|---|---|---|---|---|---|---|
| GBM | 150 | *PDGFRA* | 0.775 | – | 0.200 | 155 | 2 | suite_bulk_oof_hardened |
| GBM | 150 | *PIK3CA* | 0.602 | – | 0.122 | 155 | 2 | suite_bulk_oof_hardened |
| GBM | 150 | *PIK3R1* | 0.498 | – | 0.102 | 155 | 2 | suite_bulk_oof_hardened |
| GBM | 150 | *PTEN* | 0.588 | – | 0.388 | 155 | 2 | suite_bulk_oof_hardened |
| GBM | 150 | *RB1* | 0.710 | – | 0.293 | 155 | 2 | suite_bulk_oof_hardened |
| GBM | 150 | *STAG2* | 0.479 | – | 0.059 | 155 | 2 | suite_bulk_oof_hardened |
| GBM | 150 | *TERT* | 0.052 | – | 0.010 | 155 | 2 | suite_bulk_oof_hardened |
| GBM | 150 | *TP53* | 0.831 | – | 0.745 | 155 | 2 | suite_bulk_oof_hardened |
| LUAD | 5 | *ARID1A* | 0.614 | – | 0.077 | 567 | 5 | suite_bulk_oof_hardened |
| LUAD | 5 | *ATM* | 0.602 | – | 0.093 | 567 | 5 | suite_bulk_oof_hardened |
| LUAD | 5 | *BRAF* | 0.617 | – | 0.106 | 567 | 5 | suite_bulk_oof_hardened |
| LUAD | 5 | *CDKN2A* | 0.502 | – | 0.041 | 567 | 5 | suite_bulk_oof_hardened |
| LUAD | 5 | *EGFR* | 0.553 | – | 0.146 | 567 | 5 | suite_bulk_oof_hardened |
| LUAD | 5 | *ERBB2* | 0.699 | – | 0.055 | 454 | 5 | suite_bulk_oof_hardened |
| LUAD | 5 | *KEAP1* | 0.801 | – | 0.406 | 567 | 5 | suite_bulk_oof_hardened |
| LUAD | 5 | *KRAS* | 0.689 | – | 0.391 | 567 | 5 | suite_bulk_oof_hardened |
| LUAD | 5 | *MET* | 0.582 | – | 0.091 | 567 | 5 | suite_bulk_oof_hardened |
| LUAD | 5 | *MGA* | 0.611 | – | 0.096 | 567 | 5 | suite_bulk_oof_hardened |
| LUAD | 5 | *NF1* | 0.637 | – | 0.148 | 567 | 5 | suite_bulk_oof_hardened |
| LUAD | 5 | *PIK3CA* | 0.555 | – | 0.063 | 567 | 5 | suite_bulk_oof_hardened |
| LUAD | 5 | *PTPRD* | 0.667 | – | 0.225 | 567 | 5 | suite_bulk_oof_hardened |
| LUAD | 5 | *RB1* | 0.768 | – | 0.192 | 567 | 5 | suite_bulk_oof_hardened |
| LUAD | 5 | *RBM10* | 0.561 | – | 0.079 | 567 | 5 | suite_bulk_oof_hardened |
| LUAD | 5 | *SETD2* | 0.562 | – | 0.068 | 567 | 5 | suite_bulk_oof_hardened |
| LUAD | 5 | *SMARCA4* | 0.759 | – | 0.184 | 567 | 5 | suite_bulk_oof_hardened |
| LUAD | 5 | *STK11* | 0.750 | – | 0.268 | 567 | 5 | suite_bulk_oof_hardened |
| LUAD | 5 | *TP53* | 0.825 | – | 0.787 | 567 | 5 | suite_bulk_oof_hardened |
| LUAD | 5 | *U2AF1* | 0.597 | – | 0.035 | 567 | 5 | suite_bulk_oof_hardened |
| LUAD | 10 | *ARID1A* | 0.550 | – | 0.063 | 567 | 5 | suite_bulk_oof_hardened |
| LUAD | 10 | *ATM* | 0.722 | – | 0.192 | 567 | 5 | suite_bulk_oof_hardened |
| LUAD | 10 | *BRAF* | 0.591 | – | 0.116 | 567 | 5 | suite_bulk_oof_hardened |
| LUAD | 10 | *CDKN2A* | 0.508 | – | 0.039 | 567 | 5 | suite_bulk_oof_hardened |
| LUAD | 10 | *EGFR* | 0.831 | – | 0.480 | 567 | 5 | suite_bulk_oof_hardened |
| LUAD | 10 | *ERBB2* | 0.618 | – | 0.018 | 454 | 5 | suite_bulk_oof_hardened |
| LUAD | 10 | *KEAP1* | 0.865 | – | 0.485 | 567 | 5 | suite_bulk_oof_hardened |
| LUAD | 10 | *KRAS* | 0.786 | – | 0.554 | 567 | 5 | suite_bulk_oof_hardened |
| LUAD | 10 | *MET* | 0.534 | – | 0.058 | 567 | 5 | suite_bulk_oof_hardened |
| LUAD | 10 | *MGA* | 0.622 | – | 0.119 | 567 | 5 | suite_bulk_oof_hardened |
| LUAD | 10 | *NF1* | 0.616 | – | 0.133 | 567 | 5 | suite_bulk_oof_hardened |
| LUAD | 10 | *PIK3CA* | 0.569 | – | 0.058 | 567 | 5 | suite_bulk_oof_hardened |
| LUAD | 10 | *PTPRD* | 0.676 | – | 0.260 | 567 | 5 | suite_bulk_oof_hardened |
| LUAD | 10 | *RB1* | 0.807 | – | 0.223 | 567 | 5 | suite_bulk_oof_hardened |
| LUAD | 10 | *RBM10* | 0.478 | – | 0.056 | 567 | 5 | suite_bulk_oof_hardened |
| LUAD | 10 | *SETD2* | 0.610 | – | 0.089 | 567 | 5 | suite_bulk_oof_hardened |
| LUAD | 10 | *SMARCA4* | 0.788 | – | 0.236 | 567 | 5 | suite_bulk_oof_hardened |
| LUAD | 10 | *STK11* | 0.877 | – | 0.463 | 567 | 5 | suite_bulk_oof_hardened |
| LUAD | 10 | *TP53* | 0.871 | – | 0.835 | 567 | 5 | suite_bulk_oof_hardened |
| LUAD | 10 | *U2AF1* | 0.713 | – | 0.054 | 567 | 5 | suite_bulk_oof_hardened |
| LUAD | 25 | *ARID1A* | 0.494 | – | 0.055 | 567 | 5 | suite_bulk_oof_hardened |
| LUAD | 25 | *ATM* | 0.712 | – | 0.210 | 567 | 5 | suite_bulk_oof_hardened |
| LUAD | 25 | *BRAF* | 0.591 | – | 0.107 | 567 | 5 | suite_bulk_oof_hardened |
| LUAD | 25 | *CDKN2A* | 0.492 | – | 0.037 | 567 | 5 | suite_bulk_oof_hardened |
| LUAD | 25 | *EGFR* | 0.842 | – | 0.467 | 567 | 5 | suite_bulk_oof_hardened |
| LUAD | 25 | *ERBB2* | 0.788 | – | 0.073 | 454 | 5 | suite_bulk_oof_hardened |
| LUAD | 25 | *KEAP1* | 0.934 | – | 0.712 | 567 | 5 | suite_bulk_oof_hardened |
| LUAD | 25 | *KRAS* | 0.840 | – | 0.605 | 567 | 5 | suite_bulk_oof_hardened |
| LUAD | 25 | *MET* | 0.642 | – | 0.075 | 567 | 5 | suite_bulk_oof_hardened |
| LUAD | 25 | *MGA* | 0.625 | – | 0.159 | 567 | 5 | suite_bulk_oof_hardened |
| LUAD | 25 | *NF1* | 0.661 | – | 0.183 | 567 | 5 | suite_bulk_oof_hardened |
| LUAD | 25 | *PIK3CA* | 0.641 | – | 0.087 | 567 | 5 | suite_bulk_oof_hardened |
| LUAD | 25 | *PTPRD* | 0.661 | – | 0.243 | 567 | 5 | suite_bulk_oof_hardened |
| LUAD | 25 | *RB1* | 0.802 | – | 0.317 | 567 | 5 | suite_bulk_oof_hardened |
| LUAD | 25 | *RBM10* | 0.548 | – | 0.076 | 567 | 5 | suite_bulk_oof_hardened |
| LUAD | 25 | *SETD2* | 0.632 | – | 0.147 | 567 | 5 | suite_bulk_oof_hardened |
| LUAD | 25 | *SMARCA4* | 0.796 | – | 0.311 | 567 | 5 | suite_bulk_oof_hardened |
| LUAD | 25 | *STK11* | 0.879 | – | 0.435 | 567 | 5 | suite_bulk_oof_hardened |
| LUAD | 25 | *TP53* | 0.886 | – | 0.837 | 567 | 5 | suite_bulk_oof_hardened |
| LUAD | 25 | *U2AF1* | 0.738 | – | 0.064 | 567 | 5 | suite_bulk_oof_hardened |
| LUAD | 50 | *ARID1A* | 0.398 | – | 0.046 | 567 | 5 | suite_bulk_oof_hardened |
| LUAD | 50 | *ATM* | 0.671 | – | 0.193 | 567 | 5 | suite_bulk_oof_hardened |
| LUAD | 50 | *BRAF* | 0.623 | – | 0.117 | 567 | 5 | suite_bulk_oof_hardened |
| LUAD | 50 | *CDKN2A* | 0.358 | – | 0.028 | 567 | 5 | suite_bulk_oof_hardened |

Continued on next page

*Supplementary Table S9 continued*

| Cancer | $k$ | Driver | AUROC mean | AUROC SD | AUPRC mean | Samples | CV folds | Implementation |
|---|---|---|---|---|---|---|---|---|
| LUAD | 50 | *EGFR* | 0.829 | – | 0.506 | 567 | 5 | suite_bulk_oof_hardened |
| LUAD | 50 | *ERBB2* | 0.781 | – | 0.102 | 454 | 5 | suite_bulk_oof_hardened |
| LUAD | 50 | *KEAP1* | 0.907 | – | 0.697 | 567 | 5 | suite_bulk_oof_hardened |
| LUAD | 50 | *KRAS* | 0.833 | – | 0.637 | 567 | 5 | suite_bulk_oof_hardened |
| LUAD | 50 | *MET* | 0.687 | – | 0.086 | 567 | 5 | suite_bulk_oof_hardened |
| LUAD | 50 | *MGA* | 0.646 | – | 0.175 | 567 | 5 | suite_bulk_oof_hardened |
| LUAD | 50 | *NF1* | 0.660 | – | 0.166 | 567 | 5 | suite_bulk_oof_hardened |
| LUAD | 50 | *PIK3CA* | 0.603 | – | 0.075 | 567 | 5 | suite_bulk_oof_hardened |
| LUAD | 50 | *PTPRD* | 0.672 | – | 0.274 | 567 | 5 | suite_bulk_oof_hardened |
| LUAD | 50 | *RB1* | 0.792 | – | 0.336 | 567 | 5 | suite_bulk_oof_hardened |
| LUAD | 50 | *RBM10* | 0.520 | – | 0.070 | 567 | 5 | suite_bulk_oof_hardened |
| LUAD | 50 | *SETD2* | 0.675 | – | 0.233 | 567 | 5 | suite_bulk_oof_hardened |
| LUAD | 50 | *SMARCA4* | 0.837 | – | 0.318 | 567 | 5 | suite_bulk_oof_hardened |
| LUAD | 50 | *STK11* | 0.846 | – | 0.401 | 567 | 5 | suite_bulk_oof_hardened |
| LUAD | 50 | *TP53* | 0.880 | – | 0.827 | 567 | 5 | suite_bulk_oof_hardened |
| LUAD | 50 | *U2AF1* | 0.806 | – | 0.116 | 567 | 5 | suite_bulk_oof_hardened |
| LUAD | 75 | *ARID1A* | 0.399 | – | 0.045 | 567 | 5 | suite_bulk_oof_hardened |
| LUAD | 75 | *ATM* | 0.668 | – | 0.227 | 567 | 5 | suite_bulk_oof_hardened |
| LUAD | 75 | *BRAF* | 0.613 | – | 0.145 | 567 | 5 | suite_bulk_oof_hardened |
| LUAD | 75 | *CDKN2A* | 0.371 | – | 0.030 | 567 | 5 | suite_bulk_oof_hardened |
| LUAD | 75 | *EGFR* | 0.804 | – | 0.516 | 567 | 5 | suite_bulk_oof_hardened |
| LUAD | 75 | *ERBB2* | 0.723 | – | 0.044 | 454 | 5 | suite_bulk_oof_hardened |
| LUAD | 75 | *KEAP1* | 0.896 | – | 0.698 | 567 | 5 | suite_bulk_oof_hardened |
| LUAD | 75 | *KRAS* | 0.858 | – | 0.679 | 567 | 5 | suite_bulk_oof_hardened |
| LUAD | 75 | *MET* | 0.669 | – | 0.077 | 567 | 5 | suite_bulk_oof_hardened |
| LUAD | 75 | *MGA* | 0.572 | – | 0.137 | 567 | 5 | suite_bulk_oof_hardened |
| LUAD | 75 | *NF1* | 0.602 | – | 0.144 | 567 | 5 | suite_bulk_oof_hardened |
| LUAD | 75 | *PIK3CA* | 0.485 | – | 0.048 | 567 | 5 | suite_bulk_oof_hardened |
| LUAD | 75 | *PTPRD* | 0.620 | – | 0.248 | 567 | 5 | suite_bulk_oof_hardened |
| LUAD | 75 | *RB1* | 0.767 | – | 0.248 | 567 | 5 | suite_bulk_oof_hardened |
| LUAD | 75 | *RBM10* | 0.507 | – | 0.098 | 567 | 5 | suite_bulk_oof_hardened |
| LUAD | 75 | *SETD2* | 0.644 | – | 0.239 | 567 | 5 | suite_bulk_oof_hardened |
| LUAD | 75 | *SMARCA4* | 0.821 | – | 0.302 | 567 | 5 | suite_bulk_oof_hardened |
| LUAD | 75 | *STK11* | 0.840 | – | 0.375 | 567 | 5 | suite_bulk_oof_hardened |
| LUAD | 75 | *TP53* | 0.887 | – | 0.832 | 567 | 5 | suite_bulk_oof_hardened |
| LUAD | 75 | *U2AF1* | 0.839 | – | 0.325 | 567 | 5 | suite_bulk_oof_hardened |
| LUAD | 100 | *ARID1A* | 0.433 | – | 0.059 | 567 | 5 | suite_bulk_oof_hardened |
| LUAD | 100 | *ATM* | 0.675 | – | 0.229 | 567 | 5 | suite_bulk_oof_hardened |
| LUAD | 100 | *BRAF* | 0.563 | – | 0.091 | 567 | 5 | suite_bulk_oof_hardened |
| LUAD | 100 | *CDKN2A* | 0.431 | – | 0.031 | 567 | 5 | suite_bulk_oof_hardened |
| LUAD | 100 | *EGFR* | 0.793 | – | 0.508 | 567 | 5 | suite_bulk_oof_hardened |
| LUAD | 100 | *ERBB2* | 0.733 | – | 0.090 | 454 | 5 | suite_bulk_oof_hardened |
| LUAD | 100 | *KEAP1* | 0.888 | – | 0.689 | 567 | 5 | suite_bulk_oof_hardened |
| LUAD | 100 | *KRAS* | 0.854 | – | 0.641 | 567 | 5 | suite_bulk_oof_hardened |
| LUAD | 100 | *MET* | 0.605 | – | 0.081 | 567 | 5 | suite_bulk_oof_hardened |
| LUAD | 100 | *MGA* | 0.598 | – | 0.139 | 567 | 5 | suite_bulk_oof_hardened |
| LUAD | 100 | *NF1* | 0.595 | – | 0.153 | 567 | 5 | suite_bulk_oof_hardened |
| LUAD | 100 | *PIK3CA* | 0.504 | – | 0.050 | 567 | 5 | suite_bulk_oof_hardened |
| LUAD | 100 | *PTPRD* | 0.624 | – | 0.254 | 567 | 5 | suite_bulk_oof_hardened |
| LUAD | 100 | *RB1* | 0.715 | – | 0.234 | 567 | 5 | suite_bulk_oof_hardened |
| LUAD | 100 | *RBM10* | 0.527 | – | 0.146 | 567 | 5 | suite_bulk_oof_hardened |
| LUAD | 100 | *SETD2* | 0.670 | – | 0.273 | 567 | 5 | suite_bulk_oof_hardened |
| LUAD | 100 | *SMARCA4* | 0.823 | – | 0.352 | 567 | 5 | suite_bulk_oof_hardened |
| LUAD | 100 | *STK11* | 0.818 | – | 0.349 | 567 | 5 | suite_bulk_oof_hardened |
| LUAD | 100 | *TP53* | 0.885 | – | 0.832 | 567 | 5 | suite_bulk_oof_hardened |
| LUAD | 100 | *U2AF1* | 0.869 | – | 0.403 | 567 | 5 | suite_bulk_oof_hardened |
| LUAD | 150 | *ARID1A* | 0.522 | – | 0.083 | 567 | 5 | suite_bulk_oof_hardened |
| LUAD | 150 | *ATM* | 0.653 | – | 0.215 | 567 | 5 | suite_bulk_oof_hardened |
| LUAD | 150 | *BRAF* | 0.593 | – | 0.114 | 567 | 5 | suite_bulk_oof_hardened |
| LUAD | 150 | *CDKN2A* | 0.456 | – | 0.035 | 567 | 5 | suite_bulk_oof_hardened |
| LUAD | 150 | *EGFR* | 0.826 | – | 0.527 | 567 | 5 | suite_bulk_oof_hardened |
| LUAD | 150 | *ERBB2* | 0.739 | – | 0.095 | 454 | 5 | suite_bulk_oof_hardened |
| LUAD | 150 | *KEAP1* | 0.900 | – | 0.720 | 567 | 5 | suite_bulk_oof_hardened |
| LUAD | 150 | *KRAS* | 0.853 | – | 0.668 | 567 | 5 | suite_bulk_oof_hardened |
| LUAD | 150 | *MET* | 0.580 | – | 0.074 | 567 | 5 | suite_bulk_oof_hardened |
| LUAD | 150 | *MGA* | 0.619 | – | 0.152 | 567 | 5 | suite_bulk_oof_hardened |
| LUAD | 150 | *NF1* | 0.546 | – | 0.129 | 567 | 5 | suite_bulk_oof_hardened |
| LUAD | 150 | *PIK3CA* | 0.499 | – | 0.050 | 567 | 5 | suite_bulk_oof_hardened |
| LUAD | 150 | *PTPRD* | 0.613 | – | 0.277 | 567 | 5 | suite_bulk_oof_hardened |
| LUAD | 150 | *RB1* | 0.780 | – | 0.374 | 567 | 5 | suite_bulk_oof_hardened |
| LUAD | 150 | *RBM10* | 0.643 | – | 0.162 | 567 | 5 | suite_bulk_oof_hardened |
| LUAD | 150 | *SETD2* | 0.762 | – | 0.387 | 567 | 5 | suite_bulk_oof_hardened |
| LUAD | 150 | *SMARCA4* | 0.835 | – | 0.391 | 567 | 5 | suite_bulk_oof_hardened |

Continued on next page

*Supplementary Table S9 continued*

| Cancer | *k* | Driver | AUROC mean | AUROC SD | AUPRC mean | Samples | CV folds | Implementation |
|---|---|---|---|---|---|---|---|---|
| LUAD | 150 | *STK11* | 0.814 | – | 0.368 | 567 | 5 | suite_bulk_oof_hardened |
| LUAD | 150 | *TP53* | 0.880 | – | 0.831 | 567 | 5 | suite_bulk_oof_hardened |
| LUAD | 150 | *U2AF1* | 0.917 | – | 0.580 | 567 | 5 | suite_bulk_oof_hardened |
| LUAD | 250 | *ARID1A* | 0.569 | – | 0.119 | 567 | 5 | suite_bulk_oof_hardened |
| LUAD | 250 | *ATM* | 0.650 | – | 0.236 | 567 | 5 | suite_bulk_oof_hardened |
| LUAD | 250 | *BRAF* | 0.632 | – | 0.130 | 567 | 5 | suite_bulk_oof_hardened |
| LUAD | 250 | *CDKN2A* | 0.445 | – | 0.032 | 567 | 5 | suite_bulk_oof_hardened |
| LUAD | 250 | *EGFR* | 0.837 | – | 0.553 | 567 | 5 | suite_bulk_oof_hardened |
| LUAD | 250 | *ERBB2* | 0.729 | – | 0.051 | 454 | 5 | suite_bulk_oof_hardened |
| LUAD | 250 | *KEAP1* | 0.916 | – | 0.755 | 567 | 5 | suite_bulk_oof_hardened |
| LUAD | 250 | *KRAS* | 0.859 | – | 0.650 | 567 | 5 | suite_bulk_oof_hardened |
| LUAD | 250 | *MET* | 0.610 | – | 0.057 | 567 | 5 | suite_bulk_oof_hardened |
| LUAD | 250 | *MGA* | 0.653 | – | 0.173 | 567 | 5 | suite_bulk_oof_hardened |
| LUAD | 250 | *NF1* | 0.590 | – | 0.133 | 567 | 5 | suite_bulk_oof_hardened |
| LUAD | 250 | *PIK3CA* | 0.530 | – | 0.063 | 567 | 5 | suite_bulk_oof_hardened |
| LUAD | 250 | *PTPRD* | 0.657 | – | 0.276 | 567 | 5 | suite_bulk_oof_hardened |
| LUAD | 250 | *RB1* | 0.813 | – | 0.396 | 567 | 5 | suite_bulk_oof_hardened |
| LUAD | 250 | *RBM10* | 0.732 | – | 0.282 | 567 | 5 | suite_bulk_oof_hardened |
| LUAD | 250 | *SETD2* | 0.782 | – | 0.447 | 567 | 5 | suite_bulk_oof_hardened |
| LUAD | 250 | *SMARCA4* | 0.851 | – | 0.495 | 567 | 5 | suite_bulk_oof_hardened |
| LUAD | 250 | *STK11* | 0.821 | – | 0.355 | 567 | 5 | suite_bulk_oof_hardened |
| LUAD | 250 | *TP53* | 0.901 | – | 0.860 | 567 | 5 | suite_bulk_oof_hardened |
| LUAD | 250 | *U2AF1* | 0.944 | – | 0.752 | 567 | 5 | suite_bulk_oof_hardened |
| LUAD | 500 | *ARID1A* | 0.623 | – | 0.185 | 567 | 5 | suite_bulk_oof_hardened |
| LUAD | 500 | *ATM* | 0.692 | – | 0.239 | 567 | 5 | suite_bulk_oof_hardened |
| LUAD | 500 | *BRAF* | 0.660 | – | 0.134 | 567 | 5 | suite_bulk_oof_hardened |
| LUAD | 500 | *CDKN2A* | 0.447 | – | 0.038 | 567 | 5 | suite_bulk_oof_hardened |
| LUAD | 500 | *EGFR* | 0.852 | – | 0.586 | 567 | 5 | suite_bulk_oof_hardened |
| LUAD | 500 | *ERBB2* | 0.695 | – | 0.026 | 454 | 5 | suite_bulk_oof_hardened |
| LUAD | 500 | *KEAP1* | 0.928 | – | 0.742 | 567 | 5 | suite_bulk_oof_hardened |
| LUAD | 500 | *KRAS* | 0.883 | – | 0.703 | 567 | 5 | suite_bulk_oof_hardened |
| LUAD | 500 | *MET* | 0.713 | – | 0.093 | 567 | 5 | suite_bulk_oof_hardened |
| LUAD | 500 | *MGA* | 0.684 | – | 0.201 | 567 | 5 | suite_bulk_oof_hardened |
| LUAD | 500 | *NF1* | 0.589 | – | 0.134 | 567 | 5 | suite_bulk_oof_hardened |
| LUAD | 500 | *PIK3CA* | 0.571 | – | 0.076 | 567 | 5 | suite_bulk_oof_hardened |
| LUAD | 500 | *PTPRD* | 0.701 | – | 0.321 | 567 | 5 | suite_bulk_oof_hardened |
| LUAD | 500 | *RB1* | 0.800 | – | 0.365 | 567 | 5 | suite_bulk_oof_hardened |
| LUAD | 500 | *RBM10* | 0.834 | – | 0.390 | 567 | 5 | suite_bulk_oof_hardened |
| LUAD | 500 | *SETD2* | 0.837 | – | 0.549 | 567 | 5 | suite_bulk_oof_hardened |
| LUAD | 500 | *SMARCA4* | 0.859 | – | 0.558 | 567 | 5 | suite_bulk_oof_hardened |
| LUAD | 500 | *STK11* | 0.841 | – | 0.376 | 567 | 5 | suite_bulk_oof_hardened |
| LUAD | 500 | *TP53* | 0.910 | – | 0.874 | 567 | 5 | suite_bulk_oof_hardened |
| LUAD | 500 | *U2AF1* | 0.955 | – | 0.778 | 567 | 5 | suite_bulk_oof_hardened |
| PAAD | 5 | *ATM* | 0.416 | – | 0.040 | 172 | 2 | suite_bulk_oof_hardened |
| PAAD | 5 | *CDKN2A* | 0.587 | – | 0.286 | 172 | 2 | suite_bulk_oof_hardened |
| PAAD | 5 | *GNAS* | 0.558 | – | 0.188 | 172 | 2 | suite_bulk_oof_hardened |
| PAAD | 5 | *KDM6A* | 0.670 | – | 0.081 | 172 | 2 | suite_bulk_oof_hardened |
| PAAD | 5 | *KMT2D* | 0.533 | – | 0.056 | 172 | 2 | suite_bulk_oof_hardened |
| PAAD | 5 | *KRAS* | 0.831 | – | 0.882 | 172 | 2 | suite_bulk_oof_hardened |
| PAAD | 5 | *MAP2K4* | 0.644 | – | 0.136 | 172 | 2 | suite_bulk_oof_hardened |
| PAAD | 5 | *RNF43* | 0.538 | – | 0.119 | 172 | 2 | suite_bulk_oof_hardened |
| PAAD | 5 | *ROBO2* | 0.519 | – | 0.104 | 172 | 2 | suite_bulk_oof_hardened |
| PAAD | 5 | *SMAD4* | 0.681 | – | 0.355 | 172 | 2 | suite_bulk_oof_hardened |
| PAAD | 5 | *STK11* | 0.217 | – | 0.018 | 172 | 2 | suite_bulk_oof_hardened |
| PAAD | 5 | *TGFBR2* | 0.447 | – | 0.045 | 172 | 2 | suite_bulk_oof_hardened |
| PAAD | 5 | *TP53* | 0.737 | – | 0.784 | 172 | 2 | suite_bulk_oof_hardened |
| PAAD | 10 | *ATM* | 0.436 | – | 0.110 | 172 | 2 | suite_bulk_oof_hardened |
| PAAD | 10 | *CDKN2A* | 0.593 | – | 0.229 | 172 | 2 | suite_bulk_oof_hardened |
| PAAD | 10 | *GNAS* | 0.714 | – | 0.283 | 172 | 2 | suite_bulk_oof_hardened |
| PAAD | 10 | *KDM6A* | 0.707 | – | 0.084 | 172 | 2 | suite_bulk_oof_hardened |
| PAAD | 10 | *KMT2D* | 0.572 | – | 0.065 | 172 | 2 | suite_bulk_oof_hardened |
| PAAD | 10 | *KRAS* | 0.881 | – | 0.931 | 172 | 2 | suite_bulk_oof_hardened |
| PAAD | 10 | *MAP2K4* | 0.659 | – | 0.149 | 172 | 2 | suite_bulk_oof_hardened |
| PAAD | 10 | *RNF43* | 0.511 | – | 0.083 | 172 | 2 | suite_bulk_oof_hardened |
| PAAD | 10 | *ROBO2* | 0.501 | – | 0.034 | 172 | 2 | suite_bulk_oof_hardened |
| PAAD | 10 | *SMAD4* | 0.703 | – | 0.397 | 172 | 2 | suite_bulk_oof_hardened |
| PAAD | 10 | *STK11* | 0.314 | – | 0.020 | 172 | 2 | suite_bulk_oof_hardened |
| PAAD | 10 | *TGFBR2* | 0.627 | – | 0.066 | 172 | 2 | suite_bulk_oof_hardened |
| PAAD | 10 | *TP53* | 0.793 | – | 0.832 | 172 | 2 | suite_bulk_oof_hardened |
| PAAD | 25 | *ATM* | 0.440 | – | 0.040 | 172 | 2 | suite_bulk_oof_hardened |
| PAAD | 25 | *CDKN2A* | 0.572 | – | 0.222 | 172 | 2 | suite_bulk_oof_hardened |
| PAAD | 25 | *GNAS* | 0.801 | – | 0.192 | 172 | 2 | suite_bulk_oof_hardened |

Continued on next page

*Supplementary Table S9 continued*

| Cancer | *k* | Driver | AUROC mean | AUROC SD | AUPRC mean | Samples | CV folds | Implementation |
|---|---|---|---|---|---|---|---|---|
| PAAD | 25 | *KDM6A* | 0.775 | – | 0.116 | 172 | 2 | `suite_bulk_oof_hardened` |
| PAAD | 25 | *KMT2D* | 0.547 | – | 0.058 | 172 | 2 | `suite_bulk_oof_hardened` |
| PAAD | 25 | *KRAS* | 0.908 | – | 0.946 | 172 | 2 | `suite_bulk_oof_hardened` |
| PAAD | 25 | *MAP2K4* | 0.628 | – | 0.072 | 172 | 2 | `suite_bulk_oof_hardened` |
| PAAD | 25 | *RNF43* | 0.589 | – | 0.136 | 172 | 2 | `suite_bulk_oof_hardened` |
| PAAD | 25 | *ROBO2* | 0.366 | – | 0.022 | 172 | 2 | `suite_bulk_oof_hardened` |
| PAAD | 25 | *SMAD4* | 0.710 | – | 0.367 | 172 | 2 | `suite_bulk_oof_hardened` |
| PAAD | 25 | *STK11* | 0.232 | – | 0.018 | 172 | 2 | `suite_bulk_oof_hardened` |
| PAAD | 25 | *TGFBR2* | 0.527 | – | 0.052 | 172 | 2 | `suite_bulk_oof_hardened` |
| PAAD | 25 | *TP53* | 0.784 | – | 0.821 | 172 | 2 | `suite_bulk_oof_hardened` |
| PAAD | 50 | *ATM* | 0.556 | – | 0.055 | 172 | 2 | `suite_bulk_oof_hardened` |
| PAAD | 50 | *CDKN2A* | 0.496 | – | 0.195 | 172 | 2 | `suite_bulk_oof_hardened` |
| PAAD | 50 | *GNAS* | 0.813 | – | 0.292 | 172 | 2 | `suite_bulk_oof_hardened` |
| PAAD | 50 | *KDM6A* | 0.674 | – | 0.061 | 172 | 2 | `suite_bulk_oof_hardened` |
| PAAD | 50 | *KMT2D* | 0.623 | – | 0.088 | 172 | 2 | `suite_bulk_oof_hardened` |
| PAAD | 50 | *KRAS* | 0.894 | – | 0.941 | 172 | 2 | `suite_bulk_oof_hardened` |
| PAAD | 50 | *MAP2K4* | 0.595 | – | 0.095 | 172 | 2 | `suite_bulk_oof_hardened` |
| PAAD | 50 | *RNF43* | 0.711 | – | 0.247 | 172 | 2 | `suite_bulk_oof_hardened` |
| PAAD | 50 | *ROBO2* | 0.500 | – | 0.028 | 172 | 2 | `suite_bulk_oof_hardened` |
| PAAD | 50 | *SMAD4* | 0.685 | – | 0.361 | 172 | 2 | `suite_bulk_oof_hardened` |
| PAAD | 50 | *STK11* | 0.417 | – | 0.026 | 172 | 2 | `suite_bulk_oof_hardened` |
| PAAD | 50 | *TGFBR2* | 0.514 | – | 0.050 | 172 | 2 | `suite_bulk_oof_hardened` |
| PAAD | 50 | *TP53* | 0.788 | – | 0.829 | 172 | 2 | `suite_bulk_oof_hardened` |
| PAAD | 75 | *ATM* | 0.567 | – | 0.055 | 172 | 2 | `suite_bulk_oof_hardened` |
| PAAD | 75 | *CDKN2A* | 0.543 | – | 0.213 | 172 | 2 | `suite_bulk_oof_hardened` |
| PAAD | 75 | *GNAS* | 0.771 | – | 0.430 | 172 | 2 | `suite_bulk_oof_hardened` |
| PAAD | 75 | *KDM6A* | 0.704 | – | 0.090 | 172 | 2 | `suite_bulk_oof_hardened` |
| PAAD | 75 | *KMT2D* | 0.515 | – | 0.047 | 172 | 2 | `suite_bulk_oof_hardened` |
| PAAD | 75 | *KRAS* | 0.890 | – | 0.932 | 172 | 2 | `suite_bulk_oof_hardened` |
| PAAD | 75 | *MAP2K4* | 0.571 | – | 0.065 | 172 | 2 | `suite_bulk_oof_hardened` |
| PAAD | 75 | *RNF43* | 0.710 | – | 0.172 | 172 | 2 | `suite_bulk_oof_hardened` |
| PAAD | 75 | *ROBO2* | 0.452 | – | 0.025 | 172 | 2 | `suite_bulk_oof_hardened` |
| PAAD | 75 | *SMAD4* | 0.691 | – | 0.403 | 172 | 2 | `suite_bulk_oof_hardened` |
| PAAD | 75 | *STK11* | 0.414 | – | 0.026 | 172 | 2 | `suite_bulk_oof_hardened` |
| PAAD | 75 | *TGFBR2* | 0.574 | – | 0.056 | 172 | 2 | `suite_bulk_oof_hardened` |
| PAAD | 75 | *TP53* | 0.797 | – | 0.835 | 172 | 2 | `suite_bulk_oof_hardened` |
| PAAD | 100 | *ATM* | 0.558 | – | 0.053 | 172 | 2 | `suite_bulk_oof_hardened` |
| PAAD | 100 | *CDKN2A* | 0.555 | – | 0.225 | 172 | 2 | `suite_bulk_oof_hardened` |
| PAAD | 100 | *GNAS* | 0.785 | – | 0.426 | 172 | 2 | `suite_bulk_oof_hardened` |
| PAAD | 100 | *KDM6A* | 0.701 | – | 0.082 | 172 | 2 | `suite_bulk_oof_hardened` |
| PAAD | 100 | *KMT2D* | 0.514 | – | 0.048 | 172 | 2 | `suite_bulk_oof_hardened` |
| PAAD | 100 | *KRAS* | 0.884 | – | 0.930 | 172 | 2 | `suite_bulk_oof_hardened` |
| PAAD | 100 | *MAP2K4* | 0.533 | – | 0.052 | 172 | 2 | `suite_bulk_oof_hardened` |
| PAAD | 100 | *RNF43* | 0.704 | – | 0.165 | 172 | 2 | `suite_bulk_oof_hardened` |
| PAAD | 100 | *ROBO2* | 0.485 | – | 0.032 | 172 | 2 | `suite_bulk_oof_hardened` |
| PAAD | 100 | *SMAD4* | 0.693 | – | 0.399 | 172 | 2 | `suite_bulk_oof_hardened` |
| PAAD | 100 | *STK11* | 0.391 | – | 0.024 | 172 | 2 | `suite_bulk_oof_hardened` |
| PAAD | 100 | *TGFBR2* | 0.534 | – | 0.052 | 172 | 2 | `suite_bulk_oof_hardened` |
| PAAD | 100 | *TP53* | 0.805 | – | 0.842 | 172 | 2 | `suite_bulk_oof_hardened` |
| PAAD | 150 | *ATM* | 0.558 | – | 0.053 | 172 | 2 | `suite_bulk_oof_hardened` |
| PAAD | 150 | *CDKN2A* | 0.555 | – | 0.225 | 172 | 2 | `suite_bulk_oof_hardened` |
| PAAD | 150 | *GNAS* | 0.785 | – | 0.426 | 172 | 2 | `suite_bulk_oof_hardened` |
| PAAD | 150 | *KDM6A* | 0.701 | – | 0.082 | 172 | 2 | `suite_bulk_oof_hardened` |
| PAAD | 150 | *KMT2D* | 0.514 | – | 0.048 | 172 | 2 | `suite_bulk_oof_hardened` |
| PAAD | 150 | *KRAS* | 0.884 | – | 0.930 | 172 | 2 | `suite_bulk_oof_hardened` |
| PAAD | 150 | *MAP2K4* | 0.533 | – | 0.052 | 172 | 2 | `suite_bulk_oof_hardened` |
| PAAD | 150 | *RNF43* | 0.704 | – | 0.165 | 172 | 2 | `suite_bulk_oof_hardened` |
| PAAD | 150 | *ROBO2* | 0.485 | – | 0.032 | 172 | 2 | `suite_bulk_oof_hardened` |
| PAAD | 150 | *SMAD4* | 0.693 | – | 0.399 | 172 | 2 | `suite_bulk_oof_hardened` |
| PAAD | 150 | *STK11* | 0.391 | – | 0.024 | 172 | 2 | `suite_bulk_oof_hardened` |
| PAAD | 150 | *TGFBR2* | 0.534 | – | 0.052 | 172 | 2 | `suite_bulk_oof_hardened` |
| PAAD | 150 | *TP53* | 0.805 | – | 0.842 | 172 | 2 | `suite_bulk_oof_hardened` |
| SKCM | 5 | *AKT1* | 0.567 | – | 0.013 | 372 | 5 | `suite_bulk_oof_hardened` |
| SKCM | 5 | *AKT3* | 0.355 | – | 0.015 | 466 | 5 | `suite_bulk_oof_hardened` |
| SKCM | 5 | *APC* | 0.546 | – | 0.106 | 466 | 5 | `suite_bulk_oof_hardened` |
| SKCM | 5 | *ARID1A* | 0.526 | – | 0.064 | 466 | 5 | `suite_bulk_oof_hardened` |
| SKCM | 5 | *ARID2* | 0.480 | – | 0.119 | 466 | 5 | `suite_bulk_oof_hardened` |
| SKCM | 5 | *BRAF* | 0.605 | – | 0.606 | 466 | 5 | `suite_bulk_oof_hardened` |
| SKCM | 5 | *CDKN2A* | 0.516 | – | 0.132 | 466 | 5 | `suite_bulk_oof_hardened` |
| SKCM | 5 | *CTNNB1* | 0.422 | – | 0.053 | 466 | 5 | `suite_bulk_oof_hardened` |
| SKCM | 5 | *GNA11* | 0.164 | – | 0.016 | 466 | 5 | `suite_bulk_oof_hardened` |
| SKCM | 5 | *GNAQ* | 0.668 | – | 0.051 | 466 | 5 | `suite_bulk_oof_hardened` |

Continued on next page

*Supplementary Table S9 continued*

| Cancer | *k* | Driver | AUROC mean | AUROC SD | AUPRC mean | Samples | CV folds | Implementation |
|---|---|---|---|---|---|---|---|---|
| SKCM | 5 | *HRAS* | 0.251 | – | 0.010 | 466 | 5 | suite_bulk_oof_hardened |
| SKCM | 5 | *IDH1* | 0.654 | – | 0.123 | 466 | 5 | suite_bulk_oof_hardened |
| SKCM | 5 | *KDM6A* | 0.029 | – | 0.004 | 280 | 5 | suite_bulk_oof_hardened |
| SKCM | 5 | *KIT* | 0.665 | – | 0.133 | 466 | 5 | suite_bulk_oof_hardened |
| SKCM | 5 | *KRAS* | 0.498 | – | 0.024 | 466 | 5 | suite_bulk_oof_hardened |
| SKCM | 5 | *MAP2K1* | 0.538 | – | 0.075 | 466 | 5 | suite_bulk_oof_hardened |
| SKCM | 5 | *MAP2K2* | 0.451 | – | 0.019 | 466 | 5 | suite_bulk_oof_hardened |
| SKCM | 5 | *NF1* | 0.429 | – | 0.122 | 466 | 5 | suite_bulk_oof_hardened |
| SKCM | 5 | *NRAS* | 0.590 | – | 0.359 | 466 | 5 | suite_bulk_oof_hardened |
| SKCM | 5 | *PIK3CA* | 0.648 | – | 0.056 | 466 | 5 | suite_bulk_oof_hardened |
| SKCM | 5 | *PPP6C* | 0.552 | – | 0.081 | 466 | 5 | suite_bulk_oof_hardened |
| SKCM | 5 | *PREX2* | 0.606 | – | 0.280 | 466 | 5 | suite_bulk_oof_hardened |
| SKCM | 5 | *PTEN* | 0.573 | – | 0.119 | 466 | 5 | suite_bulk_oof_hardened |
| SKCM | 5 | *RAC1* | 0.473 | – | 0.081 | 466 | 5 | suite_bulk_oof_hardened |
| SKCM | 5 | *SETD2* | 0.641 | – | 0.094 | 466 | 5 | suite_bulk_oof_hardened |
| SKCM | 5 | *TP53* | 0.653 | – | 0.228 | 466 | 5 | suite_bulk_oof_hardened |
| SKCM | 10 | *AKT1* | 0.338 | – | 0.008 | 372 | 5 | suite_bulk_oof_hardened |
| SKCM | 10 | *AKT3* | 0.489 | – | 0.020 | 466 | 5 | suite_bulk_oof_hardened |
| SKCM | 10 | *APC* | 0.633 | – | 0.149 | 466 | 5 | suite_bulk_oof_hardened |
| SKCM | 10 | *ARID1A* | 0.523 | – | 0.055 | 466 | 5 | suite_bulk_oof_hardened |
| SKCM | 10 | *ARID2* | 0.553 | – | 0.154 | 466 | 5 | suite_bulk_oof_hardened |
| SKCM | 10 | *BRAF* | 0.755 | – | 0.777 | 466 | 5 | suite_bulk_oof_hardened |
| SKCM | 10 | *CDKN2A* | 0.526 | – | 0.143 | 466 | 5 | suite_bulk_oof_hardened |
| SKCM | 10 | *CTNNB1* | 0.675 | – | 0.164 | 466 | 5 | suite_bulk_oof_hardened |
| SKCM | 10 | *GNA11* | 0.329 | – | 0.019 | 466 | 5 | suite_bulk_oof_hardened |
| SKCM | 10 | *GNAQ* | 0.662 | – | 0.081 | 466 | 5 | suite_bulk_oof_hardened |
| SKCM | 10 | *HRAS* | 0.263 | – | 0.010 | 466 | 5 | suite_bulk_oof_hardened |
| SKCM | 10 | *IDH1* | 0.731 | – | 0.169 | 466 | 5 | suite_bulk_oof_hardened |
| SKCM | 10 | *KDM6A* | 0.079 | – | 0.004 | 280 | 5 | suite_bulk_oof_hardened |
| SKCM | 10 | *KIT* | 0.764 | – | 0.192 | 466 | 5 | suite_bulk_oof_hardened |
| SKCM | 10 | *KRAS* | 0.414 | – | 0.022 | 466 | 5 | suite_bulk_oof_hardened |
| SKCM | 10 | *MAP2K1* | 0.513 | – | 0.065 | 466 | 5 | suite_bulk_oof_hardened |
| SKCM | 10 | *MAP2K2* | 0.344 | – | 0.015 | 466 | 5 | suite_bulk_oof_hardened |
| SKCM | 10 | *NF1* | 0.596 | – | 0.177 | 466 | 5 | suite_bulk_oof_hardened |
| SKCM | 10 | *NRAS* | 0.650 | – | 0.386 | 466 | 5 | suite_bulk_oof_hardened |
| SKCM | 10 | *PIK3CA* | 0.670 | – | 0.057 | 466 | 5 | suite_bulk_oof_hardened |
| SKCM | 10 | *PPP6C* | 0.485 | – | 0.062 | 466 | 5 | suite_bulk_oof_hardened |
| SKCM | 10 | *PREX2* | 0.594 | – | 0.275 | 466 | 5 | suite_bulk_oof_hardened |
| SKCM | 10 | *PTEN* | 0.672 | – | 0.186 | 466 | 5 | suite_bulk_oof_hardened |
| SKCM | 10 | *RAC1* | 0.557 | – | 0.120 | 466 | 5 | suite_bulk_oof_hardened |
| SKCM | 10 | *SETD2* | 0.588 | – | 0.080 | 466 | 5 | suite_bulk_oof_hardened |
| SKCM | 10 | *TP53* | 0.736 | – | 0.311 | 466 | 5 | suite_bulk_oof_hardened |
| SKCM | 25 | *AKT1* | 0.570 | – | 0.013 | 372 | 5 | suite_bulk_oof_hardened |
| SKCM | 25 | *AKT3* | 0.311 | – | 0.013 | 466 | 5 | suite_bulk_oof_hardened |
| SKCM | 25 | *APC* | 0.592 | – | 0.131 | 466 | 5 | suite_bulk_oof_hardened |
| SKCM | 25 | *ARID1A* | 0.487 | – | 0.056 | 466 | 5 | suite_bulk_oof_hardened |
| SKCM | 25 | *ARID2* | 0.606 | – | 0.252 | 466 | 5 | suite_bulk_oof_hardened |
| SKCM | 25 | *BRAF* | 0.820 | – | 0.836 | 466 | 5 | suite_bulk_oof_hardened |
| SKCM | 25 | *CDKN2A* | 0.596 | – | 0.180 | 466 | 5 | suite_bulk_oof_hardened |
| SKCM | 25 | *CTNNB1* | 0.613 | – | 0.095 | 466 | 5 | suite_bulk_oof_hardened |
| SKCM | 25 | *GNA11* | 0.354 | – | 0.022 | 466 | 5 | suite_bulk_oof_hardened |
| SKCM | 25 | *GNAQ* | 0.727 | – | 0.129 | 466 | 5 | suite_bulk_oof_hardened |
| SKCM | 25 | *HRAS* | 0.420 | – | 0.019 | 466 | 5 | suite_bulk_oof_hardened |
| SKCM | 25 | *IDH1* | 0.642 | – | 0.146 | 466 | 5 | suite_bulk_oof_hardened |
| SKCM | 25 | *KDM6A* | 0.659 | – | 0.010 | 280 | 5 | suite_bulk_oof_hardened |
| SKCM | 25 | *KIT* | 0.777 | – | 0.141 | 466 | 5 | suite_bulk_oof_hardened |
| SKCM | 25 | *KRAS* | 0.436 | – | 0.027 | 466 | 5 | suite_bulk_oof_hardened |
| SKCM | 25 | *MAP2K1* | 0.607 | – | 0.084 | 466 | 5 | suite_bulk_oof_hardened |
| SKCM | 25 | *MAP2K2* | 0.317 | – | 0.015 | 466 | 5 | suite_bulk_oof_hardened |
| SKCM | 25 | *NF1* | 0.697 | – | 0.310 | 466 | 5 | suite_bulk_oof_hardened |
| SKCM | 25 | *NRAS* | 0.828 | – | 0.621 | 466 | 5 | suite_bulk_oof_hardened |
| SKCM | 25 | *PIK3CA* | 0.634 | – | 0.049 | 466 | 5 | suite_bulk_oof_hardened |
| SKCM | 25 | *PPP6C* | 0.460 | – | 0.062 | 466 | 5 | suite_bulk_oof_hardened |
| SKCM | 25 | *PREX2* | 0.600 | – | 0.277 | 466 | 5 | suite_bulk_oof_hardened |
| SKCM | 25 | *PTEN* | 0.734 | – | 0.248 | 466 | 5 | suite_bulk_oof_hardened |
| SKCM | 25 | *RAC1* | 0.583 | – | 0.105 | 466 | 5 | suite_bulk_oof_hardened |
| SKCM | 25 | *SETD2* | 0.583 | – | 0.071 | 466 | 5 | suite_bulk_oof_hardened |
| SKCM | 25 | *TP53* | 0.802 | – | 0.473 | 466 | 5 | suite_bulk_oof_hardened |
| SKCM | 50 | *AKT1* | 0.604 | – | 0.013 | 372 | 5 | suite_bulk_oof_hardened |
| SKCM | 50 | *AKT3* | 0.316 | – | 0.013 | 466 | 5 | suite_bulk_oof_hardened |
| SKCM | 50 | *APC* | 0.534 | – | 0.101 | 466 | 5 | suite_bulk_oof_hardened |
| SKCM | 50 | *ARID1A* | 0.493 | – | 0.056 | 466 | 5 | suite_bulk_oof_hardened |
| SKCM | 50 | *ARID2* | 0.623 | – | 0.265 | 466 | 5 | suite_bulk_oof_hardened |

Continued on next page

*Supplementary Table S9 continued*

| Cancer | $k$ | Driver | AUROC mean | AUROC SD | AUPRC mean | Samples | CV folds | Implementation |
|---|---|---|---|---|---|---|---|---|
| SKCM | 50 | *BRAF* | 0.820 | – | 0.831 | 466 | 5 | suite_bulk_oof_hardened |
| SKCM | 50 | *CDKN2A* | 0.567 | – | 0.159 | 466 | 5 | suite_bulk_oof_hardened |
| SKCM | 50 | *CTNNB1* | 0.630 | – | 0.115 | 466 | 5 | suite_bulk_oof_hardened |
| SKCM | 50 | *GNA11* | 0.279 | – | 0.018 | 466 | 5 | suite_bulk_oof_hardened |
| SKCM | 50 | *GNAQ* | 0.737 | – | 0.117 | 466 | 5 | suite_bulk_oof_hardened |
| SKCM | 50 | *HRAS* | 0.564 | – | 0.023 | 466 | 5 | suite_bulk_oof_hardened |
| SKCM | 50 | *IDH1* | 0.681 | – | 0.174 | 466 | 5 | suite_bulk_oof_hardened |
| SKCM | 50 | *KDM6A* | 0.462 | – | 0.007 | 280 | 5 | suite_bulk_oof_hardened |
| SKCM | 50 | *KIT* | 0.727 | – | 0.144 | 466 | 5 | suite_bulk_oof_hardened |
| SKCM | 50 | *KRAS* | 0.558 | – | 0.086 | 466 | 5 | suite_bulk_oof_hardened |
| SKCM | 50 | *MAP2K1* | 0.615 | – | 0.082 | 466 | 5 | suite_bulk_oof_hardened |
| SKCM | 50 | *MAP2K2* | 0.315 | – | 0.015 | 466 | 5 | suite_bulk_oof_hardened |
| SKCM | 50 | *NF1* | 0.695 | – | 0.287 | 466 | 5 | suite_bulk_oof_hardened |
| SKCM | 50 | *NRAS* | 0.844 | – | 0.682 | 466 | 5 | suite_bulk_oof_hardened |
| SKCM | 50 | *PIK3CA* | 0.598 | – | 0.040 | 466 | 5 | suite_bulk_oof_hardened |
| SKCM | 50 | *PPP6C* | 0.370 | – | 0.057 | 466 | 5 | suite_bulk_oof_hardened |
| SKCM | 50 | *PREX2* | 0.576 | – | 0.250 | 466 | 5 | suite_bulk_oof_hardened |
| SKCM | 50 | *PTEN* | 0.751 | – | 0.268 | 466 | 5 | suite_bulk_oof_hardened |
| SKCM | 50 | *RAC1* | 0.510 | – | 0.066 | 466 | 5 | suite_bulk_oof_hardened |
| SKCM | 50 | *SETD2* | 0.590 | – | 0.085 | 466 | 5 | suite_bulk_oof_hardened |
| SKCM | 50 | *TP53* | 0.806 | – | 0.522 | 466 | 5 | suite_bulk_oof_hardened |
| SKCM | 75 | *AKT1* | 0.589 | – | 0.013 | 372 | 5 | suite_bulk_oof_hardened |
| SKCM | 75 | *AKT3* | 0.375 | – | 0.017 | 466 | 5 | suite_bulk_oof_hardened |
| SKCM | 75 | *APC* | 0.504 | – | 0.099 | 466 | 5 | suite_bulk_oof_hardened |
| SKCM | 75 | *ARID1A* | 0.540 | – | 0.071 | 466 | 5 | suite_bulk_oof_hardened |
| SKCM | 75 | *ARID2* | 0.648 | – | 0.315 | 466 | 5 | suite_bulk_oof_hardened |
| SKCM | 75 | *BRAF* | 0.823 | – | 0.824 | 466 | 5 | suite_bulk_oof_hardened |
| SKCM | 75 | *CDKN2A* | 0.575 | – | 0.179 | 466 | 5 | suite_bulk_oof_hardened |
| SKCM | 75 | *CTNNB1* | 0.647 | – | 0.155 | 466 | 5 | suite_bulk_oof_hardened |
| SKCM | 75 | *GNA11* | 0.425 | – | 0.023 | 466 | 5 | suite_bulk_oof_hardened |
| SKCM | 75 | *GNAQ* | 0.650 | – | 0.069 | 466 | 5 | suite_bulk_oof_hardened |
| SKCM | 75 | *HRAS* | 0.480 | – | 0.017 | 466 | 5 | suite_bulk_oof_hardened |
| SKCM | 75 | *IDH1* | 0.743 | – | 0.192 | 466 | 5 | suite_bulk_oof_hardened |
| SKCM | 75 | *KDM6A* | 0.677 | – | 0.011 | 280 | 5 | suite_bulk_oof_hardened |
| SKCM | 75 | *KIT* | 0.746 | – | 0.149 | 466 | 5 | suite_bulk_oof_hardened |
| SKCM | 75 | *KRAS* | 0.625 | – | 0.081 | 466 | 5 | suite_bulk_oof_hardened |
| SKCM | 75 | *MAP2K1* | 0.624 | – | 0.081 | 466 | 5 | suite_bulk_oof_hardened |
| SKCM | 75 | *MAP2K2* | 0.334 | – | 0.015 | 466 | 5 | suite_bulk_oof_hardened |
| SKCM | 75 | *NF1* | 0.692 | – | 0.294 | 466 | 5 | suite_bulk_oof_hardened |
| SKCM | 75 | *NRAS* | 0.833 | – | 0.661 | 466 | 5 | suite_bulk_oof_hardened |
| SKCM | 75 | *PIK3CA* | 0.698 | – | 0.048 | 466 | 5 | suite_bulk_oof_hardened |
| SKCM | 75 | *PPP6C* | 0.392 | – | 0.055 | 466 | 5 | suite_bulk_oof_hardened |
| SKCM | 75 | *PREX2* | 0.577 | – | 0.255 | 466 | 5 | suite_bulk_oof_hardened |
| SKCM | 75 | *PTEN* | 0.722 | – | 0.228 | 466 | 5 | suite_bulk_oof_hardened |
| SKCM | 75 | *RAC1* | 0.494 | – | 0.060 | 466 | 5 | suite_bulk_oof_hardened |
| SKCM | 75 | *SETD2* | 0.599 | – | 0.130 | 466 | 5 | suite_bulk_oof_hardened |
| SKCM | 75 | *TP53* | 0.813 | – | 0.506 | 466 | 5 | suite_bulk_oof_hardened |
| SKCM | 100 | *AKT1* | 0.492 | – | 0.010 | 372 | 5 | suite_bulk_oof_hardened |
| SKCM | 100 | *AKT3* | 0.518 | – | 0.019 | 466 | 5 | suite_bulk_oof_hardened |
| SKCM | 100 | *APC* | 0.517 | – | 0.102 | 466 | 5 | suite_bulk_oof_hardened |
| SKCM | 100 | *ARID1A* | 0.549 | – | 0.072 | 466 | 5 | suite_bulk_oof_hardened |
| SKCM | 100 | *ARID2* | 0.671 | – | 0.335 | 466 | 5 | suite_bulk_oof_hardened |
| SKCM | 100 | *BRAF* | 0.822 | – | 0.828 | 466 | 5 | suite_bulk_oof_hardened |
| SKCM | 100 | *CDKN2A* | 0.576 | – | 0.175 | 466 | 5 | suite_bulk_oof_hardened |
| SKCM | 100 | *CTNNB1* | 0.647 | – | 0.116 | 466 | 5 | suite_bulk_oof_hardened |
| SKCM | 100 | *GNA11* | 0.475 | – | 0.024 | 466 | 5 | suite_bulk_oof_hardened |
| SKCM | 100 | *GNAQ* | 0.707 | – | 0.088 | 466 | 5 | suite_bulk_oof_hardened |
| SKCM | 100 | *HRAS* | 0.453 | – | 0.017 | 466 | 5 | suite_bulk_oof_hardened |
| SKCM | 100 | *IDH1* | 0.775 | – | 0.313 | 466 | 5 | suite_bulk_oof_hardened |
| SKCM | 100 | *KDM6A* | 0.581 | – | 0.008 | 280 | 5 | suite_bulk_oof_hardened |
| SKCM | 100 | *KIT* | 0.726 | – | 0.173 | 466 | 5 | suite_bulk_oof_hardened |
| SKCM | 100 | *KRAS* | 0.652 | – | 0.045 | 466 | 5 | suite_bulk_oof_hardened |
| SKCM | 100 | *MAP2K1* | 0.635 | – | 0.081 | 466 | 5 | suite_bulk_oof_hardened |
| SKCM | 100 | *MAP2K2* | 0.345 | – | 0.015 | 466 | 5 | suite_bulk_oof_hardened |
| SKCM | 100 | *NF1* | 0.690 | – | 0.280 | 466 | 5 | suite_bulk_oof_hardened |
| SKCM | 100 | *NRAS* | 0.834 | – | 0.660 | 466 | 5 | suite_bulk_oof_hardened |
| SKCM | 100 | *PIK3CA* | 0.769 | – | 0.061 | 466 | 5 | suite_bulk_oof_hardened |
| SKCM | 100 | *PPP6C* | 0.482 | – | 0.067 | 466 | 5 | suite_bulk_oof_hardened |
| SKCM | 100 | *PREX2* | 0.595 | – | 0.283 | 466 | 5 | suite_bulk_oof_hardened |
| SKCM | 100 | *PTEN* | 0.726 | – | 0.204 | 466 | 5 | suite_bulk_oof_hardened |
| SKCM | 100 | *RAC1* | 0.520 | – | 0.074 | 466 | 5 | suite_bulk_oof_hardened |
| SKCM | 100 | *SETD2* | 0.675 | – | 0.158 | 466 | 5 | suite_bulk_oof_hardened |
| SKCM | 100 | *TP53* | 0.861 | – | 0.581 | 466 | 5 | suite_bulk_oof_hardened |

Continued on next page

*Supplementary Table S9 continued*

| Cancer | $k$ | Driver | AUROC mean | AUROC SD | AUPRC mean | Samples | CV folds | Implementation |
|---|---|---|---|---|---|---|---|---|
| SKCM | 150 | *AKT1* | 0.437 | – | 0.009 | 372 | 5 | `suite_bulk_oof_hardened` |
| SKCM | 150 | *AKT3* | 0.523 | – | 0.020 | 466 | 5 | `suite_bulk_oof_hardened` |
| SKCM | 150 | *APC* | 0.566 | – | 0.113 | 466 | 5 | `suite_bulk_oof_hardened` |
| SKCM | 150 | *ARID1A* | 0.535 | – | 0.085 | 466 | 5 | `suite_bulk_oof_hardened` |
| SKCM | 150 | *ARID2* | 0.647 | – | 0.336 | 466 | 5 | `suite_bulk_oof_hardened` |
| SKCM | 150 | *BRAF* | 0.815 | – | 0.831 | 466 | 5 | `suite_bulk_oof_hardened` |
| SKCM | 150 | *CDKN2A* | 0.557 | – | 0.154 | 466 | 5 | `suite_bulk_oof_hardened` |
| SKCM | 150 | *CTNNB1* | 0.654 | – | 0.143 | 466 | 5 | `suite_bulk_oof_hardened` |
| SKCM | 150 | *GNA11* | 0.504 | – | 0.027 | 466 | 5 | `suite_bulk_oof_hardened` |
| SKCM | 150 | *GNAQ* | 0.681 | – | 0.119 | 466 | 5 | `suite_bulk_oof_hardened` |
| SKCM | 150 | *HRAS* | 0.409 | – | 0.014 | 466 | 5 | `suite_bulk_oof_hardened` |
| SKCM | 150 | *IDH1* | 0.815 | – | 0.399 | 466 | 5 | `suite_bulk_oof_hardened` |
| SKCM | 150 | *KDM6A* | 0.789 | – | 0.017 | 280 | 5 | `suite_bulk_oof_hardened` |
| SKCM | 150 | *KIT* | 0.786 | – | 0.269 | 466 | 5 | `suite_bulk_oof_hardened` |
| SKCM | 150 | *KRAS* | 0.672 | – | 0.048 | 466 | 5 | `suite_bulk_oof_hardened` |
| SKCM | 150 | *MAP2K1* | 0.592 | – | 0.073 | 466 | 5 | `suite_bulk_oof_hardened` |
| SKCM | 150 | *MAP2K2* | 0.357 | – | 0.015 | 466 | 5 | `suite_bulk_oof_hardened` |
| SKCM | 150 | *NF1* | 0.663 | – | 0.248 | 466 | 5 | `suite_bulk_oof_hardened` |
| SKCM | 150 | *NRAS* | 0.846 | – | 0.693 | 466 | 5 | `suite_bulk_oof_hardened` |
| SKCM | 150 | *PIK3CA* | 0.715 | – | 0.047 | 466 | 5 | `suite_bulk_oof_hardened` |
| SKCM | 150 | *PPP6C* | 0.505 | – | 0.075 | 466 | 5 | `suite_bulk_oof_hardened` |
| SKCM | 150 | *PREX2* | 0.586 | – | 0.293 | 466 | 5 | `suite_bulk_oof_hardened` |
| SKCM | 150 | *PTEN* | 0.700 | – | 0.195 | 466 | 5 | `suite_bulk_oof_hardened` |
| SKCM | 150 | *RAC1* | 0.544 | – | 0.076 | 466 | 5 | `suite_bulk_oof_hardened` |
| SKCM | 150 | *SETD2* | 0.709 | – | 0.148 | 466 | 5 | `suite_bulk_oof_hardened` |
| SKCM | 150 | *TP53* | 0.876 | – | 0.604 | 466 | 5 | `suite_bulk_oof_hardened` |
| SKCM | 250 | *AKT1* | 0.308 | – | 0.008 | 372 | 5 | `suite_bulk_oof_hardened` |
| SKCM | 250 | *AKT3* | 0.444 | – | 0.019 | 466 | 5 | `suite_bulk_oof_hardened` |
| SKCM | 250 | *APC* | 0.545 | – | 0.107 | 466 | 5 | `suite_bulk_oof_hardened` |
| SKCM | 250 | *ARID1A* | 0.572 | – | 0.064 | 466 | 5 | `suite_bulk_oof_hardened` |
| SKCM | 250 | *ARID2* | 0.684 | – | 0.338 | 466 | 5 | `suite_bulk_oof_hardened` |
| SKCM | 250 | *BRAF* | 0.810 | – | 0.821 | 466 | 5 | `suite_bulk_oof_hardened` |
| SKCM | 250 | *CDKN2A* | 0.591 | – | 0.174 | 466 | 5 | `suite_bulk_oof_hardened` |
| SKCM | 250 | *CTNNB1* | 0.663 | – | 0.138 | 466 | 5 | `suite_bulk_oof_hardened` |
| SKCM | 250 | *GNA11* | 0.553 | – | 0.030 | 466 | 5 | `suite_bulk_oof_hardened` |
| SKCM | 250 | *GNAQ* | 0.643 | – | 0.183 | 466 | 5 | `suite_bulk_oof_hardened` |
| SKCM | 250 | *HRAS* | 0.387 | – | 0.013 | 466 | 5 | `suite_bulk_oof_hardened` |
| SKCM | 250 | *IDH1* | 0.858 | – | 0.481 | 466 | 5 | `suite_bulk_oof_hardened` |
| SKCM | 250 | *KDM6A* | 0.602 | – | 0.009 | 280 | 5 | `suite_bulk_oof_hardened` |
| SKCM | 250 | *KIT* | 0.807 | – | 0.338 | 466 | 5 | `suite_bulk_oof_hardened` |
| SKCM | 250 | *KRAS* | 0.708 | – | 0.069 | 466 | 5 | `suite_bulk_oof_hardened` |
| SKCM | 250 | *MAP2K1* | 0.599 | – | 0.074 | 466 | 5 | `suite_bulk_oof_hardened` |
| SKCM | 250 | *MAP2K2* | 0.369 | – | 0.016 | 466 | 5 | `suite_bulk_oof_hardened` |
| SKCM | 250 | *NF1* | 0.695 | – | 0.275 | 466 | 5 | `suite_bulk_oof_hardened` |
| SKCM | 250 | *NRAS* | 0.861 | – | 0.700 | 466 | 5 | `suite_bulk_oof_hardened` |
| SKCM | 250 | *PIK3CA* | 0.703 | – | 0.043 | 466 | 5 | `suite_bulk_oof_hardened` |
| SKCM | 250 | *PPP6C* | 0.583 | – | 0.097 | 466 | 5 | `suite_bulk_oof_hardened` |
| SKCM | 250 | *PREX2* | 0.629 | – | 0.323 | 466 | 5 | `suite_bulk_oof_hardened` |
| SKCM | 250 | *PTEN* | 0.660 | – | 0.168 | 466 | 5 | `suite_bulk_oof_hardened` |
| SKCM | 250 | *RAC1* | 0.570 | – | 0.079 | 466 | 5 | `suite_bulk_oof_hardened` |
| SKCM | 250 | *SETD2* | 0.728 | – | 0.158 | 466 | 5 | `suite_bulk_oof_hardened` |
| SKCM | 250 | *TP53* | 0.890 | – | 0.603 | 466 | 5 | `suite_bulk_oof_hardened` |

**Supplementary Table S10. Controls and guardrails.** Summary of completed bulk-label permutations, solid-tumor reference-label permutations, cell-state reference fallback use, and reference-direction flags. Each row records the control family, cancer, number of tests, operational definition, summary result, and source artifact.

| Control family | Cancer | Tests | Definition | Summary metric | Source artifact |
|---|---|---|---|---|---|
| `bulk_label_permutation` | all | 158 | Driver labels permuted within each bulk cancer cohort; empirical AUROC null with 2,000 permutations. | 107 of 158 models had FDR $q<0.05$; 102 met full claim-safety criteria. | `Supplementary_Table_S1_bulk_model_audit.csv` |
| `reference_label_permutation` | BRCA | 18 | Reference/non-reference labels permuted at the observation level; directed tumour-versus-reference Cliff delta compared with 2,000-label null. | 8 positive FDR-supported drivers; 8 FDR-supported in either reported direction. | `Fig5_BRCA_Wu_permutation_null.csv` |
| `reference_label_permutation` | CRC | 25 | Reference/non-reference labels permuted at the observation level; directed tumour-versus-reference Cliff delta compared with 2,000-label null. | 20 positive FDR-supported drivers; 20 FDR-supported in either reported direction. | `Fig5_CRC_Lee_permutation_null.csv` |
| `reference_label_permutation` | GBM | 15 | Reference/non-reference labels permuted at the observation level; directed tumour-versus-reference Cliff delta compared with 2,000-label null. | 3 positive FDR-supported drivers; 3 FDR-supported in either reported direction. | `Fig5_GBM_Neftel_permutation_null.csv` |
| `reference_label_permutation` | LUAD | 20 | Reference/non-reference labels permuted at the observation level; directed tumour-versus-reference Cliff delta compared with 2,000-label null. | 15 positive FDR-supported drivers; 15 FDR-supported in either reported direction. | `Fig5_LUAD_Lambrechts_permutation_null.csv` |
| `reference_label_permutation` | PAAD | 15 | Reference/non-reference labels permuted at the observation level; directed tumour-versus-reference Cliff delta compared with 2,000-label null. | 10 positive FDR-supported drivers; 10 FDR-supported in either reported direction. | `Fig5_PAAD_Moncada_permutation_null.csv` |
| `reference_label_permutation` | SKCM | 26 | Reference/non-reference labels permuted at the observation level; directed tumour-versus-reference Cliff delta compared with 2,000-label null. | 8 positive FDR-supported drivers; 8 FDR-supported in either reported direction. | `Fig5_SKCM_JerbyArnon_permutation_null.csv` |
| `reference_guardrail_fallback` | AML | 232 | Audit of state-matched reference medians; cohort-wide reference median used when fewer than 25 same-state reference cells were available. | 0 of 232 driver-state combinations used the global fallback (0.0%). | `Reference_cellstate_guardrail_AML_vanGalen.csv` |
| `reference_guardrail_fallback` | BRCA | 144 | Audit of state-matched reference medians; cohort-wide reference median used when fewer than 25 same-state reference cells were available. | 90 of 144 driver-state combinations used the global fallback (62.5%). | `Reference_cellstate_guardrail_BRCA_Wu.csv` |
| `reference_guardrail_fallback` | CRC | 150 | Audit of state-matched reference medians; cohort-wide reference median used when fewer than 25 same-state reference cells were available. | 75 of 150 driver-state combinations used the global fallback (50.0%). | `Reference_cellstate_guardrail_CRC_Lee.csv` |
| `reference_guardrail_fallback` | GBM | 180 | Audit of state-matched reference medians; cohort-wide reference median used when fewer than 25 same-state reference cells were available. | 75 of 180 driver-state combinations used the global fallback (41.7%). | `Reference_cellstate_guardrail_GBM_Neftel.csv` |
| `reference_guardrail_fallback` | LUAD | 120 | Audit of state-matched reference medians; cohort-wide reference median used when fewer than 25 same-state reference cells were available. | 60 of 120 driver-state combinations used the global fallback (50.0%). | `Reference_cellstate_guardrail_LUAD_Lambrechts.csv` |

Continued on next page

*Supplementary Table S10 continued*

| Control family | Cancer | Tests | Definition | Summary metric | Source artifact |
|---|---|---|---|---|---|
| reference_guardrail_fallback | PAAD | 60 | Audit of state-matched reference medians; cohort-wide reference median used when fewer than 25 same-state reference cells were available. | 30 of 60 driver-state combinations used the global fallback (50.0%). | Reference_cellstate_guardrail_PAAD_Moncada.csv |
| reference_guardrail_fallback | SKCM | 208 | Audit of state-matched reference medians; cohort-wide reference median used when fewer than 25 same-state reference cells were available. | 130 of 208 driver-state combinations used the global fallback (62.5%). | Reference_cellstate_guardrail_SKCM_JerbyArnon.csv |
| reference_direction_audit | AML | 145 | Audit of whether healthy-reference scores exceeded the non-reference compartment and whether available mutation-label and tumour/reference effects disagreed in direction. | 12 reference-high flags; 5 lineage-confound flags. | Reference_direction_sensitivity_AML_vanGalen.csv |
| reference_direction_audit | BRCA | 36 | Audit of whether healthy-reference scores exceeded the non-reference compartment and whether available mutation-label and tumour/reference effects disagreed in direction. | 4 reference-high flags; 0 lineage-confound flags. | Reference_direction_sensitivity_BRCA_Wu.csv |
| reference_direction_audit | CRC | 50 | Audit of whether healthy-reference scores exceeded the non-reference compartment and whether available mutation-label and tumour/reference effects disagreed in direction. | 8 reference-high flags; 0 lineage-confound flags. | Reference_direction_sensitivity_CRC_Lee.csv |
| reference_direction_audit | GBM | 30 | Audit of whether healthy-reference scores exceeded the non-reference compartment and whether available mutation-label and tumour/reference effects disagreed in direction. | 6 reference-high flags; 0 lineage-confound flags. | Reference_direction_sensitivity_GBM_Neftel.csv |
| reference_direction_audit | LUAD | 40 | Audit of whether healthy-reference scores exceeded the non-reference compartment and whether available mutation-label and tumour/reference effects disagreed in direction. | 6 reference-high flags; 0 lineage-confound flags. | Reference_direction_sensitivity_LUAD_Lambrechts.csv |
| reference_direction_audit | PAAD | 30 | Audit of whether healthy-reference scores exceeded the non-reference compartment and whether available mutation-label and tumour/reference effects disagreed in direction. | 4 reference-high flags; 0 lineage-confound flags. | Reference_direction_sensitivity_PAAD_Moncada.csv |
| reference_direction_audit | SKCM | 52 | Audit of whether healthy-reference scores exceeded the non-reference compartment and whether available mutation-label and tumour/reference effects disagreed in direction. | 22 reference-high flags; 0 lineage-confound flags. | Reference_direction_sensitivity_SKCM_JerbyArnon.csv |

##### 3.1 Machine-readable file index

Each typeset table above has a corresponding CSV file that preserves full numerical precision and the normalized field structure for programmatic use.

**Index of machine-readable Supplementary Tables.** Each file is supplied as a separate CSV in the `Supplementary_Tables/` directory.

| Table | File | Content |
|---|---|---|
| S1 | `Supplementary_Table_S1_bulk_model_audit.csv` | Complete 158-model bulk audit: valid pooled out-of-fold class counts, prevalence, AUROC and bootstrap interval, AUPRC and prevalence baseline, Brier score, permutation result, FDR, prior tier, fold-level omissions, and claim-safety status. |
| S2 | `Supplementary_Table_S2_data_sources_accessions_and_cohort_characteristics.csv` | Bulk and single-cell source publications and DOIs, repository/project accessions, access status, analysis-input provenance, modeled or retained sizes, and single-cell reference annotations. |
| S3 | `Supplementary_Table_S3_AML_percell_truth.csv` | Complete AML direct-label validation for all 12 evaluable drivers, including score column, cell counts, apparent AUROC, Mann–Whitney result, Cliff's $\delta$, intervals, and FDR. |
| S4 | `Supplementary_Table_S4_patient_level_bridge.csv` | Patient-level genotype bridge for AML, CRC, and BRCA using median cell-state-residual scores. |
| S5 | `Supplementary_Table_S5_CNV_concordance_and_discordance.csv` | Continuous score–CNV correlations, binary adjusted Rand indices, four-way discordance counts, and group-wise mean scores and CNV burdens. |
| S6 | `Supplementary_Table_S6_near_diploid_patient_validation.csv` | Patient-level driver comparisons after restricting to patients with inferred-CNV malignant fraction below 0.10. |
| S7 | `Supplementary_Table_S7_AML_longitudinal_driver_trends.csv` | Complete patient-by-driver longitudinal summary, including first and last effects, change statistics, trend tests, and evaluation status. |
| S8 | `Supplementary_Table_S8_panel_provenance.csv` | Panel-level provenance linking manuscript items to source artifacts, analysis version, configuration hash, model object, alignment, component count, score column, subset, test, correction family, and random seed. |
| S9 | `Supplementary_Table_S9_latent_dimension_sensitivity.csv` | Cancer-, driver-, and component-count-specific cross-validated AUROC and AUPRC from the archived latent-dimension sensitivity grid. |
| S10 | `Supplementary_Table_S10_controls_and_guardrails.csv` | Summary of completed bulk-label permutations, solid-tumor reference-label permutations, cell-state reference fallback usage, and reference-direction flags. |

Additional unnumbered machine-readable files in the `Supplementary_Tables/` directory contain the cohort-specific reference-label permutation nulls, state-matched reference guardrail audits, and reference-direction audits used to construct Supplementary Figure S10. The complete per-driver source outputs remain in `Supplemental_Code.zip`.
